# The genetic architecture of language functional connectivity

**DOI:** 10.1101/2021.10.18.464351

**Authors:** Yasmina Mekki, Vincent Guillemot, Hervé Lemaitre, Amaia Carrion-Castillo, Stephanie Forkel, Vincent Frouin, Cathy Philippe

## Abstract

Language is a unique trait of the human species, of which the genetic architecture remains largely unknown. Through language disorders studies, many candidate genes were identified. However, such complex and multifactorial trait is unlikely to be driven by only few genes and case-control studies, suffering from a lack of power, struggle to uncover significant variants. In parallel, neuroimaging has significantly contributed to the understanding of structural and functional aspects of language in the human brain and the recent availability of large scale cohorts like UK Biobank have made possible to study language via image-derived endophenotypes in the general population. Because of its strong relationship with task-based fMRI activations and its easiness of acquisition, resting-state functional MRI have been more popularised, making it a good surrogate of functional neuronal processes. Taking advantage of such a synergistic system by aggregating effects across spatially distributed traits, we performed a multivariate genome-wide association study (mvGWAS) between genetic variations and resting-state functional connectivity (FC) of classical brain language areas in the inferior frontal (pars opercularis, triangularis and orbitalis), temporal and inferior parietal lobes (angular and supramarginal gyri), in 32,186 participants from UK Biobank. Twenty genomic loci were found associated with language FCs, out of which three were replicated in an independent replication sample. A locus in 3p11.1, regulating *EPHA3* gene expression, is found associated with FCs of the semantic component of the language network, while a locus in 15q14, regulating *THBS1* gene expression is found associated with FCs of the perceptual-motor language processing, bringing novel insights into the neurobiology of language.

## 1. Introduction

Language is a unique trait of the human species. Although its genetic origins are broadly accepted, they remain largely unknown. Since the seminal study that revealed the major role of *FOXP2* in language processing (Fisher et al., 1998), several candidate genes related to language disorders were identified (Landi and Perdue, 2019). Human language is a complex system – both structurally and functionally. As such a complex and multifactorial trait, it is unlikely to be associated with only a few genes but rather with many genes that are also interacting with each other. These genes likely contribute to the development of neural pathways involved in language development, together with experience-dependent contributions from the environment (Fisher and Vernes, 2015). In parallel, neuroimaging techniques provided innovative and quantitative ways to study language. Anatomically, the language system comprises perisylvian cortical regions predominantly - but not exclusively - in the left hemisphere. Amongst these regions the prominent regions are in the pars orbitalis and triangularis in the inferior frontal gyrus (also referred to as ‘Broca’s’ region), the angular and supramarginal gyri in the inferior parietal lobe (also referred to as ‘Geschwind’s’ region), and the posterior temporal regions (‘Wernicke’s’ region). These cortical regions are interconnected by a network of brain connections, most prominently the arcuate fasciculus (Catani et al., 2005; Forkel and Catani, 2018). These regions also connect to the sensory-motor system (auditory, visual, and motor cortex). Functionally, phonology, semantics, and syntax are three main language components and form a tripartite parallel architecture (e.g. Jackendoff and Jackendoff (2002); Bates et al. (2003); Vigneau et al. (2006); Price (2012)). Consequently, it has become common practice to study language in the healthy population using neuroimaging. Mapping endophenotypes based on anatomical and task-based functional MRI expanded the understanding of how the brain supports language (Price, 2012; Friederici, 2017; Ardila et al., 2016; Leroy et al., 2015; Labache et al., 2020). These MRI endophenotypes give access to biologically relevant measurements of individual variability (Forkel et al., 2014a, 2020b; Uddén et al., 2019; Dubois and Adolphs, 2016; Seghier and Price, 2018; Fedorenko, 2021; Forkel et al., 2020a) and are consequently suitable for the search of genetic associations. Traditionally, the language-brain relationship is investigated through task-based activation experiments. In the past decade, the use of resting-state functional MRI (rsfMRI) has been popularised mainly due to the strong relationship observed between the signals collected in resting-state and the cognitive task-based fMRI activations (Smith et al., 2009; Tavor et al., 2016; Cole et al., 2016; Ngo et al., 2020; Dohmatob et al., 2021). rsfMRI is paradigm free and as such it is easier to implement in very large cohorts like the UK Biobank. rsfMRI can recover specific brain functional activations making it a good surrogate of functional neuronal processes. Here, we propose to use task-free functional connectivity (FC) from the UK Biobank (Bycroft et al., 2018), using perisylvian cortical regions as areas of interest serving as a proxy for language.

As rsfMRI FC are low amplitudes and correlated to each other, we anticipate it would be really difficult to disentangle the genetic associations with each FC signal in a massively univariate manner. Language related brain regions share information across components and scales, and genetic variants are supposed to have distributed effect across regions. Thus, we take advantage of this synergistic system and perform a multivariate approach with MOSTest (van der Meer et al., 2020). This method considers the distributed nature of genetic signals shared across brain regions and aggregates effects across spatially distributed traits of interest. This approach tests each SNP independently for its simultaneous association with the brain endophenotypes, making it half multivariate and half univariate. For convenience, we will use the term “multivariate GWAS” (mvGWAS) while being aware that correlations between SNPs are not accounted for in this approach.

In this study, we use rsfMRI in a discovery sample of 32,186 healthy volunteers from the UK Biobank (Sudlow et al., 2015) and the compiled information of a large-scale meta-analysis on language components (Vigneau et al., 2006, 2011). First, we derived functional connectivity (FC) endophenotypes reflecting individual language network characteristics, based on regions of interest from the meta-analysis. Second, we performed a multivariate genetic association of language specific functional connectivity, filtered based on heritability significance. The results from this analysis were subjected to a replication study in an independent sample (N=4,754). Additionally, as the connections between different language regions are ensured by the white matter bundles (Catani et al., 2005; Catani and Forkel, 2019), we tested the potential associations of the hit SNPs with the neuroanatomical tracts underlying the hit endophenotypes using diffusion-based white matter analysis. Finally, the extensive functional annotations of each genomic risk locus allowed us to suggest two new genes with a role in different aspects of the language system.

## 2. Materials and Methods

### 2.1. Demographics and neuroimaging Data from the UK Biobank

#### UK Biobank cohort

The UK Biobank is an open-access longitudinal populationwide cohort study that includes 500k participants from all over the United Kingdom (Sudlow et al., 2015). Data collection comprises detailed genotyping and a wide variety of endophenotypes ranging from health/activity questionnaires, extended demographics to neuroimaging and clinical health records. All participants provided informed consent and the study was approved by the North West Multi-Centre Research Ethics Committee (MREC).

This study used the February 2020 release (application number #64984). This release consisted of 36,940 participants, age range between 40 to 70 years (mean age=54 ±7.45 years), with genotyping and resting-state functional MRI. To avoid any possible confounding effects related to ancestry, we restricted our analysis to individuals with British ancestry using the sample quality control information provided by UK Biobank (Bycroft et al., 2018). A final cohort of 32,186 volunteers (15,234 females, mean age = 55 ±7.51 years) were included in the study. We made use of the first ten principal components (Data field 22009) of the genotyping data’s multidimensional scaling analysis capturing population genetic diversity to account for population stratification. An independent replication dataset of 4754 non-British individuals was also drawn from the UK Biobank. The age range of these participants was 40 to 70 (mean age = 53 ±7.55 years), 2153 were female.

#### Resting-state functional MRI data

The MRI data available from the UK Biobank are described in the UK Biobank Brain Imaging Documentation (v.1.7, January 2020) as well as in (Miller et al., 2016; Alfaro-Almagro et al., 2018). Briefly, resting-state functional MRI (rsfMRI) data were acquired using the following parameters: 3T Siemens Skyra scanner, TR = 0.735s, TE = 39ms, duration = 6 min (490 time points), resolution: 2.4 × 2.4 × 2.4 mm, Field-of-view = 88 × 88 × 64 matrix. During the resting-state scan, participants were instructed to keep their eyes fixated on a crosshair, to relax, and to think of nothing particular (Miller et al., 2016). The preprocessing of the UK Biobank data includes motion correction, grand-mean intensity normalisation, high-pass temporal filtering including EPI unwarping with alignment to the T1 template and gradient non-linearity distortion correction (GDC) unwarping, brain masking, and registration to MNI space. The rsfMRI volumes were further cleaned using ICA-FIX for automatically identifying and removing artefacts.

#### Diffusion-weighted MRI data

The Diffusion-weighted MRI (dMRI) images were acquired using the following parameters; isotropic voxel size (resolution): 2 × 2 × 2 mm, five non diffusion-weighted image b=0 s/*mm*^2^, diffusion-weighting of b=1000, and 2000 s/*mm*^2^ with 50 directions each, acquisition time: 7 min. Tensor fits utilize the b=1000 s/*mm*^2^ data and the NODDI (Zhang et al., 2012) (Neurite Orientation Dispersion and Density Imaging) model is fits using AMICO (Daducci et al., 2015) (Accelerated Microstructure Imaging via Convex Optimization) tool, creating outputs including nine diffusion indices maps. These ones were subject to a TBSS-style analysis using FSL tool resulting in a white matter skeleton mask.

### 2.2. Genetic quality control

Genotyping was performed using the UK BiLEVE Axiom array by Affymetrix (Wain et al., 2015) on a subset of 49,950 participants (807,411 markers) and the UK Biobank Axiom array on 438,427 participants (825,927 markers). Both arrays are extremely similar and share 95% of common SNP probes. The imputed genotypes were obtained from the UK Biobank repository Bycroft et al. (2018). These genetic data underwent a stringent quality control protocol, excluding participants with unusual heterozygosity, high missingness (Data field 22027), sex mismatches, such as discrepancy between genetically inferred sex (Data field 22001) and self-reported sex (Data field 31). Variants with minor allele frequency (MAF) < 0.01 were filtered out from the imputed genotyping data using PLINK 1.9 (Chang et al., 2015) to retain the common variants only. Overall, 9,812,367 autosomal SNPs were considered.

### 2.3. Regions of interest for rsfMRI functional connectivity

We leverage a large-scale meta-analysis of 946 activation peaks (728 peaks in the left hemisphere, 218 peaks in the right hemisphere) obtained from a meta-analysis of 129 task-based fMRI language studies (Vigneau et al., 2006, 2011). The identified fronto-parietal-temporal activation foci revealed via a hierarchical clustering analysis, 50 distinct, albeit partially overlapping, clusters of activation foci for phonology, semantics, and sentence processing: 30 clusters in the left hemisphere and 20 in the right hemisphere.

Because this overlap could unduly increase the co-activation between regions and to avoid a deconvolution bias in the estimation of the functional connectivity, we proceeded as follow: First, because the clustering process was performed for each component independently, we checked whether pairs of clusters belonging to different language-component networks were spatially distinct considering the significance of their mean Euclidean distance with paired t-tests. We identified areas that are common to multiple language components; in the temporal lobe, the anterior part of the Superior temporal gyrus (T1a) area appears to be common to all three language components, the anterior part of the superior temporal sulcus (Pole) and Lateral/middle part of the middle temporal gyrus (T2ml) are common to semantic and sentence’s clusters and the posterior part of the left inferior temporal gyrus (T3p) to semantic and phonology clusters. In the frontal lobe, the L—R dorsal part of the pars opercularis (F3opd) and the ventral part of the pars triangularis (F3tv) are common to semantic and syntactic clusters. In these cases, we retained the larger cluster and assigned multiple labels. Second, ROIs were obtained for each cluster by building a 3D convex-hull of the peaks in the MNI space and were then subjected to a morphological opening operation. Third, overlapping areas between the convex-hull ROIs were processed as follows: the common region between two ROIs was attributed to the most representative in terms of the number of peaks. Finally, we excluded regions with less than 100 voxels. This preprocessing resulted in 25 multilabelled ROIs: 19 in the left hemisphere and 6 in the right hemisphere which are summarised in Fig. 1 and Table SI1.

**Figure 1:**
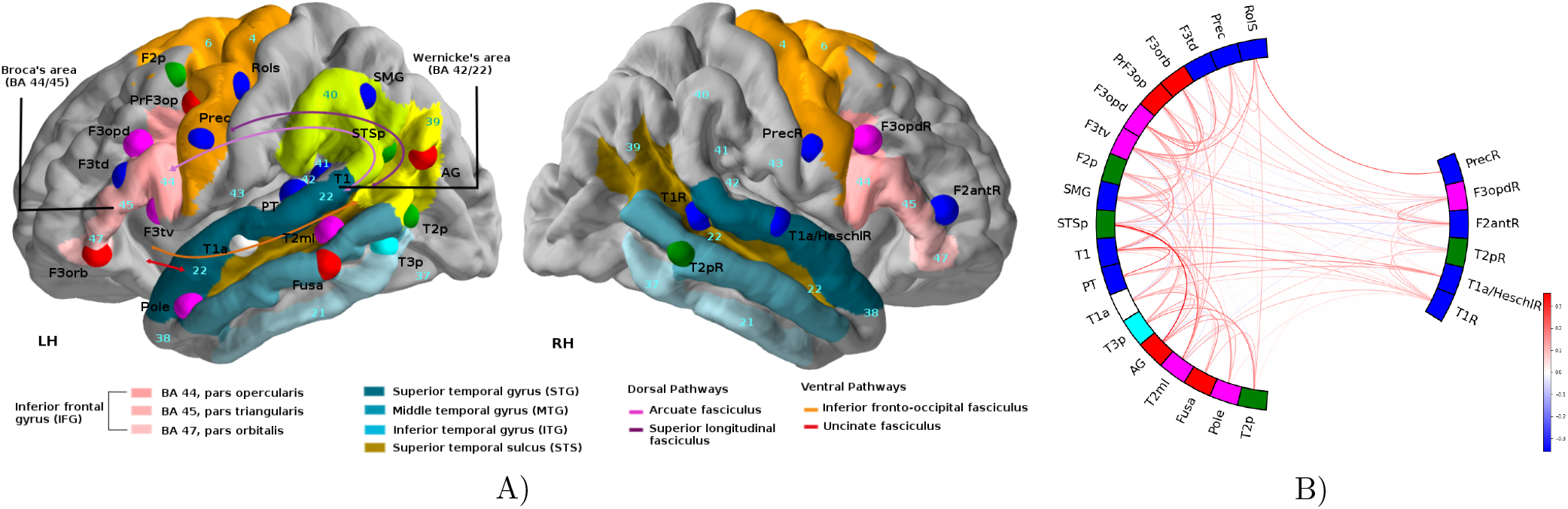
A) Overview of the regions obtained from the meta-analysis. Each language seed is color-coded according to its language category: phonology (blue), semantic (red), and syntax (green). ROIs of different components that were not spatially distinct are color-coded as pink (semantic/syntax), cyan (phonology/semantic) and white for the three language component. For the sake of ROIs figure visibility, the coordinates were modified. The exact coordinates for each ROI are available in Table ??. Different gyri and sulcus, known to be relevant for language: the inferior frontal gyrus (IFG), middle temporal gyrus (MTG), superior temporal gyrus (STG), and superior temporal sulcus (STS), are color-coded. Numbers in the left hemisphere (LH) represents language-relevant Brodmann areas (BA) which were defined on the basis of cytoarchitectonic characteristics. Numbers in the right hemisphere (RH) represents the language-relevant BA counterpart. The pars opercularis (BA 44), the pars triangularis (BA 45) represents Broca’s area. The pars orbitalis (BA 47) is located anterior to Broca’s area. BA 42 and BA 22 represents Wernicke’s area Friederici (2011). Both supramarginal gyrus (BA40) and angular gyrus (BA39), also known as Geschwind’s territory, are represented by green/yellow colors respectively. The primary motor cortex (BA4), the premotor cortex and the supplementary motor area (B6) are colored in orange. Whitin the left hemisphere, dorsal and ventral long-range fiber bundles connect language areas and are indicated by color-coded arrows. B) Mean functional connectivity of the 142 heritable endophenotypes, calculated using a shrinked estimate of partial correlation Marrelec et al. (2006) (estimated with a Ledoit-Wolf estimator Ledoit and Wolf (2004)) over 32,186 UKB rs fMRI subjects.

### 2.4. Neuroimaging endophenotypes

#### Functional connectivity endophenotypes

The preprocessed resting-state BOLD signal was masked using the 25 ROIs and averaged at each time volume. A connectome matrix was computed using Nilearn (Abraham et al., 2014) for each participant using a shrunk (Ledoit and Wolf, 2004) estimate of partial correlation (Marrelec et al., 2006). This resulted in 300 (= 25 × 24*/*2) edges connecting language ROIs for each individual. Each edge -also denoted functional connectivity (FC)-is further considered as a candidate endophenotype. See the Fig. 1b.

#### Diffusion MRI endophenotypes

We hypothesised that the hit SNPs associated with the hit FCs could be associated with neuroanatomical white matter tracts that supports the information transmission between the regions that compose these hit-FCs. Therefore, we tested the potential associations between the hit SNPs with the following white matter bundles: the corpus callosum, the left frontal aslant tract, the left arcuate anterior/long/posterior segment, the left inferior fronto-occipital fasciculus, the left uncinate tract (See section 3.3 for more details). The resulting skeletonised images are averaged across the set of 7 brain white matter structures defined by the probabilistic atlas (Rojkova et al., 2016) thresholded at 90% of probabilities. These structural white matter tracts are assessed by 9 indices: fractional anisotropy (FA) maps, tensor mode (MO), mean diffusivity (MD), intracellular volume fraction (ICVF), isotropic volume fraction (ISOVF), mean eigenvectors (L1, L2, L3), and orientation dispersion index (OD) yielding 63 = 7 × 9 dMRI endophenotypes.

### 2.5. SNP-based heritability and genetic correlation analysis

The proportion of additive genetic variance in the FC phenotypic variance, also called narrow-sense heritability, was estimated using the genotyped SNPs information using genome-based restricted maximum likelihood (GREML) (Yang et al., 2010) for each FCs, controlling for the above-mentioned covariates (refer to section 2.4). To define significantly heritable FCs, a 0.05 threshold on False Discovery Rate (FDR) adjusted p-values was applied to account for multiple testing on the 300 FCs. Similarly, the proportion of additive genetic variance in the covariance of pairs of FCs was estimated using the bivariate GREML (Lee et al., 2012). Both heritability and part of covariance explained by genetics were obtained using GCTA (Yang et al., 2011).

### 2.6. Multivariate genome-wide association studies (mvGWAS)

We performed a multivariate genome-wide association studies (mvGWAS) between the filtered imputed genotypes and the 142 significantly heritable FC endophenotypes, using the Multivariate Omnibus Statistical Test (MOSTest) (van der Meer et al., 2020). All endophenotypes were pre-residualised controlling for covariates including sex, genotype array type, age, recruitment site, and ten genetic principal components provided by UK Biobank. In addition, MOSTest performs a rank-based inverse-normal transformation of the residualised endophenotypes to ensure that the inputs are normally distributed. The distributions across the participants of all endophenotypes were visually inspected before and after co-variate adjustment. MOSTest generated summary statistics that capture the significance of the association across all heritable 142 language FC endophenotypes. To account for multiple testing over the whole genome, statistically significant SNPs were considered as those reaching the genome-wide threshold *p* = 5e−8.

### 2.7. mvGWASes replication

The multivariate genome-wide association results were replicated in an independent non-British sample considering the nominal significance threshold *p* < 0.05. Following the same pre-processing steps as for the primary sample, the non-British replication sample consists in 4,754 individuals with a mean age of 53 years (±7.55) and 2,153 female.

### 2.8. Fine-mapping: identification of genomic risk loci and functional annotation

We performed functional annotation analysis using the FUMA online platform v1.3.6a (Watanabe et al., 2017) with default parameters. The genomic positions are reported according to the GRCh37 reference. SNPs were annotated for functional consequences on gene functions using ANNOVAR (Wang et al., 2010), Combined Annotation Dependent Depletion (CADD) scores (Kircher et al., 2014), and 15-core chromatin state prediction by ChromHMM (Ernst and Kellis, 2012). In addition, they were annotated for their effects on gene expression using eQTLs of various tissue types. The eQTL module queried data from different tissue-datasets using GTEx v8 (Consortium et al., 2017), Blood eQTL browser (Westra et al., 2013), BIOS QTL browser (Zhernakova et al., 2017), BRAINEAC (Ramasamy et al., 2014), eQTLGen (Võsa et al., 2018), PsychENCODE (Wang et al., 2018), DICE (Schmiedel et al., 2018). RegulomeDB v2.0 (Boyle et al., 2012) was queried externally. Coding hit SNPs are also annotated with polymorphism phenotyping v2 (Polyphen-2) (Ramensky et al., 2002).

## 3. Results

### 3.1. SNP-based heritability of functional connectivity measures

The single-nucleotide polymorphism (SNP)-based heritability (*h*^2^) was estimated for each of the 300 FCs endophenotypes. P-values correction for multiple testing revealed 142 FCs significant SNP-based heritabilities (Table SI2), ranging from 14% for the SMG↔F3opd to 3% for the SMG↔T1 FC.

### 3.2. Multivariate genome-wide association analysis

We performed a multivariate genome-wide association study (mvGWAS) using the Multivariate Omnibus Statistical Test (MOSTest) (van der Meer et al., 2020) method, with the 142 FCs with significant SNP-based heritability. This analysis tested each SNP separately for its simultaneous association with the 142 FCs and yielded 4566 significant SNPs at a genomic threshold (see Table SI3), distributed on chromosomes 2, 3, 5, 6, 10, 11, 14, 15, 17, 18 and 22. FUMA (Watanabe et al., 2017) software was used to analyse mvGWAS results and identify lead SNPs at each associated locus. Considering the genome-wide significance threshold *p* = 5e−8, there were 20 distinct genomic loci distributed on the 11 chromosomes, associated with different aspects of language FC (Fig.2a, Table 1 and Fig. SI1, SI2, 2b, and SI3) and represented by 20 lead SNPs.

**Figure 2:**
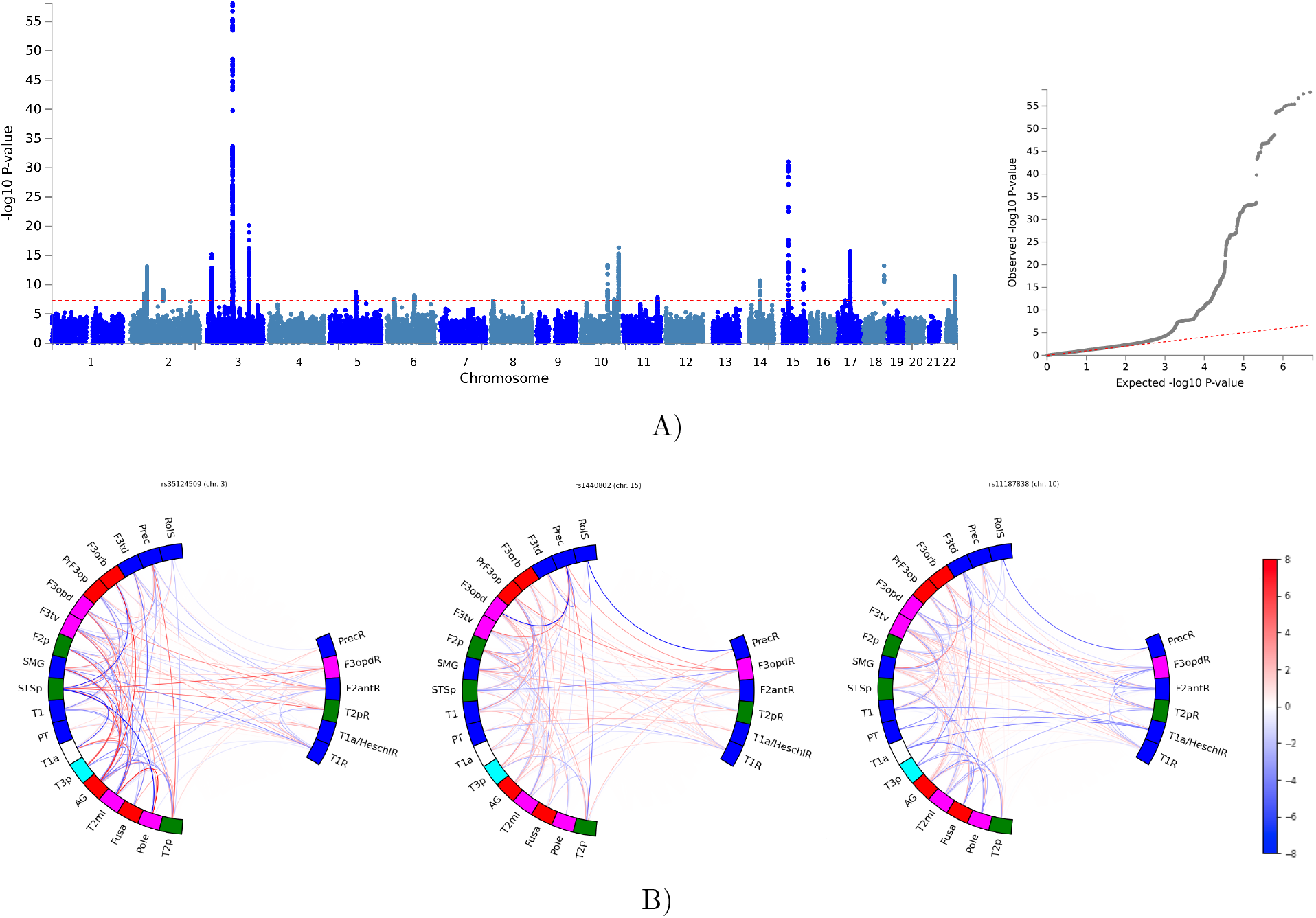
A) Multivariate GWAS analysis of the resting state functional connectivity in 32,186 participants. Manhattan plot for multivariate GWAS accross 142 FCs. The red dashed line indicates the genome-wide significance threshold *p* = 5e 8. The Quantile-quantile plot is also shown. B) Circle plot illustrating the 3 lead SNPs identified from the mvGWAS. Z-values from the univariate GWAS for each FC are mapped. The absolute Z-values scaling is clipped at 8 (*p* = 1.2e 15). Positif effects of carrying the minor allele are shown in red, and negative in blue.

**Table 1:**
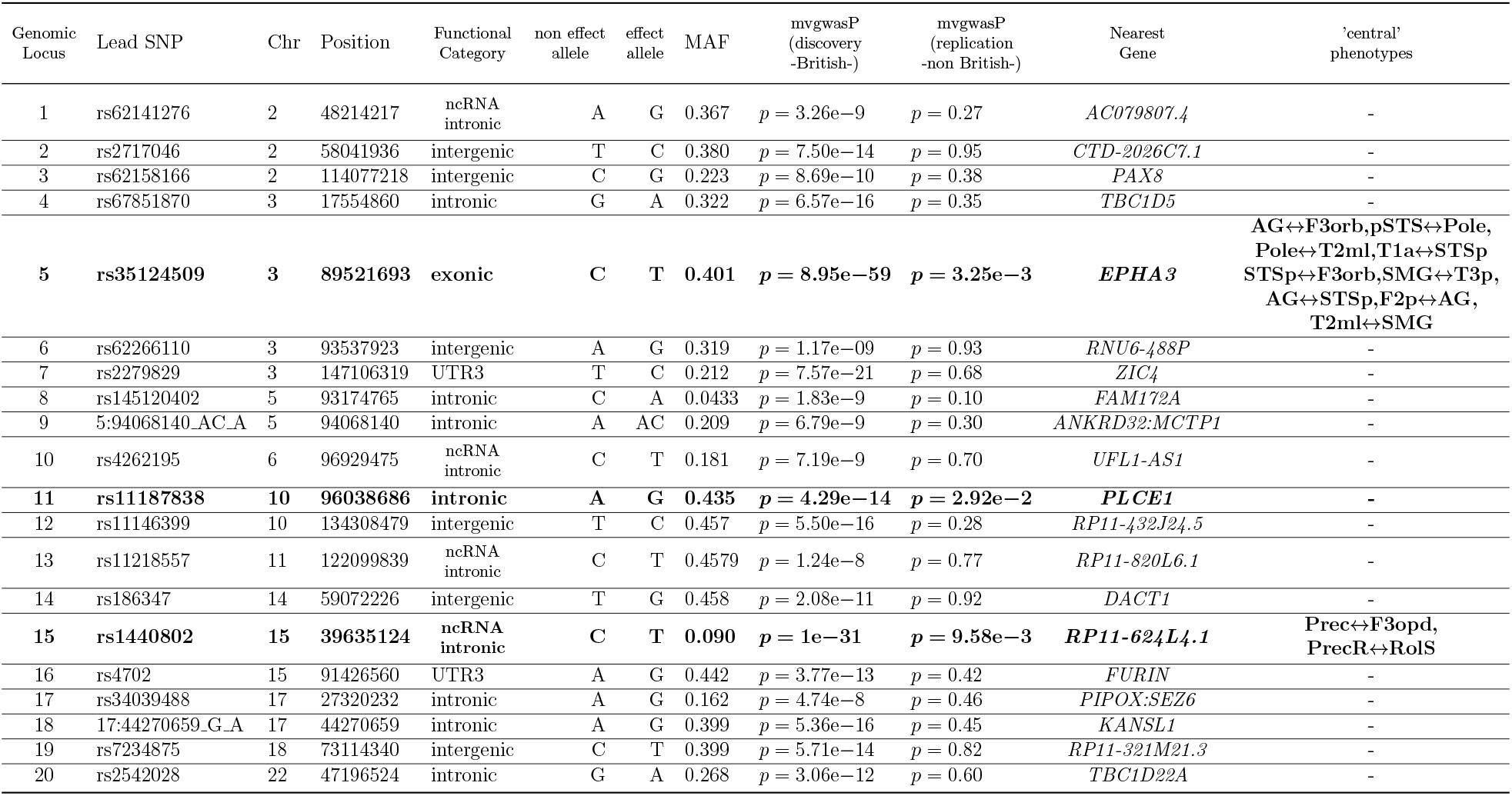
Genomic loci associated highlighted using the multivariate genome-wide association studie. Lead SNP: ID of the lead SNPs within each locus. Position: position of the SNP in the hg19 human reference genome. mvgwasP discovery -British-: MOSTest association P value obtained using the discovery sample. mvgwasP replication -non British-: MOSTest association P value obtained using the independent replication sample. Functionnal category: Functional consequence of the SNP on the gene obtained from ANNOVAR. ‘Central’ phenotypes: the phenotypes that contributed most to the multivariate association considering the genome-wide association threshold (5*e* − 8).

#### Validation of lead SNPs associated with rsfMRI FCs

The three lead SNPs were replicated at the nominal significance level (*p* < 5e−2) on multivariate test in the independent non-British replication dataset: *rs*1440802(*p* = 9.58e−3), *rs*35124509(*p* = 3.25e−3), *rs*11187838(*p* = 2.92e−2). Table SI4 summarises these results. Moreover, these lead SNP showed association at *p* < 0.05 on univariate testing of all but three specific central traits identified in the discovery mvGWAS.Here, we present three of these loci that were replicated in an independent data set (refer to section 2.3). MOSTest results highlighted the three following genomic risk regions: *i)* 15q14 locus (chr15, start=39598529, length=260kb) with its strongest association related to the imputed SNP rs1440802 (*p* = 1e−31); *ii)* 3p11.1 locus (chr3, start=89121389, length=1,381kb) with its strongest association related to the imputed SNP rs35124509 (*p* = 8.95e−59); *iii)* 10q23.33 locus (chr10, start=95988042, length=139kb) with its strongest association related to the imputed SNP rs11187838 (*p* = 4.29e−14). See Fig. 2a and Table 1.

#### Identification of central endophenotypes associated with genomic risk regions

For each lead SNP, we defined the ‘central’ endophenotypes that contributed the most in the multivariate association by using the individual univariate summary statistics performed by MOSTest and by considering the genome-wide significance threshold (*p* < 5*e* − 8) (Table SI5).

On 15q14, the lead SNP rs1440802 had two central FCs: the minor allele was associated with the partial correlation between *i)* the precentral gyrus and the dorsal pars opercularis (Prec↔F3opd). Both connected regions are in the left frontal lobe, and are labelled with a phonological linguistic component (Prec) and multi-labelled with semantic and sentence language processing (F3opd). *ii)* The (PrecR↔RolS) corresponds to the partial correlation between the precentral gyrus and the Rolandic sulcus. Both regions are identified in the right and left frontal lobes respectively, and are labelled as phonological linguistic component (Fig. 3a and Table SI5). These edges have previously been described in FC studies dedicated to language and more specifically in the perceptual motor interactions (Schwartz et al., 2008; Fridriksson et al., 2009; Turner et al., 2009; Nishitani and Hari, 2000; Schwartz et al., 2012). At the univariate level, these loci associated to central endophenotypes display an important overlap; See Fig 3d.

**Figure 3:**
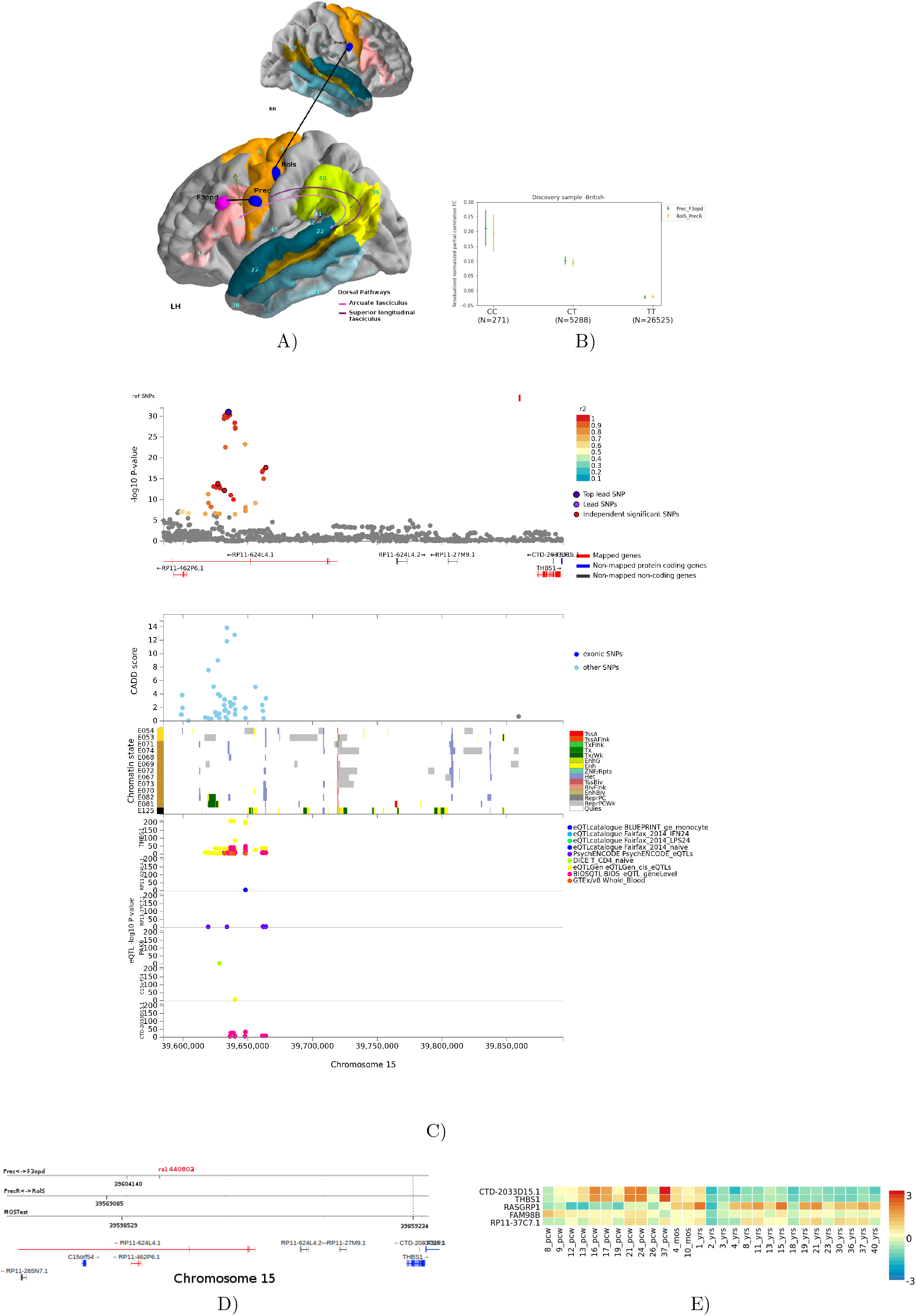
Main results for the 15q14 locus. A) The two pairs of ROIs that forms the endpoints of the associated FCs reported as black bold lines. B) Effect sizes of the SNP rs1440802 for the two connections: (Prec↔F3opd) FC in green and (PrecR↔RolS) FC in yellow. C) Locus Zoom of the genomic region identified by the mvGWAS. Chromatin state of the genomic region. Brain tissue name abbreviations are the following; E054:Ganglion Eminence derived primary cultured neurospheres, E053: Cortex derived primary cultured neurospheres, E071: Brain Hippocampus Middle, E074: Brain Substantia Nigra, E068: Brain Anterior Caudate, E069: Brain Cingulate Gyrus, E072: Brain Inferior Temporal Lobe, E067:Brain Angular Gyrus, E073: Brain Dorsolateral Prefrontal Cortex, E070: Brain Germinal Matrix, E082: Fetal Brain Female, E081: Fetal Brain Male, E125: NH-A Astrocytes Primary Cells. The state abbreviations are the following; TssA: active transcription start site (TSS), TssFlnk: Flanking Active TSS, TxFlnk: Transcription at gene 5‚ and 3‚, Tx: Strong transcription, TxWk: Weak transcription, EnhG: Genic enhancers, Enh: Enhancers, ZNF/Rpts: ZNF genes & repeats, Het: Heterochromatin, TssBiv: Bivalent/Poised TSS, BivFlnk: Flanking Bivalent TSS/Enh, EnhBiv: Bivalent Enhancer, ReprPC: Repressed PolyComb, ReprPCWk: Weak Repressed PolyComb, Quies: Quiescent/Low. Expression quantitative trait loci (eQTL) associations (data source: eQTLGen (Võsa et al., 2018), PsychENCODE(Wang et al., 2018), DICE (Schmiedel et al., 2018), BIOS QTL browser (Zhernakova et al., 2017), GTEx/v8 (Consortium et al., 2017), eQTLcatalogue). D) Overlap of the genomic region risk region identified from FUMA for MOSTest results, (Prec↔F3opd) and (PrecR↔RolS). E) Gene expression from BrainSpan for the interesting genes prioritised by FUMA.

On 3p11.1, the lead SNP rs35124509 had nine central FCs: the minor allele was associated with the partial correlation between the left posterior part of the superior temporal sulcus and the left temporal pole (Pole↔STSp), the left temporal pole and the lateral/middle part of the middle temporal gyrus (Pole↔T2ml), the angular gyrus and the pars orbitalis of the left inferior frontal gyrus (AG↔F3orb), the anterior part of the Superior temporal gyrus and the left posterior part of the superior temporal sulcus (T1a↔STSp), the left posterior part of the superior temporal sulcus and the pars orbitalis of the left inferior frontal gyrus (STSp↔F3orb), the supramarginal gyrus and the posterior part of the left inferior temporal gyrus (SMG↔T3p), the angular gyrus and the left posterior part of the superior temporal sulcus (AG↔STSp), the Posterior part of the middle frontal gyrus and the angular gyrus (F2p↔AG), the lateral/middle part of the middle temporal gyrus and the supramarginal gyrus (T2ml↔SMG) (Fig. 4a and Table SI5). These connected regions are located across the left parieto-frontal-temporal lobe, and are mainly labelled as semantic language processing. These edges have previously been described in FC studies dedicated to language and especially to the semantic component. This component typically includes the inferior frontal gyrus, the left temporal cortex (i.e. temporal pole, middle temporal gyrus, fusiform gyrus) and the left angular gyrus (Binder et al., 2009; Jackson et al., 2016; Vigneau et al., 2006). At the univariate level, these loci associated to central endophenotypes display an important overlap; See Fig 4d.

**Figure 4:**
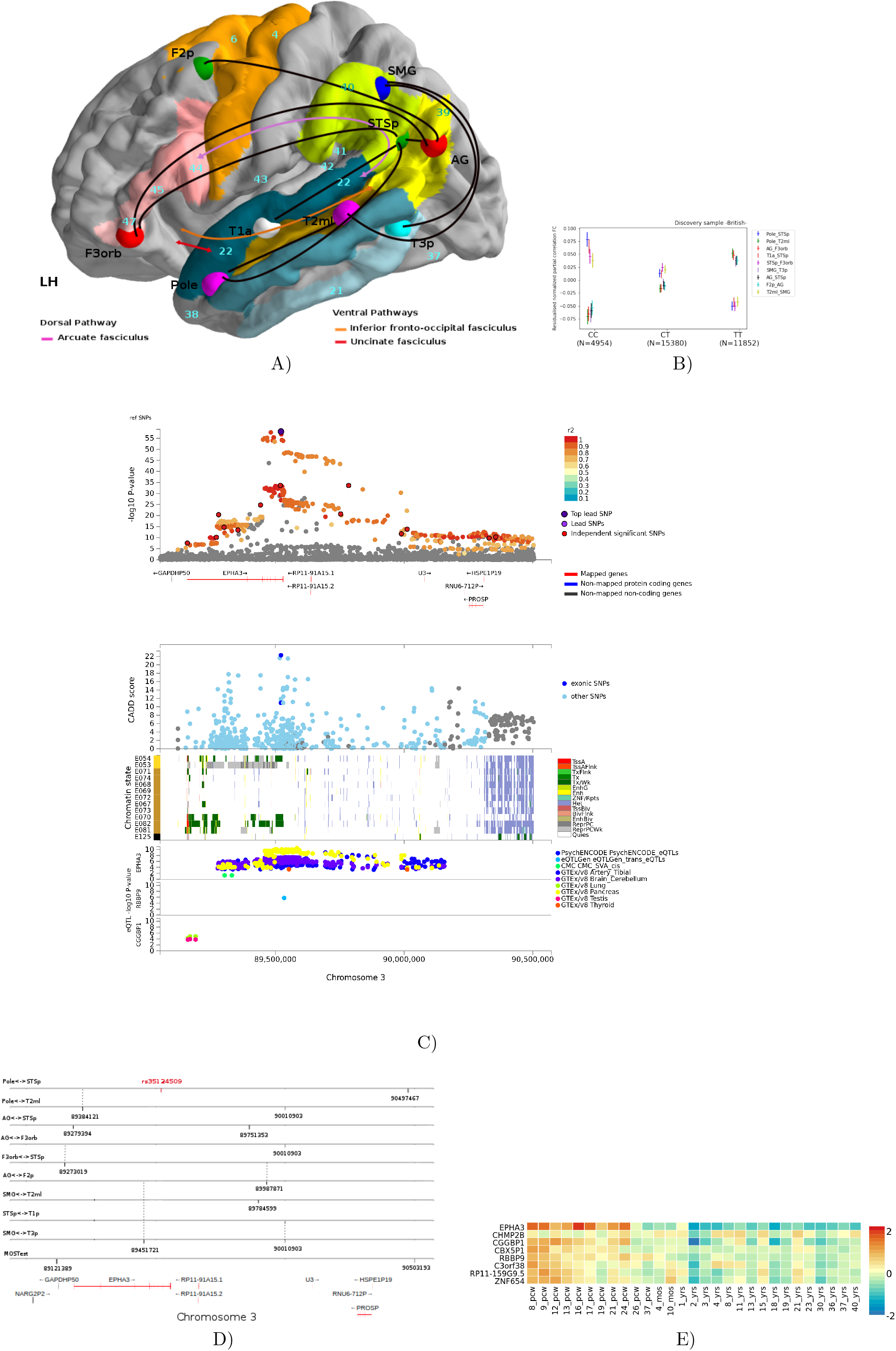
Main results for the 3p11.1 locus. A) The pairs of ROIs that forms the endpoints of the associated FCs reported as black bold lines. B) Effect sizes of the SNP rs35124509 for the nine connections: (AG↔F3orb), (Pole↔STSp), (Pole↔T2ml), (T1a↔STSp), (STSp↔F3orb), (SMG↔T3p), (AG↔STSp), (F2p↔AG) and (T2ml↔SMG) FCs. C) Locus Zoom of the genomic region identified by the mvGWAS. Chromatin state of the genomic region. Brain tissue name abbreviations are the following; E054:Ganglion Eminence derived primary cultured neurospheres, E053: Cortex derived primary cultured neurospheres, E071: Brain Hippocampus Middle, E074: Brain Substantia Nigra, E068: Brain Anterior Caudate, E069: Brain Cingulate Gyrus, E072: Brain Inferior Temporal Lobe, E067:Brain Angular Gyrus, E073: Brain Dorsolateral Prefrontal Cortex, E070: Brain Germinal Matrix, E082: Fetal Brain Female, E081: Fetal Brain Male, E125: NH-A Astrocytes Primary Cells. The state abbreviations are the following; TssA: active transcription start site (TSS), TssFlnk: Flanking Active TSS, TxFlnk: Transcription at gene 5‚ and 3‚, Tx: Strong transcription, TxWk: Weak transcription, EnhG: Genic enhancers, Enh: Enhancers, ZNF/Rpts: ZNF genes & repeats, Het: Heterochromatin, TssBiv: Bivalent/Poised TSS, BivFlnk: Flanking Bivalent TSS/Enh, EnhBiv: Bivalent Enhancer, ReprPC: Repressed PolyComb, ReprPCWk: Weak Repressed PolyComb, Quies: Quiescent/Low. Expression quantitative trait loci (eQTL) associations (data source: eQTLGen (Võsa et al., 2018), PsychENCODE(Wang et al., 2018), DICE (Schmiedel et al., 2018), BIOS QTL browser (Zhernakova et al., 2017), GTEx/v8 (Consortium et al., 2017), eQTLcatalogue). D) Overlap of the genomic region risk region identified from FUMA for MOSTest results and the nine FCs mentioned above. E) Gene expression from BrainSpan for the interesting genes prioritised by FUMA.

A locus in 10q23.33 was highlighted by the mvGWAS. At the univariate level, no endophenotype reached the genome-wide significance threshold for the leas SNP in this locus (rs11187838).

As a conclusion of the mvGWAS, we retained: *i)* a multifold link between two FCs and a locus in 15q4 region; and *ii)* a multifold link between nine FCs and a locus in 3p11.1 region. Such a multivariate approach has the advantage of leveraging the distributed nature of genetic effects and the presence of pleiotropy across endophenotypes. Loci respectively identified by MOSTest as associated with several FCs made clear that these SNPs have distributed effects, often with mixed directions, across regions and FCs. Fig. ??b shows the FCs associations with both 15q14, and 3p11.1 lead SNPs. The regional effects of all other lead SNPs can be appreciated in the Supplementary Fig. ??.

### 3.3. Downstream analyses

#### SNP-based genetic correlation of functional connectivity measures

The SNP-based genetic correlation analysis was estimated (using GCTA (Lee et al., 2012) software) for each pair of central FCs associated to 15q14 or 3p11.1 genetic loci, indicating overlapping genetic contributions among several FCs (Table SI6). For central endophenotypes associated with 3p11.1 locus, a negative genetic correlation between some FCs has been observed which indicates that variants can have antagonistic effects on the co-activations of these regions.

#### Validation of lead SNPs using diffusion imaging derived endophenotypes

We hypothesised that the genetic variants significantly associated with the language FCs could be associated with neuroanatomical tracts that support the information transmission between language areas. Therefore, we tested the potential associations between the hit SNPs with the average values of dMRI relevant white matter tracts :

3 white matter tracts to be tested with locus on 15q14: the white matter tracts linking the regions of the (Prec↔F3opd) consists of the *i)* arcuate anterior segment fasciculus (AF) dorsal pathway (Catani et al., 2005), *ii)* the frontal aslant tract (FAT) which is reported as connecting Broca’s region (BA44/45) with dorsal medial frontal areas including supplementary and pre-supplementary motor area (BA6) (Rojkova et al., 2016; Catani and Forkel, 2019) while the anatomical connectivity underlying the (PrecR↔RolS) FC endophenotype consists of the corpus callosum which interconnects both hemispheres.

5 white matter tracts to be tested with locus on 3p11.1: the nine central endophenotypes associated with the 3p11.1 locus, the anatomical connectivity underlying these connections consists of the *i)* inferior fronto-occipital fasciculus (IFOF) which connects the inferior frontal regions with the temporal and occipital cortex (Forkel et al., 2014b), *ii)* uncinate fasciculus (UF) which is reported to connect the anterior temporal lobe to the orbital region and part of the inferior frontal (Vigneau et al., 2006; Catani and De Schotten, 2008; Friederici, 2017; Catani and Forkel, 2019), and the *iii)* arcuate long/anterior/posterior segment fasciculus (AF) (Catani et al., 2005).

As the anterior segment of AF is tested with both loci, this yields a Bonferroni-corrected threshold of *p* = 6.94e−3(0.05/(3 * 9 + 5 * 9)) (Table SI7). The MO measured in the FAT and the OD measured in the anterior segment of AF are associated with the rs1440802 SNP with *p* = 3.33e−6 and *p* = 2.47e−65, respectively. The corpus callosum exhibits no significant association. The MO measured in the IFOF and UF is associated with the rs35124509 SNPs with *p* = 2.49e−7 and *p* = 2.12e−7, respectively. Both long and posterior segment of AF are associated with rs35124509 SNPs with *p* = 5.88e−6 (OD) and *p* = 3.90e−7(*L*3), while the anterior segment of AF exhibits no significant association (See Table SI7).

### 3.4. Functional annotations of genomic loci associated with language

#### Locus in 15q14 associated to (Prec↔F3opd) and (PrecR↔RolS) endophe-notypes

Four independent SNPs were identified in locus 15q14 (rs1440802, rs11629938, rs773225188, rs34680120) (Fig. 3c). Regarding eQTL annotations, we explored tissue-specific gene expression resources, including both brain tissues and blood - considered as a good proxy when brain tissues are not available (Qi et al., 2018). Significant results were obtained:

The four independant SNPs are cis-eQTL of *THBS1* gene in eQTLGen, BIOSQTL and GTEx/v8 . Additionally, rs34680120 is eQTL of *RP11-37C7.1* gene (*p*_*adj*_ < 1.02e−3) in PsychENCODE and eQTL of *CTD-2033D15.1* gene (*p*_*adj*_ < 6.0e−6) in BIOSQTL; see Fig.3c. Overall, the variants of this genomic risk region are found 72 times as eQTL of genes from different data sources. All eQTL associations are presented in more detail in Table SI8. Based on the human gene expression data from the Brainspan database, we found that *THBS1* gene has relatively high mRNA expression during early mid-prenatal to late prenatal stages, from 16 to 37 post-conceptional weeks; see Fig. 3e. Indirect predictions might be added from the following annotation. *RASGRP1*, identified by chromatin interaction mapping and which also appears to be under control of temporal expression during neurodevelopment, is reported as over-expressed in the perisylvian language areas (Johnson et al., 2009) and as up-regulated in the dorsal striatum (Cirnaru et al., 2020). Fig. 3 summarises these results, found by mvGWAS, associated to (Prec↔F3opd) and (PrecR↔RolS) FC endophenotypes. These pinpoint *THBS1* as the possible gene underlying this association signal.

#### Locus in 3p11.1 associated to semantic-language related endophenotypes

Fourteen independent SNPs were identified in locus 3p11.1 (Fig. 4c). The rs35124509 SNP is a non-synonymous variant within exon 16 of *EPHA3* protein-coding gene. The subregion around rs35124509 and rs113141104 has its chromatin state annotated as (weak) actively-transcribed states (Tx, TxWk) in the brain tissues, specifically in the Brain Germinal Matrix, the Ganglion Eminence derived primary cultured neurospheres, and in the Fetal Brain Female. Concerning the subregion around rs6551410, it has its chromatin state annotated as Weak transcription (TxWk) in the Fetal Brain Female, enhancer (enh) in the Brain Germinal Matrix and Repressed PolyComb (ReprPC) in both the Ganglion Eminence and Cortex derived primary cultured neurospheres. Additionally, the subregion around rs6551407 has its chromatin state annotated as Weak transcription (TxWk) in the Brain Germinal Matrix, Fetal Brain Male and Fetal Brain Male. Overall, this reveals a genomic region involved in fine regulation mechanisms of brain development.

Considering the rs35124509 SNP and variants in linkage disequilibrium (LD) with it in the genomic risk region, we scrutinised CADD and RDB scores, precise genomic positions and risk prediction, and we noticed some remarkable SNPs. We observed two exonic variants: *i)* The SNP rs1054750 (*LD*_*rs*35124509_ *r*^2^ > 0.99, *p*_*mvGW AS*_ = 6.65*e* − 34) is a synonymous variant within exon 16 of *EPHA3*. *ii)* the already mentioned non-synonymous lead SNP rs35124509 (*p*_*mvGW AS*_ = 8.95*e* − 59), the minor allele results in a substitution in the protein from tryptophan (W) residue (large size and aromatic) into an arginine (R) (large size and basic) at position 924 (W924R, p.Trp924Arg) in the Sterile Alpha Motif (SAM) domain. This SNP is not predicted to alter protein function (Polyphen-2 = “benign”) but is predicted to be potentially a regulatory element by several tools (RDB score = 3*a*, CADD = 22.3 - when CADD_*thresh*_ = 12.37 for deleterious effect as suggested by Kircher et al. (2014)) Moreover, we observed eight SNP (rs28623022, rs7650184, rs7650466, rs73139147, rs3762717, rs73139144, rs73139148, rs566480002) (*LD*_*rs*35124509_ *r*^2^ > 0.73, *p*_*mvGW AS*_ < 4.46*e* − 20) located in 3’-UTR of *EPHA3* which could affect its expression by modulating miRNA binding (Popp et al., 2016). The hit-SNP rs35124509 and the rest of highlighted SNPs act as eQTL for *EPHA3* in different tissues including brain cerebellum (*p*_FDR_ < 5e−2 in GTEx/v8 data source). The exhaustive eQTL associations are presented in Table SI8. Fig. 4 summarises the functional annotations in 3p11.1 associated to multiple FC endophenotypes in semantic component of language. These functional characterization supports *EPHA3* as a possible gene with a key role in language development in humans.

#### Locus in 10q23.33

Four independent SNPs were identified in locus 10q23.33 (rs11187838, rs17109875, rs11187844, rs20772180). The subregion around all four SNPs has its chromatin state annotated as (weak) actively-transcribed states (Tx, TxWk) in the brain tissues, specifically in the ganglion eminence and cortex derived primary cultured neurospheres, hippocampus (middle), substantia nigra, anterior caudate, angular gyrus, Dorsolateral/Prefrontal cortex, brain germinal matrix, fetal brain female/male and NH-A astrocytes primary cells. Two exonic variants are noteworthy: The rs2274224 (*LD*_*rs*11187838_ *r*^2^ > 0.99, *p*_*mvGW AS*_ = 5.04*e* − 14) and rs11187895 (*LD*_*rs*17109875_ *r*^2^ > 0.6, *p*_*mvGW AS*_ = 3.08*e* − 7) SNPs are nonsynonymous SNV within exon 19 of *PLCE1* and exon 11 of *NOC3L* and are both not predicted to alter protein function (Polyphen-2=”benign”) but are predicted to have a deleterious effect (CADD = 17.35, CADD = 19.24). Moreover, we observed three SNP (rs11187870, rs11187877, rs145707916) (*LD*_*rs*17109875_ *r*^2^ > 0.66, *p*_*mvGW AS*_ < 7.546*e* − 7) located in 3’-UTR of *PLCE1:NOC3L*. Regarding eQTL annotations, the variants in the 10q23.33 locus act as eQTL for *HELLS*, *NOC3L* and *PLCE1* genes in different brain tissues including brain cerebellum, brain cerebellar hemisphere, Brain nucleus accumbens basal ganglia, hippocampus (*p*_FDR_ < 5e−2 in GTEx/v8 data source). The exhaustive eQTL associations are presented in Table SI8. These functional characterisation highlight these three genes (*HELLS*, *NOC3L* and *PLCE1*) that may influence the FCs related to language processing in humans.

## 4. Discussion

In this study, we extracted individual language FC endophenotypes from the rsfMRI data of 32,186 participants from the UK Biobank cohort and conducted a multivariate genomewide association study. We found 4566 significantly associated SNPs distributed over 11 chromosomes. Three multivariate associations with lead SNPs were replicated in the non-British cohort, highlighting the robustness of these signals across different ancestries. Two functional connections, contributing in the *perceptual motor* interaction, associated with 15q14 locus located in the *RP11-624L4.1* antisense gene with modulatory effects on the expression of the *THBS1* gene. Multiple FCs in the *fronto-temporal semantic* language network were found to be associated with SNPs regulating *EPHA3* gene expression in 3p11.1 locus. Each lead SNP was found to be associated with the neuroanatomical white matter tracts that support each of these FCs.

### 4.1. Locus regulating THBS1 associated with the perceptual motor interactions process

A locus in 15q14 was associated with the precentral-opercularis FC (Prec↔F3opd) and the precentral-Rolandic FC endophenotypes (PrecR↔RolS). The L—R Prec regions in the ventral precentral gyrus are both associated with phonology language component and considered relevant for pharynx and tongue fine-movement coordination in the human and non-human primates (Vigneau et al., 2006; Kumar et al., 2016; Belyk and Brown, 2017). RolS in the dorsal Rolandic sulcus is attributed to the phonology component and matches the mouth primary motor area but also the perception of syllables (Vigneau et al., 2006; Wilson et al., 2004; Fadiga et al., 2002). F3opd in the dorsal pars opercularis (BA44/45) is associated with semantic/sentence processing. The motor theory of speech perception has been quite an old debate (Liberman and Mattingly, 1985; Galantucci et al., 2006; Flinker et al., 2015; Schwartz et al., 2008; Whalen, 2019). In this study, we report a locus in 15q14 (lead SNP rs1440802) associated with both this FC between the motor and Broca’s areas and the frontal aslant tract connecting directly (pre)supplementary motor area with the opercular part of inferior frontal gyrus (Vergani et al., 2014; Catani et al., 2012), in line with this perception–motor link.

SNPs in high LD with rs1440802 in the genomic region have been linked to several other structural features (surface area and cortical thickness) including primary motor cortex, primary somatosensory cortex (Elliott et al., 2018; van der Meer et al., 2020), supramarginal, and pars opercularis (van der Meer et al., 2020), supporting a common genetic influence of the sensory-motor interaction.

The lead SNP rs1440802 and SNPs in LD uncovered to be associated with both (Prec↔F3opd) and (PrecR↔RolS) are found to be eQTL of *THBS1* gene in the blood with high confidence. The thrombospondin-1 protein encoded by *THBS1* gene is a member of the thrombospondin family, a glycoprotein expressed in the extracellular matrix. It has been implicated in synaptogenesis (Christopherson et al., 2005) and regulates the differentiation and proliferation of neural progenitor cells (Lu and Kipnis, 2010), and has been involved in human neocortical evolution (Cáceres et al., 2003, 2007). Other members of the thrombospondin’s family, *THBS2* and *THBS4*, have been shown to be over-expressed in the adult human cerebral cortex compared to chimpanzees and macaques (Cáceres et al., 2007). Their increased expression suggests that human brain might display distinctive features involving enhanced synaptic plasticity in adulthood which may contribute to cognitive and linguistic abilities (Sherwood et al., 2008). From a developmental point of view, *THBS1* appears to be under control of temporal expression during development, as revealed by BrainSpan data (See Fig. 3e and Fig. SI4). *THBS1* expression was studied from the longitudinal transcriptomic profile resource of the developing human brain (18, 19, 21, 23 weeks of gestation) (Johnson et al., 2009). Its expression is reported as over-expressed in the neocortex, including the perisylvian language areas, compared to phylogenetically older parts of the brain such as the striatum, thalamus and cerebellum (Johnson et al., 2009). Thrombospondin-1 have been linked to Autism spectrum disorder (Lu et al., 2014), Alzheimer’s disease (Ko et al., 2015), and Schizophrenia (Park et al., 2012).

Taken together, these results indicate that *THBS1*, modulated by a lead SNP in the 15q14 locus, could be prioritised in the study of key genes playing a role in the functional connectivity part of the *perceptual motor* interaction required for language, and with the anatomical connectivity, support of their interactions.

### 4.2. Locus in EPHA3 associated with the fronto-temporal semantic network

A locus in 3p11.1 is found associated with nine fronto-parietal-temporal endophenotypes. The angular gyrus (AG) has been shown to activate during functional imaging tasks probing semantics and involved in conceptual knowledge (Vigneau et al., 2006). F3orb in the pars orbitalis in the inferior frontal gyrus is labelled semantic for its involvement in semantic retrieval in spoken and sign language (Rönnberg et al., 2004). It has also been associated with categorisation, association, and word generation tasks (Noppeney and Price, 2004; Booth et al., 2002; Gurd et al., 2002). The temporal pole region, located in the anterior temporal lobe, is associated with semantic and sentence processing (Vigneau et al., 2006) and the posterior superior temporal sulcus (pSTS) is reported to be implicated in syntactic complexity (Constable et al., 2004) but also process the semantic integration of complex linguistic material (Vigneau et al., 2006). Both pSTS and the angular gyrus overlap with the Geschwind’s territory (See Fig. 1a). The lateral/middle part of the middle temporal gyrus is devoted to verbal knowledge (Vigneau et al., 2006). These regions and their corresponding endophenotypes fit rather well with the *fronto-temporal semantic system* described in (Vigneau et al., 2006) facilitating the association of integrated input messages with internal knowledge. The anterior part of the superior temporal gyrus and the posterior part of the inferior temporal gyrus are phonological–semantic interface areas processing. (Vigneau et al., 2006) propose that these ones are transitional zones between the perception and semantic integration of language stimuli and are crucial during the development of language.

SNPs of this genomic region in high LD with the lead SNP rs35124509 have already been found associated with: rsfMRI ICA functional connectivity (edge 387, 383, 399, and ICA-features 3); see (Elliott et al., 2018). The ICA maps used for these FC estimations partially-overlap semantic language areas including the angular gyrus, the most anterior part of the STS, the anterior fusiform gyrus, the lateral-middle part of T2, the ventral part of the pars triangularis and the pars orbitalis of the left inferior frontal gyrus. Regarding cognitive traits, this locus was associated to intelligence (Savage et al., 2018). Finally, other SNPs, in strong LD with the lead SNP rs35124509, consistently act as an eQTL of *EPHA3* in brain tissues.

The ephrin type-A receptor 3 protein encoded by *EPHA3* gene belongs to the ephrin receptor family that can bind the ephrins subfamily of the tyrosine kinase protein family. EPH receptors and their ligands were found to play important roles in multiple developmental processes, including tissue morphogenesis, embryogenesis, neurogenesis, vascular network formation, neural crest cell migration, axon fasciculation, axon guidance, and topographic neural map formation (Pasquale, 2008; Gibson and Ma, 2011; Gerstmann and Zimmer, 2018). EPHA3 binds predominantly EFNA5 and plays a role in the segregation of motor and sensory axons during neuromuscular circuit development (Lawrenson et al., 2002). In (Johnson et al., 2009), *EPHA3* is reported as over-expressed in the fetal rhesus macaque monkey neocortex (NCTX) and especially in the occipital lobe compared to the other NCTX areas. Noticeably, its ligand *EFNA5* is over-expressed in perisylvian areas and is located in a human accelerated conserved non-coding sequence (haCNS704) (Johnson et al., 2009). EPH receptors have been linked to neurodevelopmental disorders, including schizophrenia (Zhang et al., 2010) and autism spectrum disorder (Casey et al., 2012). Moreover, in (Rudov et al., 2013), *EPHA3* is found *in silico*, as putative gene implicated in dyspraxia, dyslexia and specific language impairment (SLI). Finally, we observed that *EPHA3* is expressed in the human brain, in a consistent manner across developmental stages from early prenatal to late-mid prenatal (8-24pcw, BrainSpan; see Fig. 4e and Fig. SI4).

Taken together, these results indicate that *EPHA3* in the 3p11.1 locus, could be prioritised in the study of key genes playing a role in the *fronto-temporal semantic* network, and with the anatomical connectivity support of this network.

### 4.3. Locus in PLCE1, NOC3L and HELLS

A locus in 10q23.33 was highlighted by the mvGWAS. At the univariate level, no endophenotype reached the genome-wide significance threshold. But looking at the suggestive threshold *p* = 1*e* − 5, we pinpoint putative ‘central’ endophenotypes to aid interpretation of the processes underlying this association signal. Two bilateral fronto-temporal endophenotypes were the most associated to rs11187838: the precentral-Rolandic FC endophenotypes (PrecR↔RolS, *p* = 1.85*e* − 07) and the right anterior part of the superior temporal gyrus (T1aR) overlapping Heschl’s gyrus (T1a/HeschlR) and its homotopic areas of LH primary auditory regions (T1a↔T1a/HeschlR, *p* = 9.61*e* − 06). All these regions participate in an elementary audio–motor loop involved in both comprehension and production of syllables forming a bilateral fronto-temporal network activated by the auditory representation of speech sounds (Vigneau et al., 2006, 2011). SNPs of this genomic region in high LD with the lead rs11187838 act as an eQTL of *HELLS*, *NOC3L*, *PLCE1* genes in multiple brain tissues (Supplementary Table SI8). The *HELLS* gene encodes the lymphoid-specific helicase (Lsh), a member of the SNF2 helicase family of chromatin remodeling proteins. Patients with a genetic mutation of *HELLS* present psychomotor retardation including slow cognitive, motor development and psychomotor impairment (Thijssen et al., 2015). The Lsh protein might play a role as epigenetic regulator in neural cells (Han et al., 2017). Finally, we observed that the three genes (*NOC3L*, *PLCE1*, *HELLS*) are expressed in the human brain, across developmental stages from early prenatal to early mid prenatal (8-17 pcw, BrainSpan).

Taken together, these results indicate that the three highlighted genes (*PLCE1*, *NOC3L* and *HELLS*) in the 10q23.33 locus, as potential candidates in the study of key genes playing a role in the *bilateral fronto-temporal auditory-motor* network.

### 4.4. Limitations

The lack of a large, age-matched replication sample represents one major limitation of the present study in the sense that we could not reproduce all our results. Additionally, we observed that the three associations replicated in the non-British sample were not the three most significant ones. For example, rs2279829 on chr3 was found associated with *p* = 7.57e−21 but was not replicated in the non-British cohort, while rs11187838 on chr10 found associated with *p* = 4.29e−14 was replicated in the non-British cohort with *p* = 2.92e−2. This suggests that some lead SNPs found associated with language FCs with a lower p value than *p* = 4.29e−14 but not replicated in the non-British sample might be specific to the British ancestry. Nevertheless, the sample size of the discovery sample was an order of magnitude larger than the replication sample, making it difficult to compare these different results. Although multivariate methods have shown to substantially increase statistical power and gene discovery compared to univariate approaches, the results are less straightforward to interpret. We have addressed this issue by assessing each of the prioritized loci at the univariate level, to pinpoint at central endophenotypes that are contributing the most to the multivariate signal. Moreover, as such a complex trait as language may be driven by a lot of interacting genes, a multivariate approach on the SNPs side is highly desired to uncover relevant gene pathways in language development and processing. Compared to structural endophenotypes, the FCs have low amplitude which hinders the study in terms of statistical power. This observation constitute a third limitation that is somehow surpassed when working on large scale cohorts and using multivariate approaches. Another potential limitation is the UK Biobank dataset in which this study is based. It should be noted that the UKB constitutes a relatively old sample. Future studies in other developmental stages (i.e. children, adolescent, young-adult) will inform us whether the observed associations are stable across development, or whether they reflect some age-related specificity.

### 4.5. Conclusions

Thanks to imaging-genetics modern approaches that allow us to increase in statistical power and circumvent the small effect sizes, we could, at a certain level, shed lights into the genetic architecture of language functional connectivity by highlighting potential key genes related to language processing with -nearly- no recruitment bias. The neurobiology of language, but also many other neuroscience fields, could highly benefit from this type of methodology.

## Abbreviations

rsfMRI: resting-state functional magnetic resonance imaging
FC: functional connectivity
GWAS: genome-wide association study
mvGWAS: multivariate GWAS
SNP: single nucleotide polymorphism
eQTL: expression quantitative trait locus

## 5. Declaration of competing interest

The authors declare that they have no conflict of interest

## 6. Data Availability

The data used in this study are available via the UK Biobank, https://www.ukbiobank.ac.uk/.

## 7. Ethics statement

UK Biobank dataset: informed consent is obtained from all UK Biobank participants; ethical procedures are controlled by a dedicated Ethics and Guidance Council (http://www.ukbiobank.ac.uk/et that has developed with UK Biobank an Ethics and Governance Framework (given in full at http://www.ukbiobank.ac.uk/wp-content/uploads/2011/05/EGF20082.pdf), with IRB approval also obtained from the North West Multi-center Research Ethics Committee.

## 8. Code availability

This study used openly available software and codes, specifically GCTA (https://cnsgenomics.com/software/gcta/#GREML), PLINK (http://zzz.bwh.harvard.edu/plink/), MOSTest (https://github.com/precimed/mostest), and FUMA (https://fuma.ctglab.nl/). The anatomical connectivity atlas used is available at (http://www.bcblab.com/BCB/Atlas_of_Human_Brain_Connections.html).

## 9. Acknowledgements

This research was conducted using the UK Biobank resource under application #64984. This project was supported by the Marie Sk-lodowska-Curie programme awarded to Stephanie J. Forkel (Grant agreement No. 101028551). Amaia Carrion-Castillo was supported by a Juan de la Cierva fellowship from the Spanish Ministry of Science and Innovation, and a Gipuzkoa Fellows fellowship from the Basque Government.

## Supplementary tables (separate Excel file)

Table SI1: Overview of the regions obtained from the meta-analysis. Each ROIs is characterized by their abbreviated anatomical label defined by Vigneau et al. (2006, 2011) and is labelled according to the language component they belong to : phonology, semantic, and syntax.

Table SI2: Heritabilities of the 300 brain functional connectivity, estimated using the genotyped SNPs information using genome-based restricted maximum likelihood (GREML) (Yang et al., 2010) as implemented in GCTA (Yang et al., 2011) software (version 1.93.2beta). A 0.05 threshold on False Discovery Rate (FDR) adjusted p-values was applied to account for multiple testing.

Table SI3: SNPs associated with the 142 heritable functional connectivity measures using MOSTest (van der Meer et al., 2020) at the genome-wide significance threshold *p* = 5e−8.

Table SI4: Replication of the 20 lead SNP association using an independent non-British replication dataset (N=4,754) using MOSTest (van der Meer et al., 2020). We considered the nominal significance threshold *pvalue* < 0.05.

Table SI5: For each of the 20 lead SNPs identified in the multivariate GWAS, the corresponding univariate summary statistics for FCs identified as central FC (threshold on the genome-wide significance threshold *p* = 5e−8).

Table SI6: The SNP-based genetic correlation analysis was estimated (using GCTA (Lee et al., 2012) software, version 1.93.2beta) for each pair of central FCs associated to 15q14 or 3p11.1 genetic loci.

Table SI7: Univariate associations of 2 lead SNPs (rs1440802 on 15q14, rs35124509 on 3p11.1) using PLINK 1.9 (Purcell et al., 2007) with diffusion MRI indices on the following 7 white matter tracts: the corpus callosum, the left frontal aslant tract, the left arcuate anterior/long/posterior segment, the left inferior fronto-occipital fasciculus, the left uncinate tract. Significant results were considered at the Bonferroni-corrected threshold *p* = 6.94e−3(0.05/(3 ∗ 9 + 5 ∗ 9)).

Table SI8: eQTLs association, performed by FUMA, between the SNPs in the three replicated genomic risk loci and all mapped genes in the following databases : GTEx/v8 (Consortium et al., 2017), PsychENCODE (Wang et al., 2018), eQTLGen (Võsa et al., 2018), eQTLcatalogue, DICE (Schmiedel et al., 2018), BIOSQTL (Zhernakova et al., 2017). A 0.05 threshold on False Discovery Rate (FDR) adjusted p-values was applied to account for multiple testing.

Table SI9: Center of mass of the language processing regions of interests retained in both left and right hemispheres. Each ROIs is characterised by their abbreviated anatomical label defined by (Vigneau et al., 2006, 2011) and their center of mass MNI stereotactic coordinates (x, y, z, in mm).

## 11. Supplementary figures

Figure SI1: Locus Zoom of the significant loci identified by the multiariate GWAS for functional connectivity.

Figure SI2: **Genomic loci, eQTL associations and chromatin interactions identified via multivariate GWAS for functional connectivity.** Circos plot representing the genomic risk loci, and the genes associated with the loci by chromatin interactions and eQTLs. From outer layer to inner layer: Manhattan plot. Genomic risk loci are in blue. Genes mapped by chromatin interaction are in orange. Genes mapped by eQTL are in green. Genes mapped by both are in red. Chromatin interaction and eQTLs links follows the same color coding presented above.

Figure SI3: **Regional effects.** Circle plot illustrating the lead SNPs identified from the multivariate GWAS for functional connectivity. Z-values from the univariate GWAS for each FCs are mapped. The absolute Z-values scaling is clipped at 8 (*p* = 1.2e 15). Positif effects of carrying the minor allele are shown in red, and negative in blue.

Figure SI4: **Functional annotation of both genomic risk loci 15q14 and 3p11.1.** A) Gene expression heatmap constructed with GTEx/v8 (54 tissue types) and B) BrainSpan 29 different ages of brain samples. (Average of normalized expression per label).

## Notes

### Competing Interest Statement

The authors have declared no competing interest.

