## Supplemental Tables for "The genetic architecture of language functional connectivity"

| Table | Contents |
| --- | --- |
| Table S1 | Overview of the regions obtained from the meta-analysis. |
| Table S2 | Heritabilities of of the 300 brain functional connectivity, estimated using the genotyped SNPs information using genome-based restricted maximum likelihood (GREML) as implemented in GCTA. |
| Table S3 | SNPs associated with the 142 heritable functional connectivity measures using MOSTest at the genome-wide significance threshold p = 5e−8. |
| Table S4 | Replication of the 20 lead SNP association using an independent non-British replication dataset (N=4,754) using MOSTest. |
| Table S5 | For each of the 20 lead SNPs identified in the multivariate GWAS, the corresponding univariate summary statistics for FCs identified as central FC (threshold on the genome-wide significance threshold p value: 5e-8). |
| Table S6 | The SNP-based genetic correlation analysis was estimated using GCTA software, for each pair of central FCs associated to 15q14 or 3p11.1 genetic loci. |
| Table S7 | Univariate associations of 2 lead SNPs with diffusion MRI indices. |
| Table S8 | eQTLs association, performed by FUMA, between the SNPs in the three replicated genomic risk loci and all mapped genes in differents databases. |
| Table S9 | Center of mass of the language processing regions of interests retained in both left and right hemispheres. |

| Hemisphere | Regions of interest | Phonology | Semantic | Sentence |
|---|---|---|---|---|
| Left hemisphere | Planum temporale (PT) | x | | |
| | **Rolandic sulcus (RolS)** | **x** | | |
| | Superior temporal gyrus (T1) | x | | |
| | **Supramarginalis gyrus (SMG)** | **x** | | |
| | **Precentral gyrus (Prec)** | **x** | | |
| | Dorsal part of the pars triangularis of the inferior frontal gyrus (F3td) | x | | |
| | Anterior fusiform gyrus (Fusa) | | x | |
| | Opercular part of the inferior frontal gyrus (PrF3op) | | x | |
| | **Angular gyrus (AG)** | | **x** | |
| | **Pars orbitaris of the left inferior frontal gyrus (F3orb)** | | **x** | |
| | **Posterior part of the middle frontal gyrus (F2p)** | | | **x** |
| | **Posterior ending of the superior temporal gyrus (STSp)** | | | **x** |
| | Posterior part of the middle temporal gyrus (T2p) | | | x |
| | **Temporal pole (Pole)** | | **x** | **x** |
| | **Lateral/middle part of T2 (T2ml)** | | **x** | **x** |
| | **Dorsal part of the pars opercularis (F3opd)** | | **x** | **x** |
| | Ventral part of the pars triangularis (F3tv) | | x | x |
| | **Posterior part of the left inferior temporal gyrus (T3p)** | **x** | **x** | |
| | **Anterior part of T1 (T1a)** | **x** | **x** | **x** |
| Right hemisphere | Anterior part of the middle frontal gyrus (F2antR) | x | | |
| | **Precentral gyrus (PrecR)** | **x** | | |
| | Anterior part of tT1 overlapping the Heschl's gyrus (T1a/HeschlR) | x | | |
| | Superior temporal gyrus (T1R) | x | | |
| | Middle part of the STS (T2pR) | | | x |
| | Upper part of the pars opercularis of F3 (F3opdR) | | x | x |

Note: Bold phenotypes represents FCs with significant heritability after false discovery rate correction at $p<0.05$.

| Phenotype | P value | h² | SE | FDR P value |
|---|---|---|---|---|
| **F3opd_SMG** | 3.8858e-16 | 0.143085 | 0.018056 | 1.16574e-13 |
| **AG_F3tv** | 9.992e-15 | 0.133426 | 0.0179 | 1.4988e-12 |
| **T2ml_STSp** | 5.0371e-13 | 0.123611 | 0.017817 | 5.0371e-11 |
| **STSp_T1R** | 7.5961e-13 | 0.123237 | 0.017888 | 5.6970750000000005e-11 |
| **Pole_STSp** | 1.0457e-12 | 0.12179 | 0.017839 | 6.2742e-11 |
| **Pole_T2ml** | 2.1435e-12 | 0.118543 | 0.017727 | 1.07175e-10 |
| **PT_T1** | 4.9045e-12 | 0.118866 | 0.017877 | 2.1019285714285713e-10 |
| **F2p_STSp** | 1.3203e-11 | 0.115631 | 0.017798 | 4.951125e-10 |
| **T2ml_SMG** | 2.7522e-11 | 0.11216 | 0.017705 | 8.2566e-10 |
| **T2ml_AG** | 2.6209e-11 | 0.114751 | 0.017844 | 8.2566e-10 |
| **AG_F2antR** | 3.8704e-11 | 0.113222 | 0.017836 | 1.0555636363636364e-09 |
| **PrF3op_SMG** | 5.1329e-11 | 0.114377 | 0.017961 | 1.283225e-09 |
| **F3tv_STSp** | 9.2723e-11 | 0.111882 | 0.01789 | 2.1397615384615387e-09 |
| **RolS_PrecR** | 2.2724e-10 | 0.110825 | 0.017981 | 4.869428571428571e-09 |
| **PrF3op_F3tv** | 9.4977e-10 | 0.106623 | 0.017983 | 1.8995399999999997e-08 |
| **F3tv_SMG** | 1.9262e-09 | 0.103729 | 0.017911 | 3.611625e-08 |
| **F2p_SMG** | 2.8369e-09 | 0.099209 | 0.017622 | 5.0062941176470594e-08 |
| **STSp_F3orb** | 3.7081e-09 | 0.101059 | 0.017802 | 6.180166666666667e-08 |
| **SMG_F2antR** | 1.317e-08 | 0.096583 | 0.017754 | 2.0794736842105263e-07 |
| **T1_F3tv** | 1.4007e-07 | 0.091235 | 0.017907 | 2.10105e-06 |
| **Fusa_AG** | 1.794e-07 | 0.0866 | 0.01758 | 2.562857142857143e-06 |
| **F3opd_F3td** | 1.8925e-07 | 0.089293 | 0.01785 | 2.580681818181818e-06 |
| **T2p_T3p** | 3.8638e-07 | 0.084325 | 0.017579 | 5.0397391304347825e-06 |
| **Prec_F3opd** | 4.8207e-07 | 0.084597 | 0.017645 | 6.025875e-06 |
| **T2ml_F3tv** | 5.6358e-07 | 0.083375 | 0.017642 | 6.76296e-06 |
| **SMG_F3orb** | 6.2518e-07 | 0.084135 | 0.017708 | 7.213615384615385e-06 |
| **Prec_SMG** | 6.5433e-07 | 0.083548 | 0.017631 | 7.270333333333333e-06 |
| **F3tv_F3orb** | 1.0351e-06 | 0.083137 | 0.017748 | 1.1090357142857142e-05 |
| **SMG_F3td** | 1.7991e-06 | 0.081891 | 0.017835 | 1.7991e-05 |
| **T2ml_F3td** | 1.7851e-06 | 0.080841 | 0.01772 | 1.7991e-05 |
| **AG_T1a** | 2.5503e-06 | 0.078469 | 0.017596 | 2.4680322580645163e-05 |
| **F3opd_T2ml** | 2.6626e-06 | 0.078433 | 0.017624 | 2.4961875e-05 |
| **AG_SMG** | 2.877e-06 | 0.080434 | 0.017847 | 2.538529411764706e-05 |
| **F3tv_F2antR** | 2.8001e-06 | 0.078039 | 0.017578 | 2.538529411764706e-05 |
| **Pole_T1a** | 3.0043e-06 | 0.077865 | 0.017622 | 2.5751142857142858e-05 |
| **T1_PrF3op** | 4.7379e-06 | 0.076509 | 0.017603 | 3.94825e-05 |
| **AG_STSp** | 4.9848e-06 | 0.076665 | 0.017643 | 4.0417297297297304e-05 |
| **F3opd_STSp** | 5.2766e-06 | 0.077101 | 0.017715 | 4.165736842105263e-05 |
| **PrF3op_T3p** | 6.4761e-06 | 0.074129 | 0.017521 | 4.9816153846153846e-05 |
| **PrF3op_F2antR** | 7.0454e-06 | 0.074811 | 0.017603 | 5.2840500000000003e-05 |
| **T2ml_F3orb** | 9.0511e-06 | 0.07515 | 0.017779 | 6.622756097560976e-05 |
| **T1a_STSp** | 1.067e-05 | 0.072739 | 0.017536 | 7.621428571428571e-05 |
| **F3opdR_F2antR** | 1.1282e-05 | 0.073398 | 0.017628 | 7.871162790697674e-05 |
| **Pole_F3opd** | 1.5498e-05 | 0.073106 | 0.017736 | 0.00010566818181818181 |
| **AG_T3p** | 1.8595e-05 | 0.070762 | 0.01753 | 0.00012396666666666668 |
| **STSp_T2pR** | 2.3762e-05 | 0.070433 | 0.017629 | 0.0001549695652173913 |
| **F3opd_PrF3op** | 3.2513e-05 | 0.069213 | 0.017614 | 0.00020752978723404255 |

| | | | | |
|---|---|---|---|---|
| **T1_T1a/HeschlR** | 3.6045e-05 | 0.06916 | 0.017661 | 0.00022528124999999998 |
| **Prec_AG** | 4.1421e-05 | 0.06933 | 0.017751 | 0.00025359795918367345 |
| **F3tv_T2pR** | 4.251e-05 | 0.067067 | 0.017501 | 0.0002541588235294118 |
| **T1a_T1a/HeschlR** | 4.3207e-05 | 0.068425 | 0.017662 | 0.0002541588235294118 |
| **PrF3op_AG** | 4.5151e-05 | 0.067181 | 0.01758 | 0.00026048653846153847 |
| **T1_T1a** | 5.0854e-05 | 0.067867 | 0.01767 | 0.0002878528301886792 |
| **Prec_F3tv_RolOp** | 5.5864e-05 | 0.066566 | 0.017584 | 0.00031035555555555556 |
| **Pole_AG** | 5.7086e-05 | 0.065955 | 0.017525 | 0.0003113781818181818 |
| **F3opd_AG** | 6.102e-05 | 0.064642 | 0.017401 | 0.00032689285714285714 |
| **F3tv_F3opdR** | 6.2863e-05 | 0.065788 | 0.017518 | 0.0003308578947368421 |
| **F3orb_F2antR** | 7.8212e-05 | 0.066301 | 0.017712 | 0.0003846491803278689 |
| **PT_T1a/HeschlR** | 7.7413e-05 | 0.066029 | 0.017663 | 0.0003846491803278689 |
| **SMG_T3p** | 7.7787e-05 | 0.066515 | 0.017725 | 0.0003846491803278689 |
| **T1a_SMG** | 7.5932e-05 | 0.065428 | 0.017581 | 0.0003846491803278689 |
| **Prec_T1** | 8.2673e-05 | 0.065155 | 0.017575 | 0.00040003064516129033 |
| **Prec_T2p** | 8.4773e-05 | 0.064923 | 0.017579 | 0.0004036809523809524 |
| **PrF3op_STSp** | 9.1453e-05 | 0.064689 | 0.017576 | 0.0004286859375 |
| **T2ml_PrF3op** | 0.00013238 | 0.064308 | 0.01774 | 0.0006109846153846153 |
| **Fusa_T2p** | 0.00013858 | 0.063197 | 0.017621 | 0.0006299090909090909 |
| **F3opd_T3p** | 0.00017382 | 0.062951 | 0.017724 | 0.0007782985074626865 |
| **T2ml_T1R** | 0.00019324 | 0.061051 | 0.017536 | 0.000852529411764706 |
| **F2p_F3tv** | 0.00020405 | 0.060978 | 0.017568 | 0.0008836714285714286 |
| **T2ml_F3opdR** | 0.00020619 | 0.062341 | 0.017732 | 0.0008836714285714286 |
| **F3opd_F2p** | 0.0002127 | 0.059706 | 0.017397 | 0.0008987323943661971 |
| **T2ml_T1a** | 0.00021766 | 0.062438 | 0.017782 | 0.0009069166666666668 |
| **Pole_T1R** | 0.0002334 | 0.061741 | 0.017731 | 0.0009591780821917809 |
| **SMG_T2pR** | 0.00024161 | 0.059208 | 0.017447 | 0.00097036 |
| **AG_F3orb** | 0.00024259 | 0.060372 | 0.017564 | 0.00097036 |
| **AG_T2p** | 0.00026321 | 0.059726 | 0.017516 | 0.0010389868421052631 |
| **STSp_F3opdR** | 0.00061509 | 0.055036 | 0.017452 | 0.0023964545454545454 |
| **F2p_T2ml** | 0.00069194 | 0.05667 | 0.017744 | 0.0026613076923076924 |
| **T2p_RolS** | 0.00070926 | 0.05567 | 0.017608 | 0.002693392405063291 |
| **T3p_F3orb** | 0.00072091 | 0.054682 | 0.017478 | 0.0027034125 |
| **T1a/HeschlR_T1R** | 0.00081139 | 0.055158 | 0.017635 | 0.0030051481481481483 |
| **F3tv_RolS** | 0.0010102 | 0.054803 | 0.017743 | 0.0036513253012048193 |
| **F3orb_T2pR** | 0.0010072 | 0.051674 | 0.017288 | 0.0036513253012048193 |
| **T2ml_T2pR** | 0.0010751 | 0.0531 | 0.017536 | 0.0038396428571428574 |
| **F3tv_T1R** | 0.0011414 | 0.052596 | 0.017508 | 0.004028470588235294 |
| **F3td_T3p** | 0.001177 | 0.053444 | 0.017665 | 0.004072758620689655 |
| **T2p_F3opdR** | 0.0011811 | 0.050276 | 0.017205 | 0.004072758620689655 |
| **F3opd_F3orb** | 0.0013438 | 0.051631 | 0.01748 | 0.004581136363636364 |
| **F2p_PrF3op** | 0.0014604 | 0.051168 | 0.017478 | 0.004922696629213482 |
| **F3tv_T2p** | 0.0015213 | 0.051726 | 0.017601 | 0.005015274725274725 |
| **T1_RolS** | 0.0015125 | 0.051945 | 0.01762 | 0.005015274725274725 |
| **Prec_F3opdR** | 0.0015897 | 0.051569 | 0.017608 | 0.005183804347826087 |
| **T1_AG** | 0.001621 | 0.050772 | 0.017481 | 0.005229032258064516 |
| **STSp_F3td** | 0.0017185 | 0.051715 | 0.017699 | 0.0053703125 |
| **F3tv_T1a/HeschlR** | 0.0016959 | 0.051083 | 0.017605 | 0.0053703125 |
| **STSp_F2antR** | 0.0017051 | 0.050964 | 0.017571 | 0.0053703125 |
| **F3tv_T3p** | 0.0021128 | 0.049782 | 0.017543 | 0.006534432989690722 |

| | | | | |
|---|---|---|---|---|
| **Fusa_T3p** | 0.0023718 | 0.04862 | 0.017463 | 0.007260612244897958 |
| **Prec_Fusa** | 0.002608 | 0.049563 | 0.017718 | 0.007903030303030302 |
| **F3opdR_T2pR** | 0.0029553 | 0.046875 | 0.017387 | 0.0088659 |
| **F3tv_RolOp_F3td** | 0.0031344 | 0.047305 | 0.017509 | 0.009218823529411765 |
| **F3opdR_T1R** | 0.0031071 | 0.047125 | 0.017465 | 0.009218823529411765 |
| **SMG_F3opdR** | 0.0034507 | 0.046359 | 0.017414 | 0.010050582524271845 |
| **F3orb_F3opdR** | 0.0038804 | 0.043742 | 0.017142 | 0.011193461538461538 |
| **Fusa_STSp** | 0.0041801 | 0.046207 | 0.017613 | 0.011830471698113207 |
| **T1_T1R** | 0.0041501 | 0.045576 | 0.017464 | 0.011830471698113207 |
| **F3td_F2antR** | 0.0046048 | 0.043687 | 0.017254 | 0.012910654205607476 |
| **PrF3op_T1R** | 0.0046951 | 0.045926 | 0.017672 | 0.013041944444444443 |
| **F2antR_T2pR** | 0.0048831 | 0.045285 | 0.017595 | 0.013439724770642201 |
| **T1_F3orb** | 0.0049719 | 0.044522 | 0.017494 | 0.0135597272727272727 |
| **Prec_RolS** | 0.0056496 | 0.044567 | 0.01762 | 0.01526918918918919 |
| **F2p_AG** | 0.0059333 | 0.044539 | 0.017677 | 0.015892767857142855 |
| **T1a_F3orb** | 0.0060002 | 0.044457 | 0.017681 | 0.015929734513274337 |
| **T1a_T1R** | 0.0065148 | 0.042695 | 0.017421 | 0.017144210526315793 |
| **PT_T1R** | 0.0068357 | 0.042334 | 0.017404 | 0.017832260869565218 |
| **T2pR_PrecR** | 0.0078292 | 0.041986 | 0.01751 | 0.020247931034482757 |
| **Pole_F2p** | 0.0081415 | 0.041209 | 0.017393 | 0.020875641025641024 |
| **F3opd_F2antR** | 0.0083959 | 0.041398 | 0.017466 | 0.02134550847457627 |
| **Prec_F3td** | 0.0091195 | 0.041614 | 0.017623 | 0.022990336134453778 |
| **SMG_T2p** | 0.0097762 | 0.040337 | 0.01747 | 0.0244405 |
| **PT_AG** | 0.0099548 | 0.040569 | 0.017526 | 0.024681322314049587 |
| **PrF3op_F3opdR** | 0.010361 | 0.040279 | 0.017516 | 0.025477868852459017 |
| **PrF3op_F3td** | 0.011451 | 0.039057 | 0.017414 | 0.027704032258064514 |
| **Fusa_T1a** | 0.011411 | 0.039653 | 0.017505 | 0.027704032258064514 |
| **T2pR_T1R** | 0.013512 | 0.037623 | 0.01734 | 0.0324288 |
| **T2p_F2antR** | 0.013944 | 0.0387 | 0.017595 | 0.0332 |
| **T2ml_T2p** | 0.014149 | 0.036146 | 0.01709 | 0.03342283464566929 |
| **T2ml_T1** | 0.014456 | 0.037889 | 0.017498 | 0.03388125 |
| **F2antR_T1a/HeschlR** | 0.014975 | 0.038576 | 0.017672 | 0.03482558139534884 |
| **F3opd_T1R** | 0.015998 | 0.03773 | 0.017587 | 0.03691846153846153 |
| **F3td_T1R** | 0.016956 | 0.03663 | 0.017451 | 0.03883053435114503 |
| **F3opd_T1** | 0.017388 | 0.037128 | 0.017578 | 0.03922105263157895 |
| **F3opd_T1a** | 0.017311 | 0.036128 | 0.017353 | 0.03922105263157895 |
| **F2p_T2p** | 0.017929 | 0.036495 | 0.017494 | 0.04013955223880597 |
| **Prec_F2antR** | 0.0185 | 0.035961 | 0.017444 | 0.04111111111111111 |
| **Fusa_SMG** | 0.019285 | 0.035685 | 0.017409 | 0.042407299270072994 |
| **AG_PrecR** | 0.019366 | 0.036554 | 0.01762 | 0.042407299270072994 |
| **SMG_T1R** | 0.019518 | 0.035105 | 0.017327 | 0.042430434782608696 |
| **F2antR_T1R** | 0.020186 | 0.035751 | 0.017503 | 0.04356690647482014 |
| **F2p_T2pR** | 0.021678 | 0.035227 | 0.017513 | 0.04645285714285714 |
| **T1_SMG** | 0.022995 | 0.034586 | 0.017456 | 0.04892553191489362 |
| **F2p_T1** | 0.023312 | 0.034744 | 0.017515 | 0.04925070422535211 |
| STSp_SMG | 0.025033 | 0.034228 | 0.01752 | 0.052516783216783214 |
| F2p_T1a | 0.025393 | 0.034026 | 0.017483 | 0.052902083333333336 |
| F3tv_T1a | 0.025701 | 0.033396 | 0.017344 | 0.053174482758620696 |
| Prec_T3p | 0.026262 | 0.033914 | 0.017513 | 0.053963013698630144 |
| T2ml_F2antR | 0.028099 | 0.032696 | 0.017332 | 0.057344897959183676 |

| | | | | |
|---|---|---|---|---|
| Pole_F3orb | 0.029043 | 0.032337 | 0.017317 | 0.05887094594594595 |
| F3opd_T2pR | 0.032226 | 0.032059 | 0.017455 | 0.06488456375838926 |
| Prec_F3orb | 0.036763 | 0.030209 | 0.01723 | 0.07347417218543047 |
| PrF3op_T2pR | 0.036982 | 0.030994 | 0.017438 | 0.07347417218543047 |
| RolS_F3opdR | 0.037675 | 0.030696 | 0.017406 | 0.07435855263157895 |
| PT_RolS | 0.03822 | 0.030529 | 0.017403 | 0.07467290322580644 |
| AG_T1R | 0.038581 | 0.030888 | 0.017498 | 0.07467290322580644 |
| T1a_RolS | 0.038461 | 0.031088 | 0.017548 | 0.07467290322580644 |
| PrF3op_T2p | 0.040816 | 0.030008 | 0.017388 | 0.0784923076923077 |
| Pole_PT | 0.041623 | 0.028871 | 0.017104 | 0.0795343949044586 |
| PT_F2antR | 0.04302 | 0.029028 | 0.017209 | 0.08168354430379747 |
| T1_PrecR | 0.043377 | 0.02975 | 0.017435 | 0.0818433962264151 |
| Pole_PrF3op | 0.044381 | 0.029334 | 0.017387 | 0.083214375 |
| RolS_F2antR | 0.044847 | 0.029525 | 0.017457 | 0.08356583850931676 |
| T1_T2p | 0.047643 | 0.029116 | 0.017487 | 0.08822777777777778 |
| Pole_T1 | 0.048333 | 0.028583 | 0.017357 | 0.08895644171779142 |
| PrF3op_PrecR | 0.05013 | 0.028004 | 0.017274 | 0.09170121951219513 |
| F3opd_RolS | 0.051655 | 0.027777 | 0.017282 | 0.09391818181818182 |
| Pole_F3tv | 0.052115 | 0.028232 | 0.017443 | 0.09418373493975904 |
| T1_T2pR | 0.054995 | 0.027845 | 0.017463 | 0.0987934131736527 |
| Pole_T2p | 0.058384 | 0.027374 | 0.017463 | 0.10425714285714285 |
| F2p_F2antR | 0.059932 | 0.02652 | 0.017274 | 0.10638816568047336 |
| T1_F2antR | 0.061803 | 0.026279 | 0.017294 | 0.10897894736842105 |
| PT_T2ml | 0.062118 | 0.027032 | 0.017531 | 0.10897894736842105 |
| F3tv_PrecR | 0.06389 | 0.026512 | 0.017453 | 0.1101551724137931 |
| T2ml_T1a/HeschlR | 0.063623 | 0.026239 | 0.017348 | 0.1101551724137931 |
| AG_F3td | 0.063505 | 0.026448 | 0.017406 | 0.1101551724137931 |
| T2p_T1R | 0.068555 | 0.025693 | 0.017387 | 0.11752285714285715 |
| PT_T2pR | 0.070362 | 0.025275 | 0.017332 | 0.11993522727272726 |
| F3opd_F3tv | 0.072431 | 0.025427 | 0.017446 | 0.12276440677966101 |
| F2p_T1R | 0.074494 | 0.02451 | 0.017234 | 0.12555168539325842 |
| AG_RolS | 0.076881 | 0.024695 | 0.017383 | 0.12885083798882682 |
| STSp_T3p | 0.079086 | 0.024867 | 0.017534 | 0.13108176795580112 |
| T2ml_T3p | 0.079059 | 0.024715 | 0.017479 | 0.13108176795580112 |
| F2p_F3opdR | 0.079731 | 0.024479 | 0.017426 | 0.13142472527472526 |
| T1_STSp | 0.08256 | 0.024289 | 0.017475 | 0.13534426229508195 |
| T3p_T2pR | 0.084009 | 0.023893 | 0.017395 | 0.13697119565217392 |
| Fusa_PrF3op | 0.087616 | 0.023487 | 0.017379 | 0.14208 |
| STSp_T2p | 0.088833 | 0.023142 | 0.017311 | 0.14322032085561498 |
| T1_Fusa | 0.089274 | 0.023686 | 0.017521 | 0.14322032085561498 |
| T3p_RolS | 0.092383 | 0.02283 | 0.017327 | 0.14551884816753927 |
| AG_T2pR | 0.092647 | 0.022977 | 0.017401 | 0.14551884816753927 |
| Prec_PrF3op | 0.091683 | 0.022546 | 0.017192 | 0.14551884816753927 |
| Fusa_T1a/HeschlR | 0.091747 | 0.023454 | 0.017538 | 0.14551884816753927 |
| T3p_F3opdR | 0.093837 | 0.022158 | 0.017159 | 0.14662031250000002 |
| Prec_STSp | 0.096469 | 0.022214 | 0.017256 | 0.14995181347150258 |
| T2ml_RolS | 0.098784 | 0.022648 | 0.017501 | 0.15275876288659793 |
| T1a_F2antR | 0.10306 | 0.021924 | 0.017386 | 0.15855384615384616 |
| SMG_PrecR | 0.10434 | 0.021373 | 0.017227 | 0.15970408163265307 |
| Prec_T1a | 0.10545 | 0.02149 | 0.017311 | 0.15977272727272726 |

| | | | | |
|---|---|---|---|---|
| PT_STSp | 0.10512 | 0.022064 | 0.017525 | 0.15977272727272726 |
| PT_T2p | 0.10636 | 0.021461 | 0.01733 | 0.159915 |
| T3p_T1R | 0.10661 | 0.021009 | 0.017171 | 0.159915 |
| T1_F3opdR | 0.11208 | 0.020925 | 0.017331 | 0.16728358208955224 |
| F3td_T2pR | 0.1139 | 0.020993 | 0.017411 | 0.16742926829268293 |
| PT_T1a | 0.11293 | 0.021292 | 0.017501 | 0.16742926829268293 |
| F3opdR_PrecR | 0.11428 | 0.020773 | 0.017345 | 0.16742926829268293 |
| T1a_F3opdR | 0.11441 | 0.020961 | 0.017415 | 0.16742926829268293 |
| Fusa_F3tv | 0.1192 | 0.020265 | 0.017302 | 0.1711004784688995 |
| F3orb_T1R | 0.11885 | 0.020809 | 0.017515 | 0.1711004784688995 |
| PT_F3opd | 0.11843 | 0.020148 | 0.017232 | 0.1711004784688995 |
| PrF3op_F3orb | 0.11795 | 0.0207 | 0.017438 | 0.1711004784688995 |
| F3orb_RolS | 0.12439 | 0.020042 | 0.017392 | 0.17766824644549764 |
| Fusa_F2antR | 0.12496 | 0.019988 | 0.017388 | 0.17766824644549764 |
| F3opd_F3opdR | 0.12676 | 0.020131 | 0.017513 | 0.17853521126760563 |
| F2p_T3p | 0.12666 | 0.019672 | 0.017313 | 0.17853521126760563 |
| Fusa_T1R | 0.12763 | 0.019715 | 0.017365 | 0.17892056074766352 |
| F3td_F3orb | 0.12887 | 0.019991 | 0.017526 | 0.1798186046511628 |
| RolS_T1R | 0.13014 | 0.019467 | 0.017347 | 0.18075 |
| F3td_F3opdR | 0.13104 | 0.019771 | 0.017521 | 0.18116129032258063 |
| F2p_F3orb | 0.13208 | 0.019339 | 0.017359 | 0.18176146788990827 |
| Fusa_RolS | 0.13324 | 0.019122 | 0.017315 | 0.18252054794520547 |
| F3orb_T1a/HeschlR | 0.13475 | 0.01877 | 0.017207 | 0.18375 |
| T1a_T2p | 0.13873 | 0.018545 | 0.017234 | 0.1883212669683258 |
| T1R_PrecR | 0.14237 | 0.0183 | 0.01725 | 0.19239189189189188 |
| Prec_T1a/HeschlR | 0.14977 | 0.018223 | 0.017472 | 0.20148430493273542 |
| Pole_SMG | 0.15318 | 0.017792 | 0.017395 | 0.20333628318584074 |
| SMG_T1a/HeschlR | 0.15274 | 0.017593 | 0.017295 | 0.20333628318584074 |
| Prec_T2ml | 0.152 | 0.017788 | 0.017344 | 0.20333628318584074 |
| Pole_T1a/HeschlR | 0.15539 | 0.017813 | 0.017475 | 0.2053612334801762 |
| RolS_T1a/HeschlR | 0.15729 | 0.01766 | 0.017464 | 0.2069605263157895 |
| T2p_F3orb | 0.16806 | 0.016826 | 0.017432 | 0.2201659388646288 |
| T2p_T2pR | 0.17132 | 0.016129 | 0.017191 | 0.22236206896551725 |
| T3p_F2antR | 0.17166 | 0.016568 | 0.01742 | 0.22236206896551725 |
| F3opd_T2p | 0.17196 | 0.016526 | 0.017408 | 0.22236206896551725 |
| Pole_RolS | 0.17709 | 0.015945 | 0.017294 | 0.2280128755364807 |
| F2antR_PrecR | 0.19887 | 0.014783 | 0.017414 | 0.25496153846153846 |
| STSp_T1a/HeschlR | 0.2346 | 0.012696 | 0.017425 | 0.2994893617021277 |
| STSp_RolS | 0.2373 | 0.012498 | 0.017396 | 0.3015189873417722 |
| Prec_PrecR | 0.2382 | 0.012511 | 0.017438 | 0.3015189873417722 |
| Fusa_F3orb | 0.2413 | 0.012364 | 0.017457 | 0.3041596638655462 |
| Pole_T2pR | 0.25232 | 0.011454 | 0.017252 | 0.3167196652719665 |
| T3p_PrecR | 0.27593 | 0.010279 | 0.017287 | 0.3449125 |
| Fusa_F3td | 0.28163 | 0.010029 | 0.017322 | 0.35057676348547717 |
| T2pR_T1a/HeschlR | 0.2861 | 0.009608 | 0.017158 | 0.35466942148760333 |
| T1a_T3p | 0.28927 | 0.009551 | 0.017243 | 0.355672131147541 |
| Pole_PrecR | 0.28928 | 0.009771 | 0.017436 | 0.355672131147541 |
| Pole_F3td | 0.30359 | 0.00873 | 0.017136 | 0.3717428571428572 |
| T2p_F3td | 0.30575 | 0.008997 | 0.017488 | 0.3728658536585366 |
| Fusa_F3opdR | 0.30754 | 0.008559 | 0.017157 | 0.37353036437246956 |

| | | | | |
|---|---|---|---|---|
| Pole_Fusa | 0.3287 | 0.007569 | 0.017168 | 0.3976209677419355 |
| T1_T3p | 0.33369 | 0.007289 | 0.01711 | 0.40203614457831327 |
| Fusa_PrecR | 0.34545 | 0.006847 | 0.017242 | 0.41453999999999996 |
| Pole_F2antR | 0.36017 | 0.006288 | 0.0174 | 0.4304820717131474 |
| Prec_Pole | 0.36367 | 0.005992 | 0.017216 | 0.4312292490118577 |
| PrF3op_T1a | 0.36224 | 0.005962 | 0.017083 | 0.4312292490118577 |
| PT_SMG | 0.36966 | 0.005787 | 0.017315 | 0.43660629921259836 |
| T2p_T1a/HeschlR | 0.39113 | 0.004832 | 0.017353 | 0.4601529411764706 |
| PT_F3tv | 0.42601 | 0.003137 | 0.017011 | 0.49923046875 |
| Pole_F3opdR | 0.5 | 1,00E-06 | 0.017279 | 0.5 |
| STSp_PrecR | 0.5 | 1,00E-06 | 0.017026 | 0.5 |
| T1a_PrecR | 0.5 | 1,00E-06 | 0.017425 | 0.5 |
| F2p_T1a/HeschlR | 0.5 | 1,00E-06 | 0.017331 | 0.5 |
| Fusa_T2pR | 0.44619 | 0.002307 | 0.017122 | 0.5 |
| F3td_PrecR | 0.5 | 1,00E-06 | 0.017424 | 0.5 |
| T2ml_Fusa | 0.5 | 1,00E-06 | 0.017395 | 0.5 |
| PT_F2p | 0.5 | 1,00E-06 | 0.017399 | 0.5 |
| PT_F3opdR | 0.5 | 1,00E-06 | 0.017343 | 0.5 |
| PT_F3orb | 0.5 | 1,00E-06 | 0.017166 | 0.5 |
| F3td_RolS | 0.5 | 1,00E-06 | 0.01708 | 0.5 |
| Prec_T2pR | 0.4498 | 0.002227 | 0.017419 | 0.5 |
| PrF3op_RolS | 0.5 | 1,00E-06 | 0.017272 | 0.5 |
| F3orb_PrecR | 0.5 | 1,00E-06 | 0.017381 | 0.5 |
| AG_F3opdR | 0.5 | 1,00E-06 | 0.017246 | 0.5 |
| T1a_T2pR | 0.46503 | 0.001511 | 0.017201 | 0.5 |
| F2p_PrecR | 0.5 | 1,00E-06 | 0.017261 | 0.5 |
| PrF3op_T1a/HeschlR | 0.5 | 1,00E-06 | 0.017359 | 0.5 |
| F2p_RolS | 0.5 | 1,00E-06 | 0.017335 | 0.5 |
| F2p_Fusa | 0.5 | 1,00E-06 | 0.017206 | 0.5 |
| F3opdR_T1a/HeschlR | 0.43467 | 0.002787 | 0.017075 | 0.5 |
| T1a_F3td | 0.5 | 1,00E-06 | 0.01708 | 0.5 |
| Prec_T1R | 0.49947 | 2.2e-05 | 0.016903 | 0.5 |
| T2p_PrecR | 0.5 | 1,00E-06 | 0.017217 | 0.5 |
| T1a/HeschlR_PrecR | 0.5 | 1,00E-06 | 0.016987 | 0.5 |
| T2ml_PrecR | 0.5 | 1,00E-06 | 0.017292 | 0.5 |
| F3opd_T1a/HeschlR | 0.5 | 1,00E-06 | 0.017285 | 0.5 |
| PT_Fusa | 0.5 | 1,00E-06 | 0.01736 | 0.5 |
| SMG_RolS | 0.5 | 1,00E-06 | 0.0173 | 0.5 |
| Pole_T3p | 0.5 | 1,00E-06 | 0.017169 | 0.5 |
| AG_T1a/HeschlR | 0.49923 | 3.3e-05 | 0.017235 | 0.5 |
| F3opd_PrecR | 0.5 | 1,00E-06 | 0.017067 | 0.5 |
| PT_T3p | 0.5 | 1,00E-06 | 0.017343 | 0.5 |
| PT_F3td | 0.5 | 1,00E-06 | 0.017642 | 0.5 |
| RolS_T2pR | 0.5 | 1,00E-06 | 0.017092 | 0.5 |
| PT_PrecR | 0.5 | 1,00E-06 | 0.017182 | 0.5 |
| T1_F3td | 0.5 | 1,00E-06 | 0.017244 | 0.5 |
| F2p_F3td | 0.5 | 1,00E-06 | 0.017338 | 0.5 |
| Prec_F2p | 0.5 | 1,00E-06 | 0.017259 | 0.5 |
| T3p_T1a/HeschlR | 0.5 | 1,00E-06 | 0.017368 | 0.5 |
| F3opd_Fusa | 0.5 | 1,00E-06 | 0.017196 | 0.5 |

| | | | | |
|---|---|---|---|---|
| PT_PrF3op | 0.5 | 1,00E-06 | 0.017079 | 0.5 |
| Prec_PT | 0.5 | 1,00E-06 | 0.017299 | 0.5 |
| F3td_T1a/HeschlR | 0.5 | 1,00E-06 | 0.017407 | 0.5 |

| CHR | SNP | BP | A1 | A2 | P value | Z-stat | N |
|---|---|---|---|---|---|---|---|
| 2 | rs2717046 | 58041936 | C | T | 7.509071256533833e-14 | 7.478648684332633 | 31894.0 |
| 2 | rs2717066 | 58029645 | C | A | 9.79095283479027e-14 | 7.443692134154676 | 32006.0 |
| 2 | rs11695135 | 57998040 | A | C | 1.402962952252713e-13 | 7.396048960718104 | 31964.0 |
| 2 | 2:58029698_GA_G | 58029698 | GA | G | 2.2843870492019533e-13 | 7.331001736579746 | 31575.0 |
| 2 | rs762523673 | 57944587 | TA | T | 2.300842238678481e-13 | 7.330039897642196 | 30496.0 |
| 2 | rs2717079 | 58065936 | T | A | 3.5951928343580125e-13 | 7.269988057019218 | 31914.0 |
| 2 | rs10204616 | 57980955 | A | T | 3.762865538931088e-13 | 7.263827956399022 | 32036.0 |
| 2 | rs13025126 | 57956088 | G | A | 4.756659718561889e-13 | 7.23207596792804 | 31768.0 |
| 2 | rs2125925 | 57971343 | A | G | 4.972377109334335e-13 | 7.226051919294318 | 32030.0 |
| 2 | rs11679364 | 57975524 | T | G | 5.6011306506235 36e-13 | 7.209855442848077 | 32030.0 |
| 2 | rs2947349 | 58059803 | C | A | 8.159166509241092e-13 | 7.158454490341638 | 31899.0 |
| 2 | rs11125729 | 57975170 | T | C | 8.252758119625963e-13 | 7.156890471883142 | 32030.0 |
| 2 | rs3886275 | 57973344 | A | C | 8.316935801960934e-13 | 7.155828022329335 | 32028.0 |
| 2 | rs13002041 | 57972834 | C | A | 8.382083902838332e-13 | 7.1547577063762775 | 32024.0 |
| 2 | rs7567497 | 57974429 | A | T | 8.724965022511116e-13 | 7.149255738822888 | 32032.0 |
| 2 | rs2290867 | 57952870 | A | C | 9.22657307832851e-13 | 7.141577646704316 | 31821.0 |
| 2 | rs12990792 | 57993568 | G | A | 9.901417374212297e-13 | 7.1318702752907575 | 32058.0 |
| 2 | rs717817 | 57973452 | C | T | 9.946420124661571e-13 | 7.131246228859893 | 32027.0 |
| 2 | rs70954867 | 58004577 | A | AT | 1.0071693047278785e-12 | 7.129523571404032 | 31910.0 |
| 2 | rs7588492 | 57995852 | A | G | 1.050504443479496e-12 | 7.123722502441468 | 32040.0 |
| 2 | rs13402331 | 57947440 | T | C | 1.1182120128750344e-12 | 7.1151129766265475 | 31800.0 |
| 2 | rs10186791 | 57957572 | G | A | 1.2362338995419529e-12 | 7.1012612955872045 | 31848.0 |
| 2 | rs10496078 | 57993643 | G | A | 1.3432180603233644e-12 | 7.089783666725836 | 31970.0 |
| 2 | 2:58008036_CT_C | 58008036 | CT | C | 1.344804532198401e-12 | 7.089620303649722 | 30524.0 |
| 2 | rs6709720 | 57967563 | A | G | 1.3571681570540057e-12 | 7.088353630702332 | 31972.0 |
| 2 | rs13418015 | 57948635 | A | G | 1.44687051685477e-12 | 7.079488934648515 | 31798.0 |
| 2 | rs1568452 | 58012833 | C | T | 1.4806833531893423e-12 | 7.076286735313048 | 32002.0 |
| 2 | rs747713243 | 57944289 | GTTGA | G | 1.5320066842750308e-12 | 7.0715607847089315 | 30732.0 |
| 2 | rs1402399 | 58045395 | T | C | 1.7031534764195436e-12 | 7.056853227586067 | 32019.0 |
| 2 | rs10496079 | 57997136 | G | C | 1.7578655315805634e-12 | 7.0524563609521564 | 32094.0 |
| 2 | rs754784696 | 57995420 | CACTGCAAC CTCTGCCT | C | 1.8011464853968762e-12 | 7.04907222631785 | 31558.0 |
| 2 | rs6756221 | 58008155 | T | C | 1.8756343501621482e-12 | 7.043430605233335 | 32065.0 |
| 2 | 2:58025739_TA_T | 58025739 | TA | T | 1.98199304258768e-12 | 7.035744872113369 | 32067.0 |
| 2 | rs2717067 | 58095252 | G | C | 2.1869505778929644e-12 | 7.022012440295446 | 31937.0 |
| 2 | rs2953441 | 58041482 | C | T | 2.2064663280911888e-12 | 7.020771386097072 | 32012.0 |
| 2 | rs1474215 | 58032029 | C | G | 2.263595032794922e-12 | 7.017199373613589 | 32115.0 |
| 2 | rs2717018 | 58035491 | C | T | 2.3418209565171767e-12 | 7.012449084455942 | 32030.0 |
| 2 | rs2683629 | 58037369 | A | G | 2.4374518328118458e-12 | 7.0068489382227 | 32020.0 |
| 2 | rs2953439 | 58039357 | T | C | 2.441604914322157e-12 | 7.006610643753325 | 32018.0 |
| 2 | rs4671319 | 57950346 | G | A | 2.450050110564384e-12 | 7.006127301054232 | 31806.0 |
| 2 | rs12997222 | 57987201 | C | G | 2.550559138505833e-12 | 7.000497099624357 | 32032.0 |
| 2 | rs2717052 | 58047263 | C | A | 2.6097337244071257e-12 | 6.997283236956407 | 32019.0 |
| 2 | rs2717071 | 58030962 | G | A | 2.632890120618841e-12 | 6.996044995013306 | 32118.0 |
| 2 | rs7598169 | 58040729 | C | T | 2.6615181469845307e-12 | 6.994528849317216 | 32007.0 |
| 2 | rs2717075 | 58032530 | G | C | 2.8076048141908846e-12 | 6.987032870388254 | 32107.0 |
| 2 | rs2717048 | 58025937 | C | T | 2.8488681332895767e-12 | 6.98498480513149 | 32133.0 |
| 2 | rs2683630 | 58058069 | C | G | 2.8702760734587363e-12 | 6.983933675562249 | 32001.0 |
| 2 | rs2125924 | 57943567 | T | C | 2.8976660739236828e-12 | 6.982599981614809 | 31789.0 |
| 2 | rs2683628 | 58036044 | A | G | 2.901616570869728e-12 | 6.982408641977428 | 32034.0 |
| 2 | rs2683619 | 58032759 | C | T | 3.08752837905613e-12 | 6.973681416423547 | 32077.0 |
| 2 | rs2717055 | 58044220 | G | A | 3.1786612528173614e-12 | 6.969589932933111 | 32013.0 |
| 2 | rs2683621 | 58046995 | A | G | 3.2977151430537547e-12 | 6.964414853888366 | 32009.0 |

| | | | | | | | |
|---|---|---|---|---|---|---|---|
| 2 | rs2717054 | 58046683 | G | A | 3.3717263583317575e-12 | 6.961289267349797 | 32015.0 |
| 2 | rs1402398 | 58042241 | G | A | 3.4665385475904886e-12 | 6.957382092382258 | 32015.0 |
| 2 | rs2683614 | 58026312 | G | A | 3.4668157932338197e-12 | 6.9573708215549805 | 32136.0 |
| 2 | rs2717053 | 58047101 | G | C | 3.474504213900687e-12 | 6.957058616863455 | 32011.0 |
| 2 | rs7561842 | 57995586 | C | T | 3.4937292174916584e-12 | 6.956280899567088 | 32052.0 |
| 2 | rs6545665 | 58023218 | G | A | 3.803358069783314e-12 | 6.9443021077191185 | 32137.0 |
| 2 | rs10865304 | 58002818 | G | T | 3.9390187567150486e-12 | 6.939352238613156 | 32129.0 |
| 2 | rs11676055 | 58017224 | C | T | 4.018724989299623e-12 | 6.936521363730384 | 32142.0 |
| 2 | rs2683616 | 58035555 | G | A | 4.173080467304795e-12 | 6.931192391601898 | 32033.0 |
| 2 | rs13026612 | 58014723 | C | T | 4.2019843160646536e-12 | 6.930216034061496 | 32153.0 |
| 2 | rs13026830 | 58014831 | C | T | 4.399808385818316e-12 | 6.923705266978315 | 32186.0 |
| 2 | rs954593 | 58027337 | C | T | 4.570473597500106e-12 | 6.918314931281322 | 32123.0 |
| 2 | 2:57944589_TA_T | 57944589 | TA | T | 4.61693865900516e-12 | 6.916881530867037 | 31063.0 |
| 2 | rs4296411 | 58008872 | A | C | 4.6872699531209335e-12 | 6.914738561494605 | 32129.0 |
| 2 | rs2683620 | 58032553 | A | T | 4.75293794694769e-12 | 6.912765947786868 | 32094.0 |
| 2 | rs10166481 | 58016412 | G | A | 4.799590883779733e-12 | 6.911380697018104 | 32145.0 |
| 2 | rs6545664 | 58023134 | C | T | 4.8723988156808785e-12 | 6.9092449893533825 | 32126.0 |
| 2 | rs12713358 | 58007475 | A | T | 4.947716119983998e-12 | 6.90706834328845 | 32123.0 |
| 2 | rs6737913 | 58007759 | G | A | 5.122247456164546e-12 | 6.902146760628173 | 32129.0 |
| 2 | rs12991325 | 58016879 | C | T | 5.2058935066123285e-12 | 6.899846023034012 | 32129.0 |
| 2 | rs11898858 | 58006624 | G | A | 5.2430655488259674e-12 | 6.898835189190768 | 32129.0 |
| 2 | rs6545663 | 58023009 | A | T | 5.4461797015394634e-12 | 6.893432945788772 | 32134.0 |
| 2 | rs1568450 | 58011697 | A | G | 5.4533759225226845e-12 | 6.893245189793202 | 32140.0 |
| 2 | rs6761347 | 58010040 | T | A | 5.513784004873107e-12 | 6.8916785957517455 | 32136.0 |
| 2 | rs2683615 | 58027493 | C | A | 5.77640683131602e-12 | 6.885057819863727 | 32134.0 |
| 2 | rs1589407 | 58019903 | T | C | 6.114830298944072e-12 | 6.876948267629175 | 32133.0 |
| 2 | rs6730037 | 58005792 | G | A | 6.382510026711893e-12 | 6.870839661209673 | 32127.0 |
| 2 | rs10195687 | 57956533 | T | G | 6.637012260509859e-12 | 6.865260294841688 | 30956.0 |
| 2 | rs2090793 | 58021528 | G | A | 6.831652151435925e-12 | 6.861132997671249 | 32131.0 |
| 2 | 2:57958740_AT_A | 57958740 | AT | A | 7.2726263925100866e-12 | 6.852193157094956 | 30959.0 |
| 2 | rs4672223 | 58021882 | C | A | 7.382042436312638e-12 | 6.850057279921237 | 32135.0 |
| 2 | rs13016665 | 57995348 | C | A | 7.585676138068826e-12 | 6.846163532231407 | 31377.0 |
| 2 | rs13032423 | 57992112 | G | A | 7.590666038505336e-12 | 6.8460694097598696 | 32026.0 |
| 2 | rs2717076 | 58061127 | C | T | 8.978237879762524e-12 | 6.8219988607527675 | 31983.0 |
| 2 | rs34035400 | 57979248 | T | TC | 1.0663460561044457e-11 | 6.797250487503264 | 31344.0 |
| 2 | rs4671320 | 57955644 | C | G | 1.0827401565139252e-11 | 6.795051271028208 | 30905.0 |
| 2 | rs6720888 | 57945522 | A | G | 1.1898896428705287e-11 | 6.781433842511929 | 31719.0 |
| 2 | rs10496076 | 57942987 | T | C | 1.2500983103621435e-11 | 6.774300234350301 | 31700.0 |
| 2 | 2:58006919_GA_G | 58006919 | GA | G | 1.6801244888995532e-11 | 6.731422385463492 | 31353.0 |
| 2 | rs10196378 | 57951208 | T | G | 1.7949525851836266e-11 | 6.7217983441656735 | 31734.0 |
| 2 | rs2678870 | 58153375 | T | A | 2.205387733850836e-11 | 6.691735800270817 | 31457.0 |
| 2 | rs1460255 | 57978858 | T | C | 2.443785238474017e-11 | 6.676702377103402 | 32004.0 |
| 2 | rs2678869 | 58153374 | A | C | 2.536859778891077e-11 | 6.67121977566787 | 31449.0 |
| 2 | rs2678900 | 58177683 | T | G | 3.3523663684299326e-11 | 6.6301981081747074 | 31911.0 |
| 2 | rs200493119 | 58154132 | C | CG | 3.550819683793183e-11 | 6.621703825229244 | 31990.0 |
| 2 | rs540927767 | 58154133 | T | TG | 3.550819683793183e-11 | 6.621703825229244 | 31990.0 |
| 2 | rs2139054 | 58156583 | A | C | 3.7083772274730795e-11 | 6.615284549858058 | 32146.0 |
| 2 | rs2678871 | 58153602 | G | A | 3.722021311095729e-11 | 6.614741275750567 | 31315.0 |
| 2 | rs2717002 | 58142647 | T | G | 3.8680837122187476e-11 | 6.609044528366264 | 32151.0 |
| 2 | rs2683618 | 58033361 | A | G | 4.195036041680869e-11 | 6.5970202077931965 | 32001.0 |
| 2 | rs2717068 | 58094873 | A | C | 4.204053324410885e-11 | 6.596701733804897 | 32186.0 |
| 2 | rs908731 | 58135010 | C | G | 4.568074311588503e-11 | 6.58437351047073 | 32134.0 |
| 2 | rs2312141 | 58155358 | A | T | 4.671864965451051e-11 | 6.58103439066585 | 32146.0 |
| 2 | rs908730 | 58135011 | G | A | 4.771220096525593e-11 | 6.577905275304311 | 31892.0 |

| | | | | | | | |
|---|---|---|---|---|---|---|---|
| 2 | rs2717004 | 58143854 | C | A | 4.954565724890802e-11 | 6.572294757013515 | 32130.0 |
| 2 | rs2717015 | 58155253 | T | C | 5.062431743716292e-11 | 6.569088128209432 | 32147.0 |
| 2 | rs2717003 | 58143438 | A | G | 5.202366244763265e-11 | 6.565026348535002 | 32122.0 |
| 2 | rs7557715 | 57973818 | C | T | 5.4037119689802747e-11 | 6.559365741753241 | 32002.0 |
| 2 | rs2678873 | 58154601 | G | A | 5.4757340303383944e-11 | 6.5573909082168536 | 32144.0 |
| 2 | rs2465804 | 58148180 | G | A | 5.5002343775255674e-11 | 6.556724900470686 | 32151.0 |
| 2 | rs2678880 | 58158283 | C | T | 5.6194189905930995e-11 | 6.553525877227467 | 32155.0 |
| 2 | rs2678881 | 58140977 | G | A | 5.983692333840129e-11 | 6.544144492310271 | 32162.0 |
| 2 | rs2717007 | 58149116 | G | A | 6.225621735678367e-11 | 6.538217804341795 | 32152.0 |
| 2 | rs2678893 | 58167143 | A | G | 6.498793297339463e-11 | 6.531790701488059 | 31518.0 |
| 2 | rs2678915 | 58119574 | C | T | 6.677197742802412e-11 | 6.527734368726225 | 32084.0 |
| 2 | rs2678905 | 58135872 | A | G | 6.730540209240214e-11 | 6.52654210664636 | 32127.0 |
| 2 | rs1040224 | 58139865 | G | C | 7.00071125732075e-11 | 6.520641962172031 | 32148.0 |
| 2 | rs2678901 | 58138651 | A | G | 7.006641984620112e-11 | 6.520514956828962 | 32144.0 |
| 2 | rs2678886 | 58161221 | A | G | 7.212038918647845e-11 | 6.516180074849907 | 32157.0 |
| 2 | rs1040225 | 58139593 | A | G | 7.248312883468831e-11 | 6.515427080276327 | 32144.0 |
| 2 | rs11410659 | 58145483 | A | AT | 7.255176124644291e-11 | 6.515285024095081 | 32127.0 |
| 2 | rs2717010 | 58151817 | A | G | 7.264672457106099e-11 | 6.515088684442151 | 32145.0 |
| 2 | rs2678904 | 58137755 | G | T | 7.495260467269011e-11 | 6.510396686020832 | 32186.0 |
| 2 | rs2678903 | 58137930 | A | G | 7.62883407826805e-11 | 6.507742891087599 | 32156.0 |
| 2 | rs2717024 | 58161397 | C | G | 7.858665339407565e-11 | 6.503281278830475 | 32159.0 |
| 2 | rs2678890 | 58164435 | C | T | 7.904095685540438e-11 | 6.50241448175816 | 32186.0 |
| 2 | 2:57971694_GAAAA_G | 57971694 | GAAAA | G | 7.938673861258053e-11 | 6.5017579997209465 | 27885.0 |
| 2 | rs2717011 | 58152075 | T | C | 8.177266487209052e-11 | 6.4973029991758775 | 32118.0 |
| 2 | rs2459982 | 58150941 | A | C | 8.241143924385449e-11 | 6.49613184725004 | 32151.0 |
| 2 | rs2717014 | 58154249 | T | C | 8.288372022848787e-11 | 6.495271643788983 | 32157.0 |
| 2 | rs2717023 | 58161355 | C | T | 8.433023290534931e-11 | 6.492666503191044 | 32160.0 |
| 2 | rs2717001 | 58137618 | C | T | 8.982133723183451e-11 | 6.483159693547808 | 32142.0 |
| 2 | rs2678892 | 58167140 | G | A | 9.015062741680626e-11 | 6.4826077999802765 | 31772.0 |
| 2 | rs2717022 | 58160639 | G | C | 9.23177405796327e-11 | 6.479024130374065 | 32155.0 |
| 2 | rs10694743 | 58149555 | G | GCTAA | 9.313760211575412e-11 | 6.477689763362997 | 31917.0 |
| 2 | rs2678887 | 58161752 | C | A | 9.377967288290532e-11 | 6.476652754281639 | 32164.0 |
| 2 | rs2678888 | 58162191 | A | G | 9.540620104491484e-11 | 6.474056497863742 | 32161.0 |
| 2 | rs2678879 | 58158152 | A | G | 9.623318398756093e-11 | 6.472753019968161 | 32148.0 |
| 2 | rs2678883 | 58159670 | A | G | 9.727648543454776e-11 | 6.471124118938718 | 32165.0 |
| 2 | rs2717019 | 58159473 | T | G | 9.989130451977444e-11 | 6.467115492663632 | 32164.0 |
| 2 | rs2717033 | 58166769 | T | C | 1.003161196388558e-10 | 6.466473937085437 | 32092.0 |
| 2 | 2:57971686_AAAAG_A | 57971686 | AAAAG | A | 1.0730299105885727e-10 | 6.456287051652602 | 28278.0 |
| 2 | rs10192098 | 57971698 | A | G | 1.137357566425249e-10 | 6.447465891876822 | 28628.0 |
| 2 | 2:57944288_AGTTG_A | 57944288 | AGTTG | A | 1.1380448180828348e-10 | 6.447374308120757 | 28211.0 |
| 2 | rs1568253 | 58170004 | T | C | 1.2171348690034406e-10 | 6.437180098060972 | 32094.0 |
| 2 | rs112345168 | 58007905 | T | A | 1.2360602349898663e-10 | 6.434836852733957 | 32035.0 |
| 2 | rs59098844 | 58044533 | G | C | 1.2549593764805905e-10 | 6.432531590796591 | 31989.0 |
| 2 | rs1568254 | 58170161 | C | A | 1.308890180370703e-10 | 6.426135177831644 | 32060.0 |
| 2 | rs2678906 | 58134048 | T | A | 1.321051669433591e-10 | 6.424728405042805 | 32117.0 |
| 2 | rs2678889 | 58164223 | A | G | 1.3370833475592232e-10 | 6.422893167492071 | 32172.0 |
| 2 | rs2176546 | 58156772 | G | C | 1.4422191637712941e-10 | 6.411365551206032 | 31391.0 |
| 2 | rs2683634 | 58113995 | T | C | 1.518528093372511e-10 | 6.403502056206875 | 32092.0 |
| 2 | rs2678891 | 58166643 | T | G | 1.5421120136638274e-10 | 6.40114977658657 | 32170.0 |
| 2 | rs2678882 | 58159362 | T | A | 1.571753160248631e-10 | 6.398242706218817 | 32165.0 |
| 2 | rs970941 | 58166181 | G | A | 1.591276109496912e-10 | 6.396357102210738 | 32172.0 |
| 2 | rs2717036 | 58168098 | T | C | 1.7041806053127746e-10 | 6.385876893364757 | 32101.0 |
| 2 | rs11433420 | 58037203 | T | TA | 1.7240782506601732e-10 | 6.384100514460116 | 31750.0 |
| 2 | 2:58165967_TTC_T | 58165967 | TTC | T | 1.754988159125896e-10 | 6.381380349092268 | 32153.0 |

| | | | | | | | |
|---|---|---|---|---|---|---|---|
| 2 | rs2717031 | 58166468 | T | C | 1.8627206460450842e-10 | 6.372252343143579 | 32165.0 |
| 2 | rs1518394 | 58171287 | A | G | 1.8703166620754984e-10 | 6.371628355310698 | 32082.0 |
| 2 | rs2678908 | 58130327 | C | T | 1.9489515852226858e-10 | 6.365310424609946 | 32115.0 |
| 2 | rs2717034 | 58167258 | T | G | 2.1658607047346155e-10 | 6.34909442448742 | 31924.0 |
| 2 | rs2717032 | 58166760 | T | C | 2.1893019018983542e-10 | 6.347437980949673 | 32103.0 |
| 2 | rs66490669 | 58040174 | T | C | 2.3238840117095002e-10 | 6.338250752756887 | 31988.0 |
| 2 | rs2717056 | 58128804 | C | T | 2.4546089963083733e-10 | 6.3298114878031955 | 32093.0 |
| 2 | rs2678894 | 58167260 | G | C | 3.1863273414313694e-10 | 6.289430492216134 | 31892.0 |
| 2 | rs1356544 | 58123724 | T | C | 3.436154865625484e-10 | 6.2777011868663175 | 32043.0 |
| 2 | rs2717020 | 58159852 | T | C | 3.6114780196782717e-10 | 6.2699570098353075 | 32039.0 |
| 2 | rs1356545 | 58123798 | T | C | 3.6815460684928515e-10 | 6.266964260737129 | 32042.0 |
| 2 | rs34738965 | 58160798 | G | GT | 3.6938255039266993e-10 | 6.266445513735445 | 31829.0 |
| 2 | rs2717029 | 58165595 | G | T | 3.920585364773792e-10 | 6.257157057943985 | 32045.0 |
| 2 | rs1518393 | 58171220 | A | C | 4.293770582666652e-10 | 6.24295624044365 | 30817.0 |
| 2 | rs2717040 | 58169494 | T | C | 4.612037532345532e-10 | 6.231766777994716 | 32095.0 |
| 2 | rs2717030 | 58165694 | T | C | 4.780809461575725e-10 | 6.226135397721006 | 32048.0 |
| 2 | rs2678896 | 58169388 | A | G | 4.841172465276129e-10 | 6.224168281848755 | 32080.0 |
| 2 | rs2717039 | 58169166 | G | T | 5.083219249677849e-10 | 6.21651372950103 | 31989.0 |
| 2 | rs2678885 | 58160138 | T | G | 5.304242646916319e-10 | 6.20982859118433 | 32035.0 |
| 2 | rs2717021 | 58160605 | G | A | 5.838425690807059e-10 | 6.1947320767967975 | 32039.0 |
| 2 | rs2717041 | 58172107 | C | T | 5.862092206935854e-10 | 6.194094843803301 | 32045.0 |
| 2 | rs2717035 | 58167440 | C | T | 6.0612276042444e-10 | 6.188830340474175 | 32120.0 |
| 2 | rs2717025 | 58163146 | G | T | 6.232256912549632e-10 | 6.184441879343302 | 32029.0 |
| 2 | rs2953440 | 58039936 | C | T | 6.278383726728433e-10 | 6.183278422119954 | 31140.0 |
| 2 | rs2678874 | 58154915 | T | A | 6.280396846155336e-10 | 6.183227835313652 | 32037.0 |
| 2 | rs1401100 | 58168856 | G | T | 6.347140145895304e-10 | 6.181559567319177 | 32093.0 |
| 2 | rs68119977 | 58078939 | T | G | 6.428125546194884e-10 | 6.1795581471700025 | 31947.0 |
| 2 | rs2717026 | 58163178 | A | G | 6.534427397782493e-10 | 6.176968089051997 | 32031.0 |
| 2 | rs2312143 | 58155829 | G | A | 7.412868960084963e-10 | 6.157015300853003 | 32040.0 |
| 2 | rs13011472 | 57961602 | C | G | 7.509231289987038e-10 | 6.154968724815363 | 31828.0 |
| 2 | rs2717005 | 58145060 | A | G | 7.802993256855536e-10 | 6.1488841739851186 | 32050.0 |
| 2 | rs62158166 | 114077218 | G | C | 8.696490048426408e-10 | 6.13166349248832 | 31803.0 |
| 2 | rs2678895 | 58139906 | A | T | 8.841255681399849e-10 | 6.12903701618565 | 32087.0 |
| 2 | rs2717038 | 58168831 | A | G | 9.725575113458217e-10 | 6.11384996027404 | 31996.0 |
| 2 | rs2717000 | 58136220 | C | G | 9.726655354912617e-10 | 6.113832245361495 | 32110.0 |
| 2 | rs867740 | 58135023 | T | C | 9.987139503655902e-10 | 6.1096155988314065 | 32116.0 |
| 2 | rs12616641 | 114079248 | C | A | 1.0352198148425894e-09 | 6.103883150596727 | 31792.0 |
| 2 | rs2717012 | 58152280 | C | A | 1.0523498503794608e-09 | 6.101260878601733 | 32028.0 |
| 2 | rs2683632 | 58116640 | T | C | 1.0546480668889708e-09 | 6.100912237651203 | 32186.0 |
| 2 | rs765106764 | 58130295 | TA | T | 1.0712386840286953e-09 | 6.0984171889841035 | 32057.0 |
| 2 | rs951965 | 58144542 | C | A | 1.086093372448454e-09 | 6.09621497012119 | 32057.0 |
| 2 | rs2678907 | 58133522 | A | G | 1.1326451901864956e-09 | 6.089499075599543 | 32077.0 |
| 2 | rs2717063 | 58110969 | C | A | 1.1353003012507847e-09 | 6.089124188412862 | 31314.0 |
| 2 | rs2678902 | 58138581 | G | A | 1.1584979761741794e-09 | 6.08588469498637 | 32096.0 |
| 2 | rs6720288 | 58111064 | A | G | 1.170811156021617e-09 | 6.084190794909186 | 32107.0 |
| 2 | rs62158168 | 114078381 | C | G | 1.2639686162508764e-09 | 6.071911153655794 | 31798.0 |
| 2 | rs11682175 | 57987593 | T | C | 1.2785519848169445e-09 | 6.070069133558503 | 31916.0 |
| 2 | rs34967704 | 58150658 | G | GT | 1.301437898222938e-09 | 6.067219312147462 | 29452.0 |
| 2 | rs2678916 | 58119582 | G | T | 1.3377408163134379e-09 | 6.0627974634729505 | 32152.0 |
| 2 | rs2717013 | 58152537 | G | A | 1.3676377612098019e-09 | 6.059242852320146 | 32033.0 |
| 2 | rs1533725 | 58111922 | T | G | 1.4855149773372213e-09 | 6.045929011950247 | 32110.0 |
| 2 | rs2717059 | 58119657 | C | G | 1.5171172228633376e-09 | 6.042534683447893 | 32162.0 |
| 2 | rs2717060 | 58119122 | A | G | 1.53821636617549e-09 | 6.04030662097888 | 32162.0 |
| 2 | rs2717057 | 58127907 | C | T | 1.5520242839244666e-09 | 6.0388645767326485 | 32132.0 |

| | | | | | | | |
|---|---|---|---|---|---|---|---|
| 2 | rs35185782 | 58169712 | T | TG | 1.5611519081542134e-09 | 6.037918172190772 | 31929.0 |
| 2 | rs2678911 | 58115072 | G | C | 1.5899735751718672e-09 | 6.034964784362972 | 32143.0 |
| 2 | rs2678917 | 58120553 | T | A | 1.6042147323951215e-09 | 6.033524683179313 | 32161.0 |
| 2 | rs62158169 | 114081827 | C | T | 1.7560111217112142e-09 | 6.018904790252428 | 31833.0 |
| 2 | 2:58167698_GA_G | 58167698 | GA | G | 1.8173082916987509e-09 | 6.013347553692409 | 31935.0 |
| 2 | rs2678913 | 58117095 | T | C | 1.9067708757113082e-09 | 6.0055562114454135 | 32173.0 |
| 2 | rs2030237 | 58196110 | A | G | 1.950133348638669e-09 | 6.001907050700856 | 31821.0 |
| 2 | rs11387884 | 58126250 | C | CT | 1.994684086599985e-09 | 5.998239317832078 | 32149.0 |
| 2 | rs2678897 | 58169418 | G | A | 1.9993313462616354e-09 | 5.997861329942781 | 31730.0 |
| 2 | rs2465805 | 58148349 | C | T | 2.0272168303128795e-09 | 5.995611067330472 | 32045.0 |
| 2 | rs2678898 | 58171964 | T | C | 2.044298798455136e-09 | 5.994247458611572 | 31843.0 |
| 2 | rs62158170 | 114082175 | A | G | 2.145329402490224e-09 | 5.986402614078817 | 31848.0 |
| 2 | rs2717008 | 58149158 | T | C | 2.146458700537828e-09 | 5.9863169761032085 | 32045.0 |
| 2 | rs2678918 | 58121391 | C | T | 2.1469921833258327e-09 | 5.986276535805888 | 32142.0 |
| 2 | rs781157235 | 58115023 | GCTAT | G | 2.330216518509272e-09 | 5.972935776432117 | 32014.0 |
| 2 | rs7606466 | 57942325 | C | G | 2.413965327697891e-09 | 5.9671749548492565 | 31695.0 |
| 2 | rs2678914 | 58117926 | C | G | 2.647671675789205e-09 | 5.9520730330130815 | 32160.0 |
| 2 | rs2678909 | 58150768 | T | C | 2.703116108355669e-09 | 5.948681153815977 | 32099.0 |
| 2 | rs11688767 | 57988194 | A | T | 2.82050659497444e-09 | 5.9417178055990245 | 32019.0 |
| 2 | rs2139053 | 58156539 | T | C | 2.838538928832563e-09 | 5.940673248909506 | 32117.0 |
| 2 | rs62141276 | 48214217 | G | A | 3.260361758521317e-09 | 5.9179211596293415 | 31243.0 |
| 2 | 2:58114255_CAT_C | 58114255 | CAT | C | 3.4487512714418724e-09 | 5.908672705049372 | 31713.0 |
| 2 | rs7573056 | 58229428 | A | C | 3.4534302677763477e-09 | 5.908449316872248 | 31775.0 |
| 2 | rs7600005 | 48212055 | G | A | 3.4616021564957447e-09 | 5.908059874141163 | 31250.0 |
| 2 | rs199993536 | 114082628 | T | TA | 3.6096074540096334e-09 | 5.901157210496769 | 31456.0 |
| 2 | rs6717480 | 58210522 | C | A | 3.786755864022748e-09 | 5.8932487985970345 | 31813.0 |
| 2 | rs10172295 | 58201202 | G | A | 3.850876669734966e-09 | 5.890474768318746 | 31758.0 |
| 2 | 2:58225035_GA_G | 58225035 | GA | G | 4.229429511596792e-09 | 5.874960982998737 | 32045.0 |
| 2 | rs2459981 | 58127925 | C | A | 4.2365243948535626e-09 | 5.874683316100464 | 32095.0 |
| 2 | rs4672228 | 58237405 | T | C | 4.644581577154848e-09 | 5.8594301628841015 | 31713.0 |
| 2 | rs2678910 | 58112266 | A | T | 4.767318338502343e-09 | 5.85509694880216 | 32070.0 |
| 2 | 2:58047881_CAA_C | 58047881 | CAA | C | 5.51218864711555e-09 | 5.83092218691396 | 30062.0 |
| 2 | rs4672226 | 58203057 | C | T | 5.6737748794352885e-09 | 5.8260996397533376 | 31725.0 |
| 2 | rs6722461 | 57970943 | A | G | 5.854459831672264e-09 | 5.820862844514839 | 31913.0 |
| 2 | rs2678884 | 58160123 | A | C | 6.359046829817534e-09 | 5.807030731331679 | 32133.0 |
| 2 | rs2717027 | 58164454 | G | A | 6.4087698251746e-09 | 5.805725989050696 | 32141.0 |
| 2 | rs2678922 | 58127647 | A | G | 6.4483235353337515e-09 | 5.804695103377244 | 31764.0 |
| 2 | rs2678923 | 58127648 | A | C | 6.4483235353337515e-09 | 5.804695103377244 | 31764.0 |
| 2 | rs2043890 | 58350571 | G | A | 6.577663541719292e-09 | 5.801366547769947 | 31914.0 |
| 2 | rs62158161 | 114065572 | C | T | 8.105166786222006e-09 | 5.7662565111120125 | 31596.0 |
| 2 | rs6737318 | 114083120 | A | G | 8.266233075762942e-09 | 5.762937667077415 | 31861.0 |
| 2 | rs17396357 | 48252311 | C | T | 8.635586990735586e-09 | 5.7555583821790846 | 31972.0 |
| 2 | rs375442013 | 58364359 | C | CAT | 8.685063602642473e-09 | 5.754593285245908 | 31215.0 |
| 2 | rs2717042 | 58177082 | C | A | 1.0210498276436132e-08 | 5.727194664998169 | 32055.0 |
| 2 | rs7556815 | 114085785 | G | A | 1.2556660014912192e-08 | 5.691990828868012 | 31884.0 |
| 2 | rs12615370 | 114068017 | T | G | 1.3019778500892385e-08 | 5.685805138745737 | 31853.0 |
| 2 | rs62158163 | 114071717 | T | C | 1.4009022444895443e-08 | 5.673278472417987 | 31909.0 |
| 2 | rs62158206 | 114084596 | T | C | 1.4023388158134007e-08 | 5.673102966709871 | 31879.0 |
| 2 | rs11891807 | 48288816 | T | G | 1.4205839848827815e-08 | 5.670889026404753 | 32070.0 |
| 2 | rs986512 | 58275666 | G | A | 1.4279965185570747e-08 | 5.669997441742872 | 31777.0 |
| 2 | rs56163359 | 114071473 | T | C | 1.4560674025713104e-08 | 5.666661323095875 | 31907.0 |
| 2 | rs6545677 | 58211509 | A | G | 1.4686161699637607e-08 | 5.665190109903842 | 32051.0 |
| 2 | rs56093896 | 114103966 | C | A | 1.535550798847022e-08 | 5.657543304931606 | 32082.0 |
| 2 | 2:58002123_CA_C | 58002123 | CA | C | 1.5875374188645882e-08 | 5.651824387789941 | 24995.0 |

| | | | | | | | |
|---|---|---|---|---|---|---|---|
| 2 | rs6712131 | 48256981 | G | C | 1.6663787615773528e-08 | 5.643489311788851 | 32109.0 |
| 2 | rs966721 | 58197305 | T | A | 1.7083062502817758e-08 | 5.639211481577844 | 31335.0 |
| 2 | rs2678899 | 58176213 | A | G | 1.9200464167785536e-08 | 5.619055081979496 | 31990.0 |
| 2 | rs1807282 | 114110036 | A | T | 2.0499543768891307e-08 | 5.607731736120582 | 32027.0 |
| 2 | rs2863957 | 114089551 | C | A | 2.0674322918976568e-08 | 5.6062617511243795 | 32108.0 |
| 2 | rs62158211 | 114106139 | G | T | 2.072762263554e-08 | 5.605815872338141 | 32076.0 |
| 2 | rs1964463 | 114069021 | A | G | 2.102975056585652e-08 | 5.603309268224422 | 31760.0 |
| 2 | rs62158213 | 114110568 | G | A | 2.1822585877712768e-08 | 5.596894142901268 | 31982.0 |
| 2 | rs10188070 | 58299326 | A | G | 2.1986066378411718e-08 | 5.595599536817773 | 31849.0 |
| 2 | rs60758951 | 48285113 | T | C | 2.2509981461125985e-08 | 5.591512708884102 | 32081.0 |
| 2 | rs113257513 | 58283746 | A | ACT | 2.2656522335425278e-08 | 5.5903861294254025 | 31271.0 |
| 2 | rs2717006 | 58146956 | T | A | 2.3555198766426994e-08 | 5.5836282062748674 | 31174.0 |
| 2 | rs4618068 | 114109355 | C | T | 2.5349194254202704e-08 | 5.57085520189422 | 32036.0 |
| 2 | rs848294 | 58381785 | T | C | 2.5663554533216797e-08 | 5.5687077279453865 | 31906.0 |
| 2 | rs7601767 | 48277490 | G | A | 2.56914485633948e-08 | 5.568518411637247 | 32041.0 |
| 2 | rs762952217 | 58284638 | CTCTT | C | 2.619078709026455e-08 | 5.565162737409644 | 31543.0 |
| 2 | rs2312147 | 58222928 | T | C | 2.668416939665912e-08 | 5.561907522799853 | 32186.0 |
| 2 | rs6727241 | 48257032 | T | C | 2.8070161024372345e-08 | 5.553066261065291 | 32103.0 |
| 2 | rs56030357 | 48270092 | T | C | 2.821351458741609e-08 | 5.552176106725428 | 32078.0 |
| 2 | 2:114073498_GGAA_G | 114073498 | GGAA | G | 2.8406660479239024e-08 | 5.550983681510545 | 31461.0 |
| 2 | rs968653 | 58296890 | A | G | 2.90614720191729e-08 | 5.5469988094951335 | 31838.0 |
| 2 | rs1823125 | 114090412 | A | G | 2.952169680898016e-08 | 5.5442498955059145 | 32186.0 |
| 2 | rs62158160 | 114065390 | C | T | 3.039524367751013e-08 | 5.53914469618479 | 32186.0 |
| 2 | rs12620940 | 58311684 | T | C | 3.0423191153335044e-08 | 5.538983725251036 | 32186.0 |
| 2 | rs10172212 | 58379496 | A | C | 3.0698070522181826e-08 | 5.5374080849489165 | 31918.0 |
| 2 | rs75534727 | 58433639 | G | GA | 3.1625279565639913e-08 | 5.532192361892594 | 32008.0 |
| 2 | rs2717043 | 58179019 | C | T | 3.224668192509657e-08 | 5.528779242831654 | 31992.0 |
| 2 | 2:58300783_ATATATT_A | 58300783 | ATATATT | A | 3.274511813306779e-08 | 5.52608733122048 | 31753.0 |
| 2 | rs3771224 | 58460029 | C | T | 3.498155417567774e-08 | 5.514478562280622 | 32005.0 |
| 2 | rs111789636 | 114065433 | A | AC | 3.601482088335681e-08 | 5.5093563267206775 | 31682.0 |
| 2 | rs72816437 | 48274592 | A | T | 3.762596894116509e-08 | 5.501647010267156 | 32083.0 |
| 2 | rs7563432 | 48281138 | G | A | 3.9308476891759804e-08 | 5.493930710268429 | 31948.0 |
| 2 | rs2118899 | 58322226 | A | C | 4.0937073974179195e-08 | 5.486760853121716 | 32072.0 |
| 2 | rs62135450 | 48274591 | G | T | 4.127857499780008e-08 | 5.485292534398298 | 32082.0 |
| 2 | 2:114068802_CTG_C | 114068802 | CTG | C | 4.129171577574041e-08 | 5.485236269898231 | 31521.0 |
| 2 | rs11686281 | 58478157 | G | A | 4.167054967596864e-08 | 5.483621648160112 | 31735.0 |
| 2 | rs848286 | 58394543 | T | C | 4.190861254198733e-08 | 5.482614270732932 | 31944.0 |
| 2 | rs760021402 | 57947592 | A | AG | 4.2054532479580835e-08 | 5.4819995408598805 | 30285.0 |
| 2 | rs10205421 | 58297664 | G | A | 4.223263003054405e-08 | 5.481252049880153 | 31825.0 |
| 2 | 2:58295946_GAA_G | 58295946 | GAA | G | 4.394208145194627e-08 | 5.474228884612881 | 30963.0 |
| 2 | rs1100190 | 58399905 | G | C | 4.4103159897692726e-08 | 5.473580820945632 | 31956.0 |
| 2 | 2:58141384_CT_C | 58141384 | CT | C | 4.67827261922551e-08 | 5.463124115639486 | 30136.0 |
| 2 | rs12477242 | 58452792 | C | G | 4.776684140154312e-08 | 5.45942895954222 | 32088.0 |
| 2 | rs10641725 | 58385305 | T | TCTT A | 4.970693734381633e-08 | 5.452355579274296 | 31724.0 |
| 3 | rs35124509 | 89521693 | T | C | 8.951490491166745e-59 | 16.164670730505776 | 32186.0 |
| 3 | rs67316928 | 89482234 | T | C | 2.2235205955887675e-58 | 16.108500103666593 | 31898.0 |
| 3 | rs12636275 | 89523038 | C | A | 1.7184515800019518e-57 | 15.981540806482005 | 32183.0 |
| 3 | rs7637670 | 89493167 | G | A | 4.092314447437357e-56 | 15.782710354814027 | 31881.0 |
| 3 | rs987748 | 89494030 | C | A | 4.443269278981398e-56 | 15.777516905629911 | 31871.0 |
| 3 | rs11921985 | 89467357 | A | G | 5.58817602232919e-56 | 15.763037034636396 | 31882.0 |
| 3 | rs35968370 | 89466339 | A | T | 7.583394688902125e-56 | 15.743733733295667 | 31913.0 |
| 3 | rs3792572 | 89456555 | T | A | 1.2885955707060912e-55 | 15.710157027986815 | 31862.0 |
| 3 | rs6772953 | 89451721 | G | C | 4.2053007111931054e-55 | 15.63499163743103 | 31733.0 |
| 3 | rs2117137 | 89525505 | A | G | 6.634070713893656e-55 | 15.60592546327857 | 32186.0 |

| | | | | | | | |
|---|---|---|---|---|---|---|---|
| 3 | rs6771054 | 89489529 | T | C | 1.187567416322715e-54 | 15.568721771575609 | 31807.0 |
| 3 | rs7632502 | 89469453 | T | A | 1.3548350705909942e-54 | 15.560290064350387 | 31884.0 |
| 3 | rs7650184 | 89530057 | C | A | 1.4508910942888975e-54 | 15.555905297060995 | 32100.0 |
| 3 | rs7633500 | 89497082 | G | A | 3.353540540788501e-54 | 15.502173269547448 | 31754.0 |
| 3 | rs12637608 | 89551301 | T | C | 2.3682094318701987e-49 | 14.767651875512593 | 32107.0 |
| 3 | rs11918555 | 89541934 | C | T | 2.9732507515879407e-49 | 14.752306906494406 | 32145.0 |
| 3 | rs66757381 | 89561048 | C | T | 6.89975045802501e-49 | 14.695392527204035 | 32087.0 |
| 3 | rs7632427 | 89534377 | T | C | 7.745241742373457e-49 | 14.687560423691096 | 32186.0 |
| 3 | rs7652296 | 89587262 | A | G | 2.038268454556522e-48 | 14.621836584658427 | 32017.0 |
| 3 | rs724972 | 89581408 | A | T | 2.1511326499406396e-48 | 14.61816728392228 | 32029.0 |
| 3 | 3:89598352_GGT_G | 89598352 | GGT | G | 3.926244060484671e-48 | 14.577139870253431 | 31649.0 |
| 3 | rs73145338 | 89594767 | C | T | 1.2489838155625544e-47 | 14.497908197764525 | 31947.0 |
| 3 | 3:89605220_AT_A | 89605220 | AT | A | 1.3528406829227533e-47 | 14.492423510153527 | 31076.0 |
| 3 | rs7621573 | 89595252 | C | T | 1.423214629664844e-47 | 14.488940346980755 | 31919.0 |
| 3 | rs6551413 | 89640350 | C | G | 1.5924487000903963e-47 | 14.481220213742724 | 31866.0 |
| 3 | rs1520596 | 89645542 | T | C | 1.7336831258291088e-47 | 14.475378677028447 | 31804.0 |
| 3 | rs7635916 | 89633140 | G | A | 1.8185171058628946e-47 | 14.472093525538519 | 31929.0 |
| 3 | rs2346843 | 89590826 | C | T | 1.91652838353516e-47 | 14.46848289724526 | 31379.0 |
| 3 | rs73153219 | 89625902 | C | T | 2.0184888284954327e-47 | 14.464916810328946 | 31903.0 |
| 3 | rs66499884 | 89659012 | T | G | 2.1104002592478637e-47 | 14.461852616159408 | 31820.0 |
| 3 | rs6795325 | 89642054 | T | C | 2.153574542599331e-47 | 14.46045881801668 | 31873.0 |
| 3 | rs10511145 | 89597679 | C | A | 2.1770159658898093e-47 | 14.45971366159283 | 31919.0 |
| 3 | rs17027018 | 89604774 | C | T | 4.2359286376946544e-47 | 14.413823989224207 | 31904.0 |
| 3 | rs60243237 | 89694053 | C | T | 1.4182183353612012e-46 | 14.330146149982513 | 31971.0 |
| 3 | rs7637504 | 89702469 | A | T | 1.5850368649410514e-45 | 14.161530662606447 | 31992.0 |
| 3 | rs6551415 | 89705161 | G | T | 1.961831348329957e-45 | 14.14653693495054 | 31983.0 |
| 3 | rs7653735 | 89722185 | T | A | 2.160790038267325e-45 | 14.139740729766256 | 31893.0 |
| 3 | rs11925143 | 89723684 | G | A | 2.160790038267325e-45 | 14.139740729766256 | 31893.0 |
| 3 | 3:89578046_AT_A | 89578046 | AT | A | 2.2104143297724543e-45 | 14.13814272149066 | 31525.0 |
| 3 | rs59381722 | 89701482 | T | G | 2.4554615106874172e-45 | 14.130741211099329 | 31957.0 |
| 3 | rs7613412 | 89724217 | A | T | 1.0552795041617076e-44 | 14.027693656968848 | 31857.0 |
| 3 | 3:89474553_GT_G | 89474553 | GT | G | 1.6916164367285679e-44 | 13.994183344909814 | 29984.0 |
| 3 | rs138643396 | 89698092 | C | CATA | 3.995100932935117e-44 | 13.932949496821193 | 31553.0 |
| 3 | rs6806912 | 89751353 | T | C | 4.635446937112305e-44 | 13.922329682644449 | 31703.0 |
| 3 | rs55747880 | 89612941 | C | A | 1.6963013271361605e-40 | 13.323220141295652 | 30731.0 |
| 3 | rs113141104 | 89519238 | C | T | 2.130285595219879e-34 | 12.230725065484496 | 32136.0 |
| 3 | rs73154904 | 89784599 | G | A | 2.3967314601994482e-34 | 12.221148969219243 | 31956.0 |
| 3 | 3:89521831_TGAAAG_T | 89521831 | TGAAAG | T | 4.358711945226208e-34 | 12.172437081506205 | 31726.0 |
| 3 | 3:89521838_TTTG_T | 89521838 | TTTG | T | 4.358711945226208e-34 | 12.172437081506205 | 31726.0 |
| 3 | rs2881488 | 89500488 | C | G | 4.5873648618979695e-34 | 12.168263763758773 | 32066.0 |
| 3 | rs73139131 | 89524686 | A | G | 4.712229754332276e-34 | 12.166071174532108 | 32169.0 |
| 3 | rs12152372 | 89500365 | T | C | 4.7277310452630885e-34 | 12.165803013136616 | 32067.0 |
| 3 | rs73139119 | 89522285 | C | T | 4.8121989583688715e-34 | 12.16435695313519 | 32175.0 |
| 3 | rs73139121 | 89522433 | A | G | 4.8121989583688715e-34 | 12.16435695313519 | 32175.0 |
| 3 | rs73139125 | 89524056 | A | G | 4.86615850845712e-34 | 12.163446324992645 | 32168.0 |
| 3 | rs752364173 | 89521820 | TTG | T | 5.086378933853708e-34 | 12.159831025145804 | 31522.0 |
| 3 | 3:89521060_CTAT_C | 89521060 | CTAT | C | 5.2711575148126775e-34 | 12.156915572536152 | 32096.0 |
| 3 | rs1912965 | 89518332 | A | G | 5.4441508530182e-34 | 12.154276659348998 | 32174.0 |
| 3 | rs17026944 | 89521131 | G | A | 5.6822267158763476e-34 | 12.150778023459663 | 32174.0 |
| 3 | rs73139105 | 89519651 | T | C | 5.975161131929003e-34 | 12.146667797797086 | 32173.0 |
| 3 | rs73139281 | 89503890 | C | T | 6.294167261135255e-34 | 12.14241349126854 | 32072.0 |
| 3 | rs73139127 | 89524436 | T | A | 6.5079036662616175e-34 | 12.139681278125027 | 32169.0 |
| 3 | rs73139129 | 89524562 | A | G | 6.5079036662616175e-34 | 12.139681278125027 | 32169.0 |
| 3 | rs73139110 | 89521509 | C | A | 6.656015008185531e-34 | 12.137839750925092 | 32175.0 |

| | | | | | | | |
|---|---|---|---|---|---|---|---|
| 3 | rs1054750 | 89521725 | T | C | 6.656015008185531e-34 | 12.137839750925092 | 32175.0 |
| 3 | rs73139115 | 89521900 | G | T | 6.656015008185531e-34 | 12.137839750925092 | 32175.0 |
| 3 | rs73139117 | 89521965 | T | G | 6.656015008185531e-34 | 12.137839750925092 | 32175.0 |
| 3 | rs73139118 | 89522263 | A | T | 6.656015008185531e-34 | 12.137839750925092 | 32175.0 |
| 3 | rs17801380 | 89522483 | G | T | 6.656015008185531e-34 | 12.137839750925092 | 32175.0 |
| 3 | rs568140553 | 89507219 | C | CT | 6.7105871920243474e-34 | 12.137171478228593 | 32166.0 |
| 3 | rs112335559 | 89525297 | A | T | 6.839014297255552e-34 | 12.135619865381939 | 32174.0 |
| 3 | rs1499780 | 89526072 | G | C | 6.8725714606096485e-34 | 12.135219208056874 | 32145.0 |
| 3 | rs73139284 | 89503958 | A | G | 7.385252451907957e-34 | 12.12932850647392 | 32070.0 |
| 3 | rs112297761 | 89521602 | T | TC | 7.40428213065958e-34 | 12.1291177549397 | 32186.0 |
| 3 | rs6795074 | 89516652 | T | C | 7.405766000218071e-34 | 12.129101343872426 | 32173.0 |
| 3 | rs60759140 | 89499848 | G | C | 7.763155003845734e-34 | 12.12524093822266 | 32070.0 |
| 3 | 3:89525668_TTA_T | 89525668 | TTA | T | 8.06368640239917e-34 | 12.122128956075336 | 32167.0 |
| 3 | rs1512183 | 89505054 | A | T | 8.173770519243504e-34 | 12.1210177944235 | 32069.0 |
| 3 | rs144317613 | 89525080 | T | TA | 8.3388110064114e-34 | 12.119379460088535 | 32169.0 |
| 3 | rs73139286 | 89504295 | T | A | 8.524771043880922e-34 | 12.117571616383467 | 32069.0 |
| 3 | rs1912966 | 89518464 | T | A | 8.839216187399613e-34 | 12.114601996365206 | 32174.0 |
| 3 | rs73141108 | 89517119 | C | T | 9.825239001958323e-34 | 12.10592758305944 | 32172.0 |
| 3 | rs73139290 | 89505343 | G | A | 1.0297803921421346e-33 | 12.10207250311624 | 32085.0 |
| 3 | rs73141113 | 89518499 | T | C | 1.065092544241515e-33 | 12.099304856697852 | 32156.0 |
| 3 | rs73139296 | 89505734 | C | T | 1.1231023416837813e-33 | 12.09495027717014 | 32150.0 |
| 3 | rs73139276 | 89500196 | T | A | 1.2492816448238664e-33 | 12.086202969599947 | 32056.0 |
| 3 | rs73139288 | 89505327 | T | G | 1.2733488493443587e-33 | 12.084634669108985 | 32083.0 |
| 3 | rs1157608 | 89506464 | T | G | 1.2767054732686833e-33 | 12.084418283067858 | 32167.0 |
| 3 | rs73139302 | 89508186 | A | G | 1.2767054732686833e-33 | 12.084418283067858 | 32167.0 |
| 3 | rs2346836 | 89498096 | C | A | 1.3199988484344871e-33 | 12.081676914851734 | 32054.0 |
| 3 | rs59541469 | 89501671 | C | T | 1.3232551407927183e-33 | 12.081474349526589 | 32071.0 |
| 3 | rs73141104 | 89508390 | C | T | 1.4430070062537633e-33 | 12.07434957981874 | 32169.0 |
| 3 | rs7626735 | 89506958 | C | A | 1.760382248883153e-33 | 12.057984516973775 | 32167.0 |
| 3 | rs1157607 | 89506364 | C | T | 1.8259277218740954e-33 | 12.05497279599488 | 32165.0 |
| 3 | rs73139104 | 89519229 | A | C | 2.8204062080665332e-33 | 12.019095855578687 | 32089.0 |
| 3 | rs73137387 | 89464194 | T | C | 2.842747603927414e-33 | 12.018443824953088 | 32144.0 |
| 3 | 3:89502123_TTAG_T | 89502123 | TTAG | T | 2.9227864031127075e-33 | 12.016148969924753 | 32014.0 |
| 3 | rs73137381 | 89458550 | A | G | 5.622845142782373e-33 | 11.96194615138918 | 32127.0 |
| 3 | rs73137382 | 89461363 | C | G | 5.685328557449497e-33 | 11.961028585156445 | 32132.0 |
| 3 | rs112087049 | 89509550 | G | T | 6.7184655526595490e-33 | 11.947156671814835 | 31639.0 |
| 3 | rs12639506 | 89480265 | A | G | 6.734043461223811e-33 | 11.946964148289151 | 32060.0 |
| 3 | rs73137384 | 89463290 | C | T | 8.66546209322167e-33 | 11.925983443944228 | 32186.0 |
| 3 | rs1520594 | 89839468 | G | A | 1.4517297871720451e-32 | 11.882938313093266 | 31872.0 |
| 3 | rs1028013 | 89484268 | T | C | 1.851508682777583e-32 | 11.862593020508704 | 32034.0 |
| 3 | rs73137393 | 89471701 | T | C | 1.8734580303555448e-32 | 11.861606424818172 | 32052.0 |
| 3 | rs73139261 | 89492406 | A | G | 1.9480248430977406e-32 | 11.858338435873929 | 32061.0 |
| 3 | rs7650466 | 89530358 | C | T | 1.979202719921633e-32 | 11.857008702969685 | 32155.0 |
| 3 | rs12631313 | 89528461 | A | T | 2.1783627042218823e-32 | 11.848976041520336 | 32156.0 |
| 3 | rs73139135 | 89527937 | T | C | 2.201000048843261e-32 | 11.848109592133696 | 32139.0 |
| 3 | rs12636710 | 89528477 | G | A | 2.522427270223264e-32 | 11.83667964524282 | 32154.0 |
| 3 | rs73139134 | 89527709 | G | C | 2.5283320052248286e-32 | 11.83648348937165 | 32110.0 |
| 3 | rs73139147 | 89530347 | G | A | 2.6164438768368684e-32 | 11.833609247184715 | 32109.0 |
| 3 | rs73139139 | 89528313 | T | G | 2.7083798421587544e-32 | 11.83071096440336 | 32155.0 |
| 3 | rs3762717 | 89528948 | T | C | 2.774317311891175e-32 | 11.828691829388484 | 32156.0 |
| 3 | rs73139144 | 89529177 | G | A | 2.774317311891175e-32 | 11.828691829388484 | 32156.0 |
| 3 | rs73139148 | 89530956 | G | A | 3.2874115402937284e-32 | 11.814437652248008 | 32186.0 |
| 3 | rs1512185 | 89473615 | G | A | 3.700964839236591e-32 | 11.804474358473456 | 32067.0 |
| 3 | rs73139257 | 89482779 | A | G | 4.369686149111807e-32 | 11.790494337320304 | 32067.0 |

| | | | | | | | |
|---|---|---|---|---|---|---|---|
| 3 | rs2117138 | 89531522 | A | G | 4.871407475534304e-32 | 11.781337173322179 | 32096.0 |
| 3 | rs73137395 | 89478753 | A | C | 5.060927565432197e-32 | 11.778119988430715 | 32059.0 |
| 3 | rs73139133 | 89527708 | G | A | 5.834765341784511e-32 | 11.766118834088745 | 32095.0 |
| 3 | rs12634269 | 89532768 | A | C | 6.125068506026581e-32 | 11.762020560058415 | 32051.0 |
| 3 | rs12629165 | 89533017 | G | A | 1.103532834613903e-31 | 11.71221888407099 | 32092.0 |
| 3 | rs12634341 | 89533031 | A | C | 1.714240700584585e-31 | 11.674822052967135 | 32094.0 |
| 3 | rs7616322 | 89493283 | C | T | 2.7285679952090175e-31 | 11.635229014656305 | 32042.0 |
| 3 | rs6793538 | 89987871 | G | A | 4.679446434164816e-31 | 11.589114600238792 | 31188.0 |
| 3 | rs2346840 | 89533267 | T | G | 5.410663614883453e-31 | 11.576670984323128 | 32123.0 |
| 3 | rs6551412 | 89496502 | G | A | 5.959792887169778e-31 | 11.56837912162692 | 32039.0 |
| 3 | rs569866494 | 89526980 | G | GA | 2.2488388462955567e-30 | 11.453867259517214 | 31755.0 |
| 3 | 3:89519266_GA_G | 89519266 | GA | G | 4.773109150773868e-30 | 11.388467439247075 | 30599.0 |
| 3 | rs566480002 | 89529769 | T | TA | 5.416492877368453e-30 | 11.37744249145664 | 31365.0 |
| 3 | rs529961766 | 89520326 | G | GA | 1.0843304562121236e-29 | 11.31673710829741 | 30260.0 |
| 3 | rs6800436 | 89516072 | G | A | 1.449181050169444e-29 | 11.29127556806742 | 30313.0 |
| 3 | rs66708608 | 89486373 | C | T | 2.5664867205642913e-29 | 11.240935296282306 | 32013.0 |
| 3 | rs56163827 | 89612938 | C | A | 1.3964222474385536e-28 | 11.090412574898878 | 30447.0 |
| 3 | rs2048518 | 89533905 | T | C | 8.038473720589526e-28 | 10.9327390764573 | 32180.0 |
| 3 | rs73144998 | 89534592 | G | T | 8.958658375650495e-28 | 10.922902205199994 | 32183.0 |
| 3 | rs12638777 | 89537160 | C | T | 8.958658375650495e-28 | 10.922902205199994 | 32183.0 |
| 3 | rs2048521 | 89534253 | G | A | 8.975192792453661e-28 | 10.922734771102254 | 32181.0 |
| 3 | rs73145002 | 89537618 | T | C | 9.295561610160961e-28 | 10.919549603285034 | 32183.0 |
| 3 | 3:89557699_GTA_G | 89557699 | GTA | G | 1.0925478181620514e-27 | 10.904865451302326 | 32125.0 |
| 3 | rs150837250 | 89522314 | A | AGTT<br>ATAA<br>TTAG | 1.1042594287427312e-27 | 10.903895658106732 | 30757.0 |
| 3 | rs4857499 | 89541330 | C | A | 1.1923476161134748e-27 | 10.896912525664392 | 32162.0 |
| 3 | rs2048519 | 89533978 | G | A | 1.2037307074401732e-27 | 10.8960477120753 | 32181.0 |
| 3 | rs4857498 | 89540977 | G | A | 1.2110706847543614e-27 | 10.89549435881797 | 32162.0 |
| 3 | rs3762718 | 89535066 | T | G | 1.2641847900725812e-27 | 10.89158656337247 | 32185.0 |
| 3 | rs2196083 | 89536546 | T | C | 1.2641847900725812e-27 | 10.89158656337247 | 32185.0 |
| 3 | rs12634044 | 89537651 | A | G | 1.3123367016179446e-27 | 10.888182096847077 | 32185.0 |
| 3 | rs972030 | 89538498 | G | T | 1.3396586199830622e-27 | 10.886305045473962 | 32179.0 |
| 3 | rs373216501 | 89603917 | A | ATATT | 1.367574677841199e-27 | 10.884425988856693 | 27080.0 |
| 3 | rs17026984 | 89544978 | G | A | 1.5218490013063815e-27 | 10.874682379825847 | 32132.0 |
| 3 | rs7633599 | 89544591 | C | A | 1.568441373658271e-27 | 10.871931839205699 | 32152.0 |
| 3 | rs7613315 | 89544773 | G | C | 1.6283616117014822e-27 | 10.86851126198152 | 32151.0 |
| 3 | rs56371328 | 89539124 | G | A | 1.6312453876536898e-27 | 10.868349804691038 | 32181.0 |
| 3 | rs73146961 | 89550459 | T | C | 1.6581293311754257e-27 | 10.866858099295339 | 32131.0 |
| 3 | rs57902683 | 89539011 | C | CA | 1.7661352654864078e-27 | 10.86109759207028 | 32173.0 |
| 3 | rs4857500 | 89541486 | T | C | 1.7751396886867855e-27 | 10.860633230852383 | 32168.0 |
| 3 | rs1036285 | 89544061 | T | C | 1.795792777099463e-27 | 10.85957690751417 | 32161.0 |
| 3 | 3:89541782_TTA_T | 89541782 | TTA | T | 1.8969587483156426e-27 | 10.854570790683413 | 32154.0 |
| 3 | rs12633609 | 89545727 | T | G | 1.9447601976653147e-27 | 10.85229679794777 | 32134.0 |
| 3 | rs1512188 | 89546183 | A | G | 1.9447601976653147e-27 | 10.85229679794777 | 32134.0 |
| 3 | rs73146943 | 89546728 | A | G | 1.9447601976653147e-27 | 10.85229679794777 | 32134.0 |
| 3 | rs73146907 | 89538747 | G | A | 2.116253886018867e-27 | 10.844571414725406 | 32185.0 |
| 3 | 3:89540269_AT_A | 89540269 | AT | A | 2.1434407087129462e-27 | 10.843404043329228 | 32096.0 |
| 3 | rs2117136 | 89542699 | A | G | 2.257819071764473e-27 | 10.838648452146403 | 32165.0 |
| 3 | rs113675293 | 89539536 | C | T | 2.2813820776397823e-27 | 10.837698487511911 | 32182.0 |
| 3 | rs73146926 | 89542293 | A | G | 2.344895402352717e-27 | 10.835185554090504 | 32169.0 |
| 3 | rs138183278 | 89564090 | C | CT | 2.381668974199593e-27 | 10.833761268306048 | 32112.0 |
| 3 | rs138490449 | 89561661 | A | AT | 2.441484682372452e-27 | 10.831490476025945 | 32123.0 |
| 3 | rs55979407 | 89539275 | A | G | 2.4584038954619587e-27 | 10.830858174952263 | 32182.0 |
| 3 | rs1512189 | 89546332 | A | G | 2.472261533654048e-27 | 10.83034349685822 | 32107.0 |
| 3 | rs12632281 | 89547621 | G | A | 2.5413479514140863e-27 | 10.827819548596482 | 32134.0 |

| | | | | | | | |
|---|---|---|---|---|---|---|---|
| 3 | rs73146947 | 89548135 | T | G | 2.5413479514140863e-27 | 10.827819548596482 | 32134.0 |
| 3 | rs73146950 | 89548243 | T | C | 2.5413479514140863e-27 | 10.827819548596482 | 32134.0 |
| 3 | rs1567657 | 89548519 | G | A | 2.5413479514140863e-27 | 10.827819548596482 | 32134.0 |
| 3 | rs1567658 | 89548606 | T | A | 2.5413479514140863e-27 | 10.827819548596482 | 32134.0 |
| 3 | rs1039991 | 89558311 | C | G | 2.5436700424053712e-27 | 10.827735901861944 | 32131.0 |
| 3 | rs73146980 | 89558578 | C | A | 2.5436700424053712e-27 | 10.827735901861944 | 32131.0 |
| 3 | rs73146983 | 89559082 | A | G | 2.5436700424053712e-27 | 10.827735901861944 | 32131.0 |
| 3 | rs6774870 | 89564891 | C | T | 2.5436700424053712e-27 | 10.827735901861944 | 32131.0 |
| 3 | rs73146959 | 89549970 | A | C | 2.576479687754328e-27 | 10.826562053908848 | 32133.0 |
| 3 | rs112404167 | 89550630 | C | T | 2.576479687754328e-27 | 10.826562053908848 | 32133.0 |
| 3 | rs12637511 | 89550844 | T | C | 2.576479687754328e-27 | 10.826562053908848 | 32133.0 |
| 3 | rs73146972 | 89553348 | A | G | 2.576479687754328e-27 | 10.826562053908848 | 32133.0 |
| 3 | rs17026990 | 89555257 | A | C | 2.6234700920161517e-27 | 10.824906418507693 | 32132.0 |
| 3 | rs73146960 | 89550445 | T | C | 2.714492679559277e-27 | 10.821781397676618 | 32017.0 |
| 3 | rs73146954 | 89549680 | C | G | 2.785313970805014e-27 | 10.819420996241135 | 32130.0 |
| 3 | rs73146956 | 89549684 | G | T | 2.785313970805014e-27 | 10.819420996241135 | 32130.0 |
| 3 | rs111618480 | 89552134 | T | C | 2.785313970805014e-27 | 10.819420996241135 | 32130.0 |
| 3 | rs2048520 | 89534248 | T | C | 3.2622582074091816e-27 | 10.804924344952171 | 32186.0 |
| 3 | rs73147002 | 89565128 | T | G | 3.3624115351557566e-27 | 10.802148764132427 | 32130.0 |
| 3 | rs73146976 | 89557065 | T | C | 3.4152739040461376e-27 | 10.800716652592824 | 32133.0 |
| 3 | rs369052772 | 89540272 | T | A | 3.4856607424536485e-27 | 10.798843527469863 | 32146.0 |
| 3 | rs73146992 | 89562178 | A | G | 3.55820571596455e-27 | 10.796951818985901 | 32133.0 |
| 3 | rs73146995 | 89562209 | T | C | 3.55820571596455e-27 | 10.796951818985901 | 32133.0 |
| 3 | rs73146968 | 89552927 | A | G | 3.559918051182249e-27 | 10.796907631000972 | 32134.0 |
| 3 | rs6784577 | 89564308 | A | G | 3.56034626322845e-27 | 10.796896583989813 | 32107.0 |
| 3 | rs1028012 | 89549422 | C | T | 3.596139397862578e-27 | 10.795977819988911 | 32135.0 |
| 3 | rs17745025 | 89549919 | T | C | 3.596139397862578e-27 | 10.795977819988911 | 32135.0 |
| 3 | rs12634697 | 89550993 | G | A | 3.596139397862578e-27 | 10.795977819988911 | 32135.0 |
| 3 | rs12638168 | 89551544 | T | C | 3.596139397862578e-27 | 10.795977819988911 | 32135.0 |
| 3 | rs73146970 | 89553168 | A | G | 3.596139397862578e-27 | 10.795977819988911 | 32135.0 |
| 3 | 3:89544326_CGGTGGAAATCTCCTTATTAAATTGTGGGCAAGAGGCAACCCAT_C | 89544326 | CGGTGGAAATCTCCTTATTAAATTGTGGGCAAGAGGCAACCCAT | C | 3.6140614841923615e-27 | 10.795521184434161 | 32153.0 |
| 3 | rs1039990 | 89558086 | G | A | 3.663697144673911e-27 | 10.79426814613955 | 32132.0 |
| 3 | rs73146988 | 89561044 | T | C | 3.743832503749657e-27 | 10.792280251745398 | 32129.0 |
| 3 | rs12629999 | 89551518 | A | G | 3.866681469009881e-27 | 10.789313242686283 | 32099.0 |
| 3 | rs73146986 | 89560884 | A | G | 3.9481125338653936e-27 | 10.787397626151623 | 32101.0 |
| 3 | rs12629758 | 89556408 | A | G | 4.7695045012755785e-27 | 10.77000979539768 | 32135.0 |
| 3 | rs73149107 | 89565849 | C | A | 4.8838291618050594e-27 | 10.76782870644882 | 32106.0 |
| 3 | 3:89550504_GCT_G | 89550504 | GCT | G | 8.460289213847959e-27 | 10.717113072636256 | 31977.0 |
| 3 | 3:89571180_TAGAG_T | 89571180 | TAGAG | T | 1.1696079257089568e-26 | 10.68710837082095 | 32053.0 |
| 3 | rs56082199 | 89586973 | A | G | 1.3923595360475923e-26 | 10.670923254454554 | 32053.0 |
| 3 | rs12637036 | 89584170 | G | A | 1.4480352542814996e-26 | 10.667279828292077 | 32058.0 |
| 3 | rs7613900 | 89588889 | A | T | 1.4549802246022167e-26 | 10.666835124426909 | 32015.0 |
| 3 | rs73145320 | 89586535 | T | C | 1.539755857096151e-26 | 10.661570203527463 | 32049.0 |
| 3 | rs73145303 | 89568714 | T | C | 1.6013161469637508e-26 | 10.65792417451105 | 32044.0 |
| 3 | rs73145310 | 89578097 | T | C | 1.647701821151799e-26 | 10.655267555634452 | 32063.0 |
| 3 | rs73153218 | 89619808 | G | A | 1.7082761933463243e-26 | 10.651907816874218 | 31826.0 |
| 3 | rs907548 | 89574852 | C | T | 1.7267190726967966e-26 | 10.650908317236627 | 32069.0 |
| 3 | rs73145312 | 89578575 | C | T | 1.7289737472607043e-26 | 10.650786853356921 | 32062.0 |
| 3 | rs73149115 | 89568284 | C | T | 1.7565920462418609e-26 | 10.649311611969566 | 32023.0 |
| 3 | rs73145305 | 89570713 | G | T | 1.9837772710432688e-26 | 10.637982715592027 | 32068.0 |
| 3 | rs73145304 | 89568989 | C | T | 2.0477447933856712e-26 | 10.635024698625793 | 32060.0 |
| 3 | rs73149109 | 89567344 | G | A | 2.1085133089568557e-26 | 10.632298265254553 | 32023.0 |

| | | | | | | | |
|---|---|---|---|---|---|---|---|
| 3 | rs1080592 | 89567568 | T | C | 2.1085133089568557e-26 | 10.632298265254553 | 32023.0 |
| 3 | rs12633325 | 89566836 | A | C | 2.1453889243206907e-26 | 10.63068153462124 | 31972.0 |
| 3 | rs73145316 | 89580407 | C | T | 2.27813383747485e-26 | 10.625081380462733 | 32066.0 |
| 3 | rs73145307 | 89576040 | G | A | 2.388734992769955e-26 | 10.62065716033171 | 32069.0 |
| 3 | rs73145309 | 89576389 | C | T | 2.388734992769955e-26 | 10.62065716033171 | 32069.0 |
| 3 | rs113307255 | 89567826 | T | C | 2.4665604164515568e-26 | 10.617664114037382 | 32024.0 |
| 3 | rs73145328 | 89587686 | T | C | 2.5448092563605392e-26 | 10.61474772156756 | 32036.0 |
| 3 | rs73145332 | 89589327 | A | C | 2.821248872600302e-26 | 10.60511228947913 | 32000.0 |
| 3 | rs73145351 | 89607237 | C | T | 3.1918197968532016e-26 | 10.593569951098264 | 31968.0 |
| 3 | rs73145353 | 89607824 | G | T | 3.1918197968532016e-26 | 10.593569951098264 | 31968.0 |
| 3 | rs73145357 | 89611766 | T | C | 3.193786747773497e-26 | 10.593512301943477 | 31953.0 |
| 3 | rs73149113 | 89567867 | C | T | 3.3829301513281343e-26 | 10.588126962611044 | 32026.0 |
| 3 | rs73145345 | 89598696 | C | A | 3.5897451691503644e-26 | 10.582569943952826 | 31967.0 |
| 3 | rs73145336 | 89593317 | A | G | 4.4587697993596816e-26 | 10.56224325497722 | 31975.0 |
| 3 | rs2069183 | 89645017 | G | A | 4.653348539712592e-26 | 10.55823377885366 | 31917.0 |
| 3 | rs111835258 | 89601496 | G | A | 5.082531926373201e-26 | 10.549947808965586 | 31905.0 |
| 3 | rs148078210 | 89565641 | G | GGTAT | 5.75811034797151e-26 | 10.53821551275048 | 31675.0 |
| 3 | rs149898163 | 89568422 | T | TTTCC | 7.162306516475435e-26 | 10.517669723825133 | 31421.0 |
| 3 | rs2346837 | 89542599 | C | T | 1.0317838392087264e-25 | 10.483212344190592 | 32016.0 |
| 3 | rs73153241 | 89695319 | C | G | 1.327491623932497e-25 | 10.459360080939712 | 31963.0 |
| 3 | rs764684400 | 89443128 | AG | A | 1.335026547193961e-25 | 10.458823740849986 | 30348.0 |
| 3 | rs6551410 | 89442305 | T | A | 1.7743551880347218e-25 | 10.431831091972935 | 31084.0 |
| 3 | rs73153266 | 89720111 | A | G | 2.159855083524976e-25 | 10.413136701393666 | 31978.0 |
| 3 | rs7619303 | 89552554 | G | A | 2.2543435154844878e-25 | 10.40906094453326 | 31595.0 |
| 3 | rs113288647 | 89720352 | C | T | 2.2823775966376477e-25 | 10.407884226107942 | 31896.0 |
| 3 | rs4035593 | 89542601 | T | TAA | 2.3500044075822696e-25 | 10.40510355713966 | 32017.0 |
| 3 | 3:89532085_TA_T | 89532085 | TA | T | 2.513042744421292e-25 | 10.398713040480953 | 24150.0 |
| 3 | rs73146965 | 89552553 | A | G | 2.5882888048917957e-25 | 10.39590105942823 | 31579.0 |
| 3 | rs7617395 | 89722541 | G | T | 3.174711903303252e-25 | 10.376415787949199 | 31896.0 |
| 3 | rs537614374 | 89606871 | T | TA | 4.1124934029191508e-25 | 10.351669897068911 | 31637.0 |
| 3 | rs73153262 | 89719269 | G | T | 4.553872931086922e-25 | 10.341906309116991 | 31979.0 |
| 3 | rs73153258 | 89715585 | G | A | 4.575222065087582e-25 | 10.34145816031753 | 31976.0 |
| 3 | rs1915304 | 89704393 | C | T | 4.736454700573093e-25 | 10.338139100563355 | 31977.0 |
| 3 | rs73153252 | 89712192 | C | T | 5.477206684587232e-25 | 10.324202488290256 | 31974.0 |
| 3 | 3:89626776_AT_A | 89626776 | AT | A | 6.106209203019268e-25 | 10.31376372026957 | 31592.0 |
| 3 | 3:89704472_GCT_G | 89704472 | GCT | G | 6.113704232416414e-25 | 10.313645870272843 | 27881.0 |
| 3 | rs73149103 | 89565651 | G | A | 7.737303881357452e-25 | 10.290994527806085 | 31555.0 |
| 3 | rs1979837 | 89590902 | T | C | 2.4915751946711942e-24 | 10.177789234306914 | 30154.0 |
| 3 | rs12633626 | 89744949 | G | A | 4.7877431650747086e-24 | 10.114022076773246 | 31874.0 |
| 3 | rs73153279 | 89727968 | T | C | 5.8474022923107146e-24 | 10.094422852826892 | 31876.0 |
| 3 | rs73153282 | 89728408 | C | T | 7.32060551870652e-24 | 10.072352115911162 | 31879.0 |
| 3 | rs73153283 | 89732193 | G | T | 1.3213678949018154e-23 | 10.014116821815577 | 31853.0 |
| 3 | 3:89590098_GT_G | 89590098 | GT | G | 1.5469355360374415e-23 | 9.998518748257672 | 31214.0 |
| 3 | rs559264903 | 89578492 | T | TA | 1.6092464119171708e-23 | 9.994606779902801 | 31121.0 |
| 3 | rs111215243 | 89687803 | T | A | 3.149503083359805e-23 | 9.927857359632752 | 31459.0 |
| 3 | rs6787562 | 90010903 | G | C | 7.258310384788044e-23 | 9.844243642993883 | 31183.0 |
| 3 | rs819293 | 89613664 | G | A | 9.255914712165478e-23 | 9.819764907263902 | 31784.0 |
| 3 | rs6773817 | 89753400 | G | A | 1.844826967289189e-21 | 9.513426998689422 | 31935.0 |
| 3 | rs144848844 | 89400269 | A | C | 2.50370527865417e-21 | 9.481617913008145 | 29257.0 |
| 3 | rs6419884 | 89279467 | C | T | 4.17649897663145e-21 | 9.42808353144902 | 31866.0 |
| 3 | rs73153293 | 89756242 | C | T | 6.1617812355941376e-21 | 9.387197656703117 | 31920.0 |
| 3 | rs2279829 | 147106319 | C | T | 7.573075408782388e-21 | 9.365444630642157 | 31996.0 |
| 3 | rs145565219 | 89936402 | G | A | 1.1944220691778567e-20 | 9.317207737267145 | 31737.0 |
| 3 | rs6782527 | 89442369 | C | T | 3.271150066668919e-20 | 9.209674615429613 | 31621.0 |

| | | | | | | | |
|---|---|---|---|---|---|---|---|
| 3 | 3:89694510_GATA_G | 89694510 | GATA | G | 3.433193121359274e-20 | 9.204483189261083 | 30795.0 |
| 3 | rs7645408 | 89446132 | G | A | 4.045385175624901e-20 | 9.18684328542306 | 31543.0 |
| 3 | rs28623022 | 89448936 | T | A | 4.454225705859832e-20 | 9.176477747067702 | 31456.0 |
| 3 | rs9844971 | 89440509 | T | C | 5.262933536408853e-20 | 9.158488384862592 | 31675.0 |
| 3 | rs6800041 | 89448196 | T | C | 7.608277643728348e-20 | 9.118625972109804 | 31475.0 |
| 3 | rs754715969 | 147160015 | T | TAA | 1.0416242268842731e-19 | 9.084514591454818 | 27177.0 |
| 3 | rs201982816 | 89448885 | T | TTA | 2.0911080496906448e-19 | 9.008386209361358 | 30354.0 |
| 3 | rs562459635 | 89448888 | T | TG | 2.0911080496906448e-19 | 9.008386209361358 | 30354.0 |
| 3 | rs9832305 | 89392778 | T | C | 2.3009071192842657e-19 | 8.997893093699291 | 31666.0 |
| 3 | rs9825569 | 89431650 | T | C | 4.2265574396489144e-19 | 8.93087413563343 | 32151.0 |
| 3 | rs62275019 | 89433648 | T | G | 4.903641336735189e-19 | 8.914422729666835 | 32140.0 |
| 3 | 3:89420088_TGTGCACCTAGAC_T | 89420088 | TGTGCACCTAGAC | T | 5.78589359814372e-19 | 8.896070304268948 | 31929.0 |
| 3 | rs57670516 | 89419645 | T | C | 6.132004368999415e-19 | 8.889616750436222 | 32109.0 |
| 3 | rs9864984 | 89428776 | C | T | 6.155152593018303e-19 | 8.889198061030989 | 32186.0 |
| 3 | rs9849029 | 89433059 | A | G | 6.628380410151539e-19 | 8.880963329487672 | 32120.0 |
| 3 | rs9854073 | 89426589 | C | A | 8.892735216675079e-19 | 8.848218577310432 | 32137.0 |
| 3 | rs1512908 | 89340134 | G | A | 9.474330731024583e-19 | 8.841144174158819 | 31919.0 |
| 3 | rs73154916 | 89803713 | C | T | 9.656893114307028e-19 | 8.839011788737258 | 31982.0 |
| 3 | rs62275018 | 89423863 | A | G | 9.801496775915032e-19 | 8.837350843530574 | 32149.0 |
| 3 | rs7615949 | 89432683 | C | G | 1.061883570710988e-18 | 8.828395113155755 | 32129.0 |
| 3 | 3:89924704_TG_T | 89924704 | TG | T | 1.1510019724651969e-18 | 8.819375157488546 | 31633.0 |
| 3 | rs1879650 | 89856887 | G | T | 1.6387896280086018e-18 | 8.77972217451943 | 31965.0 |
| 3 | rs112407114 | 89797398 | G | C | 1.8425355728276288e-18 | 8.766532122875683 | 31939.0 |
| 3 | rs1284258 | 89823487 | C | T | 2.14719992770873e-18 | 8.749279105700076 | 32186.0 |
| 3 | rs73154925 | 89859312 | T | C | 2.1808392770051567e-18 | 8.747524553325581 | 31799.0 |
| 3 | rs73004625 | 147095294 | T | C | 2.423706873173051e-18 | 8.735597901822286 | 31731.0 |
| 3 | rs67707172 | 89447021 | C | A | 2.4966766720480126e-18 | 8.732244551160692 | 30152.0 |
| 3 | rs2175670 | 89343044 | A | G | 3.1296129582609436e-18 | 8.706659556826548 | 31912.0 |
| 3 | rs9863107 | 89343789 | T | G | 3.2462072002572397e-18 | 8.702510804966371 | 31911.0 |
| 3 | rs9852380 | 89341738 | T | G | 3.262505376219695e-18 | 8.70194262282431 | 31908.0 |
| 3 | rs59127701 | 89291916 | T | TA | 3.2754841893712824e-18 | 8.701492160079262 | 31881.0 |
| 3 | rs58858051 | 89291909 | A | T | 3.396892344367472e-18 | 8.69736173935998 | 31880.0 |
| 3 | rs9852722 | 89341946 | T | C | 3.405206809340993e-18 | 8.697084229821387 | 31907.0 |
| 3 | rs13064615 | 89799433 | A | G | 3.4107097332457774e-18 | 8.696900927946766 | 31816.0 |
| 3 | rs9829401 | 89300610 | A | C | 3.7458987955561315e-18 | 8.686253080969497 | 32159.0 |
| 3 | rs6776281 | 89290164 | T | A | 4.0302827024318935e-18 | 8.677932453670369 | 31882.0 |
| 3 | rs73151349 | 89892899 | T | G | 4.048371503249283e-18 | 8.677422993081567 | 31617.0 |
| 3 | 3:89333590_CT_C | 89333590 | CT | C | 4.083687417036498e-18 | 8.676434785195742 | 31781.0 |
| 3 | rs6796339 | 89286159 | C | T | 4.1294159829279396e-18 | 8.675167671839697 | 31868.0 |
| 3 | rs9821666 | 89339548 | G | T | 5.586271478024964e-18 | 8.640714170790641 | 31958.0 |
| 3 | rs1567731 | 89316027 | C | T | 5.812381934196596e-18 | 8.636180168783785 | 32078.0 |
| 3 | rs62278164 | 89292567 | T | C | 5.858693536768759e-18 | 8.635273033201042 | 31930.0 |
| 3 | rs62278163 | 89291836 | C | T | 5.909990595892147e-18 | 8.634276469720103 | 31956.0 |
| 3 | rs73146387 | 89291564 | G | C | 6.4079549890623786e-18 | 8.625023401740412 | 31958.0 |
| 3 | rs6773815 | 89299059 | C | T | 7.708917550385314e-18 | 8.603844844148826 | 32186.0 |
| 3 | rs9863058 | 89322805 | C | T | 8.228066782170996e-18 | 8.596365207498073 | 31978.0 |
| 3 | rs2063590 | 89323932 | G | C | 8.363882389047915e-18 | 8.594485314629805 | 32011.0 |
| 3 | rs9862195 | 89299752 | C | T | 8.653699637624356e-18 | 8.590572564227545 | 32181.0 |
| 3 | rs62278168 | 89301573 | A | G | 9.182605808791333e-18 | 8.58375421517716 | 32155.0 |
| 3 | rs62278170 | 89302617 | G | T | 9.231769796630686e-18 | 8.583140237628704 | 32176.0 |
| 3 | rs4264747 | 89882102 | A | G | 9.264580242684526e-18 | 8.582732281836911 | 31760.0 |
| 3 | rs9875278 | 89335463 | C | T | 9.271056543436971e-18 | 8.582651925874993 | 32015.0 |
| 3 | rs7610812 | 89544397 | G | T | 9.705104529365587e-18 | 8.577388900075999 | 29707.0 |
| 3 | rs62278166 | 89300779 | A | G | 9.92208086755152e-18 | 8.574844443757444 | 32169.0 |

| | | | | | | | |
|---|---|---|---|---|---|---|---|
| 3 | rs62278161 | 89291713 | C | G | 1.0022267787003767e-17 | 8.573688041050904 | 31950.0 |
| 3 | rs9847087 | 89297308 | C | G | 1.052743992685718e-17 | 8.5680255980217 | 32158.0 |
| 3 | rs4857502 | 89324919 | G | A | 1.0531021273177992e-17 | 8.567986419948426 | 32013.0 |
| 3 | rs9851954 | 89297904 | G | A | 1.071998319617582e-17 | 8.565937706431143 | 32172.0 |
| 3 | rs17026816 | 89297593 | A | G | 1.1211909758681766e-17 | 8.56076699500289 | 32166.0 |
| 3 | rs62274986 | 89327148 | G | A | 1.1308317746376124e-17 | 8.559779924041887 | 32018.0 |
| 3 | rs2137487 | 89325520 | C | G | 1.1328164069722655e-17 | 8.559577758972743 | 32017.0 |
| 3 | rs6786328 | 89298926 | A | G | 1.16381596454475e-17 | 8.556464557515906 | 32161.0 |
| 3 | rs113895673 | 89917796 | G | C | 1.1979802133523987e-17 | 8.553126943534943 | 31531.0 |
| 3 | rs4857497 | 89294149 | T | G | 1.2207436491401375e-17 | 8.550954857680994 | 32167.0 |
| 3 | rs6786204 | 89298813 | A | C | 1.2781140153784774e-17 | 8.545653098354157 | 32162.0 |
| 3 | rs9883541 | 89294657 | T | C | 1.2920815928464792e-17 | 8.544397880558034 | 32158.0 |
| 3 | rs62278171 | 89305928 | G | A | 1.4548602030378493e-17 | 8.53068313668469 | 32124.0 |
| 3 | rs13062221 | 89778009 | T | C | 1.5450578954548635e-17 | 8.523722318525921 | 31522.0 |
| 3 | rs28620812 | 89305668 | C | T | 1.698189223331339e-17 | 8.512775324696976 | 32093.0 |
| 3 | rs17738248 | 89296363 | C | A | 1.698764977855183e-17 | 8.51273603231002 | 32144.0 |
| 3 | rs62278172 | 89306038 | G | A | 1.7096344562709778e-17 | 8.511996699837841 | 32099.0 |
| 3 | rs993187 | 89314820 | A | G | 1.7623995448202314e-17 | 8.50847232418279 | 32082.0 |
| 3 | rs17800551 | 89317315 | G | T | 1.8084899014844495e-17 | 8.50547795121253 | 32016.0 |
| 3 | rs13316356 | 89309959 | C | A | 1.8744651902973194e-17 | 8.501320211293205 | 32107.0 |
| 3 | rs17800347 | 89314429 | T | G | 1.8976319374674594e-17 | 8.499894431211366 | 32083.0 |
| 3 | 3:89318905_TC_T | 89318905 | TC | T | 2.1277451210908605e-17 | 8.486597848149406 | 31900.0 |
| 3 | rs62274981 | 89310605 | G | T | 2.2533247923934874e-17 | 8.479928444534966 | 32098.0 |
| 3 | rs62274982 | 89310731 | G | A | 2.290802471486581e-17 | 8.478009002136924 | 32098.0 |
| 3 | rs9866618 | 89319577 | T | C | 2.4572131401868222e-17 | 8.46984427676608 | 32001.0 |
| 3 | rs993186 | 89314862 | T | G | 2.499367594878629e-17 | 8.467862659920447 | 32089.0 |
| 3 | rs9848971 | 89279394 | A | G | 2.5313863138568422e-17 | 8.466379420716931 | 31860.0 |
| 3 | rs373013392 | 89439750 | CA | C | 2.696684065970805e-17 | 8.45900503187241 | 29576.0 |
| 3 | rs62274984 | 89318317 | G | A | 2.754619989965204e-17 | 8.45652552156826 | 32008.0 |
| 3 | rs2137488 | 89308704 | A | G | 2.958124275774189e-17 | 8.448206261721108 | 32086.0 |
| 3 | rs28834186 | 89305272 | G | A | 4.1665012565082885e-17 | 8.408115677146737 | 32023.0 |
| 3 | rs9829367 | 89660010 | A | T | 4.665674966354006e-17 | 8.394830331937165 | 28964.0 |
| 3 | rs6794975 | 89292824 | G | A | 5.622927218930671e-17 | 8.372874935503052 | 32018.0 |
| 3 | rs1503729 | 147098210 | A | G | 6.965716865818574e-17 | 8.347611655838868 | 31870.0 |
| 3 | rs1398534 | 89329307 | A | G | 8.401730566528879e-17 | 8.325438274921273 | 31802.0 |
| 3 | rs1398531 | 89289709 | G | T | 9.920486473570192e-17 | 8.30573327027642 | 31635.0 |
| 3 | 3:89428011_CATCA_C | 89428011 | CATCA | C | 1.0768396030530895e-16 | 8.29599069920798 | 30229.0 |
| 3 | rs1512907 | 89355961 | T | C | 1.1795876848197795e-16 | 8.28515156268433 | 32009.0 |
| 3 | rs73153260 | 89718078 | G | A | 1.217972640429066e-16 | 8.28133964122349 | 31471.0 |
| 3 | rs34756405 | 89344320 | G | GA | 1.651896013077431e-16 | 8.244978729211276 | 31802.0 |
| 3 | rs9310121 | 89378987 | A | G | 1.7942389623803234e-16 | 8.23508920121678 | 31893.0 |
| 3 | rs9833400 | 89389507 | A | G | 1.7989317139833846e-16 | 8.23477649520648 | 31805.0 |
| 3 | rs6768381 | 89358071 | T | C | 1.832139780118669e-16 | 8.23258635790944 | 32047.0 |
| 3 | rs4449340 | 89323876 | C | T | 1.931290901172546e-16 | 8.22627267426393 | 32166.0 |
| 3 | rs13088663 | 89360266 | G | T | 1.9700777459114611e-16 | 8.223889402414693 | 32038.0 |
| 3 | rs9836060 | 89405548 | A | G | 1.97953915108408e-16 | 8.22331506217326 | 29812.0 |
| 3 | rs58036688 | 89369748 | C | CT | 2.0390822675046563e-16 | 8.219761532057811 | 31994.0 |
| 3 | rs772938042 | 89344535 | ACCCT | A | 2.0493209970897519e-16 | 8.219160809819899 | 31819.0 |
| 3 | rs145483821 | 89344540 | T | G | 2.0493209970897519e-16 | 8.219160809819899 | 31819.0 |
| 3 | rs9846844 | 89379639 | G | A | 2.1497145051334567e-16 | 8.21342252392949 | 31872.0 |
| 3 | rs9310122 | 89379408 | A | G | 2.194829151720302e-16 | 8.210929399727403 | 31881.0 |
| 3 | rs528144226 | 89394756 | C | CT | 2.229402377691001e-16 | 8.20905277181706 | 29234.0 |
| 3 | rs9866959 | 89379935 | T | C | 2.4037845562896364e-16 | 8.200004288383518 | 31873.0 |
| 3 | rs7619025 | 89368629 | C | A | 2.4502198462010956e-16 | 8.197703897151381 | 32116.0 |

| | | | | | | | |
|---|---|---|---|---|---|---|---|
| 3 | rs34468928 | 89431649 | A | C | 2.4755876354110865e-16 | 8.196465269918455 | 30247.0 |
| 3 | rs1512911 | 89348525 | T | C | 2.537334383798858e-16 | 8.193501906304915 | 31960.0 |
| 3 | rs59969878 | 89369856 | G | A | 2.758078411351061e-16 | 8.183460006426648 | 32107.0 |
| 3 | rs9310119 | 89378042 | C | G | 2.791650024531702e-16 | 8.182002611233706 | 31917.0 |
| 3 | rs954964 | 89355632 | A | G | 2.802865119614386e-16 | 8.181519593910629 | 31959.0 |
| 3 | rs35173319 | 89386361 | G | T | 2.8348226461549136e-16 | 8.180153608320891 | 31843.0 |
| 3 | rs62274987 | 89334443 | C | A | 2.8573927711274628e-16 | 8.179197990698547 | 32173.0 |
| 3 | rs9877514 | 89330825 | C | T | 2.9444267370133904e-16 | 8.17558135812544 | 32174.0 |
| 3 | rs12495537 | 89347020 | A | G | 2.945043814387409e-16 | 8.175556093978344 | 31955.0 |
| 3 | rs9873545 | 89377717 | A | G | 2.9920316990224404e-16 | 8.173647499802309 | 31932.0 |
| 3 | rs907712 | 89326567 | C | T | 3.03186612567399e-16 | 8.172052469307966 | 32175.0 |
| 3 | rs9289747 | 147100140 | A | C | 3.1209275528822503e-16 | 8.168559773669013 | 31914.0 |
| 3 | rs9845870 | 89345921 | T | C | 3.1299833872270804e-16 | 8.168210152396485 | 31962.0 |
| 3 | rs3852000 | 147116084 | A | G | 3.162629653508853e-16 | 8.166957991405937 | 31946.0 |
| 3 | rs1040017 | 89329632 | T | G | 3.359417418417972e-16 | 8.15966978682056 | 32149.0 |
| 3 | rs1398532 | 89367901 | T | C | 3.3616450787921624e-16 | 8.159589715652498 | 32186.0 |
| 3 | rs9809011 | 89349617 | A | G | 3.4558186461469287e-16 | 8.156251707917361 | 31934.0 |
| 3 | rs9817553 | 89384566 | T | G | 3.4848998068994305e-16 | 8.155239019239993 | 31827.0 |
| 3 | rs35448293 | 89386010 | A | G | 3.5644911563780554e-16 | 8.152509440132665 | 31821.0 |
| 3 | rs1398535 | 89326021 | G | A | 3.6600546633079487e-16 | 8.149310373353723 | 32174.0 |
| 3 | rs13076853 | 89382776 | A | C | 3.7202290092231295e-16 | 8.147337958248695 | 31838.0 |
| 3 | rs11923303 | 89380944 | G | A | 3.97422124159334e-16 | 8.139344969200003 | 31870.0 |
| 3 | rs9866945 | 89349818 | T | C | 4.189703155278116e-16 | 8.132949176904534 | 31909.0 |
| 3 | rs7644070 | 89370107 | G | A | 4.2047960137992436e-16 | 8.132513428276992 | 32113.0 |
| 3 | rs9990016 | 89364033 | G | A | 4.2105476304644517e-16 | 8.132347777834422 | 32085.0 |
| 3 | rs9833437 | 89283869 | C | T | 4.272137974683421e-16 | 8.130587783303412 | 31710.0 |
| 3 | rs1512910 | 89332453 | A | G | 4.335791725058324e-16 | 8.128794909007764 | 32186.0 |
| 3 | rs1016584 | 89343169 | C | T | 4.338585445549007e-16 | 8.128716816409355 | 31968.0 |
| 3 | rs28675360 | 89386151 | C | T | 4.740638079629221e-16 | 8.117965423038127 | 31818.0 |
| 3 | rs12636413 | 147191972 | G | C | 4.865408802094442e-16 | 8.11481114327969 | 31922.0 |
| 3 | rs13100388 | 89383148 | T | A | 4.956322786734662e-16 | 8.11256258932043 | 31850.0 |
| 3 | rs35511518 | 89383683 | T | C | 5.085838501549347e-16 | 8.109428526949532 | 31834.0 |
| 3 | rs2089076 | 147180677 | C | A | 5.101070326088081e-16 | 8.10906512345137 | 31925.0 |
| 3 | rs11128072 | 89317917 | C | T | 6.3884032510978625e-16 | 8.081673353162671 | 32063.0 |
| 3 | rs67851870 | 17554860 | A | G | 6.57350173000865e-16 | 8.078190213181902 | 31950.0 |
| 3 | rs62273364 | 147111779 | T | C | 6.724550735472953e-16 | 8.075418685477322 | 31938.0 |
| 3 | rs17800641 | 89320454 | A | G | 6.843340594797125e-16 | 8.073281820694675 | 32063.0 |
| 3 | rs112856289 | 89404980 | T | A | 6.848446444420301e-16 | 8.073190794781219 | 29824.0 |
| 3 | rs994129 | 89318462 | A | G | 6.891700170909206e-16 | 8.0724223490608 | 32061.0 |
| 3 | rs9879028 | 89273019 | G | A | 7.164296633585105e-16 | 8.067686078715036 | 31785.0 |
| 3 | rs9289750 | 147169065 | C | G | 7.265112748616537e-16 | 8.065979287593262 | 31877.0 |
| 3 | rs56318686 | 147171254 | G | A | 7.351453096910442e-16 | 8.06453601693495 | 31871.0 |
| 3 | rs56224184 | 147171246 | T | G | 7.635790380704428e-16 | 8.059898371865003 | 31878.0 |
| 3 | rs6551409 | 89302171 | G | T | 7.952287630892564e-16 | 8.054932200088661 | 32119.0 |
| 3 | rs9310117 | 89275906 | T | C | 8.017158987083697e-16 | 8.053938383547427 | 31866.0 |
| 3 | rs9853389 | 89312308 | T | C | 8.339574438911184e-16 | 8.049113702125544 | 32011.0 |
| 3 | rs9838512 | 89284596 | G | A | 8.524184048897732e-16 | 8.046433256280805 | 31918.0 |
| 3 | rs16859501 | 147200660 | G | C | 8.82173154810002e-16 | 8.042231103449511 | 31773.0 |
| 3 | rs6771477 | 89293142 | C | A | 9.446929045130196e-16 | 8.033838810474531 | 32004.0 |
| 3 | rs9836340 | 89405754 | A | G | 9.63206523303085e-16 | 8.031458505064485 | 29233.0 |
| 3 | rs9868782 | 89384121 | G | A | 1.0247886729169184e-15 | 8.023853072235863 | 31706.0 |
| 3 | 3:89302973_GT_G | 89302973 | GT | G | 1.0424410622937103e-15 | 8.021755940503175 | 32047.0 |
| 3 | rs9842599 | 89407256 | A | G | 1.070151135273004e-15 | 8.018537616317486 | 29280.0 |
| 3 | 3:147180141_CT_C | 147180141 | CT | C | 1.0895012780350197e-15 | 8.01633159742598 | 31217.0 |

| | | | | | | | |
|---|---|---|---|---|---|---|---|
| 3 | rs9817310 | 89280754 | C | T | 1.0897083731700632e-15 | 8.016308240682188 | 31880.0 |
| 3 | rs6771336 | 89293015 | C | T | 1.0923097951196077e-15 | 8.016015217231583 | 32000.0 |
| 3 | rs6770978 | 89286172 | G | A | 1.1091590005535587e-15 | 8.014133812173496 | 31841.0 |
| 3 | rs1567732 | 89315893 | G | A | 1.1184622415019417e-15 | 8.013107028642871 | 32060.0 |
| 3 | rs4857496 | 89294023 | A | T | 1.145918117882314e-15 | 8.010125107476027 | 32109.0 |
| 3 | rs9837346 | 89280847 | T | C | 1.1544399637944396e-15 | 8.009213869868061 | 31880.0 |
| 3 | rs9809046 | 89273974 | G | C | 1.1557965041419524e-15 | 8.009069427187951 | 31835.0 |
| 3 | rs9877953 | 89295480 | A | T | 1.1710152669556687e-15 | 8.007460302623857 | 32111.0 |
| 3 | rs7631682 | 89394143 | A | C | 1.440783058665348e-15 | 7.981916655272943 | 29214.0 |
| 3 | rs9833875 | 147134032 | A | G | 1.450201728718655e-15 | 7.981112534032029 | 31890.0 |
| 3 | rs1512912 | 89307931 | C | T | 1.453104338529484e-15 | 7.980865759714201 | 32061.0 |
| 3 | rs58262807 | 147109829 | T | C | 1.4719346338627097e-15 | 7.979276539973727 | 31940.0 |
| 3 | rs150021232 | 89408982 | T | C | 1.6503407680220206e-15 | 7.965142056722393 | 29889.0 |
| 3 | rs9860266 | 89394547 | A | T | 1.7672933374597318e-15 | 7.956671343700322 | 29206.0 |
| 3 | rs9867299 | 89300497 | G | A | 1.7787347082262198e-15 | 7.955872523474218 | 31952.0 |
| 3 | rs9818183 | 147174864 | T | C | 2.049509435997693e-15 | 7.9383121969436585 | 31922.0 |
| 3 | rs1566797 | 147172921 | T | G | 2.0985157931753328e-15 | 7.935380133841058 | 31880.0 |
| 3 | rs11720653 | 17439510 | T | C | 2.2004373846671023e-15 | 7.929492223330338 | 32186.0 |
| 3 | rs6765994 | 147189001 | A | G | 2.303109023185991e-15 | 7.923826511492237 | 31918.0 |
| 3 | rs28885734 | 89305269 | A | T | 2.3034253997348907e-15 | 7.923809440276942 | 31961.0 |
| 3 | rs6802591 | 147194290 | T | C | 2.328795664892687e-15 | 7.922447960454446 | 31915.0 |
| 3 | rs13321297 | 17528764 | T | A | 2.7982495804173758e-15 | 7.899588828893387 | 31990.0 |
| 3 | rs9870741 | 17407930 | T | G | 2.820351945654182e-15 | 7.898608066182803 | 32113.0 |
| 3 | rs9820816 | 147170593 | T | C | 2.8330456111279296e-15 | 7.898048217666086 | 31878.0 |
| 3 | rs17801001 | 89414555 | A | C | 3.1065918857493598e-15 | 7.886548364947201 | 29381.0 |
| 3 | rs137962487 | 147220939 | T | TTC | 3.6126883903627855e-15 | 7.86768325878388 | 31715.0 |
| 3 | rs73145317 | 89583712 | T | C | 3.70079807346588e-15 | 7.864667260354607 | 31386.0 |
| 3 | rs7428532 | 89281752 | G | A | 4.78593526501675e-15 | 7.83241321968594 | 31656.0 |
| 3 | rs9852407 | 147148217 | A | G | 4.9562633891476926e-15 | 7.8280166437604946 | 31905.0 |
| 3 | rs56778250 | 147139132 | A | G | 5.153512751591107e-15 | 7.8231073337130175 | 31908.0 |
| 3 | rs4377462 | 147164159 | T | C | 5.215831066851714e-15 | 7.821594700014725 | 31935.0 |
| 3 | rs2202286 | 147149477 | A | T | 5.310213780059834e-15 | 7.819337320963037 | 31917.0 |
| 3 | rs9680882 | 147154084 | A | G | 5.375600861317765e-15 | 7.817796480680863 | 31922.0 |
| 3 | rs62275016 | 89414583 | A | G | 5.384633121519347e-15 | 7.817585088019173 | 29386.0 |
| 3 | 3:147129057_AAC_A | 147129057 | AAC | A | 5.677729225187083e-15 | 7.8109083480712425 | 31668.0 |
| 3 | rs62275015 | 89408594 | C | A | 6.3607084979239944e-15 | 7.796580830294457 | 29297.0 |
| 3 | rs9289749 | 147142513 | A | G | 6.881875997838025e-15 | 7.786632473971232 | 31895.0 |
| 3 | rs872976 | 89326991 | C | A | 7.6798388622911146e-15 | 7.772753029236783 | 32100.0 |
| 3 | rs576273791 | 89285743 | A | AG | 8.246924495012534e-15 | 7.763727106720131 | 30860.0 |
| 3 | rs870899 | 89417171 | C | T | 9.568840028389665e-15 | 7.744858429243394 | 29488.0 |
| 3 | rs870898 | 89417064 | C | T | 1.1842286210217828e-14 | 7.717726612310463 | 29491.0 |
| 3 | rs17801031 | 89415802 | C | A | 1.2870349062480923e-14 | 7.707105411881276 | 29417.0 |
| 3 | rs535492055 | 147147025 | G | GA | 1.3033953230812089e-14 | 7.705492597513938 | 31263.0 |
| 3 | rs9289748 | 147137976 | T | G | 1.3227318874282768e-14 | 7.703611882766221 | 31902.0 |
| 3 | rs55787722 | 90012103 | C | T | 1.4987543019569104e-14 | 7.687638615981567 | 31155.0 |
| 3 | rs111211173 | 89660044 | T | C | 1.717167251768538e-14 | 7.670208697012378 | 26868.0 |
| 3 | rs9862638 | 89373598 | T | C | 2.0129562018472328e-14 | 7.649797776205621 | 31860.0 |
| 3 | rs1521800 | 90001588 | G | T | 2.157082583690568e-14 | 7.640900176030357 | 32186.0 |
| 3 | rs9880313 | 89373637 | A | G | 2.4448913188963245e-14 | 7.624759836514331 | 31857.0 |
| 3 | rs35317084 | 89354462 | C | T | 3.72354644931431e-14 | 7.5703030194439815 | 30628.0 |
| 3 | rs112252173 | 89525767 | A | T | 3.7962811043889496e-14 | 7.5677896247569345 | 26047.0 |
| 3 | rs11916858 | 147140680 | T | G | 6.068879942107278e-14 | 7.506587022094298 | 31932.0 |
| 3 | rs68153678 | 147203820 | A | G | 9.462480284891812e-14 | 7.448196516358862 | 31730.0 |
| 3 | rs12638746 | 89331055 | G | A | 1.0074834603733543e-13 | 7.439917307879063 | 31643.0 |

| | | | | | | | |
|---|---|---|---|---|---|---|---|
| 3 | rs9310120 | 89378413 | T | C | 1.1062104678739018e-13 | 7.427557803255984 | 31754.0 |
| 3 | rs9869999 | 17717340 | G | T | 1.2280475406069287e-13 | 7.413720326905844 | 31820.0 |
| 3 | rs138801589 | 89561919 | A | AGT | 1.5513721632938339e-13 | 7.3826777408603155 | 22126.0 |
| 3 | rs752023841 | 89618888 | TA | T | 1.6449582191582022e-13 | 7.37487783334365 | 22221.0 |
| 3 | rs9858940 | 147207565 | C | T | 2.0531881707418883e-13 | 7.345286503310769 | 31610.0 |
| 3 | rs112604674 | 89989681 | C | T | 2.1128943782207192e-13 | 7.341451679415924 | 31173.0 |
| 3 | rs924761 | 147224071 | G | C | 2.153897950954789e-13 | 7.338879266459442 | 31395.0 |
| 3 | rs7640489 | 147223564 | A | G | 2.1884763941897758e-13 | 7.336747061559472 | 31391.0 |
| 3 | rs9858966 | 147207613 | C | T | 2.2890964728783343e-13 | 7.3307257678749025 | 31611.0 |
| 3 | rs9883675 | 90310464 | A | G | 2.6314893301971564e-13 | 7.312023571601587 | 31657.0 |
| 3 | rs7631038 | 147220482 | A | G | 3.0904221471336187e-13 | 7.290397341398351 | 31632.0 |
| 3 | rs924763 | 147218975 | T | C | 3.161924299830873e-13 | 7.287315259977714 | 31633.0 |
| 3 | rs13085441 | 17738547 | T | C | 3.1894025473255797e-13 | 7.286148991381482 | 31883.0 |
| 3 | rs9847935 | 147212708 | A | G | 3.2558273554026603e-13 | 7.283369968645599 | 31643.0 |
| 3 | rs12632762 | 147209965 | A | C | 3.3132928390496866e-13 | 7.281010339941561 | 31639.0 |
| 3 | rs9289752 | 147212354 | G | A | 3.3338712775669663e-13 | 7.280175117567388 | 31642.0 |
| 3 | rs4611784 | 147220252 | G | C | 3.597281601836409e-13 | 7.2699095967359675 | 31630.0 |
| 3 | rs6788190 | 17786057 | T | C | 4.808135006133577e-13 | 7.230614473825928 | 31655.0 |
| 3 | rs9844783 | 147220661 | T | G | 5.596292760886334e-13 | 7.20997310894746 | 31616.0 |
| 3 | rs9682090 | 90315330 | C | T | 5.753782715324425e-13 | 7.206193021222365 | 31927.0 |
| 3 | rs6797260 | 89302486 | A | T | 6.150411580820877e-13 | 7.197105632084337 | 31635.0 |
| 3 | rs9713415 | 90307719 | C | G | 6.555171023066513e-13 | 7.188406797270695 | 31946.0 |
| 3 | rs6551461 | 90286317 | T | A | 6.872856080532412e-13 | 7.181941043501119 | 32051.0 |
| 3 | rs9310523 | 17626394 | C | T | 7.182387975948142e-13 | 7.175917459195283 | 31999.0 |
| 3 | rs9865864 | 147224629 | T | C | 7.277636008166256e-13 | 7.1741151080564585 | 31502.0 |
| 3 | rs9985472 | 17681459 | A | G | 7.877209025694796e-13 | 7.163275024742397 | 31984.0 |
| 3 | rs7637642 | 90034785 | C | G | 8.056250635593625e-13 | 7.160194786552596 | 31876.0 |
| 3 | rs28708984 | 90041154 | T | G | 8.375607221247537e-13 | 7.1548637453750885 | 31887.0 |
| 3 | rs61039064 | 90044732 | C | G | 8.560640792937313e-13 | 7.151865532335742 | 31927.0 |
| 3 | rs13075167 | 89303172 | T | C | 8.797288157993054e-13 | 7.148122359079532 | 31620.0 |
| 3 | rs9855515 | 17679106 | G | A | 8.947197393795441e-13 | 7.1458019627795935 | 31987.0 |
| 3 | 3:17727954_CTAAT_C | 17727954 | CTAAT | C | 9.236661468926549e-13 | 7.141427463626982 | 31734.0 |
| 3 | rs6804837 | 17787399 | G | A | 9.995804976644642e-13 | 7.130564597352464 | 31733.0 |
| 3 | rs4908969 | 17779407 | T | C | 1.256481209935759e-12 | 7.099016158026602 | 31843.0 |
| 3 | rs6577617 | 17715644 | G | T | 1.260188911620979e-12 | 7.098608875569947 | 31751.0 |
| 3 | rs13083917 | 17713516 | T | A | 1.3968217470519504e-12 | 7.0843661195827625 | 31932.0 |
| 3 | rs9681644 | 17659105 | C | T | 1.398782844405605e-12 | 7.0841718079927185 | 31340.0 |
| 3 | rs9631465 | 147107606 | A | G | 1.439950419803685e-12 | 7.080153337097938 | 31934.0 |
| 3 | rs9681893 | 17757024 | T | A | 1.4689738684642383e-12 | 7.0773874708478015 | 31896.0 |
| 3 | 3:17796107_TG_T | 17796107 | TG | T | 1.495777063840084e-12 | 7.074880402103785 | 31585.0 |
| 3 | rs4431138 | 17749777 | G | C | 1.5043214900468253e-12 | 7.074090447549058 | 31898.0 |
| 3 | rs12634674 | 17704685 | T | G | 1.507208674578149e-12 | 7.073824514545777 | 31935.0 |
| 3 | rs6577619 | 17715678 | C | T | 1.5442490690737242e-12 | 7.070456438785641 | 31758.0 |
| 3 | rs11717504 | 90179080 | A | T | 1.6076911427963519e-12 | 7.064867679630378 | 31740.0 |
| 3 | rs4908963 | 17758260 | T | C | 1.6195377460790889e-12 | 7.063848104993688 | 31887.0 |
| 3 | rs12631291 | 17758297 | G | A | 1.6195377460790889e-12 | 7.063848104993688 | 31887.0 |
| 3 | rs34786175 | 17717997 | C | T | 1.6491084078515494e-12 | 7.061334699156908 | 31768.0 |
| 3 | 3:17758749_CA_C | 17758749 | CA | C | 1.6788074629960413e-12 | 7.058854495169465 | 31235.0 |
| 3 | rs6802945 | 90270589 | A | G | 1.708952710959944e-12 | 7.0563806639797235 | 31957.0 |
| 3 | rs556500778 | 17720571 | GA | G | 1.760832064967557e-12 | 7.052221812364813 | 31454.0 |
| 3 | rs9819028 | 147219323 | G | A | 1.8265461440229507e-12 | 7.047123182520893 | 31421.0 |
| 3 | rs7617683 | 17728919 | T | C | 1.8653144782180828e-12 | 7.0441989657435835 | 31881.0 |
| 3 | rs35423549 | 17739615 | T | C | 1.8681049163922254e-12 | 7.043990794629867 | 31850.0 |
| 3 | rs6803312 | 17771005 | T | C | 1.890136975972773e-12 | 7.042357799922111 | 31846.0 |

| | | | | | | | |
|---|---|---|---|---|---|---|---|
| 3 | rs113165984 | 89989689 | T | C | 1.8944097059398822e-12 | 7.042043272121112 | 30928.0 |
| 3 | rs4908966 | 17773919 | A | G | 1.909539316601717e-12 | 7.0409351059565175 | 31838.0 |
| 3 | rs7431897 | 17762335 | G | T | 1.935316599957125e-12 | 7.0390667417514905 | 31885.0 |
| 3 | rs7652156 | 17780490 | C | G | 2.0115942432398438e-12 | 7.033677748388014 | 31843.0 |
| 3 | rs11929111 | 17777546 | A | G | 2.022007160615868e-12 | 7.0329576768375714 | 31844.0 |
| 3 | rs6577644 | 17766733 | A | G | 2.0998399577532326e-12 | 7.027687856842192 | 31848.0 |
| 3 | rs6577643 | 17766697 | C | G | 2.1073021945322184e-12 | 7.0271927177473446 | 31847.0 |
| 3 | rs9880456 | 17777123 | A | G | 2.1526638426927968e-12 | 7.024219359353872 | 31847.0 |
| 3 | rs10689924 | 17696951 | T | TTA | 2.1639893063691348e-12 | 7.023486588274659 | 32018.0 |
| 3 | rs4482690 | 17767046 | T | A | 2.20612838985696e-12 | 7.0207927846632945 | 31850.0 |
| 3 | rs34990533 | 17718227 | T | C | 2.214816467403188e-12 | 7.020243665938968 | 31761.0 |
| 3 | rs79003773 | 90348777 | T | G | 2.221167641618076e-12 | 7.0198435832888855 | 31446.0 |
| 3 | rs12632570 | 17763501 | A | C | 2.304114648535416e-12 | 7.014719057717426 | 31845.0 |
| 3 | rs57309958 | 17732035 | T | G | 2.3468230580647905e-12 | 7.012150647658568 | 31858.0 |
| 3 | rs9681232 | 17760883 | A | G | 2.3506260322643044e-12 | 7.011924170709113 | 31888.0 |
| 3 | rs11128839 | 17703391 | C | T | 2.351323276438534e-12 | 7.011882687001928 | 31932.0 |
| 3 | rs4909012 | 17756467 | C | A | 2.361352221301839e-12 | 7.011287329322928 | 31896.0 |
| 3 | rs4909010 | 17736023 | G | A | 2.3616550670433108e-12 | 7.011269389806065 | 31893.0 |
| 3 | rs4908959 | 17736035 | G | A | 2.3616550670433108e-12 | 7.011269389806065 | 31893.0 |
| 3 | rs12497320 | 17754558 | G | A | 2.373324845097752e-12 | 7.0105798262162775 | 31890.0 |
| 3 | rs6810235 | 17753801 | C | A | 2.383427441572933e-12 | 7.009985547191025 | 31890.0 |
| 3 | rs61081795 | 17709275 | T | C | 2.397738599300175e-12 | 7.009147917180012 | 31933.0 |
| 3 | rs4513482 | 17708780 | C | T | 2.401854078446696e-12 | 7.008907946453973 | 31933.0 |
| 3 | rs4243835 | 17706179 | A | G | 2.402835900039937e-12 | 7.008850756701626 | 31934.0 |
| 3 | rs4243836 | 17706261 | C | T | 2.402835900039937e-12 | 7.008850756701626 | 31934.0 |
| 3 | rs7632240 | 17711352 | C | A | 2.412482636786042e-12 | 7.008290063331605 | 31922.0 |
| 3 | rs4566567 | 17749177 | C | T | 2.418114650889453e-12 | 7.007963731833024 | 31898.0 |
| 3 | rs4568172 | 17749661 | A | G | 2.418114650889453e-12 | 7.007963731833024 | 31898.0 |
| 3 | rs35642812 | 17741409 | C | G | 2.4621218442585295e-12 | 7.0054392308914375 | 31893.0 |
| 3 | rs199634373 | 17741172 | A | AC | 2.484093018765783e-12 | 7.004195356346671 | 31850.0 |
| 3 | rs7633601 | 17734215 | T | C | 2.4877172135095427e-12 | 7.003991213714072 | 31904.0 |
| 3 | rs4589965 | 17709985 | C | T | 2.4987612866898425e-12 | 7.0033709201172725 | 31937.0 |
| 3 | rs10510477 | 17721297 | T | C | 2.5279906791613894e-12 | 7.001742128081176 | 31891.0 |
| 3 | rs6577622 | 17724339 | T | C | 2.545156197569473e-12 | 7.0007941757539545 | 31893.0 |
| 3 | rs11128843 | 17736281 | C | T | 2.5840509709595368e-12 | 6.99866923681335 | 31853.0 |
| 3 | rs12632939 | 17741107 | C | A | 2.5840509709595368e-12 | 6.99866923681335 | 31853.0 |
| 3 | rs7630695 | 17726483 | A | G | 2.594998193360649e-12 | 6.998076812530364 | 31897.0 |
| 3 | rs4908958 | 17728043 | G | A | 2.596245092785621e-12 | 6.99800949036788 | 31898.0 |
| 3 | rs34297417 | 17728287 | T | C | 2.596245092785621e-12 | 6.99800949036788 | 31898.0 |
| 3 | rs4128509 | 17731099 | T | C | 2.596245092785621e-12 | 6.99800949036788 | 31898.0 |
| 3 | rs12487916 | 17733795 | G | C | 2.596245092785621e-12 | 6.99800949036788 | 31898.0 |
| 3 | rs13071934 | 17757928 | G | C | 2.59792934643583e-12 | 6.997918605053169 | 31887.0 |
| 3 | rs4908964 | 17758859 | C | T | 2.59792934643583e-12 | 6.997918605053169 | 31887.0 |
| 3 | rs11128844 | 17736387 | C | T | 2.611567756004917e-12 | 6.997184773649369 | 31896.0 |
| 3 | rs4243838 | 17759733 | T | C | 2.6168836895469927e-12 | 6.996899760807812 | 31894.0 |
| 3 | rs6809062 | 17711772 | G | A | 2.639479772359708e-12 | 6.9956945783870985 | 31933.0 |
| 3 | rs6797952 | 17735318 | A | G | 2.642502887043374e-12 | 6.995534105800493 | 31901.0 |
| 3 | rs763203017 | 17688987 | TAAGAA | T | 2.6448942023750945e-12 | 6.99540729778978 | 32011.0 |
| 3 | rs8179983 | 17764309 | C | T | 2.6468850677449966e-12 | 6.995301810806179 | 31892.0 |
| 3 | rs34487714 | 17763899 | T | C | 2.6560323793145802e-12 | 6.994818133928543 | 31893.0 |
| 3 | rs11128841 | 17721451 | C | T | 2.705528724414522e-12 | 6.992228953309332 | 31892.0 |
| 3 | rs4602424 | 17722619 | A | T | 2.705528724414522e-12 | 6.992228953309332 | 31892.0 |
| 3 | rs13085167 | 17717405 | C | T | 2.709320310619848e-12 | 6.9920325331452140 | 31897.0 |
| 3 | rs35697509 | 17763957 | G | T | 2.7721460606280917e-12 | 6.988816552683426 | 31893.0 |

| | | | | | | | |
|---|---|---|---|---|---|---|---|
| 3 | rs9816920 | 17739024 | A | C | 2.8477768888937683e-12 | 6.9850385924487374 | 31898.0 |
| 3 | rs11712168 | 17736700 | G | C | 2.851514577062112e-12 | 6.984854445981207 | 31858.0 |
| 3 | rs4909009 | 17720104 | C | A | 2.8518566807052323e-12 | 6.984837603219857 | 31892.0 |
| 3 | rs6577616 | 17715524 | C | T | 2.8743415464246065e-12 | 6.9837349298245455 | 31928.0 |
| 3 | rs13097908 | 17719764 | A | G | 2.9077699423113257e-12 | 6.982111115868727 | 31896.0 |
| 3 | rs13065637 | 17710414 | T | C | 2.9110733219957922e-12 | 6.9819516463721705 | 31930.0 |
| 3 | rs11927637 | 17781315 | A | G | 2.923272014544493e-12 | 6.981364292428776 | 31840.0 |
| 3 | rs11719266 | 17781576 | A | G | 2.9397957471945867e-12 | 6.980572513770193 | 31839.0 |
| 3 | rs7622878 | 17738416 | C | T | 2.9534353141073288e-12 | 6.979922218694592 | 31893.0 |
| 3 | rs6795189 | 17711724 | T | A | 2.9989462960844103e-12 | 6.9777734963060185 | 31931.0 |
| 3 | 3:17758810_CTTTA_C | 17758810 | CTTTA | C | 3.18952794393819e-12 | 6.969109760175615 | 31795.0 |
| 3 | 3:89357752_AT_A | 89357752 | AT | A | 3.2073589858498615e-12 | 6.968325314580415 | 24303.0 |
| 3 | rs11386338 | 17676494 | C | CA | 3.3371221038424933e-12 | 6.962742185695863 | 32008.0 |
| 3 | 3:17761502_TAA_T | 17761502 | TAA | T | 3.3504953358339763e-12 | 6.962178944164588 | 31835.0 |
| 3 | rs62275606 | 90355890 | A | G | 3.363706721928822e-12 | 6.961624679466237 | 31668.0 |
| 3 | rs555285785 | 17745398 | T | G | 3.5194864238576627e-12 | 6.955245483487438 | 31822.0 |
| 3 | rs373576726 | 17745406 | GT | G | 3.5194864238576627e-12 | 6.955245483487438 | 31822.0 |
| 3 | rs4484221 | 17702593 | C | G | 3.53888525397634e-12 | 6.954470561666048 | 31946.0 |
| 3 | rs12488591 | 17795931 | T | C | 3.653202940045209e-12 | 6.949986838215603 | 31718.0 |
| 3 | rs4908972 | 17796192 | T | C | 3.653202940045209e-12 | 6.949986838215603 | 31718.0 |
| 3 | rs13064990 | 17696489 | A | G | 3.8332351814758496e-12 | 6.943197278877039 | 31691.0 |
| 3 | 3:90299284_CAT_C | 90299284 | CAT | C | 4.055873215682047e-12 | 6.935220749547025 | 32003.0 |
| 3 | rs4908939 | 17793850 | G | A | 4.0920831680837146e-12 | 6.9339641774377325 | 31711.0 |
| 3 | rs6789697 | 17700708 | G | A | 4.175326011720528e-12 | 6.931116301074304 | 31999.0 |
| 3 | rs9867064 | 17792574 | C | T | 4.359820472076567e-12 | 6.924997881050645 | 31714.0 |
| 3 | rs4435674 | 17794200 | C | T | 4.396610975366511e-12 | 6.923808199165312 | 31715.0 |
| 3 | rs4566568 | 17798924 | A | G | 4.435578236675783e-12 | 6.922558723937961 | 31715.0 |
| 3 | rs73151380 | 90000532 | A | T | 4.6644706576945485e-12 | 6.915429774991282 | 31156.0 |
| 3 | rs6809775 | 17802881 | T | C | 4.7202090346207644e-12 | 6.913745735241778 | 31712.0 |
| 3 | rs7647914 | 17742786 | A | G | 4.733587530132952e-12 | 6.913344428287775 | 31849.0 |
| 3 | rs9868785 | 90066423 | G | A | 4.778048524979328e-12 | 6.912018699552486 | 31909.0 |
| 3 | rs4444738 | 17742738 | A | G | 4.8959920673787646e-12 | 6.908559624813582 | 31843.0 |
| 3 | rs13061771 | 17743799 | C | T | 4.8959920673787646e-12 | 6.908559624813582 | 31843.0 |
| 3 | rs11128848 | 17744210 | A | G | 4.8959920673787646e-12 | 6.908559624813582 | 31843.0 |
| 3 | rs4908973 | 17800423 | T | C | 4.949623205302797e-12 | 6.90701365173695 | 31714.0 |
| 3 | rs7609916 | 17804464 | A | G | 4.961002241989991e-12 | 6.9066877514743444 | 31716.0 |
| 3 | rs34485427 | 17694291 | C | A | 5.074732058971656e-12 | 6.903470159008832 | 32013.0 |
| 3 | rs1464463 | 90056380 | G | A | 5.0889816608216386e-12 | 6.903072008537888 | 31940.0 |
| 3 | rs13317478 | 17464862 | T | C | 5.242404381643454e-12 | 6.8988531071274055 | 32037.0 |
| 3 | rs4908940 | 17797271 | A | C | 5.2715815546688474e-12 | 6.898064492419145 | 31717.0 |
| 3 | rs12487248 | 17697696 | C | T | 5.2727117825146296e-12 | 6.8980340302193035 | 32005.0 |
| 3 | rs62277724 | 90281205 | G | A | 5.49922756875775e-12 | 6.892054551300879 | 32050.0 |
| 3 | rs4908992 | 17693974 | A | G | 5.631568925574133e-12 | 6.888671814473199 | 32014.0 |
| 3 | rs13096559 | 17694643 | G | T | 5.631568925574133e-12 | 6.888671814473199 | 32014.0 |
| 3 | rs6796448 | 17801811 | C | T | 5.68439905234404e-12 | 6.88734315305969 | 31713.0 |
| 3 | rs7647928 | 17742839 | A | G | 5.9753278771336245e-12 | 6.880236492606926 | 31847.0 |
| 3 | rs4396913 | 90047036 | T | C | 6.0232202310429865e-12 | 6.879099213694418 | 31937.0 |
| 3 | rs283917 | 17328509 | A | G | 6.052223819418952e-12 | 6.8784147791061425 | 32045.0 |
| 3 | rs73143703 | 90281792 | C | T | 6.159462747593839e-12 | 6.875911742275027 | 32090.0 |
| 3 | rs55701200 | 17690427 | C | T | 6.251961412698973e-12 | 6.873786826666192 | 32015.0 |
| 3 | rs7427588 | 17806424 | A | G | 6.3273081710072575e-12 | 6.872078580023108 | 31708.0 |
| 3 | rs6768666 | 17692937 | T | C | 6.647816578809861e-12 | 6.865028100723753 | 31997.0 |
| 3 | rs11426295 | 17757500 | T | TG | 6.961950781984316e-12 | 6.8584339476238885 | 31787.0 |
| 3 | rs6551450 | 90129328 | A | C | 7.404977784896984e-12 | 6.849613500826952 | 32019.0 |

| | | | | | | | |
|---|---|---|---|---|---|---|---|
| 3 | rs11128837 | 17697175 | T | C | 7.574146631497768e-12 | 6.846381240893978 | 32009.0 |
| 3 | rs6792422 | 17701005 | G | T | 7.68104337592372e-12 | 6.844375071837683 | 32002.0 |
| 3 | rs754076021 | 17698320 | AAT | A | 8.390732971745232e-12 | 6.831711734561948 | 31886.0 |
| 3 | rs283926 | 17357820 | G | A | 8.560881359320345e-12 | 6.8288319026073205 | 32040.0 |
| 3 | rs9824116 | 90319290 | A | G | 8.687141991052624e-12 | 6.826730910655702 | 31653.0 |
| 3 | rs6768637 | 17692849 | T | C | 8.873230839460203e-12 | 6.823688287785388 | 32014.0 |
| 3 | rs9864344 | 90121955 | T | G | 8.966119244884789e-12 | 6.822192842814637 | 32009.0 |
| 3 | rs9847461 | 17607956 | C | G | 8.968948986827473e-12 | 6.822147524373449 | 31755.0 |
| 3 | rs9851877 | 90258929 | G | A | 8.980205935606082e-12 | 6.821967382448863 | 32049.0 |
| 3 | rs13071728 | 17690673 | A | C | 9.011572204724779e-12 | 6.8214666011474785 | 32017.0 |
| 3 | rs9881089 | 17624602 | C | A | 9.129622352589657e-12 | 6.819597046393375 | 31980.0 |
| 3 | rs5014632 | 17695082 | A | G | 9.409487295378278e-12 | 6.8152578028134805 | 32017.0 |
| 3 | rs57107595 | 17706556 | G | GA | 9.893030981062307e-12 | 6.808050253900029 | 31277.0 |
| 3 | rs9857910 | 17697786 | A | T | 9.916208953112847e-12 | 6.807713498964951 | 32007.0 |
| 3 | rs137965771 | 90386418 | C | A | 9.99180997743898e-12 | 6.806620419600885 | 31241.0 |
| 3 | rs11397577 | 90081678 | A | AT | 1.0051734312568243e-11 | 6.805759743267554 | 31643.0 |
| 3 | rs4470554 | 17671037 | T | C | 1.0165761288485024e-11 | 6.8041357998555485 | 32020.0 |
| 3 | rs9880971 | 17624579 | C | A | 1.0415935494109378e-11 | 6.800634527033478 | 31980.0 |
| 3 | rs283939 | 17343995 | T | C | 1.103371952287868e-11 | 6.7923294824508265 | 32059.0 |
| 3 | rs17273020 | 17415705 | C | T | 1.1072421490980412e-11 | 6.791824476437116 | 32133.0 |
| 3 | rs9843225 | 17527551 | T | C | 1.1480787335145779e-11 | 6.786598823299585 | 31832.0 |
| 3 | rs7372028 | 90294773 | G | T | 1.153899133211317e-11 | 6.785868882454113 | 31784.0 |
| 3 | rs146780908 | 17704760 | T | TTTAAAATGG | 1.1695201280349536e-11 | 6.783927527970923 | 31638.0 |
| 3 | rs9821955 | 17650745 | A | G | 1.1888215159119055e-11 | 6.7815635626716055 | 31993.0 |
| 3 | rs6779834 | 17668766 | G | A | 1.1985521488861453e-11 | 6.780386006566283 | 32007.0 |
| 3 | rs113450730 | 90135516 | T | G | 1.2460849968489966e-11 | 6.774765158393402 | 32031.0 |
| 3 | rs2733481 | 17320397 | G | A | 1.2519235942139481e-11 | 6.774089267018537 | 32186.0 |
| 3 | rs11435463 | 90122867 | A | AT | 1.2691133957532184e-11 | 6.772117110282802 | 31789.0 |
| 3 | rs4908987 | 17688490 | T | C | 1.2732371032448164e-11 | 6.771647896190382 | 31998.0 |
| 3 | rs9681617 | 17655388 | T | C | 1.2790707050047855e-11 | 6.770986658587939 | 31990.0 |
| 3 | rs9877921 | 90108491 | C | T | 1.2928665012122016e-11 | 6.769434586777487 | 32045.0 |
| 3 | 3:90306556_GC_G | 90306556 | GC | G | 1.3253358594725208e-11 | 6.765844768794855 | 32020.0 |
| 3 | 3:89312113_CTTTTTTTTT_C | 89312113 | CTTTTTTTTT | C | 1.3331935447788465e-11 | 6.7649889710855815 | 30681.0 |
| 3 | rs12714734 | 90107488 | A | C | 1.3574377814650967e-11 | 6.762379276954781 | 32019.0 |
| 3 | rs9853571 | 17640071 | T | C | 1.379527753197024e-11 | 6.760040916864492 | 31990.0 |
| 3 | rs924762 | 147223899 | C | T | 1.3976457574959524e-11 | 6.7581502278723695 | 31404.0 |
| 3 | 3:17663199_CT_C | 17663199 | CT | C | 1.7034474248273265e-11 | 6.729416554789053 | 30598.0 |
| 3 | rs7429932 | 90283951 | A | G | 1.7219558707937205e-11 | 6.7278438293478136 | 32047.0 |
| 3 | rs71105145 | 90176889 | T | TA | 1.7441798633384762e-11 | 6.725977108037249 | 31684.0 |
| 3 | rs9816940 | 90146579 | C | T | 1.8316989979939764e-11 | 6.718845574747562 | 32118.0 |
| 3 | rs75980247 | 17740272 | G | A | 1.9228896373012745e-11 | 6.711761383019455 | 31398.0 |
| 3 | rs141333755 | 90185094 | C | CA | 1.9241220694527848e-11 | 6.711667911487239 | 32066.0 |
| 3 | rs5846968 | 17796108 | G | T | 1.9628853249251715e-11 | 6.70875751335181 | 29246.0 |
| 3 | rs9875185 | 90394152 | A | G | 1.964250464264342e-11 | 6.7086560456620905 | 31262.0 |
| 3 | rs60759458 | 90045885 | G | C | 1.985507106528537e-11 | 6.707084933923726 | 31846.0 |
| 3 | rs283923 | 17325703 | A | G | 2.0031544315681183e-11 | 6.705793053980978 | 32136.0 |
| 3 | rs9857243 | 90395814 | A | G | 2.051737918191796e-11 | 6.702293226047133 | 31222.0 |
| 3 | rs9837548 | 90156142 | C | T | 2.0642054178261244e-11 | 6.701408183394675 | 32138.0 |
| 3 | rs283907 | 17336419 | T | C | 2.1194869900242616e-11 | 6.697545879302278 | 32119.0 |
| 3 | rs283909 | 17333068 | A | G | 2.175380812159352e-11 | 6.693739801336408 | 32112.0 |
| 3 | rs73143764 | 90405465 | G | C | 2.3111244716020672e-11 | 6.684880949360141 | 30977.0 |
| 3 | rs283919 | 17328070 | G | C | 2.322520095726821e-11 | 6.684160591538907 | 32088.0 |
| 3 | rs283921 | 17325889 | G | A | 2.375832435869218e-11 | 6.680835842284638 | 32129.0 |
| 3 | rs7433265 | 90275692 | A | G | 2.375850949566848e-11 | 6.6808347004387425 | 32038.0 |

| | | | | | | | |
|---|---|---|---|---|---|---|---|
| 3 | rs6763681 | 17409517 | G | A | 2.3905586058517858e-11 | 6.6799303358274535 | 32034.0 |
| 3 | rs2470349 | 17304712 | G | A | 2.4290084774249862e-11 | 6.677591566949777 | 31544.0 |
| 3 | rs76900983 | 90363402 | C | A | 2.4611963086865608e-11 | 6.675661397826332 | 31268.0 |
| 3 | rs2596678 | 17320600 | T | C | 2.4695300093957433e-11 | 6.675165687980456 | 32134.0 |
| 3 | rs2596681 | 17320850 | T | A | 2.4849840762344805e-11 | 6.674250759206821 | 32059.0 |
| 3 | rs283915 | 17329768 | T | C | 2.6021794248527122e-11 | 6.6674884444057385 | 32107.0 |
| 3 | rs7433880 | 90174960 | C | A | 2.605949271846171e-11 | 6.667275902862492 | 32173.0 |
| 3 | rs283916 | 17329413 | A | G | 2.610049512862385e-11 | 6.667045075343088 | 32106.0 |
| 3 | rs2470347 | 17313703 | T | A | 2.6388068218399025e-11 | 6.665436057033777 | 32077.0 |
| 3 | rs283908 | 17334741 | T | C | 2.6551819833526336e-11 | 6.664527492617422 | 32115.0 |
| 3 | rs9815264 | 90276511 | G | A | 2.68407497294016e-11 | 6.66293768066142 | 32056.0 |
| 3 | rs9815542 | 90276661 | G | A | 2.68407497294016e-11 | 6.66293768066142 | 32056.0 |
| 3 | rs283922 | 17325841 | T | C | 2.7065534177050327e-11 | 6.661712367054516 | 32129.0 |
| 3 | rs1840943 | 90460310 | T | A | 2.764769483309361e-11 | 6.658584697266695 | 31451.0 |
| 3 | rs73153370 | 90056125 | T | A | 2.791038350528269e-11 | 6.657194447939745 | 31839.0 |
| 3 | rs10049142 | 17387061 | A | G | 2.8134816986677296e-11 | 6.656016768164017 | 32083.0 |
| 3 | rs2596679 | 17320788 | T | C | 2.822618555432561e-11 | 6.655539957522713 | 32139.0 |
| 3 | 3:17807171_CAAAAA_C | 17807171 | CAAAAA | C | 2.867227950450393e-11 | 6.653233479894247 | 31500.0 |
| 3 | rs10049143 | 17387080 | A | C | 2.8972933195137268e-11 | 6.651698714317143 | 32082.0 |
| 3 | rs10433560 | 17803974 | C | T | 2.909320473836321e-11 | 6.6510891177300175 | 31248.0 |
| 3 | rs9857806 | 90189355 | G | A | 2.94984436179467e-11 | 6.6490531676246025 | 32168.0 |
| 3 | rs2733523 | 17322432 | G | A | 2.9840780533051234e-11 | 6.647354460573151 | 32142.0 |
| 3 | rs149460172 | 90408448 | T | G | 3.009595324679452e-11 | 6.646100632197946 | 30985.0 |
| 3 | rs6577618 | 17715648 | A | T | 3.0157270777768555e-11 | 6.645800890258384 | 31747.0 |
| 3 | rs17043492 | 17402565 | T | C | 3.0273455487688325e-11 | 6.645234570341692 | 32125.0 |
| 3 | rs2470577 | 17313311 | A | G | 3.048157134471134e-11 | 6.644225447197192 | 32078.0 |
| 3 | rs7432427 | 90299370 | A | T | 3.064798511710054e-11 | 6.643423371457423 | 31947.0 |
| 3 | rs4312680 | 17788430 | G | A | 3.096426462281933e-11 | 6.641910658054526 | 31726.0 |
| 3 | rs17043494 | 17402806 | T | C | 3.127235029439718e-11 | 6.640451604868798 | 32122.0 |
| 3 | rs6785870 | 17387479 | T | C | 3.174126815864995e-11 | 6.638257653565363 | 32100.0 |
| 3 | rs2733479 | 17320931 | G | T | 3.177578420590788e-11 | 6.638097417150288 | 32135.0 |
| 3 | 3:17392222_AG_A | 17392222 | AG | A | 3.1908026379740784e-11 | 6.6374850720345 | 32120.0 |
| 3 | rs2733487 | 17315969 | A | C | 3.1911741920947105e-11 | 6.637467903180543 | 32132.0 |
| 3 | rs79430269 | 90403309 | G | A | 3.216789963608309e-11 | 6.636288936156372 | 31514.0 |
| 3 | 3:90392328_CTAGA_C | 90392328 | CTAGA | C | 3.235795806331464e-11 | 6.635420114098173 | 31487.0 |
| 3 | rs12638179 | 90456907 | C | G | 3.2859504406008797e-11 | 6.633151135222247 | 30949.0 |
| 3 | rs2733524 | 17322397 | G | T | 3.295500207709091e-11 | 6.6327229508169925 | 32123.0 |
| 3 | rs283906 | 17337410 | C | A | 3.3399281859333326e-11 | 6.63074675964393 | 32112.0 |
| 3 | rs9857738 | 90431024 | A | G | 3.4547255327156096e-11 | 6.625757231106981 | 31036.0 |
| 3 | rs9856356 | 90406018 | A | G | 3.457995381332456e-11 | 6.625617499876428 | 30958.0 |
| 3 | rs283913 | 17330957 | A | C | 3.50936957897676e-11 | 6.623438927192893 | 32119.0 |
| 3 | rs2733485 | 17318438 | A | C | 3.5339957281176577e-11 | 6.622405674092777 | 32134.0 |
| 3 | rs283925 | 17325403 | C | G | 3.54324009761069e-11 | 6.622019620894389 | 32140.0 |
| 3 | rs2733514 | 17323931 | T | C | 3.5716148917956886e-11 | 6.620840790147581 | 32138.0 |
| 3 | rs2596674 | 17315986 | C | T | 3.5999348741899645e-11 | 6.6196733396057175 | 32137.0 |
| 3 | rs283912 | 17331582 | A | T | 3.656929522077085e-11 | 6.61735081650173 | 32121.0 |
| 3 | rs7426793 | 90251951 | C | A | 3.721093387522339e-11 | 6.6147781615916985 | 32043.0 |
| 3 | rs2596675 | 17316259 | C | G | 3.746343295219191e-11 | 6.613777649308427 | 32133.0 |
| 3 | rs283947 | 17354812 | G | A | 3.8045340154371094e-11 | 6.611496778225224 | 32170.0 |
| 3 | rs9820410 | 90431093 | G | A | 3.838508051244863e-11 | 6.6101808480995174 | 31045.0 |
| 3 | rs62277418 | 90431424 | C | A | 3.838508051244863e-11 | 6.6101808480995174 | 31045.0 |
| 3 | rs75342952 | 17386077 | C | T | 3.9025498464974724e-11 | 6.6077309908659885 | 32114.0 |
| 3 | rs283949 | 17355801 | G | C | 3.9229054402710186e-11 | 6.606960541187829 | 32139.0 |
| 3 | rs4908970 | 17788866 | A | G | 3.931509127736396e-11 | 6.606636071137466 | 31731.0 |

| | | | | | | | |
|---|---|---|---|---|---|---|---|
| 3 | rs28808578 | 90335353 | G | A | 3.945564206458915e-11 | 6.606107504958096 | 31808.0 |
| 3 | rs283935 | 17367357 | A | T | 3.966785890452737e-11 | 6.605312906710787 | 32128.0 |
| 3 | rs71266497 | 17333904 | C | CTTGA | 3.9677685395959016e-11 | 6.605276214460268 | 32035.0 |
| 3 | rs4908938 | 17787557 | G | A | 3.985713286256979e-11 | 6.604607714458687 | 31745.0 |
| 3 | rs4488868 | 17771429 | T | C | 4.04672520493996e-11 | 6.60235664813253 | 31846.0 |
| 3 | rs9682382 | 90342103 | C | A | 4.060840955115057e-11 | 6.601840571961873 | 31779.0 |
| 3 | rs4908971 | 17789074 | T | C | 4.1498024314096404e-11 | 6.598627952584303 | 31741.0 |
| 3 | rs35500376 | 17601086 | A | AT | 4.154041946158659e-11 | 6.598476540484098 | 30588.0 |
| 3 | rs2060625 | 17339152 | G | T | 4.1887601881058506e-11 | 6.597242255058798 | 32123.0 |
| 3 | rs283911 | 17331704 | C | T | 4.2026169927163466e-11 | 6.596752417643219 | 32123.0 |
| 3 | rs9714360 | 90357513 | G | T | 4.2157690476030006e-11 | 6.59628895266878 | 31557.0 |
| 3 | rs144184070 | 90358160 | G | A | 4.2270979048824655e-11 | 6.595890868049959 | 31568.0 |
| 3 | rs9713426 | 90338187 | C | T | 4.3053807824005735e-11 | 6.593168284471479 | 31805.0 |
| 3 | rs283938 | 17343457 | T | C | 4.325566879071583e-11 | 6.592474091867447 | 32149.0 |
| 3 | rs2060627 | 17339381 | C | G | 4.340977450614585e-11 | 6.5919462581266695 | 32122.0 |
| 3 | rs5023579 | 17788324 | A | T | 4.436607595809167e-11 | 6.588711217906033 | 29805.0 |
| 3 | rs13318609 | 17395320 | T | C | 4.4848352145487115e-11 | 6.587105532012856 | 32138.0 |
| 3 | rs6551451 | 90156286 | C | T | 4.505503036459839e-11 | 6.586422584398056 | 32133.0 |
| 3 | rs12488093 | 17388506 | T | C | 4.5184375657491726e-11 | 6.585996733385879 | 32141.0 |
| 3 | rs11374153 | 17779317 | C | CA | 4.543082592639431e-11 | 6.585188621885175 | 29799.0 |
| 3 | rs9810980 | 90349804 | T | C | 4.5446273026969496e-11 | 6.585138113714327 | 31513.0 |
| 3 | rs2733478 | 17321551 | A | G | 4.572838801086521e-11 | 6.584218610699789 | 32140.0 |
| 3 | rs283948 | 17355617 | C | G | 4.6465835120943494e-11 | 6.581841001725683 | 32138.0 |
| 3 | 3:90308526_TA_T | 90308526 | TA | T | 4.6534794366754424e-11 | 6.581620560489414 | 30577.0 |
| 3 | rs2596640 | 17288440 | A | G | 4.70483132025095e-11 | 6.579988981954751 | 32055.0 |
| 3 | rs2060624 | 17338719 | C | T | 4.759166941306228e-11 | 6.578281464810266 | 32123.0 |
| 3 | rs7429720 | 90228124 | T | G | 4.806174431102522e-11 | 6.576819554712597 | 32107.0 |
| 3 | rs283931 | 17361581 | T | C | 4.8327028800041327e-11 | 6.576000695476942 | 32146.0 |
| 3 | rs2470580 | 17367746 | T | C | 4.8327028800041327e-11 | 6.576000695476942 | 32146.0 |
| 3 | rs17043422 | 17383881 | C | T | 4.83594974441569e-11 | 6.575900776026603 | 32146.0 |
| 3 | rs761041249 | 17714993 | AGT | A | 4.851317501382226e-11 | 6.575428735276426 | 31791.0 |
| 3 | rs34502358 | 17377815 | T | C | 4.8738387278214067e-11 | 6.574739602508626 | 32125.0 |
| 3 | rs28408438 | 17384023 | T | C | 4.8776022803848926e-11 | 6.574624744487263 | 32145.0 |
| 3 | rs28454413 | 17383964 | C | T | 4.904708914257682e-11 | 6.573800042412517 | 32145.0 |
| 3 | rs767685 | 17324733 | A | G | 4.925991707191575e-11 | 6.573155645883522 | 32144.0 |
| 3 | rs4582042 | 17421879 | C | T | 4.9661859156779535e-11 | 6.571946047980727 | 32171.0 |
| 3 | rs2596677 | 17318844 | T | C | 5.087620877587458e-11 | 6.568348938985747 | 32141.0 |
| 3 | rs13095571 | 17532320 | T | C | 5.120063430222194e-11 | 6.5674021516154735 | 31885.0 |
| 3 | rs9877959 | 90263752 | C | A | 5.143973683473012e-11 | 6.5667081157068266 | 32014.0 |
| 3 | rs2596638 | 17287018 | C | T | 5.2303853511152606e-11 | 6.564225914395305 | 32056.0 |
| 3 | rs2596639 | 17287135 | G | A | 5.2303853511152606e-11 | 6.564225914395305 | 32056.0 |
| 3 | rs2733488 | 17315496 | T | C | 5.233774575827299e-11 | 6.564129377441825 | 32116.0 |
| 3 | rs12637691 | 90355883 | C | G | 5.294131827617789e-11 | 6.56242035795571 | 31644.0 |
| 3 | rs6802891 | 17297337 | T | C | 5.337246668791118e-11 | 6.561211189167997 | 31976.0 |
| 3 | rs147324072 | 17718175 | C | T | 5.4454217061053323e-11 | 6.5582189524137675 | 31769.0 |
| 3 | rs2084572 | 17315758 | A | G | 5.478593708087261e-11 | 6.557313021771454 | 32135.0 |
| 3 | rs2596636 | 17283809 | T | A | 5.484212164033272e-11 | 6.5571601127443095 | 32059.0 |
| 3 | rs9798932 | 90461784 | G | A | 5.493645340118397e-11 | 6.556903728533144 | 31007.0 |
| 3 | rs2167118 | 17283278 | G | A | 5.53307308631461e-11 | 6.555836763153791 | 32060.0 |
| 3 | rs36172181 | 17719478 | G | A | 5.545280485236896e-11 | 6.5555079227327955 | 31901.0 |
| 3 | rs187303 | 17357075 | G | A | 5.554132480207315e-11 | 6.555269912113168 | 32139.0 |
| 3 | rs2470583 | 17280812 | G | A | 5.666161959575846e-11 | 6.5522893391934725 | 32058.0 |
| 3 | rs56399694 | 90361837 | G | A | 5.669436518259791e-11 | 6.552203088428433 | 31769.0 |
| 3 | rs13322641 | 17378509 | G | A | 5.7152567166866246e-11 | 6.551001284838223 | 32138.0 |

| | | | | | | | |
|---|---|---|---|---|---|---|---|
| 3 | rs2596680 | 17320823 | C | G | 5.8082193903473704e-11 | 6.548591686069328 | 32103.0 |
| 3 | rs9849053 | 90152502 | G | A | 5.884024527573764e-11 | 6.5466545722031615 | 32103.0 |
| 3 | rs10470576 | 90154964 | C | T | 5.884024527573764e-11 | 6.5466545722031615 | 32103.0 |
| 3 | rs2596673 | 17270823 | A | G | 5.987155673754076e-11 | 6.544058006998358 | 31975.0 |
| 3 | rs4417835 | 147256611 | A | G | 5.997995691426401e-11 | 6.543787629794822 | 31617.0 |
| 3 | rs4908965 | 17768038 | T | C | 6.082980817290828e-11 | 6.541684301833628 | 31848.0 |
| 3 | rs2592099 | 17369313 | G | T | 6.101875790302335e-11 | 6.541220570144469 | 32145.0 |
| 3 | rs4908967 | 17774944 | C | A | 6.107466262370033e-11 | 6.5410836346842265 | 31837.0 |
| 3 | rs2733497 | 17280282 | C | T | 6.120828654130726e-11 | 6.540756826524771 | 32058.0 |
| 3 | rs2733501 | 17274801 | T | C | 6.146268317453347e-11 | 6.540136565167125 | 32049.0 |
| 3 | rs11927654 | 17723430 | G | A | 6.156634426825017e-11 | 6.539884541931246 | 31894.0 |
| 3 | rs62275565 | 90351548 | C | G | 6.230160927966493e-11 | 6.5381087714862485 | 31809.0 |
| 3 | rs1375817 | 17266023 | T | C | 6.259996445082675e-11 | 6.537394040679718 | 32050.0 |
| 3 | rs186005608 | 90483873 | C | A | 6.273977526423265e-11 | 6.537060259741365 | 31385.0 |
| 3 | rs9838264 | 90351314 | A | T | 6.303535103075735e-11 | 6.536356997602062 | 31797.0 |
| 3 | 3:17397140_AT_A | 17397140 | AT | A | 6.334691587338075e-11 | 6.535619175911579 | 32087.0 |
| 3 | rs283937 | 17341329 | C | T | 6.342638568806846e-11 | 6.5354315502904425 | 32107.0 |
| 3 | rs7630286 | 17371566 | C | T | 6.345615120523385e-11 | 6.5353613340649455 | 32076.0 |
| 3 | rs7630433 | 17371582 | G | C | 6.345615120523385e-11 | 6.5353613340649455 | 32076.0 |
| 3 | rs12490933 | 17754925 | A | G | 6.35401544222618e-11 | 6.535163345857673 | 31887.0 |
| 3 | rs2596671 | 17266560 | T | C | 6.385663563347623e-11 | 6.534419719561633 | 32051.0 |
| 3 | rs2596647 | 17300133 | A | C | 6.397313349259684e-11 | 6.534146895410168 | 32043.0 |
| 3 | rs11128852 | 17776334 | T | C | 6.397411748835413e-11 | 6.5341445930790805 | 31847.0 |
| 3 | rs11128840 | 17704787 | C | G | 6.42823485775731e-11 | 6.5334251002074515 | 31935.0 |
| 3 | rs6806610 | 17705696 | G | A | 6.42823485775731e-11 | 6.5334251002074515 | 31935.0 |
| 3 | rs6780587 | 17709796 | C | T | 6.42823485775731e-11 | 6.5334251002074515 | 31935.0 |
| 3 | rs145367119 | 90049321 | C | T | 6.444667617972763e-11 | 6.533042893912648 | 29845.0 |
| 3 | 3:17343781_GTA_G | 17343781 | GTA | G | 6.445262348399996e-11 | 6.533029079075513 | 32154.0 |
| 3 | 3:17648762_CA_C | 17648762 | CA | C | 6.506009982099186e-11 | 6.531624515366473 | 31927.0 |
| 3 | rs747681042 | 17375416 | AGGGAGAATG | A | 6.517425288645024e-11 | 6.531362010991927 | 32109.0 |
| 3 | rs1545424 | 17266007 | T | A | 6.540818060143423e-11 | 6.530825476999426 | 32051.0 |
| 3 | rs34422952 | 17800933 | A | ATTAAA | 6.543131703920895e-11 | 6.530772513550287 | 29439.0 |
| 3 | rs374899793 | 17741045 | T | TAAAG | 6.628847728675223e-11 | 6.528823120055892 | 25168.0 |
| 3 | rs34002492 | 17741698 | C | G | 6.692690333233655e-11 | 6.527387135401845 | 31893.0 |
| 3 | rs2596665 | 17259109 | A | G | 6.717322939085549e-11 | 6.526836663472451 | 32067.0 |
| 3 | rs283927 | 17358421 | G | A | 6.773880521008964e-11 | 6.525580188609568 | 32145.0 |
| 3 | rs11708672 | 17703137 | C | T | 6.795517262722006e-11 | 6.525102221458359 | 31935.0 |
| 3 | rs283928 | 17359316 | C | T | 6.809212171874403e-11 | 6.524800462174404 | 32145.0 |
| 3 | rs2348003 | 17292100 | T | C | 6.836105891443882e-11 | 6.524209598255785 | 32037.0 |
| 3 | rs2252237 | 17263417 | C | T | 6.900061119883254e-11 | 6.522813562096026 | 32057.0 |
| 3 | 3:17273959_GA_G | 17273959 | GA | G | 6.967817147894517e-11 | 6.5213482989092775 | 31991.0 |
| 3 | rs13069155 | 17767422 | A | G | 6.970333243392485e-11 | 6.5212941556126784 | 31852.0 |
| 3 | rs2348002 | 17251689 | T | C | 6.97172551096506e-11 | 6.5212642039318975 | 32011.0 |
| 3 | rs13072194 | 90189970 | C | T | 6.984375567166853e-11 | 6.520992332552599 | 32169.0 |
| 3 | rs13353435 | 17430574 | T | C | 7.008882213371768e-11 | 6.520467010131734 | 32166.0 |
| 3 | rs4435673 | 17749125 | A | G | 7.010174383523746e-11 | 6.520439361159028 | 31897.0 |
| 3 | rs7636787 | 17433760 | G | C | 7.030233092570759e-11 | 6.520010796656934 | 32161.0 |
| 3 | rs2733500 | 17276149 | G | A | 7.03363573274781e-11 | 6.519938216175518 | 32058.0 |
| 3 | rs10433561 | 17803996 | T | G | 7.163266541177574e-11 | 6.517198377211733 | 31719.0 |
| 3 | rs2596667 | 17263274 | C | T | 7.170806126390438e-11 | 6.517040518455827 | 32056.0 |
| 3 | rs1449881 | 17286817 | T | G | 7.175543002799003e-11 | 6.51694142393168 | 32053.0 |
| 3 | rs7645459 | 17808236 | C | G | 7.200158211431801e-11 | 6.516427506378317 | 31716.0 |
| 3 | rs6551407 | 89270605 | C | T | 7.256822258213377e-11 | 6.5152509717672356 | 31912.0 |
| 3 | rs62275605 | 90354891 | C | T | 7.301416131376979e-11 | 6.514331354495455 | 31794.0 |

| | | | | | | | |
|---|---|---|---|---|---|---|---|
| 3 | rs773279507 | 17281259 | TG | T | 7.311848968747522e-11 | 6.5141170008672855 | 31978.0 |
| 3 | rs7651998 | 17711127 | G | A | 7.440650764623108e-11 | 6.511494996084712 | 31930.0 |
| 3 | rs8179891 | 17727378 | T | G | 7.464157496402109e-11 | 6.511021265731821 | 31897.0 |
| 3 | rs12634893 | 17729692 | T | C | 7.464157496402109e-11 | 6.511021265731821 | 31897.0 |
| 3 | rs10510478 | 17730282 | A | G | 7.464157496402109e-11 | 6.511021265731821 | 31897.0 |
| 3 | rs4408889 | 17730418 | T | C | 7.464157496402109e-11 | 6.511021265731821 | 31897.0 |
| 3 | rs56059241 | 17732173 | C | A | 7.464157496402109e-11 | 6.511021265731821 | 31897.0 |
| 3 | rs4243837 | 17733234 | G | A | 7.464157496402109e-11 | 6.511021265731821 | 31897.0 |
| 3 | rs6789467 | 17802134 | G | C | 7.468111055804245e-11 | 6.510941733137558 | 31719.0 |
| 3 | rs4611850 | 17798649 | T | G | 7.489748884749038e-11 | 6.510507179285071 | 31721.0 |
| 3 | rs9830414 | 90358707 | T | C | 7.515138454255336e-11 | 6.509998841715555 | 31683.0 |
| 3 | rs11128842 | 17735345 | G | C | 7.51738787137695e-11 | 6.509953886002097 | 31895.0 |
| 3 | rs35460334 | 17737277 | C | T | 7.51738787137695e-11 | 6.509953886002097 | 31895.0 |
| 3 | rs2198957 | 90355214 | G | A | 7.529049329414704e-11 | 6.50972103666537 | 31686.0 |
| 3 | rs752814983 | 17336486 | AAT | A | 7.539860739704535e-11 | 6.509505475438412 | 32037.0 |
| 3 | rs2596669 | 17265041 | C | T | 7.602659528592089e-11 | 6.50825931961228 | 32057.0 |
| 3 | rs4643736 | 17792256 | G | T | 7.613633591281728e-11 | 6.508042588060771 | 31722.0 |
| 3 | rs2733491 | 17308965 | G | A | 7.616379547054446e-11 | 6.507988404765503 | 32053.0 |
| 3 | rs11128849 | 17764670 | T | C | 7.623511906189814e-11 | 6.507847758004508 | 31891.0 |
| 3 | rs11128850 | 17764748 | A | G | 7.623511906189814e-11 | 6.507847758004508 | 31891.0 |
| 3 | rs1449879 | 17277391 | A | G | 7.62503276843369e-11 | 6.507817783962687 | 32056.0 |
| 3 | rs9682018 | 90364375 | G | A | 7.62983129427688e-11 | 6.507723250119509 | 31672.0 |
| 3 | rs4349529 | 17792118 | A | G | 7.636624539659218e-11 | 6.5075895184697545 | 31723.0 |
| 3 | rs35811992 | 17793235 | G | A | 7.636624539659218e-11 | 6.5075895184697545 | 31723.0 |
| 3 | rs728022 | 17305303 | G | C | 7.683342495234359e-11 | 6.506672968808999 | 32043.0 |
| 3 | rs4615062 | 17434502 | C | A | 7.740844927700991e-11 | 6.505552293785633 | 32163.0 |
| 3 | rs13076724 | 17758473 | A | G | 7.771245117376833e-11 | 6.504963104367636 | 31885.0 |
| 3 | rs2733517 | 17323763 | G | A | 7.870091630794183e-11 | 6.5030628083722055 | 32120.0 |
| 3 | rs283918 | 17328219 | C | G | 7.900447061208541e-11 | 6.502483916239567 | 31987.0 |
| 3 | rs2596652 | 17368197 | C | A | 7.912997737849163e-11 | 6.502245204305607 | 32149.0 |
| 3 | rs11412361 | 17399660 | C | CT | 7.915182470508473e-11 | 6.502203688863753 | 31711.0 |
| 3 | rs11713917 | 17382262 | G | A | 8.03121277846429e-11 | 6.5000147670045125 | 32139.0 |
| 3 | rs1449886 | 17261930 | G | A | 8.244600275284007e-11 | 6.4960687306719125 | 32058.0 |
| 3 | rs283944 | 17346655 | C | A | 8.24940376997804e-11 | 6.495981056792062 | 32186.0 |
| 3 | rs7627064 | 17737834 | T | C | 8.330082307442581e-11 | 6.494515916963308 | 31892.0 |
| 3 | rs1597394 | 17260188 | C | T | 8.331295197656576e-11 | 6.494493996643692 | 32053.0 |
| 3 | rs776423900 | 17313583 | TGGAAA | T | 8.427472186455086e-11 | 6.492765668527159 | 32058.0 |
| 3 | rs283941 | 17344551 | C | A | 8.44660559354879e-11 | 6.4924241368056474 | 32174.0 |
| 3 | rs283946 | 17347636 | T | C | 8.44660559354879e-11 | 6.4924241368056474 | 32174.0 |
| 3 | rs283943 | 17346355 | C | T | 8.480711932367973e-11 | 6.491817209726034 | 32175.0 |
| 3 | rs9856430 | 90348280 | C | T | 8.518411878966293e-11 | 6.4911491047493755 | 31719.0 |
| 3 | rs2596637 | 17285288 | T | G | 8.674856680994849e-11 | 6.488407191614907 | 32048.0 |
| 3 | rs12629181 | 90483648 | C | A | 8.825007479941249e-11 | 6.485820686411516 | 30867.0 |
| 3 | rs283924 | 17325616 | G | A | 8.88344293380907e-11 | 6.484825689003768 | 31890.0 |
| 3 | rs2596645 | 17296356 | T | C | 8.896919256950251e-11 | 6.4845971319511975 | 32039.0 |
| 3 | 3:17331124_CA_C | 17331124 | CA | C | 8.941164694451039e-11 | 6.483849106770532 | 30840.0 |
| 3 | rs2733502 | 17274422 | C | T | 8.994297850457061e-11 | 6.482955591527452 | 32047.0 |
| 3 | rs13079096 | 17795076 | T | A | 9.151859788214483e-11 | 6.480335974153067 | 31725.0 |
| 3 | rs2596646 | 17296401 | T | C | 9.192148444934949e-11 | 6.479673216366381 | 32043.0 |
| 3 | rs2596672 | 17267023 | G | A | 9.270342514599837e-11 | 6.478394971747619 | 32048.0 |
| 3 | rs2733498 | 17276455 | T | C | 9.301248372705238e-11 | 6.477892655668027 | 32057.0 |
| 3 | rs13093375 | 17435598 | A | G | 9.325832140089187e-11 | 6.4774942570894405 | 32163.0 |
| 3 | rs7609645 | 89248378 | G | A | 9.331898841143923e-11 | 6.477396099564541 | 31960.0 |
| 3 | rs4908968 | 17778320 | T | C | 9.515356648305372e-11 | 6.474456899914402 | 31847.0 |

| | | | | | | | |
|---|---|---|---|---|---|---|---|
| 3 | rs7642719 | 17435581 | G | A | 9.542979609188414e-11 | 6.474019154873448 | 32162.0 |
| 3 | rs2060628 | 17275173 | G | C | 9.600518555400523e-11 | 6.473111290866616 | 32055.0 |
| 3 | rs2733505 | 17271758 | C | T | 9.663644914134896e-11 | 6.472121366621279 | 31990.0 |
| 3 | rs11719326 | 17781780 | A | C | 9.690580717820529e-11 | 6.471700892247704 | 31841.0 |
| 3 | rs2733508 | 17270788 | T | C | 9.912309764921869e-11 | 6.468282440836528 | 31930.0 |
| 3 | rs2596670 | 17265224 | G | A | 9.939397017671778e-11 | 6.467869964664018 | 32050.0 |
| 3 | rs6442679 | 17397851 | C | T | 9.996693781066171e-11 | 6.467001076373732 | 32137.0 |
| 3 | rs6551402 | 89246713 | G | A | 1.0053470959041452e-10 | 6.4661448568664746 | 31964.0 |
| 3 | 3:17316598_AT_A | 17316598 | AT | A | 1.0056006879252172e-10 | 6.466106724686258 | 32096.0 |
| 3 | rs6795664 | 89246622 | A | G | 1.0152840628779274e-10 | 6.4646576419887305 | 31965.0 |
| 3 | rs7645756 | 17397336 | T | C | 1.0180186903077416e-10 | 6.464250860967742 | 32132.0 |
| 3 | rs2122368 | 17261455 | G | A | 1.0188202521695844e-10 | 6.464131829555779 | 32053.0 |
| 3 | rs2733509 | 17264355 | A | G | 1.0265171733544593e-10 | 6.462993479699921 | 32052.0 |
| 3 | rs144807058 | 89392008 | T | TAC | 1.0331026410528122e-10 | 6.462026114672063 | 25352.0 |
| 3 | rs9812199 | 17555377 | C | T | 1.0463108758760251e-10 | 6.460103943644234 | 32021.0 |
| 3 | rs62245860 | 17292933 | T | C | 1.0476464230125769e-10 | 6.459910906109365 | 31923.0 |
| 3 | rs113252352 | 90356125 | A | G | 1.0517756195819036e-10 | 6.45931559768286 | 31299.0 |
| 3 | rs2733520 | 17246392 | G | A | 1.0520487906995922e-10 | 6.459276295074582 | 32186.0 |
| 3 | rs2060629 | 17279526 | T | C | 1.0529732703920684e-10 | 6.459143359145933 | 32059.0 |
| 3 | rs1375818 | 17270900 | T | C | 1.0596936753989195e-10 | 6.458180411327232 | 32047.0 |
| 3 | rs2733507 | 17270795 | T | C | 1.0661876495361552e-10 | 6.457255563009628 | 31922.0 |
| 3 | rs2733499 | 17276334 | C | T | 1.067254769967556e-10 | 6.4571041145702095 | 32056.0 |
| 3 | rs9852861 | 17513896 | C | T | 1.0684453206375884e-10 | 6.456935323205072 | 32042.0 |
| 3 | rs1597393 | 17259757 | T | C | 1.0805715752286675e-10 | 6.455226510544772 | 32061.0 |
| 3 | rs2596664 | 17254290 | T | G | 1.0848452561403641e-10 | 6.454628734666112 | 32121.0 |
| 3 | rs2596666 | 17263042 | G | A | 1.0853505987399673e-10 | 6.454558202740582 | 32053.0 |
| 3 | rs34983821 | 17757940 | G | GA | 1.0855826369887157e-10 | 6.4545258273379575 | 31722.0 |
| 3 | rs35417636 | 17262695 | A | AT | 1.1012747534210573e-10 | 6.45235191708529 | 31971.0 |
| 3 | rs9682244 | 90484101 | A | C | 1.1064395731599012e-10 | 6.451643026819402 | 30788.0 |
| 3 | rs2596663 | 17254001 | A | T | 1.111916765180496e-10 | 6.450894787722068 | 32122.0 |
| 3 | rs2167119 | 17262139 | A | G | 1.1168324389627578e-10 | 6.450226319069508 | 32053.0 |
| 3 | rs7653828 | 90180985 | C | T | 1.1269603112300756e-10 | 6.4488580783122496 | 32186.0 |
| 3 | rs2733504 | 17271759 | A | G | 1.1314234559233491e-10 | 6.448258934231408 | 31982.0 |
| 3 | rs781571315 | 17347602 | CACTTCCAG G | C | 1.164373196339529e-10 | 6.4439058659766175 | 32168.0 |
| 3 | rs62245914 | 17385662 | T | C | 1.1872194360682964e-10 | 6.440957721437262 | 32049.0 |
| 3 | rs7644389 | 17502918 | T | C | 1.1986157999087842e-10 | 6.439507769905656 | 32043.0 |
| 3 | rs1463218 | 17301087 | G | C | 1.226519262977228e-10 | 6.4360137515635785 | 32027.0 |
| 3 | rs7613953 | 17743150 | G | T | 1.2295797699565838e-10 | 6.4356352597001 22 | 31842.0 |
| 3 | rs2733494 | 17297521 | C | T | 1.240790536061838e-10 | 6.434256648638472 | 32011.0 |
| 3 | rs9839588 | 17667278 | T | C | 1.27253730329169e-10 | 6.430417742637543 | 32035.0 |
| 3 | 3:89248669_CAT_C | 89248669 | CAT | C | 1.2945724595270534e-10 | 6.427807837699769 | 31549.0 |
| 3 | rs7627732 | 17784818 | T | C | 1.3108211667526236e-10 | 6.425910961172381 | 31752.0 |
| 3 | rs9861749 | 17409919 | G | A | 1.3135187580066631e-10 | 6.425598270018632 | 32161.0 |
| 3 | rs9881850 | 17554181 | C | T | 1.3257863577630085e-10 | 6.424184143278418 | 32021.0 |
| 3 | 3:147256440_TA_T | 147256440 | TA | T | 1.3342407088300354e-10 | 6.423217006531988 | 29073.0 |
| 3 | rs11128838 | 17697368 | G | A | 1.3497570394307724e-10 | 6.4214574951102925 | 32018.0 |
| 3 | rs35204863 | 17241552 | T | TC | 1.3523174461236616e-10 | 6.421169053188461 | 31972.0 |
| 3 | rs1449883 | 17253429 | T | A | 1.3615601098307074e-10 | 6.4201322458478085 | 32127.0 |
| 3 | rs35997387 | 17739538 | C | A | 1.406713355274898e-10 | 6.415164009113338 | 31675.0 |
| 3 | rs55740247 | 90362832 | T | A | 1.4145998331070252e-10 | 6.41431227294002 | 31143.0 |
| 3 | rs1449884 | 17256687 | G | C | 1.4191907055150115e-10 | 6.413818595147129 | 32118.0 |
| 3 | rs2470586 | 17297663 | C | T | 1.4494575445808801e-10 | 6.410602404683499 | 32029.0 |
| 3 | 3:17260971_GT_G | 17260971 | GT | G | 1.4620861937288644e-10 | 6.409279833048678 | 30820.0 |
| 3 | rs2247106 | 17308770 | A | G | 1.468312758378202e-10 | 6.408631841778978 | 32055.0 |

| | | | | | | | |
|---|---|---|---|---|---|---|---|
| 3 | 3:17303337_GTT_G | 17303337 | GTT | G | 1.468748143975356e-10 | 6.408586632214668 | 31988.0 |
| 3 | rs9883518 | 17485883 | T | C | 1.476303857076324e-10 | 6.407804140984278 | 32044.0 |
| 3 | rs145581379 | 17606493 | G | GAAA ACAA | 1.4778869924531e-10 | 6.407640682962817 | 31958.0 |
| 3 | rs142250908 | 90206899 | T | A | 1.4967881172577231e-10 | 6.405702258994558 | 31755.0 |
| 3 | rs6442680 | 17418533 | T | C | 1.500136184975251e-10 | 6.405361390122372 | 32168.0 |
| 3 | rs2733510 | 17263805 | C | T | 1.5024294175920043e-10 | 6.4051283434001824 | 32007.0 |
| 3 | rs2733493 | 17302355 | T | C | 1.5069918981977796e-10 | 6.404665719319255 | 32034.0 |
| 3 | rs62276591 | 90380252 | A | T | 1.5101099861984427e-10 | 6.404350339691867 | 31242.0 |
| 3 | rs9848610 | 17419843 | A | G | 1.513257430778287e-10 | 6.404032635505778 | 32173.0 |
| 3 | rs17043541 | 17426846 | T | C | 1.5247067361737924e-10 | 6.402882360274936 | 32163.0 |
| 3 | rs17043539 | 17426748 | C | G | 1.5391735805144452e-10 | 6.401440933740537 | 32164.0 |
| 3 | rs7623979 | 17406857 | A | G | 1.580627671930674e-10 | 6.397382749866451 | 32100.0 |
| 3 | rs11128836 | 17683892 | T | C | 1.5850750311088727e-10 | 6.396953565711813 | 32019.0 |
| 3 | rs6788763 | 17519053 | T | C | 1.587737308000255e-10 | 6.396697210290648 | 32049.0 |
| 3 | rs6807020 | 17692815 | C | G | 1.6004829262172697e-10 | 6.395475701923263 | 32012.0 |
| 3 | rs62276586 | 90376687 | C | G | 1.6230832307968725e-10 | 6.393332931764972 | 31203.0 |
| 3 | rs61236237 | 17691510 | T | C | 1.6356862835539613e-10 | 6.392150644464508 | 32015.0 |
| 3 | rs7650617 | 17743260 | A | G | 1.6413822271668568e-10 | 6.391619227994566 | 31839.0 |
| 3 | rs149819143 | 90035767 | C | CA | 1.650513879002891e-10 | 6.390771017645331 | 30076.0 |
| 3 | rs9815204 | 17631880 | C | T | 1.6986029390829932e-10 | 6.38637848140959 | 32000.0 |
| 3 | rs2733519 | 17249128 | A | G | 1.7100965893087818e-10 | 6.38534663102537 | 32124.0 |
| 3 | rs9847477 | 17607991 | C | G | 1.725457884361337e-10 | 6.383978090192692 | 31856.0 |
| 3 | rs9821024 | 90330554 | C | A | 1.7423155328598603e-10 | 6.382489872453733 | 31665.0 |
| 3 | rs9812906 | 17525505 | G | A | 1.7593755022828534e-10 | 6.3809980485359885 | 32046.0 |
| 3 | rs9799070 | 90400831 | G | A | 1.7690464886866338e-10 | 6.380158625491532 | 31335.0 |
| 3 | rs6782889 | 17692938 | G | T | 1.788410330965863e-10 | 6.378491273461911 | 31995.0 |
| 3 | 3:17304858_CAA_C | 17304858 | CAA | C | 1.8270051683886703e-10 | 6.375219981164969 | 30877.0 |
| 3 | rs6784025 | 17701058 | A | T | 1.8399958589070883e-10 | 6.374134063902567 | 32001.0 |
| 3 | 3:17702996_CT_C | 17702996 | CT | C | 1.844868085627637e-10 | 6.37372871540896 | 30547.0 |
| 3 | 3:17339063_TA_T | 17339063 | TA | T | 1.903427508855578e-10 | 6.368937012808582 | 28235.0 |
| 3 | rs6799259 | 17427142 | T | C | 1.904422463645999e-10 | 6.368856849831783 | 32161.0 |
| 3 | rs12493567 | 17426616 | A | G | 1.9150382529228545e-10 | 6.368004079518408 | 32167.0 |
| 3 | rs7648883 | 17407522 | C | T | 1.9353310644691061e-10 | 6.366386725531818 | 32144.0 |
| 3 | rs9310524 | 17640706 | C | T | 1.9436869262610882e-10 | 6.3657255669560495 | 31990.0 |
| 3 | rs2348001 | 17251577 | T | G | 1.958651085658917e-10 | 6.364548435356218 | 32127.0 |
| 3 | rs2733518 | 17249603 | T | C | 1.960371879962197e-10 | 6.364413635506259 | 32124.0 |
| 3 | rs9310526 | 17643034 | T | G | 1.9748197205436042e-10 | 6.3632863902980255 | 31991.0 |
| 3 | rs2733516 | 17252724 | T | G | 1.9892821424943902e-10 | 6.362166047707905 | 32129.0 |
| 3 | rs9860854 | 17645229 | G | C | 2.01962088656118e-10 | 6.359841462768397 | 31985.0 |
| 3 | rs58595750 | 90290037 | C | CT | 2.0445910088858053e-10 | 6.357953678630509 | 27240.0 |
| 3 | rs11720651 | 89261061 | A | G | 2.074457938519988e-10 | 6.355725047737504 | 31920.0 |
| 3 | rs149092028 | 90049995 | A | C | 2.076781580452919e-10 | 6.355552977107049 | 29596.0 |
| 3 | rs7617933 | 17252067 | G | T | 2.0968266647000828e-10 | 6.35407635735964 | 32119.0 |
| 3 | rs7644020 | 17481548 | C | T | 2.1090248482686714e-10 | 6.3531845132929625 | 32049.0 |
| 3 | rs10212255 | 17451108 | A | G | 2.1430247397254985e-10 | 6.350725026915907 | 32069.0 |
| 3 | rs12494513 | 17427622 | A | G | 2.203491734283313e-10 | 6.346443671289508 | 32164.0 |
| 3 | rs9798866 | 90490026 | G | A | 2.2217509490868184e-10 | 6.345173374506732 | 30936.0 |
| 3 | rs112433497 | 90376972 | C | A | 2.2976437971891068e-10 | 6.340000524759537 | 31330.0 |
| 3 | rs4377510 | 17686288 | G | A | 2.3053616870875642e-10 | 6.339483856097232 | 32014.0 |
| 3 | rs35234876 | 17432517 | G | C | 2.359376362886403e-10 | 6.335914480107903 | 32167.0 |
| 3 | rs9856545 | 90402558 | C | A | 2.427739329665769e-10 | 6.331509682025085 | 31330.0 |
| 3 | rs73143751 | 90378437 | C | T | 2.4394195433035207e-10 | 6.330769234699491 | 31347.0 |
| 3 | rs723813 | 17240402 | C | G | 2.497525591814647e-10 | 6.327136409554676 | 32119.0 |
| 3 | rs9682919 | 90368237 | A | G | 2.533729421033016e-10 | 6.324914435285116 | 31249.0 |

| | | | | | | | |
|---|---|---|---|---|---|---|---|
| 3 | rs62277419 | 90434757 | C | T | 2.5350674407508574e-10 | 6.324832911087443 | 31043.0 |
| 3 | rs62275607 | 90356428 | T | A | 2.57607718273616e-10 | 6.322354398918903 | 31194.0 |
| 3 | rs62275608 | 90356435 | C | G | 2.57607718273616e-10 | 6.322354398918903 | 31194.0 |
| 3 | rs2198953 | 90375306 | T | G | 2.690145334006776e-10 | 6.3156579112057685 | 31241.0 |
| 3 | rs4637311 | 17686445 | C | T | 2.711350014235499e-10 | 6.3144437016411175 | 32011.0 |
| 3 | rs10687921 | 17770840 | T | TGA | 2.7446030763826254e-10 | 6.3125581348778255 | 29717.0 |
| 3 | rs6793041 | 90205657 | C | A | 2.7729895963208907e-10 | 6.310966086976216 | 28858.0 |
| 3 | rs62266068 | 90497467 | A | T | 2.8094615293357186e-10 | 6.3089437729291475 | 31046.0 |
| 3 | rs9840464 | 90369496 | C | T | 2.820314450571651e-10 | 6.308346942799133 | 31183.0 |
| 3 | rs62264901 | 90478305 | C | T | 2.8281191940284167e-10 | 6.30791912462621 | 31023.0 |
| 3 | rs2733521 | 17243164 | G | A | 2.8775618798119094e-10 | 6.305235411469523 | 32121.0 |
| 3 | rs113465892 | 17747914 | TAC | T | 2.891676184069592e-10 | 6.304477557141579 | 31319.0 |
| 3 | rs9682654 | 90473594 | G | T | 2.971113808104428e-10 | 6.3002784648407175 | 30403.0 |
| 3 | rs2198954 | 90372187 | A | G | 2.9730370065633415e-10 | 6.3001781690454735 | 31304.0 |
| 3 | rs2733528 | 17236255 | A | G | 3.0117821958917317e-10 | 6.298170969163297 | 32121.0 |
| 3 | rs7651278 | 17443745 | C | T | 3.065169848080715e-10 | 6.295446136743219 | 32168.0 |
| 3 | rs9799179 | 90414184 | C | A | 3.075560857886289e-10 | 6.294921185749429 | 31057.0 |
| 3 | rs2470581 | 17244668 | C | G | 3.2418850464484887e-10 | 6.286746302475202 | 32124.0 |
| 3 | rs2347959 | 90039465 | G | A | 3.245224241893099e-10 | 6.286586408252963 | 31638.0 |
| 3 | rs11425835 | 17471456 | T | TA | 3.276094392887528e-10 | 6.285115781495564 | 32035.0 |
| 3 | rs62276658 | 90403891 | G | T | 3.327013874945474e-10 | 6.282719335615753 | 31317.0 |
| 3 | rs2733526 | 17238792 | T | C | 3.44090281131076e-10 | 6.277486431471587 | 32122.0 |
| 3 | rs76658459 | 90357875 | C | A | 3.446358001699724e-10 | 6.277240043421522 | 31249.0 |
| 3 | rs2596657 | 17238673 | G | A | 3.4597345727539264e-10 | 6.276637487785997 | 32112.0 |
| 3 | 3:17279462_TA_T | 17279462 | TA | T | 3.5200627563014284e-10 | 6.273947928023027 | 31835.0 |
| 3 | rs9823767 | 17615737 | A | G | 3.631101345215965e-10 | 6.2691131818530845 | 31948.0 |
| 3 | rs2596658 | 17245725 | C | T | 3.639473451150596e-10 | 6.268754524751627 | 32129.0 |
| 3 | rs2733529 | 17235874 | C | T | 3.7459253199487286e-10 | 6.264263110704932 | 32108.0 |
| 3 | rs201902007 | 90423228 | G | GA | 3.8994082240492825e-10 | 6.258001999684257 | 30734.0 |
| 3 | rs759739186 | 17421533 | GTGTA | G | 4.0559207247942577e-10 | 6.2518603198392855 | 31603.0 |
| 3 | rs10656408 | 17791796 | C | CATATAT | 4.057895634302807e-10 | 6.25178431025808 | 29811.0 |
| 3 | rs10662235 | 17564583 | G | GTAT | 4.1862793810347685e-10 | 6.246919027517609 | 31844.0 |
| 3 | rs2279832 | 147101640 | T | C | 4.2303786447138963e-10 | 6.245281376143963 | 31865.0 |
| 3 | rs6772394 | 17488124 | C | T | 4.269819844730177e-10 | 6.243830760147801 | 32045.0 |
| 3 | rs9854201 | 17471528 | A | G | 4.3115906002323777e-10 | 6.242308657171183 | 32063.0 |
| 3 | rs2733527 | 17237409 | A | T | 4.435781149933546e-10 | 6.237866794871361 | 32118.0 |
| 3 | rs5846952 | 17227194 | T | TA | 4.466940896612823e-10 | 6.236771372195902 | 30893.0 |
| 3 | rs11915971 | 17779893 | A | T | 4.4952226867524217e-10 | 6.2357835619466675 | 31374.0 |
| 3 | rs13320693 | 17464812 | A | T | 4.53919788554256e-10 | 6.234259605856604 | 31984.0 |
| 3 | rs9310528 | 17661932 | C | T | 4.65980034532062e-10 | 6.230152941805053 | 31886.0 |
| 3 | 3:17398486_TAA_T | 17398486 | TAA | T | 4.825367803379669e-10 | 6.224681002575593 | 31926.0 |
| 3 | rs9681339 | 90473595 | A | T | 5.014400051011565e-10 | 6.218653263966407 | 30379.0 |
| 3 | rs9839103 | 147228736 | C | T | 5.192440872668541e-10 | 6.213175472825212 | 30662.0 |
| 3 | rs13320647 | 17589880 | A | C | 5.510672237216866e-10 | 6.2038259718734174 | 31997.0 |
| 3 | rs13327810 | 17464911 | C | A | 5.550205194263967e-10 | 6.202701499846581 | 32061.0 |
| 3 | rs9875973 | 17584573 | T | C | 5.610342300051409e-10 | 6.201005864453286 | 31970.0 |
| 3 | rs11128078 | 90030538 | C | T | 5.639810392481288e-10 | 6.200181441631368 | 31639.0 |
| 3 | rs9810675 | 17449619 | C | T | 5.813910681366148e-10 | 6.195394819039035 | 32071.0 |
| 3 | rs11708391 | 17414753 | A | C | 6.051400128786527e-10 | 6.189086161939107 | 32140.0 |
| 3 | rs34892692 | 17448866 | G | A | 6.343949519271295e-10 | 6.1816389275722035 | 32059.0 |
| 3 | rs11717644 | 17461255 | C | T | 6.467329307355059e-10 | 6.1785981117027555 | 32070.0 |
| 3 | rs2316653 | 90442925 | A | G | 6.468243081586934e-10 | 6.178575802654545 | 30932.0 |
| 3 | rs9878954 | 17509481 | T | A | 6.584988755999898e-10 | 6.175750548408445 | 32053.0 |
| 3 | 3:17462769_ACT_A | 17462769 | ACT | A | 7.197403675217658e-10 | 6.161686938799421 | 31959.0 |

| | | | | | | | |
|---|---|---|---|---|---|---|---|
| 3 | rs6442683 | 17462945 | G | A | 7.208793010605877e-10 | 6.161436605115073 | 32064.0 |
| 3 | 3:17602111_TA_T | 17602111 | TA | T | 7.385612456162229e-10 | 6.157598893192391 | 31738.0 |
| 3 | 3:17267802_CT_C | 17267802 | CT | C | 7.56675543313846e-10 | 6.153759184047128 | 28760.0 |
| 3 | rs6787756 | 17453223 | T | C | 7.615572085178094e-10 | 6.152739747768706 | 32063.0 |
| 3 | rs1463221 | 17221017 | T | C | 7.641088184674238e-10 | 6.152209429857828 | 31285.0 |
| 3 | rs9756494 | 17588575 | A | G | 7.642845150439375e-10 | 6.1521729772745966 | 31994.0 |
| 3 | rs6442681 | 17454766 | A | T | 7.745374977985819e-10 | 6.150059776681527 | 32063.0 |
| 3 | rs1875748 | 147102874 | C | T | 7.950917193115554e-10 | 6.145904382829273 | 31859.0 |
| 3 | rs9799190 | 90461761 | A | G | 8.038964801731507e-10 | 6.144156312684956 | 30886.0 |
| 3 | rs7632989 | 17462283 | G | T | 9.121844394591506e-10 | 6.1240636240694 | 32066.0 |
| 3 | rs2733530 | 17233114 | A | G | 9.504347412558647e-10 | 6.1175189289598455 | 31312.0 |
| 3 | rs9848766 | 17412485 | G | C | 9.632048006194992e-10 | 6.115391035002253 | 32156.0 |
| 3 | rs9874826 | 17410188 | A | T | 9.65556376675017e-10 | 6.115002189874545 | 32157.0 |
| 3 | rs112149405 | 89886780 | T | C | 9.66241531715435e-10 | 6.114889069708164 | 31537.0 |
| 3 | rs201959300 | 17385963 | T | C | 9.7795541039246e-10 | 6.112967096120636 | 32110.0 |
| 3 | rs283936 | 17367376 | A | T | 9.867431298342778e-10 | 6.111539918037486 | 32125.0 |
| 3 | rs9820703 | 17414107 | A | G | 9.89838248815261e-10 | 6.111040202074029 | 32159.0 |
| 3 | rs10663489 | 17526824 | T | TTTA | 9.93551795459552e-10 | 6.110442646064807 | 31959.0 |
| 3 | rs9823064 | 17520222 | A | G | 1.0594379758376213e-09 | 6.100187979985567 | 32046.0 |
| 3 | rs2198947 | 90487716 | G | C | 1.0601338807411276e-09 | 6.1000830214806445 | 30521.0 |
| 3 | rs6763593 | 17379330 | G | A | 1.0899885989996202e-09 | 6.095642359874638 | 32134.0 |
| 3 | rs9811041 | 89367555 | G | A | 1.1027216446578372e-09 | 6.093784386146092 | 31205.0 |
| 3 | rs56328245 | 90456144 | T | A | 1.1094005526337707e-09 | 6.092818163683464 | 30722.0 |
| 3 | rs9310517 | 17550666 | G | A | 1.111586516395955e-09 | 6.0925031568228745 | 32050.0 |
| 3 | rs2733515 | 17323831 | T | C | 1.132352843794062e-09 | 6.089540405665464 | 32130.0 |
| 3 | 3:17776417_C_CA | 17776417 | C | CA | 1.1494198590563922e-09 | 6.087144832585776 | 28297.0 |
| 3 | rs62266110 | 93537923 | G | A | 1.1768907828427615e-09 | 6.0833608269175325 | 31789.0 |
| 3 | rs9821318 | 17567220 | C | T | 1.1778794040157895e-09 | 6.083226259291004 | 32005.0 |
| 3 | rs9757853 | 90436845 | A | G | 1.1825386208317286e-09 | 6.082593541818183 | 30957.0 |
| 3 | rs2596656 | 17235490 | G | A | 1.2114388726085825e-09 | 6.078722382657428 | 32043.0 |
| 3 | rs2100628 | 90439303 | A | G | 1.2388983816427347e-09 | 6.075126732870567 | 30808.0 |
| 3 | rs9831141 | 90433267 | C | T | 1.2394911708167675e-09 | 6.075049970576347 | 31152.0 |
| 3 | rs9757339 | 17569829 | T | C | 1.2469750894894139e-09 | 6.074083918519257 | 31980.0 |
| 3 | rs2733489 | 17312588 | A | G | 1.2493539451106585e-09 | 6.073778030545197 | 32078.0 |
| 3 | rs10510475 | 17422679 | A | T | 1.2956195165956787e-09 | 6.0679391752891245 | 32172.0 |
| 3 | rs283942 | 17346033 | A | G | 1.3229206069877164e-09 | 6.06458832758192 | 32148.0 |
| 3 | rs1449882 | 17311336 | T | C | 1.328104640842911e-09 | 6.0639596784468806 | 32064.0 |
| 3 | rs35521582 | 17496173 | T | C | 1.3432634597981624e-09 | 6.0621350530453535 | 32029.0 |
| 3 | rs283930 | 17360961 | A | T | 1.3520189599944006e-09 | 6.06109030044814 | 32140.0 |
| 3 | rs283934 | 17367065 | T | C | 1.3520288961259135e-09 | 6.061089118566817 | 32140.0 |
| 3 | rs1992737 | 17422005 | A | G | 1.355899505192296e-09 | 6.060629360817982 | 32175.0 |
| 3 | rs9850127 | 17398386 | T | C | 1.368712803681188e-09 | 6.0591164499841605 | 32100.0 |
| 3 | rs13318886 | 17399281 | G | T | 1.4141720161802712e-09 | 6.053858091339857 | 32131.0 |
| 3 | rs17200839 | 17425832 | C | T | 1.4187071239722966e-09 | 6.053342575643916 | 32186.0 |
| 3 | rs34958195 | 17434610 | G | GT | 1.4282187406436758e-09 | 6.052266566195757 | 31092.0 |
| 3 | rs4312642 | 17434006 | A | G | 1.4579018234152366e-09 | 6.048952997151655 | 32164.0 |
| 3 | rs1470667 | 17309934 | C | A | 1.472343250946373e-09 | 6.0473645845670685 | 32054.0 |
| 3 | rs9817290 | 17408297 | A | C | 1.4861727235914304e-09 | 6.045857650499399 | 32129.0 |
| 3 | rs9866664 | 90437988 | A | G | 1.5705946810903372e-09 | 6.0369447495101545 | 30947.0 |
| 3 | rs35863421 | 17521749 | A | AG | 1.5730029574227416e-09 | 6.036697401490853 | 31873.0 |
| 3 | rs2060623 | 17338680 | T | C | 1.5762581594218193e-09 | 6.036363653774417 | 32109.0 |
| 3 | rs9713931 | 90390418 | A | G | 1.597602478388743e-09 | 6.034191774732009 | 31224.0 |
| 3 | rs11389760 | 17440236 | A | AC | 1.5985863227505005e-09 | 6.034092347263081 | 32159.0 |
| 3 | rs2348004 | 17295308 | G | A | 1.633961377025016e-09 | 6.030556395053347 | 32015.0 |

| | | | | | | | |
|---|---|---|---|---|---|---|---|
| 3 | rs2733496 | 17281330 | C | T | 1.6602153843614626e-09 | 6.027980049290748 | 32024.0 |
| 3 | rs2596641 | 17291294 | A | G | 1.6658060979982918e-09 | 6.027436554029865 | 32019.0 |
| 3 | rs6762516 | 17375264 | T | A | 1.6707422185051005e-09 | 6.026958169828683 | 32136.0 |
| 3 | rs9858006 | 90390344 | A | T | 1.7021074303941292e-09 | 6.023950214086997 | 31225.0 |
| 3 | rs9861554 | 90349123 | C | G | 1.7040276100930848e-09 | 6.0237678257257805 | 31410.0 |
| 3 | rs62245947 | 17497051 | T | G | 1.705263136152051e-09 | 6.023650575084336 | 32034.0 |
| 3 | rs1521806 | 90090619 | A | C | 1.7153297908557355e-09 | 6.022698329326974 | 31854.0 |
| 3 | rs7647905 | 17433383 | T | C | 1.7274894739377767e-09 | 6.02155533069468 | 32167.0 |
| 3 | rs34096182 | 89465849 | A | T | 1.7416575977787074e-09 | 6.020233388473454 | 30026.0 |
| 3 | rs36029966 | 17389500 | C | CA | 1.7523842740917609e-09 | 6.019239498488884 | 32072.0 |
| 3 | rs11449656 | 17371025 | T | TA | 1.7847858927003157e-09 | 6.016272900552857 | 32100.0 |
| 3 | rs1867772 | 17391339 | T | A | 1.7907626987786938e-09 | 6.015731419711689 | 32141.0 |
| 3 | rs7614318 | 17440747 | C | G | 1.8019316936768264e-09 | 6.014724245504999 | 32165.0 |
| 3 | rs283929 | 17359916 | C | T | 1.8462300987663765e-09 | 6.010788613645768 | 32142.0 |
| 3 | rs9843640 | 17441862 | C | G | 1.941805314923773e-09 | 6.002601720292059 | 32166.0 |
| 3 | rs2596643 | 17294388 | A | G | 1.9491244723370795e-09 | 6.001991050476347 | 32019.0 |
| 3 | rs2733495 | 17296298 | A | T | 1.9742851995278287e-09 | 5.999908687719621 | 32011.0 |
| 3 | 3:17442866_CATATGT_C | 17442866 | CATATGT | C | 2.1013376346676632e-09 | 5.989773279452883 | 32022.0 |
| 3 | rs528162866 | 17747475 | C | CT | 2.1352085799300925e-09 | 5.987172071963976 | 29292.0 |
| 3 | rs2348005 | 17295756 | T | C | 2.2611172495202192e-09 | 5.977842829186933 | 31999.0 |
| 3 | rs9864494 | 17473346 | A | G | 2.279827717811538e-09 | 5.976499844293282 | 32057.0 |
| 3 | rs1449888 | 17272276 | T | C | 2.283351798664924e-09 | 5.9762480976306485 | 32012.0 |
| 3 | 3:90272973_AAAAT_A | 90272973 | AAAAT | A | 2.293738966218558e-09 | 5.975508274455246 | 31144.0 |
| 3 | rs9877833 | 90138312 | G | A | 2.304407585880217e-09 | 5.974751794434964 | 31912.0 |
| 3 | rs9835988 | 17517721 | A | G | 2.395120388063961e-09 | 5.968454075208341 | 32066.0 |
| 3 | rs283940 | 17344285 | A | T | 2.414498597022578e-09 | 5.9671389003014275 | 32170.0 |
| 3 | rs1449885 | 17261756 | A | G | 2.437753730424244e-09 | 5.9655741083863605 | 32022.0 |
| 3 | 3:17346381_TA_T | 17346381 | TA | T | 2.458025378920036e-09 | 5.964221884959509 | 32158.0 |
| 3 | rs5846961 | 17467525 | G | GA | 2.467846148299688e-09 | 5.963570689973358 | 32071.0 |
| 3 | rs11419267 | 17487665 | C | CT | 2.4715838302346407e-09 | 5.96332351487291 | 32044.0 |
| 3 | rs1449887 | 17272228 | A | T | 2.5233245633718826e-09 | 5.959938774090163 | 32013.0 |
| 3 | rs283945 | 17346756 | C | T | 2.546924255988956e-09 | 5.958417333848318 | 32171.0 |
| 3 | rs7615612 | 17516671 | G | C | 2.579184589222341e-09 | 5.956359617993027 | 31446.0 |
| 3 | rs17200818 | 17410411 | T | A | 2.622271132217394e-09 | 5.953650109747441 | 32160.0 |
| 3 | rs62273833 | 147255596 | G | A | 2.642474894101568e-09 | 5.952394489936948 | 31149.0 |
| 3 | rs62276243 | 90121534 | C | T | 2.750462075101354e-09 | 5.9458379157627546 | 31841.0 |
| 3 | rs3058046 | 147090583 | G | GAA | 2.774697214046381e-09 | 5.944400928654988 | 30090.0 |
| 3 | rs9681592 | 90485493 | A | G | 2.851523816326651e-09 | 5.939925069182968 | 31188.0 |
| 3 | rs4974357 | 90127762 | T | C | 3.071391871134277e-09 | 5.9277363035417485 | 31858.0 |
| 3 | rs1446303 | 17482564 | T | C | 3.168542233341763e-09 | 5.922619072286135 | 32049.0 |
| 3 | rs2028874 | 17426715 | C | G | 3.3590318363439358e-09 | 5.913014211822586 | 32171.0 |
| 3 | rs17804073 | 90124806 | T | A | 3.439290490621876e-09 | 5.909125291642253 | 31877.0 |
| 3 | rs2596648 | 17302558 | A | G | 3.602949024415061e-09 | 5.901461777527333 | 32005.0 |
| 3 | rs2733492 | 17308296 | C | A | 3.655730159262074e-09 | 5.899062380332516 | 32043.0 |
| 3 | rs9830240 | 17398284 | G | T | 3.781867856487939e-09 | 5.893462138105296 | 31939.0 |
| 3 | rs2596650 | 17307841 | C | T | 3.784217801437365e-09 | 5.8933595401153624 | 32039.0 |
| 3 | rs7613624 | 147237920 | T | C | 3.797031045207153e-09 | 5.892801205862018 | 30816.0 |
| 3 | rs9851518 | 17455397 | A | G | 3.950817449624112e-09 | 5.886239473461068 | 32063.0 |
| 3 | rs6801690 | 17406273 | C | T | 4.084444692510904e-09 | 5.8807366942570365 | 32142.0 |
| 3 | rs2596649 | 17307395 | G | A | 4.0863896788412025e-09 | 5.880657900317372 | 32041.0 |
| 3 | rs9870604 | 147234379 | C | G | 4.160197250889698e-09 | 5.8776945188184975 | 30792.0 |
| 3 | rs2596655 | 17232635 | G | A | 4.251465012227832e-09 | 5.874100073796918 | 31239.0 |
| 3 | rs75948196 | 90420549 | T | C | 4.3411943332016224e-09 | 5.870638693351692 | 30960.0 |
| 3 | rs7652792 | 17439143 | G | A | 4.409280998516757e-09 | 5.86805834697288 | 32163.0 |

| | | | | | | | |
|---|---|---|---|---|---|---|---|
| 3 | rs2596662 | 17253663 | A | T | 4.414620385550493e-09 | 5.867857637772263 | 32080.0 |
| 3 | rs6792443 | 17458313 | G | T | 4.455959820786778e-09 | 5.866311623608463 | 32060.0 |
| 3 | rs35742571 | 17459374 | A | AT | 4.696869794338952e-09 | 5.8575706671798 | 31877.0 |
| 3 | rs73004648 | 147137674 | A | G | 4.759360259641056e-09 | 5.855374598792473 | 32043.0 |
| 3 | 3:17428891_AT_A | 17428891 | AT | A | 4.913473099116154e-09 | 5.850076386163192 | 32165.0 |
| 3 | rs75162052 | 147137597 | T | C | 5.116264616483677e-09 | 5.84334583702532 | 32041.0 |
| 3 | rs57499472 | 147239337 | T | C | 5.15222874753324e-09 | 5.842179365820226 | 30828.0 |
| 3 | rs9842092 | 147241494 | A | G | 5.233879866726416e-09 | 5.839560211412637 | 30829.0 |
| 3 | rs5846955 | 17306752 | A | AG | 5.348886418802155e-09 | 5.835937752263532 | 31962.0 |
| 3 | rs7625855 | 17470782 | T | A | 6.136832127763889e-09 | 5.81298565727831 | 32064.0 |
| 3 | rs9310516 | 17470827 | T | A | 6.136832127763889e-09 | 5.81298565727831 | 32064.0 |
| 3 | rs1374197 | 17394615 | A | G | 6.245441011947333e-09 | 5.810049402299937 | 31882.0 |
| 3 | rs2168715 | 90452164 | A | C | 6.498409661178453e-09 | 5.803398501490575 | 30903.0 |
| 3 | rs79769556 | 90423789 | A | G | 6.671965352986565e-09 | 5.7989795844944965 | 30918.0 |
| 3 | rs71105148 | 90190366 | C | CA | 7.146937653828485e-09 | 5.7874348871536885 | 28351.0 |
| 3 | rs2733525 | 17238881 | C | T | 7.193843407579309e-09 | 5.7863355751113 | 32071.0 |
| 3 | rs10865749 | 17450271 | A | T | 7.2551279569350886e-09 | 5.784909729195174 | 32072.0 |
| 3 | rs2121262 | 17470114 | C | T | 7.469262367275186e-09 | 5.7800180222270825 | 32064.0 |
| 3 | rs9813532 | 17465704 | G | A | 7.54419878536488e-09 | 5.7783383277946445 | 32059.0 |
| 3 | rs9882273 | 17464102 | C | T | 7.616666788650926e-09 | 5.7767293232286585 | 32061.0 |
| 3 | rs9824952 | 17448437 | T | A | 7.940389199298085e-09 | 5.76971885911191 | 32059.0 |
| 3 | rs12488687 | 17461342 | A | T | 8.005571972349632e-09 | 5.768340944530977 | 32059.0 |
| 3 | rs201380658 | 17393317 | AG | A | 8.087976844122914e-09 | 5.766614497257367 | 30357.0 |
| 3 | 3:17474521_CCCCA_C | 17474521 | CCCCA | C | 8.276724485687683e-09 | 5.762723673453406 | 32023.0 |
| 3 | rs6442682 | 17461731 | T | C | 8.305424831044279e-09 | 5.762139616369718 | 32070.0 |
| 3 | rs1120412 | 17452025 | G | A | 8.305779766114334e-09 | 5.762132405669299 | 32066.0 |
| 3 | rs34633537 | 17852607 | G | GA | 8.460693380244274e-09 | 5.759013505305696 | 31957.0 |
| 3 | rs2733522 | 17243002 | T | C | 8.472629166308509e-09 | 5.758775509080987 | 32071.0 |
| 3 | rs7625175 | 17455081 | C | T | 8.911219892393935e-09 | 5.750248829042838 | 32066.0 |
| 3 | rs34732565 | 90144346 | T | A | 9.08373065623957e-09 | 5.747006374034588 | 31900.0 |
| 3 | rs9681820 | 90481415 | A | T | 9.819561357021818e-09 | 5.733816420058807 | 30981.0 |
| 3 | rs2198951 | 90460470 | G | C | 9.890160353991436e-09 | 5.732601858673518 | 30910.0 |
| 3 | rs1532769 | 147090343 | C | T | 1.0184218144155572e-08 | 5.727632011390803 | 31249.0 |
| 3 | rs12632358 | 17740995 | C | T | 1.065505002779513e-08 | 5.719957821055236 | 29932.0 |
| 3 | rs9830582 | 17458832 | C | A | 1.0659810173248368e-08 | 5.719881929857707 | 32078.0 |
| 3 | rs1394041 | 147096847 | C | A | 1.0727280084204777e-08 | 5.7188097817204655 | 31806.0 |
| 3 | 3:17452696_AT_A | 17452696 | AT | A | 1.3708605455888141e-08 | 5.676989286787379 | 30456.0 |
| 3 | rs4974358 | 90141506 | A | G | 1.4208940691664729e-08 | 5.670851638774238 | 31972.0 |
| 3 | rs34185023 | 147095042 | G | A | 1.4300303536887922e-08 | 5.6697535966276185 | 31821.0 |
| 3 | rs142854083 | 17774212 | A | TAAATAAATAAATAAATAAA | 1.4438276666651231e-08 | 5.668108219232932 | 30741.0 |
| 3 | rs9865185 | 89252480 | A | C | 1.4836749275724107e-08 | 5.663440661114489 | 31788.0 |
| 3 | rs7632016 | 147100106 | A | C | 1.7641899785133312e-08 | 5.633665553340478 | 32022.0 |
| 3 | rs6577596 | 17832829 | C | T | 2.2395682062269682e-08 | 5.592396374669118 | 31716.0 |
| 3 | rs112272175 | 89269015 | C | T | 2.24688252982316e-08 | 5.591830390511554 | 31673.0 |
| 3 | rs2060626 | 17339260 | T | A | 2.2635658772900946e-08 | 5.590546092122966 | 29018.0 |
| 3 | rs2596682 | 17221849 | C | A | 2.291470943477489e-08 | 5.5884183340076179 | 31272.0 |
| 3 | rs2596676 | 17220717 | G | A | 2.2988711049416235e-08 | 5.58785829183338 | 31257.0 |
| 3 | rs10804719 | 147117260 | A | G | 2.333680813045812e-08 | 5.5852471593706134 | 32018.0 |
| 3 | rs283914 | 17330649 | T | C | 2.3381263670837704e-08 | 5.584916417063188 | 32064.0 |
| 3 | rs73004638 | 147113586 | A | T | 2.436295750902088e-08 | 5.57776421283271 | 32011.0 |
| 3 | rs2470578 | 17330092 | G | A | 2.5688301175816943e-08 | 5.5685397629165925 | 31801.0 |
| 3 | rs73004642 | 147116846 | A | C | 2.6149752873805392e-08 | 5.565436145606383 | 32017.0 |
| 3 | rs374871846 | 89979666 | AC | A | 2.7571019574792272e-08 | 5.556200476076803 | 27026.0 |

| | | | | | | | |
|---|---|---|---|---|---|---|---|
| 3 | rs62274613 | 89157751 | G | A | 2.8025679008703726e-08 | 5.553343369121187 | 32010.0 |
| 3 | rs62274656 | 89271959 | C | T | 2.8504831533809592e-08 | 5.550380614746951 | 31957.0 |
| 3 | rs62274657 | 89272043 | C | G | 2.938196051321304e-08 | 5.5450801165635095 | 31955.0 |
| 3 | rs2279830 | 147105790 | T | C | 2.996055757166998e-08 | 5.541667044712511 | 32186.0 |
| 3 | rs12107172 | 147134039 | A | G | 3.0809350587529045e-08 | 5.536774103732859 | 32023.0 |
| 3 | rs7373232 | 89268266 | A | G | 3.21498690295381e-08 | 5.529306779811202 | 31959.0 |
| 3 | rs73004644 | 147124706 | G | A | 3.2359367948082e-08 | 5.528167144065436 | 32039.0 |
| 3 | rs62274614 | 89159719 | C | T | 3.2631838258342037e-08 | 5.526695616038647 | 32004.0 |
| 3 | rs57468276 | 89269988 | A | T | 3.4479578660735746e-08 | 5.517020215133528 | 31965.0 |
| 3 | rs1463222 | 17221252 | C | A | 3.4899737659542324e-08 | 5.514890401152521 | 31266.0 |
| 3 | rs9824191 | 17453982 | T | C | 3.5010773137913804e-08 | 5.5143317094901345 | 32062.0 |
| 3 | rs62274654 | 89271419 | C | G | 3.828366102729661e-08 | 5.498591620260124 | 31960.0 |
| 3 | rs1503738 | 147093600 | A | G | 4.30290559708264e-08 | 5.477946343696077 | 31457.0 |
| 3 | rs9809894 | 89261561 | A | T | 4.322257883663679e-08 | 5.477152053945327 | 31992.0 |
| 3 | rs9852594 | 89262358 | C | A | 4.403254876593951e-08 | 5.473864626328236 | 31991.0 |
| 3 | rs9310112 | 89260495 | A | C | 4.51494112300714e-08 | 5.469426496832137 | 31998.0 |
| 3 | rs9852981 | 89253461 | T | C | 4.8516767742428295e-08 | 5.456662350193096 | 32017.0 |
| 3 | rs76407105 | 90329812 | G | A | 4.935801479370472e-08 | 5.4536077493821 | 31436.0 |
| 5 | rs145120402 | 93174765 | A | C | 1.8342886641780968e-09 | 6.011840399143533 | 32127.0 |
| 5 | 5:94068140_AC_A | 94068140 | AC | A | 6.7993445650989546e-09 | 5.7958069423521925 | 31981.0 |
| 5 | rs11955446 | 94073541 | T | C | 9.945874699859212e-09 | 5.731649302485057 | 32092.0 |
| 5 | rs369272402 | 93106457 | G | A | 1.0568052384284206e-08 | 5.72135066333192 | 32059.0 |
| 5 | rs115877304 | 92797166 | C | T | 1.1105662817024772e-08 | 5.7129158126540815 | 32184.0 |
| 5 | rs9791082 | 94075700 | T | C | 1.1245505958331708e-08 | 5.710786788866312 | 32100.0 |
| 5 | rs7442779 | 92788278 | A | G | 1.2380968628457991e-08 | 5.694395640362044 | 32186.0 |
| 5 | rs11955417 | 94073489 | T | C | 1.2896839546535178e-08 | 5.687426082530264 | 32095.0 |
| 5 | rs11135411 | 94072196 | A | T | 1.3228063419128444e-08 | 5.683092553355047 | 32085.0 |
| 5 | rs10684163 | 93998112 | G | GTTA | 1.3667150269273305e-08 | 5.6775075422171195 | 32054.0 |
| 5 | rs200712407 | 93280729 | T | TA | 1.4138860793113416e-08 | 5.671698550493665 | 32117.0 |
| 5 | rs17083079 | 92683607 | G | A | 1.4279443174335546e-08 | 5.670003704796567 | 32186.0 |
| 5 | rs72773593 | 94068179 | T | G | 1.5587772116202267e-08 | 5.654965344305387 | 32134.0 |
| 5 | 5:92797041_TG_T | 92797041 | TG | T | 1.5933130384938252e-08 | 5.651200290175116 | 32181.0 |
| 5 | rs140083818 | 93150564 | G | A | 1.661870673012065e-08 | 5.6439554769913185 | 32154.0 |
| 5 | rs880264 | 94069803 | T | C | 1.6626696663996366e-08 | 5.643872766489857 | 31943.0 |
| 5 | rs35111621 | 94067705 | A | G | 1.6689572242714282e-08 | 5.643223232545884 | 32128.0 |
| 5 | rs12523457 | 94069338 | C | T | 1.6896249002667543e-08 | 5.641104785849002 | 32119.0 |
| 5 | rs141834426 | 93385558 | G | C | 1.7166811229392963e-08 | 5.638369229636546 | 32161.0 |
| 5 | rs12109404 | 94072931 | A | G | 1.768156577594884e-08 | 5.633278416558889 | 31955.0 |
| 5 | rs2003118 | 94069530 | G | A | 1.7728439717585573e-08 | 5.632822016473777 | 32126.0 |
| 5 | rs61152231 | 94067762 | A | G | 1.784697721963519e-08 | 5.631673053977108 | 32137.0 |
| 5 | rs74765438 | 94080778 | G | GA | 1.7902745712377323e-08 | 5.631135061101539 | 31943.0 |
| 5 | rs57402656 | 94065844 | G | GT | 1.7918105660953198e-08 | 5.630987171095782 | 31991.0 |
| 5 | rs199920547 | 93961802 | C | CA | 1.84174153676272e-08 | 5.626245525115798 | 31941.0 |
| 5 | rs61238929 | 94079820 | T | C | 1.8422418248498005e-08 | 5.62619865022276 | 32015.0 |
| 5 | rs67577452 | 94068397 | A | G | 1.893610225423365e-08 | 5.621450259368147 | 32137.0 |
| 5 | rs67573880 | 94068468 | A | G | 1.893610225423365e-08 | 5.621450259368147 | 32137.0 |
| 5 | rs114072973 | 93372299 | C | A | 1.97020529407094e-08 | 5.6145973099942195 | 32154.0 |
| 5 | rs12515845 | 94069040 | T | G | 2.0298945456439636e-08 | 5.609433945501047 | 32125.0 |
| 5 | rs114891177 | 93397741 | C | T | 2.033055720835697e-08 | 5.609164617797831 | 32138.0 |
| 5 | rs1451735 | 93983612 | C | A | 2.469223853612216e-08 | 5.575427742024976 | 32186.0 |
| 5 | rs115840240 | 93509229 | G | A | 2.5653435267643315e-08 | 5.568776456594041 | 31920.0 |
| 5 | rs17083945 | 94049046 | T | C | 3.4196639092687804e-08 | 5.518468685653462 | 32186.0 |
| 5 | rs1390644 | 94000772 | G | T | 3.783847730314352e-08 | 5.500654150151289 | 32171.0 |
| 5 | rs7735473 | 94063520 | T | G | 3.8052210577467335e-08 | 5.499661006674137 | 32155.0 |

| | | | | | | | |
|---|---|---|---|---|---|---|---|
| 5 | rs78371824 | 94042149 | A | AT | 3.80733840953267e-08 | 5.4995629154516585 | 32121.0 |
| 5 | rs75522556 | 93970841 | C | CCT | 3.859468230726556e-08 | 5.497164418483997 | 32072.0 |
| 5 | rs67674687 | 94060082 | C | G | 3.888293750538777e-08 | 5.495851608610331 | 32162.0 |
| 5 | rs2201620 | 94004124 | C | T | 3.9184055835361716e-08 | 5.494490256559926 | 32141.0 |
| 5 | rs66901899 | 94062810 | A | C | 3.920829823461168e-08 | 5.494381098641886 | 32164.0 |
| 5 | rs72773517 | 93970216 | C | T | 3.975420029750141e-08 | 5.491940203508388 | 32121.0 |
| 5 | 5:92872910_GA_G | 92872910 | GA | G | 4.000339006564027e-08 | 5.490836785587875 | 32068.0 |
| 5 | rs1006626 | 94015064 | C | T | 4.049792993616618e-08 | 5.488666553165093 | 32153.0 |
| 5 | rs66578784 | 94015540 | T | C | 4.049792993616618e-08 | 5.488666553165093 | 32153.0 |
| 5 | rs67313609 | 94016254 | T | G | 4.049792993616618e-08 | 5.488666553165093 | 32153.0 |
| 5 | rs143236930 | 94006853 | T | TTATC | 4.111161485642439e-08 | 5.4860089190386345 | 32149.0 |
| 5 | rs6890092 | 94024807 | G | A | 4.280990766094194e-08 | 5.4788500024400815 | 32151.0 |
| 5 | rs12653116 | 94024851 | G | A | 4.280990766094194e-08 | 5.4788500024400815 | 32151.0 |
| 5 | 5:94000377_CT_C | 94000377 | CT | C | 4.32969448825642e-08 | 5.4768477448038695 | 32162.0 |
| 5 | rs17083887 | 94003296 | G | A | 4.3445075404239205e-08 | 5.4762430952682495 | 32153.0 |
| 5 | rs2220965 | 94003758 | G | A | 4.3445075404239205e-08 | 5.4762430952682495 | 32153.0 |
| 5 | rs72773547 | 94004908 | A | C | 4.3445075404239205e-08 | 5.4762430952682495 | 32153.0 |
| 5 | rs72773549 | 94005100 | T | C | 4.3445075404239205e-08 | 5.4762430952682495 | 32153.0 |
| 5 | rs17083888 | 94007823 | A | G | 4.3445075404239205e-08 | 5.4762430952682495 | 32153.0 |
| 5 | rs1124668 | 94013241 | T | G | 4.3445075404239205e-08 | 5.4762430952682495 | 32153.0 |
| 5 | rs67831380 | 94015736 | T | G | 4.3445075404239205e-08 | 5.4762430952682495 | 32153.0 |
| 5 | rs2270626 | 94024016 | G | C | 4.3445075404239205e-08 | 5.4762430952682495 | 32153.0 |
| 5 | rs2032813 | 94001252 | A | G | 4.662003306897137e-08 | 5.463742244618361 | 32173.0 |
| 5 | rs6556851 | 93993844 | G | A | 4.7241614353506325e-08 | 5.461391799312943 | 32170.0 |
| 5 | rs66696350 | 93994219 | C | T | 4.7241614353506325e-08 | 5.461391799312943 | 32170.0 |
| 5 | rs2132548 | 93996763 | T | G | 4.818322916965352e-08 | 5.457887676916075 | 32174.0 |
| 5 | rs2044911 | 93998902 | A | T | 4.818322916965352e-08 | 5.457887676916075 | 32174.0 |
| 5 | rs66483218 | 93988485 | G | A | 4.851861883658346e-08 | 5.456655572605651 | 32183.0 |
| 5 | rs12651956 | 94036583 | A | G | 4.868001267879463e-08 | 5.456065607480955 | 32146.0 |
| 5 | rs12655665 | 93992069 | C | T | 4.902116143185066e-08 | 5.4548247739043 | 32173.0 |
| 5 | rs7735659 | 93996182 | T | A | 4.9940132871793876e-08 | 5.4515234569137485 | 32174.0 |
| 6 | rs4262195 | 96929475 | T | C | 7.197157893193263e-09 | 5.786258158688043 | 32153.0 |
| 6 | rs144345632 | 96886894 | T | TA | 7.265397365586511e-09 | 5.784671947576885 | 32114.0 |
| 6 | rs11153023 | 96968525 | C | T | 8.185721475553405e-09 | 5.764588708368089 | 32165.0 |
| 6 | rs9398072 | 96911304 | T | C | 8.697501927762305e-09 | 5.754351502537402 | 32166.0 |
| 6 | rs11152952 | 96894303 | A | G | 8.705461725132107e-09 | 5.7541969521387655 | 32119.0 |
| 6 | rs11152953 | 96894305 | C | T | 8.705461725132107e-09 | 5.7541969521387655 | 32119.0 |
| 6 | rs12190520 | 96928620 | A | C | 9.019472416931042e-09 | 5.74820709855625 | 32155.0 |
| 6 | rs12201449 | 96959382 | C | T | 9.198334390044896e-09 | 5.7448852491926745 | 32165.0 |
| 6 | 6:96893385_CAG_C | 96893385 | CAG | C | 9.519722964580184e-09 | 5.739071136566513 | 32099.0 |
| 6 | rs4078038 | 96904366 | T | C | 9.702708214008913e-09 | 5.7358454971722175 | 32172.0 |
| 6 | rs2472895 | 96870387 | A | T | 1.0008580127052474e-08 | 5.730583402676022 | 32178.0 |
| 6 | rs2499813 | 96875338 | T | C | 1.0008580127052474e-08 | 5.730583402676022 | 32178.0 |
| 6 | rs4486027 | 96911886 | T | C | 1.0346439624473771e-08 | 5.7249496916168425 | 32164.0 |
| 6 | rs4839837 | 96908706 | G | A | 1.0385649866645693e-08 | 5.724307488594034 | 32175.0 |
| 6 | rs9400016 | 96940883 | G | T | 1.1079938990910542e-08 | 5.713310276925482 | 32173.0 |
| 6 | rs2472884 | 96863965 | C | T | 1.1507933232072181e-08 | 5.706860040868146 | 32172.0 |
| 6 | rs12194971 | 96911073 | T | C | 1.171060621960076e-08 | 5.703886510920026 | 32176.0 |
| 6 | rs35487471 | 96879715 | C | T | 1.1961567516206782e-08 | 5.700273057167312 | 32185.0 |
| 6 | rs72931061 | 96880285 | G | A | 1.1961567516206782e-08 | 5.700273057167312 | 32185.0 |
| 6 | rs34209750 | 96882215 | G | T | 1.1961567516206782e-08 | 5.700273057167312 | 32185.0 |
| 6 | rs13207333 | 96882254 | T | C | 1.1961567516206782e-08 | 5.700273057167312 | 32185.0 |
| 6 | rs4486019 | 96883814 | T | C | 1.1961567516206782e-08 | 5.700273057167312 | 32185.0 |
| 6 | rs67996804 | 96884387 | C | T | 1.1961567516206782e-08 | 5.700273057167312 | 32185.0 |

| | | | | | | | |
|---|---|---|---|---|---|---|---|
| 6 | rs34236778 | 96884556 | C | A | 1.1961567516206782e-08 | 5.700273057167312 | 32185.0 |
| 6 | rs11757063 | 96884886 | G | A | 1.1961567516206782e-08 | 5.700273057167312 | 32185.0 |
| 6 | rs11752584 | 96884952 | C | A | 1.1961567516206782e-08 | 5.700273057167312 | 32185.0 |
| 6 | rs11757102 | 96885073 | G | A | 1.1961567516206782e-08 | 5.700273057167312 | 32185.0 |
| 6 | rs35035182 | 96885300 | T | C | 1.1961567516206782e-08 | 5.700273057167312 | 32185.0 |
| 6 | rs35410524 | 96885405 | C | T | 1.1961567516206782e-08 | 5.700273057167312 | 32185.0 |
| 6 | rs12208884 | 96885572 | G | A | 1.1961567516206782e-08 | 5.700273057167312 | 32185.0 |
| 6 | rs13213297 | 96887569 | C | A | 1.1961567516206782e-08 | 5.700273057167312 | 32185.0 |
| 6 | rs13202091 | 96887771 | T | C | 1.1961567516206782e-08 | 5.700273057167312 | 32185.0 |
| 6 | rs140961595 | 96888323 | A | G | 1.1961567516206782e-08 | 5.700273057167312 | 32185.0 |
| 6 | rs148083617 | 96888353 | A | T | 1.1961567516206782e-08 | 5.700273057167312 | 32185.0 |
| 6 | rs12210834 | 96890407 | A | T | 1.1961567516206782e-08 | 5.700273057167312 | 32185.0 |
| 6 | rs12205655 | 96891733 | T | A | 1.1961567516206782e-08 | 5.700273057167312 | 32185.0 |
| 6 | rs12529248 | 96895377 | T | G | 1.1961567516206782e-08 | 5.700273057167312 | 32185.0 |
| 6 | rs12213426 | 96896035 | T | C | 1.1961567516206782e-08 | 5.700273057167312 | 32185.0 |
| 6 | rs12190507 | 96898265 | T | G | 1.1961567516206782e-08 | 5.700273057167312 | 32185.0 |
| 6 | rs12190508 | 96898269 | T | C | 1.1961567516206782e-08 | 5.700273057167312 | 32185.0 |
| 6 | rs34826652 | 96899085 | A | G | 1.1961567516206782e-08 | 5.700273057167312 | 32185.0 |
| 6 | rs28623445 | 96899116 | G | T | 1.1961567516206782e-08 | 5.700273057167312 | 32185.0 |
| 6 | rs11152959 | 96900272 | C | T | 1.1961567516206782e-08 | 5.700273057167312 | 32185.0 |
| 6 | rs11152960 | 96900398 | A | G | 1.1961567516206782e-08 | 5.700273057167312 | 32185.0 |
| 6 | rs12195315 | 96905711 | T | G | 1.1961567516206782e-08 | 5.700273057167312 | 32185.0 |
| 6 | rs11152968 | 96908653 | C | T | 1.1961567516206782e-08 | 5.700273057167312 | 32185.0 |
| 6 | rs11152969 | 96908712 | A | C | 1.1961567516206782e-08 | 5.700273057167312 | 32185.0 |
| 6 | rs11152970 | 96909077 | A | G | 1.1961567516206782e-08 | 5.700273057167312 | 32185.0 |
| 6 | rs13201795 | 96910471 | G | A | 1.1961567516206782e-08 | 5.700273057167312 | 32185.0 |
| 6 | rs11153071 | 97039741 | G | A | 1.2031878143044842e-08 | 5.699273900797943 | 31955.0 |
| 6 | rs2472888 | 96865501 | T | C | 1.2138579228209842e-08 | 5.697768400114338 | 32178.0 |
| 6 | rs139140529 | 96903530 | T | TTCT AAG | 1.2181069281380896e-08 | 5.69717246309208 | 32100.0 |
| 6 | rs12205728 | 96891838 | T | C | 1.2354949615121169e-08 | 5.694754596811173 | 32183.0 |
| 6 | rs12193732 | 96942609 | G | A | 1.2397830559683924e-08 | 5.6941634055785455 | 32175.0 |
| 6 | rs10447413 | 96943434 | G | C | 1.2397830559683924e-08 | 5.6941634055785455 | 32175.0 |
| 6 | rs112107820 | 96943946 | C | T | 1.2397830559683924e-08 | 5.6941634055785455 | 32175.0 |
| 6 | rs11153018 | 96956137 | G | A | 1.2397830559683924e-08 | 5.6941634055785455 | 32175.0 |
| 6 | rs12208682 | 96921307 | G | C | 1.2471364537930623e-08 | 5.693154216516754 | 32148.0 |
| 6 | rs146759159 | 96944004 | G | T | 1.248943798966697e-08 | 5.6929070595212385 | 32171.0 |
| 6 | rs6921291 | 97066242 | C | T | 1.2521483907412662e-08 | 5.692469680219324 | 32142.0 |
| 6 | rs2472889 | 96866803 | T | G | 1.2927599343506168e-08 | 5.687019112494964 | 32186.0 |
| 6 | rs12199452 | 96880703 | G | A | 1.3100215886116972e-08 | 5.68475260425986 | 32181.0 |
| 6 | rs11152951 | 96894135 | C | G | 1.3282625382310893e-08 | 5.682388828096013 | 32164.0 |
| 6 | rs12203657 | 96883285 | G | A | 1.3390986132219229e-08 | 5.6809995097169725 | 32186.0 |
| 6 | rs147506219 | 96889290 | C | A | 1.3908048410642165e-08 | 5.6745170266471865 | 32186.0 |
| 6 | rs142957763 | 96890032 | G | A | 1.3908048410642165e-08 | 5.6745170266471865 | 32186.0 |
| 6 | rs2472887 | 96865437 | A | G | 1.390814693875232e-08 | 5.674515813842371 | 32186.0 |
| 6 | rs11417499 | 96866499 | C | CT | 1.390814693875232e-08 | 5.674515813842371 | 32186.0 |
| 6 | rs2472890 | 96867646 | C | T | 1.390814693875232e-08 | 5.674515813842371 | 32186.0 |
| 6 | rs2142652 | 96868504 | A | C | 1.390814693875232e-08 | 5.674515813842371 | 32186.0 |
| 6 | rs2499809 | 96870110 | C | T | 1.390814693875232e-08 | 5.674515813842371 | 32186.0 |
| 6 | rs2472897 | 96870569 | G | T | 1.390814693875232e-08 | 5.674515813842371 | 32186.0 |
| 6 | rs2092095 | 96870864 | T | G | 1.390814693875232e-08 | 5.674515813842371 | 32186.0 |
| 6 | rs2092096 | 96870919 | T | C | 1.390814693875232e-08 | 5.674515813842371 | 32186.0 |
| 6 | rs2263192 | 96873037 | C | G | 1.390814693875232e-08 | 5.674515813842371 | 32186.0 |
| 6 | rs1116113 | 96873586 | G | C | 1.390814693875232e-08 | 5.674515813842371 | 32186.0 |
| 6 | rs2472904 | 96874962 | T | G | 1.390814693875232e-08 | 5.674515813842371 | 32186.0 |

| | | | | | | | |
|---|---|---|---|---|---|---|---|
| 6 | rs2472906 | 96875318 | G | T | 1.390814693875232e-08 | 5.674515813842371 | 32186.0 |
| 6 | rs2472908 | 96875998 | A | T | 1.390814693875232e-08 | 5.674515813842371 | 32186.0 |
| 6 | rs2472911 | 96877055 | A | G | 1.390814693875232e-08 | 5.674515813842371 | 32186.0 |
| 6 | rs2499814 | 96877245 | C | T | 1.390814693875232e-08 | 5.674515813842371 | 32186.0 |
| 6 | rs2499816 | 96878206 | C | T | 1.390814693875232e-08 | 5.674515813842371 | 32186.0 |
| 6 | rs2499817 | 96878321 | C | T | 1.390814693875232e-08 | 5.674515813842371 | 32186.0 |
| 6 | rs2472882 | 96863474 | C | T | 1.3945242081251579e-08 | 5.674059793419876 | 32180.0 |
| 6 | rs2472883 | 96863817 | C | T | 1.3945242081251579e-08 | 5.674059793419876 | 32180.0 |
| 6 | rs2499807 | 96864053 | T | G | 1.3945242081251579e-08 | 5.674059793419876 | 32180.0 |
| 6 | rs1303852 | 96864523 | C | T | 1.3945242081251579e-08 | 5.674059793419876 | 32180.0 |
| 6 | rs2179126 | 96871350 | A | G | 1.4353955417355738e-08 | 5.6691119544594955 | 32185.0 |
| 6 | rs2142653 | 96871520 | C | A | 1.4353955417355738e-08 | 5.6691119544594955 | 32185.0 |
| 6 | rs12213375 | 96902134 | A | G | 1.4527125929726536e-08 | 5.6670567264151845 | 32183.0 |
| 6 | rs75303492 | 96902999 | G | A | 1.4527125929726536e-08 | 5.6670567264151845 | 32183.0 |
| 6 | rs12208449 | 96903253 | C | T | 1.4527125929726536e-08 | 5.6670567264151845 | 32183.0 |
| 6 | rs12191653 | 96903898 | T | C | 1.4527125929726536e-08 | 5.6670567264151845 | 32183.0 |
| 6 | rs12210146 | 96903962 | C | T | 1.4527125929726536e-08 | 5.6670567264151845 | 32183.0 |
| 6 | rs4077932 | 96905144 | C | T | 1.4527125929726536e-08 | 5.6670567264151845 | 32183.0 |
| 6 | rs11755945 | 96908038 | G | C | 1.4527125929726536e-08 | 5.6670567264151845 | 32183.0 |
| 6 | rs2499804 | 96861216 | T | C | 1.4558212639347637e-08 | 5.666690303298651 | 32185.0 |
| 6 | rs2472881 | 96862562 | C | T | 1.4558212639347637e-08 | 5.666690303298651 | 32185.0 |
| 6 | rs2205755 | 96864704 | A | G | 1.4558212639347637e-08 | 5.666690303298651 | 32185.0 |
| 6 | rs2472886 | 96864876 | C | T | 1.4558212639347637e-08 | 5.666690303298651 | 32185.0 |
| 6 | rs2472893 | 96870048 | G | C | 1.4558418728947923e-08 | 5.666687876630551 | 32186.0 |
| 6 | rs2273622 | 97058567 | A | G | 1.477671313042931e-08 | 5.664136053485628 | 32150.0 |
| 6 | rs34242766 | 96872828 | C | CA | 1.5226315407558652e-08 | 5.658993688649467 | 32146.0 |
| 6 | rs9373978 | 97056255 | T | C | 1.5834026921500586e-08 | 5.652272530960173 | 32110.0 |
| 6 | rs13208422 | 96892119 | C | T | 1.725020750039938e-08 | 5.637534478157657 | 32184.0 |
| 6 | rs2472880 | 96860130 | G | A | 1.7356835917286702e-08 | 5.636472875880543 | 32183.0 |
| 6 | rs6568443 | 96944473 | T | C | 1.790265237875031e-08 | 5.631135960121127 | 32167.0 |
| 6 | rs2499801 | 96854594 | G | A | 1.8047232473831608e-08 | 5.629748746361807 | 32086.0 |
| 6 | rs760619 | 96857431 | A | G | 1.8514990807853505e-08 | 5.625333507415743 | 32088.0 |
| 6 | rs12208011 | 96991105 | G | A | 1.855808724781166e-08 | 5.624932178739016 | 32183.0 |
| 6 | rs10457159 | 96993139 | G | A | 1.855808724781166e-08 | 5.624932178739016 | 32183.0 |
| 6 | rs3734238 | 96996228 | A | T | 1.855808724781166e-08 | 5.624932178739016 | 32183.0 |
| 6 | rs3734239 | 96996244 | A | G | 1.855808724781166e-08 | 5.624932178739016 | 32183.0 |
| 6 | rs34756237 | 96854540 | T | TA | 1.9584276538476583e-08 | 5.615634042668363 | 31913.0 |
| 6 | rs6916232 | 96985959 | A | T | 1.98633142548538e-08 | 5.6131875166612994 | 32185.0 |
| 6 | rs12204342 | 97054314 | G | T | 2.0153973680185632e-08 | 5.6106743237043295 | 32142.0 |
| 6 | rs4265039 | 96878929 | T | C | 2.0804598821952385e-08 | 5.6051738887070055 | 32172.0 |
| 6 | rs4346856 | 96880270 | C | T | 2.0804598821952385e-08 | 5.6051738887070055 | 32172.0 |
| 6 | rs2064947 | 97052754 | C | T | 2.1775426464067423e-08 | 5.5972693493090775 | 32149.0 |
| 6 | rs926276 | 96868735 | C | T | 2.19305046288764e-08 | 5.596038480336651 | 32168.0 |
| 6 | rs12190397 | 97036465 | G | A | 2.236447747255332e-08 | 5.592638382776396 | 32090.0 |
| 6 | rs12212614 | 97036466 | T | G | 2.236447747255332e-08 | 5.592638382776396 | 32090.0 |
| 6 | rs12530462 | 97053514 | C | T | 2.2537978021701546e-08 | 5.5912969273839925 | 32148.0 |
| 6 | rs4343922 | 96876903 | C | T | 2.2554593152867565e-08 | 5.591168990433197 | 32177.0 |
| 6 | rs9486205 | 96877006 | C | T | 2.2554593152867565e-08 | 5.591168990433197 | 32177.0 |
| 6 | rs13206547 | 97006942 | T | C | 2.2677282321877e-08 | 5.5902271027116175 | 32173.0 |
| 6 | rs34051754 | 97007754 | A | T | 2.2677282321877e-08 | 5.5902271027116175 | 32173.0 |
| 6 | rs4839826 | 96853616 | C | T | 2.3026096257317047e-08 | 5.587576026554098 | 31993.0 |
| 6 | rs6568392 | 96861476 | G | T | 2.334158356131465e-08 | 5.585211601603699 | 32167.0 |
| 6 | rs11153064 | 97021959 | A | G | 2.3427604148388207e-08 | 5.584572300224953 | 32171.0 |
| 6 | rs3798294 | 97049587 | G | A | 2.3681224256591145e-08 | 5.582700582505454 | 32147.0 |

| | | | | | | | |
|---|---|---|---|---|---|---|---|
| 6 | rs3798289 | 97023632 | T | C | 2.373266964839396e-08 | 5.582323289810205 | 32170.0 |
| 6 | rs11153059 | 97014537 | T | A | 2.460685785323787e-08 | 5.576030658261048 | 32173.0 |
| 6 | rs11153058 | 97012746 | C | T | 2.4852172545471726e-08 | 5.574303791896439 | 32172.0 |
| 6 | rs12527021 | 97025496 | C | T | 2.4960861963575042e-08 | 5.5735439668401465 | 32168.0 |
| 6 | rs13210597 | 96969694 | C | A | 2.5270349098967107e-08 | 5.5713978666438475 | 32163.0 |
| 6 | 6:96969558_GAACCCCCAGCGCCGCGGTAC_G | 96969558 | GAACCCCCAGCGCCGCGGTAC | G | 2.546993140628335e-08 | 5.5700273751407945 | 31983.0 |
| 6 | rs74221529 | 28680690 | T | TA | 2.5578582020377243e-08 | 5.569285668093932 | 26930.0 |
| 6 | rs12214469 | 97019471 | G | T | 2.588914007725708e-08 | 5.567182371548926 | 32167.0 |
| 6 | rs12204068 | 97036201 | C | T | 2.715992078432288e-08 | 5.558823479990467 | 32152.0 |
| 6 | rs142257404 | 96984849 | C | CAT | 2.7337396308392496e-08 | 5.557686409026506 | 32184.0 |
| 6 | rs6920913 | 96874297 | A | G | 2.773218233667587e-08 | 5.555182523792388 | 32133.0 |
| 6 | rs35300731 | 96875349 | C | CG | 2.8015140473871862e-08 | 5.5534090831918 | 31950.0 |
| 6 | rs9386648 | 97035656 | T | C | 2.8240681141498203e-08 | 5.55200791068886 | 32154.0 |
| 6 | rs11153067 | 97039421 | A | T | 2.9214986240355743e-08 | 5.5460772061806205 | 32151.0 |
| 6 | rs12207570 | 96987884 | T | G | 2.9302218257178866e-08 | 5.545555611034255 | 32186.0 |
| 6 | rs2145535 | 97034715 | G | A | 3.0028283296374926e-08 | 5.541271727635776 | 32160.0 |
| 6 | rs12208556 | 96903486 | C | T | 3.010749273870184e-08 | 5.540810475747254 | 32138.0 |
| 6 | rs11756315 | 97008678 | G | A | 3.0828040605419376e-08 | 5.53666784150109 | 32171.0 |
| 6 | rs200992455 | 97062117 | ATAAT | A | 3.2025135601595215e-08 | 5.529988733165904 | 32017.0 |
| 6 | rs12197411 | 97039621 | G | A | 3.2446103467552235e-08 | 5.5276974123837705 | 32141.0 |
| 6 | rs12525695 | 97039818 | G | A | 3.2638826886894314e-08 | 5.526658029593632 | 32154.0 |
| 6 | rs9486719 | 97060124 | G | A | 3.3077140760372865e-08 | 5.524316140774461 | 32110.0 |
| 6 | rs10872018 | 97026429 | G | A | 3.4254891477899416e-08 | 5.518169522648507 | 32167.0 |
| 6 | rs12208124 | 97027942 | C | T | 3.44466263923172e-08 | 5.517188315395654 | 32165.0 |
| 6 | rs13202675 | 97028427 | C | T | 3.44466263923172e-08 | 5.517188315395654 | 32165.0 |
| 6 | rs3798296 | 97052286 | G | A | 3.5278327315430915e-08 | 5.512992498942244 | 32156.0 |
| 6 | rs4839833 | 96882069 | T | C | 3.64093221337398e-08 | 5.507438172353678 | 32174.0 |
| 6 | rs3798295 | 97052164 | C | T | 3.663348218542855e-08 | 5.506357216704617 | 32158.0 |
| 6 | rs6923212 | 97061940 | T | C | 3.702689707644587e-08 | 5.504475488604517 | 32119.0 |
| 6 | rs3906199 | 97049899 | C | A | 3.833782203423869e-08 | 5.49834228177656 | 32161.0 |
| 6 | rs6924957 | 96853579 | G | T | 3.924372742845882e-08 | 5.4942216868363225 | 32009.0 |
| 6 | rs1157316 | 96869710 | G | A | 4.113214715125233e-08 | 5.485920668072157 | 32172.0 |
| 6 | rs13208321 | 96860354 | A | G | 4.195875327310651e-08 | 5.482402804707823 | 32137.0 |
| 6 | 6:96890391_AAC_A | 96890391 | AAC | A | 4.649137291940286e-08 | 5.464232552773115 | 32089.0 |
| 6 | rs11751075 | 97065751 | G | A | 4.776367978584963e-08 | 5.459440712187559 | 32118.0 |
| 6 | rs11759769 | 97065212 | G | A | 4.843673574501173e-08 | 5.456955618924575 | 32121.0 |
| 10 | rs4880259 | 134298687 | T | C | 4.508961432972106e-17 | 8.39884379292765 | 29501.0 |
| 10 | rs11146399 | 134308479 | C | T | 5.503654932591339e-16 | 8.099828874727846 | 32033.0 |
| 10 | rs11146400 | 134308575 | C | T | 5.579365481328456e-16 | 8.098166542010222 | 32035.0 |
| 10 | rs4469802 | 134309194 | G | A | 7.732138798449122e-16 | 8.058365415682667 | 32022.0 |
| 10 | rs4880400 | 134313795 | T | C | 1.0224780358603788e-15 | 8.024130209862609 | 32097.0 |
| 10 | rs11146401 | 134313336 | C | A | 1.4602767365264618e-15 | 7.980258052331684 | 32120.0 |
| 10 | rs35974361 | 134307547 | C | G | 1.5083154621191118e-15 | 7.976262038123653 | 32031.0 |
| 10 | rs4454641 | 134307845 | C | T | 1.799713061243937e-15 | 7.954420903432577 | 32052.0 |
| 10 | rs796953214 | 134309294 | ATT | A | 1.821553447497812e-15 | 7.952927234292401 | 31784.0 |
| 10 | rs35263482 | 134313046 | G | C | 2.022029132986991e-15 | 7.9399867130618365 | 31992.0 |
| 10 | rs35133675 | 134312857 | C | T | 2.406803752990105e-15 | 7.9183513913494785 | 32186.0 |
| 10 | 10:134312221_CACAA_C | 134312221 | CACAA | C | 3.3161458194685373e-15 | 7.878394400038856 | 29589.0 |
| 10 | rs7915907 | 134316231 | G | A | 5.788851709847463e-15 | 7.808465322329078 | 31962.0 |
| 10 | rs67221163 | 134297909 | C | G | 6.552749215637474e-15 | 7.79282472369671 | 31711.0 |
| 10 | rs71481917 | 134297801 | C | T | 7.883924929787514e-15 | 7.769431363381061 | 30916.0 |
| 10 | rs57472801 | 134322756 | A | G | 9.539882660931271e-15 | 7.745243528791251 | 31745.0 |
| 10 | rs7897053 | 134316342 | C | T | 1.7793118674599962e-14 | 7.665647546168928 | 31956.0 |

| | | | | | | | |
|---|---|---|---|---|---|---|---|
| 10 | rs7917322 | 134306694 | A | C | 1.8653002955682653e-14 | 7.659588324802858 | 31090.0 |
| 10 | rs4356146 | 134324520 | G | A | 2.093333047833664e-14 | 7.644761295339781 | 31724.0 |
| 10 | rs11187838 | 96038686 | G | A | 4.2917313033974e-14 | 7.551833440068696 | 32147.0 |
| 10 | rs2274224 | 96039597 | G | C | 5.049986986642375e-14 | 7.530620223036226 | 32186.0 |
| 10 | rs7084359 | 134297562 | A | C | 5.877560539023701e-14 | 7.51078109325326 | 31837.0 |
| 10 | rs57866767 | 96023077 | T | C | 6.855622855055232e-14 | 7.490606197873143 | 32139.0 |
| 10 | rs3891783 | 96015793 | C | G | 6.894184578704718e-14 | 7.489870037653303 | 32139.0 |
| 10 | rs10786156 | 96014622 | C | G | 7.468100049579422e-14 | 7.479367788191028 | 32139.0 |
| 10 | rs7083220 | 134284485 | G | A | 8.010723377127049e-14 | 7.470143795428133 | 32155.0 |
| 10 | rs9645539 | 134282867 | G | T | 9.598010224717971e-14 | 7.446319635920517 | 31875.0 |
| 10 | rs11819412 | 134297803 | G | C | 9.847909623093286e-14 | 7.442926218060664 | 31133.0 |
| 10 | rs10735665 | 134308936 | G | A | 1.043280898859151e-13 | 7.435303572358428 | 32018.0 |
| 10 | rs11146402 | 134323354 | C | G | 1.1676556619761485e-13 | 7.42040178792245 | 31717.0 |
| 10 | 10:96026184_CA_C | 96026184 | CA | C | 1.2073208668189555e-13 | 7.4159763034235775 | 31917.0 |
| 10 | rs12762160 | 134330662 | C | T | 1.2559451019952514e-13 | 7.410742211508107 | 31313.0 |
| 10 | rs11379033 | 134325396 | C | CA | 1.478293105137883e-13 | 7.389096989850744 | 31120.0 |
| 10 | rs11596664 | 134280157 | C | T | 1.482216672420236e-13 | 7.38874450194884 | 31747.0 |
| 10 | rs10747062 | 134285515 | C | T | 1.485203015482186e-13 | 7.3884768269320675 | 32132.0 |
| 10 | rs10747061 | 134285403 | G | A | 2.1826563960480998e-13 | 7.337103612380594 | 32156.0 |
| 10 | rs10781576 | 134285017 | T | C | 2.19278175959502e-13 | 7.336483900656723 | 32186.0 |
| 10 | rs11146393 | 134299369 | G | C | 2.405872474501524e-13 | 7.324055423314602 | 31763.0 |
| 10 | rs61861415 | 134328497 | T | C | 2.5388030908469136e-13 | 7.3168389114944015 | 31385.0 |
| 10 | rs200898494 | 134299394 | A | AG | 3.1172050039237527e-13 | 7.289234747996606 | 31465.0 |
| 10 | rs4880397 | 134309321 | C | G | 3.11869145734844e-13 | 7.289170511538463 | 32036.0 |
| 10 | rs10870313 | 134310920 | A | G | 3.156833903200461e-13 | 7.287532405802485 | 31965.0 |
| 10 | rs11814088 | 134298002 | A | C | 3.1840617023815953e-13 | 7.286374900040142 | 31820.0 |
| 10 | rs10870314 | 134310923 | T | G | 3.765233955795692e-13 | 7.263742888352753 | 31955.0 |
| 10 | rs7080018 | 134301505 | G | A | 5.410387895424326e-13 | 7.214571989497416 | 31929.0 |
| 10 | rs4492738 | 134300136 | T | C | 5.933744352233561e-13 | 7.201995978071093 | 31939.0 |
| 10 | rs4592360 | 134300069 | A | G | 5.963998680296833e-13 | 7.2013026833470475 | 31937.0 |
| 10 | rs4309079 | 134300091 | C | T | 6.260333474957793e-13 | 7.194688911191922 | 31934.0 |
| 10 | rs7903563 | 134300816 | G | A | 7.435485968066305e-13 | 7.1711785352399655 | 31943.0 |
| 10 | rs4880396 | 134298833 | T | C | 7.460960526637285e-13 | 7.170710360150287 | 31766.0 |
| 10 | rs4501914 | 134308062 | C | T | 7.574386324413606e-13 | 7.168644673091168 | 32186.0 |
| 10 | rs11146395 | 134299444 | T | C | 7.825654384829695e-13 | 7.1641747216956935 | 31752.0 |
| 10 | rs4293055 | 134323744 | G | A | 8.019254548016124e-13 | 7.1608257231068215 | 31495.0 |
| 10 | rs543302184 | 96009182 | T | TA | 1.324066465091119e-12 | 7.091770832009837 | 30691.0 |
| 10 | rs7080472 | 96012950 | G | T | 1.484423407411613e-12 | 7.075936955168742 | 31940.0 |
| 10 | rs7096828 | 134301541 | C | T | 1.5309795102694828e-12 | 7.071653836089679 | 31966.0 |
| 10 | rs7914305 | 134312761 | C | T | 1.652873068056089e-12 | 7.061017893580643 | 31964.0 |
| 10 | rs61861416 | 134333189 | T | A | 2.0093864012123997e-12 | 7.033830894274732 | 30952.0 |
| 10 | rs11596632 | 134279989 | C | A | 2.0632144190394627e-12 | 7.0301433586987425 | 31517.0 |
| 10 | rs7918535 | 134300351 | A | G | 2.1668849492925253e-12 | 7.023299840532359 | 31951.0 |
| 10 | rs7897891 | 134284005 | T | C | 2.9129910699348014e-12 | 6.981859149156841 | 31712.0 |
| 10 | 10:134303087_AG_A | 134303087 | AG | A | 2.9544155561831077e-12 | 6.979875597084392 | 31783.0 |
| 10 | rs12572574 | 134281065 | C | T | 3.132384377583428e-12 | 6.9716529838953205 | 31599.0 |
| 10 | rs7917077 | 134305875 | T | A | 3.742350590645925e-12 | 6.946584784492347 | 31441.0 |
| 10 | rs7914354 | 134334476 | A | G | 3.796200835261439e-12 | 6.944568039094263 | 30876.0 |
| 10 | rs4593926 | 134281537 | A | G | 4.045781625797275e-12 | 6.935572911887697 | 31491.0 |
| 10 | rs34102287 | 134331173 | C | T | 4.702499223453563e-12 | 6.914278684473759 | 30935.0 |
| 10 | rs10870316 | 134321788 | G | A | 4.8995817747026744e-12 | 6.908455630628014 | 31837.0 |
| 10 | rs10870317 | 134321883 | G | C | 5.3143179118716205e-12 | 6.896917082362893 | 31833.0 |
| 10 | rs10747060 | 134285007 | A | G | 5.426028775359202e-12 | 6.893959998599807 | 31841.0 |
| 10 | rs11591553 | 134323564 | G | A | 5.5975357092005364e-12 | 6.8895342248941205 | 31722.0 |

| | | | | | | | |
|---|---|---|---|---|---|---|---|
| 10 | rs7090098 | 134320629 | G | A | 5.971001940904423e-12 | 6.880339658270646 | 31864.0 |
| 10 | rs71503747 | 134304818 | C | T | 5.997122655095898e-12 | 6.879717837933923 | 30316.0 |
| 10 | rs4497325 | 134302745 | G | A | 6.353751246874942e-12 | 6.871483790556253 | 32050.0 |
| 10 | rs5025116 | 134320913 | C | T | 6.50461582493483e-12 | 6.868136083827987 | 31854.0 |
| 10 | rs61861413 | 134326732 | G | A | 7.367172859675429e-12 | 6.850345715835326 | 31664.0 |
| 10 | rs10735666 | 134322595 | C | A | 8.071625049381277e-12 | 6.8372704950253045 | 31637.0 |
| 10 | rs11146397 | 134303122 | T | C | 8.307068403221574e-12 | 6.833148843921718 | 32143.0 |
| 10 | rs10747063 | 134317884 | C | T | 1.0237974201001445e-11 | 6.803116565445973 | 31848.0 |
| 10 | rs7086785 | 134316795 | T | C | 1.1307133896980027e-11 | 6.78879836458131 | 31909.0 |
| 10 | rs10870315 | 134316622 | A | G | 1.1415220006620805e-11 | 6.787425462792664 | 31889.0 |
| 10 | rs139032047 | 134304807 | C | A | 1.3152606812709075e-11 | 6.766949379012942 | 29973.0 |
| 10 | rs71503746 | 134304802 | G | A | 1.3219783080215665e-11 | 6.766211963506348 | 29797.0 |
| 10 | rs10781577 | 134325083 | G | T | 1.3418596867656795e-11 | 6.764050832741418 | 31588.0 |
| 10 | rs11146398 | 134303274 | T | C | 1.3469926177200972e-11 | 6.763497970450421 | 32186.0 |
| 10 | rs7894289 | 134305663 | G | A | 2.275826163692291e-11 | 6.687134553866423 | 30701.0 |
| 10 | rs7093232 | 134303666 | G | A | 2.530307144250132e-11 | 6.671599266352523 | 32037.0 |
| 10 | rs79361141 | 134304808 | C | A | 2.5735232153973962e-11 | 6.669113986323622 | 30417.0 |
| 10 | rs10747064 | 134322499 | C | T | 3.580626510439153e-11 | 6.62046831942667 | 31669.0 |
| 10 | rs11187844 | 96056629 | C | A | 7.449547612740312e-11 | 6.511315526300487 | 31901.0 |
| 10 | rs7907962 | 134287486 | A | G | 7.866417789894608e-11 | 6.503133018285553 | 30475.0 |
| 10 | rs71503745 | 134304757 | C | A | 8.617597876540119e-11 | 6.489405080311924 | 29932.0 |
| 10 | rs35364222 | 134296800 | T | C | 9.152344500176895e-11 | 6.480327983588389 | 31866.0 |
| 10 | rs7904728 | 134295644 | G | A | 1.385228174250261e-10 | 6.417508291801415 | 32120.0 |
| 10 | rs4344410 | 134294911 | C | A | 4.89945396730622e-10 | 6.222291581631027 | 32016.0 |
| 10 | rs7923337 | 134295750 | A | T | 5.722881646151891e-10 | 6.197879803253204 | 32133.0 |
| 10 | rs10747068 | 134335157 | A | G | 8.130444520354161e-10 | 6.142359772913839 | 28423.0 |
| 10 | rs7095973 | 134324510 | T | C | 8.393782519584108e-10 | 6.137296144650871 | 31678.0 |
| 10 | rs753724 | 96051417 | G | T | 8.474703943185124e-10 | 6.13577123017936 | 32180.0 |
| 10 | rs11187840 | 96050351 | A | G | 8.991669143104533e-10 | 6.126352131568465 | 32163.0 |
| 10 | rs12220091 | 96053689 | C | T | 1.0664123647871142e-09 | 6.099139106994366 | 32182.0 |
| 10 | rs75017201 | 96055152 | C | T | 1.0664123647871142e-09 | 6.099139106994366 | 32182.0 |
| 10 | rs3740365 | 96053239 | T | A | 1.0974119598630194e-09 | 6.094556606071854 | 32183.0 |
| 10 | rs769015184 | 96049708 | AC | A | 1.3089632122842845e-09 | 6.066292902156383 | 32140.0 |
| 10 | rs3781266 | 96052747 | A | G | 1.3681948132374006e-09 | 6.05917734265741 | 32182.0 |
| 10 | rs11187842 | 96052511 | C | T | 2.1051480825203594e-09 | 5.989478615108893 | 32186.0 |
| 10 | rs4485028 | 134286588 | A | G | 2.207540188100426e-09 | 5.9817491836361825 | 30571.0 |
| 10 | rs4523616 | 134286571 | T | C | 2.348132108810336e-09 | 5.97168664910004 | 30612.0 |
| 10 | rs4509684 | 134286579 | C | T | 2.3661538095337906e-09 | 5.97043945428576 | 30614.0 |
| 10 | rs4486544 | 134294345 | T | C | 2.742066681100417e-09 | 5.946338585361418 | 31672.0 |
| 10 | rs11146404 | 134326869 | A | G | 2.870438404284808e-09 | 5.938841139382259 | 31523.0 |
| 10 | rs4412685 | 134294337 | A | G | 2.886947059474323e-09 | 5.937900754646733 | 31669.0 |
| 10 | rs3740360 | 96025491 | A | C | 3.0584526883104308e-09 | 5.928429715535812 | 32179.0 |
| 10 | rs17109875 | 96026575 | T | C | 3.0584526883104308e-09 | 5.928429715535812 | 32179.0 |
| 10 | rs12360525 | 134289214 | A | T | 3.133292905811434e-09 | 5.924457887070626 | 31503.0 |
| 10 | rs760730420 | 134288420 | ATTAAG | A | 3.32579742244662e-09 | 5.9146511312185615 | 31180.0 |
| 10 | rs11187837 | 96035980 | T | C | 3.5910208638460205e-09 | 5.9020087628729465 | 32186.0 |
| 10 | rs10747067 | 134332633 | C | G | 6.378609435107791e-09 | 5.806516224210881 | 31006.0 |
| 10 | rs4576738 | 134294398 | G | A | 6.393412599686308e-09 | 5.806127912051996 | 31548.0 |
| 10 | rs2077218 | 96071561 | G | A | 7.2919590692838575e-09 | 5.784058440926818 | 31871.0 |
| 10 | rs11187845 | 96060198 | C | A | 7.771044273502751e-09 | 5.773350746165256 | 32182.0 |
| 10 | rs11146388 | 134288095 | G | A | 7.924842405929412e-09 | 5.770049130831633 | 31467.0 |
| 10 | rs200197176 | 96055963 | C | CA | 7.951285316975931e-09 | 5.769487759590603 | 32010.0 |
| 10 | rs10781575 | 134280542 | C | T | 8.547751317426325e-09 | 5.7572850348932265 | 30144.0 |
| 10 | rs11187847 | 96063440 | C | G | 8.796288004349406e-09 | 5.7524430980208985 | 32165.0 |

| | | | | | | | |
|---|---|---|---|---|---|---|---|
| 10 | rs12217792 | 96062386 | T | C | 8.922820878887288e-09 | 5.750028874857671 | 32173.0 |
| 10 | rs11146389 | 134290675 | T | G | 9.792243886330027e-09 | 5.734288659530687 | 32186.0 |
| 10 | rs4242719 | 134290895 | T | C | 9.959870496461384e-09 | 5.731410829458055 | 32180.0 |
| 10 | rs372868680 | 134304572 | G | A | 1.0336874123084594e-08 | 5.725106718615635 | 26110.0 |
| 10 | rs4880382 | 134290769 | A | G | 1.0380865178548989e-08 | 5.724385728018667 | 32184.0 |
| 10 | rs35908501 | 134290607 | C | T | 1.0518954378181126e-08 | 5.7221416554067055 | 32180.0 |
| 10 | rs10870311 | 134290526 | C | A | 1.1422431492958501e-08 | 5.708129777089063 | 32175.0 |
| 10 | rs10870312 | 134290544 | C | A | 1.1453698624559652e-08 | 5.707664379311956 | 32176.0 |
| 10 | rs10870310 | 134290153 | G | A | 1.2125665747339857e-08 | 5.6979499176098365 | 32154.0 |
| 10 | rs10747058 | 134276427 | G | T | 1.2153216817882458e-08 | 5.697562874544871 | 31302.0 |
| 10 | rs4880390 | 134296740 | T | G | 1.2898808534548391e-08 | 5.6874000034344565 | 31928.0 |
| 10 | rs11816277 | 134296796 | A | G | 1.2952465406306794e-08 | 5.686690807100817 | 31938.0 |
| 10 | rs7898918 | 134276762 | G | A | 1.3651981715908724e-08 | 5.6776975548434745 | 31568.0 |
| 10 | rs4880258 | 134293640 | G | A | 1.3769274930651107e-08 | 5.6762335604193686 | 31927.0 |
| 10 | rs56094089 | 134289658 | T | C | 1.4654100512552693e-08 | 5.665564829395205 | 31787.0 |
| 10 | rs7096883 | 96069149 | G | A | 1.4776857383835465e-08 | 5.664134379326696 | 32171.0 |
| 10 | rs4880385 | 134296241 | C | T | 1.4823078792702371e-08 | 5.663598764240249 | 32186.0 |
| 10 | rs4880383 | 134295388 | T | G | 1.6702462516060403e-08 | 5.643090363620527 | 32150.0 |
| 10 | rs4880381 | 134289543 | A | T | 1.9049440232855808e-08 | 5.620419441253746 | 31764.0 |
| 10 | rs12771233 | 134291950 | C | T | 1.927591563531653e-08 | 5.618377347359412 | 31980.0 |
| 10 | rs4880384 | 134295412 | A | G | 2.1428391460668057e-08 | 5.600054912798445 | 32163.0 |
| 10 | rs12358548 | 134273093 | T | A | 2.7301359692661493e-08 | 5.557916711914437 | 32083.0 |
| 10 | rs10787738 | 118777371 | C | T | 3.4236147452205936e-08 | 5.518265731247902 | 30800.0 |
| 11 | rs11218557 | 122099839 | T | C | 1.2464323864899471e-08 | 5.693250592916878 | 31941.0 |
| 11 | rs10892820 | 122120972 | T | G | 1.38594196355121e-08 | 5.675116630254277 | 31716.0 |
| 11 | rs4127499 | 122176383 | G | A | 1.965344634739912e-08 | 5.615024440812554 | 32186.0 |
| 11 | rs10892823 | 122128720 | T | G | 2.4543313926902302e-08 | 5.5764806933890165 | 31750.0 |
| 11 | rs17337676 | 122132835 | G | A | 2.6847885743111573e-08 | 5.560840257902767 | 31754.0 |
| 11 | rs10750202 | 122133372 | A | G | 3.9934397271703946e-08 | 5.491141618547697 | 31756.0 |
| 11 | rs10892822 | 122124710 | C | T | 4.686732286065681e-08 | 5.462803525844152 | 31759.0 |
| 14 | rs186347 | 59072226 | G | T | 2.088812305545398e-11 | 6.6996766444809115 | 31582.0 |
| 14 | rs160459 | 59074136 | A | C | 3.009831399689939e-11 | 6.6460890809515245 | 32186.0 |
| 14 | rs160458 | 59074878 | T | C | 7.369881980320526e-11 | 6.5129300836411925 | 31575.0 |
| 14 | rs160460 | 59072964 | A | G | 1.8501390581295992e-10 | 6.373291369079907 | 31654.0 |
| 14 | rs10782438 | 59072144 | T | C | 1.139520201158146e-09 | 6.088530116890116 | 31609.0 |
| 14 | rs468213 | 59071100 | C | G | 2.1961397723828772e-09 | 5.982592306756784 | 31522.0 |
| 14 | rs170239 | 59064739 | C | T | 2.4584735860110557e-09 | 5.964192110128144 | 31447.0 |
| 14 | rs149142 | 59071725 | T | C | 2.880563469249779e-09 | 5.938263761505443 | 31601.0 |
| 14 | rs221326 | 59069053 | G | T | 6.285716910176658e-09 | 5.808973146903211 | 31543.0 |
| 14 | rs183400 | 59064730 | C | T | 4.6060415643556655e-08 | 5.465884514618648 | 31430.0 |
| 15 | rs1440802 | 39635124 | T | C | 1.003264426053347e-31 | 11.720291283567304 | 32084.0 |
| 15 | rs530653020 | 39633074 | T | TA | 4.089044471473289e-31 | 11.60066166457904 | 32111.0 |
| 15 | rs5812099 | 39633877 | A | AG | 5.030460954055212e-31 | 11.582917079319868 | 32142.0 |
| 15 | rs17694988 | 39636227 | T | C | 5.0592061130992065e-31 | 11.582428729203016 | 32107.0 |
| 15 | rs71471500 | 39632013 | C | G | 7.394326587208771e-31 | 11.549857115153193 | 32137.0 |
| 15 | rs7182018 | 39632269 | A | G | 7.395323097815078e-31 | 11.549845533223973 | 32138.0 |
| 15 | rs1080066 | 39634222 | A | G | 1.6492128982617422e-30 | 11.480709647945735 | 32186.0 |
| 15 | rs2033939 | 39633904 | G | A | 1.8508740402272843e-30 | 11.470731791569252 | 32152.0 |
| 15 | rs11639391 | 39631771 | C | T | 3.915485110331933e-30 | 11.405714632088108 | 32132.0 |
| 15 | rs4924345 | 39639898 | A | C | 3.761294813836875e-29 | 11.207145251213028 | 31998.0 |
| 15 | rs4924346 | 39639992 | G | A | 3.8360538262484317e-29 | 11.205402673043961 | 31978.0 |
| 15 | 15:39640223_TTAAA_T | 39640223 | TTAAA | T | 5.480639751552537e-28 | 10.967432850121407 | 31741.0 |
| 15 | rs550614297 | 39640378 | G | GA | 8.798646845869065e-28 | 10.924538566330936 | 31531.0 |
| 15 | rs28520337 | 39647894 | T | C | 5.317029944391902e-24 | 10.103748107662494 | 31643.0 |

| | | | | | | | |
|---|---|---|---|---|---|---|---|
| 15 | rs574500639 | 39632734 | T | TAC | 2.762888912491769e-23 | 9.940910935884553 | 31822.0 |
| 15 | rs34680120 | 39664000 | C | A | 2.1600033993622744e-18 | 8.748608127196812 | 32140.0 |
| 15 | rs1837636 | 39661779 | C | T | 1.0057181932250956e-17 | 8.573287725237199 | 32186.0 |
| 15 | rs4566132 | 39661138 | G | A | 2.3385374914593398e-17 | 8.475608598799981 | 32159.0 |
| 15 | rs4923822 | 39662277 | A | G | 1.03855342042465e-15 | 8.022214778410742 | 32022.0 |
| 15 | rs11629938 | 39626935 | G | A | 1.610559054323356e-14 | 7.678425397017933 | 32071.0 |
| 15 | rs10520128 | 39627146 | G | C | 5.2713275909708294e-14 | 7.525017469153091 | 32066.0 |
| 15 | rs12591944 | 39627982 | A | C | 5.4820024315590295e-14 | 7.519895540148804 | 31996.0 |
| 15 | rs8025239 | 39623588 | G | A | 7.264782978334865e-14 | 7.482994668635992 | 32062.0 |
| 15 | rs12595360 | 39625987 | C | G | 1.0022316550340508e-13 | 7.440607703164034 | 31944.0 |
| 15 | 15:39625576_ACT_A | 39625576 | ACT | A | 1.2943951374925129e-13 | 7.406742428061791 | 31999.0 |
| 15 | rs72722978 | 39628675 | C | T | 1.9539972300765807e-13 | 7.351906333476349 | 31910.0 |
| 15 | rs4702 | 91426560 | G | A | 3.7705468908363357e-13 | 7.263552251032982 | 32186.0 |
| 15 | rs773225188 | 39631994 | AAGAG | A | 6.594332253670534e-13 | 7.187593335651549 | 31403.0 |
| 15 | rs8032326 | 39619456 | C | A | 5.0360107939395915e-12 | 6.904557636537796 | 32186.0 |
| 15 | rs16968876 | 39636877 | C | T | 8.97969466977613e-12 | 6.821975559301237 | 31917.0 |
| 15 | rs6224 | 91423543 | G | T | 4.886248673375258e-11 | 6.574361197508928 | 31804.0 |
| 15 | rs11372849 | 91419432 | T | TC | 5.2979422210413684e-11 | 6.562313106986054 | 31414.0 |
| 15 | rs16968890 | 39638991 | G | C | 1.0066687469709907e-10 | 6.465946225660579 | 31948.0 |
| 15 | rs7183988 | 91428589 | T | G | 1.7658202163711128e-10 | 6.380438160280219 | 31955.0 |
| 15 | rs17514846 | 91416550 | C | A | 2.830281105954788e-10 | 6.30780082301566 | 31337.0 |
| 15 | rs2071382 | 91428197 | T | C | 3.276211975652744e-10 | 6.285110205864173 | 31970.0 |
| 15 | rs7177338 | 91428636 | G | A | 3.312961310757099e-10 | 6.283377100366999 | 31950.0 |
| 15 | rs1894401 | 91429042 | G | A | 3.503889036264062e-10 | 6.2746645453339776 | 31981.0 |
| 15 | rs8039305 | 91422543 | T | C | 5.634662823476524e-10 | 6.200325150671808 | 31771.0 |
| 15 | rs11539637 | 91428290 | C | T | 6.233674012213628e-10 | 6.184406010904782 | 31943.0 |
| 15 | rs8033007 | 39619661 | G | C | 7.09354584270465e-10 | 6.163987693333132 | 32158.0 |
| 15 | rs978740 | 39655920 | C | T | 7.122082397681491e-10 | 6.163352267873036 | 31831.0 |
| 15 | rs56113415 | 39621418 | G | A | 5.450276098992282e-09 | 5.832806501939312 | 32117.0 |
| 15 | rs11638016 | 39621773 | T | C | 5.732272021074552e-09 | 5.8243866704669145 | 32072.0 |
| 15 | rs3910644 | 39648234 | C | T | 7.87810735134866e-09 | 5.7710457641289965 | 31154.0 |
| 17 | rs62073094 | 44316219 | C | T | 2.13287849079083e-16 | 8.2143661529309 | 30258.0 |
| 17 | rs28434724 | 44312278 | A | G | 3.417830122355649e-16 | 8.157587303086428 | 30400.0 |
| 17 | 17:44270659_G_A | 44270659 | G | A | 5.3690367959712e-16 | 8.102841047469974 | 31162.0 |
| 17 | rs9908639 | 44200192 | A | G | 7.269001835554949e-16 | 8.065913914756733 | 31520.0 |
| 17 | rs146895044 | 44166992 | A | AAAA TATG | 9.690427459411694e-16 | 8.030717476095408 | 31603.0 |
| 17 | 17:44266982_T_C | 44266982 | T | C | 1.378355579070789e-15 | 7.987381037311898 | 31307.0 |
| 17 | rs2696461 | 44285844 | C | T | 4.1734793872419705e-15 | 7.8496055100203215 | 30674.0 |
| 17 | rs56807059 | 44189270 | A | C | 4.651963238220654e-15 | 7.835980972849088 | 31590.0 |
| 17 | rs11655181 | 44332093 | G | A | 4.7281277156267305e-15 | 7.833940453520714 | 30363.0 |
| 17 | rs2016034 | 44189598 | A | G | 5.448534290102809e-15 | 7.816099428898584 | 31452.0 |
| 17 | rs8068300 | 44202690 | T | C | 5.635245115901575e-15 | 7.811854818021024 | 31563.0 |
| 17 | 17:44224272_G_A | 44224272 | G | A | 5.841373000630384e-15 | 7.807326664453987 | 31551.0 |
| 17 | rs2732634 | 44319047 | A | G | 6.12657229696107e-15 | 7.801314276785388 | 30344.0 |
| 17 | 17:44222424_G_C | 44222424 | G | C | 6.883752693979011e-15 | 7.786598008032664 | 31556.0 |
| 17 | rs62073102 | 44319735 | C | G | 7.761565907087313e-15 | 7.771412545015654 | 30253.0 |
| 17 | rs8070420 | 44185479 | G | T | 8.077938286891876e-15 | 7.76635118980803 | 31610.0 |
| 17 | rs60521849 | 44172812 | G | A | 8.438923542335159e-15 | 7.760809138525095 | 31613.0 |
| 17 | rs2158258 | 44163809 | A | C | 9.194383518591518e-15 | 7.749929148398963 | 32174.0 |
| 17 | rs11079732 | 44161581 | T | C | 1.1688228860338078e-14 | 7.719395950440285 | 32174.0 |
| 17 | rs8065825 | 44175489 | A | G | 1.1700608596082432e-14 | 7.719261008151095 | 31614.0 |
| 17 | rs8064986 | 44119367 | A | G | 1.1816714426070542e-14 | 7.718002219102359 | 32132.0 |
| 17 | rs11652924 | 44182267 | T | C | 1.3278333333467694e-14 | 7.70312021510791 | 31588.0 |
| 17 | rs11654640 | 44162230 | G | A | 1.3310946129784842e-14 | 7.7028068721159295 | 32186.0 |

| | | | | | | | |
|---|---|---|---|---|---|---|---|
| 17 | rs6503455 | 44140748 | C | T | 1.3783520164396154e-14 | 7.6983492636561985 | 32079.0 |
| 17 | rs9904766 | 44186952 | T | C | 1.3982135245140943e-14 | 7.696520523128266 | 31584.0 |
| 17 | rs72836329 | 44161763 | C | T | 1.4045405462530557e-14 | 7.695943329908069 | 32133.0 |
| 17 | rs7521 | 44105395 | A | G | 1.4350380161318967e-14 | 7.693196551425887 | 31932.0 |
| 17 | rs7220988 | 44109474 | G | A | 1.4406409059492114e-14 | 7.692698174437496 | 32186.0 |
| 17 | rs35173354 | 44212633 | T | TA | 1.448100275080826e-14 | 7.69203761505191 | 31107.0 |
| 17 | rs143437112 | 44222045 | T | TA | 1.4776354172690798e-14 | 7.689454612264255 | 31219.0 |
| 17 | rs113564729 | 44179095 | G | A | 1.888184334855864e-14 | 7.6580220776557075 | 31591.0 |
| 17 | 17:44329580_AT_A | 44329580 | AT | A | 2.329237985971001e-14 | 7.6310087884511155 | 29428.0 |
| 17 | rs113042517 | 44139453 | G | A | 2.3648050221099208e-14 | 7.62905512938946 | 32138.0 |
| 17 | 17:44291980_G_A | 44291980 | G | A | 2.7589075071189735e-14 | 7.609156391840178 | 30969.0 |
| 17 | rs11654745 | 44131096 | G | A | 2.8072808355485715e-14 | 7.606909340756143 | 32146.0 |
| 17 | rs7213077 | 44119051 | T | C | 2.946716625002523e-14 | 7.600639175764982 | 32169.0 |
| 17 | rs8078067 | 44125466 | G | A | 3.0864487649235227e-14 | 7.594641853734182 | 32145.0 |
| 17 | 17:44236005_C_T | 44236005 | C | T | 3.1716502443640664e-14 | 7.5911147051925125 | 31436.0 |
| 17 | rs2532231 | 44274474 | C | T | 3.1841021524474027e-14 | 7.590607044648504 | 31058.0 |
| 17 | rs10514899 | 44119987 | C | G | 3.2417572680618755e-14 | 7.588281628652442 | 32155.0 |
| 17 | 17:44221843_A_G | 44221843 | A | G | 3.701082879538306e-14 | 7.571089028173356 | 31509.0 |
| 17 | rs62063662 | 44119840 | T | C | 4.162144509646456e-14 | 7.55582459292876 | 32145.0 |
| 17 | rs4792827 | 44131305 | T | C | 4.525871664883928e-14 | 7.544913681717287 | 32186.0 |
| 17 | rs16940845 | 44116843 | C | A | 5.54497518594104e-14 | 7.518402078825425 | 32186.0 |
| 17 | 17:44235827_C_A | 44235827 | C | A | 6.414129976668959e-14 | 7.4993372625050405 | 31424.0 |
| 17 | 17:44262950_G_A | 44262950 | G | A | 7.905298590330521e-14 | 7.471886826823125 | 31063.0 |
| 17 | rs7206949 | 44247017 | C | A | 9.121445721906327e-14 | 7.453038779941987 | 31433.0 |
| 17 | rs62071628 | 44356058 | G | A | 9.372049721569844e-14 | 7.449463594156009 | 29940.0 |
| 17 | rs916896 | 44098797 | C | T | 9.501552202862083e-14 | 7.44765273390392 | 31873.0 |
| 17 | rs2732610 | 44365490 | C | T | 1.4632985329522965e-13 | 7.39045260467646 | 28524.0 |
| 17 | 17:44286384_C_A | 44286384 | C | A | 1.5677689447218874e-13 | 7.381278310002557 | 30912.0 |
| 17 | rs28787973 | 44084349 | T | A | 1.6261729526153812e-13 | 7.376407892845621 | 31447.0 |
| 17 | rs28753217 | 44084350 | A | T | 1.6261729526153812e-13 | 7.376407892845621 | 31447.0 |
| 17 | 17:44270181_G_A | 44270181 | G | A | 1.772164883371061e-13 | 7.364947581948904 | 31009.0 |
| 17 | rs7225002 | 44189067 | A | G | 2.0855535071878594e-13 | 7.343194343001845 | 32016.0 |
| 17 | rs4988900 | 44342378 | G | A | 2.1510463799243e-13 | 7.3390566005727855 | 30044.0 |
| 17 | rs41543317 | 44087500 | A | G | 2.72179029812221e-13 | 7.307489774832258 | 32186.0 |
| 17 | rs11654982 | 44284600 | G | A | 2.8773910537242517e-13 | 7.300013383248197 | 30935.0 |
| 17 | rs56087321 | 44084403 | G | C | 8.070230228141074e-13 | 7.15995711665839 | 31994.0 |
| 17 | rs4510068 | 44184828 | G | T | 9.779838480860608e-13 | 7.1335702129148535 | 31147.0 |
| 17 | 17:44265839_C_T | 44265839 | C | T | 1.1781176311389222e-12 | 7.107911847653782 | 31488.0 |
| 17 | 17:44289832_A_G | 44289832 | A | G | 1.708862261797252e-12 | 7.056388022347883 | 31415.0 |
| 17 | rs2532351 | 44336997 | C | G | 3.341428429409897e-12 | 6.962560574598143 | 29681.0 |
| 17 | rs5820605 | 44102682 | C | CT | 4.549285068847872e-12 | 6.9189733248548215 | 31797.0 |
| 17 | rs556046991 | 44212908 | AT | A | 2.5284615680573183e-11 | 6.671706325089718 | 30379.0 |
| 17 | 17:44776768_GAC_G | 44776768 | GAC | G | 2.7670825585399068e-11 | 6.6584617624473985 | 27910.0 |
| 17 | rs7225145 | 44363740 | C | A | 1.0468120415860143e-10 | 6.4600314778510475 | 27604.0 |
| 17 | rs12946693 | 44084008 | T | C | 3.5501898495868213e-10 | 6.272621601170705 | 30782.0 |
| 17 | rs141698966 | 44113227 | T | TG | 9.486148463518452e-10 | 6.1178244482294035 | 31241.0 |
| 17 | 17:43850935_CA_C | 43850935 | CA | C | 1.7293561995738133e-09 | 6.02138055476673 | 31964.0 |
| 17 | 17:43975360_ATTACTGTATAAGCTGCTTATAT_A | 43975360 | ATTACTGTATAAGCTGCTTATAT | A | 1.817261697786532e-09 | 6.013351708151768 | 31537.0 |
| 17 | rs201474071 | 44113197 | G | GA | 2.063508245962949e-09 | 5.992727216507348 | 31754.0 |
| 17 | rs552290972 | 43947883 | T | G | 2.345121521146673e-09 | 5.971895905764653 | 30687.0 |
| 17 | 17:44125277_CT_C | 44125277 | CT | C | 2.4498426228045372e-09 | 5.964766404918502 | 31244.0 |
| 17 | rs3874943 | 44572068 | A | G | 2.5195362927716274e-09 | 5.960184288844944 | 27960.0 |
| 17 | rs55767267 | 43960325 | T | C | 2.9062622168003382e-09 | 5.936807127465631 | 31886.0 |

| | | | | | | | |
|---|---|---|---|---|---|---|---|
| 17 | rs56127309 | 43960341 | G | A | 2.9062622168003382e-09 | 5.936807127465631 | 31886.0 |
| 17 | rs56365901 | 43960323 | A | G | 3.363145642852868e-09 | 5.9128126890820525 | 31885.0 |
| 17 | rs113758975 | 44207523 | T | A | 3.627707827590935e-09 | 5.9003320268691155 | 31745.0 |
| 17 | 17:44238616_TAAAAAAAA_T | 44238616 | TAAAAAAAA | T | 3.763144394135963e-09 | 5.894281823598792 | 30703.0 |
| 17 | rs62063672 | 44125288 | A | G | 3.781960260626573e-09 | 5.893458102592508 | 31311.0 |
| 17 | rs11653367 | 44191483 | A | G | 4.224703479899445e-09 | 5.875146193674572 | 31486.0 |
| 17 | rs55657917 | 43844560 | T | G | 4.772312670729565e-09 | 5.854922931424293 | 31909.0 |
| 17 | rs111905143 | 44184808 | C | G | 4.9122925833058585e-09 | 5.850116352840077 | 32065.0 |
| 17 | rs56194509 | 43844559 | T | G | 4.929924900067284e-09 | 5.8495203763914905 | 31910.0 |
| 17 | rs112384186 | 44129398 | G | GA | 4.9344669353585786e-09 | 5.849367190484272 | 31969.0 |
| 17 | rs1987152 | 44156180 | A | G | 4.938160365611556e-09 | 5.84924272596936 | 32017.0 |
| 17 | rs1987151 | 44156181 | T | A | 4.938160365611556e-09 | 5.84924272596936 | 32017.0 |
| 17 | rs146712066 | 43970012 | C | CAG | 4.9889700481133636e-09 | 5.847539629369838 | 31857.0 |
| 17 | rs62055546 | 43964567 | A | C | 5.098216431587452e-09 | 5.8439342269399 | 31956.0 |
| 17 | rs368385820 | 44106792 | T | TA | 5.12279839108671e-09 | 5.843133326832207 | 31170.0 |
| 17 | rs78005814 | 44125175 | A | C | 5.264651549300895e-09 | 5.838583438455203 | 26395.0 |
| 17 | rs113161176 | 43974354 | G | A | 5.365659004771693e-09 | 5.835415793866591 | 32108.0 |
| 17 | rs10221243 | 44212310 | G | A | 5.5518526502616836e-09 | 5.8297257994907685 | 32040.0 |
| 17 | rs7207582 | 44210933 | A | G | 5.732365591624566e-09 | 5.824383944090744 | 32042.0 |
| 17 | 17:44127285_CTT_C | 44127285 | CTT | C | 5.859930651890654e-09 | 5.820706747588661 | 32093.0 |
| 17 | rs112166495 | 44005361 | G | A | 5.903844898784417e-09 | 5.8194588700593135 | 31862.0 |
| 17 | rs112572874 | 44072984 | A | G | 5.991871885911168e-09 | 5.816984434847938 | 32096.0 |
| 17 | rs62061820 | 44185431 | T | C | 6.0742608604278374e-09 | 5.814700313852785 | 32040.0 |
| 17 | rs8070942 | 44208674 | G | T | 6.3931664243754524e-09 | 5.806134362495151 | 32053.0 |
| 17 | rs137941099 | 44003625 | T | TAA | 6.436529570610816e-09 | 5.8050018436902455 | 31798.0 |
| 17 | rs9303525 | 44187257 | A | G | 6.510220896236532e-09 | 5.803094155298064 | 32055.0 |
| 17 | rs6503457 | 44208312 | G | T | 6.589502368966091e-09 | 5.801065063555706 | 32050.0 |
| 17 | rs200924217 | 44010463 | A | AC | 6.677133490488729e-09 | 5.798849719070576 | 32103.0 |
| 17 | rs9910284 | 44200191 | C | T | 6.787093327748542e-09 | 5.7961095610645925 | 31742.0 |
| 17 | rs2696578 | 44227623 | T | C | 6.866835012913628e-09 | 5.794149304538688 | 32033.0 |
| 17 | rs112790197 | 44182845 | T | TG | 6.938460764720714e-09 | 5.792407341025692 | 32055.0 |
| 17 | rs2532387 | 44313522 | G | T | 7.018141668292813e-09 | 5.790489904675497 | 31628.0 |
| 17 | 17:44212782_T_C | 44212782 | T | C | 7.029339306356394e-09 | 5.790222143617128 | 32050.0 |
| 17 | rs2668669 | 44284734 | G | C | 7.089644127740483e-09 | 5.7887872122919575 | 31844.0 |
| 17 | rs2696441 | 44274985 | C | A | 7.237490140509601e-09 | 5.7853188864642275 | 31999.0 |
| 17 | rs17689182 | 43783573 | C | T | 7.464685432067008e-09 | 5.7801211442091445 | 31999.0 |
| 17 | rs2696459 | 44285142 | G | C | 7.580921467356801e-09 | 5.777521106106148 | 31613.0 |
| 17 | rs62062271 | 44091988 | T | C | 7.826731463699745e-09 | 5.772148019227184 | 31960.0 |
| 17 | rs62062324 | 44114362 | A | T | 7.827904281718493e-09 | 5.772122778376504 | 31749.0 |
| 17 | rs62062325 | 44114364 | C | A | 7.827904281718493e-09 | 5.772122778376504 | 31749.0 |
| 17 | rs62057121 | 43900760 | G | A | 7.885584212843481e-09 | 5.770885933083626 | 31758.0 |
| 17 | rs4471723 | 44205690 | C | T | 7.950994239275908e-09 | 5.769493929153912 | 32064.0 |
| 17 | rs2732666 | 44286198 | G | A | 7.976385425108773e-09 | 5.768956572248392 | 31885.0 |
| 17 | rs79857651 | 44013475 | G | A | 8.00516699267329e-09 | 5.768349471769907 | 32036.0 |
| 17 | rs113756354 | 44013483 | A | G | 8.00516699267329e-09 | 5.768349471769907 | 32036.0 |
| 17 | rs113141256 | 43991468 | T | TA | 8.005737952935895e-09 | 5.768337449770316 | 32140.0 |
| 17 | 17:44183002_C_A | 44183002 | C | A | 8.007451073610568e-09 | 5.7683013837230765 | 32081.0 |
| 17 | rs16940904 | 44186063 | C | T | 8.139461127296319e-09 | 5.7655445204118765 | 31950.0 |
| 17 | rs2696658 | 44245766 | G | A | 8.292381343594003e-09 | 5.762404809987661 | 32030.0 |
| 17 | rs2532252 | 44257783 | A | G | 8.33967854983302e-09 | 5.761445110299026 | 31604.0 |
| 17 | rs2004674 | 44075110 | A | T | 8.380549158328217e-09 | 5.760620064166237 | 32045.0 |
| 17 | rs2532402 | 44304130 | C | G | 8.432364624567446e-09 | 5.7595796820975576 | 31640.0 |
| 17 | rs62061813 | 44181118 | G | A | 8.781696169491658e-09 | 5.7527236756998015 | 32062.0 |
| 17 | rs111952388 | 44181454 | C | T | 8.781696169491658e-09 | 5.7527236756998015 | 32062.0 |

| | | | | | | | |
|---|---|---|---|---|---|---|---|
| 17 | rs62061814 | 44181508 | T | G | 8.781696169491658e-09 | 5.7527236756998015 | 32062.0 |
| 17 | rs7220752 | 44138201 | T | C | 8.849983965520486e-09 | 5.751414492192211 | 32136.0 |
| 17 | rs62062789 | 44025033 | T | C | 8.990046245816046e-09 | 5.748759732385103 | 32031.0 |
| 17 | rs62062790 | 44025045 | C | T | 8.990046245816046e-09 | 5.748759732385103 | 32031.0 |
| 17 | rs201722973 | 44161747 | C | CCAA CAA | 9.019076026939812e-09 | 5.748214531246369 | 32176.0 |
| 17 | rs2532315 | 44213934 | G | A | 9.071923182049246e-09 | 5.747226387073598 | 31708.0 |
| 17 | rs62054372 | 43798775 | G | C | 9.080642081525762e-09 | 5.7470638977292285 | 31764.0 |
| 17 | 17:43798790_TTTC_T | 43798790 | TTTC | T | 9.080642081525762e-09 | 5.7470638977292285 | 31764.0 |
| 17 | rs62061848 | 44192946 | C | T | 9.09920515535972e-09 | 5.746718452517286 | 31889.0 |
| 17 | rs62063298 | 44048323 | G | T | 9.100629644840502e-09 | 5.746691972121361 | 31939.0 |
| 17 | rs62063299 | 44048336 | T | C | 9.100629644840502e-09 | 5.746691972121361 | 31939.0 |
| 17 | rs62055545 | 43964561 | C | T | 9.115275716936577e-09 | 5.746419943967272 | 31871.0 |
| 17 | rs9907738 | 44151546 | T | G | 9.152838703954849e-09 | 5.745724207134223 | 32138.0 |
| 17 | 17:44024749_CT_C | 44024749 | CT | C | 9.199580751573985e-09 | 5.7448623226173545 | 25804.0 |
| 17 | rs8080583 | 44162597 | C | A | 9.208616587039008e-09 | 5.744696200439411 | 32138.0 |
| 17 | rs112439933 | 44005226 | A | C | 9.25017531913764e-09 | 5.743934185472909 | 32124.0 |
| 17 | 17:44263341_GA_G | 44263341 | GA | G | 9.260577265792153e-09 | 5.743743977624972 | 31990.0 |
| 17 | rs113313477 | 44005186 | G | A | 9.358106492034937e-09 | 5.741970613508898 | 32119.0 |
| 17 | rs7221390 | 44116950 | T | G | 9.42750567347102e-09 | 5.740719642018024 | 32129.0 |
| 17 | rs2066899 | 44155732 | C | T | 9.42750567347102e-09 | 5.740719642018024 | 32140.0 |
| 17 | rs8077487 | 44115440 | T | C | 9.43488254132351e-09 | 5.740587195258987 | 32128.0 |
| 17 | rs62061817 | 44182706 | A | G | 9.474748270521084e-09 | 5.739873169936041 | 32062.0 |
| 17 | rs62062133 | 44114372 | T | C | 9.502340489183279e-09 | 5.7393806805580665 | 31771.0 |
| 17 | rs7218319 | 44126365 | C | T | 9.528927600124829e-09 | 5.738907444500238 | 32134.0 |
| 17 | rs2097760 | 44168677 | A | G | 9.587635626554305e-09 | 5.737867003672412 | 32137.0 |
| 17 | 17:44292021_C_CAAA | 44292021 | C | CAAA | 9.631488714112438e-09 | 5.7370938591622345 | 31895.0 |
| 17 | rs34579536 | 44108906 | A | G | 9.676088810140791e-09 | 5.736311046424043 | 32186.0 |
| 17 | rs2696694 | 44343370 | C | G | 9.746831908265065e-09 | 5.735076541377636 | 30735.0 |
| 17 | rs2696690 | 44344178 | T | G | 9.833930999654512e-09 | 5.733568523087111 | 31132.0 |
| 17 | 17:43996716_GC_G | 43996716 | GC | G | 1.0003248259976625e-08 | 5.730673783696318 | 31925.0 |
| 17 | rs4383188 | 44197603 | T | C | 1.0028645654448288e-08 | 5.730243688830643 | 32075.0 |
| 17 | rs17573907 | 44099082 | C | T | 1.0057248730507933e-08 | 5.729760572748196 | 32174.0 |
| 17 | rs62062286 | 44096486 | C | T | 1.0169719399655156e-08 | 5.727873765069468 | 32139.0 |
| 17 | rs62056842 | 43991515 | T | G | 1.0174125328745758e-08 | 5.727800264704152 | 32128.0 |
| 17 | 17:44104278_TG_T | 44104278 | TG | T | 1.0201686751161462e-08 | 5.727341181821134 | 32150.0 |
| 17 | rs78556223 | 44026478 | A | G | 1.0261254101117566e-08 | 5.726353089473003 | 32014.0 |
| 17 | rs2004673 | 44075099 | T | C | 1.0384436298507755e-08 | 5.724327329595718 | 32046.0 |
| 17 | rs55703644 | 44274766 | T | C | 1.0397933653611111e-08 | 5.724106783894618 | 32042.0 |
| 17 | rs111657460 | 44166362 | G | GA | 1.0456250366158628e-08 | 5.7231570806458585 | 32134.0 |
| 17 | rs62061717 | 44012248 | T | A | 1.0469765016451486e-08 | 5.722937725776888 | 32009.0 |
| 17 | rs7350980 | 44110271 | G | A | 1.0477569726427747e-08 | 5.72281117366795 | 32131.0 |
| 17 | rs1029463 | 44150161 | T | G | 1.0509516905837769e-08 | 5.7222941086111065 | 31902.0 |
| 17 | 17:44161091_GCTCCCCTGG TAAGTCCTAAA_G | 44161091 | GCTCCCCTG GTAAGTCCT AAA | G | 1.0578876443579533e-08 | 5.72117676316835 | 31621.0 |
| 17 | rs4597358 | 44110670 | C | G | 1.0670238098857621e-08 | 5.71971579204759 | 32128.0 |
| 17 | rs62062290 | 44099129 | C | A | 1.0761763244878894e-08 | 5.718264346205183 | 32182.0 |
| 17 | rs62057063 | 43856730 | C | T | 1.0860218739791629e-08 | 5.716716329061586 | 32130.0 |
| 17 | rs111324579 | 44082181 | A | G | 1.0897173786277804e-08 | 5.716138802390118 | 32155.0 |
| 17 | rs2532343 | 44344097 | G | C | 1.0919684784044884e-08 | 5.715787936794063 | 31285.0 |
| 17 | rs76924476 | 44199029 | T | C | 1.0960879223779716e-08 | 5.715147679782088 | 32053.0 |
| 17 | rs113343439 | 44314217 | A | G | 1.109803958939614e-08 | 5.7130326190496525 | 32135.0 |
| 17 | 17:44617964_TA_T | 44617964 | TA | T | 1.127573554916828e-08 | 5.710329945665555 | 24240.0 |
| 17 | rs1029464 | 44149348 | A | C | 1.1311031160568889e-08 | 5.70979804621208 | 32085.0 |
| 17 | rs62062277 | 44093753 | T | G | 1.1376948637395873e-08 | 5.7088089847934285 | 32117.0 |
| 17 | rs55686102 | 44202608 | C | T | 1.1401643569912871e-08 | 5.708439882015022 | 32098.0 |

| | | | | | | | |
|---|---|---|---|---|---|---|---|
| 17 | rs78765934 | 44261613 | T | TCC | 1.1454599412845998e-08 | 5.707650989770802 | 31949.0 |
| 17 | rs17577159 | 44188477 | T | G | 1.1460606536654154e-08 | 5.707561724543995 | 32104.0 |
| 17 | rs62061823 | 44188645 | C | T | 1.1460606536654154e-08 | 5.707561724543995 | 32104.0 |
| 17 | rs35095455 | 44249911 | T | TA | 1.1463367602877696e-08 | 5.707520710641857 | 31998.0 |
| 17 | rs2532246 | 44262418 | G | A | 1.1494596762329644e-08 | 5.707057488280466 | 31935.0 |
| 17 | rs62063786 | 44061023 | G | A | 1.1525582796761422e-08 | 5.7065990793488535 | 32186.0 |
| 17 | 17:44058699_TAGG_T | 44058699 | TAGG | T | 1.165265937673128e-08 | 5.704731544771029 | 31776.0 |
| 17 | rs62063300 | 44048350 | T | C | 1.1661328529001888e-08 | 5.704604863856237 | 31949.0 |
| 17 | rs4581739 | 44204739 | T | C | 1.1731491510169164e-08 | 5.703582938351105 | 32095.0 |
| 17 | rs113638513 | 44204749 | A | G | 1.1731491510169164e-08 | 5.703582938351105 | 32095.0 |
| 17 | 17:44003580_GA_G | 44003580 | GA | G | 1.173656053059945e-08 | 5.703509338157732 | 32113.0 |
| 17 | rs17577369 | 44192923 | A | G | 1.174637227729239e-08 | 5.703366963154106 | 31867.0 |
| 17 | rs111697872 | 43840726 | C | CTT | 1.1750282586896844e-08 | 5.7033102541508045 | 31913.0 |
| 17 | rs62063280 | 44037299 | G | C | 1.1892244391697479e-08 | 5.701263783491683 | 32098.0 |
| 17 | rs113201171 | 44082249 | A | T | 1.1999309197316695e-08 | 5.69973601782045 | 32168.0 |
| 17 | rs17661027 | 44193097 | C | A | 1.20386153221461e-08 | 5.699178459477285 | 32088.0 |
| 17 | rs2696605 | 44213921 | A | C | 1.213082158106812e-08 | 5.697877422455728 | 31737.0 |
| 17 | rs77926909 | 43994252 | T | C | 1.2141386274598999e-08 | 5.697728967878548 | 32144.0 |
| 17 | rs2696629 | 44327438 | C | G | 1.2173651456289728e-08 | 5.697276354859926 | 31479.0 |
| 17 | rs75210307 | 43940012 | T | C | 1.2175116136244289e-08 | 5.6972558361318635 | 31670.0 |
| 17 | rs74944644 | 43940020 | T | C | 1.2175116136244289e-08 | 5.6972558361318635 | 31670.0 |
| 17 | rs2532247 | 44262403 | C | T | 1.2242418763008577e-08 | 5.6963155703547415 | 31931.0 |
| 17 | rs1918793 | 44228169 | C | T | 1.228459207976397e-08 | 5.695728935359227 | 31985.0 |
| 17 | rs12150087 | 44210988 | C | G | 1.2309389618270796e-08 | 5.6953849120653794 | 32098.0 |
| 17 | rs1468241 | 44196153 | A | G | 1.2355984669810972e-08 | 5.694740303247832 | 32095.0 |
| 17 | rs62055934 | 43848638 | G | A | 1.2375669197289433e-08 | 5.694468691456607 | 31958.0 |
| 17 | rs77635460 | 44186593 | T | C | 1.2388195796965373e-08 | 5.6942960649313 | 32100.0 |
| 17 | rs2696569 | 44266290 | G | C | 1.2420840781774168e-08 | 5.693846986729873 | 31550.0 |
| 17 | rs78929339 | 44189910 | A | G | 1.2435347419987253e-08 | 5.693647795210402 | 32005.0 |
| 17 | rs78590209 | 44098862 | A | G | 1.2442650981671116e-08 | 5.693547595001906 | 32186.0 |
| 17 | rs55744103 | 44316280 | T | C | 1.245004682320814e-08 | 5.693446186990813 | 32135.0 |
| 17 | rs62061816 | 44182336 | C | A | 1.245797572558895e-08 | 5.693337534912724 | 32101.0 |
| 17 | rs111854640 | 44276431 | T | TAGAA | 1.2534611686377576e-08 | 5.692290819441827 | 31731.0 |
| 17 | 17:43972025_TC_T | 43972025 | TC | T | 1.2553865033433435e-08 | 5.692028829174291 | 32150.0 |
| 17 | rs111750615 | 44156132 | T | TTTA | 1.2591479197284734e-08 | 5.6915181182271075 | 32021.0 |
| 17 | rs9709338 | 44094871 | A | G | 1.2608410416095024e-08 | 5.691288716131664 | 32018.0 |
| 17 | 17:44265192_AT_A | 44265192 | AT | A | 1.2621525157595216e-08 | 5.691111229334996 | 31785.0 |
| 17 | rs55857174 | 44137386 | T | TAA | 1.2685146846171165e-08 | 5.690252747941858 | 31940.0 |
| 17 | rs17650842 | 44037491 | A | G | 1.2748271867040671e-08 | 5.6894050929992295 | 32121.0 |
| 17 | rs113138968 | 44314261 | C | T | 1.2764146342859638e-08 | 5.689192569226142 | 32157.0 |
| 17 | rs9899964 | 44314342 | T | C | 1.2764146342859638e-08 | 5.689192569226142 | 32157.0 |
| 17 | rs974294 | 44150163 | T | G | 1.2817956641416064e-08 | 5.688474075282652 | 31961.0 |
| 17 | rs77095128 | 44204304 | T | G | 1.2883096075054548e-08 | 5.687608221567945 | 32098.0 |
| 17 | rs17660907 | 44191085 | G | A | 1.2918755151845635e-08 | 5.687136029855043 | 32108.0 |
| 17 | rs79772780 | 44189855 | G | A | 1.292012507414096e-08 | 5.68711791483531 | 31916.0 |
| 17 | rs55909047 | 44209957 | A | G | 1.2929078741216775e-08 | 5.686999562921853 | 32106.0 |
| 17 | rs2732631 | 44289232 | G | T | 1.2972287866391269e-08 | 5.686429530498604 | 31747.0 |
| 17 | rs62061810 | 44178884 | G | A | 1.2977974158376992e-08 | 5.686354652009177 | 32107.0 |
| 17 | 17:44251554_ATG_A | 44251554 | ATG | A | 1.2978799791092566e-08 | 5.686343782524999 | 31316.0 |
| 17 | rs56257094 | 44201667 | G | A | 1.2988895022601303e-08 | 5.686210932740496 | 32099.0 |
| 17 | rs55672516 | 44201680 | A | C | 1.2988895022601303e-08 | 5.686210932740496 | 32099.0 |
| 17 | rs55669501 | 44202564 | G | A | 1.2988895022601303e-08 | 5.686210932740496 | 32099.0 |
| 17 | rs55885063 | 44202952 | C | G | 1.2988895022601303e-08 | 5.686210932740496 | 32099.0 |
| 17 | rs56273589 | 44203131 | C | G | 1.2988895022601303e-08 | 5.686210932740496 | 32099.0 |

| | | | | | | | |
|---|---|---|---|---|---|---|---|
| 17 | rs62063171 | 44203311 | C | T | 1.2988895022601303e-08 | 5.686210932740496 | 32099.0 |
| 17 | rs62063172 | 44203366 | T | C | 1.2988895022601303e-08 | 5.686210932740496 | 32099.0 |
| 17 | rs76380490 | 44203413 | C | T | 1.2988895022601303e-08 | 5.686210932740496 | 32099.0 |
| 17 | rs113120855 | 44203520 | C | T | 1.2988895022601303e-08 | 5.686210932740496 | 32099.0 |
| 17 | 17:44238130_AT_A | 44238130 | AT | A | 1.2992292422576544e-08 | 5.686166246682251 | 31893.0 |
| 17 | rs34847058 | 44285460 | G | GA | 1.3000008675557232e-08 | 5.6860647968385445 | 31769.0 |
| 17 | 17:44048939_AG_A | 44048939 | AG | A | 1.3000835683577132e-08 | 5.6860539271783255 | 31987.0 |
| 17 | rs79301522 | 44207932 | A | G | 1.3066240341538411e-08 | 5.685196410957714 | 32095.0 |
| 17 | rs62064668 | 44082981 | G | A | 1.30851847717582e-08 | 5.6849488106018375 | 32143.0 |
| 17 | rs35014258 | 44192592 | A | AT | 1.3086664494636077e-08 | 5.684929485551902 | 32133.0 |
| 17 | rs111327992 | 44126691 | G | A | 1.3097489958919875e-08 | 5.684788170490058 | 32157.0 |
| 17 | rs2532289 | 44238571 | C | G | 1.326458476114663e-08 | 5.682621199805998 | 31987.0 |
| 17 | rs2532256 | 44255017 | T | C | 1.3280432943912072e-08 | 5.682417051363528 | 32069.0 |
| 17 | 17:44285467_A_G | 44285467 | A | G | 1.3302971166540527e-08 | 5.682127132413973 | 31784.0 |
| 17 | rs111541901 | 43994358 | C | T | 1.3306355134979376e-08 | 5.6820836441664735 | 32160.0 |
| 17 | rs1881195 | 44257473 | T | C | 1.3307577331118151e-08 | 5.6820679400511205 | 31889.0 |
| 17 | rs62063174 | 44205122 | C | G | 1.3329501532715356e-08 | 5.681786471647012 | 32108.0 |
| 17 | rs17661348 | 44207066 | A | G | 1.3329501532715356e-08 | 5.681786471647012 | 32108.0 |
| 17 | rs62063202 | 44208945 | C | T | 1.3329501532715356e-08 | 5.681786471647012 | 32108.0 |
| 17 | rs62063203 | 44209313 | T | C | 1.3329501532715356e-08 | 5.681786471647012 | 32108.0 |
| 17 | rs111724389 | 44209364 | G | A | 1.3329501532715356e-08 | 5.681786471647012 | 32108.0 |
| 17 | rs112401617 | 44209474 | A | T | 1.3329501532715356e-08 | 5.681786471647012 | 32108.0 |
| 17 | rs111424940 | 44211502 | G | A | 1.3329501532715356e-08 | 5.681786471647012 | 32108.0 |
| 17 | rs62063207 | 44211902 | A | G | 1.3329501532715356e-08 | 5.681786471647012 | 32108.0 |
| 17 | rs62063208 | 44212032 | C | A | 1.3329501532715356e-08 | 5.681786471647012 | 32108.0 |
| 17 | rs2532316 | 44213712 | G | A | 1.334589722172127e-08 | 5.681576273470894 | 32086.0 |
| 17 | rs2942163 | 44282007 | T | C | 1.334589722172127e-08 | 5.681576273470894 | 32068.0 |
| 17 | rs62062778 | 44022528 | A | G | 1.334702868792924e-08 | 5.681561776953956 | 32160.0 |
| 17 | rs2696630 | 44327482 | G | A | 1.3350706603100502e-08 | 5.681514663192817 | 31412.0 |
| 17 | rs2532274 | 44247164 | A | G | 1.3379693137836802e-08 | 5.681143788998659 | 31986.0 |
| 17 | rs62061771 | 44169406 | G | A | 1.3403438095770652e-08 | 5.680840560412812 | 32144.0 |
| 17 | rs111689007 | 43981103 | C | A | 1.3432726718631917e-08 | 5.680467256014563 | 32180.0 |
| 17 | rs17577954 | 44209628 | T | C | 1.3454473026867577e-08 | 5.680190594980607 | 32105.0 |
| 17 | rs75686108 | 44022376 | C | T | 1.3477586574570167e-08 | 5.679897015494713 | 32163.0 |
| 17 | rs112003140 | 43980998 | G | A | 1.3497592621304009e-08 | 5.679643300948688 | 32177.0 |
| 17 | rs78729125 | 43970108 | C | A | 1.3587309242380043e-08 | 5.6785099987296475 | 32170.0 |
| 17 | rs77727624 | 43790770 | A | G | 1.3593835915044665e-08 | 5.67842783756996 | 31843.0 |
| 17 | rs2532236 | 44271430 | A | G | 1.3618723038718737e-08 | 5.678114896757539 | 32075.0 |
| 17 | rs2532423 | 44285952 | T | C | 1.3620839061405951e-08 | 5.678088314660332 | 32012.0 |
| 17 | rs1528075 | 44220454 | T | G | 1.3733354060271753e-08 | 5.676680615381238 | 32090.0 |
| 17 | rs62055489 | 43950441 | C | A | 1.3743927742705082e-08 | 5.676548902230616 | 31916.0 |
| 17 | rs1816 | 44276618 | A | G | 1.377151504909594e-08 | 5.6762057184594354 | 31850.0 |
| 17 | rs62056859 | 43994435 | A | G | 1.3778809492125065e-08 | 5.676115087844011 | 32162.0 |
| 17 | rs62061847 | 44191682 | G | A | 1.3829879194351662e-08 | 5.675481869139421 | 32109.0 |
| 17 | 17:44282744_G_C | 44282744 | G | C | 1.3830172112089745e-08 | 5.675478243776992 | 32064.0 |
| 17 | rs112402370 | 44092744 | A | G | 1.3845314478917476e-08 | 5.6752909323857805 | 31806.0 |
| 17 | rs17577313 | 44191835 | A | G | 1.3917340317897564e-08 | 5.6744026871640685 | 32103.0 |
| 17 | rs62055876 | 43832940 | A | G | 1.3964181600160998e-08 | 5.673827419119629 | 32109.0 |
| 17 | rs56119705 | 44209985 | T | C | 1.3977494546333131e-08 | 5.673664262001751 | 32106.0 |
| 17 | rs2668657 | 44282892 | T | C | 1.399437548706368e-08 | 5.673457594183887 | 31571.0 |
| 17 | rs12185233 | 43923654 | G | C | 1.4088921056165821e-08 | 5.6723045615076515 | 32186.0 |
| 17 | rs740710 | 44214815 | C | G | 1.41123066038136e-08 | 5.672020522139162 | 32088.0 |
| 17 | rs1918802 | 44215045 | T | C | 1.41123066038136e-08 | 5.672020522139162 | 32088.0 |
| 17 | rs2696603 | 44215823 | A | G | 1.41123066038136e-08 | 5.672020522139162 | 32088.0 |

| | | | | | | | |
|---|---|---|---|---|---|---|---|
| 17 | rs2532313 | 44217293 | G | A | 1.41123066038136e-08 | 5.672020522139162 | 32088.0 |
| 17 | rs2532308 | 44220942 | G | A | 1.41123066038136e-08 | 5.672020522139162 | 32088.0 |
| 17 | 17:44221350_G_A | 44221350 | G | A | 1.41123066038136e-08 | 5.672020522139162 | 32088.0 |
| 17 | rs2696590 | 44221602 | G | C | 1.41123066038136e-08 | 5.672020522139162 | 32088.0 |
| 17 | rs1406074 | 44222067 | T | G | 1.41123066038136e-08 | 5.672020522139162 | 32088.0 |
| 17 | rs1918798 | 44222335 | G | A | 1.41123066038136e-08 | 5.672020522139162 | 32088.0 |
| 17 | rs2696458 | 44284420 | C | T | 1.4118781666943559e-08 | 5.671941957262657 | 31913.0 |
| 17 | rs9944484 | 43990662 | G | A | 1.417518781689061e-08 | 5.671259032654301 | 32174.0 |
| 17 | rs12150320 | 44213204 | C | T | 1.4198814017473974e-08 | 5.670973767934006 | 32089.0 |
| 17 | 17:44279371_TCAAA_T | 44279371 | TCAAA | T | 1.4222077426296612e-08 | 5.6706933337770975 | 32049.0 |
| 17 | rs382104 | 44305026 | G | A | 1.4259155330449068e-08 | 5.670247289362929 | 32024.0 |
| 17 | rs2696600 | 44216226 | A | G | 1.4298749085903576e-08 | 5.669772221705316 | 32089.0 |
| 17 | rs62062279 | 44093964 | T | C | 1.4320354377379992e-08 | 5.669513528161421 | 32175.0 |
| 17 | rs62062280 | 44093993 | C | T | 1.4320354377379992e-08 | 5.669513528161421 | 32175.0 |
| 17 | rs112073200 | 44201791 | G | C | 1.4325811238024417e-08 | 5.669448249760468 | 32094.0 |
| 17 | 17:43841356_CTT_C | 43841356 | CTT | C | 1.4338450687139852e-08 | 5.6692971414373945 | 31769.0 |
| 17 | rs77084869 | 43982903 | A | G | 1.4355961733755425e-08 | 5.669088005413522 | 32179.0 |
| 17 | rs74573628 | 43982961 | T | C | 1.4355961733755425e-08 | 5.669088005413522 | 32179.0 |
| 17 | rs62056805 | 43983363 | G | A | 1.4355961733755425e-08 | 5.669088005413522 | 32179.0 |
| 17 | rs62054833 | 43933673 | T | C | 1.4363458412531882e-08 | 5.668998547639752 | 32155.0 |
| 17 | rs740711 | 44214814 | G | A | 1.4416347892423576e-08 | 5.668368703900398 | 32014.0 |
| 17 | 17:44155925_TACAA_T | 44155925 | TACAA | T | 1.442316326742013e-08 | 5.668287705112364 | 32133.0 |
| 17 | rs17770186 | 43964191 | C | A | 1.4424384265299169e-08 | 5.668273197828037 | 32183.0 |
| 17 | rs62060915 | 44251548 | G | A | 1.4435886969544584e-08 | 5.668136586990582 | 32080.0 |
| 17 | rs2696664 | 44251698 | G | A | 1.4435886969544584e-08 | 5.668136586990582 | 32080.0 |
| 17 | rs2732643 | 44251907 | C | T | 1.4435886969544584e-08 | 5.668136586990582 | 32080.0 |
| 17 | rs17573593 | 44082603 | A | C | 1.4444952955703834e-08 | 5.668028990110214 | 32177.0 |
| 17 | rs62061853 | 44196125 | T | C | 1.4454942188110674e-08 | 5.667910511898155 | 32105.0 |
| 17 | rs62063163 | 44201865 | C | T | 1.4461877474290664e-08 | 5.667828302065617 | 32095.0 |
| 17 | rs62063164 | 44201887 | C | T | 1.4461877474290664e-08 | 5.667828302065617 | 32095.0 |
| 17 | rs62063165 | 44202147 | A | G | 1.4461877474290664e-08 | 5.667828302065617 | 32095.0 |
| 17 | rs62063166 | 44202231 | A | G | 1.4461877474290664e-08 | 5.667828302065617 | 32095.0 |
| 17 | rs17577024 | 44186252 | G | A | 1.4501205034190557e-08 | 5.667362842215812 | 32110.0 |
| 17 | rs17577052 | 44186301 | T | C | 1.4501205034190557e-08 | 5.667362842215812 | 32110.0 |
| 17 | rs62061822 | 44186894 | A | C | 1.4501205034190557e-08 | 5.667362842215812 | 32110.0 |
| 17 | rs17577094 | 44187492 | A | G | 1.4501205034190557e-08 | 5.667362842215812 | 32110.0 |
| 17 | rs62061824 | 44188755 | T | C | 1.4501205034190557e-08 | 5.667362842215812 | 32110.0 |
| 17 | rs55794067 | 44188764 | T | C | 1.4501205034190557e-08 | 5.667362842215812 | 32110.0 |
| 17 | rs17660847 | 44189373 | C | T | 1.4501205034190557e-08 | 5.667362842215812 | 32110.0 |
| 17 | rs17660865 | 44189408 | T | C | 1.4501205034190557e-08 | 5.667362842215812 | 32110.0 |
| 17 | rs17660936 | 44191159 | C | T | 1.4501205034190557e-08 | 5.667362842215812 | 32110.0 |
| 17 | rs4548919 | 44192395 | G | T | 1.4501205034190557e-08 | 5.667362842215812 | 32110.0 |
| 17 | rs4606752 | 44192618 | A | G | 1.4501205034190557e-08 | 5.667362842215812 | 32110.0 |
| 17 | rs62061849 | 44193882 | C | T | 1.4501205034190557e-08 | 5.667362842215812 | 32110.0 |
| 17 | rs17661045 | 44194152 | T | A | 1.4501205034190557e-08 | 5.667362842215812 | 32110.0 |
| 17 | rs17577447 | 44194381 | T | A | 1.4501205034190557e-08 | 5.667362842215812 | 32110.0 |
| 17 | rs62061850 | 44194519 | T | C | 1.4501205034190557e-08 | 5.667362842215812 | 32110.0 |
| 17 | rs62061852 | 44195424 | A | C | 1.4501205034190557e-08 | 5.667362842215812 | 32110.0 |
| 17 | rs727425 | 44195495 | G | A | 1.4501205034190557e-08 | 5.667362842215812 | 32110.0 |
| 17 | rs17661141 | 44196015 | C | G | 1.4501205034190557e-08 | 5.667362842215812 | 32110.0 |
| 17 | rs1468240 | 44196447 | T | A | 1.4501205034190557e-08 | 5.667362842215812 | 32110.0 |
| 17 | rs112813897 | 44196799 | G | A | 1.4501205034190557e-08 | 5.667362842215812 | 32110.0 |
| 17 | 17:44243543_TA_T | 44243543 | TA | T | 1.450703646504456e-08 | 5.667293928949545 | 32076.0 |
| 17 | rs112102887 | 44179749 | G | A | 1.4521778749498228e-08 | 5.6671198310949435 | 32110.0 |

| | | | | | | | |
|---|---|---|---|---|---|---|---|
| 17 | rs112893440 | 44179969 | G | A | 1.4521778749498228e-08 | 5.6671198310949435 | 32110.0 |
| 17 | rs17576842 | 44179992 | A | G | 1.4521778749498228e-08 | 5.6671198310949435 | 32110.0 |
| 17 | rs62061772 | 44169437 | T | C | 1.4582540474274828e-08 | 5.6664040776776154 | 32151.0 |
| 17 | rs56240678 | 44176170 | C | G | 1.4582694747662427e-08 | 5.666402264076862 | 32111.0 |
| 17 | rs62061808 | 44177337 | C | T | 1.4582694747662427e-08 | 5.666402264076862 | 32111.0 |
| 17 | rs77604208 | 44177669 | C | T | 1.4582694747662427e-08 | 5.666402264076862 | 32111.0 |
| 17 | rs17660464 | 44177993 | C | A | 1.4582694747662427e-08 | 5.666402264076862 | 32111.0 |
| 17 | rs17576779 | 44178272 | C | T | 1.4582694747662427e-08 | 5.666402264076862 | 32111.0 |
| 17 | rs17660488 | 44178337 | T | C | 1.4582694747662427e-08 | 5.666402264076862 | 32111.0 |
| 17 | rs62061809 | 44178839 | T | C | 1.4582694747662427e-08 | 5.666402264076862 | 32111.0 |
| 17 | rs62061811 | 44180173 | T | G | 1.4582694747662427e-08 | 5.666402264076862 | 32111.0 |
| 17 | rs17576870 | 44180442 | A | G | 1.4582694747662427e-08 | 5.666402264076862 | 32111.0 |
| 17 | rs17660595 | 44180954 | A | G | 1.4582694747662427e-08 | 5.666402264076862 | 32111.0 |
| 17 | rs62061812 | 44180997 | C | T | 1.4582694747662427e-08 | 5.666402264076862 | 32111.0 |
| 17 | rs62061815 | 44181681 | A | G | 1.4582694747662427e-08 | 5.666402264076862 | 32111.0 |
| 17 | rs17576954 | 44181933 | T | C | 1.4582694747662427e-08 | 5.666402264076862 | 32111.0 |
| 17 | rs77924366 | 44182181 | T | C | 1.4582694747662427e-08 | 5.666402264076862 | 32111.0 |
| 17 | rs62061818 | 44182917 | C | G | 1.4582694747662427e-08 | 5.666402264076862 | 32111.0 |
| 17 | rs111676341 | 44183403 | A | G | 1.4582694747662427e-08 | 5.666402264076862 | 32111.0 |
| 17 | rs17576989 | 44183616 | C | A | 1.4582694747662427e-08 | 5.666402264076862 | 32111.0 |
| 17 | rs79923708 | 44183790 | G | A | 1.4582694747662427e-08 | 5.666402264076862 | 32111.0 |
| 17 | rs76761511 | 44185245 | C | A | 1.4582694747662427e-08 | 5.666402264076862 | 32111.0 |
| 17 | rs62056775 | 43969952 | T | C | 1.4599932017891488e-08 | 5.6661997441695355 | 32176.0 |
| 17 | rs62056776 | 43969995 | C | T | 1.4599932017891488e-08 | 5.6661997441695355 | 32176.0 |
| 17 | rs17770343 | 43970154 | T | C | 1.4599932017891488e-08 | 5.6661997441695355 | 32176.0 |
| 17 | rs34200189 | 44343125 | C | T | 1.4610593096461796e-08 | 5.666074603857435 | 31522.0 |
| 17 | rs111880194 | 44194621 | C | G | 1.4642931968502232e-08 | 5.665695550326655 | 32107.0 |
| 17 | rs62063301 | 44048941 | G | C | 1.4668150199158358e-08 | 5.665400523841102 | 32107.0 |
| 17 | rs62063302 | 44048943 | G | A | 1.4668150199158358e-08 | 5.665400523841102 | 32107.0 |
| 17 | rs55788678 | 43992233 | C | T | 1.4694861718536931e-08 | 5.665088564316528 | 32027.0 |
| 17 | 17:43989129_CA_C | 43989129 | CA | C | 1.4757582417822114e-08 | 5.664358219202338 | 32178.0 |
| 17 | rs199995459 | 43989149 | A | AAC | 1.4757582417822114e-08 | 5.664358219202338 | 32178.0 |
| 17 | rs66619507 | 43989151 | A | AC | 1.4757582417822114e-08 | 5.664358219202338 | 32178.0 |
| 17 | rs113906906 | 43989159 | G | GC | 1.4757582417822114e-08 | 5.664358219202338 | 32178.0 |
| 17 | 17:43989509_TA_T | 43989509 | TA | T | 1.4757582417822114e-08 | 5.664358219202338 | 32178.0 |
| 17 | 17:43774255_TAGCC_T | 43774255 | TAGCC | T | 1.4760548399984342e-08 | 5.664323756817423 | 32091.0 |
| 17 | rs2696682 | 44310101 | A | G | 1.480239452093904e-08 | 5.663838253360081 | 32023.0 |
| 17 | rs2532314 | 44217112 | G | T | 1.4802864189849892e-08 | 5.663832811777213 | 32090.0 |
| 17 | rs1918799 | 44218242 | C | T | 1.4802864189849892e-08 | 5.663832811777213 | 32090.0 |
| 17 | rs113089855 | 44256811 | A | AACAAG | 1.4844200020717588e-08 | 5.663354550642258 | 32062.0 |
| 17 | rs2696703 | 44260967 | G | A | 1.4846240996840432e-08 | 5.663330969800586 | 32075.0 |
| 17 | rs62055525 | 43958736 | G | A | 1.491132984132945e-08 | 5.662580599102138 | 32145.0 |
| 17 | rs62062772 | 44020314 | C | T | 1.4922056133213005e-08 | 5.662457247483374 | 32177.0 |
| 17 | rs62062773 | 44020545 | A | G | 1.4922056133213005e-08 | 5.662457247483374 | 32177.0 |
| 17 | rs2732590 | 44266972 | A | G | 1.494679748327177e-08 | 5.662173051792737 | 32072.0 |
| 17 | rs62062771 | 44020282 | G | A | 1.4961345195018435e-08 | 5.662006160305352 | 32173.0 |
| 17 | rs761294768 | 44262203 | CTTTTTT | C | 1.4980235746671896e-08 | 5.661789682796763 | 32064.0 |
| 17 | rs75839508 | 44022319 | G | A | 1.5007240173080026e-08 | 5.661480683828107 | 32167.0 |
| 17 | 17:44061608_GT_G | 44061608 | GT | G | 1.5033921448897772e-08 | 5.661175912496357 | 32162.0 |
| 17 | rs80120913 | 44177755 | C | G | 1.504139417959669e-08 | 5.6610906481590275 | 32110.0 |
| 17 | rs2696697 | 44257788 | T | C | 1.5054228211434793e-08 | 5.6609443070020085 | 31675.0 |
| 17 | rs430688 | 43685707 | T | C | 1.5055077120938775e-08 | 5.6609346315114415 | 31906.0 |
| 17 | rs111878933 | 43856341 | C | T | 1.509470970387831e-08 | 5.660483505956402 | 32059.0 |
| 17 | rs111735741 | 43856372 | G | A | 1.509470970387831e-08 | 5.660483505956402 | 32059.0 |

| | | | | | | | |
|---|---|---|---|---|---|---|---|
| 17 | rs62063281 | 44038785 | A | G | 1.5103542661074415e-08 | 5.66038311990287 | 32186.0 |
| 17 | rs1122381 | 44205839 | A | G | 1.515376313671144e-08 | 5.659813448622416 | 32105.0 |
| 17 | rs55709241 | 44041101 | T | C | 1.5153976735995342e-08 | 5.6598110295974315 | 31917.0 |
| 17 | rs56280951 | 44041107 | G | A | 1.5153976735995342e-08 | 5.6598110295974315 | 31917.0 |
| 17 | rs111678697 | 44197032 | T | C | 1.5193008097457935e-08 | 5.659369552062273 | 32109.0 |
| 17 | rs1966345 | 44197806 | T | C | 1.5193008097457935e-08 | 5.659369552062273 | 32109.0 |
| 17 | rs62061854 | 44198080 | C | T | 1.5193008097457935e-08 | 5.659369552062273 | 32109.0 |
| 17 | rs112560196 | 44200078 | A | T | 1.5193008097457935e-08 | 5.659369552062273 | 32109.0 |
| 17 | rs75652502 | 44200510 | C | A | 1.5193008097457935e-08 | 5.659369552062273 | 32109.0 |
| 17 | rs62061859 | 44201109 | C | G | 1.5193008097457935e-08 | 5.659369552062273 | 32109.0 |
| 17 | rs55802590 | 44201344 | G | C | 1.5193008097457935e-08 | 5.659369552062273 | 32109.0 |
| 17 | 17:44325593_TAA_T | 44325593 | TAA | T | 1.5240835097416602e-08 | 5.658830088017088 | 31817.0 |
| 17 | rs62062777 | 44022068 | G | A | 1.5288378848105168e-08 | 5.658295446172117 | 32175.0 |
| 17 | rs34256996 | 44197775 | G | GCA | 1.5305786867496383e-08 | 5.658100092283383 | 32050.0 |
| 17 | rs3087534 | 44198806 | T | C | 1.5309130568436687e-08 | 5.658062593769796 | 32105.0 |
| 17 | rs62062797 | 44027366 | T | G | 1.532742573560103e-08 | 5.657857560185351 | 32003.0 |
| 17 | rs74863825 | 43963120 | C | T | 1.5380055840294728e-08 | 5.657269058414851 | 32177.0 |
| 17 | rs62056761 | 43969686 | T | A | 1.5380055840294728e-08 | 5.657269058414851 | 32177.0 |
| 17 | rs17770337 | 43969779 | G | T | 1.5380055840294728e-08 | 5.657269058414851 | 32177.0 |
| 17 | rs62056773 | 43969833 | A | G | 1.5380055840294728e-08 | 5.657269058414851 | 32177.0 |
| 17 | rs62056774 | 43969901 | G | A | 1.5380055840294728e-08 | 5.657269058414851 | 32177.0 |
| 17 | rs62056777 | 43970303 | G | A | 1.5380055840294728e-08 | 5.657269058414851 | 32177.0 |
| 17 | rs2696604 | 44214094 | A | G | 1.5414010192337196e-08 | 5.656890423202205 | 32086.0 |
| 17 | rs62062798 | 44027383 | A | G | 1.5434602942729444e-08 | 5.6566611816965855 | 32000.0 |
| 17 | 17:43947176_AC_A | 43947176 | AC | A | 1.5440476000529367e-08 | 5.656595856402808 | 31848.0 |
| 17 | rs62064663 | 44080039 | T | G | 1.5445045449826602e-08 | 5.6565450476761425 | 32163.0 |
| 17 | rs2732670 | 44283479 | T | C | 1.545799942736159e-08 | 5.656401088833977 | 31925.0 |
| 17 | rs17577496 | 44194835 | T | C | 1.5460504365957716e-08 | 5.656373264722557 | 32109.0 |
| 17 | rs55933304 | 43992305 | G | A | 1.5474452169318255e-08 | 5.6562184167035685 | 32115.0 |
| 17 | 17:44126473_AG_A | 44126473 | AG | A | 1.5517626734911468e-08 | 5.6557399520549465 | 32128.0 |
| 17 | rs78773690 | 44126478 | T | C | 1.5517626734911468e-08 | 5.6557399520549465 | 32128.0 |
| 17 | rs111164082 | 44047037 | T | A | 1.553961406386533e-08 | 5.655496783070992 | 31773.0 |
| 17 | rs62054832 | 43933579 | C | A | 1.555165986548028e-08 | 5.655363704118821 | 32137.0 |
| 17 | rs28416808 | 44021218 | C | T | 1.5553357964488248e-08 | 5.655344952005117 | 32173.0 |
| 17 | rs17576695 | 44172327 | T | A | 1.555609721710122e-08 | 5.655314706619028 | 32112.0 |
| 17 | rs35631660 | 43832337 | A | G | 1.558845615664487e-08 | 5.6549578072032975 | 32159.0 |
| 17 | rs529311580 | 43699113 | A | G | 1.5595430186242067e-08 | 5.654880982160933 | 31863.0 |
| 17 | rs17661385 | 44207778 | A | G | 1.5599550144886507e-08 | 5.654835612886558 | 32105.0 |
| 17 | rs112915466 | 43958744 | A | G | 1.5600429207557104e-08 | 5.654825934093138 | 32151.0 |
| 17 | rs56343551 | 43959092 | G | T | 1.5647970816190766e-08 | 5.654303271475945 | 32176.0 |
| 17 | rs55975673 | 43959113 | A | G | 1.5647970816190766e-08 | 5.654303271475945 | 32176.0 |
| 17 | rs112638515 | 44016601 | A | AG | 1.5654695307121113e-08 | 5.654229468348861 | 32173.0 |
| 17 | rs62055947 | 43852605 | C | T | 1.5661091714153657e-08 | 5.6541592946015165 | 32097.0 |
| 17 | rs10514901 | 44194735 | G | T | 1.5667656209341226e-08 | 5.654087305729983 | 32110.0 |
| 17 | rs62061851 | 44194891 | C | T | 1.5667656209341226e-08 | 5.654087305729983 | 32110.0 |
| 17 | rs56227067 | 43939609 | A | T | 1.5671739693320283e-08 | 5.65404253939495 | 32153.0 |
| 17 | rs79115768 | 44021061 | C | T | 1.5672401978272502e-08 | 5.654035279978718 | 32176.0 |
| 17 | rs62062774 | 44021463 | T | C | 1.5672401978272502e-08 | 5.654035279978718 | 32176.0 |
| 17 | 17:43951599_CAGAT_C | 43951599 | CAGAT | C | 1.5688968027099458e-08 | 5.653853793615996 | 32111.0 |
| 17 | rs62056841 | 43990041 | A | C | 1.5700021600430884e-08 | 5.653732801685147 | 32163.0 |
| 17 | rs2532417 | 44289220 | C | T | 1.5706270246174382e-08 | 5.65366444088261 | 32036.0 |
| 17 | rs113271363 | 43996721 | C | G | 1.5715730800532446e-08 | 5.653560991738008 | 32177.0 |
| 17 | rs2055794 | 43951717 | G | A | 1.5730846078731442e-08 | 5.653395834846817 | 32174.0 |
| 17 | rs76618565 | 44068481 | G | A | 1.5740432005334742e-08 | 5.653291174196489 | 32181.0 |

| | | | | | | | |
|---|---|---|---|---|---|---|---|
| 17 | rs2696599 | 44217226 | A | G | 1.574203945539724e-08 | 5.653273629865727 | 32087.0 |
| 17 | rs62062776 | 44022043 | A | C | 1.5761119426375143e-08 | 5.6530655164923935 | 32171.0 |
| 17 | rs2055795 | 43951995 | C | T | 1.5776833035174098e-08 | 5.652894305129718 | 32175.0 |
| 17 | rs2055796 | 43952191 | A | T | 1.5776833035174098e-08 | 5.652894305129718 | 32175.0 |
| 17 | rs62055494 | 43953016 | C | T | 1.5776833035174098e-08 | 5.652894305129718 | 32175.0 |
| 17 | rs62055496 | 43953307 | C | T | 1.5776833035174098e-08 | 5.652894305129718 | 32175.0 |
| 17 | rs62055497 | 43953529 | A | G | 1.5776833035174098e-08 | 5.652894305129718 | 32175.0 |
| 17 | rs113952577 | 44010661 | A | G | 1.578255593575453e-08 | 5.652831991047036 | 32175.0 |
| 17 | rs17564871 | 44011018 | A | G | 1.578255593575453e-08 | 5.652831991047036 | 32175.0 |
| 17 | rs62060800 | 44145592 | A | G | 1.5795120116126962e-08 | 5.652695262299527 | 32175.0 |
| 17 | rs62060801 | 44145860 | A | G | 1.5795120116126962e-08 | 5.652695262299527 | 32175.0 |
| 17 | rs17659743 | 44146165 | C | G | 1.5795120116126962e-08 | 5.652695262299527 | 32175.0 |
| 17 | rs62060802 | 44146336 | A | C | 1.5795120116126962e-08 | 5.652695262299527 | 32175.0 |
| 17 | rs75975083 | 44146357 | T | C | 1.5795120116126962e-08 | 5.652695262299527 | 32175.0 |
| 17 | rs77950913 | 44146451 | T | C | 1.5795120116126962e-08 | 5.652695262299527 | 32175.0 |
| 17 | rs79782347 | 44146567 | T | C | 1.5795120116126962e-08 | 5.652695262299527 | 32175.0 |
| 17 | rs147317628 | 44277476 | C | CTTT AG | 1.580396539591499e-08 | 5.652599067554778 | 31685.0 |
| 17 | rs76159289 | 44098861 | G | T | 1.586188540758542e-08 | 5.651970461495058 | 32186.0 |
| 17 | rs10445337 | 44067400 | T | C | 1.5949071794986193e-08 | 5.651028418588432 | 32186.0 |
| 17 | rs79406732 | 43940497 | C | T | 1.5950138624160674e-08 | 5.651016922575581 | 32148.0 |
| 17 | rs62061727 | 44017124 | G | T | 1.5971152568881555e-08 | 5.650790631135135 | 32186.0 |
| 17 | rs2532239 | 44266227 | T | C | 1.597761953287015e-08 | 5.6507210489673625 | 32038.0 |
| 17 | rs17649641 | 43997372 | T | C | 1.60194073006468e-08 | 5.6502720862161375 | 32170.0 |
| 17 | rs146122400 | 44782224 | A | AGT | 1.6028996400598114e-08 | 5.6501692225445925 | 28928.0 |
| 17 | rs55960528 | 43968994 | T | C | 1.6048756396278798e-08 | 5.649957442535808 | 32170.0 |
| 17 | rs55877243 | 43969019 | T | C | 1.6048756396278798e-08 | 5.649957442535808 | 32170.0 |
| 17 | 17:44279557_TA_T | 44279557 | TA | T | 1.60544064964506e-08 | 5.649896933501612 | 32024.0 |
| 17 | rs76402917 | 44126477 | G | A | 1.6106252284915896e-08 | 5.649342661229283 | 32108.0 |
| 17 | rs62061710 | 44007727 | C | T | 1.6142403389254844e-08 | 5.648957201942871 | 32181.0 |
| 17 | rs62061711 | 44007764 | T | C | 1.6142403389254844e-08 | 5.648957201942871 | 32181.0 |
| 17 | 17:44027836_GC_G | 44027836 | GC | G | 1.615888785845711e-08 | 5.648781715373413 | 32155.0 |
| 17 | rs55719714 | 43939638 | G | A | 1.619925910331536e-08 | 5.6483526737219325 | 32152.0 |
| 17 | rs56299858 | 43939659 | G | A | 1.619925910331536e-08 | 5.6483526737219325 | 32152.0 |
| 17 | 17:44177304_TA_T | 44177304 | TA | T | 1.620729958474991e-08 | 5.64826734827466 | 31914.0 |
| 17 | rs62055948 | 43852621 | T | C | 1.622487680616186e-08 | 5.648080962194964 | 32095.0 |
| 17 | rs767058 | 43998767 | T | C | 1.6232986886003778e-08 | 5.647995030296722 | 32156.0 |
| 17 | rs62055486 | 43949092 | T | G | 1.6235100728731368e-08 | 5.647972639523085 | 32173.0 |
| 17 | rs796654785 | 44284962 | A | AG | 1.623647201431751e-08 | 5.6479581157630445 | 31910.0 |
| 17 | rs62063849 | 44073062 | T | G | 1.625459507484153e-08 | 5.647766279993343 | 32152.0 |
| 17 | rs111372048 | 44136577 | A | C | 1.627039029984253e-08 | 5.647599254115653 | 32174.0 |
| 17 | rs17575822 | 44144345 | A | G | 1.627039029984253e-08 | 5.647599254115653 | 32174.0 |
| 17 | rs17575850 | 44144387 | C | A | 1.627039029984253e-08 | 5.647599254115653 | 32174.0 |
| 17 | rs17659731 | 44144634 | G | A | 1.627039029984253e-08 | 5.647599254115653 | 32174.0 |
| 17 | rs75641221 | 44145373 | T | C | 1.627039029984253e-08 | 5.647599254115653 | 32174.0 |
| 17 | rs55876817 | 43958139 | A | C | 1.6341080297847714e-08 | 5.646853670320683 | 32155.0 |
| 17 | rs76475392 | 44243351 | T | TCG | 1.6379073763134556e-08 | 5.646454237654327 | 32075.0 |
| 17 | rs72245049 | 44243354 | A | AG | 1.6379073763134556e-08 | 5.646454237654327 | 32075.0 |
| 17 | rs2316773 | 43946536 | C | T | 1.6380514611336868e-08 | 5.646439107453866 | 32173.0 |
| 17 | rs17577650 | 44205500 | A | G | 1.639441076516199e-08 | 5.646293251665482 | 32104.0 |
| 17 | rs62063200 | 44206482 | C | T | 1.639441076516199e-08 | 5.646293251665482 | 32104.0 |
| 17 | rs10514903 | 44206646 | C | G | 1.639441076516199e-08 | 5.646293251665482 | 32104.0 |
| 17 | rs17661428 | 44208144 | C | G | 1.639441076516199e-08 | 5.646293251665482 | 32104.0 |
| 17 | rs17577877 | 44208218 | A | G | 1.639441076516199e-08 | 5.646293251665482 | 32104.0 |
| 17 | rs75676226 | 43947171 | G | A | 1.6403527380979158e-08 | 5.6461976277234776 | 31836.0 |

| | | | | | | | |
|---|---|---|---|---|---|---|---|
| 17 | rs75229280 | 44196653 | G | A | 1.6404335422150246e-08 | 5.646189154691317 | 32105.0 |
| 17 | rs17660251 | 44166646 | C | T | 1.6409184489090754e-08 | 5.646138316414134 | 32173.0 |
| 17 | rs113246445 | 43851209 | A | AT | 1.6421197753907394e-08 | 5.646012430534867 | 32157.0 |
| 17 | rs147319425 | 44003133 | A | AAAG | 1.6432988373213812e-08 | 5.645888964677875 | 32180.0 |
| 17 | rs113909426 | 44196822 | G | A | 1.644345660958568e-08 | 5.645779418277972 | 32107.0 |
| 17 | rs62062274 | 44093319 | G | A | 1.649636026160344e-08 | 5.64522683422776 | 32186.0 |
| 17 | rs138939613 | 43841624 | G | GATGGATGA | 1.654430832282164e-08 | 5.644727496617726 | 32123.0 |
| 17 | rs62062769 | 44019643 | G | A | 1.661043824904894e-08 | 5.644041111640657 | 32182.0 |
| 17 | rs62062770 | 44019680 | T | C | 1.661043824904894e-08 | 5.644041111640657 | 32182.0 |
| 17 | rs17426195 | 43832367 | G | A | 1.6611431584547413e-08 | 5.644030821718526 | 32160.0 |
| 17 | rs2696456 | 44283761 | A | T | 1.664125875655792e-08 | 5.643722121303632 | 31990.0 |
| 17 | rs62056837 | 43988884 | T | C | 1.6643483274856193e-08 | 5.6436991198831175 | 32174.0 |
| 17 | rs62064422 | 43668017 | G | A | 1.6686919721310232e-08 | 5.643250586274861 | 31737.0 |
| 17 | 17:43914979_AC_A | 43914979 | AC | A | 1.6687389276927867e-08 | 5.6432457437454255 | 32032.0 |
| 17 | rs2732630 | 44289150 | A | C | 1.6721584235246092e-08 | 5.642893446214223 | 32034.0 |
| 17 | rs55944735 | 43958172 | A | G | 1.6723995747974846e-08 | 5.642868627741804 | 32148.0 |
| 17 | rs17649571 | 43994952 | A | G | 1.6734881206496987e-08 | 5.642756641523667 | 32177.0 |
| 17 | rs112310745 | 44082432 | G | A | 1.674718719253342e-08 | 5.642630126520812 | 31927.0 |
| 17 | rs62062783 | 44023520 | T | G | 1.6791184328379537e-08 | 5.642178539361438 | 32177.0 |
| 17 | rs34232924 | 44023828 | C | CGTG | 1.6791184328379537e-08 | 5.642178539361438 | 32177.0 |
| 17 | rs35163442 | 44023842 | G | A | 1.6791184328379537e-08 | 5.642178539361438 | 32177.0 |
| 17 | rs3785884 | 44057595 | G | A | 1.6822866179420575e-08 | 5.64185406755858 | 32160.0 |
| 17 | rs75257002 | 43843395 | C | T | 1.685757051576501e-08 | 5.641499321132812 | 32091.0 |
| 17 | rs55721252 | 43958099 | T | C | 1.690250584143878e-08 | 5.641041046144041 | 32158.0 |
| 17 | rs62055888 | 43838720 | C | T | 1.6916241320605208e-08 | 5.640901200316831 | 31975.0 |
| 17 | rs138190012 | 44197417 | T | TGAA | 1.6920167753531742e-08 | 5.640861244165512 | 32028.0 |
| 17 | rs2532305 | 44223060 | G | A | 1.6962463221676373e-08 | 5.640431407199746 | 32090.0 |
| 17 | rs1918797 | 44223243 | G | A | 1.6962463221676373e-08 | 5.640431407199746 | 32090.0 |
| 17 | rs1918796 | 44223387 | T | C | 1.6962463221676373e-08 | 5.640431407199746 | 32090.0 |
| 17 | rs1918795 | 44224686 | A | T | 1.6962463221676373e-08 | 5.640431407199746 | 32090.0 |
| 17 | rs1406071 | 44225485 | T | C | 1.6962463221676373e-08 | 5.640431407199746 | 32090.0 |
| 17 | rs1406070 | 44225802 | C | T | 1.6962463221676373e-08 | 5.640431407199746 | 32090.0 |
| 17 | rs2696585 | 44226030 | G | A | 1.6962463221676373e-08 | 5.640431407199746 | 32090.0 |
| 17 | rs2532304 | 44226126 | A | C | 1.6962463221676373e-08 | 5.640431407199746 | 32090.0 |
| 17 | rs367820416 | 43982069 | T | G | 1.7030172111224256e-08 | 5.639745462887162 | 32186.0 |
| 17 | rs62056803 | 43982159 | A | G | 1.7030172111224256e-08 | 5.639745462887162 | 32186.0 |
| 17 | rs572543580 | 43914981 | C | G | 1.7044368887318875e-08 | 5.639601974300356 | 32033.0 |
| 17 | rs9897621 | 44113392 | T | A | 1.705168145707174e-08 | 5.6395281105287145 | 30255.0 |
| 17 | rs56042957 | 43992244 | G | A | 1.712810750627347e-08 | 5.638757971735994 | 32102.0 |
| 17 | rs55810795 | 43992283 | A | G | 1.712810750627347e-08 | 5.638757971735994 | 32102.0 |
| 17 | rs62055893 | 43842494 | A | G | 1.7151848427921437e-08 | 5.638519415734012 | 32139.0 |
| 17 | rs62055477 | 43945687 | C | A | 1.7153597130754428e-08 | 5.6385018569178875 | 32180.0 |
| 17 | rs78637438 | 43945855 | G | A | 1.7153597130754428e-08 | 5.6385018569178875 | 32180.0 |
| 17 | rs2316771 | 43945931 | T | C | 1.7153597130754428e-08 | 5.6385018569178875 | 32180.0 |
| 17 | rs62055478 | 43945938 | C | T | 1.7153597130754428e-08 | 5.6385018569178875 | 32180.0 |
| 17 | rs12150170 | 44090646 | A | G | 1.7175138596374174e-08 | 5.638285700425402 | 32185.0 |
| 17 | rs62062266 | 44090704 | A | G | 1.7175138596374174e-08 | 5.638285700425402 | 32185.0 |
| 17 | rs17573858 | 44090796 | T | C | 1.7175138596374174e-08 | 5.638285700425402 | 32185.0 |
| 17 | rs62062267 | 44090854 | C | T | 1.7175138596374174e-08 | 5.638285700425402 | 32185.0 |
| 17 | rs62062268 | 44090858 | C | T | 1.7175138596374174e-08 | 5.638285700425402 | 32185.0 |
| 17 | rs62062269 | 44091092 | G | C | 1.7175138596374174e-08 | 5.638285700425402 | 32185.0 |
| 17 | 17:44163547_ACG_A | 44163547 | ACG | A | 1.7175621622093454e-08 | 5.63828085655253 | 32166.0 |
| 17 | rs111362359 | 43954416 | A | AC | 1.7209466550353988e-08 | 5.6379417821939 | 32125.0 |
| 17 | rs78834738 | 44022434 | G | A | 1.7231258534628826e-08 | 5.6377238024285194 | 32035.0 |

| | | | | | | | |
|---|---|---|---|---|---|---|---|
| 17 | rs55900347 | 44176092 | G | A | 1.7247923419902477e-08 | 5.63755728831928 | 32109.0 |
| 17 | rs1815 | 44276578 | T | C | 1.7289563660488493e-08 | 5.637141904552935 | 31827.0 |
| 17 | rs62062781 | 44022632 | G | A | 1.732168042233269e-08 | 5.636822185701022 | 32160.0 |
| 17 | 17:44303564_C_T | 44303564 | C | T | 1.733111974265017e-08 | 5.636728327756746 | 31022.0 |
| 17 | rs2696601 | 44216158 | T | C | 1.7387372817817178e-08 | 5.636170014200799 | 32181.0 |
| 17 | rs2696587 | 44222460 | T | G | 1.7399598834126198e-08 | 5.63604890267097 | 32091.0 |
| 17 | rs1467963 | 43978934 | T | G | 1.7414036377790996e-08 | 5.635905990011746 | 32185.0 |
| 17 | rs62056800 | 43979468 | T | A | 1.7414036377790996e-08 | 5.635905990011746 | 32185.0 |
| 17 | rs62056801 | 43979504 | G | A | 1.7414036377790996e-08 | 5.635905990011746 | 32185.0 |
| 17 | rs113834264 | 43979622 | G | A | 1.7414036377790996e-08 | 5.635905990011746 | 32185.0 |
| 17 | rs2099151 | 43979822 | A | T | 1.7414036377790996e-08 | 5.635905990011746 | 32185.0 |
| 17 | rs2082068 | 43979971 | C | T | 1.7414036377790996e-08 | 5.635905990011746 | 32185.0 |
| 17 | rs1984937 | 43980530 | A | C | 1.7414036377790996e-08 | 5.635905990011746 | 32185.0 |
| 17 | rs62056802 | 43980762 | A | G | 1.7414036377790996e-08 | 5.635905990011746 | 32185.0 |
| 17 | rs113589236 | 43981795 | G | A | 1.7414036377790996e-08 | 5.635905990011746 | 32185.0 |
| 17 | rs113029914 | 43981831 | A | T | 1.7414036377790996e-08 | 5.635905990011746 | 32185.0 |
| 17 | rs112275277 | 43981958 | C | T | 1.7414036377790996e-08 | 5.635905990011746 | 32185.0 |
| 17 | rs62056804 | 43982286 | C | T | 1.7414036377790996e-08 | 5.635905990011746 | 32185.0 |
| 17 | rs75617973 | 43983159 | T | C | 1.7414036377790996e-08 | 5.635905990011746 | 32185.0 |
| 17 | rs62056806 | 43983801 | A | G | 1.7414036377790996e-08 | 5.635905990011746 | 32185.0 |
| 17 | rs62056807 | 43984253 | G | C | 1.7414036377790996e-08 | 5.635905990011746 | 32185.0 |
| 17 | rs1467966 | 43984399 | C | T | 1.7414036377790996e-08 | 5.635905990011746 | 32185.0 |
| 17 | rs79492906 | 43984864 | C | T | 1.7414036377790996e-08 | 5.635905990011746 | 32185.0 |
| 17 | rs62056811 | 43985247 | C | T | 1.7414036377790996e-08 | 5.635905990011746 | 32185.0 |
| 17 | rs74509629 | 43985597 | C | T | 1.7414036377790996e-08 | 5.635905990011746 | 32185.0 |
| 17 | rs79290980 | 43985662 | T | C | 1.7414036377790996e-08 | 5.635905990011746 | 32185.0 |
| 17 | rs17649700 | 43997901 | G | C | 1.7422361653633985e-08 | 5.635823633045801 | 32174.0 |
| 17 | rs62058962 | 43998151 | C | T | 1.7422361653633985e-08 | 5.635823633045801 | 32174.0 |
| 17 | rs1467969 | 43998328 | C | T | 1.7422361653633985e-08 | 5.635823633045801 | 32174.0 |
| 17 | rs1467970 | 43998574 | T | G | 1.7422361653633985e-08 | 5.635823633045801 | 32174.0 |
| 17 | rs767057 | 43998822 | A | G | 1.7422361653633985e-08 | 5.635823633045801 | 32174.0 |
| 17 | rs17649518 | 43994386 | C | A | 1.743179351684591e-08 | 5.635730375435699 | 32156.0 |
| 17 | rs62055937 | 43848968 | G | T | 1.7474237499830987e-08 | 5.635311315761906 | 32091.0 |
| 17 | rs112535292 | 44344300 | G | T | 1.7476816839613408e-08 | 5.6352858811884365 | 30814.0 |
| 17 | rs146453014 | 44172493 | A | AT | 1.748154659903445e-08 | 5.635239251043206 | 32104.0 |
| 17 | rs2532262 | 44252510 | T | C | 1.7485540230205692e-08 | 5.63519988783904 | 32000.0 |
| 17 | rs2668668 | 44284641 | T | G | 1.7495497482224488e-08 | 5.635101782245767 | 31975.0 |
| 17 | rs62061726 | 44016341 | G | A | 1.750170821264075e-08 | 5.635040617375357 | 32185.0 |
| 17 | rs77561671 | 43976627 | G | T | 1.7508597887566464e-08 | 5.634972790740013 | 32185.0 |
| 17 | rs62060768 | 44134984 | G | A | 1.7540127432221545e-08 | 5.634662722848171 | 32179.0 |
| 17 | rs1122380 | 44206079 | C | T | 1.754635369997522e-08 | 5.634601556477394 | 32100.0 |
| 17 | rs62055515 | 43956370 | G | T | 1.7559182941920917e-08 | 5.634475589431555 | 32169.0 |
| 17 | rs62055516 | 43956390 | T | C | 1.7559182941920917e-08 | 5.634475589431555 | 32169.0 |
| 17 | rs62055517 | 43956425 | C | T | 1.7559182941920917e-08 | 5.634475589431555 | 32169.0 |
| 17 | rs62055518 | 43956505 | A | C | 1.7559182941920917e-08 | 5.634475589431555 | 32169.0 |
| 17 | rs56364632 | 43956642 | T | C | 1.7559182941920917e-08 | 5.634475589431555 | 32169.0 |
| 17 | rs55768605 | 43956734 | C | A | 1.7559182941920917e-08 | 5.634475589431555 | 32169.0 |
| 17 | rs535791904 | 43958207 | C | CAAAAAAAT | 1.7568070113343078e-08 | 5.634388380957434 | 32171.0 |
| 17 | rs55780786 | 44209875 | T | C | 1.757609704295297e-08 | 5.634309650720012 | 32106.0 |
| 17 | rs3089 | 44209919 | C | G | 1.757609704295297e-08 | 5.634309650720012 | 32106.0 |
| 17 | rs113260576 | 44193908 | C | CCAACAA | 1.765699868539633e-08 | 5.633518089782053 | 32182.0 |
| 17 | rs17573266 | 44072881 | C | G | 1.766723790285813e-08 | 5.633418158024321 | 32182.0 |
| 17 | rs62062782 | 44023087 | G | A | 1.770470713891394e-08 | 5.6330529481280625 | 32176.0 |
| 17 | rs1476554 | 44159102 | C | T | 1.7717587672235054e-08 | 5.6329275758485675 | 32186.0 |

| | | | | | | | |
|---|---|---|---|---|---|---|---|
| 17 | rs145503157 | 43859929 | C | CACG | 1.771908166179805e-08 | 5.632913039875293 | 32139.0 |
| 17 | rs17564983 | 44011825 | A | C | 1.7720575774907004e-08 | 5.632898503890213 | 32174.0 |
| 17 | 17:44007773_CAAAA_C | 44007773 | CAAAA | C | 1.7739636550946084e-08 | 5.632713169045707 | 32175.0 |
| 17 | rs2732669 | 44284542 | G | A | 1.7746742763866594e-08 | 5.632644122240447 | 32013.0 |
| 17 | rs111555239 | 44006453 | C | CT | 1.775796878438311e-08 | 5.632535100426687 | 32174.0 |
| 17 | rs112791677 | 44001549 | G | GC | 1.7774321327612724e-08 | 5.632376411933286 | 32182.0 |
| 17 | rs55895732 | 43968937 | T | C | 1.7780191869506874e-08 | 5.632319477550741 | 32184.0 |
| 17 | rs41384744 | 44137070 | A | G | 1.77845023072101e-08 | 5.632277685175954 | 32177.0 |
| 17 | rs1560310 | 43978534 | G | A | 1.7799565646333607e-08 | 5.632131713941879 | 32186.0 |
| 17 | rs56371647 | 43855503 | C | T | 1.7825095593688503e-08 | 5.6318845898029695 | 32133.0 |
| 17 | rs62063851 | 44074435 | C | G | 1.7839691302145328e-08 | 5.631743461006758 | 32165.0 |
| 17 | rs55967721 | 43948977 | G | A | 1.784044332973342e-08 | 5.631736192540767 | 32095.0 |
| 17 | rs77924424 | 44012915 | T | C | 1.7895676838661088e-08 | 5.6312031636569895 | 32180.0 |
| 17 | rs76475191 | 44290849 | T | A | 1.7904856785074718e-08 | 5.631114727784213 | 32064.0 |
| 17 | rs56338487 | 43960812 | A | T | 1.7910895430667437e-08 | 5.631056577931129 | 32186.0 |
| 17 | rs62055538 | 43960997 | T | C | 1.7910895430667437e-08 | 5.631056577931129 | 32186.0 |
| 17 | rs62055539 | 43961044 | A | G | 1.7910895430667437e-08 | 5.631056577931129 | 32186.0 |
| 17 | rs62055540 | 43961142 | T | C | 1.7910895430667437e-08 | 5.631056577931129 | 32186.0 |
| 17 | rs17770108 | 43961292 | G | T | 1.7910895430667437e-08 | 5.631056577931129 | 32186.0 |
| 17 | rs62055541 | 43962017 | G | C | 1.7910895430667437e-08 | 5.631056577931129 | 32186.0 |
| 17 | rs77129180 | 43963046 | G | A | 1.7910895430667437e-08 | 5.631056577931129 | 32186.0 |
| 17 | rs80254317 | 43963282 | C | G | 1.7910895430667437e-08 | 5.631056577931129 | 32186.0 |
| 17 | rs112425426 | 43963416 | C | G | 1.7910895430667437e-08 | 5.631056577931129 | 32186.0 |
| 17 | rs62055542 | 43963844 | C | T | 1.7910895430667437e-08 | 5.631056577931129 | 32186.0 |
| 17 | rs17770120 | 43963943 | C | A | 1.7910895430667437e-08 | 5.631056577931129 | 32186.0 |
| 17 | rs17770150 | 43964163 | G | A | 1.7910895430667437e-08 | 5.631056577931129 | 32186.0 |
| 17 | rs17691449 | 43964283 | G | C | 1.7910895430667437e-08 | 5.631056577931129 | 32186.0 |
| 17 | rs62055548 | 43964899 | A | G | 1.7910895430667437e-08 | 5.631056577931129 | 32186.0 |
| 17 | rs113327263 | 43965063 | T | C | 1.7910895430667437e-08 | 5.631056577931129 | 32186.0 |
| 17 | rs62055549 | 43965595 | G | C | 1.7910895430667437e-08 | 5.631056577931129 | 32186.0 |
| 17 | rs62055552 | 43965955 | A | G | 1.7910895430667437e-08 | 5.631056577931129 | 32186.0 |
| 17 | rs62055553 | 43966178 | C | A | 1.7910895430667437e-08 | 5.631056577931129 | 32186.0 |
| 17 | rs62055554 | 43966260 | A | G | 1.7910895430667437e-08 | 5.631056577931129 | 32186.0 |
| 17 | rs62055555 | 43966282 | G | T | 1.7910895430667437e-08 | 5.631056577931129 | 32186.0 |
| 17 | rs111941646 | 43966608 | A | T | 1.7910895430667437e-08 | 5.631056577931129 | 32186.0 |
| 17 | rs62055556 | 43966749 | T | C | 1.7910895430667437e-08 | 5.631056577931129 | 32186.0 |
| 17 | rs62055557 | 43966795 | C | A | 1.7910895430667437e-08 | 5.631056577931129 | 32186.0 |
| 17 | rs62055558 | 43966955 | A | G | 1.7910895430667437e-08 | 5.631056577931129 | 32186.0 |
| 17 | rs62055559 | 43967138 | T | G | 1.7910895430667437e-08 | 5.631056577931129 | 32186.0 |
| 17 | rs17691466 | 43968219 | A | G | 1.7910895430667437e-08 | 5.631056577931129 | 32186.0 |
| 17 | rs17691508 | 43968463 | C | G | 1.7910895430667437e-08 | 5.631056577931129 | 32186.0 |
| 17 | rs55943606 | 43968874 | A | G | 1.7910895430667437e-08 | 5.631056577931129 | 32186.0 |
| 17 | rs112297913 | 43969258 | G | GC | 1.7910895430667437e-08 | 5.631056577931129 | 32186.0 |
| 17 | rs17691610 | 43970662 | G | T | 1.7910895430667437e-08 | 5.631056577931129 | 32186.0 |
| 17 | rs76594404 | 43971457 | G | C | 1.7910895430667437e-08 | 5.631056577931129 | 32186.0 |
| 17 | rs80233201 | 43971481 | T | C | 1.7910895430667437e-08 | 5.631056577931129 | 32186.0 |
| 17 | rs62056778 | 43971604 | C | G | 1.7910895430667437e-08 | 5.631056577931129 | 32186.0 |
| 17 | rs11575895 | 43971785 | A | G | 1.7910895430667437e-08 | 5.631056577931129 | 32186.0 |
| 17 | rs62056779 | 43971937 | C | A | 1.7910895430667437e-08 | 5.631056577931129 | 32186.0 |
| 17 | rs11575896 | 43972142 | G | A | 1.7910895430667437e-08 | 5.631056577931129 | 32186.0 |
| 17 | rs111972148 | 43973121 | G | C | 1.7910895430667437e-08 | 5.631056577931129 | 32186.0 |
| 17 | rs76324150 | 43973233 | C | T | 1.7910895430667437e-08 | 5.631056577931129 | 32186.0 |
| 17 | rs113347741 | 43973408 | C | T | 1.7910895430667437e-08 | 5.631056577931129 | 32186.0 |
| 17 | rs78720789 | 43973413 | A | G | 1.7910895430667437e-08 | 5.631056577931129 | 32186.0 |

| | | | | | | | |
|---|---|---|---|---|---|---|---|
| 17 | rs62056780 | 43973652 | C | T | 1.7910895430667437e-08 | 5.631056577931129 | 32186.0 |
| 17 | rs62056781 | 43973899 | T | C | 1.7910895430667437e-08 | 5.631056577931129 | 32186.0 |
| 17 | rs74496580 | 43974065 | C | A | 1.7910895430667437e-08 | 5.631056577931129 | 32186.0 |
| 17 | rs62056782 | 43974230 | C | T | 1.7910895430667437e-08 | 5.631056577931129 | 32186.0 |
| 17 | rs80346216 | 43974476 | G | T | 1.7910895430667437e-08 | 5.631056577931129 | 32186.0 |
| 17 | rs76632685 | 43974477 | A | T | 1.7910895430667437e-08 | 5.631056577931129 | 32186.0 |
| 17 | rs62056783 | 43974549 | G | C | 1.7910895430667437e-08 | 5.631056577931129 | 32186.0 |
| 17 | rs62056784 | 43974889 | C | G | 1.7910895430667437e-08 | 5.631056577931129 | 32186.0 |
| 17 | rs62056785 | 43975263 | T | C | 1.7910895430667437e-08 | 5.631056577931129 | 32186.0 |
| 17 | rs62056786 | 43975285 | T | C | 1.7910895430667437e-08 | 5.631056577931129 | 32186.0 |
| 17 | rs62056789 | 43975415 | A | C | 1.7910895430667437e-08 | 5.631056577931129 | 32186.0 |
| 17 | rs62056790 | 43975417 | G | A | 1.7910895430667437e-08 | 5.631056577931129 | 32186.0 |
| 17 | rs62056791 | 43976064 | T | C | 1.7910895430667437e-08 | 5.631056577931129 | 32186.0 |
| 17 | rs62056792 | 43976644 | G | A | 1.7910895430667437e-08 | 5.631056577931129 | 32186.0 |
| 17 | rs62056793 | 43977018 | A | G | 1.7910895430667437e-08 | 5.631056577931129 | 32186.0 |
| 17 | rs62056794 | 43977049 | A | G | 1.7910895430667437e-08 | 5.631056577931129 | 32186.0 |
| 17 | rs1864324 | 43977702 | C | T | 1.7910895430667437e-08 | 5.631056577931129 | 32186.0 |
| 17 | rs1864325 | 43977827 | C | T | 1.7910895430667437e-08 | 5.631056577931129 | 32186.0 |
| 17 | rs62056795 | 43978041 | C | T | 1.7910895430667437e-08 | 5.631056577931129 | 32186.0 |
| 17 | rs62056796 | 43978120 | A | G | 1.7910895430667437e-08 | 5.631056577931129 | 32186.0 |
| 17 | rs62059007 | 44005615 | A | G | 1.7917376615223474e-08 | 5.6309941877745135 | 32182.0 |
| 17 | rs113139571 | 44099856 | G | GC | 1.7930534825883564e-08 | 5.630867589603918 | 32162.0 |
| 17 | rs12150336 | 44125704 | C | T | 1.7937148986130492e-08 | 5.6308039873142635 | 32175.0 |
| 17 | rs35134656 | 44103927 | C | CT | 1.7962367713696416e-08 | 5.63056169080707 | 32168.0 |
| 17 | rs34043286 | 44117119 | A | G | 1.797044504782219e-08 | 5.6304841552320575 | 32186.0 |
| 17 | rs113518470 | 43850932 | C | T | 1.7971581208575324e-08 | 5.63047325176488 | 32155.0 |
| 17 | rs12150515 | 44090685 | G | T | 1.7975937144094745e-08 | 5.630431455079186 | 32184.0 |
| 17 | rs754512 | 44055647 | A | T | 1.798408362717603e-08 | 5.63035331318761 | 32174.0 |
| 17 | rs62060762 | 44133396 | G | A | 1.801847763417664e-08 | 5.630023780837887 | 32182.0 |
| 17 | rs62060763 | 44133403 | T | C | 1.801847763417664e-08 | 5.630023780837887 | 32182.0 |
| 17 | rs75944932 | 44072423 | A | C | 1.8041780737843905e-08 | 5.6298008584552655 | 32179.0 |
| 17 | 17:44012257_CG_C | 44012257 | CG | C | 1.8045012574997114e-08 | 5.629769964101674 | 32158.0 |
| 17 | 17:44250669_TA_T | 44250669 | TA | T | 1.8048118208213343e-08 | 5.629740281241128 | 32001.0 |
| 17 | rs10445370 | 43935118 | G | A | 1.8095274032255358e-08 | 5.629290186119532 | 32180.0 |
| 17 | rs56347602 | 43939255 | T | C | 1.811193112930293e-08 | 5.629131468855238 | 32180.0 |
| 17 | rs62061730 | 44017342 | A | G | 1.811193112930293e-08 | 5.629131468855238 | 32178.0 |
| 17 | rs1076222 | 44109769 | C | G | 1.814350636491146e-08 | 5.6288309926228415 | 32175.0 |
| 17 | rs111459873 | 44113787 | T | TG | 1.815593479385414e-08 | 5.6287128604646375 | 32185.0 |
| 17 | rs62060844 | 44153836 | C | T | 1.8160780955131198e-08 | 5.628666819002093 | 32165.0 |
| 17 | rs74825921 | 44153909 | A | G | 1.8176092951851927e-08 | 5.628521424132329 | 32185.0 |
| 17 | 17:43849572_AT_A | 43849572 | AT | A | 1.8213277140201245e-08 | 5.628168836668515 | 32143.0 |
| 17 | rs62060796 | 44143029 | G | T | 1.8215963419373868e-08 | 5.62814339194367 | 32183.0 |
| 17 | rs55894109 | 43957221 | C | T | 1.8217882428089383e-08 | 5.628125217118069 | 32185.0 |
| 17 | rs56249780 | 43957328 | C | T | 1.8217882428089383e-08 | 5.628125217118069 | 32185.0 |
| 17 | rs56387391 | 43959261 | G | A | 1.8217882428089383e-08 | 5.628125217118069 | 32185.0 |
| 17 | rs2019816 | 43959661 | G | A | 1.8217882428089383e-08 | 5.628125217118069 | 32185.0 |
| 17 | rs2006305 | 43959752 | A | T | 1.8217882428089383e-08 | 5.628125217118069 | 32185.0 |
| 17 | rs2019822 | 43959918 | T | A | 1.8217882428089383e-08 | 5.628125217118069 | 32185.0 |
| 17 | rs62055529 | 43960020 | A | G | 1.8217882428089383e-08 | 5.628125217118069 | 32185.0 |
| 17 | rs79580365 | 43960172 | C | T | 1.8217882428089383e-08 | 5.628125217118069 | 32185.0 |
| 17 | rs55937610 | 43960283 | A | G | 1.8217882428089383e-08 | 5.628125217118069 | 32185.0 |
| 17 | rs56194412 | 43960600 | T | C | 1.8217882428089383e-08 | 5.628125217118069 | 32185.0 |
| 17 | rs79412431 | 43940021 | G | A | 1.8241438406225568e-08 | 5.627902271089016 | 31659.0 |
| 17 | rs12150570 | 44126218 | G | A | 1.825284304586086e-08 | 5.6277944320676605 | 32178.0 |

| | | | | | | | |
|---|---|---|---|---|---|---|---|
| 17 | 17:44168295_CTTA_C | 44168295 | CTTA | C | 1.8265344918702985e-08 | 5.6276762930681885 | 32032.0 |
| 17 | rs369585371 | 44215416 | A | AG | 1.828388878168477e-08 | 5.627501203578409 | 32132.0 |
| 17 | rs111321973 | 43953170 | C | T | 1.828440236913505e-08 | 5.627496356786163 | 32183.0 |
| 17 | rs112950348 | 44033188 | G | A | 1.8291144515970435e-08 | 5.627432742516301 | 31709.0 |
| 17 | rs41382552 | 44118999 | A | G | 1.82980818390558e-08 | 5.627367310460005 | 32177.0 |
| 17 | rs77768866 | 44121917 | C | T | 1.82980818390558e-08 | 5.627367310460005 | 32177.0 |
| 17 | rs112281640 | 44081218 | C | CT | 1.8314214660058785e-08 | 5.6272152405904325 | 32010.0 |
| 17 | rs79765413 | 44022433 | T | C | 1.831640105485157e-08 | 5.627194641385243 | 32036.0 |
| 17 | rs62055488 | 43950394 | C | T | 1.832218980611719e-08 | 5.6271401139629456 | 32183.0 |
| 17 | rs62055490 | 43950533 | A | G | 1.832218980611719e-08 | 5.6271401139629456 | 32183.0 |
| 17 | rs10491140 | 43950976 | G | T | 1.832218980611719e-08 | 5.6271401139629456 | 32183.0 |
| 17 | rs62055491 | 43951120 | T | C | 1.832218980611719e-08 | 5.6271401139629456 | 32183.0 |
| 17 | rs34416056 | 43951220 | T | G | 1.832218980611719e-08 | 5.6271401139629456 | 32183.0 |
| 17 | rs62055492 | 43951345 | C | A | 1.832218980611719e-08 | 5.6271401139629456 | 32183.0 |
| 17 | 17:44045574_AC_A | 44045574 | AC | A | 1.8343430581064684e-08 | 5.62694017866005 | 32179.0 |
| 17 | rs62061712 | 44009876 | C | A | 1.8356186635108443e-08 | 5.626820216406343 | 32184.0 |
| 17 | rs7501759 | 44009921 | A | G | 1.8356186635108443e-08 | 5.626820216406343 | 32184.0 |
| 17 | rs2316782 | 44010040 | A | G | 1.8356186635108443e-08 | 5.626820216406343 | 32184.0 |
| 17 | rs2316783 | 44010118 | G | C | 1.8356186635108443e-08 | 5.626820216406343 | 32184.0 |
| 17 | rs4255816 | 44010364 | C | G | 1.8356186635108443e-08 | 5.626820216406343 | 32184.0 |
| 17 | rs62061713 | 44010405 | A | G | 1.8356186635108443e-08 | 5.626820216406343 | 32184.0 |
| 17 | rs62061714 | 44010452 | C | A | 1.8356186635108443e-08 | 5.626820216406343 | 32184.0 |
| 17 | rs78077519 | 44010984 | G | A | 1.8356186635108443e-08 | 5.626820216406343 | 32184.0 |
| 17 | rs17650258 | 44011073 | G | A | 1.8356186635108443e-08 | 5.626820216406343 | 32184.0 |
| 17 | rs62061715 | 44011526 | A | G | 1.8356186635108443e-08 | 5.626820216406343 | 32184.0 |
| 17 | rs17564948 | 44011749 | A | G | 1.8356186635108443e-08 | 5.626820216406343 | 32184.0 |
| 17 | rs62061716 | 44012096 | A | G | 1.8356186635108443e-08 | 5.626820216406343 | 32184.0 |
| 17 | rs17650335 | 44012343 | G | A | 1.8356186635108443e-08 | 5.626820216406343 | 32184.0 |
| 17 | rs17650417 | 44013103 | G | T | 1.8356186635108443e-08 | 5.626820216406343 | 32184.0 |
| 17 | rs62061720 | 44013189 | C | T | 1.8356186635108443e-08 | 5.626820216406343 | 32184.0 |
| 17 | rs12150111 | 44013938 | A | G | 1.8356186635108443e-08 | 5.626820216406343 | 32184.0 |
| 17 | rs12150460 | 44014137 | G | T | 1.8356186635108443e-08 | 5.626820216406343 | 32184.0 |
| 17 | rs12150195 | 44014263 | T | C | 1.8356186635108443e-08 | 5.626820216406343 | 32184.0 |
| 17 | rs55874169 | 44014384 | G | A | 1.8356186635108443e-08 | 5.626820216406343 | 32184.0 |
| 17 | rs62063856 | 44076466 | A | G | 1.83700477573754e-08 | 5.626689953452677 | 32146.0 |
| 17 | rs80348388 | 43952634 | A | T | 1.8389535135268136e-08 | 5.62650697784233 | 32184.0 |
| 17 | rs62055493 | 43952944 | T | C | 1.8389535135268136e-08 | 5.62650697784233 | 32184.0 |
| 17 | rs62055495 | 43953043 | A | C | 1.8389535135268136e-08 | 5.62650697784233 | 32184.0 |
| 17 | rs62055498 | 43953814 | G | A | 1.8389535135268136e-08 | 5.62650697784233 | 32184.0 |
| 17 | rs62055499 | 43953991 | C | G | 1.8389535135268136e-08 | 5.62650697784233 | 32184.0 |
| 17 | rs62055500 | 43954134 | G | A | 1.8389535135268136e-08 | 5.62650697784233 | 32184.0 |
| 17 | rs62055501 | 43954227 | G | A | 1.8389535135268136e-08 | 5.62650697784233 | 32184.0 |
| 17 | rs62055502 | 43954284 | G | A | 1.8389535135268136e-08 | 5.62650697784233 | 32184.0 |
| 17 | rs916793 | 43954686 | G | A | 1.8389535135268136e-08 | 5.62650697784233 | 32184.0 |
| 17 | rs4441322 | 43955030 | A | G | 1.8389535135268136e-08 | 5.62650697784233 | 32184.0 |
| 17 | rs113414067 | 43955093 | T | C | 1.8389535135268136e-08 | 5.62650697784233 | 32184.0 |
| 17 | rs55905252 | 43955365 | C | A | 1.8389535135268136e-08 | 5.62650697784233 | 32184.0 |
| 17 | 17:43848646_AG_A | 43848646 | AG | A | 1.839631529652097e-08 | 5.6264433600561485 | 31962.0 |
| 17 | rs78681971 | 44106219 | C | T | 1.8409689081199895e-08 | 5.626317941472553 | 32181.0 |
| 17 | rs74548327 | 43973080 | A | G | 1.841570069675768e-08 | 5.626261593706863 | 32184.0 |
| 17 | rs62064664 | 44081462 | T | C | 1.841932152285706e-08 | 5.626227663783895 | 32180.0 |
| 17 | rs62063673 | 44125484 | T | G | 1.8439378014487505e-08 | 5.626039836261129 | 32177.0 |
| 17 | rs2158072 | 43950195 | T | C | 1.844222647739435e-08 | 5.626013176711074 | 32182.0 |
| 17 | rs2040845 | 43767188 | G | C | 1.84429386605418e-08 | 5.626006511817354 | 31947.0 |

| | | | | | | | |
|---|---|---|---|---|---|---|---|
| 17 | rs62062768 | 44019479 | C | G | 1.8472680175585146e-08 | 5.625728401767181 | 32182.0 |
| 17 | rs62061719 | 44012989 | C | T | 1.847832275616277e-08 | 5.625675687610405 | 32183.0 |
| 17 | rs17650381 | 44013024 | T | C | 1.847832275616277e-08 | 5.625675687610405 | 32183.0 |
| 17 | rs62061721 | 44013346 | T | C | 1.847832275616277e-08 | 5.625675687610405 | 32183.0 |
| 17 | rs56318865 | 44147411 | G | A | 1.851396839060105e-08 | 5.625343039523072 | 32185.0 |
| 17 | rs56041778 | 44147511 | G | C | 1.851396839060105e-08 | 5.625343039523072 | 32185.0 |
| 17 | rs55947022 | 44147574 | G | A | 1.851396839060105e-08 | 5.625343039523072 | 32185.0 |
| 17 | rs55929190 | 44147599 | G | C | 1.851396839060105e-08 | 5.625343039523072 | 32185.0 |
| 17 | rs56323408 | 44147721 | C | T | 1.851396839060105e-08 | 5.625343039523072 | 32185.0 |
| 17 | rs12150125 | 43986432 | G | A | 1.855033259128316e-08 | 5.6250043259200275 | 32121.0 |
| 17 | rs62060794 | 44142566 | C | T | 1.8568182475249623e-08 | 5.624838299380213 | 32183.0 |
| 17 | rs10445336 | 43934948 | A | G | 1.8577635395850848e-08 | 5.624750437996989 | 32171.0 |
| 17 | rs55836887 | 43668815 | A | G | 1.8614710239923808e-08 | 5.6244062595973165 | 31812.0 |
| 17 | rs62062323 | 44113224 | C | T | 1.862013393913532e-08 | 5.6243559653685296 | 32184.0 |
| 17 | rs9896485 | 44050828 | G | C | 1.8620330005472286e-08 | 5.624354147502191 | 31249.0 |
| 17 | 17:44221042_C_T | 44221042 | C | T | 1.862170252651901e-08 | 5.624341422432661 | 32158.0 |
| 17 | rs56040418 | 43958079 | C | T | 1.8629808959205825e-08 | 5.624266283742441 | 32172.0 |
| 17 | rs17564153 | 43995098 | G | A | 1.8642891143266973e-08 | 5.624145091641967 | 32171.0 |
| 17 | rs62063305 | 44050282 | G | A | 1.864544321857753e-08 | 5.624121459086771 | 32185.0 |
| 17 | rs17651134 | 44050340 | G | A | 1.864544321857753e-08 | 5.624121459086771 | 32185.0 |
| 17 | rs62063306 | 44050395 | C | A | 1.864544321857753e-08 | 5.624121459086771 | 32185.0 |
| 17 | rs62063774 | 44050463 | T | C | 1.864544321857753e-08 | 5.624121459086771 | 32185.0 |
| 17 | rs62063775 | 44050823 | G | T | 1.864544321857753e-08 | 5.624121459086771 | 32185.0 |
| 17 | rs62063776 | 44050850 | A | G | 1.864544321857753e-08 | 5.624121459086771 | 32185.0 |
| 17 | rs1800547 | 44051846 | A | G | 1.864544321857753e-08 | 5.624121459086771 | 32185.0 |
| 17 | rs17651213 | 44051924 | G | A | 1.864544321857753e-08 | 5.624121459086771 | 32185.0 |
| 17 | rs17572361 | 44052009 | T | C | 1.864544321857753e-08 | 5.624121459086771 | 32185.0 |
| 17 | rs17651243 | 44052284 | G | A | 1.864544321857753e-08 | 5.624121459086771 | 32185.0 |
| 17 | rs62063777 | 44053171 | A | G | 1.864544321857753e-08 | 5.624121459086771 | 32185.0 |
| 17 | rs2217394 | 44053448 | A | G | 1.864544321857753e-08 | 5.624121459086771 | 32185.0 |
| 17 | rs62063778 | 44053989 | G | A | 1.864544321857753e-08 | 5.624121459086771 | 32185.0 |
| 17 | rs17651285 | 44054237 | G | A | 1.864544321857753e-08 | 5.624121459086771 | 32185.0 |
| 17 | rs112578465 | 44125066 | C | T | 1.865722646795108e-08 | 5.624012385350765 | 32176.0 |
| 17 | rs62063680 | 44131559 | C | A | 1.866010792830025e-08 | 5.623985722780873 | 32181.0 |
| 17 | rs62060850 | 44157982 | A | T | 1.8664627460828654e-08 | 5.623943910943562 | 32184.0 |
| 17 | rs62060851 | 44157992 | T | C | 1.8664627460828654e-08 | 5.623943910943562 | 32184.0 |
| 17 | rs77138434 | 44159555 | C | A | 1.8664627460828654e-08 | 5.623943910943562 | 32184.0 |
| 17 | rs111913701 | 44159631 | T | G | 1.8664627460828654e-08 | 5.623943910943562 | 32184.0 |
| 17 | rs111519055 | 44159672 | A | G | 1.8664627460828654e-08 | 5.623943910943562 | 32184.0 |
| 17 | rs77925835 | 44159695 | C | T | 1.8664627460828654e-08 | 5.623943910943562 | 32184.0 |
| 17 | rs17659953 | 44159725 | C | T | 1.8664627460828654e-08 | 5.623943910943562 | 32184.0 |
| 17 | rs17576165 | 44159849 | T | C | 1.8664627460828654e-08 | 5.623943910943562 | 32184.0 |
| 17 | rs17576200 | 44159988 | G | A | 1.8664627460828654e-08 | 5.623943910943562 | 32184.0 |
| 17 | rs62060852 | 44160050 | G | A | 1.8664627460828654e-08 | 5.623943910943562 | 32184.0 |
| 17 | rs62060853 | 44160592 | G | A | 1.8664627460828654e-08 | 5.623943910943562 | 32184.0 |
| 17 | rs17660017 | 44160674 | G | C | 1.8664627460828654e-08 | 5.623943910943562 | 32184.0 |
| 17 | rs55987700 | 44161146 | T | G | 1.8664627460828654e-08 | 5.623943910943562 | 32184.0 |
| 17 | rs113788190 | 44161302 | A | G | 1.8664627460828654e-08 | 5.623943910943562 | 32184.0 |
| 17 | rs112364920 | 44161360 | A | T | 1.8664627460828654e-08 | 5.623943910943562 | 32184.0 |
| 17 | 17:44161478_CTCACACCTGTAATCCCAGCACTTTGGGAAGCTGAGGCGGGTGGA_C | 44161478 | CTCACACCTGTAATCCCAGCACTTTGGGAAGCTGAGGCGGGTGGA | C | 1.8664627460828654e-08 | 5.623943910943562 | 32184.0 |
| 17 | rs113955100 | 44161522 | A | C | 1.8664627460828654e-08 | 5.623943910943562 | 32184.0 |
| 17 | rs62060855 | 44161860 | C | T | 1.8664627460828654e-08 | 5.623943910943562 | 32184.0 |

| | | | | | | | |
|---|---|---|---|---|---|---|---|
| 17 | rs62060856 | 44161875 | A | C | 1.8664627460828654e-08 | 5.623943910943562 | 32184.0 |
| 17 | rs2316955 | 44161985 | T | G | 1.8664627460828654e-08 | 5.623943910943562 | 32184.0 |
| 17 | rs17660065 | 44162284 | T | C | 1.8664627460828654e-08 | 5.623943910943562 | 32184.0 |
| 17 | rs55864131 | 44163411 | T | C | 1.8664627460828654e-08 | 5.623943910943562 | 32184.0 |
| 17 | 17:44163551_AGT_A | 44163551 | AGT | A | 1.8664627460828654e-08 | 5.623943910943562 | 32184.0 |
| 17 | 17:44179168_AACC_A | 44179168 | AACC | A | 1.8664627460828654e-08 | 5.623943910943562 | 32184.0 |
| 17 | rs62056840 | 43989689 | T | C | 1.8711195191868134e-08 | 5.623513667524141 | 32174.0 |
| 17 | rs17770296 | 43969136 | C | T | 1.8721113715652386e-08 | 5.623422163716851 | 32179.0 |
| 17 | rs62061858 | 44200033 | T | C | 1.872144223176493e-08 | 5.623419133781455 | 32083.0 |
| 17 | rs55653657 | 43956778 | A | G | 1.875300611349402e-08 | 5.623128257595542 | 32166.0 |
| 17 | rs12150527 | 43986462 | A | G | 1.876327560670554e-08 | 5.623033721817364 | 32168.0 |
| 17 | 17:44104410_TCTC_T | 44104410 | TCTC | T | 1.8833003425946265e-08 | 5.62239316816773 | 32180.0 |
| 17 | rs78938131 | 44157622 | T | G | 1.883472178328147e-08 | 5.622377411592308 | 32185.0 |
| 17 | 17:44157676_CT_C | 44157676 | CT | C | 1.883472178328147e-08 | 5.622377411592308 | 32185.0 |
| 17 | rs62055547 | 43964755 | C | T | 1.8837101301106704e-08 | 5.622355594772682 | 32184.0 |
| 17 | rs17691556 | 43969168 | T | A | 1.8853303073623486e-08 | 5.622207118488356 | 32181.0 |
| 17 | rs12150576 | 44015624 | G | A | 1.888117559214277e-08 | 5.621951978813066 | 32185.0 |
| 17 | rs12150242 | 44015816 | A | G | 1.888117559214277e-08 | 5.621951978813066 | 32185.0 |
| 17 | rs12150104 | 44015929 | G | A | 1.888117559214277e-08 | 5.621951978813066 | 32185.0 |
| 17 | rs62061725 | 44016157 | G | C | 1.888117559214277e-08 | 5.621951978813066 | 32185.0 |
| 17 | rs113537106 | 44016462 | A | G | 1.888117559214277e-08 | 5.621951978813066 | 32185.0 |
| 17 | rs62061728 | 44017180 | T | G | 1.888117559214277e-08 | 5.621951978813066 | 32185.0 |
| 17 | rs62061731 | 44017666 | C | A | 1.888117559214277e-08 | 5.621951978813066 | 32185.0 |
| 17 | rs62061732 | 44017941 | G | T | 1.888117559214277e-08 | 5.621951978813066 | 32185.0 |
| 17 | rs62062791 | 44025273 | G | A | 1.8881374353589016e-08 | 5.621950160702569 | 32182.0 |
| 17 | rs62062792 | 44025347 | T | C | 1.8881374353589016e-08 | 5.621950160702569 | 32182.0 |
| 17 | rs78026984 | 44025592 | G | C | 1.8881374353589016e-08 | 5.621950160702569 | 32182.0 |
| 17 | rs113395365 | 44025650 | T | C | 1.8881374353589016e-08 | 5.621950160702569 | 32182.0 |
| 17 | rs62063681 | 44131592 | A | G | 1.8882235676899308e-08 | 5.6219422822216085 | 32182.0 |
| 17 | rs62063682 | 44131622 | G | A | 1.8882235676899308e-08 | 5.6219422822216085 | 32182.0 |
| 17 | rs62063683 | 44131717 | G | A | 1.8882235676899308e-08 | 5.6219422822216085 | 32182.0 |
| 17 | rs79180862 | 44131798 | C | T | 1.8882235676899308e-08 | 5.6219422822216085 | 32182.0 |
| 17 | rs55711631 | 43958431 | T | C | 1.8884687346446983e-08 | 5.621919858833741 | 32180.0 |
| 17 | rs55886302 | 43936011 | A | G | 1.8888862529085026e-08 | 5.621881678405983 | 32181.0 |
| 17 | rs56136075 | 43936298 | C | A | 1.8888862529085026e-08 | 5.621881678405983 | 32181.0 |
| 17 | rs76563578 | 43933879 | C | G | 1.890517429304776e-08 | 5.6217325921463015 | 32159.0 |
| 17 | 17:43934587_GT_G | 43934587 | GT | G | 1.891857883326624e-08 | 5.621610170793235 | 32175.0 |
| 17 | rs62056760 | 43968572 | G | A | 1.892355813878035e-08 | 5.621564717107423 | 32178.0 |
| 17 | 17:44133818_TA_T | 44133818 | TA | T | 1.8937041906624586e-08 | 5.621441688552141 | 32167.0 |
| 17 | rs74457229 | 43972581 | C | T | 1.8949138811566547e-08 | 5.6213313863698735 | 32186.0 |
| 17 | rs757055 | 43946875 | C | T | 1.8960711069626185e-08 | 5.6212259319996285 | 32170.0 |
| 17 | 17:44290910_A_G | 44290910 | A | G | 1.900000194852798e-08 | 5.620868352211257 | 31936.0 |
| 17 | 17:44293963_T_A | 44293963 | T | A | 1.900266862499195e-08 | 5.620844109255179 | 32062.0 |
| 17 | rs62061729 | 44017283 | A | G | 1.900720281730497e-08 | 5.620802896154489 | 32184.0 |
| 17 | rs62062293 | 44100553 | C | A | 1.9018476214117096e-08 | 5.620700469066808 | 32182.0 |
| 17 | rs7687 | 44103296 | T | C | 1.902434898094589e-08 | 5.620647134019864 | 32185.0 |
| 17 | rs17652748 | 44103616 | C | T | 1.902434898094589e-08 | 5.620647134019864 | 32185.0 |
| 17 | rs2158257 | 44104343 | A | C | 1.902434898094589e-08 | 5.620647134019864 | 32185.0 |
| 17 | rs17574228 | 44104509 | T | C | 1.902434898094589e-08 | 5.620647134019864 | 32185.0 |
| 17 | rs17574361 | 44108202 | A | G | 1.902434898094589e-08 | 5.620647134019864 | 32185.0 |
| 17 | rs17652961 | 44108355 | G | A | 1.902434898094589e-08 | 5.620647134019864 | 32185.0 |
| 17 | rs17574425 | 44109188 | C | G | 1.902434898094589e-08 | 5.620647134019864 | 32185.0 |
| 17 | rs55881134 | 44109248 | G | A | 1.902434898094589e-08 | 5.620647134019864 | 32185.0 |
| 17 | rs1467968 | 43986396 | C | T | 1.9047256403058693e-08 | 5.620439247035908 | 32172.0 |

| | | | | | | | |
|---|---|---|---|---|---|---|---|
| 17 | rs4371195 | 43986420 | A | G | 1.9047256403058693e-08 | 5.620439247035908 | 32172.0 |
| 17 | rs17585012 | 44227510 | A | T | 1.905648042870355e-08 | 5.6203556066936216 | 32113.0 |
| 17 | rs147091683 | 44101775 | A | AT | 1.9065575058303763e-08 | 5.62027317814784 | 32170.0 |
| 17 | rs78229689 | 44106806 | G | A | 1.906805006892691e-08 | 5.620250752668966 | 32183.0 |
| 17 | rs733969 | 44089727 | G | A | 1.9071395178441992e-08 | 5.620220447923149 | 32185.0 |
| 17 | rs62063684 | 44132269 | T | C | 1.9078957245938833e-08 | 5.620151959008602 | 32184.0 |
| 17 | rs62063685 | 44132379 | A | G | 1.9078957245938833e-08 | 5.620151959008602 | 32184.0 |
| 17 | rs62063687 | 44132592 | A | G | 1.9078957245938833e-08 | 5.620151959008602 | 32184.0 |
| 17 | rs62060760 | 44132887 | C | T | 1.9078957245938833e-08 | 5.620151959008602 | 32184.0 |
| 17 | rs17575437 | 44133031 | T | A | 1.9078957245938833e-08 | 5.620151959008602 | 32184.0 |
| 17 | rs17653889 | 44133070 | A | T | 1.9078957245938833e-08 | 5.620151959008602 | 32184.0 |
| 17 | rs62060761 | 44133162 | A | C | 1.9078957245938833e-08 | 5.620151959008602 | 32184.0 |
| 17 | rs111259120 | 44133534 | C | T | 1.9078957245938833e-08 | 5.620151959008602 | 32184.0 |
| 17 | rs62060764 | 44133633 | A | G | 1.9078957245938833e-08 | 5.620151959008602 | 32184.0 |
| 17 | rs62060765 | 44133648 | A | C | 1.9078957245938833e-08 | 5.620151959008602 | 32184.0 |
| 17 | rs17575507 | 44134095 | A | G | 1.9078957245938833e-08 | 5.620151959008602 | 32184.0 |
| 17 | rs876944 | 44134391 | G | T | 1.9078957245938833e-08 | 5.620151959008602 | 32184.0 |
| 17 | rs62060767 | 44134910 | G | A | 1.9078957245938833e-08 | 5.620151959008602 | 32184.0 |
| 17 | rs62060769 | 44135026 | G | C | 1.9078957245938833e-08 | 5.620151959008602 | 32184.0 |
| 17 | rs1107820 | 44135359 | T | C | 1.9078957245938833e-08 | 5.620151959008602 | 32184.0 |
| 17 | rs17575556 | 44135827 | G | A | 1.9078957245938833e-08 | 5.620151959008602 | 32184.0 |
| 17 | rs17653998 | 44135985 | A | G | 1.9078957245938833e-08 | 5.620151959008602 | 32184.0 |
| 17 | rs62060785 | 44136634 | G | A | 1.9078957245938833e-08 | 5.620151959008602 | 32184.0 |
| 17 | rs17654016 | 44137925 | T | C | 1.9078957245938833e-08 | 5.620151959008602 | 32184.0 |
| 17 | rs2316951 | 44139083 | G | C | 1.9078957245938833e-08 | 5.620151959008602 | 32184.0 |
| 17 | rs2316952 | 44139190 | G | C | 1.9078957245938833e-08 | 5.620151959008602 | 32184.0 |
| 17 | rs17575683 | 44139662 | C | T | 1.9078957245938833e-08 | 5.620151959008602 | 32184.0 |
| 17 | rs112411928 | 44139762 | C | G | 1.9078957245938833e-08 | 5.620151959008602 | 32184.0 |
| 17 | rs2316953 | 44140341 | G | A | 1.9078957245938833e-08 | 5.620151959008602 | 32184.0 |
| 17 | rs2838 | 44141347 | A | G | 1.9078957245938833e-08 | 5.620151959008602 | 32184.0 |
| 17 | rs12150127 | 44141732 | G | A | 1.9078957245938833e-08 | 5.620151959008602 | 32184.0 |
| 17 | rs62060792 | 44141955 | T | A | 1.9078957245938833e-08 | 5.620151959008602 | 32184.0 |
| 17 | rs76307183 | 44142121 | G | A | 1.9078957245938833e-08 | 5.620151959008602 | 32184.0 |
| 17 | rs62060793 | 44142427 | A | G | 1.9078957245938833e-08 | 5.620151959008602 | 32184.0 |
| 17 | rs62060795 | 44142663 | C | T | 1.9078957245938833e-08 | 5.620151959008602 | 32184.0 |
| 17 | rs17575773 | 44142879 | C | T | 1.9078957245938833e-08 | 5.620151959008602 | 32184.0 |
| 17 | rs62060797 | 44143344 | G | A | 1.9078957245938833e-08 | 5.620151959008602 | 32184.0 |
| 17 | rs12150344 | 44143484 | G | A | 1.9078957245938833e-08 | 5.620151959008602 | 32184.0 |
| 17 | rs62060798 | 44144152 | T | C | 1.9078957245938833e-08 | 5.620151959008602 | 32184.0 |
| 17 | rs62060799 | 44144251 | G | T | 1.9078957245938833e-08 | 5.620151959008602 | 32184.0 |
| 17 | 17:44092729_TAAA_T | 44092729 | TAAA | T | 1.9093956231335654e-08 | 5.620016192598079 | 31794.0 |
| 17 | rs12150506 | 44090536 | G | A | 1.9108564560327418e-08 | 5.61988406179908 | 32185.0 |
| 17 | rs17572467 | 44054388 | T | C | 1.9111514359379523e-08 | 5.6198573930791484 | 32186.0 |
| 17 | rs17572495 | 44054596 | T | G | 1.9111514359379523e-08 | 5.6198573930791484 | 32186.0 |
| 17 | rs62063779 | 44054671 | C | T | 1.9111514359379523e-08 | 5.6198573930791484 | 32186.0 |
| 17 | rs62063780 | 44056238 | A | G | 1.9111514359379523e-08 | 5.6198573930791484 | 32186.0 |
| 17 | rs2163129 | 44056433 | T | C | 1.9111514359379523e-08 | 5.6198573930791484 | 32186.0 |
| 17 | rs2163130 | 44056434 | G | A | 1.9111514359379523e-08 | 5.6198573930791484 | 32186.0 |
| 17 | rs1981997 | 44056767 | G | A | 1.9111514359379523e-08 | 5.6198573930791484 | 32186.0 |
| 17 | rs1981998 | 44056833 | G | A | 1.9111514359379523e-08 | 5.6198573930791484 | 32186.0 |
| 17 | rs17572627 | 44058629 | T | A | 1.9111514359379523e-08 | 5.6198573930791484 | 32186.0 |
| 17 | rs62063781 | 44058691 | G | T | 1.9111514359379523e-08 | 5.6198573930791484 | 32186.0 |
| 17 | rs17651483 | 44058861 | C | A | 1.9111514359379523e-08 | 5.6198573930791484 | 32186.0 |
| 17 | rs62063784 | 44059775 | A | G | 1.9111514359379523e-08 | 5.6198573930791484 | 32186.0 |

| | | | | | | | |
|---|---|---|---|---|---|---|---|
| 17 | rs56234850 | 44060248 | G | A | 1.9111514359379523e-08 | 5.6198573930791484 | 32186.0 |
| 17 | rs63750417 | 44060775 | C | T | 1.9111514359379523e-08 | 5.6198573930791484 | 32186.0 |
| 17 | rs62063787 | 44061036 | T | C | 1.9111514359379523e-08 | 5.6198573930791484 | 32186.0 |
| 17 | rs62063788 | 44061696 | C | T | 1.9111514359379523e-08 | 5.6198573930791484 | 32186.0 |
| 17 | rs56363731 | 44062337 | G | A | 1.9111514359379523e-08 | 5.6198573930791484 | 32186.0 |
| 17 | rs62063790 | 44062915 | G | T | 1.9111514359379523e-08 | 5.6198573930791484 | 32186.0 |
| 17 | rs62063791 | 44062993 | A | G | 1.9111514359379523e-08 | 5.6198573930791484 | 32186.0 |
| 17 | rs1529534 | 44063140 | A | G | 1.9111514359379523e-08 | 5.6198573930791484 | 32186.0 |
| 17 | rs17572795 | 44063393 | G | A | 1.9111514359379523e-08 | 5.6198573930791484 | 32186.0 |
| 17 | rs17572823 | 44063563 | T | C | 1.9111514359379523e-08 | 5.6198573930791484 | 32186.0 |
| 17 | rs17572851 | 44063766 | A | G | 1.9111514359379523e-08 | 5.6198573930791484 | 32186.0 |
| 17 | rs62063792 | 44063787 | A | G | 1.9111514359379523e-08 | 5.6198573930791484 | 32186.0 |
| 17 | rs62063793 | 44064334 | T | C | 1.9111514359379523e-08 | 5.6198573930791484 | 32186.0 |
| 17 | rs1529535 | 44064759 | A | C | 1.9111514359379523e-08 | 5.6198573930791484 | 32186.0 |
| 17 | rs919461 | 44065669 | A | T | 1.9111514359379523e-08 | 5.6198573930791484 | 32186.0 |
| 17 | rs919462 | 44065740 | C | T | 1.9111514359379523e-08 | 5.6198573930791484 | 32186.0 |
| 17 | rs62063794 | 44066088 | T | A | 1.9111514359379523e-08 | 5.6198573930791484 | 32186.0 |
| 17 | rs112385572 | 44066172 | A | G | 1.9111514359379523e-08 | 5.6198573930791484 | 32186.0 |
| 17 | rs17651700 | 44066634 | C | G | 1.9111514359379523e-08 | 5.6198573930791484 | 32186.0 |
| 17 | rs372810927 | 44066868 | C | A | 1.9111514359379523e-08 | 5.6198573930791484 | 32186.0 |
| 17 | rs113134013 | 44066995 | C | T | 1.9111514359379523e-08 | 5.6198573930791484 | 32186.0 |
| 17 | rs76357066 | 44067046 | G | A | 1.9111514359379523e-08 | 5.6198573930791484 | 32186.0 |
| 17 | rs79447161 | 44067508 | A | G | 1.9111514359379523e-08 | 5.6198573930791484 | 32186.0 |
| 17 | rs17651754 | 44067546 | T | C | 1.9111514359379523e-08 | 5.6198573930791484 | 32186.0 |
| 17 | rs10445338 | 44067682 | G | A | 1.9111514359379523e-08 | 5.6198573930791484 | 32186.0 |
| 17 | rs111652694 | 44067831 | T | C | 1.9111514359379523e-08 | 5.6198573930791484 | 32186.0 |
| 17 | rs75743061 | 44068112 | C | G | 1.9111514359379523e-08 | 5.6198573930791484 | 32186.0 |
| 17 | rs62063795 | 44068238 | C | T | 1.9111514359379523e-08 | 5.6198573930791484 | 32186.0 |
| 17 | rs62063796 | 44068289 | G | A | 1.9111514359379523e-08 | 5.6198573930791484 | 32186.0 |
| 17 | rs62063797 | 44068294 | G | C | 1.9111514359379523e-08 | 5.6198573930791484 | 32186.0 |
| 17 | rs77290642 | 44068480 | T | C | 1.9111514359379523e-08 | 5.6198573930791484 | 32186.0 |
| 17 | rs62063798 | 44068543 | G | A | 1.9111514359379523e-08 | 5.6198573930791484 | 32186.0 |
| 17 | rs1052551 | 44068924 | G | A | 1.9111514359379523e-08 | 5.6198573930791484 | 32186.0 |
| 17 | rs62063799 | 44069180 | C | G | 1.9111514359379523e-08 | 5.6198573930791484 | 32186.0 |
| 17 | rs62063800 | 44069198 | C | T | 1.9111514359379523e-08 | 5.6198573930791484 | 32186.0 |
| 17 | rs62063801 | 44069376 | G | T | 1.9111514359379523e-08 | 5.6198573930791484 | 32186.0 |
| 17 | rs17651857 | 44069687 | A | G | 1.9111514359379523e-08 | 5.6198573930791484 | 32186.0 |
| 17 | rs74759276 | 44069768 | A | G | 1.9111514359379523e-08 | 5.6198573930791484 | 32186.0 |
| 17 | rs17651887 | 44069840 | G | A | 1.9111514359379523e-08 | 5.6198573930791484 | 32186.0 |
| 17 | rs78599197 | 44069970 | C | G | 1.9111514359379523e-08 | 5.6198573930791484 | 32186.0 |
| 17 | rs62063842 | 44070111 | G | C | 1.9111514359379523e-08 | 5.6198573930791484 | 32186.0 |
| 17 | rs62064674 | 44088772 | C | A | 1.9111514359379523e-08 | 5.6198573930791484 | 32186.0 |
| 17 | rs76723223 | 44089108 | T | C | 1.9111514359379523e-08 | 5.6198573930791484 | 32186.0 |
| 17 | rs733968 | 44089715 | C | T | 1.9111514359379523e-08 | 5.6198573930791484 | 32186.0 |
| 17 | rs4283261 | 44089921 | C | T | 1.9111514359379523e-08 | 5.6198573930791484 | 32186.0 |
| 17 | rs62062265 | 44090035 | C | G | 1.9111514359379523e-08 | 5.6198573930791484 | 32186.0 |
| 17 | rs75666751 | 44090148 | A | C | 1.9111514359379523e-08 | 5.6198573930791484 | 32186.0 |
| 17 | rs4306559 | 44090236 | T | C | 1.9111514359379523e-08 | 5.6198573930791484 | 32186.0 |
| 17 | rs747152 | 44090339 | T | C | 1.9111514359379523e-08 | 5.6198573930791484 | 32186.0 |
| 17 | rs62062270 | 44091195 | T | C | 1.9111514359379523e-08 | 5.6198573930791484 | 32186.0 |
| 17 | rs75534191 | 44091724 | G | A | 1.9111514359379523e-08 | 5.6198573930791484 | 32186.0 |
| 17 | rs62062272 | 44092108 | T | C | 1.9111514359379523e-08 | 5.6198573930791484 | 32186.0 |
| 17 | rs62062273 | 44092243 | C | T | 1.9111514359379523e-08 | 5.6198573930791484 | 32186.0 |
| 17 | rs62062275 | 44093496 | A | G | 1.9111514359379523e-08 | 5.6198573930791484 | 32186.0 |

| | | | | | | | |
|---|---|---|---|---|---|---|---|
| 17 | rs62062276 | 44093627 | A | G | 1.9111514359379523e-08 | 5.6198573930791484 | 32186.0 |
| 17 | rs62062278 | 44093860 | A | G | 1.9111514359379523e-08 | 5.6198573930791484 | 32186.0 |
| 17 | rs62062281 | 44094112 | A | G | 1.9111514359379523e-08 | 5.6198573930791484 | 32186.0 |
| 17 | rs62062282 | 44094374 | A | G | 1.9111514359379523e-08 | 5.6198573930791484 | 32186.0 |
| 17 | rs17652502 | 44094471 | G | A | 1.9111514359379523e-08 | 5.6198573930791484 | 32186.0 |
| 17 | rs2316948 | 44094667 | C | G | 1.9111514359379523e-08 | 5.6198573930791484 | 32186.0 |
| 17 | rs62062283 | 44095055 | C | A | 1.9111514359379523e-08 | 5.6198573930791484 | 32186.0 |
| 17 | rs111976319 | 44095335 | C | A | 1.9111514359379523e-08 | 5.6198573930791484 | 32186.0 |
| 17 | rs62062284 | 44095570 | G | A | 1.9111514359379523e-08 | 5.6198573930791484 | 32186.0 |
| 17 | rs62062285 | 44095661 | T | G | 1.9111514359379523e-08 | 5.6198573930791484 | 32186.0 |
| 17 | rs55726761 | 44099669 | G | A | 1.9111514359379523e-08 | 5.6198573930791484 | 32186.0 |
| 17 | rs62062292 | 44099809 | A | C | 1.9111514359379523e-08 | 5.6198573930791484 | 32186.0 |
| 17 | rs77747656 | 44099827 | C | G | 1.9111514359379523e-08 | 5.6198573930791484 | 32186.0 |
| 17 | rs62062294 | 44100666 | A | C | 1.9111514359379523e-08 | 5.6198573930791484 | 32186.0 |
| 17 | rs62062295 | 44100906 | A | G | 1.9111514359379523e-08 | 5.6198573930791484 | 32186.0 |
| 17 | rs62062296 | 44100916 | C | T | 1.9111514359379523e-08 | 5.6198573930791484 | 32186.0 |
| 17 | rs62062297 | 44101004 | A | G | 1.9111514359379523e-08 | 5.6198573930791484 | 32186.0 |
| 17 | rs7350928 | 44108100 | C | T | 1.9111514359379523e-08 | 5.6198573930791484 | 32186.0 |
| 17 | rs36076725 | 44110532 | G | A | 1.9111514359379523e-08 | 5.6198573930791484 | 32186.0 |
| 17 | rs35833914 | 44110541 | G | A | 1.9111514359379523e-08 | 5.6198573930791484 | 32186.0 |
| 17 | rs77009866 | 44110888 | A | T | 1.9111514359379523e-08 | 5.6198573930791484 | 32186.0 |
| 17 | rs62062321 | 44111109 | A | G | 1.9111514359379523e-08 | 5.6198573930791484 | 32186.0 |
| 17 | rs17574604 | 44111613 | A | G | 1.9111514359379523e-08 | 5.6198573930791484 | 32186.0 |
| 17 | rs17653162 | 44111827 | C | A | 1.9111514359379523e-08 | 5.6198573930791484 | 32186.0 |
| 17 | rs62062322 | 44112385 | A | G | 1.9111514359379523e-08 | 5.6198573930791484 | 32186.0 |
| 17 | rs17653193 | 44112441 | G | A | 1.9111514359379523e-08 | 5.6198573930791484 | 32186.0 |
| 17 | rs17653211 | 44112638 | T | C | 1.9111514359379523e-08 | 5.6198573930791484 | 32186.0 |
| 17 | rs17653255 | 44112743 | C | A | 1.9111514359379523e-08 | 5.6198573930791484 | 32186.0 |
| 17 | rs4608377 | 44113359 | T | C | 1.9111514359379523e-08 | 5.6198573930791484 | 32186.0 |
| 17 | rs12185225 | 44113491 | T | C | 1.9111514359379523e-08 | 5.6198573930791484 | 32186.0 |
| 17 | rs12150438 | 44113963 | C | T | 1.9111514359379523e-08 | 5.6198573930791484 | 32186.0 |
| 17 | rs12150162 | 44114218 | T | C | 1.9111514359379523e-08 | 5.6198573930791484 | 32186.0 |
| 17 | rs62062134 | 44114432 | A | C | 1.9111514359379523e-08 | 5.6198573930791484 | 32186.0 |
| 17 | rs17574796 | 44115074 | T | C | 1.9111514359379523e-08 | 5.6198573930791484 | 32186.0 |
| 17 | rs17574824 | 44115107 | C | T | 1.9111514359379523e-08 | 5.6198573930791484 | 32186.0 |
| 17 | rs12185243 | 44115351 | T | C | 1.9111514359379523e-08 | 5.6198573930791484 | 32186.0 |
| 17 | rs11079729 | 44115569 | C | A | 1.9111514359379523e-08 | 5.6198573930791484 | 32186.0 |
| 17 | rs12150542 | 44115730 | G | A | 1.9111514359379523e-08 | 5.6198573930791484 | 32186.0 |
| 17 | rs12150090 | 44115886 | C | T | 1.9111514359379523e-08 | 5.6198573930791484 | 32186.0 |
| 17 | rs41437445 | 44116238 | G | A | 1.9111514359379523e-08 | 5.6198573930791484 | 32186.0 |
| 17 | rs62062136 | 44116312 | T | C | 1.9111514359379523e-08 | 5.6198573930791484 | 32186.0 |
| 17 | rs10514897 | 44117397 | A | G | 1.9111514359379523e-08 | 5.6198573930791484 | 32186.0 |
| 17 | rs75546272 | 44130668 | T | A | 1.9111514359379523e-08 | 5.6198573930791484 | 32186.0 |
| 17 | rs62063686 | 44132458 | G | A | 1.9111514359379523e-08 | 5.6198573930791484 | 32186.0 |
| 17 | rs111990897 | 44138377 | GA | GAA | 1.9111514359379523e-08 | 5.6198573930791484 | 32186.0 |
| 17 | rs111990897 | 44138377 | G | GA | 1.9111514359379523e-08 | 5.6198573930791484 | 32186.0 |
| 17 | rs150514510 | 44142231 | T | TA | 1.9111514359379523e-08 | 5.6198573930791484 | 32186.0 |
| 17 | rs146149698 | 44142491 | C | CCA | 1.9111514359379523e-08 | 5.6198573930791484 | 32186.0 |
| 17 | rs55831516 | 44148267 | C | T | 1.9111514359379523e-08 | 5.6198573930791484 | 32186.0 |
| 17 | rs62060809 | 44148613 | G | A | 1.9111514359379523e-08 | 5.6198573930791484 | 32186.0 |
| 17 | rs62060810 | 44148674 | A | G | 1.9111514359379523e-08 | 5.6198573930791484 | 32186.0 |
| 17 | rs1117253 | 44149297 | A | C | 1.9111514359379523e-08 | 5.6198573930791484 | 32186.0 |
| 17 | rs12150625 | 44149448 | C | G | 1.9111514359379523e-08 | 5.6198573930791484 | 32186.0 |
| 17 | rs12150627 | 44149537 | C | G | 1.9111514359379523e-08 | 5.6198573930791484 | 32186.0 |

| | | | | | | | |
|---|---|---|---|---|---|---|---|
| 17 | rs12150628 | 44149581 | C | T | 1.9111514359379523e-08 | 5.6198573930791484 | 32186.0 |
| 17 | rs12150611 | 44149741 | G | A | 1.9111514359379523e-08 | 5.6198573930791484 | 32186.0 |
| 17 | rs74850776 | 44150042 | T | C | 1.9111514359379523e-08 | 5.6198573930791484 | 32186.0 |
| 17 | rs974295 | 44150152 | A | T | 1.9111514359379523e-08 | 5.6198573930791484 | 32186.0 |
| 17 | rs974293 | 44150233 | G | A | 1.9111514359379523e-08 | 5.6198573930791484 | 32186.0 |
| 17 | rs974292 | 44150464 | T | C | 1.9111514359379523e-08 | 5.6198573930791484 | 32186.0 |
| 17 | rs974291 | 44150480 | A | T | 1.9111514359379523e-08 | 5.6198573930791484 | 32186.0 |
| 17 | rs12150551 | 44150920 | T | C | 1.9111514359379523e-08 | 5.6198573930791484 | 32186.0 |
| 17 | rs12150558 | 44151156 | T | C | 1.9111514359379523e-08 | 5.6198573930791484 | 32186.0 |
| 17 | rs62060812 | 44151667 | A | G | 1.9111514359379523e-08 | 5.6198573930791484 | 32186.0 |
| 17 | rs62060834 | 44151697 | A | T | 1.9111514359379523e-08 | 5.6198573930791484 | 32186.0 |
| 17 | rs113989747 | 44151969 | C | CA | 1.9111514359379523e-08 | 5.6198573930791484 | 32186.0 |
| 17 | rs62060835 | 44152390 | C | T | 1.9111514359379523e-08 | 5.6198573930791484 | 32186.0 |
| 17 | rs62060836 | 44152447 | G | A | 1.9111514359379523e-08 | 5.6198573930791484 | 32186.0 |
| 17 | rs112597362 | 44152736 | G | GCT | 1.9111514359379523e-08 | 5.6198573930791484 | 32186.0 |
| 17 | rs62060837 | 44152760 | T | G | 1.9111514359379523e-08 | 5.6198573930791484 | 32186.0 |
| 17 | rs368572568 | 44152812 | T | CCCA AGTG AGGC ACCT GGGT | 1.9111514359379523e-08 | 5.6198573930791484 | 32186.0 |
| 17 | rs62060838 | 44153144 | A | G | 1.9111514359379523e-08 | 5.6198573930791484 | 32186.0 |
| 17 | rs62060839 | 44153227 | A | G | 1.9111514359379523e-08 | 5.6198573930791484 | 32186.0 |
| 17 | rs62060841 | 44153386 | A | T | 1.9111514359379523e-08 | 5.6198573930791484 | 32186.0 |
| 17 | rs62060842 | 44153607 | T | A | 1.9111514359379523e-08 | 5.6198573930791484 | 32186.0 |
| 17 | rs62060843 | 44153626 | C | T | 1.9111514359379523e-08 | 5.6198573930791484 | 32186.0 |
| 17 | rs76154201 | 44154033 | T | C | 1.9111514359379523e-08 | 5.6198573930791484 | 32186.0 |
| 17 | rs75530705 | 44154048 | C | T | 1.9111514359379523e-08 | 5.6198573930791484 | 32186.0 |
| 17 | rs112197756 | 44154105 | A | G | 1.9111514359379523e-08 | 5.6198573930791484 | 32186.0 |
| 17 | rs62060845 | 44154393 | A | G | 1.9111514359379523e-08 | 5.6198573930791484 | 32186.0 |
| 17 | rs62060846 | 44154461 | A | C | 1.9111514359379523e-08 | 5.6198573930791484 | 32186.0 |
| 17 | rs74481389 | 44154607 | T | C | 1.9111514359379523e-08 | 5.6198573930791484 | 32186.0 |
| 17 | rs62060847 | 44154946 | C | T | 1.9111514359379523e-08 | 5.6198573930791484 | 32186.0 |
| 17 | rs56100031 | 44155157 | C | T | 1.9111514359379523e-08 | 5.6198573930791484 | 32186.0 |
| 17 | rs55972730 | 44155707 | C | T | 1.9111514359379523e-08 | 5.6198573930791484 | 32186.0 |
| 17 | rs2316954 | 44156167 | G | A | 1.9111514359379523e-08 | 5.6198573930791484 | 32186.0 |
| 17 | rs56167560 | 43792411 | A | G | 1.9118823739493804e-08 | 5.619791327215463 | 32057.0 |
| 17 | rs55849973 | 43792418 | T | C | 1.9118823739493804e-08 | 5.619791327215463 | 32057.0 |
| 17 | rs113912415 | 44007785 | A | G | 1.912036643596991e-08 | 5.619777386680962 | 32172.0 |
| 17 | rs62054841 | 43937104 | G | A | 1.91266726383752e-08 | 5.6197204122097055 | 32182.0 |
| 17 | rs62054842 | 43937162 | C | T | 1.91266726383752e-08 | 5.6197204122097055 | 32182.0 |
| 17 | rs62054843 | 43937184 | T | C | 1.91266726383752e-08 | 5.6197204122097055 | 32182.0 |
| 17 | rs62054844 | 43937333 | T | G | 1.91266726383752e-08 | 5.6197204122097055 | 32182.0 |
| 17 | rs55888391 | 43938107 | A | G | 1.91266726383752e-08 | 5.6197204122097055 | 32182.0 |
| 17 | rs76329216 | 43938499 | G | A | 1.91266726383752e-08 | 5.6197204122097055 | 32182.0 |
| 17 | rs62054846 | 43938634 | A | G | 1.91266726383752e-08 | 5.6197204122097055 | 32182.0 |
| 17 | rs55646970 | 43938796 | A | G | 1.91266726383752e-08 | 5.6197204122097055 | 32182.0 |
| 17 | rs55673092 | 43938882 | G | A | 1.91266726383752e-08 | 5.6197204122097055 | 32182.0 |
| 17 | rs62056835 | 43988623 | G | A | 1.913861975881557e-08 | 5.619612523884325 | 32179.0 |
| 17 | rs2106784 | 43949892 | C | T | 1.9141842706753038e-08 | 5.619583430292 | 32181.0 |
| 17 | rs112454267 | 44005329 | G | A | 1.9156755765267813e-08 | 5.619448871812402 | 32179.0 |
| 17 | rs62056856 | 43992939 | T | C | 1.915944401391537e-08 | 5.619424626933757 | 32182.0 |
| 17 | rs739642 | 43946838 | C | A | 1.9175514009628392e-08 | 5.619279763099554 | 32182.0 |
| 17 | rs56398500 | 43947488 | A | G | 1.9175514009628392e-08 | 5.619279763099554 | 32182.0 |
| 17 | rs62055483 | 43948040 | C | T | 1.9175514009628392e-08 | 5.619279763099554 | 32182.0 |
| 17 | rs78238042 | 43948159 | G | A | 1.9175514009628392e-08 | 5.619279763099554 | 32182.0 |

| | | | | | | | |
|---|---|---|---|---|---|---|---|
| 17 | rs56327054 | 43943957 | G | C | 1.9177868465523206e-08 | 5.6192585486321125 | 32175.0 |
| 17 | rs733966 | 44089563 | C | T | 1.919004889041742e-08 | 5.6191488391278455 | 32184.0 |
| 17 | rs733967 | 44089690 | T | C | 1.919624295318227e-08 | 5.619093074923172 | 32185.0 |
| 17 | rs113796169 | 44005254 | T | A | 1.9197320381597044e-08 | 5.619083376782888 | 32180.0 |
| 17 | rs753235 | 43946223 | G | A | 1.9202506335282376e-08 | 5.6190367044092735 | 32133.0 |
| 17 | rs111739681 | 43898459 | C | T | 1.9206211428960207e-08 | 5.6190033669250425 | 32173.0 |
| 17 | rs35168513 | 44275172 | A | ACT | 1.9207761039635642e-08 | 5.618989425776863 | 31737.0 |
| 17 | rs62063671 | 44124837 | A | G | 1.924822900987232e-08 | 5.618625739814208 | 32177.0 |
| 17 | 17:44296522_T_C | 44296522 | T | C | 1.9255522118162867e-08 | 5.618560275556184 | 32071.0 |
| 17 | rs10445371 | 44065410 | G | A | 1.9264236882173603e-08 | 5.618482081823107 | 32185.0 |
| 17 | rs919464 | 44065901 | C | T | 1.9264236882173603e-08 | 5.618482081823107 | 32185.0 |
| 17 | rs62060858 | 44164090 | G | C | 1.9283436506755184e-08 | 5.618309933175943 | 32183.0 |
| 17 | rs62060859 | 44164133 | G | A | 1.9283436506755184e-08 | 5.618309933175943 | 32183.0 |
| 17 | rs55692232 | 44164323 | G | A | 1.9283436506755184e-08 | 5.618309933175943 | 32183.0 |
| 17 | rs3912061 | 44165098 | C | T | 1.9283436506755184e-08 | 5.618309933175943 | 32183.0 |
| 17 | rs74348235 | 44165852 | A | G | 1.9283436506755184e-08 | 5.618309933175943 | 32183.0 |
| 17 | rs17660167 | 44166311 | C | T | 1.9283436506755184e-08 | 5.618309933175943 | 32183.0 |
| 17 | rs17660294 | 44167101 | T | C | 1.9283436506755184e-08 | 5.618309933175943 | 32183.0 |
| 17 | rs62061766 | 44167464 | G | A | 1.9283436506755184e-08 | 5.618309933175943 | 32183.0 |
| 17 | rs79923630 | 44167661 | G | T | 1.9283436506755184e-08 | 5.618309933175943 | 32183.0 |
| 17 | rs62061767 | 44167729 | A | G | 1.9283436506755184e-08 | 5.618309933175943 | 32183.0 |
| 17 | rs111364350 | 44168508 | C | T | 1.9283436506755184e-08 | 5.618309933175943 | 32183.0 |
| 17 | rs62061770 | 44169377 | G | C | 1.9283436506755184e-08 | 5.618309933175943 | 32183.0 |
| 17 | rs111970616 | 44169581 | T | C | 1.9283436506755184e-08 | 5.618309933175943 | 32183.0 |
| 17 | rs17660337 | 44169605 | T | C | 1.9283436506755184e-08 | 5.618309933175943 | 32183.0 |
| 17 | rs62061788 | 44169759 | G | A | 1.9283436506755184e-08 | 5.618309933175943 | 32183.0 |
| 17 | rs62061789 | 44169769 | C | T | 1.9283436506755184e-08 | 5.618309933175943 | 32183.0 |
| 17 | rs17660398 | 44170612 | C | T | 1.9283436506755184e-08 | 5.618309933175943 | 32183.0 |
| 17 | rs62061792 | 44171554 | G | A | 1.929581803535456e-08 | 5.618199005529698 | 32030.0 |
| 17 | rs56289364 | 43947928 | A | G | 1.9303331940816704e-08 | 5.618131721212616 | 32181.0 |
| 17 | rs55673340 | 43948733 | C | T | 1.9303331940816704e-08 | 5.618131721212616 | 32181.0 |
| 17 | rs77092481 | 43949008 | C | T | 1.9303331940816704e-08 | 5.618131721212616 | 32181.0 |
| 17 | rs17577975 | 44210442 | C | T | 1.9307597866133944e-08 | 5.618093532703923 | 32077.0 |
| 17 | rs17573607 | 44083081 | G | A | 1.931125510932152e-08 | 5.6180607996316265 | 32164.0 |
| 17 | rs78136862 | 44085743 | T | C | 1.9312542078659842e-08 | 5.618049282425286 | 32184.0 |
| 17 | rs62064670 | 44085852 | T | A | 1.9312542078659842e-08 | 5.618049282425286 | 32184.0 |
| 17 | rs62064671 | 44086100 | T | C | 1.9312542078659842e-08 | 5.618049282425286 | 32184.0 |
| 17 | rs3912060 | 44165435 | C | T | 1.9319655794434408e-08 | 5.6179856345724035 | 32178.0 |
| 17 | rs17660132 | 44165803 | T | C | 1.9319655794434408e-08 | 5.6179856345724035 | 32178.0 |
| 17 | rs62061764 | 44166197 | A | G | 1.9319655794434408e-08 | 5.6179856345724035 | 32178.0 |
| 17 | rs12149995 | 44092801 | G | A | 1.9326433150640832e-08 | 5.6179250173592505 | 32182.0 |
| 17 | rs112746008 | 44126650 | C | T | 1.9334975938915028e-08 | 5.617848639378409 | 32184.0 |
| 17 | rs74977736 | 44126977 | G | C | 1.9334975938915028e-08 | 5.617848639378409 | 32184.0 |
| 17 | rs12150447 | 44128125 | A | C | 1.9334975938915028e-08 | 5.617848639378409 | 32184.0 |
| 17 | rs12150064 | 44128407 | C | A | 1.9334975938915028e-08 | 5.617848639378409 | 32184.0 |
| 17 | rs17653836 | 44128876 | T | A | 1.9334975938915028e-08 | 5.617848639378409 | 32184.0 |
| 17 | rs17575423 | 44129253 | T | C | 1.9334975938915028e-08 | 5.617848639378409 | 32184.0 |
| 17 | rs62063676 | 44129800 | G | A | 1.9334975938915028e-08 | 5.617848639378409 | 32184.0 |
| 17 | rs62063677 | 44129885 | C | T | 1.9334975938915028e-08 | 5.617848639378409 | 32184.0 |
| 17 | rs62063678 | 44129949 | C | G | 1.9334975938915028e-08 | 5.617848639378409 | 32184.0 |
| 17 | rs79649865 | 44130587 | T | G | 1.9334975938915028e-08 | 5.617848639378409 | 32184.0 |
| 17 | rs111853996 | 44130878 | A | G | 1.9334975938915028e-08 | 5.617848639378409 | 32184.0 |
| 17 | rs112674766 | 44130924 | A | T | 1.9334975938915028e-08 | 5.617848639378409 | 32184.0 |
| 17 | rs112074397 | 44131022 | T | C | 1.9334975938915028e-08 | 5.617848639378409 | 32184.0 |

| | | | | | | | |
|---|---|---|---|---|---|---|---|
| 17 | rs78104015 | 44052733 | A | G | 1.9340401853207505e-08 | 5.617800145253169 | 32176.0 |
| 17 | rs74829364 | 44084533 | T | C | 1.934691492127344e-08 | 5.617741952129459 | 32183.0 |
| 17 | rs62064669 | 44084676 | C | T | 1.934691492127344e-08 | 5.617741952129459 | 32183.0 |
| 17 | rs62063669 | 44122626 | C | A | 1.935166538924977e-08 | 5.617699519524156 | 32185.0 |
| 17 | rs77105679 | 44122896 | G | A | 1.935166538924977e-08 | 5.617699519524156 | 32185.0 |
| 17 | 17:43954982_TTCTCTTCCTCA_T | 43954982 | TTCTCTTCCTCA | T | 1.937781341552966e-08 | 5.617466138396959 | 31883.0 |
| 17 | rs112116311 | 43989898 | T | G | 1.9393246795865605e-08 | 5.617328533033664 | 32177.0 |
| 17 | rs78209533 | 43989909 | A | G | 1.9393246795865605e-08 | 5.617328533033664 | 32177.0 |
| 17 | rs55760800 | 43996908 | T | G | 1.9393246795865605e-08 | 5.617328533033664 | 32177.0 |
| 17 | rs56303672 | 43996960 | A | G | 1.9393246795865605e-08 | 5.617328533033664 | 32177.0 |
| 17 | rs17564020 | 43991781 | G | T | 1.9406173869043003e-08 | 5.617213355925144 | 32181.0 |
| 17 | rs17572613 | 44058004 | A | G | 1.9420471517372392e-08 | 5.6170860540482375 | 32185.0 |
| 17 | rs75690239 | 43941710 | C | T | 1.9424558456633766e-08 | 5.61704968191712 | 32180.0 |
| 17 | rs62062786 | 44024660 | G | C | 1.9425239695167358e-08 | 5.617043619888079 | 32181.0 |
| 17 | rs62062787 | 44024865 | T | C | 1.9425239695167358e-08 | 5.617043619888079 | 32181.0 |
| 17 | rs62062788 | 44024906 | A | G | 1.9425239695167358e-08 | 5.617043619888079 | 32181.0 |
| 17 | rs113762994 | 43847095 | C | T | 1.943927842230604e-08 | 5.616918741633143 | 32143.0 |
| 17 | rs370644144 | 43933956 | G | GTGTTTGTTTGTT | 1.9455442080906158e-08 | 5.61677506994172 | 32157.0 |
| 17 | rs754513 | 44055638 | T | A | 1.9460150432686044e-08 | 5.616733241257862 | 32170.0 |
| 17 | rs56101773 | 43957269 | T | C | 1.946813671332537e-08 | 5.616662314135753 | 32182.0 |
| 17 | rs111447859 | 43987184 | T | C | 1.9488424016150845e-08 | 5.616482267102056 | 32168.0 |
| 17 | rs10445335 | 43934896 | T | A | 1.951386362079759e-08 | 5.6162567510909644 | 32169.0 |
| 17 | rs79252344 | 44120608 | A | G | 1.954070618216852e-08 | 5.616019107488698 | 32184.0 |
| 17 | rs62056838 | 43989087 | A | G | 1.9561136227795475e-08 | 5.615838447741403 | 32180.0 |
| 17 | rs62056839 | 43989221 | T | C | 1.9561136227795475e-08 | 5.615838447741403 | 32180.0 |
| 17 | rs17564780 | 44005413 | A | G | 1.9583235189436923e-08 | 5.61564323616627 | 32180.0 |
| 17 | rs62059008 | 44006085 | A | G | 1.9583235189436923e-08 | 5.61564323616627 | 32180.0 |
| 17 | rs62059009 | 44006130 | T | C | 1.9583235189436923e-08 | 5.61564323616627 | 32180.0 |
| 17 | rs62061705 | 44006252 | T | C | 1.9583235189436923e-08 | 5.61564323616627 | 32180.0 |
| 17 | rs62063783 | 44059215 | C | T | 1.9583784543512533e-08 | 5.615638386162144 | 32182.0 |
| 17 | rs17564829 | 44006601 | T | C | 1.962379061674828e-08 | 5.615285544836743 | 32181.0 |
| 17 | rs62061709 | 44007493 | C | T | 1.962379061674828e-08 | 5.615285544836743 | 32181.0 |
| 17 | rs17572248 | 44049867 | A | G | 1.965608748663744e-08 | 5.615001205443412 | 32184.0 |
| 17 | rs62064672 | 44086521 | G | A | 1.9656914529265967e-08 | 5.6149939301757925 | 32185.0 |
| 17 | rs17572893 | 44064208 | G | A | 1.965836193597276e-08 | 5.614981198450344 | 32182.0 |
| 17 | rs62065457 | 43574611 | G | A | 1.966442839593437e-08 | 5.614927846359504 | 32021.0 |
| 17 | rs393208 | 43704310 | T | G | 1.967587694323588e-08 | 5.614827204482522 | 32146.0 |
| 17 | rs62060840 | 44153258 | A | G | 1.9677118744681155e-08 | 5.6148162914739 | 32179.0 |
| 17 | rs9901937 | 44063723 | A | C | 1.96887125980155e-08 | 5.614714436405967 | 32182.0 |
| 17 | rs41374248 | 44120442 | A | G | 1.969478824181389e-08 | 5.614661083520031 | 32185.0 |
| 17 | rs41399444 | 44121469 | C | T | 1.969478824181389e-08 | 5.614661083520031 | 32185.0 |
| 17 | rs62063668 | 44122484 | C | T | 1.969478824181389e-08 | 5.614661083520031 | 32185.0 |
| 17 | rs62641967 | 44047216 | T | G | 1.970017500843308e-08 | 5.6146137933291 | 32177.0 |
| 17 | rs62060786 | 44136763 | C | T | 1.971731176470958e-08 | 5.614463433943207 | 32180.0 |
| 17 | rs62060787 | 44136803 | G | T | 1.971731176470958e-08 | 5.614463433943207 | 32180.0 |
| 17 | rs55714296 | 44137189 | A | T | 1.971731176470958e-08 | 5.614463433943207 | 32180.0 |
| 17 | rs113093579 | 43934116 | T | G | 1.9725470675126963e-08 | 5.6143918915339 | 32154.0 |
| 17 | rs739643 | 43946707 | T | C | 1.9738061248380567e-08 | 5.614281545901298 | 32181.0 |
| 17 | rs62062785 | 44024562 | G | T | 1.973861486093812e-08 | 5.614276695528217 | 32184.0 |
| 17 | rs111306685 | 44025407 | C | CT | 1.973861486093812e-08 | 5.614276695528217 | 32184.0 |
| 17 | rs111582708 | 44082955 | A | AT | 1.9748582490996094e-08 | 5.614189388587956 | 32166.0 |
| 17 | rs112902492 | 43958213 | A | AAT | 1.9752598625061206e-08 | 5.614154223172274 | 32179.0 |
| 17 | rs75945623 | 43937505 | C | T | 1.979682899040038e-08 | 5.613767399040232 | 32176.0 |
| 17 | rs1467962 | 43978604 | A | G | 1.981113099816641e-08 | 5.613642497927036 | 32185.0 |

| | | | | | | | |
|---|---|---|---|---|---|---|---|
| 17 | rs1560311 | 43978625 | T | C | 1.981113099816641e-08 | 5.613642497927036 | 32185.0 |
| 17 | rs62063276 | 44036408 | T | G | 1.985235792437288e-08 | 5.613282947624162 | 32112.0 |
| 17 | rs12150469 | 44092497 | T | C | 1.9857856559198564e-08 | 5.613235047460249 | 32184.0 |
| 17 | rs17572169 | 44045974 | C | T | 1.9864679725845048e-08 | 5.613175626825616 | 32179.0 |
| 17 | rs62063296 | 44046202 | C | T | 1.9864679725845048e-08 | 5.613175626825616 | 32179.0 |
| 17 | rs74373419 | 44047609 | C | T | 1.9864679725845048e-08 | 5.613175626825616 | 32179.0 |
| 17 | rs62063297 | 44047802 | A | T | 1.9864679725845048e-08 | 5.613175626825616 | 32179.0 |
| 17 | rs62062801 | 44028011 | T | G | 1.987443109472128e-08 | 5.613090739862548 | 32167.0 |
| 17 | rs75242405 | 44049329 | C | T | 1.988850919945701e-08 | 5.612968259392109 | 32182.0 |
| 17 | rs76640332 | 44189858 | G | A | 1.9891995388303748e-08 | 5.6129379423145895 | 31996.0 |
| 17 | rs62062799 | 44027521 | G | T | 1.9901480853701113e-08 | 5.6128554796039545 | 32169.0 |
| 17 | rs111962225 | 43934016 | G | C | 1.991013323158656e-08 | 5.612780292683762 | 32154.0 |
| 17 | rs112206871 | 44041817 | A | G | 1.9910691576158745e-08 | 5.612775441903873 | 32170.0 |
| 17 | 17:44104576_CCTT_C | 44104576 | CCTT | C | 1.991250630205309e-08 | 5.612759676860153 | 32049.0 |
| 17 | rs62063853 | 44074613 | A | G | 1.9914740033488536e-08 | 5.612740273710363 | 32043.0 |
| 17 | rs55930267 | 43792091 | C | T | 1.9920884061901257e-08 | 5.612686914939994 | 31593.0 |
| 17 | 17:44294983_C_A | 44294983 | C | A | 1.993233925669153e-08 | 5.6125874731707865 | 32087.0 |
| 17 | rs430789 | 44295157 | A | G | 1.993233925669153e-08 | 5.6125874731707865 | 32087.0 |
| 17 | rs55893711 | 44119320 | C | T | 1.993331743834289e-08 | 5.612578984213675 | 32184.0 |
| 17 | rs10514898 | 44119636 | A | C | 1.993331743834289e-08 | 5.612578984213675 | 32184.0 |
| 17 | rs62063670 | 44123248 | T | C | 1.993331743834289e-08 | 5.612578984213675 | 32184.0 |
| 17 | rs74571696 | 44123685 | G | A | 1.993331743834289e-08 | 5.612578984213675 | 32184.0 |
| 17 | rs79698033 | 44123717 | C | T | 1.993331743834289e-08 | 5.612578984213675 | 32184.0 |
| 17 | rs62055469 | 43943655 | A | C | 1.993492455324273e-08 | 5.612565038061111 | 32172.0 |
| 17 | rs62055470 | 43943657 | A | G | 1.993492455324273e-08 | 5.612565038061111 | 32172.0 |
| 17 | rs62055471 | 43943755 | T | C | 1.993492455324273e-08 | 5.612565038061111 | 32172.0 |
| 17 | rs78962882 | 44084844 | G | C | 1.9998542366428748e-08 | 5.612013853157038 | 32180.0 |
| 17 | rs56249080 | 43939432 | G | T | 2.002561736780426e-08 | 5.611779791507313 | 32181.0 |
| 17 | rs62054857 | 43939936 | G | A | 2.002561736780426e-08 | 5.611779791507313 | 32181.0 |
| 17 | rs62054858 | 43940188 | T | C | 2.002561736780426e-08 | 5.611779791507313 | 32181.0 |
| 17 | rs62054859 | 43940685 | G | A | 2.002561736780426e-08 | 5.611779791507313 | 32181.0 |
| 17 | rs62055460 | 43940825 | G | C | 2.002561736780426e-08 | 5.611779791507313 | 32181.0 |
| 17 | rs62055461 | 43941350 | C | T | 2.002561736780426e-08 | 5.611779791507313 | 32181.0 |
| 17 | rs62055462 | 43941476 | A | C | 2.002561736780426e-08 | 5.611779791507313 | 32181.0 |
| 17 | rs62055463 | 43941491 | A | G | 2.002561736780426e-08 | 5.611779791507313 | 32181.0 |
| 17 | rs77652965 | 43941608 | C | G | 2.002561736780426e-08 | 5.611779791507313 | 32181.0 |
| 17 | rs62055464 | 43941858 | T | G | 2.002561736780426e-08 | 5.611779791507313 | 32181.0 |
| 17 | rs62055465 | 43942046 | T | A | 2.002561736780426e-08 | 5.611779791507313 | 32181.0 |
| 17 | rs62055466 | 43942656 | G | A | 2.002561736780426e-08 | 5.611779791507313 | 32181.0 |
| 17 | rs62055467 | 43942820 | T | C | 2.002561736780426e-08 | 5.611779791507313 | 32181.0 |
| 17 | rs56036719 | 43943974 | C | T | 2.002561736780426e-08 | 5.611779791507313 | 32181.0 |
| 17 | rs55781174 | 43944158 | C | T | 2.002561736780426e-08 | 5.611779791507313 | 32181.0 |
| 17 | rs12150223 | 43944204 | T | C | 2.002561736780426e-08 | 5.611779791507313 | 32181.0 |
| 17 | rs62055476 | 43945495 | A | G | 2.002561736780426e-08 | 5.611779791507313 | 32181.0 |
| 17 | rs1078830 | 43946112 | T | C | 2.002561736780426e-08 | 5.611779791507313 | 32181.0 |
| 17 | rs2950705 | 44343121 | A | G | 2.002645965348912e-08 | 5.611772514930292 | 31534.0 |
| 17 | rs80020809 | 44172391 | C | T | 2.0027652951007043e-08 | 5.611762206441108 | 32108.0 |
| 17 | rs76090253 | 44002969 | A | C | 2.0051533547878417e-08 | 5.611556035410863 | 32182.0 |
| 17 | rs55788597 | 44003397 | T | A | 2.0051533547878417e-08 | 5.611556035410863 | 32182.0 |
| 17 | rs56197117 | 44003446 | A | G | 2.0051533547878417e-08 | 5.611556035410863 | 32182.0 |
| 17 | rs56249311 | 44003599 | C | T | 2.0051533547878417e-08 | 5.611556035410863 | 32182.0 |
| 17 | rs62059004 | 44003893 | A | G | 2.0051533547878417e-08 | 5.611556035410863 | 32182.0 |
| 17 | rs76500077 | 44004076 | T | C | 2.0051533547878417e-08 | 5.611556035410863 | 32182.0 |
| 17 | rs62063303 | 44049133 | T | C | 2.0080015992242932e-08 | 5.611310446231721 | 32181.0 |

| | | | | | | | |
|---|---|---|---|---|---|---|---|
| 17 | rs62061821 | 44186055 | G | A | 2.0086773200486548e-08 | 5.611252232006368 | 32053.0 |
| 17 | 17:44155299_CT_C | 44155299 | CT | C | 2.010064653789345e-08 | 5.611132770971892 | 32183.0 |
| 17 | 17:43914990_AAC_A | 43914990 | AAC | A | 2.0103535063175102e-08 | 5.611107908422871 | 32137.0 |
| 17 | rs62056850 | 43992581 | C | T | 2.0109454288619142e-08 | 5.611056970409697 | 32172.0 |
| 17 | rs55979424 | 43900081 | C | T | 2.013392444311482e-08 | 5.61084654636672 | 32144.0 |
| 17 | rs55691306 | 43978295 | G | C | 2.01396406982654e-08 | 5.610797426853619 | 32184.0 |
| 17 | 17:44201358_TG_T | 44201358 | TG | T | 2.0173406078816967e-08 | 5.6105075583402835 | 32103.0 |
| 17 | rs55662347 | 44070545 | G | A | 2.0197740191931856e-08 | 5.610298947133199 | 32185.0 |
| 17 | rs55913645 | 44070956 | T | A | 2.0197740191931856e-08 | 5.610298947133199 | 32185.0 |
| 17 | rs17573175 | 44071089 | C | G | 2.0197740191931856e-08 | 5.610298947133199 | 32185.0 |
| 17 | rs62063845 | 44071294 | T | C | 2.0197740191931856e-08 | 5.610298947133199 | 32185.0 |
| 17 | rs55886080 | 44072041 | T | A | 2.0197740191931856e-08 | 5.610298947133199 | 32185.0 |
| 17 | rs77527347 | 44058017 | G | T | 2.0224158250830104e-08 | 5.610072746768979 | 32185.0 |
| 17 | rs62054834 | 43934314 | T | C | 2.0243089568463703e-08 | 5.609910826790427 | 32157.0 |
| 17 | rs2316774 | 43946642 | T | C | 2.026189625937031e-08 | 5.609750118247364 | 32176.0 |
| 17 | rs55929431 | 44000202 | G | A | 2.026210926334569e-08 | 5.609748298897105 | 32177.0 |
| 17 | rs79860128 | 43990072 | A | G | 2.028334967753644e-08 | 5.609566969393926 | 32183.0 |
| 17 | rs17563965 | 43990919 | A | G | 2.028334967753644e-08 | 5.609566969393926 | 32183.0 |
| 17 | rs17563986 | 43991272 | A | G | 2.028334967753644e-08 | 5.609566969393926 | 32183.0 |
| 17 | rs55682376 | 43991888 | G | A | 2.028334967753644e-08 | 5.609566969393926 | 32183.0 |
| 17 | rs62056851 | 43992806 | A | G | 2.028334967753644e-08 | 5.609566969393926 | 32183.0 |
| 17 | rs75046472 | 43997499 | A | T | 2.028334967753644e-08 | 5.609566969393926 | 32183.0 |
| 17 | rs17564223 | 43997520 | C | T | 2.028334967753644e-08 | 5.609566969393926 | 32183.0 |
| 17 | rs767056 | 43998910 | C | T | 2.028334967753644e-08 | 5.609566969393926 | 32183.0 |
| 17 | rs767059 | 43999100 | G | C | 2.028334967753644e-08 | 5.609566969393926 | 32183.0 |
| 17 | rs62058964 | 43999406 | A | G | 2.028334967753644e-08 | 5.609566969393926 | 32183.0 |
| 17 | rs62058965 | 43999529 | G | C | 2.028334967753644e-08 | 5.609566969393926 | 32183.0 |
| 17 | rs1052590 | 44102638 | A | G | 2.028725915378252e-08 | 5.609533614268576 | 32175.0 |
| 17 | rs62063854 | 44074619 | G | A | 2.0311656746170576e-08 | 5.609325598175318 | 32041.0 |
| 17 | rs17689918 | 43910088 | G | A | 2.0318988751940103e-08 | 5.609263132229311 | 32176.0 |
| 17 | rs2950013 | 43668455 | T | G | 2.032767662100854e-08 | 5.609189143156892 | 31782.0 |
| 17 | rs17652449 | 44088937 | G | C | 2.033465803901513e-08 | 5.609129709090339 | 32184.0 |
| 17 | rs10445369 | 43934776 | A | G | 2.0357042910289763e-08 | 5.608939276160251 | 32167.0 |
| 17 | rs62062802 | 44028045 | G | T | 2.0360395590319967e-08 | 5.6089107716938535 | 32163.0 |
| 17 | rs62063846 | 44071674 | A | G | 2.0360538269564084e-08 | 5.608909558736831 | 32184.0 |
| 17 | rs55736025 | 44072017 | C | A | 2.0360538269564084e-08 | 5.608909558736831 | 32184.0 |
| 17 | rs62063850 | 44073739 | G | A | 2.0360538269564084e-08 | 5.608909558736831 | 32184.0 |
| 17 | rs17652121 | 44073973 | T | C | 2.0360538269564084e-08 | 5.608909558736831 | 32184.0 |
| 17 | 17:44127471_GTGGTTTCACCA_G | 44127471 | GTGGTTTCACCA | G | 2.038116572968801e-08 | 5.608734285582871 | 32133.0 |
| 17 | rs62056848 | 43992456 | G | A | 2.042491507366072e-08 | 5.6083631132386325 | 32182.0 |
| 17 | rs62056849 | 43992463 | G | C | 2.042491507366072e-08 | 5.6083631132386325 | 32182.0 |
| 17 | rs76627340 | 43937460 | G | A | 2.04477555551307e-08 | 5.6081696396352765 | 32179.0 |
| 17 | rs79889833 | 43937633 | A | G | 2.04477555551307e-08 | 5.6081696396352765 | 32179.0 |
| 17 | rs77625338 | 43937671 | C | T | 2.04477555551307e-08 | 5.6081696396352765 | 32179.0 |
| 17 | rs77489223 | 43937736 | C | T | 2.04477555551307e-08 | 5.6081696396352765 | 32179.0 |
| 17 | rs17574040 | 44102865 | A | C | 2.0474781140858214e-08 | 5.607940986317135 | 32177.0 |
| 17 | rs112887308 | 44128754 | G | GA | 2.049200399932537e-08 | 5.607795423001585 | 32184.0 |
| 17 | rs77426526 | 43962982 | G | A | 2.050378112164755e-08 | 5.607695954055176 | 32186.0 |
| 17 | rs113894932 | 44080408 | G | C | 2.0532677512599388e-08 | 5.6074521314949335 | 32178.0 |
| 17 | rs56189701 | 43987801 | T | C | 2.054807645063801e-08 | 5.6073223340511 | 32149.0 |
| 17 | rs12150254 | 44090455 | T | C | 2.054822041934163e-08 | 5.60732112098644 | 32183.0 |
| 17 | rs62062793 | 44026321 | T | C | 2.0554484015322007e-08 | 5.6072683525940885 | 32176.0 |
| 17 | rs62062289 | 44097235 | G | C | 2.0556428270556028e-08 | 5.607251976164834 | 32178.0 |
| 17 | rs1052587 | 44102604 | T | C | 2.0556428270556028e-08 | 5.607251976164834 | 32178.0 |

| | | | | | | | |
|---|---|---|---|---|---|---|---|
| 17 | rs1052594 | 44102689 | G | C | 2.0556428270556028e-08 | 5.607251976164834 | 32178.0 |
| 17 | rs2696591 | 44318663 | G | A | 2.056528771538364e-08 | 5.607177372241979 | 32064.0 |
| 17 | rs62064665 | 44081527 | T | C | 2.0600690783049118e-08 | 5.6068795599886085 | 32180.0 |
| 17 | rs9468 | 44101563 | T | C | 2.061122948970679e-08 | 5.606791003861852 | 32101.0 |
| 17 | rs62057066 | 43858307 | T | C | 2.0617367472556333e-08 | 5.606739447010944 | 31951.0 |
| 17 | rs62062800 | 44027753 | A | G | 2.0652135015803413e-08 | 5.606447693092217 | 32169.0 |
| 17 | 17:43994508_GAGA_G | 43994508 | GAGA | G | 2.0762018034686628e-08 | 5.605528728155333 | 32152.0 |
| 17 | rs17653906 | 44133142 | G | A | 2.0774020052264144e-08 | 5.605428640016315 | 32178.0 |
| 17 | rs62055468 | 43943000 | A | C | 2.0802925515605065e-08 | 5.605187819594768 | 32178.0 |
| 17 | rs62061793 | 44171565 | C | A | 2.085019095825281e-08 | 5.604794735473286 | 32029.0 |
| 17 | rs62063291 | 44043378 | T | C | 2.0851797495909337e-08 | 5.604781389873217 | 32186.0 |
| 17 | rs62056908 | 43783590 | G | A | 2.090715099540381e-08 | 5.604322173845836 | 31470.0 |
| 17 | rs7502888 | 43986992 | G | C | 2.091828292479801e-08 | 5.6042299652444445 | 32177.0 |
| 17 | rs111240522 | 44066874 | G | A | 2.093088659058055e-08 | 5.604125623359512 | 32186.0 |
| 17 | rs79193046 | 43967562 | T | C | 2.0995340010145992e-08 | 5.603592984958767 | 32185.0 |
| 17 | rs17576631 | 44170018 | G | C | 2.1007327134716083e-08 | 5.603494099320055 | 32113.0 |
| 17 | rs112596352 | 44170238 | G | A | 2.1007327134716083e-08 | 5.603494099320055 | 32113.0 |
| 17 | rs62061790 | 44170414 | G | A | 2.1007327134716083e-08 | 5.603494099320055 | 32113.0 |
| 17 | rs62061791 | 44171024 | T | C | 2.1007327134716083e-08 | 5.603494099320055 | 32113.0 |
| 17 | rs111295615 | 44184428 | T | C | 2.100909267676697e-08 | 5.603479539425275 | 32071.0 |
| 17 | rs55825513 | 44176215 | G | A | 2.101262419733787e-08 | 5.603450419600186 | 32144.0 |
| 17 | rs56294117 | 44042783 | G | A | 2.1033678242930907e-08 | 5.603276912993716 | 32178.0 |
| 17 | rs112415880 | 44042868 | G | A | 2.1033678242930907e-08 | 5.603276912993716 | 32178.0 |
| 17 | rs62063292 | 44043636 | C | T | 2.1033678242930907e-08 | 5.603276912993716 | 32178.0 |
| 17 | rs62063293 | 44043819 | G | A | 2.1033678242930907e-08 | 5.603276912993716 | 32178.0 |
| 17 | rs17650991 | 44044508 | A | C | 2.1033678242930907e-08 | 5.603276912993716 | 32178.0 |
| 17 | rs75145092 | 44044823 | T | A | 2.1033678242930907e-08 | 5.603276912993716 | 32178.0 |
| 17 | rs113857334 | 44044940 | T | C | 2.1033678242930907e-08 | 5.603276912993716 | 32178.0 |
| 17 | rs77513497 | 44045474 | A | G | 2.1033678242930907e-08 | 5.603276912993716 | 32178.0 |
| 17 | rs11079723 | 43841729 | T | C | 2.109304985955393e-08 | 5.602788537467838 | 32149.0 |
| 17 | rs56166491 | 44042221 | A | G | 2.1149179294952453e-08 | 5.602328056910987 | 32177.0 |
| 17 | rs112542080 | 44216966 | G | GC | 2.1151697081037352e-08 | 5.602307429046258 | 32077.0 |
| 17 | rs12150516 | 43944519 | C | T | 2.1154585491671302e-08 | 5.602283767642748 | 32180.0 |
| 17 | rs62055475 | 43945106 | T | C | 2.1154585491671302e-08 | 5.602283767642748 | 32180.0 |
| 17 | rs5026246 | 43572896 | T | G | 2.11574002124724e-08 | 5.602260712911808 | 32186.0 |
| 17 | rs62064675 | 44088926 | C | T | 2.124982272241781e-08 | 5.6015053506988055 | 32186.0 |
| 17 | rs62064666 | 44082514 | T | C | 2.1269099254153042e-08 | 5.6013482070846905 | 32149.0 |
| 17 | rs62064662 | 44079721 | A | G | 2.129920190068361e-08 | 5.601103084561831 | 32181.0 |
| 17 | rs62063304 | 44049900 | A | G | 2.1350188287842538e-08 | 5.600688674246862 | 32182.0 |
| 17 | rs55711941 | 44040823 | C | T | 2.1355943887602283e-08 | 5.600641953887786 | 32177.0 |
| 17 | rs1967981 | 43999785 | C | G | 2.137726032023736e-08 | 5.600469026822507 | 32182.0 |
| 17 | rs1476127 | 43999793 | G | A | 2.137726032023736e-08 | 5.600469026822507 | 32182.0 |
| 17 | rs8079501 | 44000234 | A | T | 2.137726032023736e-08 | 5.600469026822507 | 32182.0 |
| 17 | rs56002706 | 44000505 | C | T | 2.137726032023736e-08 | 5.600469026822507 | 32182.0 |
| 17 | rs8078495 | 44000610 | G | A | 2.137726032023736e-08 | 5.600469026822507 | 32182.0 |
| 17 | rs17649866 | 44000914 | T | C | 2.137726032023736e-08 | 5.600469026822507 | 32182.0 |
| 17 | rs62058968 | 44001228 | G | A | 2.137726032023736e-08 | 5.600469026822507 | 32182.0 |
| 17 | rs17564493 | 44001379 | C | T | 2.137726032023736e-08 | 5.600469026822507 | 32182.0 |
| 17 | rs17649918 | 44001621 | A | G | 2.137726032023736e-08 | 5.600469026822507 | 32182.0 |
| 17 | rs17649954 | 44001661 | A | G | 2.137726032023736e-08 | 5.600469026822507 | 32182.0 |
| 17 | rs55685451 | 44001957 | T | G | 2.137726032023736e-08 | 5.600469026822507 | 32182.0 |
| 17 | rs17564591 | 44002062 | A | C | 2.137726032023736e-08 | 5.600469026822507 | 32182.0 |
| 17 | rs62059003 | 44002431 | C | T | 2.137726032023736e-08 | 5.600469026822507 | 32182.0 |
| 17 | rs17650063 | 44002555 | A | G | 2.137726032023736e-08 | 5.600469026822507 | 32182.0 |

| | | | | | | | |
|---|---|---|---|---|---|---|---|
| 17 | rs17564703 | 44002595 | C | T | 2.137726032023736e-08 | 5.600469026822507 | 32182.0 |
| 17 | rs17650651 | 44030311 | T | G | 2.13861670382034e-08 | 5.600396821693861 | 32148.0 |
| 17 | rs16940799 | 44102933 | T | C | 2.1424379914569368e-08 | 5.6000873678437175 | 32176.0 |
| 17 | rs62059005 | 44004472 | A | G | 2.1482651885109493e-08 | 5.599616502311225 | 32180.0 |
| 17 | rs55823844 | 43978320 | G | A | 2.1506049322335973e-08 | 5.599427788550347 | 32186.0 |
| 17 | rs62063294 | 44044157 | G | A | 2.1570200328179856e-08 | 5.598911394423793 | 32177.0 |
| 17 | rs17650973 | 44044220 | A | T | 2.1570200328179856e-08 | 5.598911394423793 | 32177.0 |
| 17 | rs62063295 | 44044431 | C | T | 2.1570200328179856e-08 | 5.598911394423793 | 32177.0 |
| 17 | rs150246512 | 43956305 | CTT | C | 2.160162792718521e-08 | 5.5986589566214535 | 32161.0 |
| 17 | rs62621252 | 43922942 | T | C | 2.1621822655600665e-08 | 5.598496933367898 | 32186.0 |
| 17 | rs62061706 | 44006896 | T | C | 2.1645671698472042e-08 | 5.5983057804527965 | 32176.0 |
| 17 | rs17649635 | 43995990 | A | G | 2.1689272191517988e-08 | 5.597956845264715 | 32180.0 |
| 17 | rs56113217 | 43926948 | A | G | 2.1707648046854728e-08 | 5.597809987030146 | 32136.0 |
| 17 | rs62054835 | 43934672 | A | C | 2.1710838798950122e-08 | 5.597784499114674 | 32167.0 |
| 17 | rs55996632 | 43935888 | C | T | 2.17152458316056e-08 | 5.5977493014574655 | 32186.0 |
| 17 | rs73984689 | 44184404 | G | A | 2.1722694190157173e-08 | 5.597689829396645 | 32063.0 |
| 17 | rs74903707 | 44028707 | G | A | 2.1746728410265376e-08 | 5.597498060997684 | 32163.0 |
| 17 | rs759528501 | 44275649 | CTAT | C | 2.178580151748716e-08 | 5.59718673640575 | 32000.0 |
| 17 | rs56240566 | 44062689 | A | C | 2.1863618826398868e-08 | 5.59656831925668 | 32026.0 |
| 17 | rs2266497 | 44294786 | A | G | 2.187869440075451e-08 | 5.596448760189417 | 32086.0 |
| 17 | rs17573245 | 44072854 | T | C | 2.1879459930072432e-08 | 5.596442691179933 | 32183.0 |
| 17 | rs17565025 | 44012463 | T | C | 2.190504370122556e-08 | 5.596239985081231 | 32163.0 |
| 17 | rs16940671 | 43908151 | C | T | 2.191769321168216e-08 | 5.596139844993807 | 32122.0 |
| 17 | rs56269136 | 43926992 | C | T | 2.200243741791044e-08 | 5.595470409292952 | 32141.0 |
| 17 | rs79600142 | 43897722 | T | C | 2.2047745026196213e-08 | 5.595113528535139 | 32182.0 |
| 17 | rs2469922 | 44298178 | A | G | 2.214916955414281e-08 | 5.594317197521585 | 32099.0 |
| 17 | rs55707339 | 43842462 | T | C | 2.217040886929405e-08 | 5.594150886115777 | 32129.0 |
| 17 | rs62063782 | 44058716 | G | T | 2.2250973642615435e-08 | 5.593521438082791 | 32176.0 |
| 17 | rs2732594 | 44267170 | G | T | 2.225665583103918e-08 | 5.593477127020964 | 31842.0 |
| 17 | rs17649553 | 43994648 | C | T | 2.226241730961857e-08 | 5.593432208846341 | 32152.0 |
| 17 | 17:43974343_TCGC_T | 43974343 | TCGC | T | 2.234995315041119e-08 | 5.592751138388787 | 31964.0 |
| 17 | rs62057103 | 43887480 | T | C | 2.2360508253951333e-08 | 5.5926691896621765 | 32058.0 |
| 17 | rs140819255 | 43966910 | A | AAGG | 2.2376467460916907e-08 | 5.592545355319115 | 32150.0 |
| 17 | rs2696607 | 44319167 | G | A | 2.2392516148421694e-08 | 5.592420913082024 | 32031.0 |
| 17 | rs112137135 | 43900817 | A | G | 2.2398860446931987e-08 | 5.592371742984064 | 32174.0 |
| 17 | rs2429444 | 44298279 | C | T | 2.2408419383122358e-08 | 5.592297684063107 | 32078.0 |
| 17 | rs2696581 | 44227186 | G | A | 2.2415787192851507e-08 | 5.592240622062571 | 32154.0 |
| 17 | rs2696580 | 44227196 | T | C | 2.2415787192851507e-08 | 5.592240622062571 | 32154.0 |
| 17 | rs79348289 | 44172698 | T | C | 2.246224332649256e-08 | 5.591881248687842 | 32106.0 |
| 17 | rs75412750 | 44172773 | C | T | 2.246224332649256e-08 | 5.591881248687842 | 32106.0 |
| 17 | rs62061795 | 44173179 | C | A | 2.246224332649256e-08 | 5.591881248687842 | 32106.0 |
| 17 | rs17576709 | 44173356 | C | G | 2.246224332649256e-08 | 5.591881248687842 | 32106.0 |
| 17 | rs62061796 | 44173408 | G | A | 2.246224332649256e-08 | 5.591881248687842 | 32106.0 |
| 17 | rs62061802 | 44174351 | T | A | 2.246224332649256e-08 | 5.591881248687842 | 32106.0 |
| 17 | rs62061803 | 44174513 | C | A | 2.246224332649256e-08 | 5.591881248687842 | 32106.0 |
| 17 | rs79346219 | 44184375 | A | G | 2.2462871758023704e-08 | 5.59187639224148 | 32079.0 |
| 17 | rs5820609 | 44149210 | G | GA | 2.247740904625917e-08 | 5.5917640865518985 | 32112.0 |
| 17 | rs17659881 | 44157597 | A | G | 2.2482361687240428e-08 | 5.591725841750847 | 32186.0 |
| 17 | 17:44172205_CA_C | 44172205 | CA | C | 2.2506511298593898e-08 | 5.591539473059084 | 32165.0 |
| 17 | rs17763634 | 43921754 | T | C | 2.254030042787556e-08 | 5.591279039291943 | 32175.0 |
| 17 | rs9897399 | 43804317 | A | G | 2.2544162955445285e-08 | 5.591249292536657 | 32174.0 |
| 17 | rs62056812 | 43987154 | G | A | 2.2554255885159875e-08 | 5.591171586493722 | 32141.0 |
| 17 | rs62063789 | 44061898 | T | C | 2.2590800910157905e-08 | 5.590890506349883 | 32171.0 |
| 17 | 17:43965429_TC_T | 43965429 | TC | T | 2.259498795112708e-08 | 5.590858330587542 | 32081.0 |

| | | | | | | | |
|---|---|---|---|---|---|---|---|
| 17 | rs62056809 | 43984669 | A | G | 2.26084232170706e-08 | 5.590755124921866 | 32071.0 |
| 17 | rs62062794 | 44026394 | A | G | 2.2648300874907278e-08 | 5.590449146980902 | 32173.0 |
| 17 | rs112647192 | 43994623 | G | A | 2.2661212566787092e-08 | 5.590350188710159 | 32178.0 |
| 17 | rs17572147 | 44044985 | A | G | 2.2719841432509105e-08 | 5.589901530546952 | 32172.0 |
| 17 | rs17691328 | 43955487 | C | T | 2.2725561525377265e-08 | 5.589857817648454 | 32177.0 |
| 17 | rs1724404 | 44323425 | C | T | 2.2728024777920486e-08 | 5.589838996784283 | 31820.0 |
| 17 | 17:43897026_TGGAG_T | 43897026 | TGGAG | T | 2.276810923761204e-08 | 5.5895330031844255 | 32179.0 |
| 17 | rs55822644 | 43792090 | G | C | 2.2813685885644554e-08 | 5.589185718397294 | 31606.0 |
| 17 | rs62054840 | 43936812 | G | A | 2.2827009214957102e-08 | 5.589084324540231 | 32081.0 |
| 17 | rs56301449 | 43960918 | T | C | 2.282812648724831e-08 | 5.589075824430315 | 32173.0 |
| 17 | rs8712 | 44101871 | A | G | 2.2838663404886002e-08 | 5.5889956803384075 | 32091.0 |
| 17 | rs574194744 | 43837868 | T | TTTATGTTATG | 2.289029924635972e-08 | 5.5886034548351615 | 32011.0 |
| 17 | rs112480703 | 44189799 | T | C | 2.2890379266655776e-08 | 5.588602847668277 | 31855.0 |
| 17 | rs17651549 | 44061278 | C | T | 2.2915118677303253e-08 | 5.588415232115014 | 32186.0 |
| 17 | rs113277432 | 44033169 | T | C | 2.295873656098267e-08 | 5.588084926816107 | 31726.0 |
| 17 | rs17661015 | 44192957 | T | C | 2.297864818525149e-08 | 5.587934344436785 | 31798.0 |
| 17 | rs113651837 | 43895168 | T | TG | 2.302673054622678e-08 | 5.587571241393355 | 32097.0 |
| 17 | rs17690176 | 43916773 | A | C | 2.309815100705541e-08 | 5.587033252441702 | 32177.0 |
| 17 | rs2429445 | 44298209 | T | C | 2.311034475758325e-08 | 5.586941561967222 | 32098.0 |
| 17 | rs200860866 | 43773525 | T | TAAAA | 2.312408037286556e-08 | 5.586838333718744 | 32116.0 |
| 17 | rs80143279 | 43776742 | T | C | 2.3125131072791592e-08 | 5.586830439769359 | 32087.0 |
| 17 | rs62062795 | 44027090 | C | T | 2.3128687635217146e-08 | 5.586803721761003 | 32171.0 |
| 17 | rs878887 | 43912582 | C | T | 2.3134346907889454e-08 | 5.586761215756437 | 32181.0 |
| 17 | rs878888 | 43912635 | A | G | 2.3134346907889454e-08 | 5.586761215756437 | 32181.0 |
| 17 | rs4525537 | 43912723 | T | C | 2.3134346907889454e-08 | 5.586761215756437 | 32181.0 |
| 17 | rs4640231 | 43912786 | G | C | 2.3134346907889454e-08 | 5.586761215756437 | 32181.0 |
| 17 | rs4482334 | 43912830 | T | C | 2.3134346907889454e-08 | 5.586761215756437 | 32181.0 |
| 17 | rs62055945 | 43851851 | G | A | 2.3137823991302453e-08 | 5.586735104875023 | 32168.0 |
| 17 | rs56043078 | 43853235 | T | G | 2.3137823991302453e-08 | 5.586735104875023 | 32168.0 |
| 17 | rs77965652 | 43853457 | A | G | 2.3137823991302453e-08 | 5.586735104875023 | 32168.0 |
| 17 | rs55849949 | 43853526 | A | G | 2.3137823991302453e-08 | 5.586735104875023 | 32168.0 |
| 17 | rs55991914 | 43854267 | C | T | 2.3137823991302453e-08 | 5.586735104875023 | 32168.0 |
| 17 | rs62055955 | 43854340 | A | G | 2.3137823991302453e-08 | 5.586735104875023 | 32168.0 |
| 17 | rs62056879 | 43781426 | T | C | 2.3143566343446548e-08 | 5.586691991475843 | 32145.0 |
| 17 | 17:44316878_CA_C | 44316878 | CA | C | 2.3160558943850765e-08 | 5.586564472363511 | 32166.0 |
| 17 | rs2696670 | 44317103 | T | C | 2.3160558943850765e-08 | 5.586564472363511 | 32166.0 |
| 17 | rs62063675 | 44126575 | C | T | 2.3178859883039577e-08 | 5.586427236494498 | 32159.0 |
| 17 | rs12150235 | 44015745 | A | G | 2.3211123442934252e-08 | 5.586185553070111 | 32182.0 |
| 17 | rs55780945 | 44040120 | C | T | 2.322069747951333e-08 | 5.58611389755841 | 32168.0 |
| 17 | rs17652066 | 44073145 | G | C | 2.322832702468717e-08 | 5.586056815843847 | 32171.0 |
| 17 | rs2696582 | 44226561 | T | G | 2.3229301186047573e-08 | 5.586049528803357 | 32056.0 |
| 17 | 17:44264269_GT_G | 44264269 | GT | G | 2.3266999489102326e-08 | 5.585767760962954 | 30519.0 |
| 17 | rs2316784 | 44021699 | G | T | 2.329425428993004e-08 | 5.585564326683442 | 31990.0 |
| 17 | rs4327091 | 44021717 | G | A | 2.329425428993004e-08 | 5.585564326683442 | 31990.0 |
| 17 | rs10445334 | 43934826 | T | G | 2.3294742720806736e-08 | 5.5855606830633295 | 32166.0 |
| 17 | rs17689882 | 43906828 | G | A | 2.3343065536021888e-08 | 5.585200568283975 | 32186.0 |
| 17 | rs62056810 | 43984672 | T | C | 2.3389279808623214e-08 | 5.584856843204876 | 32077.0 |
| 17 | rs77044307 | 44172408 | C | G | 2.339402061029441e-08 | 5.584821620120801 | 32105.0 |
| 17 | rs16940672 | 43908152 | C | T | 2.33954920851012e-08 | 5.584810688804753 | 32122.0 |
| 17 | rs62062779 | 44022577 | A | C | 2.3428869861497443e-08 | 5.5845629105173815 | 32118.0 |
| 17 | rs79892630 | 43781505 | C | T | 2.3504715502436075e-08 | 5.584001145571412 | 32168.0 |
| 17 | rs56303031 | 43853922 | G | A | 2.352763939962284e-08 | 5.583831701657067 | 32166.0 |
| 17 | rs17564619 | 44002138 | A | G | 2.3550667496219512e-08 | 5.583661648804454 | 32181.0 |
| 17 | rs62062784 | 44024552 | C | G | 2.3648205331295163e-08 | 5.582943157672171 | 32186.0 |

| | | | | | | | |
|---|---|---|---|---|---|---|---|
| 17 | rs1078268 | 44075901 | A | G | 2.3649775099762645e-08 | 5.5829316178498045 | 32115.0 |
| 17 | rs17650579 | 44030115 | C | T | 2.3662006267274658e-08 | 5.582841728452546 | 32151.0 |
| 17 | rs62062803 | 44028183 | T | C | 2.3721100561467915e-08 | 5.5824080665057805 | 32148.0 |
| 17 | rs62055950 | 43853133 | T | C | 2.3727564999845942e-08 | 5.582360691034778 | 32167.0 |
| 17 | rs142222997 | 44057770 | G | GCAG GAGA ATTG TTAC | 2.372806233586224e-08 | 5.582357046762584 | 32153.0 |
| 17 | rs1078269 | 44075837 | T | C | 2.3731875584831315e-08 | 5.582329107317793 | 32114.0 |
| 17 | rs2458207 | 44296251 | T | C | 2.3774940628084777e-08 | 5.582013874473717 | 32078.0 |
| 17 | 17:44303525_G_A | 44303525 | G | A | 2.3774940628084777e-08 | 5.582013874473717 | 32144.0 |
| 17 | 17:44129997_AAAC_A | 44129997 | AAAC | A | 2.3817749485558042e-08 | 5.5817010656830774 | 32168.0 |
| 17 | rs2532385 | 44313950 | C | A | 2.3830232287878798e-08 | 5.581609955298768 | 32165.0 |
| 17 | rs2696675 | 44314066 | A | G | 2.3830232287878798e-08 | 5.581609955298768 | 32165.0 |
| 17 | rs74872829 | 43938394 | A | G | 2.3835893270730723e-08 | 5.581568651771797 | 32148.0 |
| 17 | rs62054384 | 43804306 | T | C | 2.384072279783808e-08 | 5.581533422217611 | 32176.0 |
| 17 | rs62054388 | 43805590 | C | T | 2.385055134869469e-08 | 5.5814617480830275 | 32173.0 |
| 17 | rs62057107 | 43896032 | C | T | 2.38519676536656e-08 | 5.581451422124736 | 32172.0 |
| 17 | rs62055946 | 43851971 | C | A | 2.391277977796735e-08 | 5.581008615132913 | 32167.0 |
| 17 | rs62062813 | 44032956 | G | A | 2.3916538589328623e-08 | 5.580981281009078 | 32096.0 |
| 17 | rs76761706 | 43776741 | T | C | 2.3939354463456367e-08 | 5.5808154530965375 | 32079.0 |
| 17 | rs140713557 | 43953179 | A | G | 2.3976507177097885e-08 | 5.580545751672484 | 32178.0 |
| 17 | rs62063852 | 44074581 | A | G | 2.3991752770560137e-08 | 5.5804351972119575 | 32128.0 |
| 17 | rs62070943 | 44304203 | T | C | 2.404040105290708e-08 | 5.580082876244683 | 32148.0 |
| 17 | 17:44304298_T_A | 44304298 | T | A | 2.404040105290708e-08 | 5.580082876244683 | 32148.0 |
| 17 | rs681485 | 43707619 | C | A | 2.4123072421269024e-08 | 5.579485737115928 | 32186.0 |
| 17 | rs76070179 | 44304490 | T | C | 2.4219465851401122e-08 | 5.5787919858737505 | 32148.0 |
| 17 | 17:44303606_AG_A | 44303606 | AG | A | 2.42214956783502e-08 | 5.578777405874453 | 32146.0 |
| 17 | 17:43839150_AT_A | 43839150 | AT | A | 2.4311312895873574e-08 | 5.578133444061779 | 32080.0 |
| 17 | rs112947543 | 44025678 | G | GCT | 2.4313520056940785e-08 | 5.578117648480986 | 32007.0 |
| 17 | rs62057061 | 43856639 | C | G | 2.435124190939233e-08 | 5.577847906411751 | 32186.0 |
| 17 | rs17650597 | 44030195 | A | T | 2.4439226834316985e-08 | 5.577220315103852 | 32152.0 |
| 17 | rs17650633 | 44030231 | T | C | 2.4439226834316985e-08 | 5.577220315103852 | 32152.0 |
| 17 | rs2696684 | 44244397 | G | A | 2.454251257903789e-08 | 5.576486375966784 | 32035.0 |
| 17 | rs55690782 | 44137311 | G | A | 2.4555539067902188e-08 | 5.576394023877961 | 32172.0 |
| 17 | rs55649944 | 44119463 | C | T | 2.457577783777457e-08 | 5.576250634164416 | 32001.0 |
| 17 | rs56406462 | 44119474 | T | C | 2.457577783777457e-08 | 5.576250634164416 | 32001.0 |
| 17 | rs62055544 | 43964539 | A | G | 2.458873579733959e-08 | 5.576158888448607 | 31836.0 |
| 17 | rs112836774 | 44084002 | T | C | 2.4595603655313872e-08 | 5.576110281256318 | 31863.0 |
| 17 | 17:44301546_T_C | 44301546 | T | C | 2.463255062043791e-08 | 5.575849015336753 | 32123.0 |
| 17 | rs9891103 | 44091886 | C | T | 2.4677992479175507e-08 | 5.57552819987605 | 32099.0 |
| 17 | rs62057065 | 43858187 | G | T | 2.468281688747977e-08 | 5.575494173656655 | 32159.0 |
| 17 | rs56146262 | 43931907 | C | A | 2.4692813245862743e-08 | 5.5754236905678999 | 32143.0 |
| 17 | rs2316765 | 43912454 | T | C | 2.4730162305002508e-08 | 5.575160591759628 | 32183.0 |
| 17 | rs878886 | 43912490 | C | G | 2.4730162305002508e-08 | 5.575160591759628 | 32183.0 |
| 17 | rs56127111 | 43913315 | C | T | 2.4730162305002508e-08 | 5.575160591759628 | 32183.0 |
| 17 | 17:44303668_G_T | 44303668 | G | T | 2.473344269343896e-08 | 5.575137502072276 | 32138.0 |
| 17 | rs2696444 | 44288672 | G | A | 2.4741559171207692e-08 | 5.575080385349303 | 32028.0 |
| 17 | rs3029044 | 43907641 | C | CAGG TGG | 2.475244280072053e-08 | 5.575003824349705 | 32182.0 |
| 17 | rs56406407 | 43792099 | T | C | 2.4791870414042102e-08 | 5.5747267437098165 | 31597.0 |
| 17 | rs450237 | 43709225 | T | C | 2.481775657989016e-08 | 5.57454505912033 | 32012.0 |
| 17 | rs199504614 | 43781720 | T | TAAA AAAA AGAG | 2.4830492936351112e-08 | 5.5744557352514414 | 32172.0 |
| 17 | rs62055938 | 43849656 | C | A | 2.4836993545544104e-08 | 5.574410161677257 | 32164.0 |
| 17 | rs62055939 | 43849787 | A | T | 2.4836993545544104e-08 | 5.574410161677257 | 32164.0 |
| 17 | rs62055940 | 43849896 | A | G | 2.4836993545544104e-08 | 5.574410161677257 | 32164.0 |
| 17 | rs76885724 | 43850519 | C | T | 2.4836993545544104e-08 | 5.574410161677257 | 32164.0 |

| | | | | | | | |
|---|---|---|---|---|---|---|---|
| 17 | rs62055942 | 43850966 | A | G | 2.4836993545544104e-08 | 5.574410161677257 | 32164.0 |
| 17 | rs62055943 | 43851018 | T | C | 2.4836993545544104e-08 | 5.574410161677257 | 32164.0 |
| 17 | rs62055957 | 43854536 | C | A | 2.4836993545544104e-08 | 5.574410161677257 | 32164.0 |
| 17 | rs16940677 | 43911898 | C | T | 2.4893662721145408e-08 | 5.5740133628557516 | 32180.0 |
| 17 | rs78328427 | 43916932 | G | A | 2.489453153383575e-08 | 5.574007286235749 | 32180.0 |
| 17 | rs55920545 | 43803773 | A | AG | 2.490913203201084e-08 | 5.5739051987113335 | 32169.0 |
| 17 | rs4074462 | 43855228 | G | T | 2.4937574932236062e-08 | 5.573706490967775 | 32165.0 |
| 17 | rs62063855 | 44075535 | T | C | 2.502746128510316e-08 | 5.573079969850911 | 32104.0 |
| 17 | rs62062805 | 44030773 | A | G | 2.5059971511023737e-08 | 5.572853906412462 | 32130.0 |
| 17 | rs76839282 | 44029178 | G | C | 2.5082980865804005e-08 | 5.572694080269824 | 32155.0 |
| 17 | rs373685009 | 43790598 | G | GAA | 2.5097426962805417e-08 | 5.572593808387332 | 32091.0 |
| 17 | rs79501144 | 43900697 | C | T | 2.5126869687052632e-08 | 5.572389616635845 | 32177.0 |
| 17 | rs62057123 | 43901238 | A | G | 2.5126869687052632e-08 | 5.572389616635845 | 32177.0 |
| 17 | rs62055901 | 43845041 | G | A | 2.518032094527859e-08 | 5.5720195131503285 | 32156.0 |
| 17 | rs17689608 | 43896528 | C | G | 2.51884053682048e-08 | 5.571963601942153 | 32174.0 |
| 17 | rs62057108 | 43896616 | T | C | 2.51884053682048e-08 | 5.571963601942153 | 32174.0 |
| 17 | rs62057109 | 43896637 | T | C | 2.51884053682048e-08 | 5.571963601942153 | 32174.0 |
| 17 | rs62057110 | 43896734 | T | C | 2.51884053682048e-08 | 5.571963601942153 | 32174.0 |
| 17 | rs62057113 | 43897449 | A | T | 2.51884053682048e-08 | 5.571963601942153 | 32174.0 |
| 17 | rs17762361 | 43781778 | A | G | 2.52341477860469e-08 | 5.571647578787278 | 32153.0 |
| 17 | rs17650818 | 44036787 | T | C | 2.5308741236099104e-08 | 5.571133421503205 | 32140.0 |
| 17 | rs12150260 | 44090538 | T | G | 2.537106751487499e-08 | 5.57070494582015 | 32096.0 |
| 17 | rs56971664 | 43918613 | T | C | 2.5416159702107087e-08 | 5.570395586080085 | 32177.0 |
| 17 | rs10514904 | 44207887 | C | T | 2.5450318931372344e-08 | 5.570161587441014 | 32069.0 |
| 17 | rs2532392 | 44309856 | G | A | 2.545333775221642e-08 | 5.5701409224791645 | 32110.0 |
| 17 | rs2469927 | 44310164 | T | C | 2.545333775221642e-08 | 5.5701409224791645 | 32110.0 |
| 17 | rs3108507 | 44310234 | T | C | 2.545333775221642e-08 | 5.5701409224791645 | 32110.0 |
| 17 | rs62056808 | 43984646 | T | C | 2.5454758495237828e-08 | 5.570131197782981 | 32075.0 |
| 17 | rs62055879 | 43836578 | G | A | 2.546701563074139e-08 | 5.570047322059152 | 32151.0 |
| 17 | rs2469934 | 43704419 | T | C | 2.549421513684253e-08 | 5.569861335356797 | 32166.0 |
| 17 | rs373853 | 43704491 | G | A | 2.549421513684253e-08 | 5.569861335356797 | 32166.0 |
| 17 | rs71363552 | 43834671 | A | AC | 2.550488938084501e-08 | 5.569788398867682 | 32101.0 |
| 17 | rs62057062 | 43856710 | G | A | 2.5542729238989054e-08 | 5.569530079745685 | 32165.0 |
| 17 | 17:44028469_CA_C | 44028469 | CA | C | 2.5553958215731657e-08 | 5.5694534950073376 | 32137.0 |
| 17 | 17:43918301_GCCAGGAGTTCGAGA_G | 43918301 | GCCAGGAGTTCGAGA | G | 2.5564478629099384e-08 | 5.569381772495007 | 32144.0 |
| 17 | rs2873268 | 43930955 | G | A | 2.5593922667447655e-08 | 5.569181191062268 | 32155.0 |
| 17 | rs62071573 | 44342994 | C | T | 2.559722604042624e-08 | 5.5691587014887975 | 31379.0 |
| 17 | rs55787105 | 43853109 | G | A | 2.561008638971493e-08 | 5.569071174231227 | 32161.0 |
| 17 | rs76294809 | 43849327 | G | A | 2.5616518939105715e-08 | 5.5690274104419775 | 32163.0 |
| 17 | rs2732593 | 44267164 | C | T | 2.5628226667605576e-08 | 5.568947784384376 | 31825.0 |
| 17 | rs2696568 | 44266342 | T | C | 2.566427646851236e-08 | 5.568702825666077 | 31947.0 |
| 17 | rs62055900 | 43845002 | C | G | 2.5693026633242585e-08 | 5.568507707266639 | 32152.0 |
| 17 | rs17334923 | 43827244 | C | T | 2.5735178848569517e-08 | 5.5682220160543725 | 32163.0 |
| 17 | 17:44219831_T_A | 44219831 | T | A | 2.5765983774991066e-08 | 5.568013519377291 | 32083.0 |
| 17 | rs78917495 | 43858482 | C | T | 2.5820587215582816e-08 | 5.567644541173142 | 32155.0 |
| 17 | rs56070245 | 43847868 | T | C | 2.5856000493129383e-08 | 5.567405643486248 | 32168.0 |
| 17 | rs56387266 | 43847912 | T | C | 2.5856000493129383e-08 | 5.567405643486248 | 32168.0 |
| 17 | rs150334020 | 44078680 | T | TA | 2.586474849754748e-08 | 5.567346678426727 | 32174.0 |
| 17 | rs62055885 | 43838014 | A | G | 2.5918201297065425e-08 | 5.566986804988088 | 32148.0 |
| 17 | rs113793114 | 44037079 | G | A | 2.593745994501312e-08 | 5.566857321727959 | 32141.0 |
| 17 | rs76830096 | 43857129 | A | C | 2.5944877811156505e-08 | 5.566807473462673 | 32167.0 |
| 17 | rs62063277 | 44036462 | T | A | 2.598643818123028e-08 | 5.5665284421954295 | 32141.0 |
| 17 | rs56180212 | 43960911 | A | G | 2.600139412278229e-08 | 5.5664281357753085 | 32159.0 |
| 17 | rs62057070 | 43859640 | A | G | 2.6033963780078528e-08 | 5.566209891380319 | 32182.0 |

| | | | | | | | |
|---|---|---|---|---|---|---|---|
| 17 | rs62057071 | 43859691 | C | G | 2.6033963780078528e-08 | 5.566209891380319 | 32182.0 |
| 17 | rs141412361 | 44102443 | C | CCT | 2.6062756301698208e-08 | 5.566017177544046 | 32161.0 |
| 17 | rs62057069 | 43859065 | T | C | 2.6139658657488088e-08 | 5.565503466489539 | 32179.0 |
| 17 | 17:44107321_CT_C | 44107321 | CT | C | 2.6149959486481147e-08 | 5.565434767917551 | 31867.0 |
| 17 | rs62057064 | 43857033 | C | T | 2.6183076687161224e-08 | 5.565214079570914 | 32163.0 |
| 17 | rs62062780 | 44022593 | C | T | 2.618344184388564e-08 | 5.565211647728252 | 32115.0 |
| 17 | rs55787734 | 43799052 | A | G | 2.6210751445570054e-08 | 5.565029866553965 | 32013.0 |
| 17 | rs17763596 | 43921210 | G | T | 2.62352533429196e-08 | 5.564866930688727 | 32183.0 |
| 17 | rs62061707 | 44006954 | T | C | 2.6285424143734787e-08 | 5.56453375870656 | 32031.0 |
| 17 | rs242561 | 44026548 | T | C | 2.631549835776017e-08 | 5.564334338933666 | 32165.0 |
| 17 | rs62054382 | 43802525 | C | T | 2.6346524752505835e-08 | 5.564128836965827 | 32180.0 |
| 17 | rs392116 | 43703555 | A | G | 2.636104051000369e-08 | 5.564032773106753 | 32165.0 |
| 17 | rs385383 | 43704031 | C | A | 2.636104051000369e-08 | 5.564032773106753 | 32165.0 |
| 17 | rs17660228 | 44166500 | G | A | 2.636729021058442e-08 | 5.563991429008754 | 32160.0 |
| 17 | 17:43721690_CAAAAAAAA_C | 43721690 | CAAAAAAAA | C | 2.6396628050929803e-08 | 5.563797475274549 | 32173.0 |
| 17 | rs57631148 | 44304740 | A | T | 2.6430604210742e-08 | 5.56357311864762 | 32149.0 |
| 17 | rs445354 | 44305252 | G | A | 2.6430604210742e-08 | 5.56357311864762 | 32149.0 |
| 17 | rs62064661 | 44078892 | G | A | 2.643687007575164e-08 | 5.563531773488172 | 32177.0 |
| 17 | rs4309444 | 43895797 | T | C | 2.646471548422541e-08 | 5.563348151185262 | 32167.0 |
| 17 | rs34303488 | 43848181 | C | T | 2.6478279774897752e-08 | 5.563258771435863 | 32167.0 |
| 17 | rs62055899 | 43844977 | T | A | 2.6499054642428954e-08 | 5.5631219648328365 | 32152.0 |
| 17 | 17:44304910_G_C | 44304910 | G | C | 2.6583055225480688e-08 | 5.562569863564368 | 32139.0 |
| 17 | rs62055894 | 43843943 | G | A | 2.6601407164700777e-08 | 5.562449469176508 | 32163.0 |
| 17 | rs62055903 | 43846668 | C | A | 2.660446702829371e-08 | 5.5624294033664885 | 32166.0 |
| 17 | rs111374028 | 43846820 | G | A | 2.660446702829371e-08 | 5.5624294033664885 | 32166.0 |
| 17 | rs113934115 | 43847039 | C | T | 2.660446702829371e-08 | 5.5624294033664885 | 32166.0 |
| 17 | rs10445362 | 43914554 | C | A | 2.66054870594836e-08 | 5.562422714758152 | 32184.0 |
| 17 | rs10445363 | 43914558 | G | A | 2.66054870594836e-08 | 5.562422714758152 | 32184.0 |
| 17 | rs62054802 | 43914598 | G | C | 2.66054870594836e-08 | 5.562422714758152 | 32184.0 |
| 17 | rs62054803 | 43914728 | G | T | 2.66054870594836e-08 | 5.562422714758152 | 32184.0 |
| 17 | rs62054805 | 43915054 | A | G | 2.66054870594836e-08 | 5.562422714758152 | 32184.0 |
| 17 | rs62054806 | 43915312 | C | T | 2.66054870594836e-08 | 5.562422714758152 | 32184.0 |
| 17 | rs10445364 | 43916356 | G | A | 2.66054870594836e-08 | 5.562422714758152 | 32184.0 |
| 17 | rs10445333 | 43916509 | A | G | 2.66054870594836e-08 | 5.562422714758152 | 32184.0 |
| 17 | rs17763533 | 43918190 | T | C | 2.66054870594836e-08 | 5.562422714758152 | 32184.0 |
| 17 | rs112583797 | 43918418 | A | G | 2.66054870594836e-08 | 5.562422714758152 | 32184.0 |
| 17 | rs74922289 | 43918524 | A | G | 2.66054870594836e-08 | 5.562422714758152 | 32184.0 |
| 17 | rs56150806 | 43919301 | T | C | 2.66054870594836e-08 | 5.562422714758152 | 32184.0 |
| 17 | rs17690314 | 43919884 | T | G | 2.66054870594836e-08 | 5.562422714758152 | 32184.0 |
| 17 | rs56233672 | 43920300 | T | C | 2.66054870594836e-08 | 5.562422714758152 | 32184.0 |
| 17 | rs56354807 | 43920378 | C | T | 2.66054870594836e-08 | 5.562422714758152 | 32184.0 |
| 17 | rs55718212 | 43920411 | T | C | 2.66054870594836e-08 | 5.562422714758152 | 32184.0 |
| 17 | rs17690326 | 43920974 | T | C | 2.66054870594836e-08 | 5.562422714758152 | 32184.0 |
| 17 | rs62054814 | 43921572 | T | A | 2.66054870594836e-08 | 5.562422714758152 | 32184.0 |
| 17 | rs78074121 | 43896690 | T | C | 2.66708488476045e-08 | 5.561994638627192 | 32170.0 |
| 17 | rs55962674 | 43799033 | C | T | 2.6696237436012492e-08 | 5.5618286347598564 | 32166.0 |
| 17 | rs55653963 | 43799509 | A | C | 2.6696237436012492e-08 | 5.5618286347598564 | 32166.0 |
| 17 | rs62054378 | 43799667 | C | A | 2.6696237436012492e-08 | 5.5618286347598564 | 32166.0 |
| 17 | rs77819001 | 43799804 | C | A | 2.6696237436012492e-08 | 5.5618286347598564 | 32166.0 |
| 17 | rs76667867 | 43799810 | G | C | 2.6696237436012492e-08 | 5.5618286347598564 | 32166.0 |
| 17 | rs2532294 | 44233433 | G | C | 2.6718017935680107e-08 | 5.561686344505663 | 32038.0 |
| 17 | rs2532293 | 44233589 | T | C | 2.6718017935680107e-08 | 5.561686344505663 | 32038.0 |
| 17 | rs17662235 | 44233776 | T | C | 2.6718017935680107e-08 | 5.561686344505663 | 32038.0 |
| 17 | rs17585214 | 44233811 | C | T | 2.6718017935680107e-08 | 5.561686344505663 | 32038.0 |

| | | | | | | | |
|---|---|---|---|---|---|---|---|
| 17 | rs1528074 | 44234060 | A | G | 2.6718017935680107e-08 | 5.561686344505663 | 32038.0 |
| 17 | rs1406068 | 44234526 | C | T | 2.6718017935680107e-08 | 5.561686344505663 | 32038.0 |
| 17 | rs79772576 | 44106220 | A | G | 2.6763962929126413e-08 | 5.5613865583460536 | 32186.0 |
| 17 | rs4479288 | 43793828 | G | A | 2.6765362063046315e-08 | 5.561377436984622 | 32153.0 |
| 17 | 17:44106515_AAAG_A | 44106515 | AAAG | A | 2.67688135665658e-08 | 5.561354937606549 | 32166.0 |
| 17 | rs12185235 | 43923703 | C | T | 2.6773851678981403e-08 | 5.561322100625653 | 32176.0 |
| 17 | rs112493812 | 44314179 | G | A | 2.680372623749612e-08 | 5.561127509873339 | 32152.0 |
| 17 | rs62056880 | 43782304 | T | C | 2.6812694974362482e-08 | 5.56106913223524 | 32174.0 |
| 17 | rs8073146 | 43893751 | A | G | 2.6818488846764685e-08 | 5.561031429909475 | 32046.0 |
| 17 | rs113263403 | 44016011 | A | AT | 2.6818488846764685e-08 | 5.561031429909475 | 32173.0 |
| 17 | rs1526126 | 43812218 | C | T | 2.681951692003017e-08 | 5.5610247407788655 | 32170.0 |
| 17 | rs1526128 | 43779624 | C | T | 2.68303607716788e-08 | 5.560954200703983 | 32067.0 |
| 17 | rs1526129 | 43779657 | T | C | 2.6843266813114168e-08 | 5.560870281977056 | 32178.0 |
| 17 | rs75916678 | 43849366 | T | C | 2.6911357863414442e-08 | 5.560428181899663 | 32139.0 |
| 17 | rs62056878 | 43780561 | C | T | 2.6970594733975595e-08 | 5.560044452075112 | 32149.0 |
| 17 | 17:43898362_AAGG_A | 43898362 | AAGG | A | 2.6970594733975595e-08 | 5.560044452075112 | 32126.0 |
| 17 | rs144416125 | 44296523 | G | A | 2.697943017417813e-08 | 5.5599872871818805 | 32081.0 |
| 17 | rs436667 | 43709415 | C | T | 2.701630643279994e-08 | 5.559748895446765 | 32186.0 |
| 17 | rs56268325 | 43847741 | T | C | 2.702459156395899e-08 | 5.559695378498542 | 32167.0 |
| 17 | rs192252295 | 43702296 | T | C | 2.7073128366718446e-08 | 5.559382179514583 | 32116.0 |
| 17 | rs62054383 | 43802907 | T | C | 2.7080864437636863e-08 | 5.559332310432877 | 32176.0 |
| 17 | rs62055928 | 43847374 | G | A | 2.7107296623152422e-08 | 5.559162024716722 | 32165.0 |
| 17 | rs62054809 | 43918239 | C | T | 2.71114525506186e-08 | 5.559135265385506 | 32176.0 |
| 17 | rs113399561 | 44205350 | A | ATT | 2.711183039317192e-08 | 5.5591328327170455 | 31828.0 |
| 17 | rs2696586 | 44225234 | T | C | 2.7114097556874947e-08 | 5.559118236699367 | 32064.0 |
| 17 | rs611351 | 43692935 | A | G | 2.7114947791168254e-08 | 5.559112763189669 | 32158.0 |
| 17 | 17:44296230_T_A | 44296230 | T | A | 2.718589254668265e-08 | 5.558656631501583 | 32062.0 |
| 17 | rs62057073 | 43861117 | C | T | 2.7222950526661038e-08 | 5.558418830239973 | 32186.0 |
| 17 | rs2532416 | 44289291 | A | C | 2.7238224081290686e-08 | 5.5583209111554215 | 31953.0 |
| 17 | rs2532271 | 44248042 | G | A | 2.7241545530461914e-08 | 5.558299624327044 | 32037.0 |
| 17 | rs60814418 | 43850645 | C | T | 2.727820271026079e-08 | 5.558064859341171 | 32137.0 |
| 17 | rs17689104 | 43782492 | A | G | 2.7297785649227886e-08 | 5.557939568995677 | 32175.0 |
| 17 | rs56367860 | 44034673 | T | G | 2.7366148514128024e-08 | 5.557502870549703 | 32126.0 |
| 17 | rs77017444 | 44028796 | T | C | 2.742397808762933e-08 | 5.5571342839068665 | 32149.0 |
| 17 | rs17762882 | 43898887 | T | C | 2.742646190220844e-08 | 5.557118469788735 | 32171.0 |
| 17 | rs17689653 | 43898963 | A | T | 2.742646190220844e-08 | 5.557118469788735 | 32171.0 |
| 17 | rs62061708 | 44006956 | A | G | 2.7432767960248893e-08 | 5.557078326195367 | 32037.0 |
| 17 | 17:44293205_A_G | 44293205 | A | G | 2.7467572365122473e-08 | 5.556856926578536 | 31186.0 |
| 17 | rs12185268 | 43923683 | A | G | 2.7476758758618253e-08 | 5.55679853501457 | 32186.0 |
| 17 | rs17689471 | 43892973 | T | C | 2.7525132621144225e-08 | 5.5564913679226215 | 32173.0 |
| 17 | rs62054387 | 43805525 | A | C | 2.7525803736099644e-08 | 5.556487110123924 | 32182.0 |
| 17 | rs55757142 | 43894485 | G | GA | 2.7547863578619313e-08 | 5.556347210460618 | 32096.0 |
| 17 | rs62056934 | 43790005 | A | G | 2.7548823095162643e-08 | 5.556341127841767 | 32115.0 |
| 17 | rs56023973 | 43917776 | G | T | 2.755947592873085e-08 | 5.556273610633731 | 32182.0 |
| 17 | rs17763515 | 43917818 | G | A | 2.755947592873085e-08 | 5.556273610633731 | 32182.0 |
| 17 | rs10514879 | 43802971 | C | T | 2.7592035258077573e-08 | 5.556067407854644 | 32178.0 |
| 17 | rs56082319 | 43844798 | G | A | 2.7607607893446665e-08 | 5.5559688676349115 | 32163.0 |
| 17 | rs1918792 | 44228529 | G | A | 2.762222708768328e-08 | 5.555876409651645 | 32039.0 |
| 17 | rs1918791 | 44228609 | C | G | 2.762222708768328e-08 | 5.555876409651645 | 32039.0 |
| 17 | rs2696576 | 44228770 | A | G | 2.762222708768328e-08 | 5.555876409651645 | 32039.0 |
| 17 | rs2696575 | 44228824 | A | G | 2.762222708768328e-08 | 5.555876409651645 | 32039.0 |
| 17 | rs2532300 | 44229986 | T | C | 2.762222708768328e-08 | 5.555876409651645 | 32039.0 |
| 17 | rs2696573 | 44230647 | C | T | 2.762222708768328e-08 | 5.555876409651645 | 32039.0 |
| 17 | rs2532299 | 44231117 | T | C | 2.762222708768328e-08 | 5.555876409651645 | 32039.0 |

| | | | | | | | |
|---|---|---|---|---|---|---|---|
| 17 | rs2532298 | 44231295 | G | A | 2.762222708768328e-08 | 5.555876409651645 | 32039.0 |
| 17 | rs2532297 | 44231326 | G | A | 2.762222708768328e-08 | 5.555876409651645 | 32039.0 |
| 17 | rs2696572 | 44231617 | A | T | 2.762222708768328e-08 | 5.555876409651645 | 32039.0 |
| 17 | rs2109092 | 44232959 | G | A | 2.762222708768328e-08 | 5.555876409651645 | 32039.0 |
| 17 | 17:43777957_AATC_A | 43777957 | AATC | A | 2.7666708336910344e-08 | 5.555595383113613 | 32072.0 |
| 17 | 17:43797157_TA_T | 43797157 | TA | T | 2.76892645768165e-08 | 5.5554530433141265 | 32182.0 |
| 17 | rs62063859 | 44078616 | G | A | 2.7736942509953992e-08 | 5.555152544465806 | 32181.0 |
| 17 | rs62064660 | 44078832 | G | A | 2.7736942509953992e-08 | 5.555152544465806 | 32181.0 |
| 17 | rs111520035 | 44039410 | A | G | 2.774814979779533e-08 | 5.555081981252034 | 32154.0 |
| 17 | rs11079724 | 43841912 | C | T | 2.7779573168309848e-08 | 5.554884281111722 | 32154.0 |
| 17 | rs17651093 | 44049550 | G | A | 2.7831473765036232e-08 | 5.554558223188331 | 32181.0 |
| 17 | rs111985258 | 43795859 | A | G | 2.784882739917257e-08 | 5.554449333118957 | 32183.0 |
| 17 | rs55943044 | 43872228 | G | A | 2.786842356637587e-08 | 5.554326450792828 | 32171.0 |
| 17 | rs4617909 | 43806451 | C | T | 2.788463456233309e-08 | 5.554224859322027 | 32179.0 |
| 17 | rs2532291 | 44238424 | G | A | 2.7914070927013094e-08 | 5.554040533205699 | 32039.0 |
| 17 | rs17662403 | 44238938 | T | C | 2.7914070927013094e-08 | 5.554040533205699 | 32039.0 |
| 17 | rs17585426 | 44238966 | T | C | 2.7914070927013094e-08 | 5.554040533205699 | 32039.0 |
| 17 | rs2532288 | 44239958 | C | T | 2.7914070927013094e-08 | 5.554040533205699 | 32039.0 |
| 17 | rs1918789 | 44240107 | T | C | 2.7914070927013094e-08 | 5.554040533205699 | 32039.0 |
| 17 | rs2141299 | 44240986 | C | A | 2.7914070927013094e-08 | 5.554040533205699 | 32039.0 |
| 17 | rs2696567 | 44241304 | C | G | 2.7914070927013094e-08 | 5.554040533205699 | 32039.0 |
| 17 | rs2532286 | 44241664 | T | C | 2.7914070927013094e-08 | 5.554040533205699 | 32039.0 |
| 17 | rs4792844 | 44242969 | T | C | 2.7914070927013094e-08 | 5.554040533205699 | 32039.0 |
| 17 | rs4792843 | 44243179 | A | G | 2.7914070927013094e-08 | 5.554040533205699 | 32039.0 |
| 17 | rs17585608 | 44243979 | C | T | 2.7914070927013094e-08 | 5.554040533205699 | 32039.0 |
| 17 | rs17585644 | 44244581 | T | C | 2.7914070927013094e-08 | 5.554040533205699 | 32039.0 |
| 17 | rs2696657 | 44244926 | T | G | 2.7914070927013094e-08 | 5.554040533205699 | 32039.0 |
| 17 | rs2696659 | 44245876 | A | G | 2.7914070927013094e-08 | 5.554040533205699 | 32039.0 |
| 17 | rs1880752 | 43806264 | C | T | 2.7948792458349235e-08 | 5.553823354455409 | 32180.0 |
| 17 | rs2902662 | 43806925 | G | A | 2.7948792458349235e-08 | 5.553823354455409 | 32180.0 |
| 17 | rs2864087 | 43807063 | C | T | 2.7948792458349235e-08 | 5.553823354455409 | 32180.0 |
| 17 | rs79226879 | 43807485 | T | C | 2.7948792458349235e-08 | 5.553823354455409 | 32180.0 |
| 17 | rs17563599 | 43807955 | A | C | 2.7948792458349235e-08 | 5.553823354455409 | 32180.0 |
| 17 | rs17649138 | 43808067 | G | A | 2.7948792458349235e-08 | 5.553823354455409 | 32180.0 |
| 17 | rs62054389 | 43808321 | G | A | 2.7948792458349235e-08 | 5.553823354455409 | 32180.0 |
| 17 | rs62054390 | 43808443 | A | C | 2.7948792458349235e-08 | 5.553823354455409 | 32180.0 |
| 17 | rs55947210 | 43808763 | A | G | 2.7948792458349235e-08 | 5.553823354455409 | 32180.0 |
| 17 | rs62054392 | 43808945 | C | T | 2.7948792458349235e-08 | 5.553823354455409 | 32180.0 |
| 17 | rs4390635 | 43809016 | C | T | 2.7948792458349235e-08 | 5.553823354455409 | 32180.0 |
| 17 | rs62054393 | 43809264 | T | G | 2.7948792458349235e-08 | 5.553823354455409 | 32180.0 |
| 17 | rs75986348 | 43809328 | A | G | 2.7948792458349235e-08 | 5.553823354455409 | 32180.0 |
| 17 | rs17649162 | 43810371 | G | C | 2.7948792458349235e-08 | 5.553823354455409 | 32180.0 |
| 17 | rs62054394 | 43810608 | T | A | 2.7948792458349235e-08 | 5.553823354455409 | 32180.0 |
| 17 | rs113790915 | 43810782 | G | C | 2.7948792458349235e-08 | 5.553823354455409 | 32180.0 |
| 17 | rs62054395 | 43811060 | T | G | 2.7948792458349235e-08 | 5.553823354455409 | 32180.0 |
| 17 | rs62054396 | 43811510 | T | C | 2.7948792458349235e-08 | 5.553823354455409 | 32180.0 |
| 17 | rs17563718 | 43811673 | C | T | 2.7948792458349235e-08 | 5.553823354455409 | 32180.0 |
| 17 | rs62054397 | 43811792 | C | T | 2.7948792458349235e-08 | 5.553823354455409 | 32180.0 |
| 17 | rs4569335 | 43811950 | A | C | 2.7948792458349235e-08 | 5.553823354455409 | 32180.0 |
| 17 | rs1526125 | 43812177 | C | T | 2.7948792458349235e-08 | 5.553823354455409 | 32180.0 |
| 17 | rs62055895 | 43844044 | A | G | 2.7988623494998248e-08 | 5.553574538587728 | 32164.0 |
| 17 | rs62055896 | 43844201 | C | T | 2.7988623494998248e-08 | 5.553574538587728 | 32164.0 |
| 17 | rs55725840 | 43844486 | T | A | 2.7988623494998248e-08 | 5.553574538587728 | 32164.0 |
| 17 | rs56109643 | 43844859 | A | G | 2.7988623494998248e-08 | 5.553574538587728 | 32164.0 |

| | | | | | | | |
|---|---|---|---|---|---|---|---|
| 17 | rs112320018 | 44030263 | A | AC | 2.802294804667896e-08 | 5.5533603960016205 | 32146.0 |
| 17 | rs2316772 | 43946423 | A | C | 2.8049304493849017e-08 | 5.553196137168629 | 32125.0 |
| 17 | rs4277389 | 43895653 | A | G | 2.813704101660211e-08 | 5.552650422004256 | 32168.0 |
| 17 | rs2696660 | 44246211 | A | G | 2.8147230750427633e-08 | 5.552587149604854 | 32038.0 |
| 17 | rs2532278 | 44246405 | C | A | 2.8147230750427633e-08 | 5.552587149604854 | 32038.0 |
| 17 | 17:44246624_C_A | 44246624 | C | A | 2.8147230750427633e-08 | 5.552587149604854 | 32038.0 |
| 17 | rs2532275 | 44246997 | G | A | 2.8147230750427633e-08 | 5.552587149604854 | 32038.0 |
| 17 | rs2532273 | 44247314 | C | T | 2.8147230750427633e-08 | 5.552587149604854 | 32038.0 |
| 17 | rs2532272 | 44247673 | G | A | 2.8147230750427633e-08 | 5.552587149604854 | 32038.0 |
| 17 | rs2316764 | 43895602 | T | G | 2.8155757632764207e-08 | 5.552534219637443 | 32166.0 |
| 17 | rs111355814 | 43977748 | G | GGCT GGCC CATA | 2.826890429788565e-08 | 5.551833339108727 | 31660.0 |
| 17 | rs79065019 | 44124535 | T | C | 2.829992034462458e-08 | 5.5516416873125145 | 32186.0 |
| 17 | rs35282504 | 43832618 | G | A | 2.8334125833916365e-08 | 5.551430563752907 | 32170.0 |
| 17 | rs76669427 | 43836051 | T | A | 2.8367680598491984e-08 | 5.551223696755415 | 32169.0 |
| 17 | rs62057067 | 43858326 | C | T | 2.837577904524846e-08 | 5.551173804945358 | 32168.0 |
| 17 | rs62057158 | 43907143 | G | A | 2.8379532752253477e-08 | 5.551150684303316 | 32182.0 |
| 17 | rs62057068 | 43858629 | A | G | 2.842421944773628e-08 | 5.5508756680609945 | 32173.0 |
| 17 | rs77692262 | 43917086 | A | G | 2.8435896790239756e-08 | 5.550803871090021 | 32183.0 |
| 17 | rs1881193 | 44248769 | T | C | 2.8523229733062368e-08 | 5.550267818504508 | 32036.0 |
| 17 | rs17688944 | 43779419 | T | A | 2.8553724155116826e-08 | 5.550081017748939 | 32180.0 |
| 17 | rs17571739 | 44032915 | T | C | 2.8617796758346897e-08 | 5.549689155760318 | 32094.0 |
| 17 | 17:43811990_CTTAG_C | 43811990 | CTTAG | C | 2.8620984162194948e-08 | 5.5496696841335 | 32171.0 |
| 17 | rs56288604 | 43931380 | A | G | 2.8633139331440204e-08 | 5.549595448362006 | 32157.0 |
| 17 | rs78506181 | 43897480 | A | G | 2.863473382720533e-08 | 5.549585712500298 | 32182.0 |
| 17 | rs62055875 | 43832864 | T | C | 2.8641910135092776e-08 | 5.5495419010570775 | 32169.0 |
| 17 | rs2316769 | 43930798 | A | G | 2.8674824105146826e-08 | 5.549341097237346 | 32167.0 |
| 17 | rs62054442 | 43829353 | A | G | 2.867771831618432e-08 | 5.549323450733422 | 32164.0 |
| 17 | rs2532344 | 44343930 | A | G | 2.8698185560563112e-08 | 5.549198707709687 | 30783.0 |
| 17 | rs62060864 | 44235893 | C | T | 2.8740463228104864e-08 | 5.548941308382294 | 32037.0 |
| 17 | rs1528072 | 44236725 | C | A | 2.8740463228104864e-08 | 5.548941308382294 | 32037.0 |
| 17 | rs2532292 | 44237068 | A | T | 2.8740463228104864e-08 | 5.548941308382294 | 32037.0 |
| 17 | rs2696571 | 44237372 | G | C | 2.8740463228104864e-08 | 5.548941308382294 | 32037.0 |
| 17 | rs2696570 | 44237790 | G | A | 2.8740463228104864e-08 | 5.548941308382294 | 32037.0 |
| 17 | rs76527351 | 44212527 | A | G | 2.8802540815049885e-08 | 5.548564026085875 | 32130.0 |
| 17 | rs17689116 | 43782558 | T | C | 2.8855912127436797e-08 | 5.548240287197462 | 32170.0 |
| 17 | rs2903705 | 43573231 | C | G | 2.8857016691709816e-08 | 5.548233593286502 | 32136.0 |
| 17 | 17:44270809_A_G | 44270809 | A | G | 2.8864247590265453e-08 | 5.548189778534773 | 32030.0 |
| 17 | rs62065452 | 43573061 | T | C | 2.8881830094765843e-08 | 5.548083283899564 | 31991.0 |
| 17 | rs56302057 | 44034486 | A | G | 2.890334525218248e-08 | 5.547953055313376 | 32129.0 |
| 17 | rs112954158 | 43671066 | G | GA | 2.8906764993926668e-08 | 5.547932364703136 | 31900.0 |
| 17 | rs58879558 | 44095467 | T | C | 2.8956395831721048e-08 | 5.547632348167942 | 32165.0 |
| 17 | rs62061798 | 44173812 | G | A | 2.8966876284587245e-08 | 5.547569058033651 | 32113.0 |
| 17 | rs62057152 | 43904110 | G | C | 2.9020142110379302e-08 | 5.547247735439372 | 32184.0 |
| 17 | rs17689824 | 43904397 | C | T | 2.9020142110379302e-08 | 5.547247735439372 | 32184.0 |
| 17 | rs62057153 | 43904528 | T | C | 2.9020142110379302e-08 | 5.547247735439372 | 32184.0 |
| 17 | rs62057154 | 43904610 | C | T | 2.9020142110379302e-08 | 5.547247735439372 | 32184.0 |
| 17 | rs62057155 | 43904673 | C | T | 2.9020142110379302e-08 | 5.547247735439372 | 32184.0 |
| 17 | rs62057156 | 43904948 | G | C | 2.9020142110379302e-08 | 5.547247735439372 | 32184.0 |
| 17 | rs78872653 | 43905134 | G | A | 2.9020142110379302e-08 | 5.547247735439372 | 32184.0 |
| 17 | rs62057157 | 43905313 | G | C | 2.9020142110379302e-08 | 5.547247735439372 | 32184.0 |
| 17 | rs17763086 | 43905481 | T | G | 2.9020142110379302e-08 | 5.547247735439372 | 32184.0 |
| 17 | rs62055871 | 43831112 | G | A | 2.9047922733010406e-08 | 5.547080377637774 | 32177.0 |
| 17 | rs56201245 | 44034340 | C | T | 2.9052976554661964e-08 | 5.547049948778516 | 32130.0 |
| 17 | rs56369036 | 43841571 | T | A | 2.9058435649910683e-08 | 5.547017085552438 | 32161.0 |

| | | | | | | | |
|---|---|---|---|---|---|---|---|
| 17 | 17:43775212_CCTT_C | 43775212 | CCTT | C | 2.9064603614845525e-08 | 5.546979962205985 | 32115.0 |
| 17 | rs2668653 | 44288156 | T | C | 2.9097386121135187e-08 | 5.546782781174061 | 31923.0 |
| 17 | rs2458210 | 44298102 | G | A | 2.9146120817988486e-08 | 5.546490048538688 | 32090.0 |
| 17 | rs117646503 | 43893260 | T | C | 2.9180820748682804e-08 | 5.546281907246488 | 32182.0 |
| 17 | rs17762769 | 43893403 | G | A | 2.9180820748682804e-08 | 5.546281907246488 | 32182.0 |
| 17 | rs28364025 | 43894102 | T | C | 2.9180820748682804e-08 | 5.546281907246488 | 32182.0 |
| 17 | rs28364023 | 43894159 | C | T | 2.9180820748682804e-08 | 5.546281907246488 | 32182.0 |
| 17 | rs55779147 | 43894510 | G | A | 2.9180820748682804e-08 | 5.546281907246488 | 32182.0 |
| 17 | rs17688922 | 43779351 | G | A | 2.9185592615749843e-08 | 5.5462533028447165 | 32155.0 |
| 17 | rs112460810 | 43900215 | T | TAA | 2.9338288559312564e-08 | 5.545340372403554 | 32180.0 |
| 17 | 17:44094133_CA_C | 44094133 | CA | C | 2.9348496079512536e-08 | 5.545279508719148 | 32061.0 |
| 17 | rs750783539 | 44137009 | AAAAAAGAG | A | 2.937873074165259e-08 | 5.545099351000823 | 32164.0 |
| 17 | 17:43796396_TG_T | 43796396 | TG | T | 2.9405517968571584e-08 | 5.544939885562178 | 32177.0 |
| 17 | rs62056875 | 43778003 | T | A | 2.9443388031965017e-08 | 5.544714683859642 | 32181.0 |
| 17 | rs17762165 | 43778602 | C | T | 2.9443388031965017e-08 | 5.544714683859642 | 32181.0 |
| 17 | rs62063282 | 44039043 | G | A | 2.9500795814255632e-08 | 5.544373832652044 | 32134.0 |
| 17 | 17:44102741_TCA_T | 44102741 | TCA | T | 2.9531903242561263e-08 | 5.544189405097115 | 32130.0 |
| 17 | rs4792845 | 44242788 | C | T | 2.953858063963084e-08 | 5.544149841182759 | 32033.0 |
| 17 | rs78440279 | 43816052 | C | T | 2.954844534789572e-08 | 5.544091408164735 | 32173.0 |
| 17 | rs17688682 | 43775929 | C | G | 2.9549678663409538e-08 | 5.544084104024076 | 32180.0 |
| 17 | rs12150454 | 43776061 | C | T | 2.957805884768067e-08 | 5.5439161079665435 | 32178.0 |
| 17 | rs4401083 | 43806015 | G | A | 2.9599464461553525e-08 | 5.543789501200519 | 32178.0 |
| 17 | rs62055878 | 43835854 | C | A | 2.9608937170911635e-08 | 5.5437335017684655 | 32173.0 |
| 17 | rs62057143 | 43901528 | T | C | 2.9619030933488715e-08 | 5.543673850006902 | 32186.0 |
| 17 | rs62057146 | 43903106 | A | G | 2.9630673845602518e-08 | 5.543605067626999 | 32181.0 |
| 17 | 17:43777835_TATCCTGCTTCCC_T | 43777835 | TATCCTGCTTCCC | T | 2.966831213164293e-08 | 5.543382892647902 | 32117.0 |
| 17 | rs739644 | 43895008 | C | G | 2.9678632164847268e-08 | 5.543322022309524 | 32183.0 |
| 17 | rs12150390 | 43896228 | T | C | 2.9678632164847268e-08 | 5.543322022309524 | 32183.0 |
| 17 | rs62057112 | 43897202 | T | A | 2.9678632164847268e-08 | 5.543322022309524 | 32183.0 |
| 17 | rs78587102 | 43897246 | A | G | 2.9678632164847268e-08 | 5.543322022309524 | 32183.0 |
| 17 | rs79959255 | 44077708 | G | T | 2.970878679982161e-08 | 5.543144279737182 | 32177.0 |
| 17 | rs62054380 | 43800987 | C | A | 2.9758938773474378e-08 | 5.542849052729998 | 32177.0 |
| 17 | rs17563501 | 43801695 | C | T | 2.9758938773474378e-08 | 5.542849052729998 | 32177.0 |
| 17 | rs62063271 | 44036047 | G | A | 2.9783683345720328e-08 | 5.542703567940234 | 32130.0 |
| 17 | rs62054811 | 43918651 | C | G | 2.9791245330351006e-08 | 5.542659130927782 | 32181.0 |
| 17 | rs17650901 | 44039691 | A | G | 2.9824208534493324e-08 | 5.542465554710308 | 32157.0 |
| 17 | rs62056881 | 43782455 | G | T | 2.988868315964689e-08 | 5.542087527195354 | 32170.0 |
| 17 | rs62057144 | 43901558 | A | G | 2.9914053545363036e-08 | 5.541938992461254 | 32186.0 |
| 17 | rs2696671 | 44316888 | A | G | 2.9937154771922944e-08 | 5.541803849132683 | 32186.0 |
| 17 | rs41280118 | 43903089 | C | T | 3.004017785172413e-08 | 5.541202388074288 | 32184.0 |
| 17 | rs62057148 | 43903485 | G | C | 3.004017785172413e-08 | 5.541202388074288 | 32184.0 |
| 17 | rs62057149 | 43903546 | A | G | 3.004017785172413e-08 | 5.541202388074288 | 32184.0 |
| 17 | rs62057150 | 43903548 | C | T | 3.004017785172413e-08 | 5.541202388074288 | 32184.0 |
| 17 | rs17690679 | 43924803 | A | G | 3.0080731249699966e-08 | 5.540966181243062 | 32183.0 |
| 17 | rs1876831 | 43907745 | C | T | 3.030663213519985e-08 | 5.539656029052926 | 32183.0 |
| 17 | rs16940665 | 43907896 | T | C | 3.030663213519985e-08 | 5.539656029052926 | 32183.0 |
| 17 | rs55763795 | 43908773 | G | C | 3.030663213519985e-08 | 5.539656029052926 | 32183.0 |
| 17 | rs55865707 | 43908826 | T | G | 3.030663213519985e-08 | 5.539656029052926 | 32183.0 |
| 17 | rs62062806 | 44031420 | G | A | 3.0310530619863484e-08 | 5.539633502363817 | 32153.0 |
| 17 | rs55978005 | 44031540 | G | T | 3.0310530619863484e-08 | 5.539633502363817 | 32153.0 |
| 17 | rs17571857 | 44035706 | A | G | 3.034310743012957e-08 | 5.539445372961671 | 32131.0 |
| 17 | rs56280222 | 43803788 | T | C | 3.040403043909232e-08 | 5.539094071019646 | 32184.0 |
| 17 | rs62054820 | 43926755 | A | G | 3.043533033969961e-08 | 5.538913851057498 | 32173.0 |
| 17 | rs112570965 | 43801092 | T | C | 3.050300775687821e-08 | 5.538524788941274 | 32179.0 |

| | | | | | | | |
|---|---|---|---|---|---|---|---|
| 17 | rs17649019 | 43801593 | G | A | 3.050300775687821e-08 | 5.538524788941274 | 32179.0 |
| 17 | rs62054763 | 43910262 | C | G | 3.052146188077649e-08 | 5.5384188456800425 | 32182.0 |
| 17 | rs16940674 | 43910507 | C | T | 3.052146188077649e-08 | 5.5384188456800425 | 32182.0 |
| 17 | rs1876830 | 43911352 | C | T | 3.052146188077649e-08 | 5.5384188456800425 | 32182.0 |
| 17 | rs1876829 | 43911443 | T | C | 3.052146188077649e-08 | 5.5384188456800425 | 32182.0 |
| 17 | rs1876828 | 43911525 | C | T | 3.052146188077649e-08 | 5.5384188456800425 | 32182.0 |
| 17 | rs1876827 | 43911832 | T | C | 3.052146188077649e-08 | 5.5384188456800425 | 32182.0 |
| 17 | rs28364021 | 43912282 | C | T | 3.052146188077649e-08 | 5.5384188456800425 | 32182.0 |
| 17 | rs17763199 | 43910183 | G | A | 3.0536742532620314e-08 | 5.5383311680243095 | 32175.0 |
| 17 | rs4335809 | 43902216 | C | T | 3.055564147739159e-08 | 5.5382227881065464 | 32185.0 |
| 17 | rs4341787 | 43902505 | C | T | 3.055564147739159e-08 | 5.5382227881065464 | 32185.0 |
| 17 | rs4523962 | 43902522 | G | A | 3.055564147739159e-08 | 5.5382227881065464 | 32185.0 |
| 17 | rs3885074 | 43902738 | C | A | 3.055564147739159e-08 | 5.5382227881065464 | 32185.0 |
| 17 | rs3885075 | 43902799 | A | G | 3.055564147739159e-08 | 5.5382227881065464 | 32185.0 |
| 17 | rs41280116 | 43902842 | C | T | 3.055564147739159e-08 | 5.5382227881065464 | 32185.0 |
| 17 | rs34283254 | 43902861 | C | A | 3.055564147739159e-08 | 5.5382227881065464 | 32185.0 |
| 17 | rs1396862 | 43902997 | G | A | 3.055564147739159e-08 | 5.5382227881065464 | 32185.0 |
| 17 | rs80190331 | 44035233 | A | T | 3.0602300802622316e-08 | 5.537955488763862 | 32135.0 |
| 17 | rs62054435 | 43825339 | C | G | 3.066661136535968e-08 | 5.5375877173152395 | 32178.0 |
| 17 | 17:43855501_AC_A | 43855501 | AC | A | 3.067311366456745e-08 | 5.537550574414557 | 32096.0 |
| 17 | rs62054440 | 43827391 | G | C | 3.068729549323072e-08 | 5.537469590445952 | 32166.0 |
| 17 | rs12150451 | 43827431 | A | G | 3.068729549323072e-08 | 5.537469590445952 | 32166.0 |
| 17 | rs62073112 | 44324572 | A | G | 3.069700255167919e-08 | 5.537414180151142 | 31843.0 |
| 17 | rs12150048 | 43827471 | G | C | 3.0739706633931027e-08 | 5.5371706163824275 | 32165.0 |
| 17 | rs56026128 | 43931244 | G | C | 3.0760011461312155e-08 | 5.537054922431726 | 32141.0 |
| 17 | rs62055933 | 43848461 | G | A | 3.076022526622324e-08 | 5.537053704596691 | 32122.0 |
| 17 | rs62055883 | 43837446 | C | G | 3.0785036520777606e-08 | 5.536912435170517 | 32154.0 |
| 17 | rs62054815 | 43923266 | G | A | 3.0796165172341164e-08 | 5.536849107134931 | 32186.0 |
| 17 | rs17762308 | 43780948 | T | C | 3.080772602152142e-08 | 5.536783343168619 | 32158.0 |
| 17 | rs16940681 | 43912159 | G | C | 3.0820576397410083e-08 | 5.5367102718116685 | 32170.0 |
| 17 | rs2907480 | 44327712 | C | T | 3.083761123624525e-08 | 5.536613451804422 | 31636.0 |
| 17 | rs4340365 | 43930980 | C | T | 3.0898003771258036e-08 | 5.536270619519828 | 32147.0 |
| 17 | rs55638417 | 43900434 | A | G | 3.093141415698994e-08 | 5.5360812368822145 | 32175.0 |
| 17 | rs55746658 | 44034575 | T | C | 3.095001467656803e-08 | 5.535975888117147 | 32131.0 |
| 17 | rs62063275 | 44036287 | A | G | 3.098864842187824e-08 | 5.5357572722282775 | 32136.0 |
| 17 | rs1034927 | 44334419 | C | T | 3.112007583139072e-08 | 5.535015541607523 | 32186.0 |
| 17 | rs17662115 | 44227531 | T | C | 3.115956714970728e-08 | 5.534793260027852 | 31968.0 |
| 17 | rs17426174 | 43830938 | G | C | 3.1185235748722455e-08 | 5.5346489277712525 | 32177.0 |
| 17 | rs62054839 | 43936804 | A | G | 3.11929295304855e-08 | 5.534605688767506 | 32077.0 |
| 17 | rs62055550 | 43965680 | G | A | 3.124868367697357e-08 | 5.534292659626446 | 32170.0 |
| 17 | rs62055551 | 43965687 | C | G | 3.124868367697357e-08 | 5.534292659626446 | 32170.0 |
| 17 | rs368673667 | 43708418 | C | G | 3.126410378125077e-08 | 5.534206179794 | 32156.0 |
| 17 | rs16940668 | 43907966 | G | A | 3.1270838841138594e-08 | 5.534168420862933 | 32180.0 |
| 17 | 17:43814370_AC_A | 43814370 | AC | A | 3.1281270126084984e-08 | 5.5341099552641975 | 32168.0 |
| 17 | rs12150608 | 43775402 | T | C | 3.1318350493474866e-08 | 5.53390227904208 | 32178.0 |
| 17 | rs17690987 | 43931626 | T | C | 3.136342677908584e-08 | 5.533650141036731 | 32157.0 |
| 17 | rs62054426 | 43820741 | A | T | 3.136800328595502e-08 | 5.533624561620443 | 32179.0 |
| 17 | rs62054427 | 43821062 | A | G | 3.136800328595502e-08 | 5.533624561620443 | 32179.0 |
| 17 | rs74464991 | 43821685 | C | A | 3.136800328595502e-08 | 5.533624561620443 | 32179.0 |
| 17 | rs2004260 | 43822398 | T | C | 3.136800328595502e-08 | 5.533624561620443 | 32179.0 |
| 17 | rs75643244 | 43823326 | A | G | 3.136800328595502e-08 | 5.533624561620443 | 32179.0 |
| 17 | rs62054431 | 43823880 | C | T | 3.136800328595502e-08 | 5.533624561620443 | 32179.0 |
| 17 | rs62054432 | 43823889 | C | A | 3.136800328595502e-08 | 5.533624561620443 | 32179.0 |
| 17 | rs111497072 | 43824201 | G | A | 3.136800328595502e-08 | 5.533624561620443 | 32179.0 |

| | | | | | | | |
|---|---|---|---|---|---|---|---|
| 17 | rs62054433 | 43824348 | T | C | 3.136800328595502e-08 | 5.533624561620443 | 32179.0 |
| 17 | rs62054434 | 43824382 | T | G | 3.136800328595502e-08 | 5.533624561620443 | 32179.0 |
| 17 | rs12150683 | 43824848 | A | G | 3.136800328595502e-08 | 5.533624561620443 | 32179.0 |
| 17 | rs12150332 | 43824908 | C | T | 3.136800328595502e-08 | 5.533624561620443 | 32179.0 |
| 17 | rs75715199 | 43825478 | G | A | 3.136800328595502e-08 | 5.533624561620443 | 32179.0 |
| 17 | rs62054438 | 43825711 | A | C | 3.136800328595502e-08 | 5.533624561620443 | 32179.0 |
| 17 | rs62054439 | 43825725 | G | T | 3.136800328595502e-08 | 5.533624561620443 | 32179.0 |
| 17 | rs17334797 | 43825912 | A | G | 3.136800328595502e-08 | 5.533624561620443 | 32179.0 |
| 17 | rs12150363 | 43826351 | A | G | 3.136800328595502e-08 | 5.533624561620443 | 32179.0 |
| 17 | rs12150672 | 43826637 | G | A | 3.136800328595502e-08 | 5.533624561620443 | 32179.0 |
| 17 | rs17334894 | 43827209 | G | A | 3.136800328595502e-08 | 5.533624561620443 | 32179.0 |
| 17 | rs12150604 | 43828221 | T | A | 3.136800328595502e-08 | 5.533624561620443 | 32179.0 |
| 17 | rs17334944 | 43828617 | G | A | 3.136800328595502e-08 | 5.533624561620443 | 32179.0 |
| 17 | rs17426064 | 43828698 | C | T | 3.136800328595502e-08 | 5.533624561620443 | 32179.0 |
| 17 | rs17426106 | 43828935 | G | C | 3.136800328595502e-08 | 5.533624561620443 | 32179.0 |
| 17 | 17:43928629_GCACA_G | 43928629 | GCACA | G | 3.140409342597686e-08 | 5.533422970180063 | 32084.0 |
| 17 | rs12150547 | 43775546 | A | G | 3.145387941932075e-08 | 5.5331452453247865 | 32181.0 |
| 17 | rs4566212 | 43895751 | G | A | 3.148525367313643e-08 | 5.5329704469901655 | 32176.0 |
| 17 | rs55645418 | 43832115 | A | G | 3.1530784388935636e-08 | 5.532717077733919 | 32178.0 |
| 17 | rs7350923 | 43834970 | C | T | 3.158504562753334e-08 | 5.532415588010599 | 32176.0 |
| 17 | rs11079717 | 43835351 | T | C | 3.158504562753334e-08 | 5.532415588010599 | 32176.0 |
| 17 | rs757501 | 43756685 | A | G | 3.160984986014779e-08 | 5.532277936469312 | 32178.0 |
| 17 | rs62057118 | 43899727 | A | G | 3.164214588107248e-08 | 5.532098866066528 | 32162.0 |
| 17 | rs1912151 | 43902944 | C | T | 3.166655409573922e-08 | 5.531963648453353 | 32185.0 |
| 17 | rs413917 | 43723189 | G | A | 3.1694943579553196e-08 | 5.531806502376847 | 32186.0 |
| 17 | rs17761985 | 43776787 | C | T | 3.1699347282376545e-08 | 5.5317821385205805 | 32181.0 |
| 17 | rs56380663 | 43804619 | C | T | 3.17041920479587e-08 | 5.531755338240398 | 32186.0 |
| 17 | rs62054429 | 43823125 | G | A | 3.172732478973945e-08 | 5.531627427259713 | 32178.0 |
| 17 | rs28439278 | 43838071 | C | T | 3.1733606530115284e-08 | 5.531592708407307 | 32159.0 |
| 17 | rs17688767 | 43777542 | A | G | 3.1756980893607535e-08 | 5.531463578049686 | 32182.0 |
| 17 | rs62056874 | 43777578 | C | T | 3.1756980893607535e-08 | 5.531463578049686 | 32182.0 |
| 17 | rs17688773 | 43777710 | T | C | 3.1756980893607535e-08 | 5.531463578049686 | 32182.0 |
| 17 | rs12150091 | 43776242 | A | G | 3.1787326549620783e-08 | 5.5312960726607 | 32180.0 |
| 17 | rs12150464 | 43776371 | C | T | 3.1787326549620783e-08 | 5.5312960726607 | 32180.0 |
| 17 | rs12150435 | 43776391 | G | C | 3.1787326549620783e-08 | 5.5312960726607 | 32180.0 |
| 17 | rs11079725 | 43923934 | T | C | 3.182189999875133e-08 | 5.5311054191659235 | 32186.0 |
| 17 | rs12373139 | 43924130 | G | A | 3.182189999875133e-08 | 5.5311054191659235 | 32186.0 |
| 17 | rs112916880 | 44036129 | C | T | 3.184489570237336e-08 | 5.53097872179116 | 32135.0 |
| 17 | rs36047693 | 43833824 | G | C | 3.1848545555517685e-08 | 5.5309586206829655 | 32177.0 |
| 17 | rs34195569 | 43833921 | G | A | 3.1848545555517685e-08 | 5.5309586206829655 | 32177.0 |
| 17 | rs34211253 | 43834074 | C | G | 3.1848545555517685e-08 | 5.5309586206829655 | 32177.0 |
| 17 | rs34008514 | 43834314 | A | G | 3.1848545555517685e-08 | 5.5309586206829655 | 32177.0 |
| 17 | rs12943085 | 43834482 | G | A | 3.1848545555517685e-08 | 5.5309586206829655 | 32177.0 |
| 17 | rs188710438 | 43573649 | C | T | 3.1865804956464104e-08 | 5.530863596956962 | 32111.0 |
| 17 | rs77849344 | 43840864 | G | A | 3.190245622421044e-08 | 5.530661974432441 | 32156.0 |
| 17 | rs17762073 | 43777745 | C | A | 3.1929722289630223e-08 | 5.530512126614388 | 32181.0 |
| 17 | rs2668716 | 44343371 | G | A | 3.1967111652889e-08 | 5.530306845250906 | 31286.0 |
| 17 | rs17650771 | 44035367 | A | G | 3.1981768434872526e-08 | 5.5302264376659584 | 32133.0 |
| 17 | rs12373123 | 43924073 | T | C | 3.1998542980034895e-08 | 5.530134455819222 | 32186.0 |
| 17 | rs41457044 | 43911424 | C | T | 3.200298801233503e-08 | 5.530110089687952 | 32181.0 |
| 17 | rs62054824 | 43929992 | C | T | 3.2013325051264305e-08 | 5.530053438304592 | 32163.0 |
| 17 | rs62054424 | 43819384 | G | A | 3.205737746719399e-08 | 5.529812211053032 | 32179.0 |
| 17 | rs17563923 | 43819450 | G | A | 3.205737746719399e-08 | 5.529812211053032 | 32179.0 |
| 17 | rs62054425 | 43819480 | C | T | 3.205737746719399e-08 | 5.529812211053032 | 32179.0 |

| | | | | | | | |
|---|---|---|---|---|---|---|---|
| 17 | rs17690661 | 43924521 | G | A | 3.205982656449533e-08 | 5.529798809443776 | 32183.0 |
| 17 | rs55943825 | 43924776 | C | T | 3.205982656449533e-08 | 5.529798809443776 | 32183.0 |
| 17 | rs62056870 | 43773784 | C | A | 3.208053989774584e-08 | 5.529685504528321 | 32182.0 |
| 17 | rs62054381 | 43801185 | A | G | 3.210148928028927e-08 | 5.529570980551921 | 32178.0 |
| 17 | rs12948216 | 43836673 | C | T | 3.2149344196696114e-08 | 5.529309643841941 | 32167.0 |
| 17 | rs62055692 | 43754909 | A | G | 3.21583875823211e-08 | 5.529260300119018 | 32177.0 |
| 17 | rs62054831 | 43932381 | C | T | 3.2207555709221176e-08 | 5.5289922577673245 | 32147.0 |
| 17 | rs2696626 | 44327070 | G | A | 3.221270026443182e-08 | 5.528964234926182 | 31104.0 |
| 17 | rs968027 | 43781250 | C | T | 3.226553441521052e-08 | 5.5286766936748775 | 32128.0 |
| 17 | rs17563433 | 43800351 | A | C | 3.2320361043739246e-08 | 5.528378791195427 | 32178.0 |
| 17 | rs77370070 | 43800404 | T | G | 3.2320361043739246e-08 | 5.528378791195427 | 32178.0 |
| 17 | rs17687740 | 43756376 | G | C | 3.235617647888565e-08 | 5.528184451324166 | 32174.0 |
| 17 | rs62063264 | 44033401 | C | T | 3.2405753402895314e-08 | 5.527915783640228 | 32024.0 |
| 17 | rs62055869 | 43830685 | G | A | 3.2407891041817087e-08 | 5.527904208297964 | 32178.0 |
| 17 | rs17688410 | 43772251 | C | T | 3.241036637528847e-08 | 5.5278908052607365 | 32183.0 |
| 17 | rs4479290 | 44021744 | T | C | 3.2434229084855484e-08 | 5.5277616482060665 | 32041.0 |
| 17 | rs62057122 | 43901001 | C | T | 3.244076056065516e-08 | 5.527726312623166 | 32168.0 |
| 17 | rs56205560 | 43804389 | T | A | 3.247591751529355e-08 | 5.527536230360159 | 32184.0 |
| 17 | rs2532230 | 44275619 | A | G | 3.2489674936846025e-08 | 5.527461902772695 | 31899.0 |
| 17 | rs55989971 | 43798160 | G | A | 3.251675538544118e-08 | 5.527315683667462 | 32178.0 |
| 17 | rs55938136 | 43798360 | A | G | 3.251675538544118e-08 | 5.527315683667462 | 32178.0 |
| 17 | 17:43908476_GT_G | 43908476 | GT | G | 3.2520367782570045e-08 | 5.527296187696557 | 32180.0 |
| 17 | rs201825478 | 43774408 | A | AAGAAAGAAAGAG | 3.253764499177018e-08 | 5.527202972292327 | 32177.0 |
| 17 | rs9906974 | 43822772 | T | C | 3.261567362456833e-08 | 5.526782583068363 | 31556.0 |
| 17 | rs112556326 | 43575946 | G | T | 3.2631188067848216e-08 | 5.526699113309064 | 32108.0 |
| 17 | rs1724397 | 43665809 | C | T | 3.2657363834575606e-08 | 5.526558371446995 | 32025.0 |
| 17 | rs1724398 | 43665838 | A | G | 3.2657363834575606e-08 | 5.526558371446995 | 32025.0 |
| 17 | rs56046792 | 43797246 | C | T | 3.270126285686974e-08 | 5.526322580653758 | 32177.0 |
| 17 | rs56323722 | 43798015 | A | C | 3.270126285686974e-08 | 5.526322580653758 | 32177.0 |
| 17 | 17:44327642_AC_A | 44327642 | AC | A | 3.271920207256715e-08 | 5.526226313752642 | 32105.0 |
| 17 | rs62055868 | 43830640 | C | T | 3.275761075372185e-08 | 5.526020373960684 | 32177.0 |
| 17 | rs75403953 | 43798546 | A | G | 3.2784682422357345e-08 | 5.525875361680325 | 32179.0 |
| 17 | rs16940676 | 43911036 | G | A | 3.281109271106217e-08 | 5.525734004041506 | 32182.0 |
| 17 | rs62056873 | 43776945 | T | C | 3.2830458310880055e-08 | 5.525630422304684 | 32181.0 |
| 17 | rs17573447 | 44079567 | C | T | 3.285770300749895e-08 | 5.525484797555556 | 32166.0 |
| 17 | 17:44184819_CAAAGGT_C | 44184819 | CAAAGGT | C | 3.286363365760183e-08 | 5.525453113352868 | 31707.0 |
| 17 | rs80072429 | 43818611 | G | A | 3.2907461710856224e-08 | 5.525219135966747 | 32182.0 |
| 17 | rs77114624 | 43818657 | T | C | 3.2907461710856224e-08 | 5.525219135966747 | 32182.0 |
| 17 | rs76311426 | 43818690 | G | T | 3.2907461710856224e-08 | 5.525219135966747 | 32182.0 |
| 17 | rs17563861 | 43818906 | A | G | 3.2907461710856224e-08 | 5.525219135966747 | 32182.0 |
| 17 | rs17563889 | 43818946 | A | T | 3.2907461710856224e-08 | 5.525219135966747 | 32182.0 |
| 17 | rs62054825 | 43930033 | G | A | 3.2910088645755264e-08 | 5.525205121599192 | 32164.0 |
| 17 | rs56385754 | 43927255 | G | T | 3.2914543483234385e-08 | 5.525181358081311 | 32170.0 |
| 17 | rs62063263 | 44033394 | G | C | 3.291968441453706e-08 | 5.525153938598417 | 32020.0 |
| 17 | rs113927637 | 43807911 | A | G | 3.293236874253001e-08 | 5.5250863036944455 | 32174.0 |
| 17 | rs62066460 | 43574935 | C | T | 3.2948831223202825e-08 | 5.5249985607356775 | 32109.0 |
| 17 | rs62054804 | 43914809 | C | T | 3.2956607975052136e-08 | 5.524957126411263 | 32183.0 |
| 17 | rs377660109 | 44263022 | C | CAT | 3.2959925078014505e-08 | 5.5249394558613965 | 32010.0 |
| 17 | rs1052553 | 44073889 | A | G | 3.2962670514064715e-08 | 5.5249248319448725 | 32164.0 |
| 17 | rs2907478 | 44331297 | G | T | 3.306544203108806e-08 | 5.524378254500624 | 32123.0 |
| 17 | rs79322350 | 43790401 | A | G | 3.325161480797955e-08 | 5.523392301160449 | 32172.0 |
| 17 | rs4479289 | 43793951 | G | A | 3.3306233598385544e-08 | 5.523104061036136 | 32175.0 |
| 17 | rs4609898 | 43794182 | G | A | 3.3306233598385544e-08 | 5.523104061036136 | 32175.0 |
| 17 | rs4605208 | 43794209 | G | C | 3.3306233598385544e-08 | 5.523104061036136 | 32175.0 |

| | | | | | | | |
|---|---|---|---|---|---|---|---|
| 17 | rs75310534 | 43794286 | C | T | 3.3306233598385544e-08 | 5.523104061036136 | 32175.0 |
| 17 | rs113856644 | 43932277 | G | A | 3.338236114446653e-08 | 5.5227030765720215 | 32160.0 |
| 17 | rs17563683 | 43811072 | A | G | 3.339104911279691e-08 | 5.5226573710858835 | 32179.0 |
| 17 | rs79882507 | 43837782 | A | G | 3.3464928294926025e-08 | 5.522269174454172 | 32158.0 |
| 17 | rs4564621 | 43895501 | G | C | 3.3480373817520515e-08 | 5.522188121348282 | 32177.0 |
| 17 | rs2316763 | 43895530 | C | T | 3.348873821734327e-08 | 5.522144242822124 | 32175.0 |
| 17 | rs4566211 | 43895696 | G | A | 3.348873821734327e-08 | 5.522144242822124 | 32175.0 |
| 17 | rs56226437 | 44042361 | T | C | 3.3512797273595925e-08 | 5.5220180914605415 | 32064.0 |
| 17 | rs55907036 | 44042366 | G | A | 3.3512797273595925e-08 | 5.5220180914605415 | 32064.0 |
| 17 | rs76007856 | 43795238 | G | A | 3.354909220221227e-08 | 5.521827948598748 | 32176.0 |
| 17 | rs111415173 | 43795573 | C | T | 3.354909220221227e-08 | 5.521827948598748 | 32176.0 |
| 17 | rs113991678 | 43795634 | C | T | 3.354909220221227e-08 | 5.521827948598748 | 32176.0 |
| 17 | rs112995313 | 43795768 | T | C | 3.354909220221227e-08 | 5.521827948598748 | 32176.0 |
| 17 | rs1880750 | 43796541 | T | A | 3.354909220221227e-08 | 5.521827948598748 | 32176.0 |
| 17 | rs113155081 | 43752078 | A | AC | 3.3603956552632214e-08 | 5.521540902185014 | 32142.0 |
| 17 | rs189777036 | 43574229 | T | C | 3.3608853181987685e-08 | 5.521515305466088 | 32110.0 |
| 17 | rs62063857 | 44076665 | A | G | 3.362751341341905e-08 | 5.521417793820744 | 32186.0 |
| 17 | rs12150141 | 43775145 | G | A | 3.363521371588008e-08 | 5.521377570112356 | 32180.0 |
| 17 | rs62057117 | 43899657 | C | G | 3.3677946758356666e-08 | 5.521154509726183 | 32165.0 |
| 17 | rs418891 | 43693538 | G | T | 3.369080041555169e-08 | 5.52108746917642 | 32171.0 |
| 17 | rs55973918 | 43792252 | C | A | 3.3701437528500406e-08 | 5.521032008168292 | 32175.0 |
| 17 | rs1568949 | 43792975 | G | A | 3.3701437528500406e-08 | 5.521032008168292 | 32175.0 |
| 17 | rs1105571 | 43793200 | A | T | 3.3701437528500406e-08 | 5.521032008168292 | 32175.0 |
| 17 | rs1105570 | 43793342 | A | G | 3.3701437528500406e-08 | 5.521032008168292 | 32175.0 |
| 17 | rs1105569 | 43793388 | C | T | 3.3701437528500406e-08 | 5.521032008168292 | 32175.0 |
| 17 | rs7502937 | 43793665 | A | G | 3.3701437528500406e-08 | 5.521032008168292 | 32175.0 |
| 17 | rs78917479 | 43899611 | C | T | 3.373699636252597e-08 | 5.5208467305881275 | 32169.0 |
| 17 | rs62057116 | 43899655 | T | C | 3.373699636252597e-08 | 5.5208467305881275 | 32169.0 |
| 17 | rs62057101 | 43885291 | G | A | 3.375853667036053e-08 | 5.52073458796366 | 32174.0 |
| 17 | rs2942164 | 43721283 | G | C | 3.3783137183402664e-08 | 5.520606598240686 | 32186.0 |
| 17 | rs62056866 | 43771493 | G | A | 3.3796499102382603e-08 | 5.520537117722296 | 32183.0 |
| 17 | rs62056867 | 43771627 | A | G | 3.3796499102382603e-08 | 5.520537117722296 | 32183.0 |
| 17 | rs56200760 | 43772018 | G | T | 3.3796499102382603e-08 | 5.520537117722296 | 32183.0 |
| 17 | rs17688391 | 43772109 | C | A | 3.3796499102382603e-08 | 5.520537117722296 | 32183.0 |
| 17 | rs55927630 | 43772407 | C | T | 3.3796499102382603e-08 | 5.520537117722296 | 32183.0 |
| 17 | rs17688434 | 43772540 | G | A | 3.3796499102382603e-08 | 5.520537117722296 | 32183.0 |
| 17 | rs17688452 | 43773085 | G | A | 3.3796499102382603e-08 | 5.520537117722296 | 32183.0 |
| 17 | rs10491144 | 43773124 | A | C | 3.3796499102382603e-08 | 5.520537117722296 | 32183.0 |
| 17 | rs10491143 | 43773248 | A | T | 3.3796499102382603e-08 | 5.520537117722296 | 32183.0 |
| 17 | rs62056869 | 43773447 | T | C | 3.3796499102382603e-08 | 5.520537117722296 | 32183.0 |
| 17 | rs17688511 | 43773607 | T | G | 3.3796499102382603e-08 | 5.520537117722296 | 32183.0 |
| 17 | rs17688534 | 43773877 | A | T | 3.3796499102382603e-08 | 5.520537117722296 | 32183.0 |
| 17 | rs17688558 | 43773943 | A | G | 3.3796499102382603e-08 | 5.520537117722296 | 32183.0 |
| 17 | rs56162163 | 43774497 | A | T | 3.3796499102382603e-08 | 5.520537117722296 | 32183.0 |
| 17 | rs56391096 | 43774845 | C | T | 3.3796499102382603e-08 | 5.520537117722296 | 32183.0 |
| 17 | rs17761838 | 43774959 | T | C | 3.3796499102382603e-08 | 5.520537117722296 | 32183.0 |
| 17 | rs62056872 | 43775015 | T | C | 3.3796499102382603e-08 | 5.520537117722296 | 32183.0 |
| 17 | rs12150621 | 43838482 | G | T | 3.382734503984589e-08 | 5.520376823919091 | 32158.0 |
| 17 | rs81632 | 43710127 | T | A | 3.3871374600418596e-08 | 5.520148265634709 | 32184.0 |
| 17 | 17:43827156_GA_G | 43827156 | GA | G | 3.387583939455478e-08 | 5.520125104899122 | 32170.0 |
| 17 | rs566127718 | 43575452 | A | G | 3.390922612006399e-08 | 5.519952007926791 | 32106.0 |
| 17 | rs200133133 | 43576046 | T | TG | 3.395465554493667e-08 | 5.5197167396412015 | 32104.0 |
| 17 | rs17769490 | 43925605 | G | A | 3.3957717700750335e-08 | 5.5197008924435 | 32178.0 |
| 17 | rs77063768 | 43925729 | A | G | 3.3957717700750335e-08 | 5.5197008924435 | 32178.0 |

| | | | | | | | |
|---|---|---|---|---|---|---|---|
| 17 | rs62054818 | 43926056 | A | C | 3.3957717700750335e-08 | 5.5197008924435 | 32178.0 |
| 17 | rs62064421 | 43667934 | C | G | 3.4014534385989547e-08 | 5.519407107238791 | 31979.0 |
| 17 | rs55942528 | 43932028 | A | G | 3.403530393327792e-08 | 5.519299831771972 | 32135.0 |
| 17 | rs62056946 | 43798401 | G | A | 3.404829120970183e-08 | 5.519232784278897 | 32176.0 |
| 17 | rs78052495 | 43768347 | C | T | 3.405183403679462e-08 | 5.519214498555403 | 32024.0 |
| 17 | rs62057114 | 43899401 | C | T | 3.405277885168598e-08 | 5.519209622359312 | 32168.0 |
| 17 | rs62057115 | 43899417 | G | C | 3.405277885168598e-08 | 5.519209622359312 | 32168.0 |
| 17 | rs149079649 | 44203806 | C | CA | 3.4081253445104184e-08 | 5.5190627263297936 | 32131.0 |
| 17 | rs62061856 | 44200015 | T | C | 3.4094139656052484e-08 | 5.518996287480888 | 32092.0 |
| 17 | rs76493835 | 43807642 | T | G | 3.410703064076069e-08 | 5.518929848385509 | 32178.0 |
| 17 | rs62061807 | 44177097 | C | T | 3.4140166990875507e-08 | 5.51875917801984 | 32147.0 |
| 17 | rs56254732 | 43932049 | C | T | 3.41418246361071e-08 | 5.51875064445886 | 32133.0 |
| 17 | rs113316734 | 44084157 | T | C | 3.4170253677776526e-08 | 5.5186043542097405 | 32173.0 |
| 17 | rs62055718 | 43766418 | C | T | 3.4261501372117504e-08 | 5.518135607820934 | 31989.0 |
| 17 | rs62056877 | 43778828 | T | G | 3.4265541236767136e-08 | 5.5181148827298845 | 32125.0 |
| 17 | rs56317731 | 44032073 | A | G | 3.434584110884289e-08 | 5.517703423751748 | 32149.0 |
| 17 | rs80184151 | 43879308 | A | G | 3.435751477737279e-08 | 5.517643685217931 | 32186.0 |
| 17 | rs111411180 | 44156401 | T | TA | 3.4360016783490094e-08 | 5.517630884077618 | 32130.0 |
| 17 | rs78471340 | 43768346 | G | A | 3.437121842756125e-08 | 5.51757358362311 | 32030.0 |
| 17 | rs1608955 | 43815713 | C | T | 3.4372410302365666e-08 | 5.5175674878192895 | 32179.0 |
| 17 | rs1608956 | 43815820 | T | A | 3.4372410302365666e-08 | 5.5175674878192895 | 32179.0 |
| 17 | rs62054819 | 43926149 | A | G | 3.447350920302588e-08 | 5.517051165704892 | 32166.0 |
| 17 | rs62073097 | 44316389 | G | A | 3.448570367878971e-08 | 5.51698898659477 | 32114.0 |
| 17 | rs62053958 | 43751223 | A | T | 3.452518535700596e-08 | 5.51678781740648 | 32179.0 |
| 17 | rs62055660 | 43751308 | T | C | 3.452518535700596e-08 | 5.51678781740648 | 32179.0 |
| 17 | rs113486004 | 43751427 | T | C | 3.452518535700596e-08 | 5.51678781740648 | 32179.0 |
| 17 | rs62056910 | 43784222 | T | C | 3.4601162349024815e-08 | 5.516401322382399 | 32164.0 |
| 17 | rs62056909 | 43783803 | A | G | 3.4633928426983526e-08 | 5.516234895299768 | 32023.0 |
| 17 | rs3865315 | 44165169 | C | T | 3.4781933242383125e-08 | 5.515485039811344 | 32065.0 |
| 17 | rs56080482 | 44032597 | G | A | 3.480111074447326e-08 | 5.515388104980955 | 32146.0 |
| 17 | rs62056906 | 43782708 | T | C | 3.482041931728536e-08 | 5.5152905599679185 | 32161.0 |
| 17 | rs62056864 | 43770995 | G | A | 3.483840986387038e-08 | 5.515199720696769 | 32184.0 |
| 17 | rs62056865 | 43771193 | T | C | 3.483840986387038e-08 | 5.515199720696769 | 32184.0 |
| 17 | rs2668639 | 44324539 | T | C | 3.488070536548372e-08 | 5.5149863380461515 | 31828.0 |
| 17 | rs28768976 | 43688317 | A | G | 3.490042015133093e-08 | 5.514886961829708 | 32137.0 |
| 17 | rs62055956 | 43854449 | C | T | 3.493164746751046e-08 | 5.514729665832344 | 32126.0 |
| 17 | rs17571718 | 44032768 | T | C | 3.494436409502258e-08 | 5.51466564962393 | 32139.0 |
| 17 | rs17687849 | 43759719 | A | G | 3.4991758298159716e-08 | 5.514427263444762 | 32184.0 |
| 17 | rs77310586 | 43765381 | G | C | 3.4991758298159716e-08 | 5.514427263444762 | 32184.0 |
| 17 | rs17688126 | 43765080 | A | G | 3.5014932832370335e-08 | 5.514310812776519 | 32184.0 |
| 17 | rs62053959 | 43751246 | C | T | 3.5021487521131554e-08 | 5.514277889413644 | 32178.0 |
| 17 | rs62060790 | 44137676 | A | T | 3.503581491383578e-08 | 5.5142059455582055 | 32155.0 |
| 17 | rs62060791 | 44137678 | G | A | 3.503581491383578e-08 | 5.5142059455582055 | 32155.0 |
| 17 | rs113728380 | 43760199 | C | T | 3.5068497983300534e-08 | 5.514041936875175 | 32181.0 |
| 17 | rs739645 | 43894990 | T | G | 3.509865604902733e-08 | 5.513890730513327 | 32181.0 |
| 17 | rs241036 | 43731719 | A | C | 3.516404509446507e-08 | 5.5135633155887005 | 32186.0 |
| 17 | rs140510364 | 44107606 | A | AT | 3.523736878826704e-08 | 5.5131968720476285 | 31987.0 |
| 17 | rs10451283 | 43814020 | T | C | 3.526559127360855e-08 | 5.513056024032913 | 32174.0 |
| 17 | rs7210219 | 44018519 | T | C | 3.530093111026578e-08 | 5.512879810021799 | 30978.0 |
| 17 | rs17688875 | 43778406 | A | G | 3.532124580270651e-08 | 5.5127785928854385 | 32153.0 |
| 17 | rs56319902 | 43871982 | C | T | 3.535100423181964e-08 | 5.512630424600003 | 32186.0 |
| 17 | rs112169475 | 43760192 | A | G | 3.5387776916655774e-08 | 5.512447499099965 | 32182.0 |
| 17 | rs56329743 | 43931245 | G | A | 3.5439690083240964e-08 | 5.51218957097041 | 32153.0 |
| 17 | rs62055717 | 43766415 | A | G | 3.5458731311301466e-08 | 5.512095057368865 | 31991.0 |

| | | | | | | | |
|---|---|---|---|---|---|---|---|
| 17 | rs12150455 | 43827508 | A | G | 3.5469177530096854e-08 | 5.512043227117552 | 32178.0 |
| 17 | rs76728632 | 43818160 | A | C | 3.5517638218624244e-08 | 5.511802976699446 | 32181.0 |
| 17 | rs62054823 | 43929029 | A | G | 3.554743556415087e-08 | 5.511655410124129 | 32126.0 |
| 17 | rs62057119 | 43899736 | G | A | 3.5555073535069737e-08 | 5.511617603615784 | 32164.0 |
| 17 | rs17762954 | 43899786 | C | T | 3.5555073535069737e-08 | 5.511617603615784 | 32164.0 |
| 17 | rs647483 | 43729432 | C | T | 3.557701083873297e-08 | 5.511509061906429 | 32184.0 |
| 17 | rs78555354 | 44032879 | A | G | 3.5588477737216466e-08 | 5.511452351650585 | 32135.0 |
| 17 | rs75500942 | 43816106 | C | A | 3.561895041544179e-08 | 5.511301733217561 | 32180.0 |
| 17 | rs62054418 | 43816306 | G | A | 3.561895041544179e-08 | 5.511301733217561 | 32180.0 |
| 17 | rs17762535 | 43787130 | T | C | 3.565624324437751e-08 | 5.5111175745458745 | 32174.0 |
| 17 | rs62056922 | 43787563 | C | G | 3.565624324437751e-08 | 5.5111175745458745 | 32174.0 |
| 17 | rs56397077 | 43788419 | C | A | 3.565624324437751e-08 | 5.5111175745458745 | 32174.0 |
| 17 | rs62056925 | 43788649 | T | C | 3.565624324437751e-08 | 5.5111175745458745 | 32174.0 |
| 17 | rs62056926 | 43788666 | C | G | 3.565624324437751e-08 | 5.5111175745458745 | 32174.0 |
| 17 | rs62056935 | 43790446 | A | G | 3.565624324437751e-08 | 5.5111175745458745 | 32174.0 |
| 17 | rs75076131 | 43790649 | C | A | 3.565624324437751e-08 | 5.5111175745458745 | 32174.0 |
| 17 | rs62056936 | 43790911 | A | T | 3.565624324437751e-08 | 5.5111175745458745 | 32174.0 |
| 17 | rs62056937 | 43791181 | A | C | 3.565624324437751e-08 | 5.5111175745458745 | 32174.0 |
| 17 | rs62056939 | 43791332 | T | C | 3.565624324437751e-08 | 5.5111175745458745 | 32174.0 |
| 17 | rs62056940 | 43791344 | T | A | 3.565624324437751e-08 | 5.5111175745458745 | 32174.0 |
| 17 | rs78803763 | 43791405 | C | A | 3.565624324437751e-08 | 5.5111175745458745 | 32174.0 |
| 17 | rs77387379 | 43791429 | G | A | 3.565624324437751e-08 | 5.5111175745458745 | 32174.0 |
| 17 | rs75990322 | 43791468 | T | C | 3.565624324437751e-08 | 5.5111175745458745 | 32174.0 |
| 17 | rs113661667 | 43791610 | T | C | 3.565624324437751e-08 | 5.5111175745458745 | 32174.0 |
| 17 | rs74579538 | 43791652 | G | A | 3.565624324437751e-08 | 5.5111175745458745 | 32174.0 |
| 17 | rs650927 | 43709867 | T | G | 3.5678612575392644e-08 | 5.511007200394684 | 32185.0 |
| 17 | rs671740 | 43709893 | A | G | 3.5678612575392644e-08 | 5.511007200394684 | 32185.0 |
| 17 | rs16940633 | 43710371 | G | A | 3.5678612575392644e-08 | 5.511007200394684 | 32185.0 |
| 17 | rs17686610 | 43710384 | A | C | 3.5678612575392644e-08 | 5.511007200394684 | 32185.0 |
| 17 | rs2942170 | 43711312 | C | T | 3.5678612575392644e-08 | 5.511007200394684 | 32185.0 |
| 17 | rs2040844 | 43767219 | A | C | 3.5684547079835254e-08 | 5.510977929787433 | 32185.0 |
| 17 | rs62055884 | 43837567 | C | T | 3.5700871953977704e-08 | 5.5108974353708335 | 32163.0 |
| 17 | 17:44254993_GT_G | 44254993 | GT | G | 3.5773308938814616e-08 | 5.5105406943972515 | 31691.0 |
| 17 | rs62056920 | 43787226 | G | A | 3.5780498566236814e-08 | 5.510505324819525 | 32173.0 |
| 17 | rs62056921 | 43787306 | T | C | 3.5780498566236814e-08 | 5.510505324819525 | 32173.0 |
| 17 | rs12150658 | 43826305 | G | A | 3.5793269842244536e-08 | 5.510442513155876 | 32179.0 |
| 17 | rs17688032 | 43763202 | G | A | 3.581138046817069e-08 | 5.510353478769785 | 32186.0 |
| 17 | 17:43819597_AC_A | 43819597 | AC | A | 3.5916873027400046e-08 | 5.5098357289458235 | 32172.0 |
| 17 | rs62053956 | 43750315 | A | G | 3.5921228482972986e-08 | 5.509814384403341 | 32182.0 |
| 17 | rs55821155 | 44032470 | C | G | 3.5928447234313165e-08 | 5.509779013391217 | 32147.0 |
| 17 | rs138325961 | 43971238 | T | TTAG ATAC ATCA | 3.593404896875489e-08 | 5.509751570316446 | 32100.0 |
| 17 | rs1724413 | 43670073 | G | T | 3.60517583535248e-08 | 5.509175865898538 | 31694.0 |
| 17 | rs113072604 | 43741627 | A | AT | 3.611150518105404e-08 | 5.508884347493561 | 32172.0 |
| 17 | rs62056927 | 43788767 | G | A | 3.6140413404223836e-08 | 5.508743465559549 | 32173.0 |
| 17 | rs62056929 | 43789144 | G | A | 3.6140413404223836e-08 | 5.508743465559549 | 32173.0 |
| 17 | 17:43809559_GT_G | 43809559 | GT | G | 3.6186262468029605e-08 | 5.508520247758817 | 32058.0 |
| 17 | rs1526127 | 43812398 | C | G | 3.6200680356514286e-08 | 5.508450110443748 | 32182.0 |
| 17 | rs62054398 | 43812599 | A | G | 3.6200680356514286e-08 | 5.508450110443748 | 32182.0 |
| 17 | rs62054399 | 43812624 | G | A | 3.6200680356514286e-08 | 5.508450110443748 | 32182.0 |
| 17 | rs62054400 | 43812649 | T | C | 3.6200680356514286e-08 | 5.508450110443748 | 32182.0 |
| 17 | rs62054411 | 43812684 | A | G | 3.6200680356514286e-08 | 5.508450110443748 | 32182.0 |
| 17 | rs62054412 | 43813002 | G | A | 3.6200680356514286e-08 | 5.508450110443748 | 32182.0 |
| 17 | rs17563787 | 43813252 | C | G | 3.6200680356514286e-08 | 5.508450110443748 | 32182.0 |
| 17 | rs62054413 | 43813370 | G | C | 3.6200680356514286e-08 | 5.508450110443748 | 32182.0 |

| | | | | | | | |
|---|---|---|---|---|---|---|---|
| 17 | rs62054414 | 43813415 | G | A | 3.6200680356514286e-08 | 5.508450110443748 | 32182.0 |
| 17 | rs76421814 | 43814427 | T | C | 3.6200680356514286e-08 | 5.508450110443748 | 32182.0 |
| 17 | rs56006248 | 43815141 | A | G | 3.6200680356514286e-08 | 5.508450110443748 | 32182.0 |
| 17 | rs56298110 | 43815336 | A | G | 3.6200680356514286e-08 | 5.508450110443748 | 32182.0 |
| 17 | rs62054417 | 43815859 | A | C | 3.6200680356514286e-08 | 5.508450110443748 | 32182.0 |
| 17 | rs62054419 | 43816605 | A | C | 3.6200680356514286e-08 | 5.508450110443748 | 32182.0 |
| 17 | rs62054420 | 43816655 | C | G | 3.6200680356514286e-08 | 5.508450110443748 | 32182.0 |
| 17 | rs62054421 | 43816657 | G | A | 3.6200680356514286e-08 | 5.508450110443748 | 32182.0 |
| 17 | rs17563800 | 43817459 | C | T | 3.6200680356514286e-08 | 5.508450110443748 | 32182.0 |
| 17 | rs56295996 | 43817557 | C | T | 3.6200680356514286e-08 | 5.508450110443748 | 32182.0 |
| 17 | rs56384960 | 43817608 | G | A | 3.6200680356514286e-08 | 5.508450110443748 | 32182.0 |
| 17 | rs56178230 | 43817832 | G | A | 3.6200680356514286e-08 | 5.508450110443748 | 32182.0 |
| 17 | rs80157557 | 43817898 | G | A | 3.6200680356514286e-08 | 5.508450110443748 | 32182.0 |
| 17 | rs17563827 | 43818222 | C | A | 3.6200680356514286e-08 | 5.508450110443748 | 32182.0 |
| 17 | 17:43926093_GATGA_G | 43926093 | GATGA | G | 3.6203062984104864e-08 | 5.508438522513081 | 31484.0 |
| 17 | rs885639 | 43928614 | T | G | 3.6209584645258666e-08 | 5.5084068081381785 | 32159.0 |
| 17 | rs56167344 | 43797087 | A | T | 3.621686016360773e-08 | 5.508371434346075 | 32142.0 |
| 17 | rs17688056 | 43763241 | C | T | 3.625752906811529e-08 | 5.508173827739405 | 32183.0 |
| 17 | rs34739559 | 43949342 | G | A | 3.629409390450489e-08 | 5.507996345873084 | 31935.0 |
| 17 | rs111920741 | 44333412 | G | A | 3.6301008842286265e-08 | 5.507962800992825 | 31907.0 |
| 17 | rs62056863 | 43770595 | A | G | 3.632163559271736e-08 | 5.507862775885974 | 32185.0 |
| 17 | rs77301847 | 43810902 | A | G | 3.6341770317166585e-08 | 5.507765189877524 | 31746.0 |
| 17 | rs113520245 | 44033132 | C | T | 3.6386105209390584e-08 | 5.507550498787291 | 31688.0 |
| 17 | rs1706722 | 43684094 | T | C | 3.643541432590775e-08 | 5.5073120178830655 | 31744.0 |
| 17 | rs56108117 | 43757776 | T | C | 3.643806449323019e-08 | 5.5072992093559 | 32182.0 |
| 17 | rs56378021 | 43757777 | C | T | 3.643806449323019e-08 | 5.5072992093559 | 32182.0 |
| 17 | 17:43913360_TGAGGGCAGGAGG_T | 43913360 | TGAGGGCAGGAGG | T | 3.652474054644246e-08 | 5.506880792431387 | 32128.0 |
| 17 | rs62055693 | 43756910 | T | C | 3.6525752536457107e-08 | 5.506875912876061 | 32185.0 |
| 17 | 17:43769819_AAAATT_A | 43769819 | AAAATT | A | 3.659121067296127e-08 | 5.506560568794176 | 32129.0 |
| 17 | rs757502 | 43756506 | C | T | 3.660958996470194e-08 | 5.506472125066298 | 32170.0 |
| 17 | rs17652036 | 44073027 | G | A | 3.6609843535396464e-08 | 5.506470905149759 | 32140.0 |
| 17 | rs62054428 | 43822511 | G | A | 3.6682562446931016e-08 | 5.506121395638491 | 32176.0 |
| 17 | rs76171147 | 43823229 | T | C | 3.670835892459555e-08 | 5.505997571260946 | 32186.0 |
| 17 | rs383241 | 43705356 | G | A | 3.671191845388397e-08 | 5.505980491969278 | 32186.0 |
| 17 | rs376120 | 43705518 | A | G | 3.671191845388397e-08 | 5.505980491969278 | 32186.0 |
| 17 | rs594591 | 43707620 | G | T | 3.671191845388397e-08 | 5.505980491969278 | 32186.0 |
| 17 | rs2942169 | 43711539 | G | C | 3.671191845388397e-08 | 5.505980491969278 | 32186.0 |
| 17 | rs241045 | 43712337 | C | T | 3.671191845388397e-08 | 5.505980491969278 | 32186.0 |
| 17 | rs241044 | 43712464 | G | A | 3.671191845388397e-08 | 5.505980491969278 | 32186.0 |
| 17 | rs241042 | 43712657 | T | C | 3.671191845388397e-08 | 5.505980491969278 | 32186.0 |
| 17 | rs241041 | 43713925 | A | G | 3.671191845388397e-08 | 5.505980491969278 | 32186.0 |
| 17 | rs241039 | 43714673 | A | T | 3.671191845388397e-08 | 5.505980491969278 | 32186.0 |
| 17 | rs2942168 | 43714850 | G | A | 3.671191845388397e-08 | 5.505980491969278 | 32186.0 |
| 17 | rs2942167 | 43715018 | T | C | 3.671191845388397e-08 | 5.505980491969278 | 32186.0 |
| 17 | rs2942165 | 43715619 | G | C | 3.671191845388397e-08 | 5.505980491969278 | 32186.0 |
| 17 | rs117368197 | 43715924 | A | G | 3.671191845388397e-08 | 5.505980491969278 | 32186.0 |
| 17 | rs413778 | 43716885 | A | G | 3.671191845388397e-08 | 5.505980491969278 | 32186.0 |
| 17 | rs389217 | 43717131 | C | T | 3.671191845388397e-08 | 5.505980491969278 | 32186.0 |
| 17 | rs3108317 | 43719870 | G | A | 3.671191845388397e-08 | 5.505980491969278 | 32186.0 |
| 17 | rs1706746 | 43721457 | C | T | 3.671191845388397e-08 | 5.505980491969278 | 32186.0 |
| 17 | rs3418 | 43723462 | C | T | 3.671191845388397e-08 | 5.505980491969278 | 32186.0 |
| 17 | rs393675 | 43723605 | T | C | 3.671191845388397e-08 | 5.505980491969278 | 32186.0 |
| 17 | rs449501 | 43724555 | G | A | 3.671191845388397e-08 | 5.505980491969278 | 32186.0 |
| 17 | rs448830 | 43725212 | G | A | 3.671191845388397e-08 | 5.505980491969278 | 32186.0 |

| | | | | | | | |
|---|---|---|---|---|---|---|---|
| 17 | rs434428 | 43725684 | G | A | 3.671191845388397e-08 | 5.505980491969278 | 32186.0 |
| 17 | rs434598 | 43725762 | G | A | 3.671191845388397e-08 | 5.505980491969278 | 32186.0 |
| 17 | rs434971 | 43725992 | G | A | 3.671191845388397e-08 | 5.505980491969278 | 32186.0 |
| 17 | rs385691 | 43726125 | A | C | 3.671191845388397e-08 | 5.505980491969278 | 32186.0 |
| 17 | rs455028 | 43726574 | T | C | 3.671191845388397e-08 | 5.505980491969278 | 32186.0 |
| 17 | rs241035 | 43731896 | C | T | 3.671191845388397e-08 | 5.505980491969278 | 32186.0 |
| 17 | rs62055700 | 43758742 | T | C | 3.6743714956888776e-08 | 5.50582799757174 | 32184.0 |
| 17 | rs62053957 | 43750939 | T | C | 3.67951551414605e-08 | 5.505581563880859 | 32183.0 |
| 17 | rs62055751 | 43767538 | A | G | 3.685049374559409e-08 | 5.5053168268806525 | 32185.0 |
| 17 | rs75691066 | 44045585 | C | A | 3.6852663045387044e-08 | 5.505306456918514 | 32142.0 |
| 17 | rs424243 | 43727887 | A | T | 3.6873469153714466e-08 | 5.505207026976764 | 32184.0 |
| 17 | rs62053953 | 43749454 | G | A | 3.690910877234945e-08 | 5.505036835795454 | 32186.0 |
| 17 | rs62056876 | 43778752 | C | T | 3.691958991414744e-08 | 5.504986815140759 | 32152.0 |
| 17 | rs34579278 | 43833981 | A | G | 3.696359136511305e-08 | 5.504776970871898 | 32172.0 |
| 17 | rs453997 | 43727061 | C | T | 3.6976264090753444e-08 | 5.5047165791877655 | 32185.0 |
| 17 | rs422112 | 43728137 | G | A | 3.6976264090753444e-08 | 5.5047165791877655 | 32185.0 |
| 17 | rs413844 | 43729384 | G | A | 3.6976264090753444e-08 | 5.5047165791877655 | 32185.0 |
| 17 | rs241038 | 43730598 | G | A | 3.6976264090753444e-08 | 5.5047165791877655 | 32185.0 |
| 17 | rs241037 | 43730636 | G | T | 3.6976264090753444e-08 | 5.5047165791877655 | 32185.0 |
| 17 | rs241032 | 43734145 | T | C | 3.6994064293298334e-08 | 5.504631786479503 | 32178.0 |
| 17 | rs674692 | 43695967 | A | G | 3.701494860621404e-08 | 5.504532352792246 | 32143.0 |
| 17 | rs675726 | 43696254 | A | G | 3.701494860621404e-08 | 5.504532352792246 | 32143.0 |
| 17 | rs143345034 | 43696920 | C | T | 3.701494860621404e-08 | 5.504532352792246 | 32143.0 |
| 17 | 17:44130328_TC_T | 44130328 | TC | T | 3.704251306492258e-08 | 5.504401197084408 | 32098.0 |
| 17 | rs2732625 | 44332429 | C | T | 3.7063167523765334e-08 | 5.504302982181274 | 32126.0 |
| 17 | rs413519 | 43685826 | A | G | 3.7130983089119044e-08 | 5.5039808823220415 | 32070.0 |
| 17 | rs143143028 | 44170129 | C | CAA | 3.7255885438319916e-08 | 5.5033891307420095 | 31881.0 |
| 17 | rs17689378 | 43881790 | C | T | 3.732559695772039e-08 | 5.503059693555584 | 32186.0 |
| 17 | rs62055932 | 43848412 | A | G | 3.7389870292894336e-08 | 5.502756483975331 | 32143.0 |
| 17 | rs77555455 | 44035516 | G | A | 3.739103519525221e-08 | 5.502750993211269 | 32128.0 |
| 17 | rs2950009 | 43666244 | A | G | 3.742133513926251e-08 | 5.502608232754802 | 32030.0 |
| 17 | rs2950010 | 43666251 | C | G | 3.743649413013168e-08 | 5.502536852099861 | 32119.0 |
| 17 | rs12150610 | 43775479 | T | C | 3.744232611291778e-08 | 5.502509397926059 | 32178.0 |
| 17 | rs2873269 | 43931122 | C | T | 3.7539786899742076e-08 | 5.502051212058276 | 32157.0 |
| 17 | rs757500 | 43756833 | C | A | 3.7586593861779556e-08 | 5.501831571534178 | 32186.0 |
| 17 | rs55651627 | 43758011 | G | C | 3.7586593861779556e-08 | 5.501831571534178 | 32186.0 |
| 17 | rs112155389 | 43758078 | G | A | 3.7586593861779556e-08 | 5.501831571534178 | 32186.0 |
| 17 | rs17687838 | 43759048 | T | C | 3.7586593861779556e-08 | 5.501831571534178 | 32186.0 |
| 17 | rs76333921 | 43759426 | C | T | 3.7586593861779556e-08 | 5.501831571534178 | 32186.0 |
| 17 | rs76453925 | 43759431 | C | T | 3.7586593861779556e-08 | 5.501831571534178 | 32186.0 |
| 17 | rs17761100 | 43760389 | A | G | 3.7586593861779556e-08 | 5.501831571534178 | 32186.0 |
| 17 | rs17761124 | 43760516 | A | T | 3.7586593861779556e-08 | 5.501831571534178 | 32186.0 |
| 17 | rs62055704 | 43760577 | G | A | 3.7586593861779556e-08 | 5.501831571534178 | 32186.0 |
| 17 | rs62055705 | 43760599 | A | C | 3.7586593861779556e-08 | 5.501831571534178 | 32186.0 |
| 17 | rs62055706 | 43760685 | T | C | 3.7586593861779556e-08 | 5.501831571534178 | 32186.0 |
| 17 | rs75115093 | 43761381 | A | G | 3.7586593861779556e-08 | 5.501831571534178 | 32186.0 |
| 17 | rs62055707 | 43761482 | T | C | 3.7586593861779556e-08 | 5.501831571534178 | 32186.0 |
| 17 | rs2049515 | 43761856 | C | T | 3.7586593861779556e-08 | 5.501831571534178 | 32186.0 |
| 17 | rs62055709 | 43762429 | C | T | 3.7586593861779556e-08 | 5.501831571534178 | 32186.0 |
| 17 | rs62055711 | 43762554 | A | G | 3.7586593861779556e-08 | 5.501831571534178 | 32186.0 |
| 17 | rs17688002 | 43762594 | A | T | 3.7586593861779556e-08 | 5.501831571534178 | 32186.0 |
| 17 | rs2040846 | 43762846 | A | G | 3.7586593861779556e-08 | 5.501831571534178 | 32186.0 |
| 17 | rs4486953 | 43763007 | T | C | 3.7586593861779556e-08 | 5.501831571534178 | 32186.0 |
| 17 | rs17688068 | 43763935 | A | G | 3.7586593861779556e-08 | 5.501831571534178 | 32186.0 |

| | | | | | | | |
|---|---|---|---|---|---|---|---|
| 17 | rs62055712 | 43764565 | A | G | 3.7586593861779556e-08 | 5.501831571534178 | 32186.0 |
| 17 | rs62055713 | 43764685 | C | T | 3.7586593861779556e-08 | 5.501831571534178 | 32186.0 |
| 17 | rs17688090 | 43764987 | G | A | 3.7586593861779556e-08 | 5.501831571534178 | 32186.0 |
| 17 | rs17761387 | 43765450 | G | A | 3.7586593861779556e-08 | 5.501831571534178 | 32186.0 |
| 17 | rs62055716 | 43765778 | C | T | 3.7586593861779556e-08 | 5.501831571534178 | 32186.0 |
| 17 | rs17688205 | 43766352 | G | C | 3.7586593861779556e-08 | 5.501831571534178 | 32186.0 |
| 17 | rs17688249 | 43766754 | C | T | 3.7586593861779556e-08 | 5.501831571534178 | 32186.0 |
| 17 | rs62055753 | 43767631 | G | A | 3.7586593861779556e-08 | 5.501831571534178 | 32186.0 |
| 17 | rs17688296 | 43767815 | C | T | 3.7586593861779556e-08 | 5.501831571534178 | 32186.0 |
| 17 | rs62055754 | 43768230 | C | G | 3.7586593861779556e-08 | 5.501831571534178 | 32186.0 |
| 17 | rs55972157 | 43768703 | C | T | 3.7586593861779556e-08 | 5.501831571534178 | 32186.0 |
| 17 | rs17761581 | 43769088 | T | C | 3.7586593861779556e-08 | 5.501831571534178 | 32186.0 |
| 17 | rs56283089 | 43769321 | A | C | 3.7586593861779556e-08 | 5.501831571534178 | 32186.0 |
| 17 | rs56312737 | 43769516 | C | A | 3.7586593861779556e-08 | 5.501831571534178 | 32186.0 |
| 17 | rs74765707 | 43770416 | T | C | 3.7586593861779556e-08 | 5.501831571534178 | 32186.0 |
| 17 | rs62056861 | 43770443 | T | C | 3.7586593861779556e-08 | 5.501831571534178 | 32186.0 |
| 17 | rs62056862 | 43770525 | C | T | 3.7586593861779556e-08 | 5.501831571534178 | 32186.0 |
| 17 | rs56290582 | 43788371 | C | T | 3.7591668047084514e-08 | 5.501807776982367 | 32172.0 |
| 17 | rs241034 | 43733048 | A | T | 3.765091648253655e-08 | 5.501530171542176 | 32185.0 |
| 17 | rs241033 | 43733983 | G | A | 3.765091648253655e-08 | 5.501530171542176 | 32185.0 |
| 17 | rs241031 | 43734304 | T | C | 3.765091648253655e-08 | 5.501530171542176 | 32185.0 |
| 17 | rs241030 | 43734503 | A | G | 3.765091648253655e-08 | 5.501530171542176 | 32185.0 |
| 17 | rs241029 | 43734643 | T | C | 3.765091648253655e-08 | 5.501530171542176 | 32185.0 |
| 17 | rs241028 | 43735276 | C | T | 3.765091648253655e-08 | 5.501530171542176 | 32185.0 |
| 17 | rs241027 | 43735478 | A | G | 3.765091648253655e-08 | 5.501530171542176 | 32185.0 |
| 17 | rs241026 | 43735555 | G | A | 3.765091648253655e-08 | 5.501530171542176 | 32185.0 |
| 17 | rs241024 | 43736763 | C | T | 3.765091648253655e-08 | 5.501530171542176 | 32185.0 |
| 17 | rs241023 | 43737040 | G | A | 3.765091648253655e-08 | 5.501530171542176 | 32185.0 |
| 17 | rs241021 | 43738627 | T | G | 3.765091648253655e-08 | 5.501530171542176 | 32185.0 |
| 17 | rs241020 | 43738676 | C | A | 3.765091648253655e-08 | 5.501530171542176 | 32185.0 |
| 17 | 17:43746187_TC_T | 43746187 | TC | T | 3.7681553138283726e-08 | 5.501386791223204 | 32180.0 |
| 17 | rs55980621 | 43768496 | C | T | 3.769094457454167e-08 | 5.501342861704282 | 32184.0 |
| 17 | rs55669046 | 43768501 | C | T | 3.769094457454167e-08 | 5.501342861704282 | 32184.0 |
| 17 | rs1706748 | 43722509 | C | G | 3.772461605432588e-08 | 5.501185446710036 | 32182.0 |
| 17 | rs113871181 | 43852733 | G | A | 3.777544001042181e-08 | 5.500948101331852 | 32105.0 |
| 17 | 17:43734255_TA_T | 43734255 | TA | T | 3.7782762090692286e-08 | 5.500913933100693 | 32184.0 |
| 17 | rs62055697 | 43758148 | C | T | 3.779335541677192e-08 | 5.500864511079586 | 32185.0 |
| 17 | rs17761046 | 43758382 | T | C | 3.779335541677192e-08 | 5.500864511079586 | 32185.0 |
| 17 | rs56168907 | 43837091 | T | C | 3.780159668190928e-08 | 5.500826071635549 | 32162.0 |
| 17 | rs62056912 | 43784948 | G | T | 3.783287717461231e-08 | 5.500680245058068 | 32167.0 |
| 17 | rs62054822 | 43927708 | A | G | 3.790522027956596e-08 | 5.500343436040021 | 32170.0 |
| 17 | rs4609899 | 43807113 | C | T | 3.7935271938506986e-08 | 5.500203707251651 | 32165.0 |
| 17 | rs4471726 | 43807134 | T | C | 3.794380613714663e-08 | 5.5001640460432375 | 32166.0 |
| 17 | 17:43893146_CCCAGTTCAGTGTGT_C | 43893146 | CCCAGTTCAGTGTGT | C | 3.796324486816303e-08 | 5.500073740194902 | 32117.0 |
| 17 | rs3108318 | 43719829 | G | T | 3.800872753996719e-08 | 5.499862617989188 | 32095.0 |
| 17 | rs62064424 | 43668239 | T | C | 3.81007759663926e-08 | 5.499436094727134 | 31951.0 |
| 17 | rs62055719 | 43766497 | T | C | 3.8127285228853264e-08 | 5.499313444526871 | 32181.0 |
| 17 | rs77220413 | 43764378 | G | A | 3.814246034497955e-08 | 5.4992432711489965 | 32186.0 |
| 17 | rs62073155 | 44333121 | G | A | 3.81560569934822e-08 | 5.4991804199771614 | 32146.0 |
| 17 | rs62054817 | 43925966 | G | A | 3.8157905448128256e-08 | 5.499171877082409 | 32174.0 |
| 17 | rs56110538 | 43768727 | G | A | 3.816741317204504e-08 | 5.499127942130751 | 32184.0 |
| 17 | rs62056928 | 43788782 | C | T | 3.8168205587118066e-08 | 5.499124280879916 | 32171.0 |
| 17 | rs113448888 | 44108482 | A | AGCCCTCT | 3.8181150648238686e-08 | 5.499064480343653 | 32114.0 |
| 17 | rs112245406 | 43756171 | A | AAAAC | 3.8223846968633615e-08 | 5.498867381244335 | 32116.0 |

| | | | | | | | |
|---|---|---|---|---|---|---|---|
| 17 | rs112695020 | 44037106 | C | CA | 3.823522291139548e-08 | 5.498814902480753 | 32073.0 |
| 17 | rs17425752 | 43906726 | A | C | 3.826129425875895e-08 | 5.498694688919536 | 32145.0 |
| 17 | rs56220242 | 43792326 | C | T | 3.832939999970981e-08 | 5.498381031463467 | 32167.0 |
| 17 | rs56160448 | 43792358 | A | G | 3.832939999970981e-08 | 5.498381031463467 | 32167.0 |
| 17 | rs1880749 | 43792586 | A | C | 3.832939999970981e-08 | 5.498381031463467 | 32167.0 |
| 17 | rs370739771 | 44255146 | C | CT | 3.834677667024303e-08 | 5.498301090649825 | 30569.0 |
| 17 | rs190796367 | 44333293 | A | G | 3.835062445724713e-08 | 5.498283393780254 | 32148.0 |
| 17 | rs55747440 | 43837115 | A | G | 3.8399218703425435e-08 | 5.4980600455786215 | 32163.0 |
| 17 | rs241022 | 43737730 | T | C | 3.842367137684136e-08 | 5.497947760184788 | 32184.0 |
| 17 | rs111273167 | 43740967 | C | T | 3.844840534351232e-08 | 5.497834253582108 | 32186.0 |
| 17 | rs62053940 | 43741028 | A | G | 3.844840534351232e-08 | 5.497834253582108 | 32186.0 |
| 17 | rs17760577 | 43741452 | C | T | 3.844840534351232e-08 | 5.497834253582108 | 32186.0 |
| 17 | rs17760631 | 43743045 | T | C | 3.844840534351232e-08 | 5.497834253582108 | 32186.0 |
| 17 | rs4528616 | 43683597 | T | G | 3.846849653923763e-08 | 5.497742105219738 | 32092.0 |
| 17 | rs62056915 | 43785627 | A | T | 3.8553098626559806e-08 | 5.4973545887057735 | 32166.0 |
| 17 | rs56100013 | 43768467 | C | T | 3.856256939395242e-08 | 5.4973112595665565 | 32184.0 |
| 17 | rs2532372 | 43671235 | C | T | 3.8580450010123496e-08 | 5.497229483159062 | 31987.0 |
| 17 | rs62053954 | 43749949 | G | A | 3.8580583478084046e-08 | 5.497228872885958 | 32184.0 |
| 17 | rs62053955 | 43749980 | T | C | 3.8580583478084046e-08 | 5.497228872885958 | 32184.0 |
| 17 | rs17687625 | 43750238 | A | G | 3.8580583478084046e-08 | 5.497228872885958 | 32184.0 |
| 17 | rs1724412 | 43670018 | G | A | 3.87528662132884e-08 | 5.496442823870502 | 31783.0 |
| 17 | rs444071 | 43707951 | C | T | 3.8766005491175384e-08 | 5.4963830143810215 | 32185.0 |
| 17 | rs396593 | 43707955 | C | A | 3.8766005491175384e-08 | 5.4963830143810215 | 32185.0 |
| 17 | rs17762912 | 43899161 | A | C | 3.8766944177995385e-08 | 5.496378742266991 | 32172.0 |
| 17 | rs2532384 | 44317210 | C | A | 3.879551772597827e-08 | 5.496248747452748 | 32144.0 |
| 17 | rs366858 | 43726588 | C | T | 3.879954382647845e-08 | 5.496230438248186 | 32185.0 |
| 17 | rs62057111 | 43897130 | A | G | 3.88376778606795e-08 | 5.496057110184426 | 32179.0 |
| 17 | rs370670279 | 43669049 | A | AC | 3.883875259033097e-08 | 5.496052227679467 | 32124.0 |
| 17 | rs184745911 | 43698899 | A | G | 3.890678846924988e-08 | 5.4957434065365325 | 32098.0 |
| 17 | rs78487840 | 43757161 | C | T | 3.9030924295999643e-08 | 5.49518128956491 | 32186.0 |
| 17 | rs62062796 | 44027316 | G | A | 3.9054690350353065e-08 | 5.495073868895997 | 32155.0 |
| 17 | rs2693349 | 43666511 | A | T | 3.9099830822971936e-08 | 5.494870011991601 | 32027.0 |
| 17 | rs75788861 | 43822769 | C | T | 3.913108062409524e-08 | 5.494729019781338 | 31566.0 |
| 17 | rs2158474 | 43761939 | G | C | 3.926148379578634e-08 | 5.494141845481507 | 32184.0 |
| 17 | rs62055708 | 43761974 | C | A | 3.926148379578634e-08 | 5.494141845481507 | 32184.0 |
| 17 | rs76927677 | 43761985 | A | T | 3.926148379578634e-08 | 5.494141845481507 | 32184.0 |
| 17 | rs17761207 | 43762255 | T | C | 3.926148379578634e-08 | 5.494141845481507 | 32184.0 |
| 17 | 17:44288114_TA_T | 44288114 | TA | T | 3.9287696984024674e-08 | 5.494024042082318 | 31785.0 |
| 17 | rs55711776 | 43669250 | A | G | 3.933364532677643e-08 | 5.493817731672099 | 31998.0 |
| 17 | rs56400821 | 43760080 | C | A | 3.935160393948267e-08 | 5.493737160097438 | 32178.0 |
| 17 | rs56357543 | 43894547 | C | T | 3.936058625753368e-08 | 5.493696874174197 | 32172.0 |
| 17 | rs56099546 | 43894609 | A | G | 3.936058625753368e-08 | 5.493696874174197 | 32172.0 |
| 17 | 17:43742454_AC_A | 43742454 | AC | A | 3.937392734410112e-08 | 5.493637055514959 | 32183.0 |
| 17 | rs62053951 | 43749168 | G | C | 3.951837561155437e-08 | 5.492990635015013 | 32185.0 |
| 17 | rs62053952 | 43749261 | T | C | 3.951837561155437e-08 | 5.492990635015013 | 32185.0 |
| 17 | rs17687534 | 43749579 | T | C | 3.951837561155437e-08 | 5.492990635015013 | 32185.0 |
| 17 | rs17687571 | 43750010 | G | A | 3.951837561155437e-08 | 5.492990635015013 | 32185.0 |
| 17 | rs17760817 | 43750137 | G | C | 3.951837561155437e-08 | 5.492990635015013 | 32185.0 |
| 17 | rs62055661 | 43752039 | C | T | 3.951837561155437e-08 | 5.492990635015013 | 32185.0 |
| 17 | rs62055662 | 43752566 | C | T | 3.951837561155437e-08 | 5.492990635015013 | 32185.0 |
| 17 | rs62055663 | 43753454 | A | G | 3.951837561155437e-08 | 5.492990635015013 | 32185.0 |
| 17 | rs62055664 | 43753642 | A | G | 3.951837561155437e-08 | 5.492990635015013 | 32185.0 |
| 17 | rs76717812 | 43754010 | T | C | 3.951837561155437e-08 | 5.492990635015013 | 32185.0 |
| 17 | rs17687667 | 43754099 | G | A | 3.951837561155437e-08 | 5.492990635015013 | 32185.0 |

| | | | | | | | |
|---|---|---|---|---|---|---|---|
| 17 | rs79073784 | 43754209 | G | C | 3.951837561155437e-08 | 5.492990635015013 | 32185.0 |
| 17 | rs62055714 | 43764796 | G | T | 3.956773477604288e-08 | 5.492770272931184 | 32185.0 |
| 17 | rs62055715 | 43764847 | G | A | 3.956773477604288e-08 | 5.492770272931184 | 32185.0 |
| 17 | rs35879086 | 43830382 | G | A | 3.9633732303241736e-08 | 5.49247604564554 | 32167.0 |
| 17 | rs111957446 | 44179723 | T | C | 3.97476323012416e-08 | 5.491969377374328 | 32123.0 |
| 17 | rs17760733 | 43746276 | G | T | 3.9996438449544734e-08 | 5.490867477019694 | 32183.0 |
| 17 | rs79675109 | 43746418 | T | A | 3.9996438449544734e-08 | 5.490867477019694 | 32183.0 |
| 17 | rs17687504 | 43746721 | A | G | 3.9996438449544734e-08 | 5.490867477019694 | 32183.0 |
| 17 | rs62053945 | 43746839 | T | C | 3.9996438449544734e-08 | 5.490867477019694 | 32183.0 |
| 17 | rs62053946 | 43747677 | C | T | 3.9996438449544734e-08 | 5.490867477019694 | 32183.0 |
| 17 | rs62053947 | 43747683 | G | A | 3.9996438449544734e-08 | 5.490867477019694 | 32183.0 |
| 17 | rs62053949 | 43748101 | A | G | 3.9996438449544734e-08 | 5.490867477019694 | 32183.0 |
| 17 | rs62053950 | 43748412 | A | G | 3.9996438449544734e-08 | 5.490867477019694 | 32183.0 |
| 17 | 17:44029510_TGATG_T | 44029510 | TGATG | T | 4.002548430873058e-08 | 5.4907392736844205 | 32081.0 |
| 17 | rs17571781 | 44033802 | T | C | 4.003849248186777e-08 | 5.490681887132295 | 32125.0 |
| 17 | 17:43950444_CA_C | 43950444 | CA | C | 4.0118431297076336e-08 | 5.490329627353512 | 31808.0 |
| 17 | rs56323832 | 43768633 | G | A | 4.013396612499538e-08 | 5.490261250302444 | 32182.0 |
| 17 | rs74514268 | 43836191 | G | A | 4.0208670951907525e-08 | 5.4899327926152415 | 32170.0 |
| 17 | rs62054762 | 43909022 | G | T | 4.0247465547154884e-08 | 5.489762456238568 | 32003.0 |
| 17 | rs8072451 | 43893716 | C | T | 4.042731559810085e-08 | 5.488974858383769 | 32065.0 |
| 17 | rs62066125 | 43702701 | C | G | 4.044827941667506e-08 | 5.488883274990339 | 32163.0 |
| 17 | rs2696462 | 44286089 | G | A | 4.055101600489803e-08 | 5.488435120166723 | 31729.0 |
| 17 | rs56013908 | 43836920 | T | C | 4.05515765769718e-08 | 5.488432677875389 | 32156.0 |
| 17 | rs62053948 | 43748046 | C | T | 4.0677757903938884e-08 | 5.487883764449598 | 32180.0 |
| 17 | rs62054760 | 43908989 | C | T | 4.068338112855907e-08 | 5.487859340769485 | 32004.0 |
| 17 | rs368303226 | 43798784 | A | AGAG | 4.089847669958917e-08 | 5.486927552503543 | 32154.0 |
| 17 | rs2188888 | 43953695 | T | C | 4.104457204460076e-08 | 5.486297377263427 | 32034.0 |
| 17 | rs62055665 | 43754505 | A | C | 4.1044713857461073e-08 | 5.486296766617754 | 32184.0 |
| 17 | rs62055690 | 43754576 | C | G | 4.1044713857461073e-08 | 5.486296766617754 | 32184.0 |
| 17 | rs62055691 | 43754850 | T | C | 4.1044713857461073e-08 | 5.486296766617754 | 32184.0 |
| 17 | rs439558 | 43717803 | T | C | 4.1169410925656896e-08 | 5.485760611682926 | 32186.0 |
| 17 | rs62056911 | 43784624 | T | C | 4.118406369397058e-08 | 5.485697713184396 | 32169.0 |
| 17 | rs62054761 | 43909008 | C | T | 4.127579494575778e-08 | 5.485304439894855 | 32004.0 |
| 17 | rs191247549 | 44326105 | G | C | 4.129105538327725e-08 | 5.48523909706774 | 32126.0 |
| 17 | rs17689218 | 43785808 | T | C | 4.142292413622057e-08 | 5.484675428952448 | 32168.0 |
| 17 | rs62056916 | 43785938 | C | T | 4.142292413622057e-08 | 5.484675428952448 | 32168.0 |
| 17 | rs62056917 | 43786013 | G | T | 4.142292413622057e-08 | 5.484675428952448 | 32168.0 |
| 17 | rs62064409 | 43666431 | T | C | 4.146172010652132e-08 | 5.484509928228432 | 32031.0 |
| 17 | 17:44035222_GAA_G | 44035222 | GAA | G | 4.149582119602732e-08 | 5.484364579467988 | 32046.0 |
| 17 | rs112972619 | 43776425 | C | T | 4.155979891084071e-08 | 5.484092200043572 | 31852.0 |
| 17 | 17:44078816_TA_T | 44078816 | TA | T | 4.167537509517772e-08 | 5.4836011737573305 | 31993.0 |
| 17 | rs62055689 | 43754569 | G | A | 4.168286100130112e-08 | 5.483569415374904 | 32183.0 |
| 17 | rs112637166 | 43751101 | A | G | 4.181826603659983e-08 | 5.482995922588413 | 32171.0 |
| 17 | 17:44195611_CA_C | 44195611 | CA | C | 4.1849040813125735e-08 | 5.4828658306155999 | 31957.0 |
| 17 | rs55840731 | 43769277 | C | G | 4.18507752632684e-08 | 5.48285850146226 | 32176.0 |
| 17 | rs643024 | 43690468 | C | T | 4.197816036238208e-08 | 5.482321022044027 | 32103.0 |
| 17 | rs3919455 | 43953707 | C | T | 4.212032256983217e-08 | 5.481723057240609 | 32033.0 |
| 17 | rs2214258 | 43953719 | G | C | 4.212032256983217e-08 | 5.481723057240609 | 32033.0 |
| 17 | rs62056919 | 43786698 | C | T | 4.217271725820815e-08 | 5.481503167308666 | 32163.0 |
| 17 | rs74398257 | 43756340 | T | C | 4.251293393354541e-08 | 5.480081758105996 | 32180.0 |
| 17 | rs147268270 | 44101849 | T | TC | 4.254420934036169e-08 | 5.479951644968195 | 31742.0 |
| 17 | rs62064423 | 43668187 | A | G | 4.256007590205618e-08 | 5.479885671748519 | 31990.0 |
| 17 | rs28439143 | 43987723 | G | A | 4.261315342896555e-08 | 5.479665148389719 | 32186.0 |
| 17 | 17:43920219_AAAG_A | 43920219 | AAAG | A | 4.264376492817247e-08 | 5.479538086606801 | 32131.0 |

| | | | | | | | |
|---|---|---|---|---|---|---|---|
| 17 | rs62056930 | 43789640 | A | C | 4.266865262942536e-08 | 5.479434848244572 | 32153.0 |
| 17 | rs17650872 | 44039516 | G | T | 4.268147018764252e-08 | 5.4793817016367665 | 32145.0 |
| 17 | rs62064418 | 43667836 | C | A | 4.268928043358622e-08 | 5.479349324890362 | 31759.0 |
| 17 | 17:44260931_GGAGAGA_G | 44260931 | GGAGAGA | G | 4.27413353404337e-08 | 5.479133682048863 | 31787.0 |
| 17 | 17:44260939_GA_G | 44260939 | GA | G | 4.27413353404337e-08 | 5.479133682048863 | 31787.0 |
| 17 | rs62056918 | 43786676 | T | G | 4.2761702315941496e-08 | 5.479049379042454 | 32164.0 |
| 17 | rs62064419 | 43667915 | T | C | 4.2946619323637904e-08 | 5.478285746781464 | 32006.0 |
| 17 | rs55801356 | 43837237 | A | G | 4.317447103219554e-08 | 5.477349183733881 | 32158.0 |
| 17 | rs1568951 | 43792895 | T | G | 4.3175514161612873e-08 | 5.47734490707798 | 32174.0 |
| 17 | rs1568950 | 43792896 | T | C | 4.3175514161612873e-08 | 5.47734490707798 | 32174.0 |
| 17 | rs150592114 | 44333331 | A | G | 4.327965310880003e-08 | 5.476918458263115 | 30979.0 |
| 17 | 17:43798584_CA_C | 43798584 | CA | C | 4.340710172944503e-08 | 5.476397908128688 | 32148.0 |
| 17 | rs2696594 | 44220790 | A | G | 4.346466346195538e-08 | 5.476163289038521 | 31802.0 |
| 17 | rs62064415 | 43667537 | C | T | 4.364397191563788e-08 | 5.475434361689466 | 31662.0 |
| 17 | rs2902661 | 43807099 | G | A | 4.412045088701528e-08 | 5.473511390894092 | 32148.0 |
| 17 | rs17687462 | 43744990 | C | T | 4.4309897692307736e-08 | 5.4727524113866615 | 32177.0 |
| 17 | rs56328224 | 43572419 | C | T | 4.431387206882576e-08 | 5.472736522583146 | 32026.0 |
| 17 | rs62053939 | 43740856 | C | T | 4.446807164251538e-08 | 5.4721211261583775 | 32186.0 |
| 17 | 17:43733707_GC_G | 43733707 | GC | G | 4.453883873954013e-08 | 5.471839393432158 | 32175.0 |
| 17 | rs62055752 | 43767628 | G | A | 4.457018952365929e-08 | 5.471714720703196 | 32173.0 |
| 17 | 17:43760759_AT_A | 43760759 | AT | A | 4.4732976041531197e-08 | 5.4710687309534665 | 32130.0 |
| 17 | 17:44204299_AC_A | 44204299 | AC | A | 4.484620571542011e-08 | 5.4706207414131125 | 31744.0 |
| 17 | rs17652337 | 44083323 | C | T | 4.485486780954675e-08 | 5.470586515285016 | 31978.0 |
| 17 | rs62073156 | 44333568 | T | C | 4.4857033589093564e-08 | 5.470577958742771 | 32136.0 |
| 17 | rs55946323 | 44031946 | A | G | 4.488241177755955e-08 | 5.47047772465789 | 32144.0 |
| 17 | rs138070769 | 43781726 | A | AAAG AG | 4.510149429803841e-08 | 5.4696147104339765 | 32125.0 |
| 17 | rs78568882 | 43784935 | C | G | 4.514724219225904e-08 | 5.4694350124412825 | 32157.0 |
| 17 | rs62066086 | 43688387 | A | G | 4.525915774430455e-08 | 5.468996150583633 | 32097.0 |
| 17 | rs62066087 | 43688579 | A | G | 4.525915774430455e-08 | 5.468996150583633 | 32097.0 |
| 17 | 17:43762168_CT_C | 43762168 | CT | C | 4.5378854669623086e-08 | 5.468527937918856 | 32179.0 |
| 17 | rs62055698 | 43758238 | C | T | 4.5459347250077873e-08 | 5.468213751802933 | 32181.0 |
| 17 | rs61667602 | 43785349 | T | C | 4.5714113343674044e-08 | 5.467222868059473 | 32159.0 |
| 17 | rs2532309 | 44220785 | G | A | 4.5750847771906457e-08 | 5.46708043547786 | 31795.0 |
| 17 | rs62055889 | 43838919 | G | A | 4.599119910644416e-08 | 5.466151233765216 | 32150.0 |
| 17 | rs56155809 | 44119500 | C | CTT | 4.602037804174151e-08 | 5.466038748182516 | 31824.0 |
| 17 | 17:43814930_GCGTA_G | 43814930 | GCGTA | G | 4.610611659230668e-08 | 5.465708623372866 | 32135.0 |
| 17 | rs62063266 | 44033488 | C | T | 4.611994464945056e-08 | 5.465655436029004 | 32125.0 |
| 17 | rs62055701 | 43758787 | G | A | 4.638377046783034e-08 | 5.464643623725539 | 32019.0 |
| 17 | rs62055702 | 43758790 | C | T | 4.638377046783034e-08 | 5.464643623725539 | 32019.0 |
| 17 | rs7502718 | 43793651 | G | A | 4.707730273118991e-08 | 5.462010198384951 | 32169.0 |
| 17 | rs62055886 | 43838678 | G | A | 4.726923999486996e-08 | 5.461288032935883 | 32114.0 |
| 17 | rs62055887 | 43838710 | T | C | 4.726923999486996e-08 | 5.461288032935883 | 32114.0 |
| 17 | rs34039488 | 27320232 | G | A | 4.7422230508020914e-08 | 5.460714437942859 | 31978.0 |
| 17 | rs112904481 | 43766494 | A | AC | 4.744020356224431e-08 | 5.460647170802476 | 32167.0 |
| 17 | rs2942166 | 43715427 | T | C | 4.7649999746248186e-08 | 5.459863795780676 | 32185.0 |
| 17 | 17:43683792_ATGG_A | 43683792 | ATGG | A | 4.786400132129488e-08 | 5.45906815502009 | 32167.0 |
| 17 | rs111370985 | 43852742 | T | C | 4.7930481536743276e-08 | 5.458821688800632 | 32089.0 |
| 17 | rs11079721 | 43840107 | C | A | 4.794451496122485e-08 | 5.458769704177463 | 32117.0 |
| 17 | rs532036186 | 43959596 | CT | C | 4.80154039030534e-08 | 5.45850733242473 | 31650.0 |
| 17 | rs62064420 | 43667922 | G | C | 4.804501261146834e-08 | 5.458397856827787 | 32061.0 |
| 17 | rs568589031 | 44311099 | G | A | 4.806619628244283e-08 | 5.458319572191846 | 31729.0 |
| 17 | rs2532389 | 44311107 | G | C | 4.806619628244283e-08 | 5.458319572191846 | 31729.0 |
| 17 | rs62073222 | 44374710 | A | G | 4.8359899613955495e-08 | 5.4572376190285725 | 23951.0 |
| 17 | rs375647 | 43693341 | A | G | 4.843906169489807e-08 | 5.456947089103856 | 32128.0 |

| | | | | | | | |
|---|---|---|---|---|---|---|---|
| 17 | rs2141300 | 44258914 | A | G | 4.8449737439503776e-08 | 5.456907943659128 | 31742.0 |
| 17 | rs117365970 | 43893259 | G | A | 4.8755638029570635e-08 | 5.45578981578513 | 32110.0 |
| 17 | rs62064417 | 43667792 | C | T | 4.8940456012793466e-08 | 5.455117559806073 | 31503.0 |
| 17 | rs62063265 | 44033468 | G | A | 4.897315200719213e-08 | 5.4549988877965445 | 32123.0 |
| 17 | rs62063262 | 44033002 | T | C | 4.9018692344638335e-08 | 5.454833724413245 | 32128.0 |
| 17 | rs62057147 | 43903298 | G | A | 4.9356982113501856e-08 | 5.453611468056416 | 32143.0 |
| 17 | rs34097347 | 43949448 | C | A | 4.944897372052344e-08 | 5.453280502415764 | 31939.0 |
| 17 | rs389978 | 43717415 | A | G | 4.952289240128637e-08 | 5.4530149913641885 | 32182.0 |
| 17 | rs113343751 | 44333532 | C | G | 4.957507520864914e-08 | 5.4528277853545495 | 32143.0 |
| 17 | rs111891516 | 44198962 | T | TA | 4.961569905761597e-08 | 5.4526821793281295 | 32052.0 |
| 17 | rs200422537 | 44331803 | A | ATTTT | 4.9667806688046064e-08 | 5.452495581640127 | 31892.0 |
| 17 | rs2316956 | 44162056 | T | A | 4.970901815494881e-08 | 5.452348137502433 | 31923.0 |
| 17 | rs62063269 | 44034833 | G | A | 4.9720994564540985e-08 | 5.452305311176965 | 32104.0 |
| 17 | rs2732677 | 44342123 | G | C | 4.97636192699658e-08 | 5.452152970989569 | 30487.0 |
| 17 | rs2532261 | 44253203 | A | C | 4.984314392428153e-08 | 5.451869088630539 | 31629.0 |
| 18 | rs7234875 | 73114340 | T | C | 5.719962038078088e-14 | 7.51433802050141 | 32186.0 |
| 18 | rs7231353 | 73109128 | A | C | 2.4874962193190062e-12 | 7.004003653460147 | 31869.0 |
| 18 | rs7407699 | 73110452 | G | C | 3.7131410785140635e-12 | 6.947690644649823 | 32029.0 |
| 18 | rs6566091 | 73107145 | A | G | 1.2580119089949947e-11 | 6.773387748219499 | 32011.0 |
| 18 | rs10460110 | 73105287 | T | C | 1.6529841541532062e-11 | 6.733791136562622 | 32019.0 |
| 18 | rs4274504 | 73102466 | T | C | 1.722861821551898e-11 | 6.727767273229808 | 32037.0 |
| 18 | rs11151123 | 73107776 | C | T | 3.1837315227321664e-11 | 6.637812189163254 | 32044.0 |
| 22 | rs2542028 | 47196524 | A | G | 3.0626057109556033e-12 | 6.9748209684906515 | 31938.0 |
| 22 | rs801593 | 47195050 | G | C | 8.481045943624013e-12 | 6.8301760991401625 | 32186.0 |
| 22 | rs801592 | 47196383 | T | G | 9.846628833208065e-12 | 6.808726764044822 | 32182.0 |
| 22 | rs809031 | 47191699 | A | G | 1.0080353444030328e-11 | 6.805350466848145 | 32111.0 |
| 22 | rs801597 | 47190735 | T | G | 1.1371712070918635e-11 | 6.787976559775465 | 32104.0 |
| 22 | rs710119 | 47199148 | A | G | 2.0987560288025753e-11 | 6.698982583455219 | 32029.0 |
| 22 | rs710122 | 47198055 | G | A | 2.3010809873499833e-11 | 6.6855187223354395 | 32099.0 |
| 22 | rs910868 | 47198932 | C | G | 6.802133167180513e-11 | 6.5249563697014015 | 31887.0 |
| 22 | rs910867 | 47198930 | T | A | 7.45428869124417e-11 | 6.5112199734893705 | 31892.0 |
| 22 | rs801652 | 47202691 | C | T | 9.809235692532744e-11 | 6.469862160141076 | 31944.0 |
| 22 | rs801656 | 47201690 | A | G | 1.1916106927980164e-10 | 6.4403974202827134 | 31548.0 |
| 22 | rs4823877 | 47200353 | T | C | 1.2204880708498223e-10 | 6.436762336726692 | 31496.0 |
| 22 | rs4823878 | 47200354 | T | C | 1.5181624236894216e-10 | 6.4035388087640355 | 31482.0 |
| 22 | rs910865 | 47198133 | G | C | 2.631400757667044e-10 | 6.319071170837108 | 31919.0 |
| 22 | rs17822045 | 47208009 | C | T | 2.6328342883298037e-10 | 6.318986995635189 | 31913.0 |
| 22 | rs62233790 | 47206644 | A | G | 3.460402905686263e-10 | 6.2766074420229065 | 31910.0 |
| 22 | rs710118 | 47199389 | G | C | 3.8948464124822953e-10 | 6.258184596646344 | 31748.0 |
| 22 | 22:47168498_CATATACAT_C | 47168498 | CATATACAT | C | 1.5683446648449386e-09 | 6.037176177246598 | 30616.0 |
| 22 | rs139577 | 47208945 | G | A | 4.892453819870287e-09 | 5.850789401327728 | 31282.0 |
| 22 | rs145865429 | 47206351 | A | AT | 1.9613525990791032e-08 | 5.615376009043104 | 29117.0 |

| Note: | Bold rows represents FCs extracted from non british replication dataset significantly associated with the lead SNP identified in the discovery sample at the p=0.05 threshold. | | | | | | |
|---|---|---|---|---|---|---|---|
| CHR | SNP | BP | A1 | A2 | P value | Z-stat | N |
| **10** | **rs11187838** | **96038686** | **G** | **A** | **0.029236832632864824** | **2.180278380453736** | **4734.0** |
| 10 | rs11146399 | 134308479 | C | T | 0.28794094063036857 | 1.0626494769648172 | 4682.0 |
| 3 | rs67851870 | 17554860 | A | G | 0.3508043494236116 | 0.9330302511281164 | 4710.0 |
| **3** | **rs35124509** | **89521693** | **T** | **C** | **0.003255078387998134** | **2.9425627224280984** | **4754.0** |
| 3 | rs62266110 | 93537923 | G | A | 0.9361934712998444 | 0.08005505195419887 | 4652.0 |
| 3 | rs2279829 | 147106319 | C | T | 0.6847156850862024 | 0.40603663165285736 | 4682.0 |
| 6 | rs4262195 | 96929475 | T | C | 0.7098970843773096 | 0.3719943119238445 | 4749.0 |
| **15** | **rs1440802** | **39635124** | **T** | **C** | **0.009584804947221632** | **2.590458959618914** | **4692.0** |
| 15 | rs4702 | 91426560 | G | A | 0.4220213678843896 | 0.8029193209635919 | 4754.0 |
| 2 | rs62141276 | 48214217 | G | A | 0.27258732810218 | 1.0971236842802248 | 4563.0 |
| 2 | rs2717046 | 58041936 | C | T | 0.9522042271394622 | 0.059938988788391635 | 4680.0 |
| 2 | rs62158166 | 114077218 | G | C | 0.3888361442755477 | 0.8617299430041099 | 4642.0 |
| 18 | rs7234875 | 73114340 | T | C | 0.8250427443389228 | 0.2210638154437743 | 4754.0 |
| 5 | rs145120402 | 93174765 | A | C | 0.10257126040076994 | 1.632513958457583 | 4744.0 |
| 5 | 5:94068140_AC_A | 94068140 | AC | A | 0.30022036800097285 | 1.0359609344902436 | 4694.0 |
| 14 | rs186347 | 59072226 | G | T | 0.9203976014436072 | 0.0999328939702151 | 4629.0 |
| 22 | rs2542028 | 47196524 | A | G | 0.6040247767444131 | 0.5186214087336436 | 4729.0 |
| 11 | rs11218557 | 122099839 | T | C | 0.7774024921231244 | 0.2827056404346454 | 4651.0 |
| 17 | rs34039488 | 27320232 | G | A | 0.4619048505517788 | 0.7357138640842442 | 4706.0 |
| 17 | 17:44270659_G_A | 44270659 | G | A | 0.4504527168736029 | 0.7546604931459203 | 4551.0 |

| Genomic Locus | Lead SNP | chr | Functional category | Non effect allele | Effect allele | MAF | mvGWAS P value | Nearest gene | pheno_label | univariate P value | Beta | univariate Z value |
|---|---|---|---|---|---|---|---|---|---|---|---|---|
| 5 | rs35124509 | 3 | exonic | C | T | 0.4018 | 8,9515E-59 | EPHA3 | Pole_STSp | 1,4963E-15 | -6,4371E-02 | -7,98 |
| | | | | | | | | | Pole_T2ml | 5,9579E-15 | 6,2982E-02 | 7,80 |
| | | | | | | | | | AG_F3orb | 4,0019E-13 | 5,8557E-02 | 7,26 |
| | | | | | | | | | T1a_STSp | 1,8916E-11 | -5,4194E-02 | -6,71 |
| | | | | | | | | | STSp_F3orb | 2,0435E-11 | -5,4103E-02 | -6,70 |
| | | | | | | | | | SMG_T3p | 1,0825E-09 | 4,9217E-02 | 6,10 |
| | | | | | | | | | AG_STSp | 2,0124E-09 | 4,8411E-02 | 6,00 |
| | | | | | | | | | F2p_AG | 6,6439E-09 | 4,6821E-02 | 5,80 |
| | | | | | | | | | T2ml_SMG | 2,7656E-08 | -4,4853E-02 | -5,56 |
| 11 | rs11187838 | 10 | intronic | A | G | 0.4354 | 4,2917E-14 | PLCE1 | RolS_PrecR | 1,8523E-07 | -4,1372E-02 | -5,21 |
| | | | | | | | | | T1a_T1a/HeschlR | 9,6142E-06 | -3,5124E-02 | -4,43 |
| 15 | rs1440802 | 15 | NcRNA intronic | C | T | 0.09091 | 1,0033E-31 | RP11-624L4.1 | Prec_F3opd | 2,7889E-19 | -1,2300E-01 | -8,98 |
| | | | | | | | | | RolS_PrecR | 8,4664E-17 | -1,1409E-01 | -8,32 |

Note: Bold phenotypes represents pair of FCs with significant genetic correlation after false discovery rate correction at $p<0.05$.

| Phenotype 1 | Phenotype 2 | P value | rG | SE | FDR |
|---|---|---|---|---|---|
| **T1a_STSp** | **Pole_STSp** | **2.29E-05** | **54.8%** | **0.132** | **0.000847** |
| **SMG_T3p** | **AG_STSp** | **9.78E-05** | **65.6%** | **0.192** | **0.00181** |
| **STSp_F3orb** | **Pole_STSp** | **0.000221** | **40.4%** | **0.111** | **0.00272** |
| **AG_F3orb** | **T2ml_SMG** | **0.00101** | **-46.9%** | **0.158** | **0.00939** |
| **AG_F3orb** | **SMG_T3p** | **0.00376** | **53.6%** | **0.211** | **0.0139** |
| **Pole_T2ml** | **STSp_F3orb** | **0.00325** | **-31.4%** | **0.113** | **0.0139** |
| **AG_F3orb** | **T1a_STSp** | **0.00284** | **-52.5%** | **0.196** | **0.0139** |
| **Pole_T2ml** | **AG_STSp** | **0.00285** | **36.2%** | **0.132** | **0.0139** |
| **Pole_T2ml** | **T2ml_SMG** | **0.00252** | **-30.5%** | **0.109** | **0.0139** |
| **Pole_T2ml** | **Pole_STSp** | **0.00353** | **-28.7%** | **0.099** | **0.0139** |
| **AG_F3orb** | **Pole_STSp** | **0.00634** | **-36.5%** | **0.15** | **0.018** |
| **Pole_T2ml** | **SMG_T3p** | **0.00681** | **35.1%** | **0.146** | **0.018** |
| **F2p_AG** | **AG_STSp** | **0.00639** | **54.5%** | **0.25** | **0.018** |
| **SMG_T3p** | **STSp_F3orb** | **0.00553** | **-39.6%** | **0.16** | **0.018** |
| **F2p_AG** | **T1a_STSp** | **0.0107** | **-51.2%** | **0.225** | **0.0258** |
| **AG_F3orb** | **STSp_F3orb** | **0.0112** | **-38.2%** | **0.142** | **0.0258** |
| **T1a_STSp** | **STSp_F3orb** | **0.0121** | **32.9%** | **0.153** | **0.0263** |
| **AG_F3orb** | **F2p_AG** | **0.0139** | **53.5%** | **0.243** | **0.0285** |
| **Pole_STSp** | **T2ml_SMG** | **0.0168** | **22.9%** | **0.109** | **0.0316** |
| **T1a_STSp** | **SMG_T3p** | **0.0171** | **-38.3%** | **0.187** | **0.0316** |
| **F2p_AG** | **T2ml_SMG** | **0.024** | **-34.9%** | **0.18** | **0.0404** |
| **SMG_T3p** | **T2ml_SMG** | **0.0233** | **-29.0%** | **0.144** | **0.0404** |
| STSp_F3orb | T2ml_SMG | 0.0398 | 20.7% | 0.12 | 0.064 |
| AG_STSp | T2ml_SMG | 0.0484 | -22.3% | 0.137 | 0.0746 |
| T1a_STSp | T2ml_SMG | 0.056 | 21.9% | 0.137 | 0.0828 |
| SMG_T3p | Pole_STSp | 0.0605 | -21.8% | 0.142 | 0.0861 |
| AG_STSp | Pole_STSp | 0.0706 | -19.3% | 0.127 | 0.0968 |
| AG_STSp | STSp_F3orb | 0.0866 | -19.6% | 0.139 | 0.114 |
| F2p_AG | SMG_T3p | 0.0902 | 31.1% | 0.232 | 0.115 |
| Prec_F3opd | RolS_PrecR | 0.098 | 16.9% | 0.131 | 0.121 |
| Pole_T2ml | T1a_STSp | 0.108 | -16.6% | 0.135 | 0.129 |
| AG_F3orb | AG_STSp | 0.191 | 16.0% | 0.184 | 0.221 |
| F2p_AG | Pole_STSp | 0.208 | 14.0% | 0.176 | 0.233 |
| Pole_T2ml | F2p_AG | 0.215 | -13.7% | 0.176 | 0.234 |
| T1a_STSp | AG_STSp | 0.222 | -12.8% | 0.164 | 0.234 |
| F2p_AG | STSp_F3orb | 0.388 | 5.4% | 0.189 | 0.399 |
| Pole_T2ml | AG_F3orb | 0.456 | 1.6% | 0.148 | 0.456 |

Note: Bold phenotypes represents tracts significantly associated with the lead SNP after Bonferroni correction (p=6.94e-4 (0.05/(3*9+5*9))

| CHR | SNP | Pheno | Metric | Beta | P | Z |
|---|---|---|---|---|---|---|
| 15 | rs1440802 | Corpus callosum | L3 | -3,4428E-02 | 1,2803E-02 | -2,4892 |
| | | | ISOVF | -3,3499E-02 | 1,5433E-02 | -2,4221 |
| | | | FA | 3,2574E-02 | 1,8515E-02 | 2,3551 |
| | | | MD | -3,2090E-02 | 2,0335E-02 | -2,3201 |
| | | | L2 | -3,1503E-02 | 2,2745E-02 | -2,2777 |
| | | | L1 | -2,4891E-02 | 7,1920E-02 | -1,7996 |
| | | | ICVF | 2,0728E-02 | 1,3398E-01 | 1,4986 |
| | | | MO | -7,1922E-03 | 6,0309E-01 | -0,5200 |
| | | | OD | -7,1261E-03 | 6,0642E-01 | -0,5152 |
| | | Frontal Aslant Tract Left | **MO** | **-6,4280E-02** | **3,3396E-06** | **-4,6487** |
| | | | OD | 5,6007E-02 | 5,1196E-05 | 4,0501 |
| | | | L1 | -5,4971E-02 | 7,0339E-05 | -3,9751 |
| | | | ISOVF | -3,4517E-02 | 1,2573E-02 | -2,4956 |
| | | | MD | -3,0007E-02 | 3,0042E-02 | -2,1695 |
| | | | L3 | -1,9929E-02 | 1,4963E-01 | -1,4408 |
| | | | ICVF | 9,3029E-03 | 5,0123E-01 | 0,6726 |
| | | | FA | -8,9362E-03 | 5,1825E-01 | -0,6460 |
| | | | L2 | -3,8335E-03 | 7,8167E-01 | -0,2771 |
| | | Arcuate Anterior Segment Left | **OD** | **-2,3504E-01** | **2,4732E-65** | **-17,0702** |
| | | | MO | 2,0941E-01 | 3,8548E-52 | 15,1943 |
| | | | FA | 1,4842E-01 | 5,9537E-27 | 10,7496 |
| | | | L2 | -1,4630E-01 | 3,1250E-26 | -10,5956 |
| | | | L1 | 1,0825E-01 | 4,7482E-15 | 7,8334 |
| | | | L3 | -4,8700E-02 | 4,2921E-04 | -3,5214 |
| | | | ISOVF | -4,6469E-02 | 7,7922E-04 | -3,3601 |
| | | | MD | -3,2275E-02 | 1,9622E-02 | -2,3335 |
| | | | ICVF | 1,8871E-02 | 1,7246E-01 | 1,3644 |
| | | Inferior Fronto Occipital fasciculus Left | **MO** | **4,1907E-02** | **2,4978E-07** | **5,1579** |
| | | | OD | -3,2217E-02 | 7,3529E-05 | -3,9646 |
| | | | L1 | 2,0604E-02 | 1,1242E-02 | 2,5351 |
| | | | FA | 1,1022E-02 | 1,7507E-01 | 1,3561 |
| | | | MD | 8,8824E-03 | 2,7448E-01 | 1,0928 |
| | | | ISOVF | 4,8581E-03 | 5,5004E-01 | 0,5977 |
| | | | L2 | -4,7179E-03 | 5,6162E-01 | -0,5804 |
| | | | L3 | 4,6885E-03 | 5,6406E-01 | 0,5768 |
| | | | ICVF | -3,7015E-03 | 6,4883E-01 | -0,4554 |
| | | Uncinate Left | **MO** | **4,2154E-02** | **2,1220E-07** | **5,1883** |
| | | | OD | -2,3885E-02 | 3,2932E-03 | -2,9390 |
| | | | L1 | 2,2805E-02 | 5,0163E-03 | 2,8060 |
| | | | FA | 1,2425E-02 | 1,2634E-01 | 1,5287 |
| | | | MD | 9,7547E-03 | 2,3009E-01 | 1,2001 |
| | | | ICVF | -8,3742E-03 | 3,0287E-01 | -1,0303 |
| | | | L3 | 7,2253E-03 | 3,7404E-01 | 0,8889 |
| | | | L2 | -5,9861E-03 | 4,6144E-01 | -0,7365 |
| | | | ISOVF | -1,5708E-04 | 9,8458E-01 | -0,0193 |

| | | | | | | |
|---|---|---|---|---|---|---|
| 3 | rs35124509 | Arcuate Anterior Segment Left | MO | -2,3605E-02 | 3,6791E-03 | -2,9044 |
| | | | L2 | 2,2630E-02 | 5,3612E-03 | 2,7845 |
| | | | FA | -1,6705E-02 | 3,9844E-02 | -2,0554 |
| | | | ISOVF | 1,2345E-02 | 1,2880E-01 | 1,5189 |
| | | | MD | 1,0160E-02 | 2,1131E-01 | 1,2500 |
| | | | ICVF | -1,0146E-02 | 2,1193E-01 | -1,2483 |
| | | | L3 | 7,1300E-03 | 3,8037E-01 | 0,8772 |
| | | | OD | 5,8026E-03 | 4,7529E-01 | 0,7139 |
| | | | L1 | -7,7840E-04 | 9,2371E-01 | -0,0958 |
| | | Arcuate Long Segment Left | **OD** | **-3,6813E-02** | **5,8822E-06** | **-4,5306** |
| | | | L1 | 2,0134E-02 | 1,3239E-02 | 2,4773 |
| | | | MO | 1,4725E-02 | 7,0039E-02 | 1,8117 |
| | | | MD | 1,2833E-02 | 1,1437E-01 | 1,5788 |
| | | | ISOVF | 1,2218E-02 | 1,3278E-01 | 1,5032 |
| | | | L3 | 7,4946E-03 | 3,5649E-01 | 0,9221 |
| | | | ICVF | -6,6475E-03 | 4,1345E-01 | -0,8178 |
| | | | FA | 3,6834E-03 | 6,5043E-01 | 0,4532 |
| | | | L2 | 1,7721E-03 | 8,2741E-01 | 0,2180 |
| | | Arcuate Posterior Segment Left | **L3** | **4,1222E-02** | **3,9039E-07** | **5,0736** |
| | | | ICVF | -3,8005E-02 | 2,9062E-06 | -4,6773 |
| | | | MD | 3,6748E-02 | 6,1105E-06 | 4,5225 |
| | | | FA | -3,6428E-02 | 7,3549E-06 | -4,4832 |
| | | | L2 | 3,0797E-02 | 1,5079E-04 | 3,7898 |
| | | | L1 | 2,1984E-02 | 6,8311E-03 | 2,7050 |
| | | | MO | 1,6926E-02 | 3,7298E-02 | 2,0825 |
| | | | ISOVF | 1,4388E-02 | 7,6693E-02 | 1,7702 |
| | | | OD | 4,6444E-03 | 5,6773E-01 | 0,5714 |

| rs ID | chr | position | Data base | tissue | gene | tested Allele | P value | signed stats | FDR p value |
|---|---|---|---|---|---|---|---|---|---|
| rs73145317 | 3 | 89583712 | GTEx/v8 | Pancreas | EPHA3 | C | 3.35696E-11 | 0.45158 | 3.49803E-07 |
| rs73139276 | 3 | 89500196 | GTEx/v8 | Pancreas | EPHA3 | A | 9.15352E-11 | 0.390181 | 3.49803E-07 |
| rs73139288 | 3 | 89505327 | GTEx/v8 | Pancreas | EPHA3 | G | 9.30731E-11 | 0.388198 | 3.49803E-07 |
| rs73139129 | 3 | 89524562 | GTEx/v8 | Pancreas | EPHA3 | G | 9.30731E-11 | 0.388198 | 3.49803E-07 |
| rs972030 | 3 | 89538498 | GTEx/v8 | Pancreas | EPHA3 | T | 1.06317E-10 | 0.383465 | 3.49803E-07 |
| rs4857500 | 3 | 89541486 | GTEx/v8 | Pancreas | EPHA3 | C | 1.06317E-10 | 0.383465 | 3.49803E-07 |
| rs1512189 | 3 | 89546332 | GTEx/v8 | Pancreas | EPHA3 | G | 1.07905E-10 | 0.384892 | 3.49803E-07 |
| rs12629999 | 3 | 89551518 | GTEx/v8 | Pancreas | EPHA3 | G | 1.07905E-10 | 0.384892 | 3.49803E-07 |
| rs73146986 | 3 | 89560884 | GTEx/v8 | Pancreas | EPHA3 | G | 1.07905E-10 | 0.384892 | 3.49803E-07 |
| rs6784577 | 3 | 89564308 | GTEx/v8 | Pancreas | EPHA3 | G | 1.07905E-10 | 0.384892 | 3.49803E-07 |
| rs73145303 | 3 | 89568714 | GTEx/v8 | Pancreas | EPHA3 | C | 1.07905E-10 | 0.384892 | 3.49803E-07 |
| rs73139133 | 3 | 89527708 | GTEx/v8 | Pancreas | EPHA3 | A | 1.35747E-10 | 0.384023 | 3.49803E-07 |
| rs144317613 | 3 | 89525080 | GTEx/v8 | Pancreas | EPHA3 | TA | 1.49946E-10 | 0.383137 | 3.49803E-07 |
| rs73149103 | 3 | 89565651 | GTEx/v8 | Pancreas | EPHA3 | A | 1.5191E-10 | 0.38131 | 3.49803E-07 |
| rs73139261 | 3 | 89492406 | GTEx/v8 | Pancreas | EPHA3 | G | 1.57576E-10 | 0.386012 | 3.49803E-07 |
| rs12637036 | 3 | 89584170 | GTEx/v8 | Pancreas | EPHA3 | A | 1.7134E-10 | 0.385057 | 3.49803E-07 |
| rs56082199 | 3 | 89586973 | GTEx/v8 | Pancreas | EPHA3 | G | 1.7134E-10 | 0.385057 | 3.49803E-07 |
| rs73145336 | 3 | 89593317 | GTEx/v8 | Pancreas | EPHA3 | G | 1.7134E-10 | 0.385057 | 3.49803E-07 |
| rs1036285 | 3 | 89544061 | GTEx/v8 | Pancreas | EPHA3 | C | 1.81497E-10 | 0.361236 | 3.49803E-07 |
| rs17026984 | 3 | 89544978 | GTEx/v8 | Pancreas | EPHA3 | A | 1.81497E-10 | 0.361236 | 3.49803E-07 |
| rs111618480 | 3 | 89552134 | GTEx/v8 | Pancreas | EPHA3 | C | 1.81497E-10 | 0.361236 | 3.49803E-07 |
| rs2048518 | 3 | 89533905 | GTEx/v8 | Pancreas | EPHA3 | C | 1.83068E-10 | 0.364599 | 3.49803E-07 |
| rs2048520 | 3 | 89534248 | GTEx/v8 | Pancreas | EPHA3 | C | 1.83068E-10 | 0.364599 | 3.49803E-07 |
| rs2048521 | 3 | 89534253 | GTEx/v8 | Pancreas | EPHA3 | A | 1.83068E-10 | 0.364599 | 3.49803E-07 |
| rs73144998 | 3 | 89534592 | GTEx/v8 | Pancreas | EPHA3 | T | 1.83068E-10 | 0.364599 | 3.49803E-07 |
| rs73145002 | 3 | 89537618 | GTEx/v8 | Pancreas | EPHA3 | C | 1.83068E-10 | 0.364599 | 3.49803E-07 |
| rs56371328 | 3 | 89539124 | GTEx/v8 | Pancreas | EPHA3 | A | 1.83068E-10 | 0.364599 | 3.49803E-07 |
| rs4857498 | 3 | 89540977 | GTEx/v8 | Pancreas | EPHA3 | A | 1.83068E-10 | 0.364599 | 3.49803E-07 |
| rs4857499 | 3 | 89541330 | GTEx/v8 | Pancreas | EPHA3 | A | 1.83068E-10 | 0.364599 | 3.49803E-07 |
| rs2346837 | 3 | 89542599 | GTEx/v8 | Pancreas | EPHA3 | T | 1.83068E-10 | 0.364599 | 3.49803E-07 |
| rs12633609 | 3 | 89545727 | GTEx/v8 | Pancreas | EPHA3 | G | 1.83068E-10 | 0.364599 | 3.49803E-07 |
| rs1512188 | 3 | 89546183 | GTEx/v8 | Pancreas | EPHA3 | G | 1.83068E-10 | 0.364599 | 3.49803E-07 |
| rs73146943 | 3 | 89546728 | GTEx/v8 | Pancreas | EPHA3 | G | 1.83068E-10 | 0.364599 | 3.49803E-07 |
| rs12632281 | 3 | 89547621 | GTEx/v8 | Pancreas | EPHA3 | A | 1.83068E-10 | 0.364599 | 3.49803E-07 |
| rs73146947 | 3 | 89548135 | GTEx/v8 | Pancreas | EPHA3 | G | 1.83068E-10 | 0.364599 | 3.49803E-07 |
| rs73146950 | 3 | 89548243 | GTEx/v8 | Pancreas | EPHA3 | C | 1.83068E-10 | 0.364599 | 3.49803E-07 |
| rs1567657 | 3 | 89548519 | GTEx/v8 | Pancreas | EPHA3 | A | 1.83068E-10 | 0.364599 | 3.49803E-07 |
| rs1567658 | 3 | 89548606 | GTEx/v8 | Pancreas | EPHA3 | A | 1.83068E-10 | 0.364599 | 3.49803E-07 |
| rs73146959 | 3 | 89549970 | GTEx/v8 | Pancreas | EPHA3 | C | 1.83068E-10 | 0.364599 | 3.49803E-07 |
| rs73146960 | 3 | 89550445 | GTEx/v8 | Pancreas | EPHA3 | C | 1.83068E-10 | 0.364599 | 3.49803E-07 |
| rs112404167 | 3 | 89550630 | GTEx/v8 | Pancreas | EPHA3 | T | 1.83068E-10 | 0.364599 | 3.49803E-07 |
| rs12637511 | 3 | 89550844 | GTEx/v8 | Pancreas | EPHA3 | C | 1.83068E-10 | 0.364599 | 3.49803E-07 |
| rs73146972 | 3 | 89553348 | GTEx/v8 | Pancreas | EPHA3 | G | 1.83068E-10 | 0.364599 | 3.49803E-07 |
| rs17026990 | 3 | 89555257 | GTEx/v8 | Pancreas | EPHA3 | C | 1.83068E-10 | 0.364599 | 3.49803E-07 |
| rs73146976 | 3 | 89557065 | GTEx/v8 | Pancreas | EPHA3 | C | 1.83068E-10 | 0.364599 | 3.49803E-07 |
| rs1039991 | 3 | 89558311 | GTEx/v8 | Pancreas | EPHA3 | G | 1.83068E-10 | 0.364599 | 3.49803E-07 |
| rs73146980 | 3 | 89558578 | GTEx/v8 | Pancreas | EPHA3 | A | 1.83068E-10 | 0.364599 | 3.49803E-07 |
| rs73146983 | 3 | 89559082 | GTEx/v8 | Pancreas | EPHA3 | G | 1.83068E-10 | 0.364599 | 3.49803E-07 |
| rs138490449 | 3 | 89561661 | GTEx/v8 | Pancreas | EPHA3 | AT | 1.83068E-10 | 0.364599 | 3.49803E-07 |
| rs138183278 | 3 | 89564090 | GTEx/v8 | Pancreas | EPHA3 | CT | 1.83068E-10 | 0.364599 | 3.49803E-07 |
| rs6774870 | 3 | 89564891 | GTEx/v8 | Pancreas | EPHA3 | T | 1.83068E-10 | 0.364599 | 3.49803E-07 |
| rs113307255 | 3 | 89567826 | GTEx/v8 | Pancreas | EPHA3 | C | 1.83068E-10 | 0.364599 | 3.49803E-07 |

| | | | | | | | | | |
|---|---|---|---|---|---|---|---|---|---|
| rs2346836 | 3 | 89498096 | GTEx/v8 | Pancreas | EPHA3 | A | 2.29119E-10 | 0.368817 | 3.49803E-07 |
| rs60759140 | 3 | 89499848 | GTEx/v8 | Pancreas | EPHA3 | C | 2.29119E-10 | 0.368817 | 3.49803E-07 |
| rs12152372 | 3 | 89500365 | GTEx/v8 | Pancreas | EPHA3 | C | 2.29119E-10 | 0.368817 | 3.49803E-07 |
| rs2881488 | 3 | 89500488 | GTEx/v8 | Pancreas | EPHA3 | G | 2.29119E-10 | 0.368817 | 3.49803E-07 |
| rs59541469 | 3 | 89501671 | GTEx/v8 | Pancreas | EPHA3 | T | 2.29119E-10 | 0.368817 | 3.49803E-07 |
| 3:89502123_TTAG_T | 3 | 89502123 | GTEx/v8 | Pancreas | EPHA3 | T | 2.29119E-10 | 0.368817 | 3.49803E-07 |
| rs73139290 | 3 | 89505343 | GTEx/v8 | Pancreas | EPHA3 | A | 2.29119E-10 | 0.368817 | 3.49803E-07 |
| rs73139296 | 3 | 89505734 | GTEx/v8 | Pancreas | EPHA3 | T | 2.29119E-10 | 0.368817 | 3.49803E-07 |
| rs1157608 | 3 | 89506464 | GTEx/v8 | Pancreas | EPHA3 | G | 2.29119E-10 | 0.368817 | 3.49803E-07 |
| rs73139302 | 3 | 89508186 | GTEx/v8 | Pancreas | EPHA3 | G | 2.29119E-10 | 0.368817 | 3.49803E-07 |
| rs73141104 | 3 | 89508390 | GTEx/v8 | Pancreas | EPHA3 | T | 2.29119E-10 | 0.368817 | 3.49803E-07 |
| rs1912965 | 3 | 89518332 | GTEx/v8 | Pancreas | EPHA3 | G | 2.29119E-10 | 0.368817 | 3.49803E-07 |
| rs113141104 | 3 | 89519238 | GTEx/v8 | Pancreas | EPHA3 | T | 2.29119E-10 | 0.368817 | 3.49803E-07 |
| rs73139105 | 3 | 89519651 | GTEx/v8 | Pancreas | EPHA3 | C | 2.29119E-10 | 0.368817 | 3.49803E-07 |
| rs112297761 | 3 | 89521602 | GTEx/v8 | Pancreas | EPHA3 | TC | 2.29119E-10 | 0.368817 | 3.49803E-07 |
| rs73139119 | 3 | 89522285 | GTEx/v8 | Pancreas | EPHA3 | T | 2.29119E-10 | 0.368817 | 3.49803E-07 |
| rs73139121 | 3 | 89522433 | GTEx/v8 | Pancreas | EPHA3 | G | 2.29119E-10 | 0.368817 | 3.49803E-07 |
| rs73139131 | 3 | 89524686 | GTEx/v8 | Pancreas | EPHA3 | G | 2.29119E-10 | 0.368817 | 3.49803E-07 |
| rs1512183 | 3 | 89505054 | GTEx/v8 | Pancreas | EPHA3 | T | 2.32543E-10 | 0.364957 | 3.49803E-07 |
| rs17026944 | 3 | 89521131 | GTEx/v8 | Pancreas | EPHA3 | A | 2.32543E-10 | 0.364957 | 3.49803E-07 |
| rs73139125 | 3 | 89524056 | GTEx/v8 | Pancreas | EPHA3 | G | 2.32543E-10 | 0.364957 | 3.49803E-07 |
| rs2048519 | 3 | 89533978 | GTEx/v8 | Pancreas | EPHA3 | A | 2.81579E-10 | 0.37063 | 3.49803E-07 |
| rs3762718 | 3 | 89535066 | GTEx/v8 | Pancreas | EPHA3 | G | 2.81579E-10 | 0.37063 | 3.49803E-07 |
| rs2196083 | 3 | 89536546 | GTEx/v8 | Pancreas | EPHA3 | C | 2.81579E-10 | 0.37063 | 3.49803E-07 |
| rs12634044 | 3 | 89537651 | GTEx/v8 | Pancreas | EPHA3 | G | 2.81579E-10 | 0.37063 | 3.49803E-07 |
| rs73146907 | 3 | 89538747 | GTEx/v8 | Pancreas | EPHA3 | A | 2.81579E-10 | 0.37063 | 3.49803E-07 |
| rs57902683 | 3 | 89539011 | GTEx/v8 | Pancreas | EPHA3 | CA | 2.81579E-10 | 0.37063 | 3.49803E-07 |
| rs55979407 | 3 | 89539275 | GTEx/v8 | Pancreas | EPHA3 | G | 2.81579E-10 | 0.37063 | 3.49803E-07 |
| rs113675293 | 3 | 89539536 | GTEx/v8 | Pancreas | EPHA3 | T | 2.81579E-10 | 0.37063 | 3.49803E-07 |
| rs73146926 | 3 | 89542293 | GTEx/v8 | Pancreas | EPHA3 | G | 2.81579E-10 | 0.37063 | 3.49803E-07 |
| rs4035593 | 3 | 89542601 | GTEx/v8 | Pancreas | EPHA3 | TAA | 2.81579E-10 | 0.37063 | 3.49803E-07 |
| rs2117136 | 3 | 89542699 | GTEx/v8 | Pancreas | EPHA3 | G | 2.81579E-10 | 0.37063 | 3.49803E-07 |
| rs7633599 | 3 | 89544591 | GTEx/v8 | Pancreas | EPHA3 | A | 2.81579E-10 | 0.37063 | 3.49803E-07 |
| rs7613315 | 3 | 89544773 | GTEx/v8 | Pancreas | EPHA3 | C | 2.81579E-10 | 0.37063 | 3.49803E-07 |
| rs1028012 | 3 | 89549422 | GTEx/v8 | Pancreas | EPHA3 | T | 2.81579E-10 | 0.37063 | 3.49803E-07 |
| rs17745025 | 3 | 89549919 | GTEx/v8 | Pancreas | EPHA3 | C | 2.81579E-10 | 0.37063 | 3.49803E-07 |
| rs73146961 | 3 | 89550459 | GTEx/v8 | Pancreas | EPHA3 | C | 2.81579E-10 | 0.37063 | 3.49803E-07 |
| rs12634697 | 3 | 89550993 | GTEx/v8 | Pancreas | EPHA3 | A | 2.81579E-10 | 0.37063 | 3.49803E-07 |
| rs12638168 | 3 | 89551544 | GTEx/v8 | Pancreas | EPHA3 | C | 2.81579E-10 | 0.37063 | 3.49803E-07 |
| rs73146970 | 3 | 89553168 | GTEx/v8 | Pancreas | EPHA3 | G | 2.81579E-10 | 0.37063 | 3.49803E-07 |
| rs12629758 | 3 | 89556408 | GTEx/v8 | Pancreas | EPHA3 | G | 2.81579E-10 | 0.37063 | 3.49803E-07 |
| rs1039990 | 3 | 89558086 | GTEx/v8 | Pancreas | EPHA3 | A | 2.81579E-10 | 0.37063 | 3.49803E-07 |
| rs73146988 | 3 | 89561044 | GTEx/v8 | Pancreas | EPHA3 | C | 2.81579E-10 | 0.37063 | 3.49803E-07 |
| rs73146992 | 3 | 89562178 | GTEx/v8 | Pancreas | EPHA3 | G | 2.81579E-10 | 0.37063 | 3.49803E-07 |
| rs73146995 | 3 | 89562209 | GTEx/v8 | Pancreas | EPHA3 | C | 2.81579E-10 | 0.37063 | 3.49803E-07 |
| rs73147002 | 3 | 89565128 | GTEx/v8 | Pancreas | EPHA3 | G | 2.81579E-10 | 0.37063 | 3.49803E-07 |
| rs148078210 | 3 | 89565641 | GTEx/v8 | Pancreas | EPHA3 | GGTAT | 2.81579E-10 | 0.37063 | 3.49803E-07 |
| rs12633325 | 3 | 89566836 | GTEx/v8 | Pancreas | EPHA3 | C | 2.81579E-10 | 0.37063 | 3.49803E-07 |
| rs73149113 | 3 | 89567867 | GTEx/v8 | Pancreas | EPHA3 | T | 2.81579E-10 | 0.37063 | 3.49803E-07 |
| rs73137381 | 3 | 89458550 | GTEx/v8 | Pancreas | EPHA3 | G | 3.0069E-10 | 0.379833 | 3.49803E-07 |
| rs73137382 | 3 | 89461363 | GTEx/v8 | Pancreas | EPHA3 | G | 3.0069E-10 | 0.379833 | 3.49803E-07 |
| rs73137393 | 3 | 89471701 | GTEx/v8 | Pancreas | EPHA3 | C | 3.0069E-10 | 0.379833 | 3.49803E-07 |
| rs73137387 | 3 | 89464194 | GTEx/v8 | Pancreas | EPHA3 | C | 3.01355E-10 | 0.378092 | 3.49803E-07 |
| rs12639506 | 3 | 89480265 | GTEx/v8 | Pancreas | EPHA3 | G | 3.01355E-10 | 0.378092 | 3.49803E-07 |

| | | | | | | | | | |
|---|---|---|---|---|---|---|---|---|---|
| rs12631313 | 3 | 89528461 | GTEx/v8 | Pancreas | EPHA3 | T | 3.17901E-10 | 0.363797 | 3.49803E-07 |
| rs7650466 | 3 | 89530358 | GTEx/v8 | Pancreas | EPHA3 | T | 3.17901E-10 | 0.363797 | 3.49803E-07 |
| rs2117138 | 3 | 89531522 | GTEx/v8 | Pancreas | EPHA3 | G | 3.17901E-10 | 0.363797 | 3.49803E-07 |
| rs12634269 | 3 | 89532768 | GTEx/v8 | Pancreas | EPHA3 | C | 3.23497E-10 | 0.359939 | 3.49803E-07 |
| rs73146965 | 3 | 89552553 | GTEx/v8 | Pancreas | EPHA3 | G | 3.46782E-10 | 0.359264 | 3.49803E-07 |
| rs73145304 | 3 | 89568989 | GTEx/v8 | Pancreas | EPHA3 | T | 3.558E-10 | 0.35898 | 3.49803E-07 |
| rs73145310 | 3 | 89578097 | GTEx/v8 | Pancreas | EPHA3 | C | 3.558E-10 | 0.35898 | 3.49803E-07 |
| rs73145312 | 3 | 89578575 | GTEx/v8 | Pancreas | EPHA3 | T | 3.558E-10 | 0.35898 | 3.49803E-07 |
| rs73145320 | 3 | 89586535 | GTEx/v8 | Pancreas | EPHA3 | C | 3.558E-10 | 0.35898 | 3.49803E-07 |
| rs1157607 | 3 | 89506364 | GTEx/v8 | Pancreas | EPHA3 | T | 4.14668E-10 | 0.372333 | 3.49803E-07 |
| rs7626735 | 3 | 89506958 | GTEx/v8 | Pancreas | EPHA3 | A | 4.14668E-10 | 0.372333 | 3.49803E-07 |
| rs568140553 | 3 | 89507219 | GTEx/v8 | Pancreas | EPHA3 | CT | 4.14668E-10 | 0.372333 | 3.49803E-07 |
| rs6795074 | 3 | 89516652 | GTEx/v8 | Pancreas | EPHA3 | C | 4.14668E-10 | 0.372333 | 3.49803E-07 |
| rs73141108 | 3 | 89517119 | GTEx/v8 | Pancreas | EPHA3 | T | 4.14668E-10 | 0.372333 | 3.49803E-07 |
| rs1912966 | 3 | 89518464 | GTEx/v8 | Pancreas | EPHA3 | A | 4.14668E-10 | 0.372333 | 3.49803E-07 |
| rs73139104 | 3 | 89519229 | GTEx/v8 | Pancreas | EPHA3 | C | 4.14668E-10 | 0.372333 | 3.49803E-07 |
| 3:89519266_GA_G | 3 | 89519266 | GTEx/v8 | Pancreas | EPHA3 | G | 4.14668E-10 | 0.372333 | 3.49803E-07 |
| 3:89521060_CTAT_C | 3 | 89521060 | GTEx/v8 | Pancreas | EPHA3 | C | 4.14668E-10 | 0.372333 | 3.49803E-07 |
| rs73139110 | 3 | 89521509 | GTEx/v8 | Pancreas | EPHA3 | A | 4.14668E-10 | 0.372333 | 3.49803E-07 |
| rs1054750 | 3 | 89521725 | GTEx/v8 | Pancreas | EPHA3 | C | 4.14668E-10 | 0.372333 | 3.49803E-07 |
| rs73139115 | 3 | 89521900 | GTEx/v8 | Pancreas | EPHA3 | T | 4.14668E-10 | 0.372333 | 3.49803E-07 |
| rs73139117 | 3 | 89521965 | GTEx/v8 | Pancreas | EPHA3 | G | 4.14668E-10 | 0.372333 | 3.49803E-07 |
| rs73139118 | 3 | 89522263 | GTEx/v8 | Pancreas | EPHA3 | T | 4.14668E-10 | 0.372333 | 3.49803E-07 |
| rs150837250 | 3 | 89522314 | GTEx/v8 | Pancreas | EPHA3 | AGTTATAATTAG | 4.14668E-10 | 0.372333 | 3.49803E-07 |
| rs17801380 | 3 | 89522483 | GTEx/v8 | Pancreas | EPHA3 | T | 4.14668E-10 | 0.372333 | 3.49803E-07 |
| rs73139127 | 3 | 89524436 | GTEx/v8 | Pancreas | EPHA3 | A | 4.14668E-10 | 0.372333 | 3.49803E-07 |
| rs1499780 | 3 | 89526072 | GTEx/v8 | Pancreas | EPHA3 | C | 4.14668E-10 | 0.372333 | 3.49803E-07 |
| rs73145305 | 3 | 89570713 | GTEx/v8 | Pancreas | EPHA3 | T | 4.85418E-10 | 0.370693 | 3.49803E-07 |
| 3:89571180_TAGAG_T | 3 | 89571180 | GTEx/v8 | Pancreas | EPHA3 | T | 4.85418E-10 | 0.370693 | 3.49803E-07 |
| rs907548 | 3 | 89574852 | GTEx/v8 | Pancreas | EPHA3 | T | 4.85418E-10 | 0.370693 | 3.49803E-07 |
| rs73145307 | 3 | 89576040 | GTEx/v8 | Pancreas | EPHA3 | A | 4.85418E-10 | 0.370693 | 3.49803E-07 |
| rs73145309 | 3 | 89576389 | GTEx/v8 | Pancreas | EPHA3 | T | 4.85418E-10 | 0.370693 | 3.49803E-07 |
| rs73145316 | 3 | 89580407 | GTEx/v8 | Pancreas | EPHA3 | T | 4.85418E-10 | 0.370693 | 3.49803E-07 |
| rs73145332 | 3 | 89589327 | GTEx/v8 | Pancreas | EPHA3 | C | 4.85418E-10 | 0.370693 | 3.49803E-07 |
| rs112087049 | 3 | 89509550 | GTEx/v8 | Pancreas | EPHA3 | T | 4.96309E-10 | 0.397999 | 3.49803E-07 |
| rs73139281 | 3 | 89503890 | GTEx/v8 | Pancreas | EPHA3 | T | 5.01652E-10 | 0.365232 | 3.49803E-07 |
| rs73139284 | 3 | 89503958 | GTEx/v8 | Pancreas | EPHA3 | G | 5.01652E-10 | 0.365232 | 3.49803E-07 |
| rs73139286 | 3 | 89504295 | GTEx/v8 | Pancreas | EPHA3 | A | 5.01652E-10 | 0.365232 | 3.49803E-07 |
| rs73149107 | 3 | 89565849 | GTEx/v8 | Pancreas | EPHA3 | A | 5.51674E-10 | 0.364495 | 3.49803E-07 |
| rs73153260 | 3 | 89718078 | GTEx/v8 | Pancreas | EPHA3 | A | 5.52434E-10 | 0.429495 | 3.49803E-07 |
| rs73139134 | 3 | 89527709 | GTEx/v8 | Pancreas | EPHA3 | C | 5.82105E-10 | 0.366907 | 3.49803E-07 |
| rs73139135 | 3 | 89527937 | GTEx/v8 | Pancreas | EPHA3 | C | 5.82105E-10 | 0.366907 | 3.49803E-07 |
| rs73139139 | 3 | 89528313 | GTEx/v8 | Pancreas | EPHA3 | G | 5.82105E-10 | 0.366907 | 3.49803E-07 |
| rs12636710 | 3 | 89528477 | GTEx/v8 | Pancreas | EPHA3 | A | 5.82105E-10 | 0.366907 | 3.49803E-07 |
| rs3762717 | 3 | 89528948 | GTEx/v8 | Pancreas | EPHA3 | C | 5.82105E-10 | 0.366907 | 3.49803E-07 |
| rs73139147 | 3 | 89530347 | GTEx/v8 | Pancreas | EPHA3 | A | 5.82105E-10 | 0.366907 | 3.49803E-07 |
| rs73139148 | 3 | 89530956 | GTEx/v8 | Pancreas | EPHA3 | A | 5.82105E-10 | 0.366907 | 3.49803E-07 |
| rs12629165 | 3 | 89533017 | GTEx/v8 | Pancreas | EPHA3 | A | 5.82105E-10 | 0.366907 | 3.49803E-07 |
| rs752364173 | 3 | 89521820 | GTEx/v8 | Pancreas | EPHA3 | T | 6.57419E-10 | 0.352038 | 3.49803E-07 |
| 3:89521831_TGAAAG_T | 3 | 89521831 | GTEx/v8 | Pancreas | EPHA3 | T | 6.57419E-10 | 0.352038 | 3.49803E-07 |
| 3:89521838_TTTG_T | 3 | 89521838 | GTEx/v8 | Pancreas | EPHA3 | T | 6.57419E-10 | 0.352038 | 3.49803E-07 |

| | | | | | | | | | |
|---|---|---|---|---|---|---|---|---|---|
| 3:89544326_CGGTGGAAATCTCCTTATTAAATTGTGGGCAAGAGGCAACCCAT_C | 3 | 89544326 | GTEx/v8 | Pancreas | EPHA3 | C | 7.02079E-10 | 0.363859 | 3.49803E-07 |
| rs7619303 | 3 | 89552554 | GTEx/v8 | Pancreas | EPHA3 | A | 7.96E-10 | 0.346987 | 3.49803E-07 |
| rs73149109 | 3 | 89567344 | GTEx/v8 | Pancreas | EPHA3 | A | 8.72692E-10 | 0.347768 | 3.49803E-07 |
| rs1080592 | 3 | 89567568 | GTEx/v8 | Pancreas | EPHA3 | C | 8.72692E-10 | 0.347768 | 3.49803E-07 |
| rs12633626 | 3 | 89744949 | GTEx/v8 | Pancreas | EPHA3 | A | 9.26126E-10 | 0.377684 | 3.49803E-07 |
| 3:89694510_GATA_G | 3 | 89694510 | GTEx/v8 | Artery Tibial | EPHA3 | G | 1.01112E-09 | -0.175842 | 7.80657E-07 |
| 3:89626776_AT_A | 3 | 89626776 | GTEx/v8 | Pancreas | EPHA3 | A | 1.1209E-09 | 0.36937 | 3.49803E-07 |
| rs73139144 | 3 | 89529177 | GTEx/v8 | Pancreas | EPHA3 | A | 1.27678E-09 | 0.360298 | 3.49803E-07 |
| rs12634341 | 3 | 89533031 | GTEx/v8 | Pancreas | EPHA3 | C | 1.33245E-09 | 0.355856 | 3.49803E-07 |
| rs2346840 | 3 | 89533267 | GTEx/v8 | Pancreas | EPHA3 | G | 1.33245E-09 | 0.355856 | 3.49803E-07 |
| rs73153266 | 3 | 89720111 | GTEx/v8 | Pancreas | EPHA3 | G | 1.69953E-09 | 0.360744 | 3.49803E-07 |
| rs66708608 | 3 | 89486373 | GTEx/v8 | Pancreas | EPHA3 | T | 1.80707E-09 | 0.343463 | 3.49803E-07 |
| rs73145345 | 3 | 89598696 | GTEx/v8 | Pancreas | EPHA3 | A | 2.0732E-09 | 0.365608 | 3.49803E-07 |
| rs111835258 | 3 | 89601496 | GTEx/v8 | Pancreas | EPHA3 | A | 2.0732E-09 | 0.365608 | 3.49803E-07 |
| rs73145351 | 3 | 89607237 | GTEx/v8 | Pancreas | EPHA3 | T | 2.0732E-09 | 0.365608 | 3.49803E-07 |
| rs73145353 | 3 | 89607824 | GTEx/v8 | Pancreas | EPHA3 | T | 2.0732E-09 | 0.365608 | 3.49803E-07 |
| rs73145357 | 3 | 89611766 | GTEx/v8 | Pancreas | EPHA3 | C | 2.0732E-09 | 0.365608 | 3.49803E-07 |
| rs819293 | 3 | 89613664 | GTEx/v8 | Pancreas | EPHA3 | A | 2.07428E-09 | 0.356596 | 3.49803E-07 |
| rs73153262 | 3 | 89719269 | GTEx/v8 | Pancreas | EPHA3 | T | 2.21545E-09 | 0.362082 | 3.49803E-07 |
| rs113288647 | 3 | 89720352 | GTEx/v8 | Pancreas | EPHA3 | T | 2.21545E-09 | 0.362082 | 3.49803E-07 |
| rs2069183 | 3 | 89645017 | GTEx/v8 | Pancreas | EPHA3 | A | 2.26183E-09 | 0.357285 | 3.49803E-07 |
| rs73153241 | 3 | 89695319 | GTEx/v8 | Pancreas | EPHA3 | G | 2.4658E-09 | 0.355974 | 3.49803E-07 |
| rs73153252 | 3 | 89712192 | GTEx/v8 | Pancreas | EPHA3 | T | 3.18359E-09 | 0.354961 | 3.49803E-07 |
| rs7617395 | 3 | 89722541 | GTEx/v8 | Pancreas | EPHA3 | T | 3.18359E-09 | 0.354961 | 3.49803E-07 |
| rs73153218 | 3 | 89619808 | GTEx/v8 | Pancreas | EPHA3 | A | 3.88508E-09 | 0.357171 | 3.49803E-07 |
| rs73153258 | 3 | 89715585 | GTEx/v8 | Pancreas | EPHA3 | A | 4.43857E-09 | 0.356589 | 3.49803E-07 |
| rs73137384 | 3 | 89463290 | GTEx/v8 | Pancreas | EPHA3 | T | 5.37938E-09 | 0.332038 | 3.49803E-07 |
| rs6773817 | 3 | 89753400 | GTEx/v8 | Artery Tibial | EPHA3 | A | 5.93098E-09 | -0.168368 | 7.80657E-07 |
| rs73137395 | 3 | 89478753 | GTEx/v8 | Pancreas | EPHA3 | C | 6.16802E-09 | 0.343206 | 3.49803E-07 |
| rs73139257 | 3 | 89482779 | GTEx/v8 | Pancreas | EPHA3 | G | 6.16802E-09 | 0.343206 | 3.49803E-07 |
| rs13064615 | 3 | 89799433 | GTEx/v8 | Artery Tibial | EPHA3 | G | 7.12193E-09 | -0.168807 | 7.80657E-07 |
| rs112149405 | 3 | 89886780 | GTEx/v8 | Pancreas | EPHA3 | C | 8.46419E-09 | 0.408213 | 3.49803E-07 |
| rs73153282 | 3 | 89728408 | GTEx/v8 | Pancreas | EPHA3 | T | 9.61374E-09 | 0.344833 | 3.49803E-07 |
| rs73151380 | 3 | 90000532 | GTEx/v8 | Pancreas | EPHA3 | T | 9.80386E-09 | 0.393419 | 3.49803E-07 |
| rs73153279 | 3 | 89727968 | GTEx/v8 | Pancreas | EPHA3 | C | 1.0246E-08 | 0.342672 | 3.49803E-07 |
| rs73153283 | 3 | 89732193 | GTEx/v8 | Pancreas | EPHA3 | T | 1.0246E-08 | 0.342672 | 3.49803E-07 |
| rs1284258 | 3 | 89823487 | GTEx/v8 | Artery Tibial | EPHA3 | T | 1.04495E-08 | -0.16652 | 7.80657E-07 |
| rs112604674 | 3 | 89989681 | GTEx/v8 | Pancreas | EPHA3 | T | 1.15883E-08 | 0.392034 | 3.49803E-07 |
| rs1512185 | 3 | 89473615 | GTEx/v8 | Pancreas | EPHA3 | A | 1.17088E-08 | 0.324867 | 3.49803E-07 |
| rs73153293 | 3 | 89756242 | GTEx/v8 | Pancreas | EPHA3 | T | 1.51089E-08 | 0.368874 | 3.49803E-07 |
| rs6551412 | 3 | 89496502 | GTEx/v8 | Pancreas | EPHA3 | A | 2.13147E-08 | 0.340301 | 3.49803E-07 |
| rs4264747 | 3 | 89882102 | GTEx/v8 | Artery Tibial | EPHA3 | G | 2.39483E-08 | -0.162136 | 7.80657E-07 |
| rs1879650 | 3 | 89856887 | GTEx/v8 | Artery Tibial | EPHA3 | T | 2.83016E-08 | -0.161113 | 7.80657E-07 |
| rs7616322 | 3 | 89493283 | GTEx/v8 | Pancreas | EPHA3 | T | 3.30748E-08 | 0.319057 | 3.49803E-07 |
| rs13062221 | 3 | 89778009 | GTEx/v8 | Artery Tibial | EPHA3 | C | 3.38194E-08 | -0.160623 | 7.80657E-07 |
| rs112407114 | 3 | 89797398 | GTEx/v8 | Pancreas | EPHA3 | C | 4.05413E-08 | 0.351939 | 3.49803E-07 |
| rs2196083 | 3 | 89536546 | GTEx/v8 | Brain Cerebellum | EPHA3 | C | 5.10725E-08 | -0.379543 | 0.000131411 |
| rs2048519 | 3 | 89533978 | GTEx/v8 | Brain Cerebellum | EPHA3 | A | 5.20018E-08 | -0.380424 | 0.000131411 |
| rs3762718 | 3 | 89535066 | GTEx/v8 | Brain Cerebellum | EPHA3 | G | 5.20018E-08 | -0.380424 | 0.000131411 |
| rs12634044 | 3 | 89537651 | GTEx/v8 | Brain Cerebellum | EPHA3 | G | 5.20018E-08 | -0.380424 | 0.000131411 |

| | | | | | | | | | |
|---|---|---|---|---|---|---|---|---|---|
| rs73146907 | 3 | 89538747 | GTEx/v8 | Brain Cerebellum | EPHA3 | A | 5.20018E-08 | -0.380424 | 0.000131411 |
| rs57902683 | 3 | 89539011 | GTEx/v8 | Brain Cerebellum | EPHA3 | CA | 5.20018E-08 | -0.380424 | 0.000131411 |
| rs55979407 | 3 | 89539275 | GTEx/v8 | Brain Cerebellum | EPHA3 | G | 5.20018E-08 | -0.380424 | 0.000131411 |
| rs113675293 | 3 | 89539536 | GTEx/v8 | Brain Cerebellum | EPHA3 | T | 5.20018E-08 | -0.380424 | 0.000131411 |
| rs4857500 | 3 | 89541486 | GTEx/v8 | Brain Cerebellum | EPHA3 | C | 5.20018E-08 | -0.380424 | 0.000131411 |
| rs73146926 | 3 | 89542293 | GTEx/v8 | Brain Cerebellum | EPHA3 | G | 5.20018E-08 | -0.380424 | 0.000131411 |
| rs4035593 | 3 | 89542601 | GTEx/v8 | Brain Cerebellum | EPHA3 | TAA | 5.20018E-08 | -0.380424 | 0.000131411 |
| rs2117136 | 3 | 89542699 | GTEx/v8 | Brain Cerebellum | EPHA3 | G | 5.20018E-08 | -0.380424 | 0.000131411 |
| 3:89544326_CGGTGGAAATCTCCTTATTAAATTGTGGGCAAGAGGCAACCCAT_C | 3 | 89544326 | GTEx/v8 | Brain Cerebellum | EPHA3 | C | 5.20018E-08 | -0.380424 | 0.000131411 |
| rs7633599 | 3 | 89544591 | GTEx/v8 | Brain Cerebellum | EPHA3 | A | 5.20018E-08 | -0.380424 | 0.000131411 |
| rs7613315 | 3 | 89544773 | GTEx/v8 | Brain Cerebellum | EPHA3 | C | 5.20018E-08 | -0.380424 | 0.000131411 |
| rs1028012 | 3 | 89549422 | GTEx/v8 | Brain Cerebellum | EPHA3 | T | 5.20018E-08 | -0.380424 | 0.000131411 |
| rs17745025 | 3 | 89549919 | GTEx/v8 | Brain Cerebellum | EPHA3 | C | 5.20018E-08 | -0.380424 | 0.000131411 |
| rs73146961 | 3 | 89550459 | GTEx/v8 | Brain Cerebellum | EPHA3 | C | 5.20018E-08 | -0.380424 | 0.000131411 |
| rs12634697 | 3 | 89550993 | GTEx/v8 | Brain Cerebellum | EPHA3 | A | 5.20018E-08 | -0.380424 | 0.000131411 |
| rs12638168 | 3 | 89551544 | GTEx/v8 | Brain Cerebellum | EPHA3 | C | 5.20018E-08 | -0.380424 | 0.000131411 |
| rs73146970 | 3 | 89553168 | GTEx/v8 | Brain Cerebellum | EPHA3 | G | 5.20018E-08 | -0.380424 | 0.000131411 |
| rs12629758 | 3 | 89556408 | GTEx/v8 | Brain Cerebellum | EPHA3 | G | 5.20018E-08 | -0.380424 | 0.000131411 |
| rs1039990 | 3 | 89558086 | GTEx/v8 | Brain Cerebellum | EPHA3 | A | 5.20018E-08 | -0.380424 | 0.000131411 |
| rs73146988 | 3 | 89561044 | GTEx/v8 | Brain Cerebellum | EPHA3 | C | 5.20018E-08 | -0.380424 | 0.000131411 |
| rs73146992 | 3 | 89562178 | GTEx/v8 | Brain Cerebellum | EPHA3 | G | 5.20018E-08 | -0.380424 | 0.000131411 |
| rs73146995 | 3 | 89562209 | GTEx/v8 | Brain Cerebellum | EPHA3 | C | 5.20018E-08 | -0.380424 | 0.000131411 |
| rs73147002 | 3 | 89565128 | GTEx/v8 | Brain Cerebellum | EPHA3 | G | 5.20018E-08 | -0.380424 | 0.000131411 |
| rs148078210 | 3 | 89565641 | GTEx/v8 | Brain Cerebellum | EPHA3 | GGTAT | 5.20018E-08 | -0.380424 | 0.000131411 |
| rs73149103 | 3 | 89565651 | GTEx/v8 | Brain Cerebellum | EPHA3 | A | 5.20018E-08 | -0.380424 | 0.000131411 |
| rs73149107 | 3 | 89565849 | GTEx/v8 | Brain Cerebellum | EPHA3 | A | 5.20018E-08 | -0.380424 | 0.000131411 |
| rs12633325 | 3 | 89566836 | GTEx/v8 | Brain Cerebellum | EPHA3 | C | 5.20018E-08 | -0.380424 | 0.000131411 |
| rs73149113 | 3 | 89567867 | GTEx/v8 | Brain Cerebellum | EPHA3 | T | 5.20018E-08 | -0.380424 | 0.000131411 |
| rs73145305 | 3 | 89570713 | GTEx/v8 | Brain Cerebellum | EPHA3 | T | 5.20018E-08 | -0.380424 | 0.000131411 |
| 3:89571180_TAGAG_T | 3 | 89571180 | GTEx/v8 | Brain Cerebellum | EPHA3 | T | 5.20018E-08 | -0.380424 | 0.000131411 |
| rs907548 | 3 | 89574852 | GTEx/v8 | Brain Cerebellum | EPHA3 | T | 5.20018E-08 | -0.380424 | 0.000131411 |
| rs73145307 | 3 | 89576040 | GTEx/v8 | Brain Cerebellum | EPHA3 | A | 5.20018E-08 | -0.380424 | 0.000131411 |
| rs73145309 | 3 | 89576389 | GTEx/v8 | Brain Cerebellum | EPHA3 | T | 5.20018E-08 | -0.380424 | 0.000131411 |
| rs73145316 | 3 | 89580407 | GTEx/v8 | Brain Cerebellum | EPHA3 | T | 5.20018E-08 | -0.380424 | 0.000131411 |
| rs12637036 | 3 | 89584170 | GTEx/v8 | Brain Cerebellum | EPHA3 | A | 5.20018E-08 | -0.380424 | 0.000131411 |
| rs73145332 | 3 | 89589327 | GTEx/v8 | Brain Cerebellum | EPHA3 | C | 5.20018E-08 | -0.380424 | 0.000131411 |
| rs73145336 | 3 | 89593317 | GTEx/v8 | Brain Cerebellum | EPHA3 | G | 5.20018E-08 | -0.380424 | 0.000131411 |
| rs2346840 | 3 | 89533267 | GTEx/v8 | Brain Cerebellum | EPHA3 | G | 5.26884E-08 | -0.389826 | 0.000131411 |
| rs73139276 | 3 | 89500196 | GTEx/v8 | Brain Cerebellum | EPHA3 | A | 5.55208E-08 | -0.397101 | 0.000131411 |
| rs73139288 | 3 | 89505327 | GTEx/v8 | Brain Cerebellum | EPHA3 | G | 5.55208E-08 | -0.397101 | 0.000131411 |
| rs1157607 | 3 | 89506364 | GTEx/v8 | Brain Cerebellum | EPHA3 | T | 5.55208E-08 | -0.397101 | 0.000131411 |

| | | | | | | | | | |
|---|---|---|---|---|---|---|---|---|---|
| rs7626735 | 3 | 89506958 | GTEx/v8 | Brain Cerebellum | EPHA3 | A | 5.55208E-08 | -0.397101 | 0.000131411 |
| rs568140553 | 3 | 89507219 | GTEx/v8 | Brain Cerebellum | EPHA3 | CT | 5.55208E-08 | -0.397101 | 0.000131411 |
| rs6795074 | 3 | 89516652 | GTEx/v8 | Brain Cerebellum | EPHA3 | C | 5.55208E-08 | -0.397101 | 0.000131411 |
| rs73141108 | 3 | 89517119 | GTEx/v8 | Brain Cerebellum | EPHA3 | T | 5.55208E-08 | -0.397101 | 0.000131411 |
| rs1912966 | 3 | 89518464 | GTEx/v8 | Brain Cerebellum | EPHA3 | A | 5.55208E-08 | -0.397101 | 0.000131411 |
| rs73139104 | 3 | 89519229 | GTEx/v8 | Brain Cerebellum | EPHA3 | C | 5.55208E-08 | -0.397101 | 0.000131411 |
| 3:89519266_GA_G | 3 | 89519266 | GTEx/v8 | Brain Cerebellum | EPHA3 | G | 5.55208E-08 | -0.397101 | 0.000131411 |
| 3:89521060_CTAT_C | 3 | 89521060 | GTEx/v8 | Brain Cerebellum | EPHA3 | C | 5.55208E-08 | -0.397101 | 0.000131411 |
| rs73139110 | 3 | 89521509 | GTEx/v8 | Brain Cerebellum | EPHA3 | A | 5.55208E-08 | -0.397101 | 0.000131411 |
| rs1054750 | 3 | 89521725 | GTEx/v8 | Brain Cerebellum | EPHA3 | C | 5.55208E-08 | -0.397101 | 0.000131411 |
| rs73139115 | 3 | 89521900 | GTEx/v8 | Brain Cerebellum | EPHA3 | T | 5.55208E-08 | -0.397101 | 0.000131411 |
| rs73139117 | 3 | 89521965 | GTEx/v8 | Brain Cerebellum | EPHA3 | G | 5.55208E-08 | -0.397101 | 0.000131411 |
| rs73139118 | 3 | 89522263 | GTEx/v8 | Brain Cerebellum | EPHA3 | T | 5.55208E-08 | -0.397101 | 0.000131411 |
| rs150837250 | 3 | 89522314 | GTEx/v8 | Brain Cerebellum | EPHA3 | AGTTATAATTAG | 5.55208E-08 | -0.397101 | 0.000131411 |
| rs17801380 | 3 | 89522483 | GTEx/v8 | Brain Cerebellum | EPHA3 | T | 5.55208E-08 | -0.397101 | 0.000131411 |
| rs73139127 | 3 | 89524436 | GTEx/v8 | Brain Cerebellum | EPHA3 | A | 5.55208E-08 | -0.397101 | 0.000131411 |
| rs73139129 | 3 | 89524562 | GTEx/v8 | Brain Cerebellum | EPHA3 | G | 5.55208E-08 | -0.397101 | 0.000131411 |
| rs1499780 | 3 | 89526072 | GTEx/v8 | Brain Cerebellum | EPHA3 | C | 5.55208E-08 | -0.397101 | 0.000131411 |
| rs73137381 | 3 | 89458550 | GTEx/v8 | Brain Cerebellum | EPHA3 | G | 7.06753E-08 | -0.399375 | 0.000131411 |
| rs73137382 | 3 | 89461363 | GTEx/v8 | Brain Cerebellum | EPHA3 | G | 7.06753E-08 | -0.399375 | 0.000131411 |
| rs73137387 | 3 | 89464194 | GTEx/v8 | Brain Cerebellum | EPHA3 | C | 7.06753E-08 | -0.399375 | 0.000131411 |
| rs73137393 | 3 | 89471701 | GTEx/v8 | Brain Cerebellum | EPHA3 | C | 7.06753E-08 | -0.399375 | 0.000131411 |
| rs12639506 | 3 | 89480265 | GTEx/v8 | Brain Cerebellum | EPHA3 | G | 7.06753E-08 | -0.399375 | 0.000131411 |
| rs112087049 | 3 | 89509550 | GTEx/v8 | Brain Cerebellum | EPHA3 | T | 7.14504E-08 | -0.411656 | 0.000131411 |
| rs144317613 | 3 | 89525080 | GTEx/v8 | Brain Cerebellum | EPHA3 | TA | 7.4538E-08 | -0.396948 | 0.000131411 |
| rs1512189 | 3 | 89546332 | GTEx/v8 | Brain Cerebellum | EPHA3 | G | 7.98666E-08 | -0.377125 | 0.000131411 |
| rs6784577 | 3 | 89564308 | GTEx/v8 | Brain Cerebellum | EPHA3 | G | 7.98666E-08 | -0.377125 | 0.000131411 |
| rs56082199 | 3 | 89586973 | GTEx/v8 | Brain Cerebellum | EPHA3 | G | 7.98666E-08 | -0.377125 | 0.000131411 |
| rs12629165 | 3 | 89533017 | GTEx/v8 | Brain Cerebellum | EPHA3 | A | 8.19033E-08 | -0.386321 | 0.000131411 |
| rs12634341 | 3 | 89533031 | GTEx/v8 | Brain Cerebellum | EPHA3 | C | 8.19033E-08 | -0.386321 | 0.000131411 |
| rs73153218 | 3 | 89619808 | GTEx/v8 | Artery Tibial | EPHA3 | A | 8.97526E-08 | 0.177382 | 7.80657E-07 |
| rs148078210 | 3 | 89565641 | GTEx/v8 | Artery Tibial | EPHA3 | GGTAT | 1.02345E-07 | 0.171217 | 7.80657E-07 |
| rs73154925 | 3 | 89859312 | GTEx/v8 | Pancreas | EPHA3 | C | 1.30179E-07 | 0.328276 | 3.49803E-07 |
| rs972030 | 3 | 89538498 | GTEx/v8 | Brain Cerebellum | EPHA3 | T | 1.33265E-07 | -0.369879 | 0.000131411 |
| rs11128078 | 3 | 90030538 | GTEx/v8 | Artery Tibial | EPHA3 | T | 1.43479E-07 | -0.157932 | 7.80657E-07 |
| rs12634341 | 3 | 89533031 | GTEx/v8 | Artery Tibial | EPHA3 | C | 1.51111E-07 | 0.169578 | 7.80657E-07 |
| rs73139133 | 3 | 89527708 | GTEx/v8 | Brain Cerebellum | EPHA3 | A | 1.51833E-07 | -0.378354 | 0.000131411 |
| rs73139134 | 3 | 89527709 | GTEx/v8 | Brain Cerebellum | EPHA3 | C | 1.51833E-07 | -0.378354 | 0.000131411 |
| rs73139135 | 3 | 89527937 | GTEx/v8 | Brain Cerebellum | EPHA3 | C | 1.51833E-07 | -0.378354 | 0.000131411 |
| rs73139139 | 3 | 89528313 | GTEx/v8 | Brain Cerebellum | EPHA3 | G | 1.51833E-07 | -0.378354 | 0.000131411 |
| rs12636710 | 3 | 89528477 | GTEx/v8 | Brain Cerebellum | EPHA3 | A | 1.51833E-07 | -0.378354 | 0.000131411 |
| rs3762717 | 3 | 89528948 | GTEx/v8 | Brain Cerebellum | EPHA3 | C | 1.51833E-07 | -0.378354 | 0.000131411 |
| rs73139144 | 3 | 89529177 | GTEx/v8 | Brain Cerebellum | EPHA3 | A | 1.51833E-07 | -0.378354 | 0.000131411 |
| rs73139147 | 3 | 89530347 | GTEx/v8 | Brain Cerebellum | EPHA3 | A | 1.51833E-07 | -0.378354 | 0.000131411 |

| | | | | | | | | | |
|---|---|---|---|---|---|---|---|---|---|
| rs73139148 | 3 | 89530956 | GTEx/v8 | Brain Cerebellum | EPHA3 | A | 1.51833E-07 | -0.378354 | 0.000131411 |
| rs6551412 | 3 | 89496502 | GTEx/v8 | Brain Cerebellum | EPHA3 | A | 1.56124E-07 | -0.387105 | 0.000131411 |
| rs819293 | 3 | 89613664 | GTEx/v8 | Artery Tibial | EPHA3 | A | 1.56358E-07 | 0.170458 | 7.80657E-07 |
| rs2347959 | 3 | 90039465 | GTEx/v8 | Artery Tibial | EPHA3 | A | 1.56916E-07 | -0.156664 | 7.80657E-07 |
| rs113165984 | 3 | 89989689 | GTEx/v8 | Pancreas | EPHA3 | C | 1.6939E-07 | 0.400186 | 3.49803E-07 |
| rs73146986 | 3 | 89560884 | GTEx/v8 | Brain Cerebellum | EPHA3 | G | 1.86176E-07 | -0.371667 | 0.000131411 |
| rs12629165 | 3 | 89533017 | GTEx/v8 | Artery Tibial | EPHA3 | A | 2.12155E-07 | 0.17036 | 7.80657E-07 |
| rs73146907 | 3 | 89538747 | GTEx/v8 | Artery Tibial | EPHA3 | A | 2.33274E-07 | 0.166443 | 7.80657E-07 |
| rs57902683 | 3 | 89539011 | GTEx/v8 | Artery Tibial | EPHA3 | CA | 2.33274E-07 | 0.166443 | 7.80657E-07 |
| rs55979407 | 3 | 89539275 | GTEx/v8 | Artery Tibial | EPHA3 | G | 2.33274E-07 | 0.166443 | 7.80657E-07 |
| rs113675293 | 3 | 89539536 | GTEx/v8 | Artery Tibial | EPHA3 | T | 2.33274E-07 | 0.166443 | 7.80657E-07 |
| rs73146926 | 3 | 89542293 | GTEx/v8 | Artery Tibial | EPHA3 | G | 2.33274E-07 | 0.166443 | 7.80657E-07 |
| rs4035593 | 3 | 89542601 | GTEx/v8 | Artery Tibial | EPHA3 | TAA | 2.33274E-07 | 0.166443 | 7.80657E-07 |
| 3:89544326_CGGTGGAAATCTCCTTATTAAATTGTGGGCAAGAGGCAACCCAT_C | 3 | 89544326 | GTEx/v8 | Artery Tibial | EPHA3 | C | 2.33274E-07 | 0.166443 | 7.80657E-07 |
| rs7633599 | 3 | 89544591 | GTEx/v8 | Artery Tibial | EPHA3 | A | 2.33274E-07 | 0.166443 | 7.80657E-07 |
| rs7613315 | 3 | 89544773 | GTEx/v8 | Artery Tibial | EPHA3 | C | 2.33274E-07 | 0.166443 | 7.80657E-07 |
| rs1028012 | 3 | 89549422 | GTEx/v8 | Artery Tibial | EPHA3 | T | 2.33274E-07 | 0.166443 | 7.80657E-07 |
| rs17745025 | 3 | 89549919 | GTEx/v8 | Artery Tibial | EPHA3 | C | 2.33274E-07 | 0.166443 | 7.80657E-07 |
| rs73146961 | 3 | 89550459 | GTEx/v8 | Artery Tibial | EPHA3 | C | 2.33274E-07 | 0.166443 | 7.80657E-07 |
| rs12634697 | 3 | 89550993 | GTEx/v8 | Artery Tibial | EPHA3 | A | 2.33274E-07 | 0.166443 | 7.80657E-07 |
| rs12638168 | 3 | 89551544 | GTEx/v8 | Artery Tibial | EPHA3 | C | 2.33274E-07 | 0.166443 | 7.80657E-07 |
| rs73146970 | 3 | 89553168 | GTEx/v8 | Artery Tibial | EPHA3 | G | 2.33274E-07 | 0.166443 | 7.80657E-07 |
| rs12629758 | 3 | 89556408 | GTEx/v8 | Artery Tibial | EPHA3 | G | 2.33274E-07 | 0.166443 | 7.80657E-07 |
| rs1039990 | 3 | 89558086 | GTEx/v8 | Artery Tibial | EPHA3 | A | 2.33274E-07 | 0.166443 | 7.80657E-07 |
| rs73146988 | 3 | 89561044 | GTEx/v8 | Artery Tibial | EPHA3 | C | 2.33274E-07 | 0.166443 | 7.80657E-07 |
| rs73146992 | 3 | 89562178 | GTEx/v8 | Artery Tibial | EPHA3 | G | 2.33274E-07 | 0.166443 | 7.80657E-07 |
| rs73146995 | 3 | 89562209 | GTEx/v8 | Artery Tibial | EPHA3 | C | 2.33274E-07 | 0.166443 | 7.80657E-07 |
| rs73147002 | 3 | 89565128 | GTEx/v8 | Artery Tibial | EPHA3 | G | 2.33274E-07 | 0.166443 | 7.80657E-07 |
| rs73149107 | 3 | 89565849 | GTEx/v8 | Artery Tibial | EPHA3 | A | 2.33274E-07 | 0.166443 | 7.80657E-07 |
| rs12633325 | 3 | 89566836 | GTEx/v8 | Artery Tibial | EPHA3 | C | 2.33274E-07 | 0.166443 | 7.80657E-07 |
| rs73149113 | 3 | 89567867 | GTEx/v8 | Artery Tibial | EPHA3 | T | 2.33274E-07 | 0.166443 | 7.80657E-07 |
| rs73153218 | 3 | 89619808 | GTEx/v8 | Brain Cerebellum | EPHA3 | A | 2.3627E-07 | -0.37069 | 0.000131411 |
| rs73139261 | 3 | 89492406 | GTEx/v8 | Brain Cerebellum | EPHA3 | G | 2.49711E-07 | -0.388801 | 0.000131411 |
| rs7632427 | 3 | 89534377 | GTEx/v8 | Brain Cerebellum | EPHA3 | C | 2.51175E-07 | -0.326002 | 0.000131411 |
| rs12629999 | 3 | 89551518 | GTEx/v8 | Brain Cerebellum | EPHA3 | G | 2.55552E-07 | -0.36834 | 0.000131411 |
| rs73145303 | 3 | 89568714 | GTEx/v8 | Brain Cerebellum | EPHA3 | C | 2.55552E-07 | -0.36834 | 0.000131411 |
| rs1521806 | 3 | 90090619 | GTEx/v8 | Artery Tibial | EPHA3 | C | 2.55872E-07 | -0.151229 | 7.80657E-07 |
| rs2346840 | 3 | 89533267 | GTEx/v8 | Artery Tibial | EPHA3 | G | 2.57258E-07 | 0.166189 | 7.80657E-07 |
| rs73146986 | 3 | 89560884 | GTEx/v8 | Artery Tibial | EPHA3 | G | 2.62731E-07 | 0.172075 | 7.80657E-07 |
| rs2117136 | 3 | 89542699 | GTEx/v8 | Artery Tibial | EPHA3 | G | 2.64986E-07 | 0.165797 | 7.80657E-07 |
| rs73145305 | 3 | 89570713 | GTEx/v8 | Artery Tibial | EPHA3 | T | 2.73133E-07 | 0.167896 | 7.80657E-07 |
| 3:89571180_TAGAG_T | 3 | 89571180 | GTEx/v8 | Artery Tibial | EPHA3 | T | 2.73133E-07 | 0.167896 | 7.80657E-07 |
| rs907548 | 3 | 89574852 | GTEx/v8 | Artery Tibial | EPHA3 | T | 2.73133E-07 | 0.167896 | 7.80657E-07 |
| rs73145307 | 3 | 89576040 | GTEx/v8 | Artery Tibial | EPHA3 | A | 2.73133E-07 | 0.167896 | 7.80657E-07 |
| rs73145309 | 3 | 89576389 | GTEx/v8 | Artery Tibial | EPHA3 | T | 2.73133E-07 | 0.167896 | 7.80657E-07 |
| rs73145316 | 3 | 89580407 | GTEx/v8 | Artery Tibial | EPHA3 | T | 2.73133E-07 | 0.167896 | 7.80657E-07 |
| rs73145332 | 3 | 89589327 | GTEx/v8 | Artery Tibial | EPHA3 | C | 2.73133E-07 | 0.167896 | 7.80657E-07 |
| rs972030 | 3 | 89538498 | GTEx/v8 | Artery Tibial | EPHA3 | T | 2.77633E-07 | 0.169134 | 7.80657E-07 |
| rs4857500 | 3 | 89541486 | GTEx/v8 | Artery Tibial | EPHA3 | C | 2.77633E-07 | 0.169134 | 7.80657E-07 |

| | | | | | | | | | |
|---|---|---|---|---|---|---|---|---|---|
| rs9854073 | 3 | 89426589 | GTEx/v8 | Brain Cerebellum | EPHA3 | A | 2.78818E-07 | -0.344028 | 0.000131411 |
| rs9849029 | 3 | 89433059 | GTEx/v8 | Brain Cerebellum | EPHA3 | G | 2.78818E-07 | -0.344028 | 0.000131411 |
| rs73154916 | 3 | 89803713 | GTEx/v8 | Pancreas | EPHA3 | T | 2.84441E-07 | 0.320594 | 3.49803E-07 |
| rs73139133 | 3 | 89527708 | GTEx/v8 | Artery Tibial | EPHA3 | A | 3.03609E-07 | 0.170551 | 7.80657E-07 |
| rs3762718 | 3 | 89535066 | GTEx/v8 | Artery Tibial | EPHA3 | G | 3.11902E-07 | 0.163772 | 7.80657E-07 |
| rs9868782 | 3 | 89384121 | GTEx/v8 | Artery Tibial | EPHA3 | A | 3.16316E-07 | 0.159098 | 7.80657E-07 |
| rs2048519 | 3 | 89533978 | GTEx/v8 | Artery Tibial | EPHA3 | A | 3.1845E-07 | 0.164341 | 7.80657E-07 |
| rs2196083 | 3 | 89536546 | GTEx/v8 | Artery Tibial | EPHA3 | C | 3.1845E-07 | 0.164341 | 7.80657E-07 |
| rs12634044 | 3 | 89537651 | GTEx/v8 | Artery Tibial | EPHA3 | G | 3.1845E-07 | 0.164341 | 7.80657E-07 |
| rs73145353 | 3 | 89607824 | GTEx/v8 | Brain Cerebellum | EPHA3 | T | 3.24758E-07 | -0.368663 | 0.000131411 |
| 3:89626776_AT_A | 3 | 89626776 | GTEx/v8 | Brain Cerebellum | EPHA3 | A | 3.24758E-07 | -0.368663 | 0.000131411 |
| rs7632502 | 3 | 89469453 | GTEx/v8 | Brain Cerebellum | EPHA3 | A | 3.25559E-07 | -0.324027 | 0.000131411 |
| rs1512189 | 3 | 89546332 | GTEx/v8 | Artery Tibial | EPHA3 | G | 3.55683E-07 | 0.169577 | 7.80657E-07 |
| rs6784577 | 3 | 89564308 | GTEx/v8 | Artery Tibial | EPHA3 | G | 3.55683E-07 | 0.169577 | 7.80657E-07 |
| rs73149103 | 3 | 89565651 | GTEx/v8 | Artery Tibial | EPHA3 | A | 3.55683E-07 | 0.169577 | 7.80657E-07 |
| rs12637036 | 3 | 89584170 | GTEx/v8 | Artery Tibial | EPHA3 | A | 3.55683E-07 | 0.169577 | 7.80657E-07 |
| rs56082199 | 3 | 89586973 | GTEx/v8 | Artery Tibial | EPHA3 | G | 3.55683E-07 | 0.169577 | 7.80657E-07 |
| rs73145336 | 3 | 89593317 | GTEx/v8 | Artery Tibial | EPHA3 | G | 3.55683E-07 | 0.169577 | 7.80657E-07 |
| rs73139134 | 3 | 89527709 | GTEx/v8 | Artery Tibial | EPHA3 | C | 3.83286E-07 | 0.167029 | 7.80657E-07 |
| rs73139135 | 3 | 89527937 | GTEx/v8 | Artery Tibial | EPHA3 | C | 3.83286E-07 | 0.167029 | 7.80657E-07 |
| rs73139139 | 3 | 89528313 | GTEx/v8 | Artery Tibial | EPHA3 | G | 3.83286E-07 | 0.167029 | 7.80657E-07 |
| rs12636710 | 3 | 89528477 | GTEx/v8 | Artery Tibial | EPHA3 | A | 3.83286E-07 | 0.167029 | 7.80657E-07 |
| rs3762717 | 3 | 89528948 | GTEx/v8 | Artery Tibial | EPHA3 | C | 3.83286E-07 | 0.167029 | 7.80657E-07 |
| rs73139144 | 3 | 89529177 | GTEx/v8 | Artery Tibial | EPHA3 | A | 3.83286E-07 | 0.167029 | 7.80657E-07 |
| rs73139147 | 3 | 89530347 | GTEx/v8 | Artery Tibial | EPHA3 | A | 3.83286E-07 | 0.167029 | 7.80657E-07 |
| rs73139148 | 3 | 89530956 | GTEx/v8 | Artery Tibial | EPHA3 | A | 3.83286E-07 | 0.167029 | 7.80657E-07 |
| rs34732565 | 3 | 90144346 | GTEx/v8 | Artery Tibial | EPHA3 | A | 3.86359E-07 | -0.150768 | 7.80657E-07 |
| rs73145345 | 3 | 89598696 | GTEx/v8 | Artery Tibial | EPHA3 | A | 4.57662E-07 | 0.168804 | 7.80657E-07 |
| rs111835258 | 3 | 89601496 | GTEx/v8 | Artery Tibial | EPHA3 | A | 4.57662E-07 | 0.168804 | 7.80657E-07 |
| rs73145351 | 3 | 89607237 | GTEx/v8 | Artery Tibial | EPHA3 | T | 4.57662E-07 | 0.168804 | 7.80657E-07 |
| rs73145353 | 3 | 89607824 | GTEx/v8 | Artery Tibial | EPHA3 | T | 4.57662E-07 | 0.168804 | 7.80657E-07 |
| rs73145357 | 3 | 89611766 | GTEx/v8 | Artery Tibial | EPHA3 | C | 4.57662E-07 | 0.168804 | 7.80657E-07 |
| rs73139288 | 3 | 89505327 | GTEx/v8 | Artery Tibial | EPHA3 | G | 4.65482E-07 | 0.166579 | 7.80657E-07 |
| rs73146960 | 3 | 89550445 | GTEx/v8 | Brain Cerebellum | EPHA3 | C | 4.77005E-07 | -0.346034 | 0.000131411 |
| rs73145345 | 3 | 89598696 | GTEx/v8 | Brain Cerebellum | EPHA3 | A | 4.99521E-07 | -0.364614 | 0.000131411 |
| rs111835258 | 3 | 89601496 | GTEx/v8 | Brain Cerebellum | EPHA3 | A | 4.99521E-07 | -0.364614 | 0.000131411 |
| rs73145351 | 3 | 89607237 | GTEx/v8 | Brain Cerebellum | EPHA3 | T | 4.99521E-07 | -0.364614 | 0.000131411 |
| rs73145357 | 3 | 89611766 | GTEx/v8 | Brain Cerebellum | EPHA3 | C | 4.99521E-07 | -0.364614 | 0.000131411 |
| rs7633500 | 3 | 89497082 | GTEx/v8 | Brain Cerebellum | EPHA3 | A | 5.31007E-07 | -0.33354 | 0.000131411 |
| rs3792572 | 3 | 89456555 | GTEx/v8 | Brain Cerebellum | EPHA3 | A | 5.45397E-07 | -0.318779 | 0.000131411 |
| rs35968370 | 3 | 89466339 | GTEx/v8 | Brain Cerebellum | EPHA3 | T | 5.45397E-07 | -0.318779 | 0.000131411 |
| rs11921985 | 3 | 89467357 | GTEx/v8 | Brain Cerebellum | EPHA3 | G | 5.45397E-07 | -0.318779 | 0.000131411 |
| rs6782527 | 3 | 89442369 | GTEx/v8 | Pancreas | EPHA3 | T | 5.49846E-07 | 0.301598 | 3.49803E-07 |
| rs12629999 | 3 | 89551518 | GTEx/v8 | Artery Tibial | EPHA3 | G | 5.60355E-07 | 0.167557 | 7.80657E-07 |
| rs73145303 | 3 | 89568714 | GTEx/v8 | Artery Tibial | EPHA3 | C | 5.60355E-07 | 0.167557 | 7.80657E-07 |
| rs35317084 | 3 | 89354462 | GTEx/v8 | Artery Tibial | EPHA3 | T | 5.74001E-07 | -0.150164 | 7.80657E-07 |
| rs9854073 | 3 | 89426589 | GTEx/v8 | Artery Tibial | EPHA3 | A | 6.43643E-07 | 0.155106 | 7.80657E-07 |
| rs9849029 | 3 | 89433059 | GTEx/v8 | Artery Tibial | EPHA3 | G | 6.43643E-07 | 0.155106 | 7.80657E-07 |
| rs73139129 | 3 | 89524562 | GTEx/v8 | Artery Tibial | EPHA3 | G | 6.47764E-07 | 0.164125 | 7.80657E-07 |
| 3:89626776_AT_A | 3 | 89626776 | GTEx/v8 | Artery Tibial | EPHA3 | A | 7.076E-07 | 0.165047 | 7.80657E-07 |

| | | | | | | | | | |
|---|---|---|---|---|---|---|---|---|---|
| rs73137387 | 3 | 89464194 | GTEx/v8 | Artery Tibial | EPHA3 | C | 7.18472E-07 | 0.163632 | 7.80657E-07 |
| rs73153293 | 3 | 89756242 | GTEx/v8 | Artery Tibial | EPHA3 | T | 7.34654E-07 | 0.172034 | 7.80657E-07 |
| rs1157607 | 3 | 89506364 | GTEx/v8 | Artery Tibial | EPHA3 | T | 7.37407E-07 | 0.163625 | 7.80657E-07 |
| rs7626735 | 3 | 89506958 | GTEx/v8 | Artery Tibial | EPHA3 | A | 7.41145E-07 | 0.163105 | 7.80657E-07 |
| rs6551412 | 3 | 89496502 | GTEx/v8 | Artery Tibial | EPHA3 | A | 7.48723E-07 | 0.162863 | 7.80657E-07 |
| rs6772953 | 3 | 89451721 | GTEx/v8 | Brain Cerebellum | EPHA3 | C | 7.62522E-07 | -0.312523 | 0.000131411 |
| rs112087049 | 3 | 89509550 | GTEx/v8 | Artery Tibial | EPHA3 | T | 7.66988E-07 | 0.168789 | 7.80657E-07 |
| rs73137395 | 3 | 89478753 | GTEx/v8 | Brain Cerebellum | EPHA3 | C | 7.73151E-07 | -0.359363 | 0.000131411 |
| rs73139257 | 3 | 89482779 | GTEx/v8 | Brain Cerebellum | EPHA3 | G | 7.73151E-07 | -0.359363 | 0.000131411 |
| rs73139276 | 3 | 89500196 | GTEx/v8 | Artery Tibial | EPHA3 | A | 7.79111E-07 | 0.164089 | 7.80657E-07 |
| rs62276243 | 3 | 90121534 | GTEx/v8 | Artery Tibial | EPHA3 | T | 9.02879E-07 | -0.145657 | 7.80657E-07 |
| rs4974357 | 3 | 90127762 | GTEx/v8 | Artery Tibial | EPHA3 | C | 9.02879E-07 | -0.145657 | 7.80657E-07 |
| rs7645408 | 3 | 89446132 | GTEx/v8 | Brain Cerebellum | EPHA3 | A | 9.04261E-07 | -0.328249 | 0.000131411 |
| rs28623022 | 3 | 89448936 | GTEx/v8 | Brain Cerebellum | EPHA3 | A | 9.04261E-07 | -0.328249 | 0.000131411 |
| rs73137381 | 3 | 89458550 | GTEx/v8 | Artery Tibial | EPHA3 | G | 9.14616E-07 | 0.164052 | 7.80657E-07 |
| rs73137382 | 3 | 89461363 | GTEx/v8 | Artery Tibial | EPHA3 | G | 9.14616E-07 | 0.164052 | 7.80657E-07 |
| rs57670516 | 3 | 89419645 | GTEx/v8 | Brain Cerebellum | EPHA3 | C | 9.44625E-07 | -0.331888 | 0.000131411 |
| 3:89420088_TGTGCACCTAGAC_T | 3 | 89420088 | GTEx/v8 | Brain Cerebellum | EPHA3 | T | 9.44625E-07 | -0.331888 | 0.000131411 |
| rs12639506 | 3 | 89480265 | GTEx/v8 | Artery Tibial | EPHA3 | G | 9.92726E-07 | 0.1612 | 7.80657E-07 |
| rs57670516 | 3 | 89419645 | GTEx/v8 | Artery Tibial | EPHA3 | C | 1.00519E-06 | 0.152977 | 7.80657E-07 |
| 3:89420088_TGTGCACCTAGAC_T | 3 | 89420088 | GTEx/v8 | Artery Tibial | EPHA3 | T | 1.00519E-06 | 0.152977 | 7.80657E-07 |
| rs6795074 | 3 | 89516652 | GTEx/v8 | Artery Tibial | EPHA3 | C | 1.01653E-06 | 0.161217 | 7.80657E-07 |
| rs73141108 | 3 | 89517119 | GTEx/v8 | Artery Tibial | EPHA3 | T | 1.01653E-06 | 0.161217 | 7.80657E-07 |
| rs1912966 | 3 | 89518464 | GTEx/v8 | Artery Tibial | EPHA3 | A | 1.01653E-06 | 0.161217 | 7.80657E-07 |
| rs73139104 | 3 | 89519229 | GTEx/v8 | Artery Tibial | EPHA3 | C | 1.01653E-06 | 0.161217 | 7.80657E-07 |
| 3:89519266_GA_G | 3 | 89519266 | GTEx/v8 | Artery Tibial | EPHA3 | G | 1.01653E-06 | 0.161217 | 7.80657E-07 |
| 3:89521060_CTAT_C | 3 | 89521060 | GTEx/v8 | Artery Tibial | EPHA3 | C | 1.01653E-06 | 0.161217 | 7.80657E-07 |
| rs73139110 | 3 | 89521509 | GTEx/v8 | Artery Tibial | EPHA3 | A | 1.01653E-06 | 0.161217 | 7.80657E-07 |
| rs1054750 | 3 | 89521725 | GTEx/v8 | Artery Tibial | EPHA3 | C | 1.01653E-06 | 0.161217 | 7.80657E-07 |
| rs73139115 | 3 | 89521900 | GTEx/v8 | Artery Tibial | EPHA3 | T | 1.01653E-06 | 0.161217 | 7.80657E-07 |
| rs73139117 | 3 | 89521965 | GTEx/v8 | Artery Tibial | EPHA3 | G | 1.01653E-06 | 0.161217 | 7.80657E-07 |
| rs73139118 | 3 | 89522263 | GTEx/v8 | Artery Tibial | EPHA3 | T | 1.01653E-06 | 0.161217 | 7.80657E-07 |
| rs150837250 | 3 | 89522314 | GTEx/v8 | Artery Tibial | EPHA3 | AGTTATAATTAG | 1.01653E-06 | 0.161217 | 7.80657E-07 |
| rs17801380 | 3 | 89522483 | GTEx/v8 | Artery Tibial | EPHA3 | T | 1.01653E-06 | 0.161217 | 7.80657E-07 |
| rs73139127 | 3 | 89524436 | GTEx/v8 | Artery Tibial | EPHA3 | A | 1.01653E-06 | 0.161217 | 7.80657E-07 |
| rs568140553 | 3 | 89507219 | GTEx/v8 | Artery Tibial | EPHA3 | CT | 1.01848E-06 | 0.160739 | 7.80657E-07 |
| rs73139261 | 3 | 89492406 | GTEx/v8 | Artery Tibial | EPHA3 | G | 1.0393E-06 | 0.164521 | 7.80657E-07 |
| rs1499780 | 3 | 89526072 | GTEx/v8 | Artery Tibial | EPHA3 | C | 1.05303E-06 | 0.161305 | 7.80657E-07 |
| rs2048518 | 3 | 89533905 | GTEx/v8 | Brain Cerebellum | EPHA3 | C | 1.06295E-06 | -0.335412 | 0.000131411 |
| rs2048520 | 3 | 89534248 | GTEx/v8 | Brain Cerebellum | EPHA3 | C | 1.06295E-06 | -0.335412 | 0.000131411 |
| rs2048521 | 3 | 89534253 | GTEx/v8 | Brain Cerebellum | EPHA3 | A | 1.06295E-06 | -0.335412 | 0.000131411 |
| rs73144998 | 3 | 89534592 | GTEx/v8 | Brain Cerebellum | EPHA3 | T | 1.06295E-06 | -0.335412 | 0.000131411 |
| rs73145002 | 3 | 89537618 | GTEx/v8 | Brain Cerebellum | EPHA3 | C | 1.06295E-06 | -0.335412 | 0.000131411 |
| rs56371328 | 3 | 89539124 | GTEx/v8 | Brain Cerebellum | EPHA3 | A | 1.06295E-06 | -0.335412 | 0.000131411 |
| rs4857498 | 3 | 89540977 | GTEx/v8 | Brain Cerebellum | EPHA3 | A | 1.06295E-06 | -0.335412 | 0.000131411 |
| rs4857499 | 3 | 89541330 | GTEx/v8 | Brain Cerebellum | EPHA3 | A | 1.06295E-06 | -0.335412 | 0.000131411 |
| rs1036285 | 3 | 89544061 | GTEx/v8 | Brain Cerebellum | EPHA3 | C | 1.06295E-06 | -0.335412 | 0.000131411 |

| | | | | | | | | | |
|---|---|---|---|---|---|---|---|---|---|
| rs17026984 | 3 | 89544978 | GTEx/v8 | Brain Cerebellum | EPHA3 | A | 1.06295E-06 | -0.335412 | 0.000131411 |
| rs12633609 | 3 | 89545727 | GTEx/v8 | Brain Cerebellum | EPHA3 | G | 1.06295E-06 | -0.335412 | 0.000131411 |
| rs1512188 | 3 | 89546183 | GTEx/v8 | Brain Cerebellum | EPHA3 | G | 1.06295E-06 | -0.335412 | 0.000131411 |
| rs73146943 | 3 | 89546728 | GTEx/v8 | Brain Cerebellum | EPHA3 | G | 1.06295E-06 | -0.335412 | 0.000131411 |
| rs12632281 | 3 | 89547621 | GTEx/v8 | Brain Cerebellum | EPHA3 | A | 1.06295E-06 | -0.335412 | 0.000131411 |
| rs73146947 | 3 | 89548135 | GTEx/v8 | Brain Cerebellum | EPHA3 | G | 1.06295E-06 | -0.335412 | 0.000131411 |
| rs73146950 | 3 | 89548243 | GTEx/v8 | Brain Cerebellum | EPHA3 | C | 1.06295E-06 | -0.335412 | 0.000131411 |
| rs1567657 | 3 | 89548519 | GTEx/v8 | Brain Cerebellum | EPHA3 | A | 1.06295E-06 | -0.335412 | 0.000131411 |
| rs1567658 | 3 | 89548606 | GTEx/v8 | Brain Cerebellum | EPHA3 | A | 1.06295E-06 | -0.335412 | 0.000131411 |
| rs73146959 | 3 | 89549970 | GTEx/v8 | Brain Cerebellum | EPHA3 | C | 1.06295E-06 | -0.335412 | 0.000131411 |
| rs112404167 | 3 | 89550630 | GTEx/v8 | Brain Cerebellum | EPHA3 | T | 1.06295E-06 | -0.335412 | 0.000131411 |
| rs12637511 | 3 | 89550844 | GTEx/v8 | Brain Cerebellum | EPHA3 | C | 1.06295E-06 | -0.335412 | 0.000131411 |
| rs111618480 | 3 | 89552134 | GTEx/v8 | Brain Cerebellum | EPHA3 | C | 1.06295E-06 | -0.335412 | 0.000131411 |
| rs73146972 | 3 | 89553348 | GTEx/v8 | Brain Cerebellum | EPHA3 | G | 1.06295E-06 | -0.335412 | 0.000131411 |
| rs17026990 | 3 | 89555257 | GTEx/v8 | Brain Cerebellum | EPHA3 | C | 1.06295E-06 | -0.335412 | 0.000131411 |
| rs73146976 | 3 | 89557065 | GTEx/v8 | Brain Cerebellum | EPHA3 | C | 1.06295E-06 | -0.335412 | 0.000131411 |
| rs1039991 | 3 | 89558311 | GTEx/v8 | Brain Cerebellum | EPHA3 | G | 1.06295E-06 | -0.335412 | 0.000131411 |
| rs73146980 | 3 | 89558578 | GTEx/v8 | Brain Cerebellum | EPHA3 | A | 1.06295E-06 | -0.335412 | 0.000131411 |
| rs73146983 | 3 | 89559082 | GTEx/v8 | Brain Cerebellum | EPHA3 | G | 1.06295E-06 | -0.335412 | 0.000131411 |
| rs138490449 | 3 | 89561661 | GTEx/v8 | Brain Cerebellum | EPHA3 | AT | 1.06295E-06 | -0.335412 | 0.000131411 |
| rs138183278 | 3 | 89564090 | GTEx/v8 | Brain Cerebellum | EPHA3 | CT | 1.06295E-06 | -0.335412 | 0.000131411 |
| rs6774870 | 3 | 89564891 | GTEx/v8 | Brain Cerebellum | EPHA3 | T | 1.06295E-06 | -0.335412 | 0.000131411 |
| rs113307255 | 3 | 89567826 | GTEx/v8 | Brain Cerebellum | EPHA3 | C | 1.06295E-06 | -0.335412 | 0.000131411 |
| rs73145304 | 3 | 89568989 | GTEx/v8 | Brain Cerebellum | EPHA3 | T | 1.06295E-06 | -0.335412 | 0.000131411 |
| rs73145310 | 3 | 89578097 | GTEx/v8 | Brain Cerebellum | EPHA3 | C | 1.06295E-06 | -0.335412 | 0.000131411 |
| rs73145312 | 3 | 89578575 | GTEx/v8 | Brain Cerebellum | EPHA3 | T | 1.06295E-06 | -0.335412 | 0.000131411 |
| rs73145320 | 3 | 89586535 | GTEx/v8 | Brain Cerebellum | EPHA3 | C | 1.06295E-06 | -0.335412 | 0.000131411 |
| rs67316928 | 3 | 89482234 | GTEx/v8 | Brain Cerebellum | EPHA3 | C | 1.0649E-06 | -0.313684 | 0.000131411 |
| rs17804073 | 3 | 90124806 | GTEx/v8 | Artery Tibial | EPHA3 | A | 1.07195E-06 | -0.145091 | 7.80657E-07 |
| rs12636275 | 3 | 89523038 | GTEx/v8 | Brain Cerebellum | EPHA3 | A | 1.08168E-06 | -0.311412 | 0.000131411 |
| rs73137393 | 3 | 89471701 | GTEx/v8 | Artery Tibial | EPHA3 | C | 1.13522E-06 | 0.162812 | 7.80657E-07 |
| rs7650184 | 3 | 89530057 | GTEx/v8 | Brain Cerebellum | EPHA3 | A | 1.16781E-06 | -0.311417 | 0.000131411 |
| rs17738248 | 3 | 89296363 | GTEx/v8 | Artery Tibial | EPHA3 | A | 1.18583E-06 | 0.1608 | 7.80657E-07 |
| rs9868782 | 3 | 89384121 | GTEx/v8 | Brain Cerebellum | EPHA3 | A | 1.19018E-06 | -0.33257 | 0.000131411 |
| rs1521800 | 3 | 90001588 | GTEx/v8 | Artery Tibial | EPHA3 | T | 1.24942E-06 | -0.143263 | 7.80657E-07 |
| rs73145338 | 3 | 89594767 | GTEx/v8 | Brain Cerebellum | EPHA3 | T | 1.25465E-06 | -0.306581 | 0.000131411 |
| rs7621573 | 3 | 89595252 | GTEx/v8 | Brain Cerebellum | EPHA3 | T | 1.25465E-06 | -0.306581 | 0.000131411 |
| rs10511145 | 3 | 89597679 | GTEx/v8 | Brain Cerebellum | EPHA3 | A | 1.25465E-06 | -0.306581 | 0.000131411 |
| 3:89598352_GGT_G | 3 | 89598352 | GTEx/v8 | Brain Cerebellum | EPHA3 | G | 1.25465E-06 | -0.306581 | 0.000131411 |
| rs17027018 | 3 | 89604774 | GTEx/v8 | Brain Cerebellum | EPHA3 | T | 1.25465E-06 | -0.306581 | 0.000131411 |
| rs73153219 | 3 | 89625902 | GTEx/v8 | Brain Cerebellum | EPHA3 | T | 1.25465E-06 | -0.306581 | 0.000131411 |
| rs7635916 | 3 | 89633140 | GTEx/v8 | Brain Cerebellum | EPHA3 | A | 1.25465E-06 | -0.306581 | 0.000131411 |
| rs6551413 | 3 | 89640350 | GTEx/v8 | Brain Cerebellum | EPHA3 | G | 1.25465E-06 | -0.306581 | 0.000131411 |

| | | | | | | | | | |
|---|---|---|---|---|---|---|---|---|---|
| rs6795325 | 3 | 89642054 | GTEx/v8 | Brain Cerebellum | EPHA3 | C | 1.25465E-06 | -0.306581 | 0.000131411 |
| rs66499884 | 3 | 89659012 | GTEx/v8 | Brain Cerebellum | EPHA3 | G | 1.25465E-06 | -0.306581 | 0.000131411 |
| rs2063590 | 3 | 89323932 | GTEx/v8 | Artery Tibial | EPHA3 | C | 1.27188E-06 | 0.162629 | 7.80657E-07 |
| rs73151349 | 3 | 89892899 | GTEx/v8 | Pancreas | EPHA3 | G | 1.33186E-06 | 0.312228 | 3.49803E-07 |
| rs4396913 | 3 | 90047036 | GTEx/v8 | Artery Tibial | EPHA3 | C | 1.37457E-06 | 0.15249 | 7.80657E-07 |
| rs1464463 | 3 | 90056380 | GTEx/v8 | Artery Tibial | EPHA3 | A | 1.37457E-06 | 0.15249 | 7.80657E-07 |
| rs2346836 | 3 | 89498096 | GTEx/v8 | Brain Cerebellum | EPHA3 | A | 1.39977E-06 | -0.345515 | 0.000131411 |
| rs60759140 | 3 | 89499848 | GTEx/v8 | Brain Cerebellum | EPHA3 | C | 1.39977E-06 | -0.345515 | 0.000131411 |
| rs12152372 | 3 | 89500365 | GTEx/v8 | Brain Cerebellum | EPHA3 | C | 1.39977E-06 | -0.345515 | 0.000131411 |
| rs2881488 | 3 | 89500488 | GTEx/v8 | Brain Cerebellum | EPHA3 | G | 1.39977E-06 | -0.345515 | 0.000131411 |
| rs59541469 | 3 | 89501671 | GTEx/v8 | Brain Cerebellum | EPHA3 | T | 1.39977E-06 | -0.345515 | 0.000131411 |
| 3:89502123_TTAG_T | 3 | 89502123 | GTEx/v8 | Brain Cerebellum | EPHA3 | T | 1.39977E-06 | -0.345515 | 0.000131411 |
| rs73139281 | 3 | 89503890 | GTEx/v8 | Brain Cerebellum | EPHA3 | T | 1.39977E-06 | -0.345515 | 0.000131411 |
| rs73139284 | 3 | 89503958 | GTEx/v8 | Brain Cerebellum | EPHA3 | G | 1.39977E-06 | -0.345515 | 0.000131411 |
| rs73139286 | 3 | 89504295 | GTEx/v8 | Brain Cerebellum | EPHA3 | A | 1.39977E-06 | -0.345515 | 0.000131411 |
| rs1512183 | 3 | 89505054 | GTEx/v8 | Brain Cerebellum | EPHA3 | T | 1.39977E-06 | -0.345515 | 0.000131411 |
| rs73139290 | 3 | 89505343 | GTEx/v8 | Brain Cerebellum | EPHA3 | A | 1.39977E-06 | -0.345515 | 0.000131411 |
| rs73139296 | 3 | 89505734 | GTEx/v8 | Brain Cerebellum | EPHA3 | T | 1.39977E-06 | -0.345515 | 0.000131411 |
| rs1157608 | 3 | 89506464 | GTEx/v8 | Brain Cerebellum | EPHA3 | G | 1.39977E-06 | -0.345515 | 0.000131411 |
| rs73139302 | 3 | 89508186 | GTEx/v8 | Brain Cerebellum | EPHA3 | G | 1.39977E-06 | -0.345515 | 0.000131411 |
| rs73141104 | 3 | 89508390 | GTEx/v8 | Brain Cerebellum | EPHA3 | T | 1.39977E-06 | -0.345515 | 0.000131411 |
| rs1912965 | 3 | 89518332 | GTEx/v8 | Brain Cerebellum | EPHA3 | G | 1.39977E-06 | -0.345515 | 0.000131411 |
| rs113141104 | 3 | 89519238 | GTEx/v8 | Brain Cerebellum | EPHA3 | T | 1.39977E-06 | -0.345515 | 0.000131411 |
| rs73139105 | 3 | 89519651 | GTEx/v8 | Brain Cerebellum | EPHA3 | C | 1.39977E-06 | -0.345515 | 0.000131411 |
| rs17026944 | 3 | 89521131 | GTEx/v8 | Brain Cerebellum | EPHA3 | A | 1.39977E-06 | -0.345515 | 0.000131411 |
| rs112297761 | 3 | 89521602 | GTEx/v8 | Brain Cerebellum | EPHA3 | TC | 1.39977E-06 | -0.345515 | 0.000131411 |
| rs73139119 | 3 | 89522285 | GTEx/v8 | Brain Cerebellum | EPHA3 | T | 1.39977E-06 | -0.345515 | 0.000131411 |
| rs73139121 | 3 | 89522433 | GTEx/v8 | Brain Cerebellum | EPHA3 | G | 1.39977E-06 | -0.345515 | 0.000131411 |
| rs73139125 | 3 | 89524056 | GTEx/v8 | Brain Cerebellum | EPHA3 | G | 1.39977E-06 | -0.345515 | 0.000131411 |
| rs73139131 | 3 | 89524686 | GTEx/v8 | Brain Cerebellum | EPHA3 | G | 1.39977E-06 | -0.345515 | 0.000131411 |
| rs9877921 | 3 | 90108491 | GTEx/v8 | Artery Tibial | EPHA3 | T | 1.40136E-06 | 0.152782 | 7.80657E-07 |
| rs62276239 | 3 | 90077991 | GTEx/v8 | Artery Tibial | EPHA3 | T | 1.40499E-06 | 0.158179 | 7.80657E-07 |
| rs55889178 | 3 | 90078249 | GTEx/v8 | Artery Tibial | EPHA3 | A | 1.40499E-06 | 0.158179 | 7.80657E-07 |
| rs1520596 | 3 | 89645542 | GTEx/v8 | Brain Cerebellum | EPHA3 | C | 1.45459E-06 | -0.299389 | 0.000131411 |
| rs73146965 | 3 | 89552553 | GTEx/v8 | Brain Cerebellum | EPHA3 | G | 1.4669E-06 | -0.331695 | 0.000131411 |
| rs35124509 | 3 | 89521693 | GTEx/v8 | Brain Cerebellum | EPHA3 | C | 1.48256E-06 | -0.305957 | 0.000131411 |
| rs62274981 | 3 | 89310605 | GTEx/v8 | Brain Cerebellum | EPHA3 | T | 1.48407E-06 | -0.326947 | 0.000131411 |
| rs17800347 | 3 | 89314429 | GTEx/v8 | Brain Cerebellum | EPHA3 | G | 1.48407E-06 | -0.326947 | 0.000131411 |
| rs993186 | 3 | 89314862 | GTEx/v8 | Brain Cerebellum | EPHA3 | G | 1.48407E-06 | -0.326947 | 0.000131411 |
| rs1567731 | 3 | 89316027 | GTEx/v8 | Brain Cerebellum | EPHA3 | T | 1.48407E-06 | -0.326947 | 0.000131411 |
| rs17800551 | 3 | 89317315 | GTEx/v8 | Brain Cerebellum | EPHA3 | T | 1.48407E-06 | -0.326947 | 0.000131411 |
| rs62274984 | 3 | 89318317 | GTEx/v8 | Brain Cerebellum | EPHA3 | A | 1.48407E-06 | -0.326947 | 0.000131411 |
| rs12633626 | 3 | 89744949 | GTEx/v8 | Brain Cerebellum | EPHA3 | A | 1.526E-06 | -0.348666 | 0.000131411 |
| rs60243237 | 3 | 89694053 | GTEx/v8 | Brain Cerebellum | EPHA3 | T | 1.52877E-06 | -0.305415 | 0.000131411 |

| | | | | | | | | | |
|---|---|---|---|---|---|---|---|---|---|
| rs987748 | 3 | 89494030 | GTEx/v8 | Brain Cerebellum | EPHA3 | A | 1.53426E-06 | -0.31043 | 0.000131411 |
| rs6800041 | 3 | 89448196 | GTEx/v8 | Brain Cerebellum | EPHA3 | C | 1.53723E-06 | -0.323538 | 0.000131411 |
| rs12714734 | 3 | 90107488 | GTEx/v8 | Artery Tibial | EPHA3 | C | 1.57366E-06 | 0.151478 | 7.80657E-07 |
| rs9816940 | 3 | 90146579 | GTEx/v8 | Artery Tibial | EPHA3 | T | 1.58919E-06 | 0.153017 | 7.80657E-07 |
| rs9837548 | 3 | 90156142 | GTEx/v8 | Artery Tibial | EPHA3 | T | 1.58919E-06 | 0.153017 | 7.80657E-07 |
| rs7637670 | 3 | 89493167 | GTEx/v8 | Brain Cerebellum | EPHA3 | A | 1.60292E-06 | -0.309849 | 0.000131411 |
| rs11918555 | 3 | 89541934 | GTEx/v8 | Brain Cerebellum | EPHA3 | T | 1.62669E-06 | -0.301267 | 0.000131411 |
| rs12637608 | 3 | 89551301 | GTEx/v8 | Brain Cerebellum | EPHA3 | C | 1.62669E-06 | -0.301267 | 0.000131411 |
| rs66757381 | 3 | 89561048 | GTEx/v8 | Brain Cerebellum | EPHA3 | T | 1.62669E-06 | -0.301267 | 0.000131411 |
| rs724972 | 3 | 89581408 | GTEx/v8 | Brain Cerebellum | EPHA3 | T | 1.62669E-06 | -0.301267 | 0.000131411 |
| rs7652296 | 3 | 89587262 | GTEx/v8 | Brain Cerebellum | EPHA3 | G | 1.62669E-06 | -0.301267 | 0.000131411 |
| rs993187 | 3 | 89314820 | GTEx/v8 | Artery Tibial | EPHA3 | G | 1.68692E-06 | 0.158464 | 7.80657E-07 |
| rs11925143 | 3 | 89723684 | GTEx/v8 | Brain Cerebellum | EPHA3 | A | 1.70688E-06 | -0.30234 | 0.000131411 |
| rs7632427 | 3 | 89534377 | eQTLGen | eQTLGen trans eQTLs | RBBP9 | C | 1.8006E-06 | -4.7745 | 0.0140155531990103 |
| rs7633500 | 3 | 89497082 | GTEx/v8 | Pancreas | EPHA3 | A | 1.89641E-06 | 0.259798 | 3.49803E-07 |
| rs6806912 | 3 | 89751353 | GTEx/v8 | Brain Cerebellum | EPHA3 | C | 1.9144E-06 | -0.305863 | 0.000131411 |
| rs7653735 | 3 | 89722185 | GTEx/v8 | Brain Cerebellum | EPHA3 | A | 1.95581E-06 | -0.303396 | 0.000131411 |
| rs9863058 | 3 | 89322805 | GTEx/v8 | Brain Cerebellum | EPHA3 | T | 1.9655E-06 | -0.322799 | 0.000131411 |
| rs4857502 | 3 | 89324919 | GTEx/v8 | Brain Cerebellum | EPHA3 | A | 1.9655E-06 | -0.322799 | 0.000131411 |
| rs2137487 | 3 | 89325520 | GTEx/v8 | Brain Cerebellum | EPHA3 | G | 1.9655E-06 | -0.322799 | 0.000131411 |
| rs9875278 | 3 | 89335463 | GTEx/v8 | Brain Cerebellum | EPHA3 | T | 1.9655E-06 | -0.322799 | 0.000131411 |
| rs9821666 | 3 | 89339548 | GTEx/v8 | Brain Cerebellum | EPHA3 | T | 1.9655E-06 | -0.322799 | 0.000131411 |
| rs1512908 | 3 | 89340134 | GTEx/v8 | Brain Cerebellum | EPHA3 | A | 1.9655E-06 | -0.322799 | 0.000131411 |
| rs9852380 | 3 | 89341738 | GTEx/v8 | Brain Cerebellum | EPHA3 | G | 1.9655E-06 | -0.322799 | 0.000131411 |
| rs9852722 | 3 | 89341946 | GTEx/v8 | Brain Cerebellum | EPHA3 | C | 1.9655E-06 | -0.322799 | 0.000131411 |
| rs73137384 | 3 | 89463290 | GTEx/v8 | Brain Cerebellum | EPHA3 | T | 1.97372E-06 | -0.344073 | 0.000131411 |
| rs9832305 | 3 | 89392778 | GTEx/v8 | Brain Cerebellum | EPHA3 | C | 2.03152E-06 | -0.317078 | 0.000131411 |
| rs7615949 | 3 | 89432683 | GTEx/v8 | Brain Cerebellum | EPHA3 | G | 2.05275E-06 | -0.317964 | 0.000131411 |
| rs113450730 | 3 | 90135516 | GTEx/v8 | Artery Tibial | EPHA3 | G | 2.23345E-06 | 0.156885 | 7.80657E-07 |
| rs2117138 | 3 | 89531522 | GTEx/v8 | Brain Cerebellum | EPHA3 | G | 2.25084E-06 | -0.332679 | 0.000131411 |
| rs12634269 | 3 | 89532768 | GTEx/v8 | Brain Cerebellum | EPHA3 | C | 2.25084E-06 | -0.332679 | 0.000131411 |
| rs819293 | 3 | 89613664 | GTEx/v8 | Brain Cerebellum | EPHA3 | A | 2.2568E-06 | -0.340532 | 0.000131411 |
| rs12633626 | 3 | 89744949 | GTEx/v8 | Artery Tibial | EPHA3 | A | 2.299E-06 | 0.157899 | 7.80657E-07 |
| rs2346837 | 3 | 89542599 | GTEx/v8 | Brain Cerebellum | EPHA3 | T | 2.39154E-06 | -0.326146 | 0.000131411 |
| rs11923303 | 3 | 89380944 | GTEx/v8 | Brain Cerebellum | EPHA3 | A | 2.42101E-06 | -0.296307 | 0.000131411 |
| rs9864984 | 3 | 89428776 | GTEx/v8 | Brain Cerebellum | EPHA3 | T | 2.5615E-06 | -0.315584 | 0.000131411 |
| rs9825569 | 3 | 89431650 | GTEx/v8 | Brain Cerebellum | EPHA3 | C | 2.5615E-06 | -0.315584 | 0.000131411 |
| rs62275019 | 3 | 89433648 | GTEx/v8 | Brain Cerebellum | EPHA3 | G | 2.5615E-06 | -0.315584 | 0.000131411 |
| rs7619303 | 3 | 89552554 | GTEx/v8 | Brain Cerebellum | EPHA3 | A | 2.58965E-06 | -0.320493 | 0.000131411 |
| rs73149109 | 3 | 89567344 | GTEx/v8 | Brain Cerebellum | EPHA3 | A | 2.58965E-06 | -0.320493 | 0.000131411 |
| rs1080592 | 3 | 89567568 | GTEx/v8 | Brain Cerebellum | EPHA3 | C | 2.58965E-06 | -0.320493 | 0.000131411 |
| rs62278163 | 3 | 89291836 | GTEx/v8 | Brain Cerebellum | EPHA3 | T | 2.68546E-06 | -0.32343 | 0.000131411 |
| rs6786328 | 3 | 89298926 | GTEx/v8 | Brain Cerebellum | EPHA3 | G | 2.68546E-06 | -0.32343 | 0.000131411 |
| rs62278168 | 3 | 89301573 | GTEx/v8 | Brain Cerebellum | EPHA3 | G | 2.68546E-06 | -0.32343 | 0.000131411 |

| | | | | | | | | | |
|---|---|---|---|---|---|---|---|---|---|
| rs28620812 | 3 | 89305668 | GTEx/v8 | Brain Cerebellum | EPHA3 | T | 2.68546E-06 | -0.32343 | 0.000131411 |
| rs62278172 | 3 | 89306038 | GTEx/v8 | Brain Cerebellum | EPHA3 | A | 2.68546E-06 | -0.32343 | 0.000131411 |
| rs9310120 | 3 | 89378413 | GTEx/v8 | Brain Cerebellum | EPHA3 | C | 2.72467E-06 | -0.291525 | 0.000131411 |
| rs35511518 | 3 | 89383683 | GTEx/v8 | Brain Cerebellum | EPHA3 | C | 2.88981E-06 | -0.293769 | 0.000131411 |
| rs55931912 | 3 | 90038054 | GTEx/v8 | Artery Tibial | EPHA3 | A | 2.94174E-06 | 0.160673 | 7.80657E-07 |
| rs6782527 | 3 | 89442369 | GTEx/v8 | Brain Cerebellum | EPHA3 | T | 2.94697E-06 | -0.323056 | 0.000131411 |
| rs1512185 | 3 | 89473615 | GTEx/v8 | Brain Cerebellum | EPHA3 | A | 2.95573E-06 | -0.333755 | 0.000131411 |
| rs2069183 | 3 | 89645017 | GTEx/v8 | Brain Cerebellum | EPHA3 | A | 2.97291E-06 | -0.333067 | 0.000131411 |
| rs2137488 | 3 | 89308704 | GTEx/v8 | Brain Cerebellum | EPHA3 | G | 3.01033E-06 | -0.321737 | 0.000131411 |
| rs62274982 | 3 | 89310731 | GTEx/v8 | Brain Cerebellum | EPHA3 | A | 3.01033E-06 | -0.321737 | 0.000131411 |
| rs9866618 | 3 | 89319577 | GTEx/v8 | Brain Cerebellum | EPHA3 | C | 3.01033E-06 | -0.321737 | 0.000131411 |
| rs138643396 | 3 | 89698092 | GTEx/v8 | Brain Cerebellum | EPHA3 | CATA | 3.03531E-06 | -0.29645 | 0.000131411 |
| rs59381722 | 3 | 89701482 | GTEx/v8 | Brain Cerebellum | EPHA3 | G | 3.03531E-06 | -0.29645 | 0.000131411 |
| rs7637504 | 3 | 89702469 | GTEx/v8 | Brain Cerebellum | EPHA3 | T | 3.03531E-06 | -0.29645 | 0.000131411 |
| rs6551415 | 3 | 89705161 | GTEx/v8 | Brain Cerebellum | EPHA3 | T | 3.03531E-06 | -0.29645 | 0.000131411 |
| rs9875278 | 3 | 89335463 | GTEx/v8 | Artery Tibial | EPHA3 | T | 3.09831E-06 | 0.148813 | 7.80657E-07 |
| rs35448293 | 3 | 89386010 | GTEx/v8 | Brain Cerebellum | EPHA3 | G | 3.10479E-06 | -0.294378 | 0.000131411 |
| rs144317613 | 3 | 89525080 | GTEx/v8 | Artery Tibial | EPHA3 | TA | 3.16115E-06 | 0.154826 | 7.80657E-07 |
| rs35173319 | 3 | 89386361 | GTEx/v8 | Brain Cerebellum | EPHA3 | T | 3.26506E-06 | -0.293781 | 0.000131411 |
| rs12631313 | 3 | 89528461 | GTEx/v8 | Brain Cerebellum | EPHA3 | T | 3.26615E-06 | -0.327927 | 0.000131411 |
| rs7650466 | 3 | 89530358 | GTEx/v8 | Brain Cerebellum | EPHA3 | T | 3.26615E-06 | -0.327927 | 0.000131411 |
| rs112407114 | 3 | 89797398 | GTEx/v8 | Artery Tibial | EPHA3 | C | 3.30588E-06 | 0.157525 | 7.80657E-07 |
| rs12638746 | 3 | 89331055 | GTEx/v8 | Artery Tibial | EPHA3 | A | 3.30854E-06 | -0.141292 | 7.80657E-07 |
| rs9848971 | 3 | 89279394 | GTEx/v8 | Brain Cerebellum | EPHA3 | G | 3.5827E-06 | -0.323238 | 0.000131411 |
| rs4857502 | 3 | 89324919 | GTEx/v8 | Artery Tibial | EPHA3 | A | 3.61635E-06 | 0.14766 | 7.80657E-07 |
| rs1567731 | 3 | 89316027 | GTEx/v8 | Artery Tibial | EPHA3 | T | 3.66393E-06 | 0.147438 | 7.80657E-07 |
| rs28620812 | 3 | 89305668 | GTEx/v8 | Artery Tibial | EPHA3 | T | 3.66811E-06 | 0.147499 | 7.80657E-07 |
| rs62278172 | 3 | 89306038 | GTEx/v8 | Artery Tibial | EPHA3 | A | 3.66811E-06 | 0.147499 | 7.80657E-07 |
| rs9817553 | 3 | 89384566 | GTEx/v8 | Brain Cerebellum | EPHA3 | G | 3.69048E-06 | -0.289832 | 0.000131411 |
| rs6776967 | 3 | 90025022 | GTEx/v8 | Artery Tibial | EPHA3 | T | 3.79842E-06 | 0.151056 | 7.80657E-07 |
| rs9310119 | 3 | 89378042 | GTEx/v8 | Brain Cerebellum | EPHA3 | G | 3.8657E-06 | -0.290574 | 0.000131411 |
| rs28675360 | 3 | 89386151 | GTEx/v8 | Brain Cerebellum | EPHA3 | T | 3.88293E-06 | -0.291267 | 0.000131411 |
| rs9863058 | 3 | 89322805 | GTEx/v8 | Artery Tibial | EPHA3 | T | 3.93751E-06 | 0.146765 | 7.80657E-07 |
| rs62274986 | 3 | 89327148 | GTEx/v8 | Brain Cerebellum | EPHA3 | A | 3.93801E-06 | -0.317653 | 0.000131411 |
| rs73153370 | 3 | 90056125 | GTEx/v8 | Artery Tibial | EPHA3 | A | 3.94503E-06 | 0.151255 | 7.80657E-07 |
| rs57670516 | 3 | 89419645 | GTEx/v8 | Pancreas | EPHA3 | C | 4.16777E-06 | 0.26691 | 3.49803E-07 |
| 3:89420088_TGTGCACCTAGAC_T | 3 | 89420088 | GTEx/v8 | Pancreas | EPHA3 | T | 4.16777E-06 | 0.26691 | 3.49803E-07 |
| rs9854073 | 3 | 89426589 | GTEx/v8 | Pancreas | EPHA3 | A | 4.16777E-06 | 0.26691 | 3.49803E-07 |
| rs9849029 | 3 | 89433059 | GTEx/v8 | Pancreas | EPHA3 | G | 4.16777E-06 | 0.26691 | 3.49803E-07 |
| rs2137487 | 3 | 89325520 | GTEx/v8 | Artery Tibial | EPHA3 | G | 4.23081E-06 | 0.146733 | 7.80657E-07 |
| rs9821666 | 3 | 89339548 | GTEx/v8 | Artery Tibial | EPHA3 | T | 4.23081E-06 | 0.146733 | 7.80657E-07 |
| rs1512908 | 3 | 89340134 | GTEx/v8 | Artery Tibial | EPHA3 | A | 4.23081E-06 | 0.146733 | 7.80657E-07 |
| rs9852380 | 3 | 89341738 | GTEx/v8 | Artery Tibial | EPHA3 | G | 4.23081E-06 | 0.146733 | 7.80657E-07 |
| rs9852722 | 3 | 89341946 | GTEx/v8 | Artery Tibial | EPHA3 | C | 4.23081E-06 | 0.146733 | 7.80657E-07 |
| rs1398534 | 3 | 89329307 | GTEx/v8 | Brain Cerebellum | EPHA3 | G | 4.2324E-06 | -0.313436 | 0.000131411 |
| rs17800551 | 3 | 89317315 | GTEx/v8 | Artery Tibial | EPHA3 | T | 4.30899E-06 | 0.146349 | 7.80657E-07 |
| rs62274984 | 3 | 89318317 | GTEx/v8 | Artery Tibial | EPHA3 | A | 4.30899E-06 | 0.146349 | 7.80657E-07 |

| | | | | | | | | | |
|---|---|---|---|---|---|---|---|---|---|
| rs752364173 | 3 | 89521820 | GTEx/v8 | Brain Cerebellum | EPHA3 | T | 4.47555E-06 | -0.325467 | 0.000131411 |
| 3:89521831_TGAAAG_T | 3 | 89521831 | GTEx/v8 | Brain Cerebellum | EPHA3 | T | 4.47555E-06 | -0.325467 | 0.000131411 |
| 3:89521838_TTTG_T | 3 | 89521838 | GTEx/v8 | Brain Cerebellum | EPHA3 | T | 4.47555E-06 | -0.325467 | 0.000131411 |
| rs6797260 | 3 | 89302486 | GTEx/v8 | Artery Tibial | EPHA3 | T | 4.47908E-06 | -0.139789 | 7.80657E-07 |
| rs9866618 | 3 | 89319577 | GTEx/v8 | Artery Tibial | EPHA3 | C | 4.49777E-06 | 0.146776 | 7.80657E-07 |
| rs6771054 | 3 | 89489529 | GTEx/v8 | Brain Cerebellum | EPHA3 | C | 4.69366E-06 | -0.293501 | 0.000131411 |
| rs7644070 | 3 | 89370107 | GTEx/v8 | Brain Cerebellum | EPHA3 | A | 4.75765E-06 | -0.288734 | 0.000131411 |
| rs113895673 | 3 | 89917796 | GTEx/v8 | Pancreas | EPHA3 | C | 4.8159E-06 | 0.292706 | 3.49803E-07 |
| rs62274986 | 3 | 89327148 | GTEx/v8 | Artery Tibial | EPHA3 | A | 4.83202E-06 | 0.147343 | 7.80657E-07 |
| rs6782527 | 3 | 89442369 | GTEx/v8 | Artery Tibial | EPHA3 | T | 4.95543E-06 | 0.1499 | 7.80657E-07 |
| rs55787722 | 3 | 90012103 | GTEx/v8 | Artery Tibial | EPHA3 | T | 5.00725E-06 | 0.14935 | 7.80657E-07 |
| rs62274981 | 3 | 89310605 | GTEx/v8 | Artery Tibial | EPHA3 | T | 5.03773E-06 | 0.145415 | 7.80657E-07 |
| rs17800347 | 3 | 89314429 | GTEx/v8 | Artery Tibial | EPHA3 | G | 5.03773E-06 | 0.145415 | 7.80657E-07 |
| rs993186 | 3 | 89314862 | GTEx/v8 | Artery Tibial | EPHA3 | G | 5.03773E-06 | 0.145415 | 7.80657E-07 |
| rs13100388 | 3 | 89383148 | GTEx/v8 | Brain Cerebellum | EPHA3 | A | 5.04914E-06 | -0.284963 | 0.000131411 |
| rs6419884 | 3 | 89279467 | GTEx/v8 | Brain Cerebellum | EPHA3 | T | 5.19678E-06 | 0.292251 | 0.000131411 |
| rs9833400 | 3 | 89389507 | GTEx/v8 | Brain Cerebellum | EPHA3 | G | 5.2433E-06 | -0.286472 | 0.000131411 |
| rs7619303 | 3 | 89552554 | GTEx/v8 | Artery Tibial | EPHA3 | A | 5.32619E-06 | 0.145109 | 7.80657E-07 |
| rs1036285 | 3 | 89544061 | GTEx/v8 | Artery Tibial | EPHA3 | C | 5.391E-06 | 0.145676 | 7.80657E-07 |
| rs17026984 | 3 | 89544978 | GTEx/v8 | Artery Tibial | EPHA3 | A | 5.391E-06 | 0.145676 | 7.80657E-07 |
| rs111618480 | 3 | 89552134 | GTEx/v8 | Artery Tibial | EPHA3 | C | 5.391E-06 | 0.145676 | 7.80657E-07 |
| rs62278164 | 3 | 89292567 | GTEx/v8 | Brain Cerebellum | EPHA3 | C | 5.41224E-06 | -0.317954 | 0.000131411 |
| rs9883541 | 3 | 89294657 | GTEx/v8 | Brain Cerebellum | EPHA3 | C | 5.41224E-06 | -0.317954 | 0.000131411 |
| rs62274982 | 3 | 89310731 | GTEx/v8 | Artery Tibial | EPHA3 | A | 5.71992E-06 | 0.146085 | 7.80657E-07 |
| rs2137488 | 3 | 89308704 | GTEx/v8 | Artery Tibial | EPHA3 | G | 5.78921E-06 | 0.145371 | 7.80657E-07 |
| rs12634269 | 3 | 89532768 | GTEx/v8 | Artery Tibial | EPHA3 | C | 5.8533E-06 | 0.146048 | 7.80657E-07 |
| rs62278163 | 3 | 89291836 | GTEx/v8 | Artery Tibial | EPHA3 | T | 5.90756E-06 | 0.14456 | 7.80657E-07 |
| rs6786328 | 3 | 89298926 | GTEx/v8 | Artery Tibial | EPHA3 | G | 5.90756E-06 | 0.14456 | 7.80657E-07 |
| rs62278168 | 3 | 89301573 | GTEx/v8 | Artery Tibial | EPHA3 | G | 5.90756E-06 | 0.14456 | 7.80657E-07 |
| rs112407114 | 3 | 89797398 | GTEx/v8 | Brain Cerebellum | EPHA3 | C | 6.08314E-06 | -0.333729 | 0.000131411 |
| rs62275018 | 3 | 89423863 | GTEx/v8 | Brain Cerebellum | EPHA3 | G | 6.32511E-06 | -0.305486 | 0.000131411 |
| rs9310121 | 3 | 89378987 | GTEx/v8 | Brain Cerebellum | EPHA3 | G | 6.34498E-06 | -0.28251 | 0.000131411 |
| rs9310122 | 3 | 89379408 | GTEx/v8 | Brain Cerebellum | EPHA3 | G | 6.34498E-06 | -0.28251 | 0.000131411 |
| rs9846844 | 3 | 89379639 | GTEx/v8 | Brain Cerebellum | EPHA3 | A | 6.34498E-06 | -0.28251 | 0.000131411 |
| rs73153293 | 3 | 89756242 | GTEx/v8 | Brain Cerebellum | EPHA3 | T | 6.37713E-06 | -0.339017 | 0.000131411 |
| rs145565219 | 3 | 89936402 | GTEx/v8 | Artery Tibial | EPHA3 | A | 6.39268E-06 | 0.153684 | 7.80657E-07 |
| rs7613412 | 3 | 89724217 | GTEx/v8 | Brain Cerebellum | EPHA3 | T | 6.6714E-06 | -0.290591 | 0.000131411 |
| rs62278164 | 3 | 89292567 | GTEx/v8 | Artery Tibial | EPHA3 | C | 6.71388E-06 | 0.145209 | 7.80657E-07 |
| rs9883541 | 3 | 89294657 | GTEx/v8 | Artery Tibial | EPHA3 | C | 6.71388E-06 | 0.145209 | 7.80657E-07 |
| rs73145304 | 3 | 89568989 | GTEx/v8 | Artery Tibial | EPHA3 | T | 7.14309E-06 | 0.145168 | 7.80657E-07 |
| rs73145310 | 3 | 89578097 | GTEx/v8 | Artery Tibial | EPHA3 | C | 7.14309E-06 | 0.145168 | 7.80657E-07 |
| rs73145312 | 3 | 89578575 | GTEx/v8 | Artery Tibial | EPHA3 | T | 7.14309E-06 | 0.145168 | 7.80657E-07 |
| rs73145320 | 3 | 89586535 | GTEx/v8 | Artery Tibial | EPHA3 | C | 7.14309E-06 | 0.145168 | 7.80657E-07 |
| rs13075167 | 3 | 89303172 | GTEx/v8 | Artery Tibial | EPHA3 | C | 7.14359E-06 | -0.136941 | 7.80657E-07 |
| rs113895673 | 3 | 89917796 | GTEx/v8 | Artery Tibial | EPHA3 | C | 7.21958E-06 | 0.152514 | 7.80657E-07 |
| rs9862638 | 3 | 89373598 | GTEx/v8 | Artery Tibial | EPHA3 | C | 7.31028E-06 | 0.125044 | 7.80657E-07 |
| rs1398532 | 3 | 89367901 | GTEx/v8 | Brain Cerebellum | EPHA3 | C | 7.58961E-06 | -0.281373 | 0.000131411 |
| rs59969878 | 3 | 89369856 | GTEx/v8 | Brain Cerebellum | EPHA3 | A | 7.58961E-06 | -0.281373 | 0.000131411 |
| rs993187 | 3 | 89314820 | GTEx/v8 | Pancreas | EPHA3 | G | 7.78447E-06 | 0.274968 | 3.49803E-07 |

| | | | | | | | | | |
|---|---|---|---|---|---|---|---|---|---|
| rs9866959 | 3 | 89379935 | GTEx/v8 | Brain Cerebellum | EPHA3 | C | 7.983E-06 | -0.279055 | 0.000131411 |
| rs7616322 | 3 | 89493283 | GTEx/v8 | Brain Cerebellum | EPHA3 | T | 8.01438E-06 | -0.320833 | 0.000131411 |
| rs7645408 | 3 | 89446132 | GTEx/v8 | Pancreas | EPHA3 | A | 8.33515E-06 | 0.250421 | 3.49803E-07 |
| rs28623022 | 3 | 89448936 | GTEx/v8 | Pancreas | EPHA3 | A | 8.33515E-06 | 0.250421 | 3.49803E-07 |
| rs113895673 | 3 | 89917796 | GTEx/v8 | Brain Cerebellum | EPHA3 | C | 8.55292E-06 | -0.324874 | 0.000131411 |
| 3:89924704_TG_T | 3 | 89924704 | GTEx/v8 | Brain Cerebellum | EPHA3 | T | 8.55292E-06 | -0.324874 | 0.000131411 |
| rs145565219 | 3 | 89936402 | GTEx/v8 | Brain Cerebellum | EPHA3 | A | 8.55292E-06 | -0.324874 | 0.000131411 |
| rs9848971 | 3 | 89279394 | GTEx/v8 | Artery Tibial | EPHA3 | G | 8.76038E-06 | 0.142057 | 7.80657E-07 |
| rs62275018 | 3 | 89423863 | GTEx/v8 | Pancreas | EPHA3 | G | 9.33406E-06 | 0.250575 | 3.49803E-07 |
| rs7615949 | 3 | 89432683 | GTEx/v8 | Pancreas | EPHA3 | G | 9.33406E-06 | 0.250575 | 3.49803E-07 |
| rs9873545 | 3 | 89377717 | GTEx/v8 | Brain Cerebellum | EPHA3 | G | 9.40869E-06 | -0.278237 | 0.000131411 |
| rs2063590 | 3 | 89323932 | GTEx/v8 | Pancreas | EPHA3 | C | 9.50719E-06 | 0.270994 | 3.49803E-07 |
| rs1512185 | 3 | 89473615 | GTEx/v8 | Artery Tibial | EPHA3 | A | 9.59551E-06 | 0.141066 | 7.80657E-07 |
| rs1398534 | 3 | 89329307 | GTEx/v8 | Artery Tibial | EPHA3 | G | 9.64593E-06 | 0.141256 | 7.80657E-07 |
| rs73153279 | 3 | 89727968 | GTEx/v8 | Brain Cerebellum | EPHA3 | C | 9.69777E-06 | -0.314789 | 0.000131411 |
| rs73153282 | 3 | 89728408 | GTEx/v8 | Brain Cerebellum | EPHA3 | T | 9.69777E-06 | -0.314789 | 0.000131411 |
| rs6551410 | 3 | 89442305 | GTEx/v8 | Brain Cerebellum | EPHA3 | A | 1.01844E-05 | -0.28014 | 0.000131411 |
| rs9990016 | 3 | 89364033 | GTEx/v8 | Brain Cerebellum | EPHA3 | A | 1.08374E-05 | -0.28036 | 0.000131411 |
| rs73149109 | 3 | 89567344 | GTEx/v8 | Artery Tibial | EPHA3 | A | 1.08887E-05 | 0.140148 | 7.80657E-07 |
| rs1080592 | 3 | 89567568 | GTEx/v8 | Artery Tibial | EPHA3 | C | 1.08887E-05 | 0.140148 | 7.80657E-07 |
| rs73137395 | 3 | 89478753 | GTEx/v8 | Artery Tibial | EPHA3 | C | 1.09323E-05 | 0.143333 | 7.80657E-07 |
| rs73139257 | 3 | 89482779 | GTEx/v8 | Artery Tibial | EPHA3 | G | 1.09323E-05 | 0.143333 | 7.80657E-07 |
| 3:89924704_TG_T | 3 | 89924704 | GTEx/v8 | Artery Tibial | EPHA3 | T | 1.09412E-05 | 0.149669 | 7.80657E-07 |
| rs17738248 | 3 | 89296363 | GTEx/v8 | Pancreas | EPHA3 | A | 1.09578E-05 | 0.272841 | 3.49803E-07 |
| rs73137384 | 3 | 89463290 | GTEx/v8 | Artery Tibial | EPHA3 | T | 1.17861E-05 | 0.140098 | 7.80657E-07 |
| rs2069183 | 3 | 89645017 | GTEx/v8 | Artery Tibial | EPHA3 | A | 1.20408E-05 | 0.144339 | 7.80657E-07 |
| rs954964 | 3 | 89355632 | GTEx/v8 | Brain Cerebellum | EPHA3 | G | 1.22093E-05 | -0.273862 | 0.000131411 |
| rs13088663 | 3 | 89360266 | GTEx/v8 | Brain Cerebellum | EPHA3 | T | 1.22093E-05 | -0.273862 | 0.000131411 |
| rs9839323 | 3 | 89190113 | GTEx/v8 | Lung | CGGBP1 | A | 1.23179E-05 | 0.183962 | 1.81917E-48 |
| rs993187 | 3 | 89314820 | GTEx/v8 | Brain Cerebellum | EPHA3 | G | 1.23204E-05 | -0.303884 | 0.000131411 |
| rs7615949 | 3 | 89432683 | GTEx/v8 | Artery Tibial | EPHA3 | G | 1.24482E-05 | 0.135002 | 7.80657E-07 |
| rs13323078 | 3 | 89168325 | GTEx/v8 | Lung | CGGBP1 | A | 1.27567E-05 | 0.182688 | 1.81917E-48 |
| rs145565219 | 3 | 89936402 | GTEx/v8 | Pancreas | EPHA3 | A | 1.31568E-05 | 0.28696 | 3.49803E-07 |
| rs9873545 | 3 | 89377717 | GTEx/v8 | Artery Tibial | EPHA3 | G | 1.32482E-05 | 0.122036 | 7.80657E-07 |
| rs73151349 | 3 | 89892899 | GTEx/v8 | Artery Tibial | EPHA3 | G | 1.33999E-05 | 0.148574 | 7.80657E-07 |
| rs9867299 | 3 | 89300497 | GTEx/v8 | Artery Tibial | EPHA3 | A | 1.35373E-05 | 0.122174 | 7.80657E-07 |
| rs1512183 | 3 | 89505054 | GTEx/v8 | Artery Tibial | EPHA3 | T | 1.38036E-05 | 0.140822 | 7.80657E-07 |
| rs73153283 | 3 | 89732193 | GTEx/v8 | Brain Cerebellum | EPHA3 | T | 1.40234E-05 | -0.309356 | 0.000131411 |
| rs752364173 | 3 | 89521820 | GTEx/v8 | Artery Tibial | EPHA3 | T | 1.40686E-05 | 0.138789 | 7.80657E-07 |
| 3:89521831_TGAAAG_T | 3 | 89521831 | GTEx/v8 | Artery Tibial | EPHA3 | T | 1.40686E-05 | 0.138789 | 7.80657E-07 |
| 3:89521838_TTTG_T | 3 | 89521838 | GTEx/v8 | Artery Tibial | EPHA3 | T | 1.40686E-05 | 0.138789 | 7.80657E-07 |
| rs66708608 | 3 | 89486373 | GTEx/v8 | Brain Cerebellum | EPHA3 | T | 1.46451E-05 | -0.319235 | 0.000131411 |
| rs7616322 | 3 | 89493283 | GTEx/v8 | Artery Tibial | EPHA3 | T | 1.47881E-05 | 0.138523 | 7.80657E-07 |
| rs12637608 | 3 | 89551301 | GTEx/v8 | Artery Tibial | EPHA3 | C | 1.48728E-05 | 0.125146 | 7.80657E-07 |
| rs56371328 | 3 | 89539124 | GTEx/v8 | Artery Tibial | EPHA3 | A | 1.50588E-05 | 0.139235 | 7.80657E-07 |
| rs4857498 | 3 | 89540977 | GTEx/v8 | Artery Tibial | EPHA3 | A | 1.50588E-05 | 0.139235 | 7.80657E-07 |
| rs4857499 | 3 | 89541330 | GTEx/v8 | Artery Tibial | EPHA3 | A | 1.50588E-05 | 0.139235 | 7.80657E-07 |
| rs12633609 | 3 | 89545727 | GTEx/v8 | Artery Tibial | EPHA3 | G | 1.50588E-05 | 0.139235 | 7.80657E-07 |
| rs1512188 | 3 | 89546183 | GTEx/v8 | Artery Tibial | EPHA3 | G | 1.50588E-05 | 0.139235 | 7.80657E-07 |

| | | | | | | | | | |
|---|---|---|---|---|---|---|---|---|---|
| rs73146943 | 3 | 89546728 | GTEx/v8 | Artery Tibial | EPHA3 | G | 1.50588E-05 | 0.139235 | 7.80657E-07 |
| rs12632281 | 3 | 89547621 | GTEx/v8 | Artery Tibial | EPHA3 | A | 1.50588E-05 | 0.139235 | 7.80657E-07 |
| rs73146947 | 3 | 89548135 | GTEx/v8 | Artery Tibial | EPHA3 | G | 1.50588E-05 | 0.139235 | 7.80657E-07 |
| rs73146950 | 3 | 89548243 | GTEx/v8 | Artery Tibial | EPHA3 | C | 1.50588E-05 | 0.139235 | 7.80657E-07 |
| rs1567657 | 3 | 89548519 | GTEx/v8 | Artery Tibial | EPHA3 | A | 1.50588E-05 | 0.139235 | 7.80657E-07 |
| rs1567658 | 3 | 89548606 | GTEx/v8 | Artery Tibial | EPHA3 | A | 1.50588E-05 | 0.139235 | 7.80657E-07 |
| rs73146959 | 3 | 89549970 | GTEx/v8 | Artery Tibial | EPHA3 | C | 1.50588E-05 | 0.139235 | 7.80657E-07 |
| rs73146960 | 3 | 89550445 | GTEx/v8 | Artery Tibial | EPHA3 | C | 1.50588E-05 | 0.139235 | 7.80657E-07 |
| rs112404167 | 3 | 89550630 | GTEx/v8 | Artery Tibial | EPHA3 | T | 1.50588E-05 | 0.139235 | 7.80657E-07 |
| rs12637511 | 3 | 89550844 | GTEx/v8 | Artery Tibial | EPHA3 | C | 1.50588E-05 | 0.139235 | 7.80657E-07 |
| rs73146972 | 3 | 89553348 | GTEx/v8 | Artery Tibial | EPHA3 | G | 1.50588E-05 | 0.139235 | 7.80657E-07 |
| rs17026990 | 3 | 89555257 | GTEx/v8 | Artery Tibial | EPHA3 | C | 1.50588E-05 | 0.139235 | 7.80657E-07 |
| rs73146976 | 3 | 89557065 | GTEx/v8 | Artery Tibial | EPHA3 | C | 1.50588E-05 | 0.139235 | 7.80657E-07 |
| rs1039991 | 3 | 89558311 | GTEx/v8 | Artery Tibial | EPHA3 | G | 1.50588E-05 | 0.139235 | 7.80657E-07 |
| rs73146980 | 3 | 89558578 | GTEx/v8 | Artery Tibial | EPHA3 | A | 1.50588E-05 | 0.139235 | 7.80657E-07 |
| rs73146983 | 3 | 89559082 | GTEx/v8 | Artery Tibial | EPHA3 | G | 1.50588E-05 | 0.139235 | 7.80657E-07 |
| rs138490449 | 3 | 89561661 | GTEx/v8 | Artery Tibial | EPHA3 | AT | 1.50588E-05 | 0.139235 | 7.80657E-07 |
| rs138183278 | 3 | 89564090 | GTEx/v8 | Artery Tibial | EPHA3 | CT | 1.50588E-05 | 0.139235 | 7.80657E-07 |
| rs6774870 | 3 | 89564891 | GTEx/v8 | Artery Tibial | EPHA3 | T | 1.50588E-05 | 0.139235 | 7.80657E-07 |
| rs113307255 | 3 | 89567826 | GTEx/v8 | Artery Tibial | EPHA3 | C | 1.50588E-05 | 0.139235 | 7.80657E-07 |
| rs73153282 | 3 | 89728408 | GTEx/v8 | Artery Tibial | EPHA3 | T | 1.51422E-05 | 0.141824 | 7.80657E-07 |
| rs73153241 | 3 | 89695319 | GTEx/v8 | Brain Cerebellum | EPHA3 | G | 1.52792E-05 | -0.306774 | 0.000131411 |
| rs73145317 | 3 | 89583712 | GTEx/v8 | Brain Cerebellum | EPHA3 | C | 1.53352E-05 | -0.364409 | 0.000131411 |
| rs10511145 | 3 | 89597679 | GTEx/v8 | Artery Tibial | EPHA3 | A | 1.56217E-05 | 0.125943 | 7.80657E-07 |
| 3:89598352_GGT_G | 3 | 89598352 | GTEx/v8 | Artery Tibial | EPHA3 | G | 1.56217E-05 | 0.125943 | 7.80657E-07 |
| rs17027018 | 3 | 89604774 | GTEx/v8 | Artery Tibial | EPHA3 | T | 1.56217E-05 | 0.125943 | 7.80657E-07 |
| rs2346837 | 3 | 89542599 | GTEx/v8 | Artery Tibial | EPHA3 | T | 1.58851E-05 | 0.139109 | 7.80657E-07 |
| rs35173319 | 3 | 89386361 | GTEx/v8 | Artery Tibial | EPHA3 | T | 1.60562E-05 | 0.122651 | 7.80657E-07 |
| rs73151349 | 3 | 89892899 | GTEx/v8 | Brain Cerebellum | EPHA3 | G | 1.61632E-05 | -0.318556 | 0.000131411 |
| rs73145338 | 3 | 89594767 | GTEx/v8 | Artery Tibial | EPHA3 | T | 1.61827E-05 | 0.125766 | 7.80657E-07 |
| rs7621573 | 3 | 89595252 | GTEx/v8 | Artery Tibial | EPHA3 | T | 1.61827E-05 | 0.125766 | 7.80657E-07 |
| rs1398531 | 3 | 89289709 | GTEx/v8 | Artery Tibial | EPHA3 | T | 1.65645E-05 | 0.136374 | 7.80657E-07 |
| rs11923303 | 3 | 89380944 | GTEx/v8 | Artery Tibial | EPHA3 | A | 1.6566E-05 | 0.122508 | 7.80657E-07 |
| rs2063590 | 3 | 89323932 | GTEx/v8 | Brain Cerebellum | EPHA3 | C | 1.67149E-05 | -0.303021 | 0.000131411 |
| rs9310119 | 3 | 89378042 | GTEx/v8 | Artery Tibial | EPHA3 | G | 1.73911E-05 | 0.121808 | 7.80657E-07 |
| rs59127701 | 3 | 89291916 | GTEx/v8 | Brain Cerebellum | EPHA3 | TA | 1.87081E-05 | -0.316913 | 0.000131411 |
| rs6795325 | 3 | 89642054 | GTEx/v8 | Artery Tibial | EPHA3 | C | 1.91183E-05 | 0.124312 | 7.80657E-07 |
| rs17026944 | 3 | 89521131 | GTEx/v8 | Artery Tibial | EPHA3 | A | 1.91548E-05 | 0.137791 | 7.80657E-07 |
| rs73139125 | 3 | 89524056 | GTEx/v8 | Artery Tibial | EPHA3 | G | 1.91548E-05 | 0.137791 | 7.80657E-07 |
| rs66708608 | 3 | 89486373 | GTEx/v8 | Artery Tibial | EPHA3 | T | 1.93534E-05 | 0.138853 | 7.80657E-07 |
| rs73153262 | 3 | 89719269 | GTEx/v8 | Brain Cerebellum | EPHA3 | T | 1.93534E-05 | -0.304517 | 0.000131411 |
| rs73146965 | 3 | 89552553 | GTEx/v8 | Artery Tibial | EPHA3 | G | 1.95703E-05 | 0.137552 | 7.80657E-07 |
| rs2117138 | 3 | 89531522 | GTEx/v8 | Artery Tibial | EPHA3 | G | 1.98743E-05 | 0.138908 | 7.80657E-07 |
| rs1398532 | 3 | 89367901 | GTEx/v8 | Artery Tibial | EPHA3 | C | 1.98916E-05 | 0.120029 | 7.80657E-07 |
| rs59969878 | 3 | 89369856 | GTEx/v8 | Artery Tibial | EPHA3 | A | 1.98916E-05 | 0.120029 | 7.80657E-07 |
| rs9833400 | 3 | 89389507 | GTEx/v8 | Artery Tibial | EPHA3 | G | 2.02599E-05 | 0.120321 | 7.80657E-07 |
| rs2048518 | 3 | 89533905 | GTEx/v8 | Artery Tibial | EPHA3 | C | 2.06431E-05 | 0.136366 | 7.80657E-07 |
| rs2048520 | 3 | 89534248 | GTEx/v8 | Artery Tibial | EPHA3 | C | 2.06431E-05 | 0.136366 | 7.80657E-07 |
| rs2048521 | 3 | 89534253 | GTEx/v8 | Artery Tibial | EPHA3 | A | 2.06431E-05 | 0.136366 | 7.80657E-07 |
| rs73144998 | 3 | 89534592 | GTEx/v8 | Artery Tibial | EPHA3 | T | 2.06431E-05 | 0.136366 | 7.80657E-07 |
| rs73145002 | 3 | 89537618 | GTEx/v8 | Artery Tibial | EPHA3 | C | 2.06431E-05 | 0.136366 | 7.80657E-07 |
| rs13316356 | 3 | 89309959 | GTEx/v8 | Brain Cerebellum | EPHA3 | A | 2.07124E-05 | -0.291246 | 0.000131411 |

| | | | | | | | | | |
|---|---|---|---|---|---|---|---|---|---|
| rs73145317 | 3 | 89583712 | GTEx/v8 | Artery Tibial | EPHA3 | C | 2.08052E-05 | 0.161437 | 7.80657E-07 |
| rs9864984 | 3 | 89428776 | GTEx/v8 | Artery Tibial | EPHA3 | T | 2.08171E-05 | 0.130228 | 7.80657E-07 |
| rs62275019 | 3 | 89433648 | GTEx/v8 | Artery Tibial | EPHA3 | G | 2.08171E-05 | 0.130228 | 7.80657E-07 |
| rs66499884 | 3 | 89659012 | GTEx/v8 | Artery Tibial | EPHA3 | G | 2.09777E-05 | 0.123858 | 7.80657E-07 |
| rs7644070 | 3 | 89370107 | GTEx/v8 | Artery Tibial | EPHA3 | A | 2.11239E-05 | 0.12014 | 7.80657E-07 |
| rs73153219 | 3 | 89625902 | GTEx/v8 | Artery Tibial | EPHA3 | T | 2.13843E-05 | 0.123247 | 7.80657E-07 |
| rs7635916 | 3 | 89633140 | GTEx/v8 | Artery Tibial | EPHA3 | A | 2.13843E-05 | 0.123247 | 7.80657E-07 |
| rs6551413 | 3 | 89640350 | GTEx/v8 | Artery Tibial | EPHA3 | G | 2.13843E-05 | 0.123247 | 7.80657E-07 |
| rs73153283 | 3 | 89732193 | GTEx/v8 | Artery Tibial | EPHA3 | T | 2.13896E-05 | 0.139189 | 7.80657E-07 |
| rs7619025 | 3 | 89368629 | GTEx/v8 | Brain Cerebellum | EPHA3 | A | 2.14141E-05 | -0.269323 | 0.000131411 |
| rs7645408 | 3 | 89446132 | GTEx/v8 | Artery Tibial | EPHA3 | A | 2.14353E-05 | 0.130754 | 7.80657E-07 |
| rs28623022 | 3 | 89448936 | GTEx/v8 | Artery Tibial | EPHA3 | A | 2.14353E-05 | 0.130754 | 7.80657E-07 |
| rs17738248 | 3 | 89296363 | GTEx/v8 | Brain Cerebellum | EPHA3 | A | 2.15453E-05 | -0.299737 | 0.000131411 |
| rs9832305 | 3 | 89392778 | GTEx/v8 | Artery Tibial | EPHA3 | C | 2.16588E-05 | 0.131238 | 7.80657E-07 |
| rs7632427 | 3 | 89534377 | GTEx/v8 | Artery Tibial | EPHA3 | C | 2.25104E-05 | 0.123488 | 7.80657E-07 |
| rs11918555 | 3 | 89541934 | GTEx/v8 | Artery Tibial | EPHA3 | T | 2.3178E-05 | 0.12223 | 7.80657E-07 |
| rs9825569 | 3 | 89431650 | GTEx/v8 | Artery Tibial | EPHA3 | C | 2.42845E-05 | 0.12901 | 7.80657E-07 |
| rs113450730 | 3 | 90135516 | GTEx/v8 | Pancreas | EPHA3 | G | 2.43056E-05 | 0.265272 | 3.49803E-07 |
| rs62275018 | 3 | 89423863 | GTEx/v8 | Artery Tibial | EPHA3 | G | 2.50397E-05 | 0.130801 | 7.80657E-07 |
| rs13076853 | 3 | 89382776 | GTEx/v8 | Brain Cerebellum | EPHA3 | C | 2.50427E-05 | -0.266047 | 0.000131411 |
| 3:89924704_TG_T | 3 | 89924704 | GTEx/v8 | Pancreas | EPHA3 | T | 2.51441E-05 | 0.272398 | 3.49803E-07 |
| rs9868785 | 3 | 90066423 | GTEx/v8 | Artery Tibial | EPHA3 | A | 2.55363E-05 | 0.130904 | 7.80657E-07 |
| rs28675360 | 3 | 89386151 | GTEx/v8 | Artery Tibial | EPHA3 | T | 2.58705E-05 | 0.117655 | 7.80657E-07 |
| rs2175670 | 3 | 89343044 | GTEx/v8 | Brain Cerebellum | EPHA3 | G | 2.59143E-05 | -0.28772 | 0.000131411 |
| rs9825569 | 3 | 89431650 | GTEx/v8 | Pancreas | EPHA3 | C | 2.60809E-05 | 0.232839 | 3.49803E-07 |
| rs66757381 | 3 | 89561048 | GTEx/v8 | Artery Tibial | EPHA3 | T | 2.63229E-05 | 0.121893 | 7.80657E-07 |
| rs724972 | 3 | 89581408 | GTEx/v8 | Artery Tibial | EPHA3 | T | 2.63229E-05 | 0.121893 | 7.80657E-07 |
| rs7652296 | 3 | 89587262 | GTEx/v8 | Artery Tibial | EPHA3 | G | 2.63229E-05 | 0.121893 | 7.80657E-07 |
| rs9990016 | 3 | 89364033 | GTEx/v8 | Artery Tibial | EPHA3 | A | 2.63903E-05 | 0.119686 | 7.80657E-07 |
| rs73153279 | 3 | 89727968 | GTEx/v8 | Artery Tibial | EPHA3 | C | 2.64971E-05 | 0.13768 | 7.80657E-07 |
| rs6776281 | 3 | 89290164 | GTEx/v8 | Brain Cerebellum | EPHA3 | A | 2.65931E-05 | -0.292638 | 0.000131411 |
| rs13100388 | 3 | 89383148 | GTEx/v8 | Artery Tibial | EPHA3 | A | 2.66441E-05 | 0.117524 | 7.80657E-07 |
| rs12631313 | 3 | 89528461 | GTEx/v8 | Artery Tibial | EPHA3 | T | 2.69776E-05 | 0.136535 | 7.80657E-07 |
| rs7650466 | 3 | 89530358 | GTEx/v8 | Artery Tibial | EPHA3 | T | 2.69776E-05 | 0.136535 | 7.80657E-07 |
| rs9866959 | 3 | 89379935 | GTEx/v8 | Artery Tibial | EPHA3 | C | 2.77804E-05 | 0.116908 | 7.80657E-07 |
| rs9864984 | 3 | 89428776 | GTEx/v8 | Pancreas | EPHA3 | T | 2.79974E-05 | 0.233258 | 3.49803E-07 |
| rs62275019 | 3 | 89433648 | GTEx/v8 | Pancreas | EPHA3 | G | 2.79974E-05 | 0.233258 | 3.49803E-07 |
| rs73153260 | 3 | 89718078 | GTEx/v8 | Artery Tibial | EPHA3 | A | 2.82023E-05 | 0.15922 | 7.80657E-07 |
| rs35448293 | 3 | 89386010 | GTEx/v8 | Artery Tibial | EPHA3 | G | 2.82312E-05 | 0.117986 | 7.80657E-07 |
| rs1520596 | 3 | 89645542 | GTEx/v8 | Artery Tibial | EPHA3 | C | 2.8462E-05 | 0.121829 | 7.80657E-07 |
| rs9817553 | 3 | 89384566 | GTEx/v8 | Artery Tibial | EPHA3 | G | 2.90737E-05 | 0.117745 | 7.80657E-07 |
| rs35511518 | 3 | 89383683 | GTEx/v8 | Artery Tibial | EPHA3 | C | 2.90995E-05 | 0.116405 | 7.80657E-07 |
| rs60759140 | 3 | 89499848 | GTEx/v8 | Artery Tibial | EPHA3 | C | 2.91486E-05 | 0.136034 | 7.80657E-07 |
| rs73153266 | 3 | 89720111 | GTEx/v8 | Brain Cerebellum | EPHA3 | G | 2.92949E-05 | -0.298481 | 0.000131411 |
| rs113288647 | 3 | 89720352 | GTEx/v8 | Brain Cerebellum | EPHA3 | T | 2.92949E-05 | -0.298481 | 0.000131411 |
| rs73154925 | 3 | 89859312 | GTEx/v8 | Artery Tibial | EPHA3 | C | 2.94586E-05 | 0.138919 | 7.80657E-07 |
| rs2346836 | 3 | 89498096 | GTEx/v8 | Artery Tibial | EPHA3 | A | 2.9937E-05 | 0.136087 | 7.80657E-07 |
| rs59541469 | 3 | 89501671 | GTEx/v8 | Artery Tibial | EPHA3 | T | 2.9937E-05 | 0.136087 | 7.80657E-07 |
| 3:89502123_TTAG_T | 3 | 89502123 | GTEx/v8 | Artery Tibial | EPHA3 | T | 2.9937E-05 | 0.136087 | 7.80657E-07 |
| rs6551450 | 3 | 90129328 | GTEx/v8 | Artery Tibial | EPHA3 | C | 3.03179E-05 | 0.12845 | 7.80657E-07 |
| rs9851954 | 3 | 89297904 | GTEx/v8 | Artery Tibial | EPHA3 | A | 3.07249E-05 | 0.131839 | 7.80657E-07 |
| rs73153260 | 3 | 89718078 | GTEx/v8 | Brain Cerebellum | EPHA3 | A | 3.12578E-05 | -0.356663 | 0.000131411 |

| | | | | | | | | | |
|---|---|---|---|---|---|---|---|---|---|
| rs73153252 | 3 | 89712192 | GTEx/v8 | Brain Cerebellum | EPHA3 | T | 3.13471E-05 | -0.294404 | 0.000131411 |
| rs7617395 | 3 | 89722541 | GTEx/v8 | Brain Cerebellum | EPHA3 | T | 3.13471E-05 | -0.294404 | 0.000131411 |
| rs1567731 | 3 | 89316027 | GTEx/v8 | Pancreas | EPHA3 | T | 3.15741E-05 | 0.246698 | 3.49803E-07 |
| rs73141104 | 3 | 89508390 | GTEx/v8 | Artery Tibial | EPHA3 | T | 3.18627E-05 | 0.136308 | 7.80657E-07 |
| rs9868782 | 3 | 89384121 | GTEx/v8 | Pancreas | EPHA3 | A | 3.19825E-05 | 0.23913 | 3.49803E-07 |
| rs9310121 | 3 | 89378987 | GTEx/v8 | Artery Tibial | EPHA3 | G | 3.20658E-05 | 0.117263 | 7.80657E-07 |
| rs6776281 | 3 | 89290164 | GTEx/v8 | Artery Tibial | EPHA3 | A | 3.20719E-05 | 0.13161 | 7.80657E-07 |
| rs73153241 | 3 | 89695319 | GTEx/v8 | Artery Tibial | EPHA3 | G | 3.42799E-05 | 0.137103 | 7.80657E-07 |
| rs73153258 | 3 | 89715585 | GTEx/v8 | Brain Cerebellum | EPHA3 | A | 3.43172E-05 | -0.298648 | 0.000131411 |
| rs9310122 | 3 | 89379408 | GTEx/v8 | Artery Tibial | EPHA3 | G | 3.45233E-05 | 0.117072 | 7.80657E-07 |
| rs60243237 | 3 | 89694053 | GTEx/v8 | Artery Tibial | EPHA3 | T | 3.53066E-05 | 0.120952 | 7.80657E-07 |
| rs6796339 | 3 | 89286159 | GTEx/v8 | Brain Cerebellum | EPHA3 | T | 3.56632E-05 | -0.28695 | 0.000131411 |
| rs62278161 | 3 | 89291713 | GTEx/v8 | Brain Cerebellum | EPHA3 | G | 3.56632E-05 | -0.28695 | 0.000131411 |
| rs58858051 | 3 | 89291909 | GTEx/v8 | Brain Cerebellum | EPHA3 | T | 3.56632E-05 | -0.28695 | 0.000131411 |
| rs6794975 | 3 | 89292824 | GTEx/v8 | Brain Cerebellum | EPHA3 | A | 3.56632E-05 | -0.28695 | 0.000131411 |
| rs4857497 | 3 | 89294149 | GTEx/v8 | Brain Cerebellum | EPHA3 | G | 3.56632E-05 | -0.28695 | 0.000131411 |
| rs9847087 | 3 | 89297308 | GTEx/v8 | Brain Cerebellum | EPHA3 | G | 3.56632E-05 | -0.28695 | 0.000131411 |
| rs17026816 | 3 | 89297593 | GTEx/v8 | Brain Cerebellum | EPHA3 | G | 3.56632E-05 | -0.28695 | 0.000131411 |
| rs9851954 | 3 | 89297904 | GTEx/v8 | Brain Cerebellum | EPHA3 | A | 3.56632E-05 | -0.28695 | 0.000131411 |
| rs6786204 | 3 | 89298813 | GTEx/v8 | Brain Cerebellum | EPHA3 | C | 3.56632E-05 | -0.28695 | 0.000131411 |
| rs6773815 | 3 | 89299059 | GTEx/v8 | Brain Cerebellum | EPHA3 | T | 3.56632E-05 | -0.28695 | 0.000131411 |
| rs9862195 | 3 | 89299752 | GTEx/v8 | Brain Cerebellum | EPHA3 | T | 3.56632E-05 | -0.28695 | 0.000131411 |
| rs62278166 | 3 | 89300779 | GTEx/v8 | Brain Cerebellum | EPHA3 | G | 3.56632E-05 | -0.28695 | 0.000131411 |
| rs62278170 | 3 | 89302617 | GTEx/v8 | Brain Cerebellum | EPHA3 | T | 3.56632E-05 | -0.28695 | 0.000131411 |
| rs28834186 | 3 | 89305272 | GTEx/v8 | Brain Cerebellum | EPHA3 | A | 3.56632E-05 | -0.28695 | 0.000131411 |
| rs62278171 | 3 | 89305928 | GTEx/v8 | Brain Cerebellum | EPHA3 | A | 3.56632E-05 | -0.28695 | 0.000131411 |
| rs9863058 | 3 | 89322805 | GTEx/v8 | Pancreas | EPHA3 | T | 3.69984E-05 | 0.241311 | 3.49803E-07 |
| rs6800041 | 3 | 89448196 | GTEx/v8 | Pancreas | EPHA3 | C | 3.74788E-05 | 0.228794 | 3.49803E-07 |
| rs9310120 | 3 | 89378413 | GTEx/v8 | Artery Tibial | EPHA3 | C | 3.90585E-05 | 0.11677 | 7.80657E-07 |
| rs12714734 | 3 | 90107488 | GTEx/v8 | Brain Cerebellum | EPHA3 | C | 3.96629E-05 | -0.310278 | 0.000131411 |
| rs9877921 | 3 | 90108491 | GTEx/v8 | Brain Cerebellum | EPHA3 | T | 3.96629E-05 | -0.310278 | 0.000131411 |
| rs9816940 | 3 | 90146579 | GTEx/v8 | Brain Cerebellum | EPHA3 | T | 3.96629E-05 | -0.310278 | 0.000131411 |
| rs9837548 | 3 | 90156142 | GTEx/v8 | Brain Cerebellum | EPHA3 | T | 3.96629E-05 | -0.310278 | 0.000131411 |
| rs6551451 | 3 | 90156286 | GTEx/v8 | Artery Tibial | EPHA3 | T | 4.00647E-05 | 0.126988 | 7.80657E-07 |
| rs12152372 | 3 | 89500365 | GTEx/v8 | Artery Tibial | EPHA3 | C | 4.02235E-05 | 0.132899 | 7.80657E-07 |
| rs2881488 | 3 | 89500488 | GTEx/v8 | Artery Tibial | EPHA3 | G | 4.13729E-05 | 0.132921 | 7.80657E-07 |
| rs9846844 | 3 | 89379639 | GTEx/v8 | Artery Tibial | EPHA3 | A | 4.16688E-05 | 0.115754 | 7.80657E-07 |
| rs73154916 | 3 | 89803713 | GTEx/v8 | Artery Tibial | EPHA3 | T | 4.24218E-05 | 0.136449 | 7.80657E-07 |
| rs73139302 | 3 | 89508186 | GTEx/v8 | Artery Tibial | EPHA3 | G | 4.3432E-05 | 0.133646 | 7.80657E-07 |
| rs62278170 | 3 | 89302617 | GTEx/v8 | Artery Tibial | EPHA3 | T | 4.39208E-05 | 0.129245 | 7.80657E-07 |
| rs28834186 | 3 | 89305272 | GTEx/v8 | Artery Tibial | EPHA3 | A | 4.39208E-05 | 0.129245 | 7.80657E-07 |
| rs62278171 | 3 | 89305928 | GTEx/v8 | Artery Tibial | EPHA3 | A | 4.39208E-05 | 0.129245 | 7.80657E-07 |
| rs73139290 | 3 | 89505343 | GTEx/v8 | Artery Tibial | EPHA3 | A | 4.46616E-05 | 0.133674 | 7.80657E-07 |
| rs73139296 | 3 | 89505734 | GTEx/v8 | Artery Tibial | EPHA3 | T | 4.46616E-05 | 0.133674 | 7.80657E-07 |
| rs1157608 | 3 | 89506464 | GTEx/v8 | Artery Tibial | EPHA3 | G | 4.46616E-05 | 0.133674 | 7.80657E-07 |
| rs10470576 | 3 | 90154964 | GTEx/v8 | Artery Tibial | EPHA3 | T | 4.6124E-05 | 0.126504 | 7.80657E-07 |
| rs62278161 | 3 | 89291713 | GTEx/v8 | Artery Tibial | EPHA3 | G | 4.63024E-05 | 0.129185 | 7.80657E-07 |
| rs58858051 | 3 | 89291909 | GTEx/v8 | Artery Tibial | EPHA3 | T | 4.63024E-05 | 0.129185 | 7.80657E-07 |

| | | | | | | | | | |
|---|---|---|---|---|---|---|---|---|---|
| rs4857497 | 3 | 89294149 | GTEx/v8 | Artery Tibial | EPHA3 | G | 4.63024E-05 | 0.129185 | 7.80657E-07 |
| rs9847087 | 3 | 89297308 | GTEx/v8 | Artery Tibial | EPHA3 | G | 4.63024E-05 | 0.129185 | 7.80657E-07 |
| rs17026816 | 3 | 89297593 | GTEx/v8 | Artery Tibial | EPHA3 | G | 4.63024E-05 | 0.129185 | 7.80657E-07 |
| rs6786204 | 3 | 89298813 | GTEx/v8 | Artery Tibial | EPHA3 | C | 4.63024E-05 | 0.129185 | 7.80657E-07 |
| rs6773815 | 3 | 89299059 | GTEx/v8 | Artery Tibial | EPHA3 | T | 4.63024E-05 | 0.129185 | 7.80657E-07 |
| rs9862195 | 3 | 89299752 | GTEx/v8 | Artery Tibial | EPHA3 | T | 4.63024E-05 | 0.129185 | 7.80657E-07 |
| rs6419884 | 3 | 89279467 | GTEx/v8 | Artery Tibial | EPHA3 | T | 4.90317E-05 | -0.120956 | 7.80657E-07 |
| rs9811041 | 3 | 89367555 | GTEx/v8 | Artery Tibial | EPHA3 | A | 4.97505E-05 | 0.115675 | 7.80657E-07 |
| rs9864344 | 3 | 90121955 | GTEx/v8 | Artery Tibial | EPHA3 | G | 5.13438E-05 | 0.124283 | 7.80657E-07 |
| rs9880313 | 3 | 89373637 | GTEx/v8 | Artery Tibial | EPHA3 | G | 5.25287E-05 | 0.111212 | 7.80657E-07 |
| rs6796339 | 3 | 89286159 | GTEx/v8 | Artery Tibial | EPHA3 | T | 5.25489E-05 | 0.128922 | 7.80657E-07 |
| rs2175670 | 3 | 89343044 | GTEx/v8 | Artery Tibial | EPHA3 | G | 5.32925E-05 | 0.128139 | 7.80657E-07 |
| rs62278164 | 3 | 89292567 | GTEx/v8 | Pancreas | EPHA3 | C | 5.51258E-05 | 0.241565 | 3.49803E-07 |
| rs9883541 | 3 | 89294657 | GTEx/v8 | Pancreas | EPHA3 | C | 5.51258E-05 | 0.241565 | 3.49803E-07 |
| rs6800041 | 3 | 89448196 | GTEx/v8 | Artery Tibial | EPHA3 | C | 5.5413E-05 | 0.123822 | 7.80657E-07 |
| rs2137488 | 3 | 89308704 | GTEx/v8 | Pancreas | EPHA3 | G | 5.58479E-05 | 0.239656 | 3.49803E-07 |
| rs9829401 | 3 | 89300610 | GTEx/v8 | Artery Tibial | EPHA3 | C | 5.604E-05 | 0.126952 | 7.80657E-07 |
| rs6768381 | 3 | 89358071 | GTEx/v8 | Brain Cerebellum | EPHA3 | C | 5.66641E-05 | -0.254543 | 0.000131411 |
| rs13316356 | 3 | 89309959 | GTEx/v8 | Artery Tibial | EPHA3 | A | 5.8345E-05 | 0.127116 | 7.80657E-07 |
| rs62274982 | 3 | 89310731 | GTEx/v8 | Pancreas | EPHA3 | A | 5.89355E-05 | 0.238939 | 3.49803E-07 |
| rs9866618 | 3 | 89319577 | GTEx/v8 | Pancreas | EPHA3 | C | 5.89355E-05 | 0.238939 | 3.49803E-07 |
| rs62274614 | 3 | 89159719 | GTEx/v8 | Lung | CGGBP1 | T | 5.98457E-05 | 0.159774 | 1.81917E-48 |
| rs73153266 | 3 | 89720111 | GTEx/v8 | Artery Tibial | EPHA3 | G | 5.98562E-05 | 0.131977 | 7.80657E-07 |
| rs1912965 | 3 | 89518332 | GTEx/v8 | Artery Tibial | EPHA3 | G | 6.09446E-05 | 0.13059 | 7.80657E-07 |
| rs113141104 | 3 | 89519238 | GTEx/v8 | Artery Tibial | EPHA3 | T | 6.09446E-05 | 0.13059 | 7.80657E-07 |
| rs73139105 | 3 | 89519651 | GTEx/v8 | Artery Tibial | EPHA3 | C | 6.09446E-05 | 0.13059 | 7.80657E-07 |
| rs112297761 | 3 | 89521602 | GTEx/v8 | Artery Tibial | EPHA3 | TC | 6.09446E-05 | 0.13059 | 7.80657E-07 |
| rs73139119 | 3 | 89522285 | GTEx/v8 | Artery Tibial | EPHA3 | T | 6.09446E-05 | 0.13059 | 7.80657E-07 |
| rs73139121 | 3 | 89522433 | GTEx/v8 | Artery Tibial | EPHA3 | G | 6.09446E-05 | 0.13059 | 7.80657E-07 |
| rs73139131 | 3 | 89524686 | GTEx/v8 | Artery Tibial | EPHA3 | G | 6.09446E-05 | 0.13059 | 7.80657E-07 |
| rs11397577 | 3 | 90081678 | GTEx/v8 | Artery Tibial | EPHA3 | AT | 6.18786E-05 | 0.12282 | 7.80657E-07 |
| rs35124509 | 3 | 89521693 | GTEx/v8 | Artery Tibial | EPHA3 | C | 6.27184E-05 | 0.115674 | 7.80657E-07 |
| rs9829401 | 3 | 89300610 | GTEx/v8 | Brain Cerebellum | EPHA3 | C | 6.43403E-05 | -0.278778 | 0.000131411 |
| rs73153258 | 3 | 89715585 | GTEx/v8 | Artery Tibial | EPHA3 | A | 6.46822E-05 | 0.132007 | 7.80657E-07 |
| rs113288647 | 3 | 89720352 | GTEx/v8 | Artery Tibial | EPHA3 | T | 6.46822E-05 | 0.132007 | 7.80657E-07 |
| rs62278166 | 3 | 89300779 | GTEx/v8 | Artery Tibial | EPHA3 | G | 6.53018E-05 | 0.126675 | 7.80657E-07 |
| rs9848971 | 3 | 89279394 | GTEx/v8 | Pancreas | EPHA3 | G | 6.80201E-05 | 0.237246 | 3.49803E-07 |
| rs62274986 | 3 | 89327148 | GTEx/v8 | Pancreas | EPHA3 | A | 6.81299E-05 | 0.236006 | 3.49803E-07 |
| rs138643396 | 3 | 89698092 | GTEx/v8 | Artery Tibial | EPHA3 | CATA | 6.83504E-05 | 0.115926 | 7.80657E-07 |
| rs9849053 | 3 | 90152502 | GTEx/v8 | Artery Tibial | EPHA3 | A | 6.87755E-05 | 0.123791 | 7.80657E-07 |
| rs6768381 | 3 | 89358071 | GTEx/v8 | Artery Tibial | EPHA3 | C | 6.90531E-05 | 0.112763 | 7.80657E-07 |
| rs73139281 | 3 | 89503890 | GTEx/v8 | Artery Tibial | EPHA3 | T | 6.91831E-05 | 0.129367 | 7.80657E-07 |
| rs73139284 | 3 | 89503958 | GTEx/v8 | Artery Tibial | EPHA3 | G | 6.91831E-05 | 0.129367 | 7.80657E-07 |
| rs7632502 | 3 | 89469453 | GTEx/v8 | Pancreas | EPHA3 | A | 6.92594E-05 | 0.197124 | 3.49803E-07 |
| rs112604674 | 3 | 89989681 | GTEx/v8 | Brain Cerebellum | EPHA3 | T | 6.99043E-05 | -0.338171 | 0.000131411 |
| rs62278163 | 3 | 89291836 | GTEx/v8 | Pancreas | EPHA3 | T | 7.08915E-05 | 0.237296 | 3.49803E-07 |
| rs6786328 | 3 | 89298926 | GTEx/v8 | Pancreas | EPHA3 | G | 7.08915E-05 | 0.237296 | 3.49803E-07 |
| rs62278168 | 3 | 89301573 | GTEx/v8 | Pancreas | EPHA3 | G | 7.08915E-05 | 0.237296 | 3.49803E-07 |
| rs7637504 | 3 | 89702469 | GTEx/v8 | Artery Tibial | EPHA3 | T | 7.31685E-05 | 0.11595 | 7.80657E-07 |
| rs6551415 | 3 | 89705161 | GTEx/v8 | Artery Tibial | EPHA3 | T | 7.31685E-05 | 0.11595 | 7.80657E-07 |
| rs73139286 | 3 | 89504295 | GTEx/v8 | Artery Tibial | EPHA3 | A | 7.60748E-05 | 0.129484 | 7.80657E-07 |
| rs62274981 | 3 | 89310605 | GTEx/v8 | Pancreas | EPHA3 | T | 7.6395E-05 | 0.234437 | 3.49803E-07 |
| rs17800347 | 3 | 89314429 | GTEx/v8 | Pancreas | EPHA3 | G | 7.6395E-05 | 0.234437 | 3.49803E-07 |

| | | | | | | | | | |
|---|---|---|---|---|---|---|---|---|---|
| rs993186 | 3 | 89314862 | GTEx/v8 | Pancreas | EPHA3 | G | 7.6395E-05 | 0.234437 | 3.49803E-07 |
| rs17800551 | 3 | 89317315 | GTEx/v8 | Pancreas | EPHA3 | T | 7.6395E-05 | 0.234437 | 3.49803E-07 |
| rs62274984 | 3 | 89318317 | GTEx/v8 | Pancreas | EPHA3 | A | 7.6395E-05 | 0.234437 | 3.49803E-07 |
| rs7617395 | 3 | 89722541 | GTEx/v8 | Artery Tibial | EPHA3 | T | 8.13187E-05 | 0.130073 | 7.80657E-07 |
| rs7650184 | 3 | 89530057 | GTEx/v8 | Artery Tibial | EPHA3 | A | 8.18324E-05 | 0.114079 | 7.80657E-07 |
| rs73154916 | 3 | 89803713 | GTEx/v8 | Brain Cerebellum | EPHA3 | T | 8.21604E-05 | -0.287448 | 0.000131411 |
| rs73154925 | 3 | 89859312 | GTEx/v8 | Brain Cerebellum | EPHA3 | C | 8.21604E-05 | -0.287448 | 0.000131411 |
| rs9832305 | 3 | 89392778 | GTEx/v8 | Pancreas | EPHA3 | C | 8.51643E-05 | 0.219124 | 3.49803E-07 |
| rs4857502 | 3 | 89324919 | GTEx/v8 | Pancreas | EPHA3 | A | 8.79202E-05 | 0.231574 | 3.49803E-07 |
| rs2137487 | 3 | 89325520 | GTEx/v8 | Pancreas | EPHA3 | G | 8.79202E-05 | 0.231574 | 3.49803E-07 |
| rs9875278 | 3 | 89335463 | GTEx/v8 | Pancreas | EPHA3 | T | 8.79202E-05 | 0.231574 | 3.49803E-07 |
| rs9821666 | 3 | 89339548 | GTEx/v8 | Pancreas | EPHA3 | T | 8.79202E-05 | 0.231574 | 3.49803E-07 |
| rs1512908 | 3 | 89340134 | GTEx/v8 | Pancreas | EPHA3 | A | 8.79202E-05 | 0.231574 | 3.49803E-07 |
| rs9852380 | 3 | 89341738 | GTEx/v8 | Pancreas | EPHA3 | G | 8.79202E-05 | 0.231574 | 3.49803E-07 |
| rs9852722 | 3 | 89341946 | GTEx/v8 | Pancreas | EPHA3 | C | 8.79202E-05 | 0.231574 | 3.49803E-07 |
| rs59381722 | 3 | 89701482 | GTEx/v8 | Artery Tibial | EPHA3 | G | 9.00303E-05 | 0.113931 | 7.80657E-07 |
| rs9877833 | 3 | 90138312 | GTEx/v8 | Artery Tibial | EPHA3 | A | 9.26656E-05 | 0.124891 | 7.80657E-07 |
| rs6794975 | 3 | 89292824 | GTEx/v8 | Artery Tibial | EPHA3 | A | 9.30529E-05 | 0.123722 | 7.80657E-07 |
| rs73153262 | 3 | 89719269 | GTEx/v8 | Artery Tibial | EPHA3 | T | 9.40759E-05 | 0.128978 | 7.80657E-07 |
| rs3792572 | 3 | 89456555 | GTEx/v8 | Pancreas | EPHA3 | A | 9.54529E-05 | 0.195036 | 3.49803E-07 |
| rs6806912 | 3 | 89751353 | GTEx/v8 | Artery Tibial | EPHA3 | C | 9.74849E-05 | 0.114539 | 7.80657E-07 |
| rs9862638 | 3 | 89373598 | GTEx/v8 | Brain Cerebellum | EPHA3 | C | 9.82701E-05 | -0.238562 | 0.000131411 |
| rs7637642 | 3 | 90034785 | GTEx/v8 | Artery Tibial | EPHA3 | G | 9.98799E-05 | 0.119569 | 7.80657E-07 |
| rs6796339 | 3 | 89286159 | GTEx/v8 | Pancreas | EPHA3 | T | 9.99846E-05 | 0.227202 | 3.49803E-07 |
| rs62278161 | 3 | 89291713 | GTEx/v8 | Pancreas | EPHA3 | G | 9.99846E-05 | 0.227202 | 3.49803E-07 |
| rs58858051 | 3 | 89291909 | GTEx/v8 | Pancreas | EPHA3 | T | 9.99846E-05 | 0.227202 | 3.49803E-07 |
| rs4857497 | 3 | 89294149 | GTEx/v8 | Pancreas | EPHA3 | G | 9.99846E-05 | 0.227202 | 3.49803E-07 |
| rs9847087 | 3 | 89297308 | GTEx/v8 | Pancreas | EPHA3 | G | 9.99846E-05 | 0.227202 | 3.49803E-07 |
| rs17026816 | 3 | 89297593 | GTEx/v8 | Pancreas | EPHA3 | G | 9.99846E-05 | 0.227202 | 3.49803E-07 |
| rs6786204 | 3 | 89298813 | GTEx/v8 | Pancreas | EPHA3 | C | 9.99846E-05 | 0.227202 | 3.49803E-07 |
| rs6773815 | 3 | 89299059 | GTEx/v8 | Pancreas | EPHA3 | T | 9.99846E-05 | 0.227202 | 3.49803E-07 |
| rs9862195 | 3 | 89299752 | GTEx/v8 | Pancreas | EPHA3 | T | 9.99846E-05 | 0.227202 | 3.49803E-07 |
| rs7619025 | 3 | 89368629 | GTEx/v8 | Artery Tibial | EPHA3 | A | 0.000102538 | 0.110193 | 7.80657E-07 |
| rs28620812 | 3 | 89305668 | GTEx/v8 | Pancreas | EPHA3 | T | 0.000103808 | 0.232001 | 3.49803E-07 |
| rs62278172 | 3 | 89306038 | GTEx/v8 | Pancreas | EPHA3 | A | 0.000103808 | 0.232001 | 3.49803E-07 |
| rs954964 | 3 | 89355632 | GTEx/v8 | Artery Tibial | EPHA3 | G | 0.000104111 | 0.108869 | 7.80657E-07 |
| rs1398534 | 3 | 89329307 | GTEx/v8 | Pancreas | EPHA3 | G | 0.000104464 | 0.229195 | 3.49803E-07 |
| rs6794975 | 3 | 89292824 | GTEx/v8 | Pancreas | EPHA3 | A | 0.000105087 | 0.223728 | 3.49803E-07 |
| rs4396913 | 3 | 90047036 | GTEx/v8 | Pancreas | EPHA3 | C | 0.000105226 | 0.237452 | 3.49803E-07 |
| rs1464463 | 3 | 90056380 | GTEx/v8 | Pancreas | EPHA3 | A | 0.000105226 | 0.237452 | 3.49803E-07 |
| rs73153252 | 3 | 89712192 | GTEx/v8 | Artery Tibial | EPHA3 | T | 0.000109319 | 0.128448 | 7.80657E-07 |
| rs73151380 | 3 | 90000532 | GTEx/v8 | Artery Tibial | EPHA3 | T | 0.000111011 | 0.144157 | 7.80657E-07 |
| rs13323078 | 3 | 89168325 | GTEx/v8 | Testis | CGGBP1 | A | 0.000111676 | 0.139843 | 4.33153E-08 |
| rs28708984 | 3 | 90041154 | GTEx/v8 | Artery Tibial | EPHA3 | G | 0.000115862 | 0.118686 | 7.80657E-07 |
| rs61039064 | 3 | 90044732 | GTEx/v8 | Artery Tibial | EPHA3 | G | 0.000115862 | 0.118686 | 7.80657E-07 |
| rs35968370 | 3 | 89466339 | GTEx/v8 | Pancreas | EPHA3 | T | 0.000118817 | 0.193052 | 3.49803E-07 |
| rs11921985 | 3 | 89467357 | GTEx/v8 | Pancreas | EPHA3 | G | 0.000118817 | 0.193052 | 3.49803E-07 |
| rs4974358 | 3 | 90141506 | GTEx/v8 | Artery Tibial | EPHA3 | G | 0.000123858 | 0.123031 | 7.80657E-07 |
| rs7653735 | 3 | 89722185 | GTEx/v8 | Artery Tibial | EPHA3 | A | 0.000124606 | 0.112649 | 7.80657E-07 |
| rs11925143 | 3 | 89723684 | GTEx/v8 | Artery Tibial | EPHA3 | A | 0.000124606 | 0.112649 | 7.80657E-07 |
| rs112604674 | 3 | 89989681 | GTEx/v8 | Artery Tibial | EPHA3 | T | 0.000126338 | 0.143135 | 7.80657E-07 |
| rs112149405 | 3 | 89886780 | GTEx/v8 | Artery Tibial | EPHA3 | C | 0.000132367 | 0.150956 | 7.80657E-07 |
| rs9811041 | 3 | 89367555 | GTEx/v8 | Brain Cerebellum | EPHA3 | A | 0.00013287 | -0.24166 | 0.000131411 |

| | | | | | | | | | |
|---|---|---|---|---|---|---|---|---|---|
| rs55787722 | 3 | 90012103 | GTEx/v8 | Brain Cerebellum | EPHA3 | T | 0.000144208 | -0.27232 | 0.000131411 |
| rs9839323 | 3 | 89190113 | GTEx/v8 | Testis | CGGBP1 | A | 0.000146528 | 0.137448 | 4.33153E-08 |
| rs6776281 | 3 | 89290164 | GTEx/v8 | Pancreas | EPHA3 | A | 0.000148275 | 0.221329 | 3.49803E-07 |
| rs9851954 | 3 | 89297904 | GTEx/v8 | Pancreas | EPHA3 | A | 0.000148275 | 0.221329 | 3.49803E-07 |
| rs9880313 | 3 | 89373637 | GTEx/v8 | Brain Cerebellum | EPHA3 | G | 0.000150532 | -0.230663 | 0.000131411 |
| rs67316928 | 3 | 89482234 | GTEx/v8 | Pancreas | EPHA3 | C | 0.000150556 | 0.194504 | 3.49803E-07 |
| rs1398531 | 3 | 89289709 | GTEx/v8 | Brain Cerebellum | EPHA3 | T | 0.000151381 | -0.264304 | 0.000131411 |
| rs13088663 | 3 | 89360266 | GTEx/v8 | Artery Tibial | EPHA3 | T | 0.000156515 | 0.105987 | 7.80657E-07 |
| rs55787722 | 3 | 90012103 | GTEx/v8 | Pancreas | EPHA3 | T | 0.000158209 | 0.240878 | 3.49803E-07 |
| rs12636275 | 3 | 89523038 | GTEx/v8 | Artery Tibial | EPHA3 | A | 0.000159547 | 0.109133 | 7.80657E-07 |
| rs6772953 | 3 | 89451721 | GTEx/v8 | Pancreas | EPHA3 | C | 0.000161016 | 0.188937 | 3.49803E-07 |
| rs62274614 | 3 | 89159719 | GTEx/v8 | Testis | CGGBP1 | T | 0.000161492 | 0.13353 | 4.33153E-08 |
| rs12714734 | 3 | 90107488 | GTEx/v8 | Pancreas | EPHA3 | C | 0.000165118 | 0.230208 | 3.49803E-07 |
| rs9877921 | 3 | 90108491 | GTEx/v8 | Pancreas | EPHA3 | T | 0.000165118 | 0.230208 | 3.49803E-07 |
| rs1512907 | 3 | 89355961 | GTEx/v8 | Brain Cerebellum | EPHA3 | C | 0.000169535 | -0.238335 | 0.000131411 |
| rs59127701 | 3 | 89291916 | GTEx/v8 | Pancreas | EPHA3 | TA | 0.000172713 | 0.239554 | 3.49803E-07 |
| rs4396913 | 3 | 90047036 | GTEx/v8 | Brain Cerebellum | EPHA3 | C | 0.000182088 | -0.281997 | 0.000131411 |
| rs1464463 | 3 | 90056380 | GTEx/v8 | Brain Cerebellum | EPHA3 | A | 0.000182088 | -0.281997 | 0.000131411 |
| rs9866945 | 3 | 89349818 | GTEx/v8 | Brain Cerebellum | EPHA3 | C | 0.000185539 | -0.238202 | 0.000131411 |
| rs7613412 | 3 | 89724217 | GTEx/v8 | Artery Tibial | EPHA3 | T | 0.000188152 | 0.109455 | 7.80657E-07 |
| rs13076853 | 3 | 89382776 | GTEx/v8 | Artery Tibial | EPHA3 | C | 0.000197313 | 0.105664 | 7.80657E-07 |
| rs872976 | 3 | 89326991 | GTEx/v8 | Artery Tibial | EPHA3 | A | 0.000215792 | -0.103644 | 7.80657E-07 |
| rs1520594 | 3 | 89839468 | GTEx/v8 | Brain Cerebellum | EPHA3 | A | 0.000215974 | -0.235442 | 0.000131411 |
| rs9310117 | 3 | 89275906 | GTEx/v8 | Brain Cerebellum | EPHA3 | C | 0.000216519 | 0.238833 | 0.000131411 |
| rs6771336 | 3 | 89293015 | GTEx/v8 | Brain Cerebellum | EPHA3 | T | 0.000216519 | -0.238833 | 0.000131411 |
| rs62274987 | 3 | 89334443 | GTEx/v8 | Brain Cerebellum | EPHA3 | A | 0.000221208 | -0.23515 | 0.000131411 |
| rs67316928 | 3 | 89482234 | GTEx/v8 | Artery Tibial | EPHA3 | C | 0.000251611 | 0.106047 | 7.80657E-07 |
| rs59127701 | 3 | 89291916 | GTEx/v8 | Artery Tibial | EPHA3 | TA | 0.000338775 | 0.122222 | 7.80657E-07 |
| rs60759458 | 3 | 90045885 | GTEx/v8 | Artery Tibial | EPHA3 | C | 0.000353926 | 0.110068 | 7.80657E-07 |
| rs1512907 | 3 | 89355961 | GTEx/v8 | Artery Tibial | EPHA3 | C | 0.000359544 | 0.101588 | 7.80657E-07 |
| rs13075167 | 3 | 89303172 | PsychENCODE | PsychENCODE eQTLs | EPHA3 | C | 0.000383092 | -0.0451166 | 0.0265128367062786 |
| rs6551410 | 3 | 89442305 | GTEx/v8 | Artery Tibial | EPHA3 | A | 0.000422515 | 0.101978 | 7.80657E-07 |
| rs13064615 | 3 | 89799433 | PsychENCODE | PsychENCODE eQTLs | EPHA3 | G | 0.00044599 | -0.0417326 | 0.0300805785656357 |
| rs7619303 | 3 | 89552554 | GTEx/v8 | Thyroid | EPHA3 | A | 0.000457637 | 0.10083 | 0.000440998 |
| rs62274987 | 3 | 89334443 | GTEx/v8 | Artery Tibial | EPHA3 | A | 0.000463358 | 0.100651 | 7.80657E-07 |
| rs55787722 | 3 | 90012103 | GTEx/v8 | Thyroid | EPHA3 | T | 0.000580418 | 0.101037 | 0.000440998 |
| rs4264747 | 3 | 89882102 | PsychENCODE | PsychENCODE eQTLs | EPHA3 | G | 0.000646709 | -0.0412393 | 0.0407642701527111 |
| rs6797260 | 3 | 89302486 | CMC | CMC SVA cis | EPHA3 | A | | 1.0 | 0.049 |
| rs13075167 | 3 | 89303172 | CMC | CMC SVA cis | EPHA3 | T | | 1.0 | 0.049 |
| rs12638746 | 3 | 89331055 | CMC | CMC SVA cis | EPHA3 | G | | 1.0 | 0.049 |
| rs2274224 | 10 | 96039597 | eQTLGen | eQTLGen cis eQTLs | NOC3L | C | 2.3506E-168 | 27.6564 | 0.0 |
| rs11187838 | 10 | 96038686 | eQTLGen | eQTLGen cis eQTLs | NOC3L | A | 2.7379E-167 | 27.5675 | 0.0 |
| rs10786156 | 10 | 96014622 | eQTLGen | eQTLGen cis eQTLs | NOC3L | G | 9.0949E-166 | 27.4404 | 0.0 |
| rs57866767 | 10 | 96023077 | eQTLGen | eQTLGen cis eQTLs | NOC3L | C | 5.1534E-165 | 27.3771 | 0.0 |
| rs7080472 | 10 | 96012950 | eQTLGen | eQTLGen cis eQTLs | NOC3L | T | 1.1918E-164 | 27.3464 | 0.0 |
| rs3891783 | 10 | 96015793 | eQTLGen | eQTLGen cis eQTLs | NOC3L | G | 2.3181E-163 | 27.2379 | 0.0 |
| rs2274224 | 10 | 96039597 | eQTLGen | eQTLGen cis eQTLs | HELLS | C | 1.5774E-32 | 2,5062E+06 | 0.0 |

| | | | | | | | | | |
|---|---|---|---|---|---|---|---|---|---|
| rs11187838 | 10 | 96038686 | eQTLGen | eQTLGen cis eQTLs | HELLS | A | 1.1038E-31 | 1,9080E+06 | 0.0 |
| rs7080472 | 10 | 96012950 | eQTLGen | eQTLGen cis eQTLs | HELLS | T | 2.1997E-31 | 1,6943E+06 | 0.0 |
| rs10786156 | 10 | 96014622 | eQTLGen | eQTLGen cis eQTLs | HELLS | G | 2.4504E-31 | 1,6600E+06 | 0.0 |
| rs2077218 | 10 | 96071561 | eQTLGen | eQTLGen cis eQTLs | HELLS | G | 2.6229E-31 | 1,6395E+06 | 0.0 |
| rs3891783 | 10 | 96015793 | eQTLGen | eQTLGen cis eQTLs | HELLS | G | 3.7006E-31 | 1,5318E+06 | 0.0 |
| rs57866767 | 10 | 96023077 | eQTLGen | eQTLGen cis eQTLs | HELLS | C | 4.9097E-31 | 1,4437E+06 | 0.0 |
| rs11187844 | 10 | 96056629 | eQTLGen | eQTLGen cis eQTLs | HELLS | A | 2.6845E-27 | 2,3119E+06 | 0.0 |
| rs2077218 | 10 | 96071561 | eQTLGen | eQTLGen cis eQTLs | TBC1D12 | G | 3.3973E-27 | -10.8011 | 0.0 |
| rs79193572 | 10 | 96119713 | eQTLGen | eQTLGen cis eQTLs | TBC1D12 | G | 2.1704E-26 | 1,6059E+06 | 0.0 |
| rs74825688 | 10 | 96119687 | eQTLGen | eQTLGen cis eQTLs | TBC1D12 | A | 2.2801E-26 | 10.625 | 0.0 |
| rs113544984 | 10 | 96127448 | eQTLGen | eQTLGen cis eQTLs | TBC1D12 | A | 2.8489E-26 | 1,5138E+06 | 0.0 |
| rs11187906 | 10 | 96124838 | eQTLGen | eQTLGen cis eQTLs | TBC1D12 | A | 3.4829E-26 | 1,4448E+06 | 0.0 |
| rs10509672 | 10 | 96115957 | eQTLGen | eQTLGen cis eQTLs | TBC1D12 | C | 3.8096E-26 | 1,4145E+06 | 0.0 |
| rs7101207 | 10 | 96122857 | eQTLGen | eQTLGen cis eQTLs | TBC1D12 | A | 4.1845E-26 | 1,3816E+06 | 0.0 |
| rs79746649 | 10 | 96120218 | eQTLGen | eQTLGen cis eQTLs | TBC1D12 | T | 4.2565E-26 | 1,3761E+06 | 0.0 |
| rs11187907 | 10 | 96125361 | eQTLGen | eQTLGen cis eQTLs | TBC1D12 | T | 5.3345E-26 | 1,2987E+06 | 0.0 |
| rs11187890 | 10 | 96102758 | eQTLGen | eQTLGen cis eQTLs | TBC1D12 | C | 5.7346E-26 | 1,2739E+06 | 0.0 |
| rs11187897 | 10 | 96106603 | eQTLGen | eQTLGen cis eQTLs | TBC1D12 | G | 6.6543E-26 | 1,2231E+06 | 0.0 |
| rs11187895 | 10 | 96106240 | eQTLGen | eQTLGen cis eQTLs | TBC1D12 | C | 8.3501E-26 | 1,1449E+06 | 0.0 |
| rs12217792 | 10 | 96062386 | eQTLGen | eQTLGen cis eQTLs | TBC1D12 | C | 2.4075E-25 | 10.403 | 0.0 |
| rs112252876 | 10 | 96123355 | eQTLGen | eQTLGen cis eQTLs | TBC1D12 | T | 2.4689E-25 | 7,6948E+05 | 0.0 |
| rs11187883 | 10 | 96096866 | eQTLGen | eQTLGen cis eQTLs | TBC1D12 | C | 4.22E-25 | 5,8211E+05 | 0.0 |
| rs11187882 | 10 | 96095861 | eQTLGen | eQTLGen cis eQTLs | TBC1D12 | C | 5.1026E-25 | 5,1600E+05 | 0.0 |
| rs7096883 | 10 | 96069149 | eQTLGen | eQTLGen cis eQTLs | TBC1D12 | A | 6.5107E-25 | 4,2980E+05 | 0.0 |
| rs11187877 | 10 | 96092121 | eQTLGen | eQTLGen cis eQTLs | TBC1D12 | A | 9.7611E-25 | 2,8772E+05 | 0.0 |
| rs11187870 | 10 | 96087866 | eQTLGen | eQTLGen cis eQTLs | TBC1D12 | C | 1.0962E-24 | 2,4681E+05 | 0.0 |
| rs7100626 | 10 | 96069674 | eQTLGen | eQTLGen cis eQTLs | TBC1D12 | A | 1.1616E-24 | 2,2636E+05 | 0.0 |
| rs11187852 | 10 | 96070132 | eQTLGen | eQTLGen cis eQTLs | TBC1D12 | A | 1.3016E-24 | 1,8618E+05 | 0.0 |
| rs752140 | 10 | 96071396 | eQTLGen | eQTLGen cis eQTLs | TBC1D12 | T | 1.3016E-24 | 10.241 | 0.0 |
| rs75409190 | 10 | 96072425 | eQTLGen | eQTLGen cis eQTLs | TBC1D12 | T | 1.4196E-24 | 1,5514E+05 | 0.0 |
| rs11187864 | 10 | 96082506 | eQTLGen | eQTLGen cis eQTLs | TBC1D12 | T | 1.4343E-24 | 1,5222E+05 | 0.0 |
| rs11187856 | 10 | 96076869 | eQTLGen | eQTLGen cis eQTLs | TBC1D12 | A | 1.4917E-24 | 1,3797E+05 | 0.0 |
| rs11187866 | 10 | 96085991 | eQTLGen | eQTLGen cis eQTLs | TBC1D12 | G | 1.9264E-24 | 4,7392E+04 | 0.0 |
| rs11187863 | 10 | 96081457 | eQTLGen | eQTLGen cis eQTLs | TBC1D12 | G | 2.1134E-24 | 1,4154E+04 | 0.0 |
| rs11187840 | 10 | 96050351 | eQTLGen | eQTLGen cis eQTLs | TBC1D12 | G | 2.3184E-24 | 10.185 | 0.0 |
| rs75017201 | 10 | 96055152 | eQTLGen | eQTLGen cis eQTLs | TBC1D12 | T | 3.1871E-24 | 10.1538 | 0.0 |
| rs11187847 | 10 | 96063440 | eQTLGen | eQTLGen cis eQTLs | TBC1D12 | G | 3.77E-24 | 10.1376 | 0.0 |
| rs11187845 | 10 | 96060198 | eQTLGen | eQTLGen cis eQTLs | TBC1D12 | A | 4.2536000000000004E-24 | 10.1258 | 0.0 |
| rs3781266 | 10 | 96052747 | eQTLGen | eQTLGen cis eQTLs | TBC1D12 | G | 1.5677E-23 | 2,9492E+06 | 0.0 |
| rs3740365 | 10 | 96053239 | eQTLGen | eQTLGen cis eQTLs | TBC1D12 | A | 2.2898E-23 | 2,8115E+06 | 0.0 |
| rs753724 | 10 | 96051417 | eQTLGen | eQTLGen cis eQTLs | TBC1D12 | T | 2.4225E-23 | 2,7914E+06 | 0.0 |
| rs12220091 | 10 | 96053689 | eQTLGen | eQTLGen cis eQTLs | TBC1D12 | T | 2.4371E-23 | 2,7889E+06 | 0.0 |
| rs11187842 | 10 | 96052511 | eQTLGen | eQTLGen cis eQTLs | TBC1D12 | T | 3.5754E-23 | 2,6490E+06 | 0.0 |

| | | | | | | | | | |
|---|---|---|---|---|---|---|---|---|---|
| rs7078844 | 10 | 96081140 | eQTLGen | eQTLGen cis eQTLs | HELLS | T | 4.4823E-23 | 2,5672E+06 | 0.0 |
| rs3740359 | 10 | 96100119 | eQTLGen | eQTLGen cis eQTLs | TBC1D12 | T | 6.3939E-22 | 9.623 | 0.0 |
| rs3740360 | 10 | 96025491 | eQTLGen | eQTLGen cis eQTLs | TBC1D12 | C | 1.301E-17 | 1,2917E+06 | 0.0 |
| rs17109875 | 10 | 96026575 | eQTLGen | eQTLGen cis eQTLs | TBC1D12 | C | 2.3589E-17 | 1,0404E+06 | 0.0 |
| rs11187837 | 10 | 96035980 | eQTLGen | eQTLGen cis eQTLs | TBC1D12 | C | 3.1089E-17 | 9,2245E+05 | 0.0 |
| rs3891783 | 10 | 96015793 | eQTLcatal ogue | BLUEPRINT ge T-cell | NOC3L | C | 8.90201E-17 | 0.0946773 | 4.451E-13 |
| rs2274224 | 10 | 96039597 | eQTLcatal ogue | BLUEPRINT ge T-cell | NOC3L | G | 9.14876E-17 | 0.0946486 | 4.57438E-13 |
| rs10786156 | 10 | 96014622 | eQTLcatal ogue | BLUEPRINT ge T-cell | NOC3L | C | 9.30907E-17 | 0.0946158 | 4.65454E-13 |
| rs11187838 | 10 | 96038686 | eQTLcatal ogue | BLUEPRINT ge T-cell | NOC3L | G | 9.34357E-17 | 0.094607 | 4.67178E-13 |
| rs7080472 | 10 | 96012950 | BIOSQTL | BIOS eQTL geneLevel | NOC3L | T | 1.2781E-16 | 8.2757861 | 0.0 |
| rs11187838 | 10 | 96038686 | BIOSQTL | BIOS eQTL geneLevel | NOC3L | A | 2.42798E-16 | 8.1989633 | 0.0 |
| rs57866767 | 10 | 96023077 | BIOSQTL | BIOS eQTL geneLevel | NOC3L | C | 3.3781E-16 | 8.1590575 | 0.0 |
| rs10786156 | 10 | 96014622 | BIOSQTL | BIOS eQTL geneLevel | NOC3L | G | 3.4918E-16 | 8.1550176 | 0.0 |
| rs3891783 | 10 | 96015793 | BIOSQTL | BIOS eQTL geneLevel | NOC3L | G | 3.4918E-16 | 8.1550176 | 0.0 |
| rs7080472 | 10 | 96012950 | eQTLcatal ogue | BLUEPRINT ge T-cell | NOC3L | G | 5.68476E-16 | 0.0922238 | 2.84238E-12 |
| rs3891783 | 10 | 96015793 | PsychEN CODE | PsychENCO DE eQTLs | NOC3L | G | 1.55666E-15 | 0.084497 | 6.05693225367462E-13 |
| rs7080472 | 10 | 96012950 | PsychEN CODE | PsychENCO DE eQTLs | NOC3L | T | 2.15698E-15 | 0.0843385 | 8.29043545618019E-13 |
| rs7096883 | 10 | 96069149 | eQTLcatal ogue | GEUVADIS ge LCL | PLCE1 | G | 2.57371E-15 | 0.354412 | 1.28686E-11 |
| rs57866767 | 10 | 96023077 | PsychEN CODE | PsychENCO DE eQTLs | NOC3L | C | 2.91938E-15 | 0.0836983 | 1.10909695717759E-12 |
| rs11187847 | 10 | 96063440 | eQTLcatal ogue | GEUVADIS ge LCL | PLCE1 | C | 3.28649E-15 | 0.347624 | 1.64325E-11 |
| rs11187842 | 10 | 96052511 | eQTLcatal ogue | GEUVADIS ge LCL | PLCE1 | C | 3.77209E-15 | 0.34516 | 1.88605E-11 |
| rs3781266 | 10 | 96052747 | eQTLcatal ogue | GEUVADIS ge LCL | PLCE1 | A | 3.77209E-15 | 0.34516 | 1.88605E-11 |
| rs3740365 | 10 | 96053239 | eQTLcatal ogue | GEUVADIS ge LCL | PLCE1 | T | 3.77209E-15 | 0.34516 | 1.88605E-11 |
| rs12220091 | 10 | 96053689 | eQTLcatal ogue | GEUVADIS ge LCL | PLCE1 | C | 3.77209E-15 | 0.34516 | 1.88605E-11 |
| rs11187845 | 10 | 96060198 | eQTLcatal ogue | GEUVADIS ge LCL | PLCE1 | C | 4.8482E-15 | 0.34562 | 2.4241E-11 |
| rs11187826 | 10 | 95988042 | eQTLGen | eQTLGen cis eQTLs | HELLS | G | 6.9055E-15 | 2,1778E+06 | 0.0 |
| rs7100626 | 10 | 96069674 | eQTLcatal ogue | GEUVADIS ge LCL | PLCE1 | C | 1.44145E-14 | 0.356819 | 7.20725E-11 |
| rs11187852 | 10 | 96070132 | eQTLcatal ogue | GEUVADIS ge LCL | PLCE1 | G | 1.44145E-14 | 0.356819 | 7.20725E-11 |
| rs11187877 | 10 | 96092121 | eQTLcatal ogue | GEUVADIS ge LCL | PLCE1 | G | 1.44145E-14 | 0.356819 | 7.20725E-11 |
| rs3740360 | 10 | 96025491 | eQTLGen | eQTLGen cis eQTLs | HELLS | C | 1.6881E-14 | 1,7625E+06 | 0.0 |
| rs11187863 | 10 | 96081457 | eQTLcatal ogue | GEUVADIS ge LCL | PLCE1 | T | 1.87895E-14 | 0.336752 | 9.39475E-11 |
| rs11187866 | 10 | 96085991 | eQTLcatal ogue | GEUVADIS ge LCL | PLCE1 | C | 1.87895E-14 | 0.336752 | 9.39475E-11 |
| rs11187883 | 10 | 96096866 | eQTLcatal ogue | GEUVADIS ge LCL | PLCE1 | T | 1.87895E-14 | 0.336752 | 9.39475E-11 |
| rs17109875 | 10 | 96026575 | eQTLGen | eQTLGen cis eQTLs | HELLS | C | 1.9726E-14 | 1,6898E+06 | 0.0 |
| rs752140 | 10 | 96071396 | eQTLcatal ogue | GEUVADIS ge LCL | PLCE1 | C | 2.1749E-14 | 0.337652 | 1.08745E-10 |
| rs769015184 | 10 | 96049708 | eQTLcatal ogue | GEUVADIS ge LCL | PLCE1 | AC | 2.76489E-14 | 0.339981 | 1.38245E-10 |
| rs11187840 | 10 | 96050351 | eQTLcatal ogue | GEUVADIS ge LCL | PLCE1 | A | 2.76489E-14 | 0.339981 | 1.38245E-10 |
| rs75017201 | 10 | 96055152 | eQTLcatal ogue | GEUVADIS ge LCL | PLCE1 | C | 2.76489E-14 | 0.339981 | 1.38245E-10 |
| rs34309017 | 10 | 96061619 | eQTLcatal ogue | GEUVADIS ge LCL | PLCE1 | ACTT | 3.52825E-14 | 0.340472 | 1.76413E-10 |
| rs12217792 | 10 | 96062386 | eQTLcatal ogue | GEUVADIS ge LCL | PLCE1 | T | 3.52825E-14 | 0.340472 | 1.76413E-10 |
| rs11187906 | 10 | 96124838 | eQTLcatal ogue | GEUVADIS ge LCL | PLCE1 | T | 4.8199E-14 | 0.36095 | 2.40995E-10 |
| rs113544984 | 10 | 96127448 | eQTLcatal ogue | GEUVADIS ge LCL | PLCE1 | C | 4.8199E-14 | 0.36095 | 2.40995E-10 |
| rs75409190 | 10 | 96072425 | eQTLcatal ogue | GEUVADIS ge LCL | PLCE1 | C | 5.25935E-14 | 0.351622 | 2.62967E-10 |

| | | | | | | | | | |
|---|---|---|---|---|---|---|---|---|---|
| rs771106747 | 10 | 96072873 | eQTLcatalogue | GEUVADISge LCL | PLCE1 | CTA | 5.25935E-14 | 0.351622 | 2.62967E-10 |
| rs749295068 | 10 | 96073508 | eQTLcatalogue | GEUVADISge LCL | PLCE1 | AGAT | 5.25935E-14 | 0.351622 | 2.62967E-10 |
| rs11187856 | 10 | 96076869 | eQTLcatalogue | GEUVADISge LCL | PLCE1 | G | 5.25935E-14 | 0.351622 | 2.62967E-10 |
| rs11187864 | 10 | 96082506 | eQTLcatalogue | GEUVADISge LCL | PLCE1 | C | 5.25935E-14 | 0.351622 | 2.62967E-10 |
| rs11187870 | 10 | 96087866 | eQTLcatalogue | GEUVADISge LCL | PLCE1 | G | 5.25935E-14 | 0.351622 | 2.62967E-10 |
| rs145707916 | 10 | 96092992 | eQTLcatalogue | GEUVADISge LCL | PLCE1 | G | 5.25935E-14 | 0.351622 | 2.62967E-10 |
| rs750148755 | 10 | 96095271 | eQTLcatalogue | GEUVADISge LCL | PLCE1 | CATTTT | 5.25935E-14 | 0.351622 | 2.62967E-10 |
| rs11187882 | 10 | 96095861 | eQTLcatalogue | GEUVADISge LCL | PLCE1 | A | 5.25935E-14 | 0.351622 | 2.62967E-10 |
| rs10509672 | 10 | 96115957 | BIOSQTL | BIOS eQTL geneLevel | TBC1D12 | C | 5.63913E-14 | 7.5163545 | 0.0 |
| rs7078844 | 10 | 96081140 | eQTLGen | eQTLGen cis eQTLs | TBC1D12 | T | 6.266E-14 | -7.5022 | 0.0 |
| rs74825688 | 10 | 96119687 | BIOSQTL | BIOS eQTL geneLevel | TBC1D12 | A | 6.33328E-14 | 7.501064 | 0.0 |
| rs79193572 | 10 | 96119713 | BIOSQTL | BIOS eQTL geneLevel | TBC1D12 | G | 6.33328E-14 | 7.501064 | 0.0 |
| rs79746649 | 10 | 96120218 | BIOSQTL | BIOS eQTL geneLevel | TBC1D12 | T | 6.33328E-14 | 7.501064 | 0.0 |
| rs11187890 | 10 | 96102758 | eQTLcatalogue | GEUVADISge LCL | PLCE1 | T | 6.3606E-14 | 0.358744 | 3.1803E-10 |
| rs11187890 | 10 | 96102758 | BIOSQTL | BIOS eQTL geneLevel | TBC1D12 | C | 6.43065E-14 | 7.4990833 | 0.0 |
| rs11187895 | 10 | 96106240 | BIOSQTL | BIOS eQTL geneLevel | TBC1D12 | C | 6.46015E-14 | 7.4985075 | 0.0 |
| rs11187897 | 10 | 96106603 | BIOSQTL | BIOS eQTL geneLevel | TBC1D12 | G | 6.46015E-14 | 7.4985075 | 0.0 |
| rs7101207 | 10 | 96122857 | eQTLcatalogue | GEUVADISge LCL | PLCE1 | G | 6.98985E-14 | 0.352103 | 3.49493E-10 |
| rs112252876 | 10 | 96123355 | eQTLcatalogue | GEUVADISge LCL | PLCE1 | C | 6.98985E-14 | 0.352103 | 3.49493E-10 |
| rs11187838 | 10 | 96038686 | PsychENCODE | PsychENCODE eQTLs | NOC3L | A | 7.16731E-14 | 0.0800871 | 2.4E-11 |
| rs113544984 | 10 | 96127448 | BIOSQTL | BIOS eQTL geneLevel | TBC1D12 | A | 9.62128E-14 | 7.4461691 | 0.0 |
| rs11187907 | 10 | 96125361 | BIOSQTL | BIOS eQTL geneLevel | TBC1D12 | T | 1.24034E-13 | 7.41242 | 0.0 |
| rs79746649 | 10 | 96120218 | eQTLcatalogue | GEUVADISge LCL | PLCE1 | C | 1.30501E-13 | 0.346373 | 6.52505E-10 |
| rs2274224 | 10 | 96039597 | GTEx/v8 | Cells Cultured fibroblasts | PLCE1-AS1 | C | 1.4491E-13 | -0.223116 | 3.86001E-30 |
| rs200197176 | 10 | 96055963 | eQTLcatalogue | GEUVADISge LCL | PLCE1 | C | 1.62082E-13 | 0.325176 | 8.1041E-10 |
| rs7080472 | 10 | 96012950 | GTEx/v8 | Cells Cultured fibroblasts | PLCE1-AS1 | T | 1.67957E-13 | -0.226956 | 3.86001E-30 |
| rs11187826 | 10 | 95988042 | eQTLcatalogue | GEUVADISge LCL | PLCE1 | A | 1.90711E-13 | 0.294603 | 9.53555E-10 |
| rs11187826 | 10 | 95988042 | eQTLGen | eQTLGen cis eQTLs | TBC1D12 | G | 2.0948E-13 | 5,5791E+05 | 0.0 |
| rs7101207 | 10 | 96122857 | BIOSQTL | BIOS eQTL geneLevel | TBC1D12 | A | 2.40687E-13 | 7.3240743 | 0.0 |
| rs112252876 | 10 | 96123355 | BIOSQTL | BIOS eQTL geneLevel | TBC1D12 | T | 2.40687E-13 | 7.3240743 | 0.0 |
| rs2077218 | 10 | 96071561 | BIOSQTL | BIOS eQTL geneLevel | TBC1D12 | G | 2.68338E-13 | -7.30937 | 0.0 |
| rs3740360 | 10 | 96025491 | eQTLcatalogue | GEUVADISge LCL | PLCE1 | A | 2.69319E-13 | 0.304952 | 1.34659E-09 |
| rs17109875 | 10 | 96026575 | eQTLcatalogue | GEUVADISge LCL | PLCE1 | T | 2.69319E-13 | 0.304952 | 1.34659E-09 |
| rs11187837 | 10 | 96035980 | eQTLcatalogue | GEUVADISge LCL | PLCE1 | T | 2.69319E-13 | 0.304952 | 1.34659E-09 |
| rs113544984 | 10 | 96127448 | eQTLGen | eQTLGen cis eQTLs | PDLIM1 | A | 2.9121E-13 | -7.2983 | 0.0 |
| rs10786156 | 10 | 96014622 | GTEx/v8 | Cells Cultured fibroblasts | PLCE1-AS1 | G | 3.41914E-13 | -0.220278 | 3.86001E-30 |
| rs3891783 | 10 | 96015793 | GTEx/v8 | Cells Cultured fibroblasts | PLCE1-AS1 | G | 3.41914E-13 | -0.220278 | 3.86001E-30 |
| rs57866767 | 10 | 96023077 | GTEx/v8 | Cells Cultured fibroblasts | PLCE1-AS1 | C | 3.41914E-13 | -0.220278 | 3.86001E-30 |
| rs11187895 | 10 | 96106240 | eQTLcatalogue | GEUVADISge LCL | PLCE1 | G | 4.57228E-13 | 0.341015 | 2.28614E-09 |
| rs11187897 | 10 | 96106603 | eQTLcatalogue | GEUVADISge LCL | PLCE1 | A | 4.57228E-13 | 0.341015 | 2.28614E-09 |

| | | | | | | | | | |
|---|---|---|---|---|---|---|---|---|---|
| rs778275808 | 10 | 96114212 | eQTLcatalogue | GEUVADISge LCL | PLCE1 | TCTC | 4.57228E-13 | 0.341015 | 2.28614E-09 |
| rs10509672 | 10 | 96115957 | eQTLcatalogue | GEUVADISge LCL | PLCE1 | G | 4.57228E-13 | 0.341015 | 2.28614E-09 |
| rs79193572 | 10 | 96119713 | eQTLcatalogue | GEUVADISge LCL | PLCE1 | A | 4.57228E-13 | 0.341015 | 2.28614E-09 |
| rs11187837 | 10 | 96035980 | eQTLGen | eQTLGen cis eQTLs | HELLS | C | 6.3849E-13 | 7,8530E+03 | 0.0 |
| rs3740359 | 10 | 96100119 | GTEx/v8 | Cells Cultured fibroblasts | PLCE1-AS1 | T | 1.0708E-12 | -0.356669 | 3.86001E-30 |
| rs11187907 | 10 | 96125361 | eQTLGen | eQTLGen cis eQTLs | PDLIM1 | T | 1.0715E-12 | -7.121 | 0.0 |
| rs11187890 | 10 | 96102758 | eQTLcatalogue | BLUEPRINTge monocyte | NOC3L | T | 1.08793E-12 | 0.300083 | 5.43965E-09 |
| rs778275808 | 10 | 96114212 | eQTLcatalogue | BLUEPRINTge monocyte | NOC3L | TCTC | 1.08984E-12 | 0.300056 | 5.4492E-09 |
| rs11187895 | 10 | 96106240 | eQTLcatalogue | BLUEPRINTge monocyte | NOC3L | G | 1.09049E-12 | 0.300052 | 5.45245E-09 |
| rs11187897 | 10 | 96106603 | eQTLcatalogue | BLUEPRINTge monocyte | NOC3L | A | 1.09049E-12 | 0.300052 | 5.45245E-09 |
| rs10509672 | 10 | 96115957 | eQTLcatalogue | BLUEPRINTge monocyte | NOC3L | G | 1.09049E-12 | 0.300052 | 5.45245E-09 |
| rs74825688 | 10 | 96119687 | eQTLcatalogue | BLUEPRINTge monocyte | NOC3L | G | 1.09049E-12 | 0.300052 | 5.45245E-09 |
| rs79193572 | 10 | 96119713 | eQTLcatalogue | BLUEPRINTge monocyte | NOC3L | A | 1.09049E-12 | 0.300052 | 5.45245E-09 |
| rs11187863 | 10 | 96081457 | BIOSQTL | BIOS eQTL geneLevel | TBC1D12 | G | 1.47096E-12 | 7.0772858 | 0.0 |
| rs11187864 | 10 | 96082506 | BIOSQTL | BIOS eQTL geneLevel | TBC1D12 | T | 1.47096E-12 | 7.0772858 | 0.0 |
| rs11187866 | 10 | 96085991 | BIOSQTL | BIOS eQTL geneLevel | TBC1D12 | G | 1.47096E-12 | 7.0772858 | 0.0 |
| rs11187870 | 10 | 96087866 | BIOSQTL | BIOS eQTL geneLevel | TBC1D12 | C | 1.47096E-12 | 7.0772858 | 0.0 |
| rs11187877 | 10 | 96092121 | BIOSQTL | BIOS eQTL geneLevel | TBC1D12 | A | 1.47096E-12 | 7.0772858 | 0.0 |
| rs11187895 | 10 | 96106240 | eQTLGen | eQTLGen cis eQTLs | PDLIM1 | C | 1.6108E-12 | -7.0644 | 0.0 |
| rs7100626 | 10 | 96069674 | BIOSQTL | BIOS eQTL geneLevel | TBC1D12 | A | 1.61777E-12 | 7.0640675 | 0.0 |
| rs11187852 | 10 | 96070132 | BIOSQTL | BIOS eQTL geneLevel | TBC1D12 | A | 1.61777E-12 | 7.0640675 | 0.0 |
| rs752140 | 10 | 96071396 | BIOSQTL | BIOS eQTL geneLevel | TBC1D12 | T | 1.61777E-12 | 7.0640675 | 0.0 |
| rs75409190 | 10 | 96072425 | BIOSQTL | BIOS eQTL geneLevel | TBC1D12 | T | 1.62477E-12 | 7.0635281 | 0.0 |
| rs11187856 | 10 | 96076869 | BIOSQTL | BIOS eQTL geneLevel | TBC1D12 | A | 1.62477E-12 | 7.0635281 | 0.0 |
| rs11187882 | 10 | 96095861 | BIOSQTL | BIOS eQTL geneLevel | TBC1D12 | C | 1.63181E-12 | 7.0629935 | 0.0 |
| rs11187883 | 10 | 96096866 | BIOSQTL | BIOS eQTL geneLevel | TBC1D12 | C | 1.63181E-12 | 7.0629935 | 0.0 |
| rs11187906 | 10 | 96124838 | eQTLGen | eQTLGen cis eQTLs | PDLIM1 | A | 1.665E-12 | -7.06 | 0.0 |
| rs7080472 | 10 | 96012950 | GTEx/v8 | Adrenal Gland | NOC3L | T | 1.68803E-12 | 0.425729 | 2.32596E-10 |
| rs79193572 | 10 | 96119713 | eQTLGen | eQTLGen cis eQTLs | PDLIM1 | G | 1.8573E-12 | -7.0447 | 0.0 |
| rs11187838 | 10 | 96038686 | GTEx/v8 | Cells Cultured fibroblasts | PLCE1-AS1 | A | 1.92734E-12 | -0.211574 | 3.86001E-30 |
| rs11187897 | 10 | 96106603 | eQTLGen | eQTLGen cis eQTLs | PDLIM1 | G | 2.0772E-12 | -7.029 | 0.0 |
| rs74825688 | 10 | 96119687 | eQTLcatalogue | GEUVADISge LCL | PLCE1 | G | 2.11766E-12 | 0.329638 | 1.05883E-08 |
| rs10509672 | 10 | 96115957 | eQTLGen | eQTLGen cis eQTLs | PDLIM1 | C | 2.2603E-12 | -7.0174 | 0.0 |
| rs74825688 | 10 | 96119687 | eQTLGen | eQTLGen cis eQTLs | PDLIM1 | A | 2.2831E-12 | -7.0159 | 0.0 |
| rs11187870 | 10 | 96087866 | eQTLGen | eQTLGen cis eQTLs | PDLIM1 | C | 2.5706E-12 | -6.9993 | 0.0 |
| rs11187890 | 10 | 96102758 | eQTLGen | eQTLGen cis eQTLs | PDLIM1 | C | 2.6602E-12 | -6.9945 | 0.0 |
| rs11187864 | 10 | 96082506 | GTEx/v8 | Cells EBV-transformed lymphocytes | PLCE1 | T | 2.83248E-12 | 0.986452 | 1.20882E-07 |
| rs11187870 | 10 | 96087866 | GTEx/v8 | Cells EBV-transformed lymphocytes | PLCE1 | C | 2.83248E-12 | 0.986452 | 1.20882E-07 |

| | | | | | | | | | |
|---|---|---|---|---|---|---|---|---|---|
| rs11187877 | 10 | 96092121 | GTEx/v8 | Cells EBV-transformed lymphocytes | PLCE1 | A | 2.83248E-12 | 0.986452 | 1.20882E-07 |
| rs145707916 | 10 | 96092992 | GTEx/v8 | Cells EBV-transformed lymphocytes | PLCE1 | GA | 2.83248E-12 | 0.986452 | 1.20882E-07 |
| rs750148755 | 10 | 96095271 | GTEx/v8 | Cells EBV-transformed lymphocytes | PLCE1 | C | 2.83248E-12 | 0.986452 | 1.20882E-07 |
| rs11187882 | 10 | 96095861 | GTEx/v8 | Cells EBV-transformed lymphocytes | PLCE1 | C | 2.83248E-12 | 0.986452 | 1.20882E-07 |
| rs11187890 | 10 | 96102758 | GTEx/v8 | Cells EBV-transformed lymphocytes | PLCE1 | C | 2.83248E-12 | 0.986452 | 1.20882E-07 |
| rs11187895 | 10 | 96106240 | GTEx/v8 | Cells EBV-transformed lymphocytes | PLCE1 | C | 2.83248E-12 | 0.986452 | 1.20882E-07 |
| rs11187897 | 10 | 96106603 | GTEx/v8 | Cells EBV-transformed lymphocytes | PLCE1 | G | 2.83248E-12 | 0.986452 | 1.20882E-07 |
| rs778275808 | 10 | 96114212 | GTEx/v8 | Cells EBV-transformed lymphocytes | PLCE1 | T | 2.83248E-12 | 0.986452 | 1.20882E-07 |
| rs10509672 | 10 | 96115957 | GTEx/v8 | Cells EBV-transformed lymphocytes | PLCE1 | C | 2.83248E-12 | 0.986452 | 1.20882E-07 |
| rs79193572 | 10 | 96119713 | GTEx/v8 | Cells EBV-transformed lymphocytes | PLCE1 | G | 2.83248E-12 | 0.986452 | 1.20882E-07 |
| rs79746649 | 10 | 96120218 | GTEx/v8 | Cells EBV-transformed lymphocytes | PLCE1 | T | 2.83248E-12 | 0.986452 | 1.20882E-07 |
| rs3891783 | 10 | 96015793 | GTEx/v8 | Adrenal Gland | NOC3L | G | 3.25993E-12 | 0.407875 | 2.32596E-10 |
| rs57866767 | 10 | 96023077 | GTEx/v8 | Adrenal Gland | NOC3L | C | 3.25993E-12 | 0.407875 | 2.32596E-10 |
| rs10786156 | 10 | 96014622 | GTEx/v8 | Adrenal Gland | NOC3L | G | 3.32979E-12 | 0.40716 | 2.32596E-10 |
| rs2274224 | 10 | 96039597 | GTEx/v8 | Adrenal Gland | NOC3L | C | 3.32979E-12 | 0.40716 | 2.32596E-10 |
| rs112252876 | 10 | 96123355 | eQTLGen | eQTLGen cis eQTLs | PDLIM1 | T | 3.4221E-12 | -6.959 | 0.0 |
| rs11187864 | 10 | 96082506 | eQTLGen | eQTLGen cis eQTLs | PDLIM1 | T | 3.6788E-12 | -6.9489 | 0.0 |
| rs11187883 | 10 | 96096866 | eQTLGen | eQTLGen cis eQTLs | HELLS | C | 4.6392E-12 | 2,6533E+06 | 0.0 |
| rs11187844 | 10 | 96056629 | eQTLcatalogue | GEUVADIS ge LCL | PLCE1 | C | 4.66285E-12 | 0.245293 | 2.33142E-08 |
| rs3740359 | 10 | 96100119 | eQTLGen | eQTLGen cis eQTLs | PDLIM1 | T | 4.9781E-12 | -6.9062 | 0.0 |
| rs11187840 | 10 | 96050351 | BIOSQTL | BIOS eQTL geneLevel | TBC1D12 | G | 5.02742E-12 | 6.9048625 | 0.0 |
| rs753724 | 10 | 96051417 | BIOSQTL | BIOS eQTL geneLevel | TBC1D12 | T | 5.02742E-12 | 6.9048625 | 0.0 |
| rs3740365 | 10 | 96053239 | BIOSQTL | BIOS eQTL geneLevel | TBC1D12 | A | 5.02742E-12 | 6.9048625 | 0.0 |
| rs12220091 | 10 | 96053689 | BIOSQTL | BIOS eQTL geneLevel | TBC1D12 | T | 5.02742E-12 | 6.9048625 | 0.0 |
| rs75017201 | 10 | 96055152 | BIOSQTL | BIOS eQTL geneLevel | TBC1D12 | T | 5.02742E-12 | 6.9048625 | 0.0 |
| rs11187863 | 10 | 96081457 | eQTLGen | eQTLGen cis eQTLs | HELLS | G | 5.4474E-12 | 2,5696E+06 | 0.0 |
| rs11187907 | 10 | 96125361 | eQTLGen | eQTLGen cis eQTLs | NOC3L | T | 5.7302E-12 | 2,5433E+06 | 0.0 |
| rs11187840 | 10 | 96050351 | GTEx/v8 | Cells EBV-transformed lymphocytes | PLCE1 | G | 5.80787E-12 | 0.9539 | 1.20882E-07 |
| rs75017201 | 10 | 96055152 | GTEx/v8 | Cells EBV-transformed lymphocytes | PLCE1 | T | 5.80787E-12 | 0.9539 | 1.20882E-07 |
| rs11187877 | 10 | 96092121 | eQTLGen | eQTLGen cis eQTLs | HELLS | A | 5.935E-12 | 2,5251E+06 | 0.0 |
| rs79746649 | 10 | 96120218 | eQTLGen | eQTLGen cis eQTLs | PDLIM1 | T | 5.9769E-12 | -6.8802 | 0.0 |
| rs7101207 | 10 | 96122857 | eQTLGen | eQTLGen cis eQTLs | NOC3L | A | 6.0614E-12 | 2,5141E+06 | 0.0 |
| rs7101207 | 10 | 96122857 | eQTLGen | eQTLGen cis eQTLs | PDLIM1 | A | 6.1643E-12 | -6.8757 | 0.0 |
| rs11187906 | 10 | 96124838 | eQTLGen | eQTLGen cis eQTLs | NOC3L | A | 6.2602E-12 | 2,4970E+06 | 0.0 |
| rs11187882 | 10 | 96095861 | eQTLGen | eQTLGen cis eQTLs | HELLS | C | 6.3397E-12 | 2,4907E+06 | 0.0 |
| rs113544984 | 10 | 96127448 | eQTLGen | eQTLGen cis eQTLs | NOC3L | A | 6.5108E-12 | 2,4769E+06 | 0.0 |
| rs10509672 | 10 | 96115957 | eQTLcatalogue | TwinsUK ge LCL | PLCE1 | G | 6.70783E-12 | 0.133806 | 3.35392E-08 |

| | | | | | | | | | |
|---|---|---|---|---|---|---|---|---|---|
| rs74825688 | 10 | 96119687 | eQTLcatalogue | TwinsUK ge LCL | PLCE1 | G | 6.70783E-12 | 0.133806 | 3.35392E-08 |
| rs79193572 | 10 | 96119713 | eQTLcatalogue | TwinsUK ge LCL | PLCE1 | A | 6.70783E-12 | 0.133806 | 3.35392E-08 |
| rs79746649 | 10 | 96120218 | eQTLcatalogue | TwinsUK ge LCL | PLCE1 | C | 6.70783E-12 | 0.133806 | 3.35392E-08 |
| rs2274224 | 10 | 96039597 | GTEx/v8 | Lung | NOC3L | C | 6.8562E-12 | 0.169146 | 6.99161E-23 |
| rs3891783 | 10 | 96015793 | GTEx/v8 | Lung | NOC3L | G | 7.03959E-12 | 0.169026 | 6.99161E-23 |
| rs57866767 | 10 | 96023077 | GTEx/v8 | Lung | NOC3L | C | 7.03959E-12 | 0.169026 | 6.99161E-23 |
| rs3781266 | 10 | 96052747 | BIOSQTL | BIOS eQTL geneLevel | TBC1D12 | G | 7.04201E-12 | 6.8569227 | 0.0 |
| rs11187842 | 10 | 96052511 | BIOSQTL | BIOS eQTL geneLevel | TBC1D12 | T | 7.48896E-12 | 6.8481774 | 0.0 |
| rs10786156 | 10 | 96014622 | GTEx/v8 | Lung | NOC3L | G | 7.60516E-12 | 0.168552 | 6.99161E-23 |
| rs11187845 | 10 | 96060198 | GTEx/v8 | Cells Cultured fibroblasts | PLCE1-AS1 | A | 7.6979E-12 | -0.362676 | 3.86001E-30 |
| rs11187847 | 10 | 96063440 | GTEx/v8 | Cells Cultured fibroblasts | PLCE1-AS1 | G | 7.6979E-12 | -0.362676 | 3.86001E-30 |
| rs11187838 | 10 | 96038686 | GTEx/v8 | Adrenal Gland | NOC3L | A | 8.06874E-12 | 0.400858 | 2.32596E-10 |
| rs11187845 | 10 | 96060198 | eQTLcatalogue | BLUEPRINT ge monocyte | NOC3L | C | 8.41476E-12 | 0.271954 | 4.20738E-08 |
| rs12217792 | 10 | 96062386 | eQTLcatalogue | BLUEPRINT ge monocyte | NOC3L | T | 8.41476E-12 | 0.271954 | 4.20738E-08 |
| rs11187847 | 10 | 96063440 | eQTLcatalogue | BLUEPRINT ge monocyte | NOC3L | C | 8.41476E-12 | 0.271954 | 4.20738E-08 |
| rs7096883 | 10 | 96069149 | eQTLcatalogue | BLUEPRINT ge monocyte | NOC3L | G | 8.41476E-12 | 0.271954 | 4.20738E-08 |
| rs11187866 | 10 | 96085991 | eQTLGen | eQTLGen cis eQTLs | HELLS | G | 8.6951E-12 | 2,3260E+06 | 0.0 |
| rs3740359 | 10 | 96100119 | BIOSQTL | BIOS eQTL geneLevel | TBC1D12 | T | 8.87869E-12 | 6.8237607 | 0.0 |
| rs11187856 | 10 | 96076869 | eQTLGen | eQTLGen cis eQTLs | HELLS | A | 9.335E-12 | 2,2880E+06 | 0.0 |
| 10:96112261_AAAAAAAC_A | 10 | 96112261 | eQTLcatalogue | GEUVADIS ge LCL | PLCE1 | AAAAAAAC | 1.01771E-11 | 0.313995 | 5.08855E-08 |
| rs112252876 | 10 | 96123355 | eQTLGen | eQTLGen cis eQTLs | NOC3L | T | 1.0741E-11 | 2,2146E+06 | 0.0 |
| rs11187852 | 10 | 96070132 | eQTLGen | eQTLGen cis eQTLs | HELLS | A | 1.0892E-11 | 2,2077E+06 | 0.0 |
| rs11187847 | 10 | 96063440 | BIOSQTL | BIOS eQTL geneLevel | TBC1D12 | G | 1.13699E-11 | 6.7880264 | 0.0 |
| rs7096883 | 10 | 96069149 | BIOSQTL | BIOS eQTL geneLevel | TBC1D12 | A | 1.13699E-11 | 6.7880264 | 0.0 |
| rs75409190 | 10 | 96072425 | eQTLGen | eQTLGen cis eQTLs | HELLS | T | 1.137E-11 | 2,1847E+06 | 0.0 |
| rs11187856 | 10 | 96076869 | eQTLcatalogue | BLUEPRINT ge monocyte | NOC3L | G | 1.17547E-11 | 0.273524 | 5.87735E-08 |
| rs7100626 | 10 | 96069674 | eQTLcatalogue | BLUEPRINT ge monocyte | NOC3L | C | 1.18126E-11 | 0.273463 | 5.9063E-08 |
| rs11187852 | 10 | 96070132 | eQTLcatalogue | BLUEPRINT ge monocyte | NOC3L | G | 1.18329E-11 | 0.273449 | 5.91645E-08 |
| rs752140 | 10 | 96071396 | eQTLcatalogue | BLUEPRINT ge monocyte | NOC3L | C | 1.18329E-11 | 0.273449 | 5.91645E-08 |
| rs75409190 | 10 | 96072425 | eQTLcatalogue | BLUEPRINT ge monocyte | NOC3L | C | 1.18329E-11 | 0.273449 | 5.91645E-08 |
| rs771106747 | 10 | 96072873 | eQTLcatalogue | BLUEPRINT ge monocyte | NOC3L | CTA | 1.18329E-11 | 0.273449 | 5.91645E-08 |
| rs749295068 | 10 | 96073508 | eQTLcatalogue | BLUEPRINT ge monocyte | NOC3L | AGAT | 1.18329E-11 | 0.273449 | 5.91645E-08 |
| rs11187863 | 10 | 96081457 | eQTLcatalogue | BLUEPRINT ge monocyte | NOC3L | T | 1.18329E-11 | 0.273449 | 5.91645E-08 |
| rs11187864 | 10 | 96082506 | eQTLcatalogue | BLUEPRINT ge monocyte | NOC3L | C | 1.18329E-11 | 0.273449 | 5.91645E-08 |
| rs11187866 | 10 | 96085991 | eQTLcatalogue | BLUEPRINT ge monocyte | NOC3L | C | 1.18329E-11 | 0.273449 | 5.91645E-08 |
| rs11187870 | 10 | 96087866 | eQTLcatalogue | BLUEPRINT ge monocyte | NOC3L | G | 1.18329E-11 | 0.273449 | 5.91645E-08 |

| | | | | | | | | | |
|---|---|---|---|---|---|---|---|---|---|
| rs11187877 | 10 | 96092121 | eQTLcatalogue | BLUEPRINT ge monocyte | NOC3L | G | 1.18329E-11 | 0.273449 | 5.91645E-08 |
| rs11187882 | 10 | 96095861 | eQTLcatalogue | BLUEPRINT ge monocyte | NOC3L | A | 1.18329E-11 | 0.273449 | 5.91645E-08 |
| rs34309017 | 10 | 96061619 | GTEx/v8 | Cells EBV-transformed lymphocytes | PLCE1 | A | 1.21124E-11 | 0.952596 | 1.20882E-07 |
| rs12217792 | 10 | 96062386 | GTEx/v8 | Cells EBV-transformed lymphocytes | PLCE1 | C | 1.21124E-11 | 0.952596 | 1.20882E-07 |
| rs7096883 | 10 | 96069149 | GTEx/v8 | Cells EBV-transformed lymphocytes | PLCE1 | A | 1.21124E-11 | 0.952596 | 1.20882E-07 |
| rs11187877 | 10 | 96092121 | eQTLGen | eQTLGen cis eQTLs | PDLIM1 | A | 1.2458E-11 | -6.7748 | 0.0 |
| rs12217792 | 10 | 96062386 | BIOSQTL | BIOS eQTL geneLevel | TBC1D12 | C | 1.25097E-11 | 6.7742715 | 0.0 |
| rs7100626 | 10 | 96069674 | eQTLGen | eQTLGen cis eQTLs | HELLS | A | 1.2807E-11 | 2,1218E+06 | 0.0 |
| rs11187906 | 10 | 96124838 | eQTLcatalogue | BLUEPRINT ge monocyte | NOC3L | T | 1.28242E-11 | 0.291148 | 6.4121E-08 |
| rs11187907 | 10 | 96125361 | eQTLcatalogue | BLUEPRINT ge monocyte | NOC3L | C | 1.28242E-11 | 0.291148 | 6.4121E-08 |
| rs113544984 | 10 | 96127448 | eQTLcatalogue | BLUEPRINT ge monocyte | NOC3L | C | 1.28242E-11 | 0.291148 | 6.4121E-08 |
| rs7101207 | 10 | 96122857 | eQTLcatalogue | BLUEPRINT ge monocyte | NOC3L | G | 1.29283E-11 | 0.291223 | 6.46415E-08 |
| rs7100626 | 10 | 96069674 | GTEx/v8 | Cells EBV-transformed lymphocytes | PLCE1 | A | 1.346E-11 | 0.959887 | 1.20882E-07 |
| rs11187852 | 10 | 96070132 | GTEx/v8 | Cells EBV-transformed lymphocytes | PLCE1 | A | 1.346E-11 | 0.959887 | 1.20882E-07 |
| rs75409190 | 10 | 96072425 | GTEx/v8 | Cells EBV-transformed lymphocytes | PLCE1 | T | 1.346E-11 | 0.959887 | 1.20882E-07 |
| rs771106747 | 10 | 96072873 | GTEx/v8 | Cells EBV-transformed lymphocytes | PLCE1 | C | 1.346E-11 | 0.959887 | 1.20882E-07 |
| rs749295068 | 10 | 96073508 | GTEx/v8 | Cells EBV-transformed lymphocytes | PLCE1 | A | 1.346E-11 | 0.959887 | 1.20882E-07 |
| rs11187856 | 10 | 96076869 | GTEx/v8 | Cells EBV-transformed lymphocytes | PLCE1 | A | 1.346E-11 | 0.959887 | 1.20882E-07 |
| rs752140 | 10 | 96071396 | eQTLGen | eQTLGen cis eQTLs | HELLS | T | 1.4723E-11 | 2,0484E+06 | 0.0 |
| rs74825688 | 10 | 96119687 | GTEx/v8 | Cells EBV-transformed lymphocytes | PLCE1 | A | 1.50065E-11 | 0.937552 | 1.20882E-07 |
| rs11187845 | 10 | 96060198 | BIOSQTL | BIOS eQTL geneLevel | TBC1D12 | A | 1.5135E-11 | 6.7467696 | 0.0 |
| rs11187856 | 10 | 96076869 | eQTLGen | eQTLGen cis eQTLs | PDLIM1 | A | 1.9462E-11 | -6.71 | 0.0 |
| rs3740359 | 10 | 96100119 | eQTLGen | eQTLGen cis eQTLs | NOC3L | T | 1.9489E-11 | 1,8990E+06 | 0.0 |
| rs7100626 | 10 | 96069674 | eQTLcatalogue | TwinsUK ge LCL | PLCE1 | C | 2.02485E-11 | 0.128196 | 1.01243E-07 |
| rs11187852 | 10 | 96070132 | eQTLcatalogue | TwinsUK ge LCL | PLCE1 | G | 2.02485E-11 | 0.128196 | 1.01243E-07 |
| rs752140 | 10 | 96071396 | eQTLcatalogue | TwinsUK ge LCL | PLCE1 | C | 2.02485E-11 | 0.128196 | 1.01243E-07 |
| rs75409190 | 10 | 96072425 | eQTLcatalogue | TwinsUK ge LCL | PLCE1 | C | 2.02485E-11 | 0.128196 | 1.01243E-07 |
| rs11187856 | 10 | 96076869 | eQTLcatalogue | TwinsUK ge LCL | PLCE1 | G | 2.02485E-11 | 0.128196 | 1.01243E-07 |
| rs11187863 | 10 | 96081457 | eQTLcatalogue | TwinsUK ge LCL | PLCE1 | T | 2.02485E-11 | 0.128196 | 1.01243E-07 |
| rs11187864 | 10 | 96082506 | eQTLcatalogue | TwinsUK ge LCL | PLCE1 | C | 2.02485E-11 | 0.128196 | 1.01243E-07 |
| rs11187866 | 10 | 96085991 | eQTLcatalogue | TwinsUK ge LCL | PLCE1 | C | 2.02485E-11 | 0.128196 | 1.01243E-07 |
| rs11187870 | 10 | 96087866 | eQTLcatalogue | TwinsUK ge LCL | PLCE1 | G | 2.02485E-11 | 0.128196 | 1.01243E-07 |
| rs11187877 | 10 | 96092121 | eQTLcatalogue | TwinsUK ge LCL | PLCE1 | G | 2.02485E-11 | 0.128196 | 1.01243E-07 |
| rs11187882 | 10 | 96095861 | eQTLcatalogue | TwinsUK ge LCL | PLCE1 | A | 2.02485E-11 | 0.128196 | 1.01243E-07 |
| rs11187883 | 10 | 96096866 | eQTLcatalogue | TwinsUK ge LCL | PLCE1 | T | 2.02485E-11 | 0.128196 | 1.01243E-07 |

| | | | | | | | | | |
|---|---|---|---|---|---|---|---|---|---|
| rs7101207 | 10 | 96122857 | eQTLcatalogue | TwinsUK ge LCL | PLCE1 | G | 2.19128E-11 | 0.133521 | 1.09564E-07 |
| rs112252876 | 10 | 96123355 | eQTLcatalogue | TwinsUK ge LCL | PLCE1 | C | 2.19128E-11 | 0.133521 | 1.09564E-07 |
| rs11187906 | 10 | 96124838 | eQTLcatalogue | TwinsUK ge LCL | PLCE1 | T | 2.19128E-11 | 0.133521 | 1.09564E-07 |
| rs11187907 | 10 | 96125361 | eQTLcatalogue | TwinsUK ge LCL | PLCE1 | C | 2.19128E-11 | 0.133521 | 1.09564E-07 |
| rs113544984 | 10 | 96127448 | eQTLcatalogue | TwinsUK ge LCL | PLCE1 | C | 2.19128E-11 | 0.133521 | 1.09564E-07 |
| rs75409190 | 10 | 96072425 | eQTLGen | eQTLGen cis eQTLs | PDLIM1 | T | 2.2135E-11 | -6.6911 | 0.0 |
| rs11187890 | 10 | 96102758 | eQTLcatalogue | TwinsUK ge LCL | PLCE1 | T | 2.49173E-11 | 0.128949 | 1.24586E-07 |
| rs11187895 | 10 | 96106240 | eQTLcatalogue | TwinsUK ge LCL | PLCE1 | G | 2.49173E-11 | 0.128949 | 1.24586E-07 |
| rs11187897 | 10 | 96106603 | eQTLcatalogue | TwinsUK ge LCL | PLCE1 | A | 2.49173E-11 | 0.128949 | 1.24586E-07 |
| rs11187882 | 10 | 96095861 | eQTLGen | eQTLGen cis eQTLs | PDLIM1 | C | 2.496E-11 | -6.6734 | 0.0 |
| rs7100626 | 10 | 96069674 | eQTLGen | eQTLGen cis eQTLs | PDLIM1 | A | 2.5234E-11 | -6.6719 | 0.0 |
| rs11187863 | 10 | 96081457 | eQTLGen | eQTLGen cis eQTLs | PDLIM1 | G | 3.0736E-11 | -6.6429 | 0.0 |
| rs11187907 | 10 | 96125361 | eQTLGen | eQTLGen cis eQTLs | HELLS | T | 3.4491E-11 | 1,5930E+06 | 0.0 |
| rs113544984 | 10 | 96127448 | GTEx/v8 | Cells EBV-transformed lymphocytes | PLCE1 | A | 3.47759E-11 | 0.940054 | 1.20882E-07 |
| rs11187883 | 10 | 96096866 | eQTLGen | eQTLGen cis eQTLs | PDLIM1 | C | 3.7055E-11 | -6.6153 | 0.0 |
| rs7101207 | 10 | 96122857 | eQTLGen | eQTLGen cis eQTLs | HELLS | A | 4.0021E-11 | 1,5126E+06 | 0.0 |
| rs7080472 | 10 | 96012950 | GTEx/v8 | Lung | NOC3L | T | 4.03452E-11 | 0.165385 | 6.99161E-23 |
| rs11187852 | 10 | 96070132 | eQTLGen | eQTLGen cis eQTLs | PDLIM1 | A | 4.3687E-11 | -6.5909 | 0.0 |
| rs2077218 | 10 | 96071561 | eQTLGen | eQTLGen cis eQTLs | NOC3L | G | 5.1754E-11 | -6.5657 | 0.0 |
| rs74825688 | 10 | 96119687 | eQTLGen | eQTLGen cis eQTLs | HELLS | A | 5.352E-11 | 1,3548E+06 | 0.0 |
| rs7101207 | 10 | 96122857 | GTEx/v8 | Cells EBV-transformed lymphocytes | PLCE1 | A | 5.68167E-11 | 0.931326 | 1.20882E-07 |
| rs112252876 | 10 | 96123355 | GTEx/v8 | Cells EBV-transformed lymphocytes | PLCE1 | T | 5.68167E-11 | 0.931326 | 1.20882E-07 |
| rs11187838 | 10 | 96038686 | GTEx/v8 | Lung | NOC3L | A | 5.94459E-11 | 0.161642 | 6.99161E-23 |
| rs752140 | 10 | 96071396 | eQTLGen | eQTLGen cis eQTLs | PDLIM1 | T | 5.9975E-11 | -6.5437 | 0.0 |
| rs112252876 | 10 | 96123355 | eQTLGen | eQTLGen cis eQTLs | HELLS | T | 6.0216E-11 | 1,2909E+06 | 0.0 |
| rs11187906 | 10 | 96124838 | BIOSQTL | BIOS eQTL geneLevel | TBC1D12 | A | 6.20974E-11 | 6.5386326 | 0.0 |
| rs11187906 | 10 | 96124838 | eQTLGen | eQTLGen cis eQTLs | HELLS | A | 6.7279E-11 | 1,2299E+06 | 0.0 |
| rs10509672 | 10 | 96115957 | eQTLGen | eQTLGen cis eQTLs | HELLS | C | 6.8274E-11 | 1,2219E+06 | 0.0 |
| rs753724 | 10 | 96051417 | GTEx/v8 | Cells Cultured fibroblasts | PLCE1-AS1 | T | 6.9572E-11 | -0.339785 | 3.86001E-30 |
| rs11187842 | 10 | 96052511 | GTEx/v8 | Cells Cultured fibroblasts | PLCE1-AS1 | T | 6.9572E-11 | -0.339785 | 3.86001E-30 |
| rs3781266 | 10 | 96052747 | GTEx/v8 | Cells Cultured fibroblasts | PLCE1-AS1 | G | 6.9572E-11 | -0.339785 | 3.86001E-30 |
| rs3740365 | 10 | 96053239 | GTEx/v8 | Cells Cultured fibroblasts | PLCE1-AS1 | A | 6.9572E-11 | -0.339785 | 3.86001E-30 |
| rs12220091 | 10 | 96053689 | GTEx/v8 | Cells Cultured fibroblasts | PLCE1-AS1 | T | 6.9572E-11 | -0.339785 | 3.86001E-30 |
| rs11187866 | 10 | 96085991 | eQTLGen | eQTLGen cis eQTLs | PDLIM1 | G | 7.5051E-11 | -6.5102 | 0.0 |
| rs3740359 | 10 | 96100119 | eQTLcatalogue | GEUVADIS ge LCL | PLCE1 | C | 7.60307E-11 | 0.257729 | 3.80154E-07 |
| rs79746649 | 10 | 96120218 | eQTLGen | eQTLGen cis eQTLs | HELLS | T | 7.9576E-11 | 1,1379E+06 | 0.0 |
| rs11187895 | 10 | 96106240 | eQTLGen | eQTLGen cis eQTLs | HELLS | C | 8.2265E-11 | 1,1193E+06 | 0.0 |
| rs79193572 | 10 | 96119713 | eQTLGen | eQTLGen cis eQTLs | HELLS | G | 8.493E-11 | 1,1021E+06 | 0.0 |
| rs11187897 | 10 | 96106603 | eQTLGen | eQTLGen cis eQTLs | HELLS | G | 8.5609E-11 | 1,0973E+06 | 0.0 |

| | | | | | | | | | |
|---|---|---|---|---|---|---|---|---|---|
| rs753724 | 10 | 96051417 | GTEx/v8 | Cells EBV-transformed lymphocytes | PLCE1 | T | 9.1525E-11 | 0.86599 | 1.20882E-07 |
| rs11187842 | 10 | 96052511 | GTEx/v8 | Cells EBV-transformed lymphocytes | PLCE1 | T | 9.1525E-11 | 0.86599 | 1.20882E-07 |
| rs3781266 | 10 | 96052747 | GTEx/v8 | Cells EBV-transformed lymphocytes | PLCE1 | G | 9.1525E-11 | 0.86599 | 1.20882E-07 |
| rs3740365 | 10 | 96053239 | GTEx/v8 | Cells EBV-transformed lymphocytes | PLCE1 | A | 9.1525E-11 | 0.86599 | 1.20882E-07 |
| rs12220091 | 10 | 96053689 | GTEx/v8 | Cells EBV-transformed lymphocytes | PLCE1 | T | 9.1525E-11 | 0.86599 | 1.20882E-07 |
| rs11187890 | 10 | 96102758 | eQTLGen | eQTLGen cis eQTLs | HELLS | C | 9.4938E-11 | 1,0407E+06 | 0.0 |
| rs3740359 | 10 | 96100119 | eQTLcatalogue | TwinsUK ge LCL | PLCE1 | C | 1.0089E-10 | 0.121466 | 5.0445E-07 |
| rs11187907 | 10 | 96125361 | eQTLcatalogue | GEUVADIS ge LCL | PLCE1 | C | 1.20612E-10 | 0.293466 | 6.0306E-07 |
| rs11187907 | 10 | 96125361 | GTEx/v8 | Cells EBV-transformed lymphocytes | PLCE1 | T | 1.25898E-10 | 0.900331 | 1.20882E-07 |
| rs3740359 | 10 | 96100119 | eQTLGen | eQTLGen cis eQTLs | HELLS | T | 1.3681E-10 | 8,3839E+05 | 0.0 |
| rs79746649 | 10 | 96120218 | eQTLGen | eQTLGen cis eQTLs | NOC3L | T | 1.425E-10 | 8,1611E+05 | 0.0 |
| rs12220091 | 10 | 96053689 | eQTLcatalogue | BLUEPRINT ge monocyte | NOC3L | C | 1.65134E-10 | 0.240279 | 8.2567E-07 |
| rs769015184 | 10 | 96049708 | eQTLcatalogue | BLUEPRINT ge monocyte | NOC3L | AC | 1.65229E-10 | 0.240282 | 8.26145E-07 |
| rs753724 | 10 | 96051417 | eQTLcatalogue | BLUEPRINT ge monocyte | NOC3L | G | 1.65229E-10 | 0.240282 | 8.26145E-07 |
| rs11187842 | 10 | 96052511 | eQTLcatalogue | BLUEPRINT ge monocyte | NOC3L | C | 1.65229E-10 | 0.240282 | 8.26145E-07 |
| rs3781266 | 10 | 96052747 | eQTLcatalogue | BLUEPRINT ge monocyte | NOC3L | A | 1.65229E-10 | 0.240282 | 8.26145E-07 |
| rs3740365 | 10 | 96053239 | eQTLcatalogue | BLUEPRINT ge monocyte | NOC3L | T | 1.65229E-10 | 0.240282 | 8.26145E-07 |
| rs75017201 | 10 | 96055152 | eQTLcatalogue | BLUEPRINT ge monocyte | NOC3L | C | 1.65229E-10 | 0.240282 | 8.26145E-07 |
| rs11187845 | 10 | 96060198 | GTEx/v8 | Cells EBV-transformed lymphocytes | PLCE1 | A | 1.82938E-10 | 0.86298 | 1.20882E-07 |
| rs11187847 | 10 | 96063440 | GTEx/v8 | Cells EBV-transformed lymphocytes | PLCE1 | G | 1.82938E-10 | 0.86298 | 1.20882E-07 |
| rs74825688 | 10 | 96119687 | eQTLGen | eQTLGen cis eQTLs | NOC3L | A | 1.8829E-10 | 6,5978E+05 | 0.0 |
| rs7080472 | 10 | 96012950 | GTEx/v8 | Skin Sun Exposed Lower leg | NOC3L | T | 2.07651E-10 | 0.149208 | 4.18218E-20 |
| rs113544984 | 10 | 96127448 | eQTLGen | eQTLGen cis eQTLs | HELLS | A | 2.0897E-10 | 6,0207E+05 | 0.0 |
| rs10509672 | 10 | 96115957 | eQTLGen | eQTLGen cis eQTLs | NOC3L | C | 2.1531E-10 | 5,8491E+05 | 0.0 |
| rs11187864 | 10 | 96082506 | eQTLGen | eQTLGen cis eQTLs | HELLS | T | 2.8671E-10 | 6.306 | 0.0 |
| rs79193572 | 10 | 96119713 | eQTLGen | eQTLGen cis eQTLs | NOC3L | G | 2.8782E-10 | 4,2128E+05 | 0.0 |
| rs3740360 | 10 | 96025491 | BIOSQTL | BIOS eQTL geneLevel | TBC1D12 | C | 3.29122E-10 | 6.2844446 | 0.0 |
| rs752140 | 10 | 96071396 | GTEx/v8 | Cells Cultured fibroblasts | PLCE1-AS1 | T | 3.39376E-10 | -0.338477 | 3.86001E-30 |
| rs11187837 | 10 | 96035980 | BIOSQTL | BIOS eQTL geneLevel | TBC1D12 | C | 4.05749E-10 | 6.2518755 | 0.0 |
| rs11187895 | 10 | 96106240 | eQTLGen | eQTLGen cis eQTLs | NOC3L | C | 4.1574E-10 | 2,1236E+05 | 0.0 |
| rs2274224 | 10 | 96039597 | eQTLcatalogue | BLUEPRINT ge monocyte | NOC3L | G | 4.18698E-10 | 0.135701 | 2.09349E-06 |
| rs3891783 | 10 | 96015793 | eQTLcatalogue | BLUEPRINT ge monocyte | NOC3L | C | 4.20614E-10 | 0.135685 | 2.10307E-06 |
| rs10786156 | 10 | 96014622 | eQTLcatalogue | BLUEPRINT ge monocyte | NOC3L | C | 4.21663E-10 | 0.135662 | 2.10832E-06 |

| | | | | | | | | | |
|---|---|---|---|---|---|---|---|---|---|
| rs11187838 | 10 | 96038686 | eQTLcatalogue | BLUEPRINT ge monocyte | NOC3L | G | 4.23274E-10 | 0.135644 | 2.11637E-06 |
| rs17109875 | 10 | 96026575 | BIOSQTL | BIOS eQTL geneLevel | TBC1D12 | C | 5.44194E-10 | 6.2059539 | 0.0 |
| rs11187863 | 10 | 96081457 | GTEx/v8 | Cells Cultured fibroblasts | PLCE1-AS1 | G | 5.69704E-10 | -0.333231 | 3.86001E-30 |
| rs11187866 | 10 | 96085991 | GTEx/v8 | Cells Cultured fibroblasts | PLCE1-AS1 | G | 5.69704E-10 | -0.333231 | 3.86001E-30 |
| rs11187883 | 10 | 96096866 | GTEx/v8 | Cells Cultured fibroblasts | PLCE1-AS1 | C | 5.69704E-10 | -0.333231 | 3.86001E-30 |
| rs34309017 | 10 | 96061619 | GTEx/v8 | Cells Cultured fibroblasts | PLCE1-AS1 | A | 6.42319E-10 | -0.337731 | 3.86001E-30 |
| rs12217792 | 10 | 96062386 | GTEx/v8 | Cells Cultured fibroblasts | PLCE1-AS1 | C | 6.42319E-10 | -0.337731 | 3.86001E-30 |
| rs11187890 | 10 | 96102758 | eQTLGen | eQTLGen cis eQTLs | NOC3L | C | 7.6705E-10 | 6.1517 | 1.316E-05 |
| rs75409190 | 10 | 96072425 | GTEx/v8 | Cells Cultured fibroblasts | PLCE1-AS1 | T | 8.07901E-10 | -0.344158 | 3.86001E-30 |
| rs771106747 | 10 | 96072873 | GTEx/v8 | Cells Cultured fibroblasts | PLCE1-AS1 | C | 8.07901E-10 | -0.344158 | 3.86001E-30 |
| rs749295068 | 10 | 96073508 | GTEx/v8 | Cells Cultured fibroblasts | PLCE1-AS1 | A | 8.07901E-10 | -0.344158 | 3.86001E-30 |
| rs11187856 | 10 | 96076869 | GTEx/v8 | Cells Cultured fibroblasts | PLCE1-AS1 | A | 8.07901E-10 | -0.344158 | 3.86001E-30 |
| rs11187863 | 10 | 96081457 | GTEx/v8 | Cells EBV-transformed lymphocytes | PLCE1 | G | 1.09697E-09 | 0.828697 | 1.20882E-07 |
| rs11187866 | 10 | 96085991 | GTEx/v8 | Cells EBV-transformed lymphocytes | PLCE1 | G | 1.09697E-09 | 0.828697 | 1.20882E-07 |
| rs11187883 | 10 | 96096866 | GTEx/v8 | Cells EBV-transformed lymphocytes | PLCE1 | C | 1.09697E-09 | 0.828697 | 1.20882E-07 |
| rs7096883 | 10 | 96069149 | GTEx/v8 | Cells Cultured fibroblasts | PLCE1-AS1 | A | 1.15214E-09 | -0.331235 | 3.86001E-30 |
| rs7096883 | 10 | 96069149 | eQTLcatalogue | TwinsUK ge LCL | PLCE1 | G | 1.16776E-09 | 0.112521 | 5.8388E-06 |
| rs7080472 | 10 | 96012950 | eQTLcatalogue | BLUEPRINT ge monocyte | NOC3L | G | 1.24509E-09 | 0.131762 | 6.22545E-06 |
| rs11187870 | 10 | 96087866 | GTEx/v8 | Cells Cultured fibroblasts | PLCE1-AS1 | C | 1.34499E-09 | -0.342164 | 3.86001E-30 |
| rs11187897 | 10 | 96106603 | eQTLGen | eQTLGen cis eQTLs | NOC3L | G | 1.3646E-09 | 6.0596 | 1.3094E-05 |
| rs3891783 | 10 | 96015793 | GTEx/v8 | Skin Sun Exposed Lower leg | NOC3L | G | 1.41912E-09 | 0.141003 | 4.18218E-20 |
| rs57866767 | 10 | 96023077 | GTEx/v8 | Skin Sun Exposed Lower leg | NOC3L | C | 1.41912E-09 | 0.141003 | 4.18218E-20 |
| rs7100626 | 10 | 96069674 | GTEx/v8 | Cells Cultured fibroblasts | PLCE1-AS1 | A | 1.47359E-09 | -0.337229 | 3.86001E-30 |
| rs11187852 | 10 | 96070132 | GTEx/v8 | Cells Cultured fibroblasts | PLCE1-AS1 | A | 1.47359E-09 | -0.337229 | 3.86001E-30 |
| rs11187882 | 10 | 96095861 | GTEx/v8 | Cells Cultured fibroblasts | PLCE1-AS1 | C | 1.4751E-09 | -0.337177 | 3.86001E-30 |
| rs74825688 | 10 | 96119687 | GTEx/v8 | Cells Cultured fibroblasts | PLCE1-AS1 | A | 1.4816E-09 | -0.33994 | 3.86001E-30 |
| rs10786156 | 10 | 96014622 | GTEx/v8 | Nerve Tibial | NOC3L | G | 1.48976E-09 | 0.168244 | 1.32457E-19 |
| rs752140 | 10 | 96071396 | eQTLGen | eQTLGen cis eQTLs | NOC3L | T | 1.5184E-09 | 6.0425 | 1.3080E-05 |
| rs79193572 | 10 | 96119713 | GTEx/v8 | Cells Cultured fibroblasts | PLCE1-AS1 | G | 1.55958E-09 | -0.341493 | 3.86001E-30 |
| rs10786156 | 10 | 96014622 | GTEx/v8 | Skin Sun Exposed Lower leg | NOC3L | G | 1.65539E-09 | 0.140345 | 4.18218E-20 |
| rs2274224 | 10 | 96039597 | GTEx/v8 | Nerve Tibial | NOC3L | C | 1.95095E-09 | 0.166488 | 1.32457E-19 |
| rs12217792 | 10 | 96062386 | eQTLcatalogue | TwinsUK ge LCL | PLCE1 | T | 2.09527E-09 | 0.108906 | 1.04764E-05 |

| | | | | | | | | | |
|---|---|---|---|---|---|---|---|---|---|
| rs11187847 | 10 | 96063440 | eQTLcatalogue | TwinsUK ge LCL | PLCE1 | C | 2.09527E-09 | 0.108906 | 1.04764E-05 |
| rs3891783 | 10 | 96015793 | GTEx/v8 | Nerve Tibial | NOC3L | G | 2.42323E-09 | 0.166243 | 1.32457E-19 |
| rs57866767 | 10 | 96023077 | GTEx/v8 | Nerve Tibial | NOC3L | C | 2.42323E-09 | 0.166243 | 1.32457E-19 |
| rs7100626 | 10 | 96069674 | eQTLGen | eQTLGen cis eQTLs | NOC3L | A | 2.5503E-09 | 2,8063E+06 | 2.603E-05 |
| rs11187852 | 10 | 96070132 | eQTLGen | eQTLGen cis eQTLs | NOC3L | A | 2.6071E-09 | 2,7931E+06 | 2.602E-05 |
| rs11187864 | 10 | 96082506 | GTEx/v8 | Cells Cultured fibroblasts | PLCE1-AS1 | T | 2.65113E-09 | -0.330345 | 3.86001E-30 |
| rs11187877 | 10 | 96092121 | GTEx/v8 | Cells Cultured fibroblasts | PLCE1-AS1 | A | 2.65113E-09 | -0.330345 | 3.86001E-30 |
| rs145707916 | 10 | 96092992 | GTEx/v8 | Cells Cultured fibroblasts | PLCE1-AS1 | GA | 2.65113E-09 | -0.330345 | 3.86001E-30 |
| rs750148755 | 10 | 96095271 | GTEx/v8 | Cells Cultured fibroblasts | PLCE1-AS1 | C | 2.65113E-09 | -0.330345 | 3.86001E-30 |
| rs778275808 | 10 | 96114212 | GTEx/v8 | Cells Cultured fibroblasts | PLCE1-AS1 | T | 2.75585E-09 | -0.334894 | 3.86001E-30 |
| rs10509672 | 10 | 96115957 | GTEx/v8 | Cells Cultured fibroblasts | PLCE1-AS1 | C | 2.75585E-09 | -0.334894 | 3.86001E-30 |
| rs79746649 | 10 | 96120218 | GTEx/v8 | Cells Cultured fibroblasts | PLCE1-AS1 | T | 2.75585E-09 | -0.334894 | 3.86001E-30 |
| rs2274224 | 10 | 96039597 | GTEx/v8 | Skin Sun Exposed Lower leg | NOC3L | C | 2.84729E-09 | 0.13795 | 4.18218E-20 |
| rs11187890 | 10 | 96102758 | GTEx/v8 | Cells Cultured fibroblasts | PLCE1-AS1 | C | 2.86262E-09 | -0.331106 | 3.86001E-30 |
| rs11187895 | 10 | 96106240 | GTEx/v8 | Cells Cultured fibroblasts | PLCE1-AS1 | C | 2.86262E-09 | -0.331106 | 3.86001E-30 |
| rs11187897 | 10 | 96106603 | GTEx/v8 | Cells Cultured fibroblasts | PLCE1-AS1 | G | 2.86262E-09 | -0.331106 | 3.86001E-30 |
| rs11187863 | 10 | 96081457 | eQTLGen | eQTLGen cis eQTLs | NOC3L | G | 2.8642E-09 | 2,7373E+06 | 2.6E-05 |
| rs11187866 | 10 | 96085991 | eQTLGen | eQTLGen cis eQTLs | NOC3L | G | 2.8993E-09 | 2,7292E+06 | 2.6E-05 |
| rs7101207 | 10 | 96122857 | GTEx/v8 | Cells Cultured fibroblasts | PLCE1-AS1 | A | 3.28002E-09 | -0.335045 | 3.86001E-30 |
| rs112252876 | 10 | 96123355 | GTEx/v8 | Cells Cultured fibroblasts | PLCE1-AS1 | T | 3.28002E-09 | -0.335045 | 3.86001E-30 |
| rs113544984 | 10 | 96127448 | eQTLcatalogue | GENCORDge LCL | PLCE1 | C | 3.28253E-09 | 0.293762 | 1.64127E-05 |
| rs11187826 | 10 | 95988042 | BIOSQTL | BIOS eQTL geneLevel | TBC1D12 | G | 3.39212E-09 | 5.9115847 | 6.013E-06 |
| rs752140 | 10 | 96071396 | GTEx/v8 | Cells EBV-transformed lymphocytes | PLCE1 | T | 3.75953E-09 | 0.803891 | 1.20882E-07 |
| rs112252876 | 10 | 96123355 | eQTLcatalogue | GENCORDge LCL | PLCE1 | C | 3.84298E-09 | 0.287768 | 1.92149E-05 |
| rs7101207 | 10 | 96122857 | eQTLcatalogue | GENCORDge LCL | PLCE1 | G | 3.89754E-09 | 0.28761 | 1.94877E-05 |
| rs11187907 | 10 | 96125361 | eQTLcatalogue | GENCORDge LCL | PLCE1 | C | 4.00496E-09 | 0.287654 | 2.00248E-05 |
| rs11187906 | 10 | 96124838 | eQTLcatalogue | GENCORDge LCL | PLCE1 | T | 4.02821E-09 | 0.287628 | 2.0141E-05 |
| rs11187883 | 10 | 96096866 | eQTLGen | eQTLGen cis eQTLs | NOC3L | C | 4.379E-09 | 2,4812E+06 | 3.237E-05 |
| rs11187840 | 10 | 96050351 | GTEx/v8 | Cells Cultured fibroblasts | PLCE1-AS1 | G | 4.40204E-09 | -0.315138 | 3.86001E-30 |
| rs75017201 | 10 | 96055152 | GTEx/v8 | Cells Cultured fibroblasts | PLCE1-AS1 | T | 4.40204E-09 | -0.315138 | 3.86001E-30 |
| rs113544984 | 10 | 96127448 | GTEx/v8 | Cells Cultured fibroblasts | PLCE1-AS1 | A | 4.55401E-09 | -0.335199 | 3.86001E-30 |
| rs11187838 | 10 | 96038686 | GTEx/v8 | Skin Sun Exposed Lower leg | NOC3L | A | 4.67744E-09 | 0.136601 | 4.18218E-20 |
| rs75409190 | 10 | 96072425 | eQTLGen | eQTLGen cis eQTLs | NOC3L | T | 5.0293E-09 | 2,3976E+06 | 3.233E-05 |
| rs11187844 | 10 | 96056629 | eQTLGen | eQTLGen cis eQTLs | TBC1D12 | A | 5.8637E-09 | 2,3041E+06 | 4.520E-05 |
| rs11187856 | 10 | 96076869 | eQTLGen | eQTLGen cis eQTLs | NOC3L | A | 6.4366E-09 | 2,2467E+06 | 4.517E-05 |

| | | | | | | | | | |
|---|---|---|---|---|---|---|---|---|---|
| rs11187877 | 10 | 96092121 | eQTLGen | eQTLGen cis eQTLs | NOC3L | A | 7.0975E-09 | 2,1872E+06 | 4.514E-05 |
| rs7096883 | 10 | 96069149 | eQTLGen | eQTLGen cis eQTLs | NOC3L | A | 8.6744E-09 | 2,0634E+06 | 5.795E-05 |
| rs3740360 | 10 | 96025491 | GTEx/v8 | Cells EBV-transformed lymphocytes | PLCE1 | C | 9.05253E-09 | 0.798488 | 1.20882E-07 |
| rs17109875 | 10 | 96026575 | GTEx/v8 | Cells EBV-transformed lymphocytes | PLCE1 | C | 9.05253E-09 | 0.798488 | 1.20882E-07 |
| rs11187837 | 10 | 96035980 | GTEx/v8 | Cells EBV-transformed lymphocytes | PLCE1 | C | 9.05253E-09 | 0.798488 | 1.20882E-07 |
| rs11187847 | 10 | 96063440 | eQTLGen | eQTLGen cis eQTLs | NOC3L | G | 9.8903E-09 | 1,9823E+06 | 5.787E-05 |
| rs12217792 | 10 | 96062386 | eQTLGen | eQTLGen cis eQTLs | NOC3L | C | 1.0186E-08 | 1,9644E+06 | 5.786E-05 |
| rs11187844 | 10 | 96056629 | eQTLcatalogue | GENCORDge LCL | PLCE1 | C | 1.04815E-08 | 0.235889 | 5.24075E-05 |
| rs11187882 | 10 | 96095861 | eQTLGen | eQTLGen cis eQTLs | NOC3L | C | 1.0753E-08 | 1,9308E+06 | 5.782E-05 |
| rs11187870 | 10 | 96087866 | eQTLGen | eQTLGen cis eQTLs | NOC3L | C | 1.1675E-08 | 1,8789E+06 | 6.421E-05 |
| rs12217792 | 10 | 96062386 | eQTLGen | eQTLGen cis eQTLs | HELLS | C | 1.1758E-08 | 1,8749E+06 | 6.420E-05 |
| rs7096883 | 10 | 96069149 | eQTLGen | eQTLGen cis eQTLs | HELLS | A | 1.1953E-08 | 1,8651E+06 | 6.418E-05 |
| rs3740359 | 10 | 96100119 | eQTLcatalogue | GENCORDge LCL | PLCE1 | C | 1.21116E-08 | 0.272238 | 6.0558E-05 |
| rs7080472 | 10 | 96012950 | GTEx/v8 | Nerve Tibial | NOC3L | T | 1.27639E-08 | 0.159332 | 1.32457E-19 |
| rs11187842 | 10 | 96052511 | eQTLcatalogue | GENCORDge LCL | PLCE1 | C | 1.39637E-08 | 0.264935 | 6.98185E-05 |
| rs3781266 | 10 | 96052747 | eQTLcatalogue | GENCORDge LCL | PLCE1 | A | 1.40013E-08 | 0.264926 | 7.00065E-05 |
| rs3740365 | 10 | 96053239 | eQTLcatalogue | GENCORDge LCL | PLCE1 | T | 1.41153E-08 | 0.2649 | 7.05765E-05 |
| rs12220091 | 10 | 96053689 | eQTLcatalogue | GENCORDge LCL | PLCE1 | C | 1.41728E-08 | 0.264887 | 7.0864E-05 |
| rs75017201 | 10 | 96055152 | eQTLcatalogue | GENCORDge LCL | PLCE1 | C | 1.45863E-08 | 0.264791 | 7.29315E-05 |
| rs11187838 | 10 | 96038686 | GTEx/v8 | Nerve Tibial | NOC3L | A | 1.48555E-08 | 0.158012 | 1.32457E-19 |
| rs11187845 | 10 | 96060198 | eQTLGen | eQTLGen cis eQTLs | NOC3L | A | 1.512E-08 | 1,7182E+06 | 7.684E-05 |
| rs3740365 | 10 | 96053239 | eQTLGen | eQTLGen cis eQTLs | HELLS | A | 1.6441E-08 | 1,6653E+06 | 8.958E-05 |
| rs753724 | 10 | 96051417 | eQTLcatalogue | GENCORDge LCL | PLCE1 | G | 1.69177E-08 | 0.264207 | 8.45885E-05 |
| rs11187840 | 10 | 96050351 | eQTLGen | eQTLGen cis eQTLs | HELLS | G | 1.7486E-08 | 1,6269E+06 | 9.593E-05 |
| rs753724 | 10 | 96051417 | eQTLGen | eQTLGen cis eQTLs | HELLS | T | 1.7567E-08 | 1,6240E+06 | 9.591E-05 |
| rs75017201 | 10 | 96055152 | eQTLGen | eQTLGen cis eQTLs | HELLS | T | 1.977E-08 | 1,5491E+06 | 1.085E-4 |
| rs752140 | 10 | 96071396 | PsychENCODE | PsychENCODE eQTLs | NOC3L | T | 1.9801E-08 | 0.0969569 | 3.523E-06 |
| rs752140 | 10 | 96071396 | eQTLcatalogue | GENCORDge LCL | PLCE1 | C | 1.98151E-08 | 0.269571 | 9.90755E-05 |
| rs11187847 | 10 | 96063440 | eQTLGen | eQTLGen cis eQTLs | HELLS | G | 2.0093E-08 | 1,5389E+06 | 1.085E-4 |
| rs11187856 | 10 | 96076869 | eQTLcatalogue | GENCORDge LCL | PLCE1 | G | 2.01876E-08 | 0.269417 | 0.000100938 |
| rs11187845 | 10 | 96060198 | eQTLcatalogue | TwinsUK ge LCL | PLCE1 | C | 2.02127E-08 | 0.0997346 | 0.000101063 |
| rs749295068 | 10 | 96073508 | eQTLcatalogue | GENCORDge LCL | PLCE1 | AGAT | 2.02287E-08 | 0.269401 | 0.000101144 |
| rs11187863 | 10 | 96081457 | eQTLcatalogue | GENCORDge LCL | PLCE1 | T | 2.02492E-08 | 0.269347 | 0.000101246 |
| rs11187864 | 10 | 96082506 | eQTLcatalogue | GENCORDge LCL | PLCE1 | C | 2.02492E-08 | 0.269347 | 0.000101246 |
| rs11187866 | 10 | 96085991 | eQTLcatalogue | GENCORDge LCL | PLCE1 | C | 2.02492E-08 | 0.269347 | 0.000101246 |
| rs11187870 | 10 | 96087866 | eQTLcatalogue | GENCORDge LCL | PLCE1 | G | 2.02492E-08 | 0.269347 | 0.000101246 |
| rs11187877 | 10 | 96092121 | eQTLcatalogue | GENCORDge LCL | PLCE1 | G | 2.02492E-08 | 0.269347 | 0.000101246 |
| rs145707916 | 10 | 96092992 | eQTLcatalogue | GENCORDge LCL | PLCE1 | G | 2.02492E-08 | 0.269347 | 0.000101246 |
| rs12217792 | 10 | 96062386 | eQTLGen | eQTLGen cis eQTLs | PDLIM1 | C | 2.0256E-08 | -5.6096 | 0.0001085152559683 |
| rs75409190 | 10 | 96072425 | eQTLcatalogue | GENCORDge LCL | PLCE1 | C | 2.02982E-08 | 0.269356 | 0.000101491 |
| rs7100626 | 10 | 96069674 | eQTLcatalogue | GENCORDge LCL | PLCE1 | C | 2.03023E-08 | 0.269338 | 0.000101511 |
| rs11187852 | 10 | 96070132 | eQTLcatalogue | GENCORDge LCL | PLCE1 | G | 2.03023E-08 | 0.269338 | 0.000101511 |

| | | | | | | | | | |
|---|---|---|---|---|---|---|---|---|---|
| rs771106747 | 10 | 96072873 | eQTLcatalogue | GENCORDge LCL | PLCE1 | CTA | 2.03044E-08 | 0.269369 | 0.000101522 |
| rs750148755 | 10 | 96095271 | eQTLcatalogue | GENCORDge LCL | PLCE1 | CATTTT | 2.04426E-08 | 0.269275 | 0.000102213 |
| rs79746649 | 10 | 96120218 | eQTLcatalogue | GENCORDge LCL | PLCE1 | C | 2.04781E-08 | 0.268979 | 0.00010239 |
| rs11187842 | 10 | 96052511 | eQTLGen | eQTLGen cis eQTLs | HELLS | T | 2.0539E-08 | 1,5246E+06 | 0.0001085014041358 |
| rs11187882 | 10 | 96095861 | eQTLcatalogue | GENCORDge LCL | PLCE1 | A | 2.05663E-08 | 0.269242 | 0.000102831 |
| rs11187890 | 10 | 96102758 | eQTLcatalogue | GENCORDge LCL | PLCE1 | T | 2.06772E-08 | 0.269218 | 0.000103386 |
| rs11187895 | 10 | 96106240 | eQTLcatalogue | GENCORDge LCL | PLCE1 | G | 2.12925E-08 | 0.268952 | 0.000106463 |
| rs11187883 | 10 | 96096866 | eQTLcatalogue | GENCORDge LCL | PLCE1 | T | 2.12993E-08 | 0.268805 | 0.000106496 |
| rs11187897 | 10 | 96106603 | eQTLcatalogue | GENCORDge LCL | PLCE1 | A | 2.13815E-08 | 0.268908 | 0.000106907 |
| rs10509672 | 10 | 96115957 | eQTLcatalogue | GENCORDge LCL | PLCE1 | G | 2.16348E-08 | 0.268776 | 0.000108174 |
| rs778275808 | 10 | 96114212 | eQTLcatalogue | GENCORDge LCL | PLCE1 | TCTC | 2.17765E-08 | 0.268773 | 0.000108882 |
| rs11187840 | 10 | 96050351 | eQTLcatalogue | GENCORDge LCL | PLCE1 | A | 2.20621E-08 | 0.262863 | 0.000110311 |
| rs79193572 | 10 | 96119713 | eQTLcatalogue | GENCORDge LCL | PLCE1 | A | 2.21705E-08 | 0.268385 | 0.000110852 |
| rs7096883 | 10 | 96069149 | eQTLGen | eQTLGen cis eQTLs | PDLIM1 | A | 2.3343E-08 | -5.5851 | 0.0001147666411629 |
| rs769015184 | 10 | 96049708 | eQTLcatalogue | GENCORDge LCL | PLCE1 | AC | 2.47513E-08 | 0.262186 | 0.000123756 |
| rs74825688 | 10 | 96119687 | eQTLcatalogue | GENCORDge LCL | PLCE1 | G | 2.62812E-08 | 0.26629 | 0.000131406 |
| rs3781266 | 10 | 96052747 | eQTLGen | eQTLGen cis eQTLs | HELLS | G | 2.6457E-08 | 1,3643E+06 | 0.0001146204788588 |
| rs17109875 | 10 | 96026575 | eQTLcatalogue | BLUEPRINTge monocyte | NOC3L | T | 2.74731E-08 | 0.184435 | 0.000137365 |
| rs11187837 | 10 | 96035980 | eQTLcatalogue | BLUEPRINTge monocyte | NOC3L | T | 2.74823E-08 | 0.184411 | 0.000137411 |
| rs3740360 | 10 | 96025491 | eQTLcatalogue | BLUEPRINTge monocyte | NOC3L | A | 2.76868E-08 | 0.184361 | 0.000138434 |
| rs12220091 | 10 | 96053689 | eQTLGen | eQTLGen cis eQTLs | HELLS | T | 2.8731E-08 | 1,3121E+06 | 0.0001272669424117 |
| rs11187907 | 10 | 96125361 | GTEx/v8 | Cells Cultured fibroblasts | PLCE1-AS1 | T | 2.91468E-08 | -0.309692 | 3.86001E-30 |
| rs10786156 | 10 | 96014622 | GTEx/v8 | Skin Not Sun Exposed Suprapubic | NOC3L | G | 3.68785E-08 | 0.151155 | 2.01564E-23 |
| rs3891783 | 10 | 96015793 | GTEx/v8 | Skin Not Sun Exposed Suprapubic | NOC3L | G | 3.68785E-08 | 0.151155 | 2.01564E-23 |
| rs57866767 | 10 | 96023077 | GTEx/v8 | Skin Not Sun Exposed Suprapubic | NOC3L | C | 3.68785E-08 | 0.151155 | 2.01564E-23 |
| rs7096883 | 10 | 96069149 | eQTLcatalogue | GENCORDge LCL | PLCE1 | G | 3.69827E-08 | 0.262001 | 0.000184913 |
| rs11187840 | 10 | 96050351 | eQTLcatalogue | TwinsUK ge LCL | PLCE1 | A | 3.7527E-08 | 0.0963516 | 0.000187635 |
| rs753724 | 10 | 96051417 | eQTLcatalogue | TwinsUK ge LCL | PLCE1 | G | 3.7527E-08 | 0.0963516 | 0.000187635 |
| rs11187842 | 10 | 96052511 | eQTLcatalogue | TwinsUK ge LCL | PLCE1 | C | 3.7527E-08 | 0.0963516 | 0.000187635 |
| rs3781266 | 10 | 96052747 | eQTLcatalogue | TwinsUK ge LCL | PLCE1 | A | 3.7527E-08 | 0.0963516 | 0.000187635 |
| rs3740365 | 10 | 96053239 | eQTLcatalogue | TwinsUK ge LCL | PLCE1 | T | 3.7527E-08 | 0.0963516 | 0.000187635 |
| rs12220091 | 10 | 96053689 | eQTLcatalogue | TwinsUK ge LCL | PLCE1 | C | 3.7527E-08 | 0.0963516 | 0.000187635 |
| rs75017201 | 10 | 96055152 | eQTLcatalogue | TwinsUK ge LCL | PLCE1 | C | 3.7527E-08 | 0.0963516 | 0.000187635 |
| rs200197176 | 10 | 96055963 | eQTLcatalogue | GENCORDge LCL | PLCE1 | C | 3.77732E-08 | 0.25545 | 0.000188866 |
| rs11187845 | 10 | 96060198 | PsychENCODE | PsychENCODE eQTLs | NOC3L | A | 3.87215E-08 | 0.097552 | 6.591E-06 |
| 10:96112261_AAAAAAAC_A | 10 | 96112261 | eQTLcatalogue | GENCORDge LCL | PLCE1 | AAAAAAAC | 3.88536E-08 | 0.25946 | 0.000194268 |
| rs75409190 | 10 | 96072425 | PsychENCODE | PsychENCODE eQTLs | NOC3L | T | 3.92785E-08 | 0.102328 | 6.68E-06 |
| rs12217792 | 10 | 96062386 | eQTLcatalogue | GENCORDge LCL | PLCE1 | T | 4.12977E-08 | 0.261741 | 0.000206488 |

| | | | | | | | | | |
|---|---|---|---|---|---|---|---|---|---|
| rs752140 | 10 | 96071396 | GTEx/v8 | Muscle Skeletal | NOC3L | T | 4.2379E-08 | 0.260214 | 7.35621E-14 |
| rs11187863 | 10 | 96081457 | GTEx/v8 | Muscle Skeletal | NOC3L | G | 4.2379E-08 | 0.260214 | 7.35621E-14 |
| rs11187866 | 10 | 96085991 | GTEx/v8 | Muscle Skeletal | NOC3L | G | 4.2379E-08 | 0.260214 | 7.35621E-14 |
| rs11187883 | 10 | 96096866 | GTEx/v8 | Muscle Skeletal | NOC3L | C | 4.2379E-08 | 0.260214 | 7.35621E-14 |
| rs34309017 | 10 | 96061619 | eQTLcatalogue | GENCORDge LCL | PLCE1 | ACTT | 4.27059E-08 | 0.261633 | 0.00021353 |
| rs7080472 | 10 | 96012950 | GTEx/v8 | Skin Not Sun Exposed Suprapubic | NOC3L | T | 4.38581E-08 | 0.151839 | 2.01564E-23 |
| rs11187845 | 10 | 96060198 | eQTLGen | eQTLGen cis eQTLs | HELLS | A | 4.4649E-08 | 1,0279E+06 | 0.0001774510425248 |
| rs11187845 | 10 | 96060198 | eQTLcatalogue | GENCORDge LCL | PLCE1 | C | 4.51948E-08 | 0.261443 | 0.000225974 |
| rs11187882 | 10 | 96095861 | PsychENCODE | PsychENCODE eQTLs | NOC3L | C | 4.61283E-08 | 0.101642 | 7.755E-06 |
| rs11187856 | 10 | 96076869 | PsychENCODE | PsychENCODE eQTLs | NOC3L | A | 4.68244E-08 | 0.101697 | 7.864E-06 |
| rs11187847 | 10 | 96063440 | PsychENCODE | PsychENCODE eQTLs | NOC3L | G | 4.94659E-08 | 0.0960576 | 8.276E-06 |
| rs11187847 | 10 | 96063440 | eQTLGen | eQTLGen cis eQTLs | PDLIM1 | G | 5.6354E-08 | -5.43 | 0.0002400657021921 |
| rs11187864 | 10 | 96082506 | PsychENCODE | PsychENCODE eQTLs | NOC3L | T | 6.74076E-08 | 0.100225 | 1.104E-05 |
| rs113544984 | 10 | 96127448 | PsychENCODE | PsychENCODE eQTLs | NOC3L | A | 6.75766E-08 | 0.102717 | 1.106E-05 |
| rs2274224 | 10 | 96039597 | GTEx/v8 | Heart Left Ventricle | NOC3L | C | 7.19999E-08 | 0.19175 | 3.09268E-10 |
| rs10786156 | 10 | 96014622 | GTEx/v8 | Heart Left Ventricle | NOC3L | G | 7.68073E-08 | 0.190819 | 3.09268E-10 |
| rs2274224 | 10 | 96039597 | GTEx/v8 | Skin Not Sun Exposed Suprapubic | NOC3L | C | 7.79403E-08 | 0.147484 | 2.01564E-23 |
| rs11187906 | 10 | 96124838 | PsychENCODE | PsychENCODE eQTLs | NOC3L | A | 7.89484E-08 | 0.102219 | 1.279E-05 |
| rs7080472 | 10 | 96012950 | GTEx/v8 | Esophagus Muscularis | NOC3L | T | 8.19561E-08 | 0.153403 | 2.723E-20 |
| rs11187864 | 10 | 96082506 | eQTLGen | eQTLGen cis eQTLs | NOC3L | T | 8.2214E-08 | 6,2944E+05 | 0.0003272704386682 |
| rs7078844 | 10 | 96081140 | eQTLGen | eQTLGen cis eQTLs | NOC3L | T | 9.9519E-08 | -5.3274 | 0.0003455425017277 |
| rs12220091 | 10 | 96053689 | PsychENCODE | PsychENCODE eQTLs | NOC3L | T | 1.00415E-07 | 0.093271 | 1.6E-05 |
| rs3781266 | 10 | 96052747 | PsychENCODE | PsychENCODE eQTLs | NOC3L | G | 1.0346E-07 | 0.0931764 | 1.645E-05 |
| rs7100626 | 10 | 96069674 | PsychENCODE | PsychENCODE eQTLs | NOC3L | A | 1.13755E-07 | 0.097166 | 1.796E-05 |
| rs75017201 | 10 | 96055152 | PsychENCODE | PsychENCODE eQTLs | NOC3L | T | 1.16476E-07 | 0.0941142 | 1.836E-05 |
| rs7096883 | 10 | 96069149 | PsychENCODE | PsychENCODE eQTLs | NOC3L | A | 1.25814E-07 | 0.0938882 | 1.972E-05 |
| rs79193572 | 10 | 96119713 | PsychENCODE | PsychENCODE eQTLs | NOC3L | G | 1.30794E-07 | 0.0996592 | 2.045E-05 |
| rs3891783 | 10 | 96015793 | GTEx/v8 | Heart Left Ventricle | NOC3L | G | 1.31784E-07 | 0.188086 | 3.09268E-10 |
| rs57866767 | 10 | 96023077 | GTEx/v8 | Heart Left Ventricle | NOC3L | C | 1.31784E-07 | 0.188086 | 3.09268E-10 |
| rs10509672 | 10 | 96115957 | PsychENCODE | PsychENCODE eQTLs | NOC3L | C | 1.32438E-07 | 0.099585 | 2.06849591120053E-05 |
| rs11187845 | 10 | 96060198 | eQTLGen | eQTLGen cis eQTLs | PDLIM1 | A | 1.3348E-07 | -5.274 | 0.0004511560874741 |
| rs11187907 | 10 | 96125361 | GTEx/v8 | Muscle Skeletal | NOC3L | T | 1.34121E-07 | 0.270996 | 7.35621E-14 |
| rs11187840 | 10 | 96050351 | PsychENCODE | PsychENCODE eQTLs | NOC3L | G | 1.38003E-07 | 0.0935981 | 2.14874417863028E-05 |
| rs11187838 | 10 | 96038686 | GTEx/v8 | Skin Not Sun Exposed Suprapubic | NOC3L | A | 1.42965E-07 | 0.146108 | 2.01564E-23 |
| rs11187838 | 10 | 96038686 | GTEx/v8 | Heart Left Ventricle | NOC3L | A | 1.43393E-07 | 0.187213 | 3.09268E-10 |
| rs11187890 | 10 | 96102758 | GTEx/v8 | Muscle Skeletal | NOC3L | C | 1.52809E-07 | 0.267003 | 7.35621E-14 |
| rs11187897 | 10 | 96106603 | PsychENCODE | PsychENCODE eQTLs | NOC3L | G | 1.5365E-07 | 0.0982801 | 2.37379907205247E-05 |
| rs74825688 | 10 | 96119687 | PsychENCODE | PsychENCODE eQTLs | NOC3L | A | 1.58469E-07 | 0.0973335 | 2.4430127491122E-05 |
| rs11187895 | 10 | 96106240 | GTEx/v8 | Muscle Skeletal | NOC3L | C | 1.60775E-07 | 0.268698 | 7.35621E-14 |
| rs11187897 | 10 | 96106603 | GTEx/v8 | Muscle Skeletal | NOC3L | G | 1.60775E-07 | 0.268698 | 7.35621E-14 |

| | | | | | | | | | |
|---|---|---|---|---|---|---|---|---|---|
| rs11187826 | 10 | 95988042 | GTEx/v8 | Cells Cultured fibroblasts | PLCE1-AS1 | G | 1.96608E-07 | -0.25953 | 3.86001E-30 |
| rs11187838 | 10 | 96038686 | GTEx/v8 | Pancreas | NOC3L | A | 2.08079E-07 | 0.254395 | 9.42454E-11 |
| rs7096883 | 10 | 96069149 | GTEx/v8 | Adipose Visceral Omentum | PLCE1-AS1 | A | 2.1392E-07 | -0.384583 | 0.000644523 |
| rs11187826 | 10 | 95988042 | eQTLcatalogue | GENCORDge T-cell | PLCE1 | A | 2.19202E-07 | 0.384369 | 0.00109601 |
| rs2274224 | 10 | 96039597 | GTEx/v8 | Esophagus Muscularis | NOC3L | C | 2.22318E-07 | 0.14706 | 2.7232299999999997E-20 |
| rs11187840 | 10 | 96050351 | eQTLGen | eQTLGen cis eQTLs | NOC3L | G | 2.3916E-07 | 5.166 | 0.0007907845579078 |
| rs3740359 | 10 | 96100119 | eQTLcatalogue | GENCORDge T-cell | PLCE1 | C | 2.42044E-07 | 0.403517 | 0.00121022 |
| rs10786156 | 10 | 96014622 | GTEx/v8 | Pancreas | NOC3L | G | 2.53785E-07 | 0.25422 | 9.42454E-11 |
| rs74825688 | 10 | 96119687 | GTEx/v8 | Muscle Skeletal | NOC3L | A | 2.59297E-07 | 0.26993 | 7.35621E-14 |
| rs3891783 | 10 | 96015793 | GTEx/v8 | Pancreas | NOC3L | G | 2.63886E-07 | 0.253623 | 9.42454E-11 |
| rs57866767 | 10 | 96023077 | GTEx/v8 | Pancreas | NOC3L | C | 2.63886E-07 | 0.253623 | 9.42454E-11 |
| rs75017201 | 10 | 96055152 | eQTLGen | eQTLGen cis eQTLs | NOC3L | T | 2.7328E-07 | 5.1411 | 0.0009131569139023 |
| rs79193572 | 10 | 96119713 | GTEx/v8 | Muscle Skeletal | NOC3L | G | 2.73463E-07 | 0.271213 | 7.35621E-14 |
| rs11187842 | 10 | 96052511 | eQTLGen | eQTLGen cis eQTLs | NOC3L | T | 2.7357E-07 | 5.1409 | 0.0009131569139023 |
| rs752140 | 10 | 96071396 | GTEx/v8 | Adipose Visceral Omentum | PLCE1-AS1 | T | 2.74207E-07 | -0.370555 | 0.000644523 |
| rs11187838 | 10 | 96038686 | GTEx/v8 | Colon Transverse | NOC3L | A | 2.82625E-07 | 0.148369 | 4.35915E-08 |
| rs3740365 | 10 | 96053239 | eQTLGen | eQTLGen cis eQTLs | NOC3L | A | 2.8395E-07 | 5.134 | 0.0009374805984975 |
| rs113544984 | 10 | 96127448 | GTEx/v8 | Muscle Skeletal | NOC3L | A | 2.88143E-07 | 0.271515 | 7.35621E-14 |
| rs11187838 | 10 | 96038686 | eQTLcatalogue | TwinsUK ge skin | NOC3L | G | 2.89308E-07 | 0.0989956 | 0.00144654 |
| rs79746649 | 10 | 96120218 | GTEx/v8 | Muscle Skeletal | NOC3L | T | 2.93423E-07 | 0.267286 | 7.35621E-14 |
| rs3740360 | 10 | 96025491 | GTEx/v8 | Cells Cultured fibroblasts | PLCE1-AS1 | C | 3.03193E-07 | -0.26005 | 3.86001E-30 |
| rs17109875 | 10 | 96026575 | GTEx/v8 | Cells Cultured fibroblasts | PLCE1-AS1 | C | 3.03193E-07 | -0.26005 | 3.86001E-30 |
| rs11187837 | 10 | 96035980 | GTEx/v8 | Cells Cultured fibroblasts | PLCE1-AS1 | C | 3.03193E-07 | -0.26005 | 3.86001E-30 |
| rs7101207 | 10 | 96122857 | GTEx/v8 | Muscle Skeletal | NOC3L | A | 3.07502E-07 | 0.26767 | 7.35621E-14 |
| rs112252876 | 10 | 96123355 | GTEx/v8 | Muscle Skeletal | NOC3L | T | 3.07502E-07 | 0.26767 | 7.35621E-14 |
| rs3781266 | 10 | 96052747 | eQTLGen | eQTLGen cis eQTLs | NOC3L | G | 3.0781E-07 | 5.1188 | 0.0009988212668279 |
| rs778275808 | 10 | 96114212 | GTEx/v8 | Muscle Skeletal | NOC3L | T | 3.11067E-07 | 0.268918 | 7.35621E-14 |
| rs10509672 | 10 | 96115957 | GTEx/v8 | Muscle Skeletal | NOC3L | C | 3.11067E-07 | 0.268918 | 7.35621E-14 |
| rs753724 | 10 | 96051417 | GTEx/v8 | Adipose Visceral Omentum | PLCE1-AS1 | T | 3.13234E-07 | -0.372223 | 0.000644523 |
| rs11187842 | 10 | 96052511 | GTEx/v8 | Adipose Visceral Omentum | PLCE1-AS1 | T | 3.13234E-07 | -0.372223 | 0.000644523 |
| rs3781266 | 10 | 96052747 | GTEx/v8 | Adipose Visceral Omentum | PLCE1-AS1 | G | 3.13234E-07 | -0.372223 | 0.000644523 |
| rs3740365 | 10 | 96053239 | GTEx/v8 | Adipose Visceral Omentum | PLCE1-AS1 | A | 3.13234E-07 | -0.372223 | 0.000644523 |
| rs12220091 | 10 | 96053689 | GTEx/v8 | Adipose Visceral Omentum | PLCE1-AS1 | T | 3.13234E-07 | -0.372223 | 0.000644523 |
| rs11187890 | 10 | 96102758 | PsychENCODE | PsychENCODE eQTLs | NOC3L | C | 3.24704E-07 | 0.09449 | 4.7507000772688E-05 |
| rs12220091 | 10 | 96053689 | eQTLGen | eQTLGen cis eQTLs | NOC3L | T | 3.249E-07 | 5.1085 | 0.0010662699150703 |
| rs753724 | 10 | 96051417 | eQTLGen | eQTLGen cis eQTLs | NOC3L | T | 3.294E-07 | 5.1058 | 0.0010722697409198 |
| rs11187870 | 10 | 96087866 | eQTLGen | eQTLGen cis eQTLs | HELLS | C | 3.3185E-07 | 5.1046 | 0.001078267335936 |
| rs112252876 | 10 | 96123355 | PsychENCODE | PsychENCODE eQTLs | NOC3L | T | 3.46649E-07 | 0.0955999 | 5.04639294195705E-05 |

| | | | | | | | | | |
|---|---|---|---|---|---|---|---|---|---|
| rs7080472 | 10 | 96012950 | eQTLcatalogue | TwinsUK ge skin | NOC3L | G | 3.47022E-07 | 0.0981732 | 0.00173511 |
| rs7100626 | 10 | 96069674 | GTEx/v8 | Adipose Visceral Omentum | PLCE1-AS1 | A | 3.47683E-07 | -0.384603 | 0.000644523 |
| rs11187852 | 10 | 96070132 | GTEx/v8 | Adipose Visceral Omentum | PLCE1-AS1 | A | 3.47683E-07 | -0.384603 | 0.000644523 |
| rs10786156 | 10 | 96014622 | eQTLcatalogue | TwinsUK ge skin | NOC3L | C | 3.48198E-07 | 0.0985294 | 0.00174099 |
| rs3891783 | 10 | 96015793 | eQTLcatalogue | TwinsUK ge skin | NOC3L | C | 3.48198E-07 | 0.0985294 | 0.00174099 |
| rs57866767 | 10 | 96023077 | eQTLcatalogue | TwinsUK ge skin | NOC3L | T | 3.48198E-07 | 0.0985294 | 0.00174099 |
| rs2274224 | 10 | 96039597 | GTEx/v8 | Pancreas | NOC3L | C | 3.48768E-07 | 0.251721 | 9.42454E-11 |
| rs79746649 | 10 | 96120218 | PsychENCODE | PsychENCODE eQTLs | NOC3L | T | 3.54686E-07 | 0.094936 | 5.1543436990261E-05 |
| rs75409190 | 10 | 96072425 | GTEx/v8 | Muscle Skeletal | NOC3L | T | 3.61127E-07 | 0.257218 | 7.35621E-14 |
| rs771106747 | 10 | 96072873 | GTEx/v8 | Muscle Skeletal | NOC3L | C | 3.61127E-07 | 0.257218 | 7.35621E-14 |
| rs749295068 | 10 | 96073508 | GTEx/v8 | Muscle Skeletal | NOC3L | A | 3.61127E-07 | 0.257218 | 7.35621E-14 |
| rs11187856 | 10 | 96076869 | GTEx/v8 | Muscle Skeletal | NOC3L | A | 3.61127E-07 | 0.257218 | 7.35621E-14 |
| rs11187882 | 10 | 96095861 | GTEx/v8 | Muscle Skeletal | NOC3L | C | 3.61127E-07 | 0.257218 | 7.35621E-14 |
| rs10786156 | 10 | 96014622 | GTEx/v8 | Esophagus Muscularis | NOC3L | G | 3.66095E-07 | 0.145079 | 2.7232299999999997E-20 |
| rs7101207 | 10 | 96122857 | PsychENCODE | PsychENCODE eQTLs | NOC3L | A | 3.79043E-07 | 0.0952988 | 5.47969109279773E-05 |
| rs11187863 | 10 | 96081457 | GTEx/v8 | Adipose Visceral Omentum | PLCE1-AS1 | G | 3.88596E-07 | -0.364458 | 0.000644523 |
| rs11187866 | 10 | 96085991 | GTEx/v8 | Adipose Visceral Omentum | PLCE1-AS1 | G | 3.88596E-07 | -0.364458 | 0.000644523 |
| rs11187883 | 10 | 96096866 | GTEx/v8 | Adipose Visceral Omentum | PLCE1-AS1 | C | 3.88596E-07 | -0.364458 | 0.000644523 |
| rs11187826 | 10 | 95988042 | GTEx/v8 | Cells EBV-transformed lymphocytes | PLCE1 | G | 3.8943E-07 | 0.687209 | 1.20882E-07 |
| rs7100626 | 10 | 96069674 | GTEx/v8 | Muscle Skeletal | NOC3L | A | 3.96229E-07 | 0.253129 | 7.35621E-14 |
| rs11187852 | 10 | 96070132 | GTEx/v8 | Muscle Skeletal | NOC3L | A | 3.96229E-07 | 0.253129 | 7.35621E-14 |
| rs11187877 | 10 | 96092121 | GTEx/v8 | Muscle Skeletal | NOC3L | A | 3.96229E-07 | 0.253129 | 7.35621E-14 |
| rs11187844 | 10 | 96056629 | eQTLcatalogue | TwinsUK ge LCL | PLCE1 | C | 3.97134E-07 | 0.0760026 | 0.00198567 |
| rs11187864 | 10 | 96082506 | GTEx/v8 | Muscle Skeletal | NOC3L | T | 4.12789E-07 | 0.254895 | 7.35621E-14 |
| rs145707916 | 10 | 96092992 | GTEx/v8 | Muscle Skeletal | NOC3L | GA | 4.12789E-07 | 0.254895 | 7.35621E-14 |
| rs750148755 | 10 | 96095271 | GTEx/v8 | Muscle Skeletal | NOC3L | C | 4.12789E-07 | 0.254895 | 7.35621E-14 |
| rs11187844 | 10 | 96056629 | PsychENCODE | PsychENCODE eQTLs | NOC3L | A | 4.17472E-07 | 0.0796474 | 5.99069241685944E-05 |
| rs3740360 | 10 | 96025491 | eQTLcatalogue | GENCORD ge LCL | PLCE1 | A | 4.30425E-07 | 0.214894 | 0.00215212 |
| rs17109875 | 10 | 96026575 | eQTLcatalogue | GENCORD ge LCL | PLCE1 | T | 4.30538E-07 | 0.214855 | 0.00215269 |
| rs11187837 | 10 | 96035980 | eQTLcatalogue | GENCORD ge LCL | PLCE1 | T | 4.31649E-07 | 0.214915 | 0.00215825 |
| rs10786156 | 10 | 96014622 | GTEx/v8 | Colon Transverse | NOC3L | G | 4.43133E-07 | 0.147768 | 4.35915E-08 |
| rs3891783 | 10 | 96015793 | GTEx/v8 | Colon Transverse | NOC3L | G | 4.43133E-07 | 0.147768 | 4.35915E-08 |
| rs57866767 | 10 | 96023077 | GTEx/v8 | Colon Transverse | NOC3L | C | 4.43133E-07 | 0.147768 | 4.35915E-08 |
| rs11187840 | 10 | 96050351 | GTEx/v8 | Adipose Visceral Omentum | PLCE1-AS1 | G | 4.88307E-07 | -0.372062 | 0.000644523 |
| rs75017201 | 10 | 96055152 | GTEx/v8 | Adipose Visceral Omentum | PLCE1-AS1 | T | 4.88307E-07 | -0.372062 | 0.000644523 |
| rs7101207 | 10 | 96122857 | GTEx/v8 | Adipose Visceral Omentum | PLCE1-AS1 | A | 4.96061E-07 | -0.391535 | 0.000644523 |
| rs112252876 | 10 | 96123355 | GTEx/v8 | Adipose Visceral Omentum | PLCE1-AS1 | T | 4.96061E-07 | -0.391535 | 0.000644523 |
| rs11187890 | 10 | 96102758 | GTEx/v8 | Adipose Visceral Omentum | PLCE1-AS1 | C | 4.97744E-07 | -0.382886 | 0.000644523 |

| | | | | | | | | | |
|---|---|---|---|---|---|---|---|---|---|
| rs3891783 | 10 | 96015793 | GTEx/v8 | Esophagus Muscularis | NOC3L | G | 4.98044E-07 | 0.143689 | 2.7232299999999997E-20 |
| rs57866767 | 10 | 96023077 | GTEx/v8 | Esophagus Muscularis | NOC3L | C | 4.98044E-07 | 0.143689 | 2.7232299999999997E-20 |
| rs11187877 | 10 | 96092121 | GTEx/v8 | Adipose Visceral Omentum | PLCE1-AS1 | A | 5.00681E-07 | -0.377836 | 0.000644523 |
| rs11187838 | 10 | 96038686 | GTEx/v8 | Skin Sun Exposed Lower leg | PLCE1 | A | 5.53023E-07 | 0.1259 | 0.000666445 |
| rs7080472 | 10 | 96012950 | GTEx/v8 | Heart Left Ventricle | NOC3L | T | 5.53501E-07 | 0.180376 | 3.09268E-10 |
| rs2274224 | 10 | 96039597 | GTEx/v8 | Colon Transverse | NOC3L | C | 5.7305E-07 | 0.145736 | 4.35915E-08 |
| rs10786156 | 10 | 96014622 | GTEx/v8 | Skin Sun Exposed Lower leg | PLCE1 | G | 5.83892E-07 | 0.125559 | 0.000666445 |
| rs11187870 | 10 | 96087866 | GTEx/v8 | Muscle Skeletal | NOC3L | C | 5.88352E-07 | 0.253513 | 7.35621E-14 |
| rs7080472 | 10 | 96012950 | GTEx/v8 | Artery Aorta | NOC3L | T | 5.98593E-07 | 0.163228 | 1.86975E-08 |
| rs11187845 | 10 | 96060198 | GTEx/v8 | Adipose Visceral Omentum | PLCE1-AS1 | A | 6.06255E-07 | -0.367463 | 0.000644523 |
| rs11187847 | 10 | 96063440 | GTEx/v8 | Adipose Visceral Omentum | PLCE1-AS1 | G | 6.06255E-07 | -0.367463 | 0.000644523 |
| rs10786156 | 10 | 96014622 | GTEx/v8 | Artery Aorta | NOC3L | G | 6.18786E-07 | 0.162455 | 1.86975E-08 |
| rs7080472 | 10 | 96012950 | GTEx/v8 | Colon Transverse | NOC3L | T | 6.21534E-07 | 0.145514 | 4.35915E-08 |
| rs3891783 | 10 | 96015793 | GTEx/v8 | Skin Sun Exposed Lower leg | PLCE1 | G | 6.68793E-07 | 0.125003 | 0.000666445 |
| rs57866767 | 10 | 96023077 | GTEx/v8 | Skin Sun Exposed Lower leg | PLCE1 | C | 6.68793E-07 | 0.125003 | 0.000666445 |
| rs7078844 | 10 | 96081140 | GTEx/v8 | Nerve Tibial | HELLS | C | 7.10111E-07 | -0.252012 | 5.533110000000001E-26 |
| rs7080472 | 10 | 96012950 | GTEx/v8 | Pancreas | NOC3L | T | 7.15314E-07 | 0.239439 | 9.42454E-11 |
| rs3891783 | 10 | 96015793 | GTEx/v8 | Artery Tibial | PLCE1 | G | 7.18774E-07 | 0.123127 | 1.21915E-07 |
| rs57866767 | 10 | 96023077 | GTEx/v8 | Artery Tibial | PLCE1 | C | 7.18774E-07 | 0.123127 | 1.21915E-07 |
| 10:96112261_AAAAAAAC_A | 10 | 96112261 | eQTLcatalogue | GENCORDge T-cell | PLCE1 | AAAAAAAC | 7.20664E-07 | 0.382443 | 0.00360332 |
| rs7080472 | 10 | 96012950 | GTEx/v8 | Artery Tibial | PLCE1 | T | 8.33992E-07 | 0.123308 | 1.21915E-07 |
| rs79746649 | 10 | 96120218 | GTEx/v8 | Adipose Visceral Omentum | PLCE1-AS1 | T | 8.5631E-07 | -0.38324 | 0.000644523 |
| rs11187838 | 10 | 96038686 | GTEx/v8 | Esophagus Muscularis | NOC3L | A | 8.82021E-07 | 0.140769 | 2.7232299999999997E-20 |
| rs79746649 | 10 | 96120218 | eQTLcatalogue | GENCORDge T-cell | PLCE1 | C | 9.21774E-07 | 0.38583 | 0.00460887 |
| rs752140 | 10 | 96071396 | eQTLcatalogue | GENCORDge T-cell | PLCE1 | C | 9.39565E-07 | 0.386047 | 0.00469783 |
| rs11187890 | 10 | 96102758 | eQTLcatalogue | GENCORDge T-cell | PLCE1 | T | 9.44539E-07 | 0.385938 | 0.00472269 |
| rs11187882 | 10 | 96095861 | eQTLcatalogue | GENCORDge T-cell | PLCE1 | A | 9.46492E-07 | 0.385876 | 0.00473246 |
| rs11187856 | 10 | 96076869 | eQTLcatalogue | GENCORDge T-cell | PLCE1 | G | 9.48129E-07 | 0.385897 | 0.00474064 |
| rs749295068 | 10 | 96073508 | eQTLcatalogue | GENCORDge T-cell | PLCE1 | AGAT | 9.48173E-07 | 0.385896 | 0.00474087 |
| rs771106747 | 10 | 96072873 | eQTLcatalogue | GENCORDge T-cell | PLCE1 | CTA | 9.50361E-07 | 0.385857 | 0.00475181 |
| rs11187863 | 10 | 96081457 | eQTLcatalogue | GENCORDge T-cell | PLCE1 | T | 9.50792E-07 | 0.385788 | 0.00475396 |
| rs11187864 | 10 | 96082506 | eQTLcatalogue | GENCORDge T-cell | PLCE1 | C | 9.50792E-07 | 0.385788 | 0.00475396 |
| rs11187866 | 10 | 96085991 | eQTLcatalogue | GENCORDge T-cell | PLCE1 | C | 9.50792E-07 | 0.385788 | 0.00475396 |
| rs11187870 | 10 | 96087866 | eQTLcatalogue | GENCORDge T-cell | PLCE1 | G | 9.50792E-07 | 0.385788 | 0.00475396 |
| rs11187877 | 10 | 96092121 | eQTLcatalogue | GENCORDge T-cell | PLCE1 | G | 9.50792E-07 | 0.385788 | 0.00475396 |
| rs145707916 | 10 | 96092992 | eQTLcatalogue | GENCORDge T-cell | PLCE1 | G | 9.50792E-07 | 0.385788 | 0.00475396 |
| rs7100626 | 10 | 96069674 | eQTLcatalogue | GENCORDge T-cell | PLCE1 | C | 9.51672E-07 | 0.385791 | 0.00475836 |
| rs11187852 | 10 | 96070132 | eQTLcatalogue | GENCORDge T-cell | PLCE1 | G | 9.51672E-07 | 0.385791 | 0.00475836 |
| rs75409190 | 10 | 96072425 | eQTLcatalogue | GENCORDge T-cell | PLCE1 | C | 9.51792E-07 | 0.385813 | 0.00475896 |
| rs750148755 | 10 | 96095271 | eQTLcatalogue | GENCORDge T-cell | PLCE1 | CATTTT | 9.54195E-07 | 0.385738 | 0.00477098 |

| | | | | | | | | | |
|---|---|---|---|---|---|---|---|---|---|
| rs3740359 | 10 | 96100119 | DICE | T CD4 TH1 | PLCE1 | T | 9.6971259049E-07 | 0.898080559827 | 0.049 |
| rs34309017 | 10 | 96061619 | GTEx/v8 | Adipose Visceral Omentum | PLCE1-AS1 | A | 9.70292E-07 | -0.366366 | 0.000644523 |
| rs12217792 | 10 | 96062386 | GTEx/v8 | Adipose Visceral Omentum | PLCE1-AS1 | C | 9.70292E-07 | -0.366366 | 0.000644523 |
| rs11187847 | 10 | 96063440 | eQTLcatalogue | GENCORDge LCL | PLCE1 | C | 9.70592E-07 | 0.233046 | 0.00485296 |
| rs11187895 | 10 | 96106240 | eQTLcatalogue | GENCORDge T-cell | PLCE1 | G | 9.72079E-07 | 0.38546 | 0.00486039 |
| rs11187897 | 10 | 96106603 | eQTLcatalogue | GENCORDge T-cell | PLCE1 | A | 9.73083E-07 | 0.385427 | 0.00486541 |
| rs11187883 | 10 | 96096866 | eQTLcatalogue | GENCORDge T-cell | PLCE1 | T | 9.73666E-07 | 0.385218 | 0.00486833 |
| rs10509672 | 10 | 96115957 | eQTLcatalogue | GENCORDge T-cell | PLCE1 | G | 9.83222E-07 | 0.385218 | 0.00491611 |
| rs778275808 | 10 | 96114212 | eQTLcatalogue | GENCORDge T-cell | PLCE1 | TCTC | 9.90757E-07 | 0.385175 | 0.00495378 |
| rs79193572 | 10 | 96119713 | eQTLcatalogue | GENCORDge T-cell | PLCE1 | A | 1.00326E-06 | 0.384625 | 0.0050163 |
| rs753724 | 10 | 96051417 | eQTLGen | eQTLGen cis eQTLs | PDLIM1 | T | 1.0311E-06 | -4.8856 | 0.0031823745410036 |
| rs2274224 | 10 | 96039597 | GTEx/v8 | Artery Aorta | NOC3L | C | 1.05756E-06 | 0.158624 | 1.86975E-08 |
| rs7080472 | 10 | 96012950 | GTEx/v8 | Colon Sigmoid | NOC3L | T | 1.10378E-06 | 0.193204 | 1.73346E-11 |
| rs74825688 | 10 | 96119687 | eQTLcatalogue | GENCORDge T-cell | PLCE1 | G | 1.1056E-06 | 0.382039 | 0.005528 |
| rs10786156 | 10 | 96014622 | GTEx/v8 | Artery Tibial | PLCE1 | G | 1.11843E-06 | 0.12099 | 1.21915E-07 |
| rs11187842 | 10 | 96052511 | eQTLGen | eQTLGen cis eQTLs | PDLIM1 | T | 1.1205E-06 | -4.869 | 0.003400403645037 |
| rs3740365 | 10 | 96053239 | eQTLGen | eQTLGen cis eQTLs | PDLIM1 | A | 1.1515E-06 | -4.8636 | 0.0034784203447854 |
| rs74825688 | 10 | 96119687 | GTEx/v8 | Adipose Visceral Omentum | PLCE1-AS1 | A | 1.16396E-06 | -0.375083 | 0.000644523 |
| rs11187845 | 10 | 96060198 | eQTLcatalogue | GENCORDge T-cell | PLCE1 | C | 1.18769E-06 | 0.380044 | 0.00593845 |
| rs34309017 | 10 | 96061619 | eQTLcatalogue | GENCORDge T-cell | PLCE1 | ACTT | 1.19242E-06 | 0.37961 | 0.0059621 |
| rs12217792 | 10 | 96062386 | eQTLcatalogue | GENCORDge T-cell | PLCE1 | T | 1.19698E-06 | 0.379327 | 0.0059849 |
| rs11187847 | 10 | 96063440 | eQTLcatalogue | GENCORDge T-cell | PLCE1 | C | 1.20172E-06 | 0.379013 | 0.0060086 |
| rs2274224 | 10 | 96039597 | GTEx/v8 | Skin Sun Exposed Lower leg | PLCE1 | C | 1.20492E-06 | 0.12172 | 0.000666445 |
| rs7096883 | 10 | 96069149 | eQTLcatalogue | GENCORDge T-cell | PLCE1 | G | 1.21204E-06 | 0.378282 | 0.0060602 |
| rs2274224 | 10 | 96039597 | GTEx/v8 | Artery Tibial | PLCE1 | C | 1.26595E-06 | 0.120273 | 1.21915E-07 |
| rs3891783 | 10 | 96015793 | GTEx/v8 | Artery Aorta | NOC3L | G | 1.312E-06 | 0.158283 | 1.86975E-08 |
| rs57866767 | 10 | 96023077 | GTEx/v8 | Artery Aorta | NOC3L | C | 1.312E-06 | 0.158283 | 1.86975E-08 |
| rs3740359 | 10 | 96100119 | GTEx/v8 | Muscle Skeletal | NOC3L | T | 1.31362E-06 | 0.221014 | 7.35621E-14 |
| rs75017201 | 10 | 96055152 | eQTLcatalogue | GENCORDge T-cell | PLCE1 | C | 1.33148E-06 | 0.367203 | 0.0066574 |
| rs12220091 | 10 | 96053689 | eQTLcatalogue | GENCORDge T-cell | PLCE1 | C | 1.33264E-06 | 0.367032 | 0.0066632 |
| rs3740365 | 10 | 96053239 | eQTLcatalogue | GENCORDge T-cell | PLCE1 | T | 1.33285E-06 | 0.367006 | 0.00666425 |
| rs753724 | 10 | 96051417 | eQTLcatalogue | GENCORDge T-cell | PLCE1 | G | 1.3329E-06 | 0.367893 | 0.0066645 |
| rs3781266 | 10 | 96052747 | eQTLcatalogue | GENCORDge T-cell | PLCE1 | A | 1.33329E-06 | 0.366955 | 0.00666645 |
| rs11187842 | 10 | 96052511 | eQTLcatalogue | GENCORDge T-cell | PLCE1 | C | 1.33345E-06 | 0.366938 | 0.00666725 |
| rs11187840 | 10 | 96050351 | eQTLcatalogue | GENCORDge T-cell | PLCE1 | A | 1.36086E-06 | 0.368475 | 0.0068043 |
| rs11187840 | 10 | 96050351 | eQTLGen | eQTLGen cis eQTLs | PDLIM1 | G | 1.3708E-06 | -4.8291 | 0.0041417591801878 |
| rs769015184 | 10 | 96049708 | eQTLcatalogue | GENCORDge T-cell | PLCE1 | AC | 1.38142E-06 | 0.36852 | 0.0069071 |
| rs11187844 | 10 | 96056629 | BIOSQTL | BIOS eQTL geneLevel | TBC1D12 | A | 1.42697E-06 | 4.8213458 | 0.0011833705299216 |
| rs75409190 | 10 | 96072425 | GTEx/v8 | Adipose Visceral Omentum | PLCE1-AS1 | T | 1.56355E-06 | -0.365914 | 0.000644523 |
| rs771106747 | 10 | 96072873 | GTEx/v8 | Adipose Visceral Omentum | PLCE1-AS1 | C | 1.56355E-06 | -0.365914 | 0.000644523 |

| | | | | | | | | | |
|---|---|---|---|---|---|---|---|---|---|
| rs11187856 | 10 | 96076869 | GTEx/v8 | Adipose Visceral Omentum | PLCE1-AS1 | A | 1.56355E-06 | -0.365914 | 0.000644523 |
| rs75017201 | 10 | 96055152 | eQTLGen | eQTLGen cis eQTLs | PDLIM1 | T | 1.5803E-06 | -4.8007 | 0.0046780776024852 |
| rs2077218 | 10 | 96071561 | GTEx/v8 | Nerve Tibial | HELLS | A | 1.61786E-06 | -0.253004 | 5.533110000000001E-26 |
| rs11187838 | 10 | 96038686 | BIOSQTL | BIOS eQTL geneLevel | PLCE1 | A | 1.65622E-06 | 4.7915542 | 0.0013530961111398 |
| rs3740359 | 10 | 96100119 | GTEx/v8 | Cells EBV-transformed lymphocytes | PLCE1 | T | 1.67783E-06 | 0.600178 | 1.20882E-07 |
| rs7080472 | 10 | 96012950 | GTEx/v8 | Thyroid | TBC1D12 | T | 1.69106E-06 | -0.142239 | 1.4449E-10 |
| rs3781266 | 10 | 96052747 | eQTLGen | eQTLGen cis eQTLs | PDLIM1 | G | 1.7971E-06 | -4.7749 | 0.0053287912890078 |
| rs7078844 | 10 | 96081140 | BIOSQTL | BIOS eQTL geneLevel | TBC1D12 | T | 1.81863E-06 | -4.77257 | 0.0014319563201813 |
| rs12220091 | 10 | 96053689 | eQTLGen | eQTLGen cis eQTLs | PDLIM1 | T | 1.9017E-06 | -4.7636 | 0.0055572444822764 |
| rs7080472 | 10 | 96012950 | GTEx/v8 | Skin Sun Exposed Lower leg | PLCE1 | T | 1.96357E-06 | 0.12091 | 0.000666445 |
| rs11187863 | 10 | 96081457 | DICE | T CD4 TH1 | PLCE1 | G | 2.1750695116E-06 | 0.887821580882 | 0.049 |
| rs11187866 | 10 | 96085991 | DICE | T CD4 TH1 | PLCE1 | G | 2.1750695116E-06 | 0.887821580882 | 0.049 |
| rs11187877 | 10 | 96092121 | DICE | T CD4 TH1 | PLCE1 | A | 2.1750695116E-06 | 0.887821580882 | 0.049 |
| rs11187883 | 10 | 96096866 | DICE | T CD4 TH1 | PLCE1 | C | 2.1750695116E-06 | 0.887821580882 | 0.049 |
| rs11187890 | 10 | 96102758 | DICE | T CD4 TH1 | PLCE1 | C | 2.1750695116E-06 | 0.887821580882 | 0.049 |
| rs11187864 | 10 | 96082506 | DICE | T CD4 TH1 | PLCE1 | T | 2.18983857164E-06 | 0.900338651896 | 0.049 |
| rs11187870 | 10 | 96087866 | DICE | T CD4 TH1 | PLCE1 | C | 2.18983857164E-06 | 0.900338651896 | 0.049 |
| rs145707916 | 10 | 96092992 | DICE | T CD4 TH1 | PLCE1 | GA | 2.18983857164E-06 | 0.900338651896 | 0.049 |
| rs750148755 | 10 | 96095271 | DICE | T CD4 TH1 | PLCE1 | C | 2.18983857164E-06 | 0.900338651896 | 0.049 |
| rs11187882 | 10 | 96095861 | DICE | T CD4 TH1 | PLCE1 | C | 2.18983857164E-06 | 0.900338651896 | 0.049 |
| rs11187895 | 10 | 96106240 | DICE | T CD4 TH1 | PLCE1 | C | 2.18983857164E-06 | 0.900338651896 | 0.049 |
| rs11187897 | 10 | 96106603 | DICE | T CD4 TH1 | PLCE1 | G | 2.18983857164E-06 | 0.900338651896 | 0.049 |
| rs11187863 | 10 | 96081457 | GTEx/v8 | Thyroid | NOC3L | G | 2.19445E-06 | -0.172139 | 0.00220997 |
| rs11187866 | 10 | 96085991 | GTEx/v8 | Thyroid | NOC3L | G | 2.19445E-06 | -0.172139 | 0.00220997 |
| rs11187883 | 10 | 96096866 | GTEx/v8 | Thyroid | NOC3L | C | 2.19445E-06 | -0.172139 | 0.00220997 |
| rs11187864 | 10 | 96082506 | GTEx/v8 | Adipose Visceral Omentum | PLCE1-AS1 | T | 2.20876E-06 | -0.359052 | 0.000644523 |
| rs145707916 | 10 | 96092992 | GTEx/v8 | Adipose Visceral Omentum | PLCE1-AS1 | GA | 2.20876E-06 | -0.359052 | 0.000644523 |
| rs750148755 | 10 | 96095271 | GTEx/v8 | Adipose Visceral Omentum | PLCE1-AS1 | C | 2.20876E-06 | -0.359052 | 0.000644523 |
| rs11187895 | 10 | 96106240 | GTEx/v8 | Adipose Visceral Omentum | PLCE1-AS1 | C | 2.23083E-06 | -0.36371 | 0.000644523 |
| rs11187897 | 10 | 96106603 | GTEx/v8 | Adipose Visceral Omentum | PLCE1-AS1 | G | 2.23083E-06 | -0.36371 | 0.000644523 |
| rs749295068 | 10 | 96073508 | GTEx/v8 | Adipose Visceral Omentum | PLCE1-AS1 | A | 2.24181E-06 | -0.362577 | 0.000644523 |
| rs11187838 | 10 | 96038686 | GTEx/v8 | Brain Nucleus accumbens basal ganglia | NOC3L | A | 2.24908E-06 | 0.20893 | 0.000164102 |
| rs752140 | 10 | 96071396 | GTEx/v8 | Thyroid | NOC3L | T | 2.28464E-06 | -0.172467 | 0.00220997 |
| rs11187837 | 10 | 96035980 | eQTLcatalogue | GENCORDge T-cell | PLCE1 | T | 2.31571E-06 | 0.335699 | 0.0115785 |
| rs17109875 | 10 | 96026575 | eQTLcatalogue | GENCORDge T-cell | PLCE1 | T | 2.32943E-06 | 0.335494 | 0.0116471 |
| rs3740360 | 10 | 96025491 | eQTLcatalogue | GENCORDge T-cell | PLCE1 | A | 2.33243E-06 | 0.335539 | 0.0116621 |
| 10:96112261_AAAAAAAC_A | 10 | 96112261 | DICE | T CD4 TH1 | PLCE1 | A | 2.33720404313E-06 | 0.948074989332 | 0.049 |
| rs11187845 | 10 | 96060198 | GTEx/v8 | Muscle Skeletal | NOC3L | A | 2.44081E-06 | 0.230282 | 7.35621E-14 |

| | | | | | | | | | |
|---|---|---|---|---|---|---|---|---|---|
| rs11187847 | 10 | 96063440 | GTEx/v8 | Muscle Skeletal | NOC3L | G | 2.44081E-06 | 0.230282 | 7.35621E-14 |
| rs7080472 | 10 | 96012950 | GTEx/v8 | Brain Cerebellar Hemisphere | NOC3L | T | 2.48212E-06 | 0.273389 | 1.25929E-05 |
| rs10786156 | 10 | 96014622 | BIOSQTL | BIOS eQTL geneLevel | PLCE1 | G | 2.48935E-06 | 4.7090961 | 0.0019017998266455 |
| rs3891783 | 10 | 96015793 | BIOSQTL | BIOS eQTL geneLevel | PLCE1 | G | 2.48935E-06 | 4.7090961 | 0.0019017998266455 |
| rs57866767 | 10 | 96023077 | BIOSQTL | BIOS eQTL geneLevel | PLCE1 | C | 2.48935E-06 | 4.7090961 | 0.0019017998266455 |
| rs11187838 | 10 | 96038686 | GTEx/v8 | Artery Aorta | NOC3L | A | 2.9258E-06 | 0.151503 | 1.86975E-08 |
| rs34309017 | 10 | 96061619 | GTEx/v8 | Muscle Skeletal | NOC3L | A | 3.04275E-06 | 0.230309 | 7.35621E-14 |
| rs12217792 | 10 | 96062386 | GTEx/v8 | Muscle Skeletal | NOC3L | C | 3.04275E-06 | 0.230309 | 7.35621E-14 |
| rs11187882 | 10 | 96095861 | GTEx/v8 | Adipose Visceral Omentum | PLCE1-AS1 | C | 3.14149E-06 | -0.35578 | 0.000644523 |
| rs7096883 | 10 | 96069149 | GTEx/v8 | Muscle Skeletal | NOC3L | A | 3.18816E-06 | 0.227145 | 7.35621E-14 |
| rs7101207 | 10 | 96122857 | eQTLcatalogue | BLUEPRINT ge T-cell | NOC3L | G | 3.24912E-06 | 0.109848 | 0.0162456 |
| rs11187906 | 10 | 96124838 | eQTLcatalogue | BLUEPRINT ge T-cell | NOC3L | T | 3.28105E-06 | 0.10976 | 0.0164052 |
| rs11187907 | 10 | 96125361 | eQTLcatalogue | BLUEPRINT ge T-cell | NOC3L | C | 3.28105E-06 | 0.10976 | 0.0164052 |
| rs113544984 | 10 | 96127448 | eQTLcatalogue | BLUEPRINT ge T-cell | NOC3L | C | 3.28105E-06 | 0.10976 | 0.0164052 |
| rs7080472 | 10 | 96012950 | eQTLcatalogue | Quach 2016 ge monocyte Pam3CSK4 | NOC3L | G | 3.41441E-06 | -0.105905 | 0.017072 |
| rs10786156 | 10 | 96014622 | GTEx/v8 | Brain Nucleus accumbens basal ganglia | NOC3L | G | 3.46741E-06 | 0.204556 | 0.000164102 |
| rs3891783 | 10 | 96015793 | GTEx/v8 | Brain Nucleus accumbens basal ganglia | NOC3L | G | 3.46741E-06 | 0.204556 | 0.000164102 |
| rs57866767 | 10 | 96023077 | GTEx/v8 | Brain Nucleus accumbens basal ganglia | NOC3L | C | 3.46741E-06 | 0.204556 | 0.000164102 |
| rs2274224 | 10 | 96039597 | GTEx/v8 | Brain Nucleus accumbens basal ganglia | NOC3L | C | 3.46741E-06 | 0.204556 | 0.000164102 |
| rs778275808 | 10 | 96114212 | DICE | T CD4 TH1 | PLCE1 | T | 3.54266627324E-06 | 0.944577708395 | 0.049 |
| rs10509672 | 10 | 96115957 | DICE | T CD4 TH1 | PLCE1 | C | 3.54266627324E-06 | 0.944577708395 | 0.049 |
| rs74825688 | 10 | 96119687 | DICE | T CD4 TH1 | PLCE1 | A | 3.54266627324E-06 | 0.944577708395 | 0.049 |
| rs79193572 | 10 | 96119713 | DICE | T CD4 TH1 | PLCE1 | G | 3.54266627324E-06 | 0.944577708395 | 0.049 |
| rs11187906 | 10 | 96124838 | DICE | T CD4 TH1 | PLCE1 | A | 3.54266627324E-06 | 0.944577708395 | 0.049 |
| rs11187907 | 10 | 96125361 | DICE | T CD4 TH1 | PLCE1 | T | 3.54266627324E-06 | 0.944577708395 | 0.049 |
| rs113544984 | 10 | 96127448 | DICE | T CD4 TH1 | PLCE1 | A | 3.54266627324E-06 | 0.944577708395 | 0.049 |
| rs79746649 | 10 | 96120218 | DICE | T CD4 TH1 | PLCE1 | T | 3.56488559879E-06 | 0.92935546315 | 0.049 |
| rs7101207 | 10 | 96122857 | DICE | T CD4 TH1 | PLCE1 | A | 3.56488559879E-06 | 0.92935546315 | 0.049 |
| rs112252876 | 10 | 96123355 | DICE | T CD4 TH1 | PLCE1 | T | 3.56488559879E-06 | 0.92935546315 | 0.049 |
| rs11187826 | 10 | 95988042 | eQTLcatalogue | TwinsUK ge LCL | PLCE1 | A | 3.67082E-06 | 0.0676536 | 0.0183541 |
| rs778275808 | 10 | 96114212 | GTEx/v8 | Adipose Visceral Omentum | PLCE1-AS1 | T | 3.89634E-06 | -0.362682 | 0.000644523 |
| rs10509672 | 10 | 96115957 | GTEx/v8 | Adipose Visceral Omentum | PLCE1-AS1 | C | 3.89634E-06 | -0.362682 | 0.000644523 |
| rs79193572 | 10 | 96119713 | GTEx/v8 | Adipose Visceral Omentum | PLCE1-AS1 | G | 3.89634E-06 | -0.362682 | 0.000644523 |
| rs7080472 | 10 | 96012950 | BIOSQTL | BIOS eQTL geneLevel | PLCE1 | T | 3.98429E-06 | 4.6122133 | 0.0028727838883029 |

| | | | | | | | | | |
|---|---|---|---|---|---|---|---|---|---|
| rs3740360 | 10 | 96025491 | eQTLcatalogue | TwinsUK ge LCL | PLCE1 | A | 4.12006E-06 | 0.0676965 | 0.0206003 |
| rs17109875 | 10 | 96026575 | eQTLcatalogue | TwinsUK ge LCL | PLCE1 | T | 4.12006E-06 | 0.0676965 | 0.0206003 |
| rs11187837 | 10 | 96035980 | eQTLcatalogue | TwinsUK ge LCL | PLCE1 | T | 4.23178E-06 | 0.0679173 | 0.0211589 |
| rs11187907 | 10 | 96125361 | GTEx/v8 | Adipose Visceral Omentum | PLCE1-AS1 | T | 4.23293E-06 | -0.353893 | 0.000644523 |
| rs11187838 | 10 | 96038686 | GTEx/v8 | Brain Cerebellar Hemisphere | NOC3L | A | 4.63289E-06 | 0.260394 | 1.25929E-05 |
| rs2274224 | 10 | 96039597 | GTEx/v8 | Brain Cerebellar Hemisphere | NOC3L | C | 4.63289E-06 | 0.260394 | 1.25929E-05 |
| rs11187870 | 10 | 96087866 | GTEx/v8 | Adipose Visceral Omentum | PLCE1-AS1 | C | 4.63963E-06 | -0.352852 | 0.000644523 |
| rs2274224 | 10 | 96039597 | GTEx/v8 | Thyroid | TBC1D12 | C | 4.82673E-06 | -0.135379 | 1.4449E-10 |
| rs10786156 | 10 | 96014622 | GTEx/v8 | Thyroid | TBC1D12 | G | 5.24533E-06 | -0.134688 | 1.4449E-10 |
| rs3891783 | 10 | 96015793 | GTEx/v8 | Thyroid | TBC1D12 | G | 5.26247E-06 | -0.134964 | 1.4449E-10 |
| rs57866767 | 10 | 96023077 | GTEx/v8 | Thyroid | TBC1D12 | C | 5.26247E-06 | -0.134964 | 1.4449E-10 |
| rs7080472 | 10 | 96012950 | GTEx/v8 | Brain Nucleus accumbens basal ganglia | NOC3L | T | 5.39627E-06 | 0.203249 | 0.000164102 |
| rs11187906 | 10 | 96124838 | BIOSQTL | BIOS eQTL geneLevel | HELLS | A | 5.58289E-06 | 4.5416722 | 0.0039545499652674 |
| rs11187906 | 10 | 96124838 | BIOSQTL | BIOS eQTL geneLevel | RP11-119K6.6 | A | 5.58289E-06 | 4.5416722 | 0.0039545499652674 |
| rs3740359 | 10 | 96100119 | GTEx/v8 | Adipose Visceral Omentum | PLCE1-AS1 | T | 5.74462E-06 | -0.311868 | 0.000644523 |
| rs11187838 | 10 | 96038686 | GTEx/v8 | Artery Tibial | PLCE1-AS1 | A | 5.76321E-06 | 0.138868 | 2.47435E-06 |
| rs200197176 | 10 | 96055963 | eQTLcatalogue | GENCORD ge T-cell | PLCE1 | C | 5.83724E-06 | 0.342589 | 0.0291862 |
| rs11187826 | 10 | 95988042 | GTEx/v8 | Adipose Visceral Omentum | PLCE1-AS1 | G | 5.84747E-06 | -0.330764 | 0.000644523 |
| rs752140 | 10 | 96071396 | GTEx/v8 | Whole Blood | TBC1D12 | T | 5.9535E-06 | 0.239486 | 0.00686763 |
| rs7080472 | 10 | 96012950 | GTEx/v8 | Artery Tibial | PLCE1-AS1 | T | 6.16553E-06 | 0.139554 | 2.47435E-06 |
| rs7078844 | 10 | 96081140 | eQTLcatalogue | BrainSeq ge brain | NOC3L | T | 6.27308E-06 | 0.0839191 | 0.0313654 |
| rs2274224 | 10 | 96039597 | GTEx/v8 | Adipose Subcutaneous | NOC3L | C | 6.31202E-06 | 0.114183 | 3.06299E-11 |
| rs10786156 | 10 | 96014622 | GTEx/v8 | Brain Cerebellar Hemisphere | NOC3L | G | 6.40412E-06 | 0.257037 | 1.25929E-05 |
| rs3891783 | 10 | 96015793 | GTEx/v8 | Brain Cerebellar Hemisphere | NOC3L | G | 6.40412E-06 | 0.257037 | 1.25929E-05 |
| rs57866767 | 10 | 96023077 | GTEx/v8 | Brain Cerebellar Hemisphere | NOC3L | C | 6.40412E-06 | 0.257037 | 1.25929E-05 |
| rs10786156 | 10 | 96014622 | GTEx/v8 | Spleen | NOC3L | G | 6.95933E-06 | 0.238458 | 1.49159E-15 |
| rs3891783 | 10 | 96015793 | GTEx/v8 | Spleen | NOC3L | G | 6.95933E-06 | 0.238458 | 1.49159E-15 |
| rs57866767 | 10 | 96023077 | GTEx/v8 | Spleen | NOC3L | C | 6.95933E-06 | 0.238458 | 1.49159E-15 |
| rs2274224 | 10 | 96039597 | GTEx/v8 | Spleen | NOC3L | C | 6.95933E-06 | 0.238458 | 1.49159E-15 |
| rs11187890 | 10 | 96102758 | eQTLcatalogue | BLUEPRINT ge T-cell | NOC3L | T | 7.12973E-06 | 0.104697 | 0.0356487 |
| rs11187895 | 10 | 96106240 | eQTLcatalogue | BLUEPRINT ge T-cell | NOC3L | G | 7.13783E-06 | 0.104684 | 0.0356892 |
| rs11187897 | 10 | 96106603 | eQTLcatalogue | BLUEPRINT ge T-cell | NOC3L | A | 7.13783E-06 | 0.104684 | 0.0356892 |
| rs10509672 | 10 | 96115957 | eQTLcatalogue | BLUEPRINT ge T-cell | NOC3L | G | 7.13783E-06 | 0.104684 | 0.0356892 |
| rs74825688 | 10 | 96119687 | eQTLcatalogue | BLUEPRINT ge T-cell | NOC3L | G | 7.13783E-06 | 0.104684 | 0.0356892 |
| rs79193572 | 10 | 96119713 | eQTLcatalogue | BLUEPRINT ge T-cell | NOC3L | A | 7.13783E-06 | 0.104684 | 0.0356892 |
| rs778275808 | 10 | 96114212 | eQTLcatalogue | BLUEPRINT ge T-cell | NOC3L | TCTC | 7.15045E-06 | 0.104677 | 0.0357522 |
| rs11187838 | 10 | 96038686 | GTEx/v8 | Spleen | NOC3L | A | 7.73019E-06 | 0.237346 | 1.49159E-15 |
| rs749295068 | 10 | 96073508 | GTEx/v8 | Whole Blood | TBC1D12 | A | 7.96594E-06 | 0.249853 | 0.00686763 |
| rs10786156 | 10 | 96014622 | GTEx/v8 | Colon Sigmoid | NOC3L | G | 8.39325E-06 | 0.178254 | 1.73346E-11 |

| | | | | | | | | | |
|---|---|---|---|---|---|---|---|---|---|
| rs3891783 | 10 | 96015793 | GTEx/v8 | Colon Sigmoid | NOC3L | G | 8.39325E-06 | 0.178254 | 1.73346E-11 |
| rs57866767 | 10 | 96023077 | GTEx/v8 | Colon Sigmoid | NOC3L | C | 8.39325E-06 | 0.178254 | 1.73346E-11 |
| rs7080472 | 10 | 96012950 | GTEx/v8 | Adipose Subcutaneo us | NOC3L | T | 9.22157E-06 | 0.113545 | 3.06299E-11 |
| rs113544984 | 10 | 96127448 | GTEx/v8 | Adipose Visceral Omentum | PLCE1-AS1 | A | 9.24866E-06 | -0.349983 | 0.000644523 |
| rs7078844 | 10 | 96081140 | GTEx/v8 | Thyroid | PLCE1-AS1 | C | 9.37332E-06 | -0.158645 | 0.00721889 |
| rs11187844 | 10 | 96056629 | eQTLcatal ogue | GENCORD ge T-cell | PLCE1 | C | 9.51488E-06 | 0.299103 | 0.0475744 |
| rs2274224 | 10 | 96039597 | eQTLcatal ogue | Fairfax 2014 LPS2 | NOC3L | G | 9.7278E-06 | -0.128461 | 0.048639 |
| rs11187890 | 10 | 96102758 | GTEx/v8 | Thyroid | NOC3L | C | 1.02717E-05 | -0.167326 | 0.00220997 |
| rs10786156 | 10 | 96014622 | GTEx/v8 | Prostate | NOC3L | G | 1.05232E-05 | 0.195047 | 7.10224E-08 |
| rs3891783 | 10 | 96015793 | GTEx/v8 | Prostate | NOC3L | G | 1.05232E-05 | 0.195047 | 7.10224E-08 |
| rs57866767 | 10 | 96023077 | GTEx/v8 | Prostate | NOC3L | C | 1.05232E-05 | 0.195047 | 7.10224E-08 |
| rs3740360 | 10 | 96025491 | eQTLGen | eQTLGen cis eQTLs | NOC3L | C | 1.0598E-05 | 7,8427E+05 | 0.0299060977059313 |
| rs7078844 | 10 | 96081140 | GTEx/v8 | Thyroid | PLCE1 | C | 1.07686E-05 | -0.152078 | 1.91428E-05 |
| rs7096883 | 10 | 96069149 | GTEx/v8 | Whole Blood | TBC1D12 | A | 1.08788E-05 | 0.237684 | 0.00686763 |
| rs11187838 | 10 | 96038686 | GTEx/v8 | Colon Sigmoid | NOC3L | A | 1.09622E-05 | 0.174007 | 1.73346E-11 |
| rs7100626 | 10 | 96069674 | GTEx/v8 | Whole Blood | TBC1D12 | A | 1.12809E-05 | 0.242268 | 0.00686763 |
| rs11187852 | 10 | 96070132 | GTEx/v8 | Whole Blood | TBC1D12 | A | 1.12809E-05 | 0.242268 | 0.00686763 |
| rs11187838 | 10 | 96038686 | GTEx/v8 | Skin Not Sun Exposed Suprapubic | RP11-76P2.4 | A | 1.15234E-05 | 0.222711 | 5.98468E-07 |
| rs2274224 | 10 | 96039597 | GTEx/v8 | Colon Sigmoid | NOC3L | C | 1.17053E-05 | 0.173093 | 1.73346E-11 |
| rs7080472 | 10 | 96012950 | GTEx/v8 | Heart Atrial Appendage | NOC3L | T | 1.20317E-05 | 0.164229 | 2.17353E-13 |
| rs17109875 | 10 | 96026575 | eQTLGen | eQTLGen cis eQTLs | NOC3L | C | 1.2032E-05 | 6,8346E+05 | 0.0338453327007472 |
| rs10786156 | 10 | 96014622 | GTEx/v8 | Skin Not Sun Exposed Suprapubic | RP11-76P2.4 | G | 1.21338E-05 | 0.219968 | 5.98468E-07 |
| rs3891783 | 10 | 96015793 | GTEx/v8 | Skin Not Sun Exposed Suprapubic | RP11-76P2.4 | G | 1.21338E-05 | 0.219968 | 5.98468E-07 |
| rs57866767 | 10 | 96023077 | GTEx/v8 | Skin Not Sun Exposed Suprapubic | RP11-76P2.4 | C | 1.21338E-05 | 0.219968 | 5.98468E-07 |
| rs10786156 | 10 | 96014622 | GTEx/v8 | Spleen | RP11-76P2.4 | G | 1.21461E-05 | 0.37517 | 9.37816E-08 |
| rs3891783 | 10 | 96015793 | GTEx/v8 | Spleen | RP11-76P2.4 | G | 1.21461E-05 | 0.37517 | 9.37816E-08 |
| rs57866767 | 10 | 96023077 | GTEx/v8 | Spleen | RP11-76P2.4 | C | 1.21461E-05 | 0.37517 | 9.37816E-08 |
| rs2274224 | 10 | 96039597 | GTEx/v8 | Spleen | RP11-76P2.4 | C | 1.21461E-05 | 0.37517 | 9.37816E-08 |
| rs11187838 | 10 | 96038686 | GTEx/v8 | Brain Hippocampu s | NOC3L | A | 1.21767E-05 | 0.237097 | 0.000170124 |
| rs2274224 | 10 | 96039597 | GTEx/v8 | Brain Hippocampu s | NOC3L | C | 1.21767E-05 | 0.237097 | 0.000170124 |
| rs7101207 | 10 | 96122857 | BIOSQTL | BIOS eQTL geneLevel | HELLS | A | 1.23339E-05 | 4.3716277 | 0.0083205004812319 |
| rs7101207 | 10 | 96122857 | BIOSQTL | BIOS eQTL geneLevel | RP11-119K6.6 | A | 1.23339E-05 | 4.3716277 | 0.0083205004812319 |
| rs112252876 | 10 | 96123355 | BIOSQTL | BIOS eQTL geneLevel | HELLS | T | 1.23339E-05 | 4.3716277 | 0.0083205004812319 |
| rs112252876 | 10 | 96123355 | BIOSQTL | BIOS eQTL geneLevel | RP11-119K6.6 | T | 1.23339E-05 | 4.3716277 | 0.0083205004812319 |
| rs2274224 | 10 | 96039597 | GTEx/v8 | Prostate | NOC3L | C | 1.26886E-05 | 0.192241 | 7.10224E-08 |
| rs7080472 | 10 | 96012950 | GTEx/v8 | Skin Not Sun Exposed Suprapubic | PLCE1 | T | 1.28165E-05 | 0.13641 | 9.09957E-06 |
| rs11187863 | 10 | 96081457 | GTEx/v8 | Whole Blood | TBC1D12 | G | 1.29458E-05 | 0.229961 | 0.00686763 |
| rs11187866 | 10 | 96085991 | GTEx/v8 | Whole Blood | TBC1D12 | G | 1.29458E-05 | 0.229961 | 0.00686763 |
| rs11187883 | 10 | 96096866 | GTEx/v8 | Whole Blood | TBC1D12 | C | 1.29458E-05 | 0.229961 | 0.00686763 |

| | | | | | | | | | |
|---|---|---|---|---|---|---|---|---|---|
| rs7096883 | 10 | 96069149 | DICE | T CD4 TH1 | PLCE1 | A | 1.33719940329E-05 | 0.827716391619 | 0.049 |
| rs7100626 | 10 | 96069674 | DICE | T CD4 TH1 | PLCE1 | A | 1.33719940329E-05 | 0.827716391619 | 0.049 |
| rs11187852 | 10 | 96070132 | DICE | T CD4 TH1 | PLCE1 | A | 1.33719940329E-05 | 0.827716391619 | 0.049 |
| rs752140 | 10 | 96071396 | DICE | T CD4 TH1 | PLCE1 | T | 1.33719940329E-05 | 0.827716391619 | 0.049 |
| rs11187864 | 10 | 96082506 | DICE | T CD4 TH1 17 | PLCE1 | T | 1.35358306881E-05 | 0.825812915876 | 0.049 |
| rs11187870 | 10 | 96087866 | DICE | T CD4 TH1 17 | PLCE1 | C | 1.35358306881E-05 | 0.825812915876 | 0.049 |
| rs145707916 | 10 | 96092992 | DICE | T CD4 TH1 17 | PLCE1 | GA | 1.35358306881E-05 | 0.825812915876 | 0.049 |
| rs750148755 | 10 | 96095271 | DICE | T CD4 TH1 17 | PLCE1 | C | 1.35358306881E-05 | 0.825812915876 | 0.049 |
| rs11187882 | 10 | 96095861 | DICE | T CD4 TH1 17 | PLCE1 | C | 1.35358306881E-05 | 0.825812915876 | 0.049 |
| rs11187895 | 10 | 96106240 | DICE | T CD4 TH1 17 | PLCE1 | C | 1.35358306881E-05 | 0.825812915876 | 0.049 |
| rs11187897 | 10 | 96106603 | DICE | T CD4 TH1 17 | PLCE1 | G | 1.35358306881E-05 | 0.825812915876 | 0.049 |
| rs7080472 | 10 | 96012950 | GTEx/v8 | Skin Sun Exposed Lower leg | RP11-76P2.4 | T | 1.35417E-05 | 0.210063 | 1.123E-06 |
| rs3740360 | 10 | 96025491 | GTEx/v8 | Adipose Visceral Omentum | PLCE1-AS1 | C | 1.36375E-05 | -0.307083 | 0.000644523 |
| rs17109875 | 10 | 96026575 | GTEx/v8 | Adipose Visceral Omentum | PLCE1-AS1 | C | 1.36375E-05 | -0.307083 | 0.000644523 |
| rs11187837 | 10 | 96035980 | GTEx/v8 | Adipose Visceral Omentum | PLCE1-AS1 | C | 1.36375E-05 | -0.307083 | 0.000644523 |
| rs3891783 | 10 | 96015793 | GTEx/v8 | Artery Tibial | PLCE1-AS1 | G | 1.3673E-05 | 0.133394 | 2.47435E-06 |
| rs57866767 | 10 | 96023077 | GTEx/v8 | Artery Tibial | PLCE1-AS1 | C | 1.3673E-05 | 0.133394 | 2.47435E-06 |
| rs11187845 | 10 | 96060198 | DICE | T CD4 TH1 | PLCE1 | A | 1.38155069706E-05 | 0.838348523246 | 0.049 |
| rs34309017 | 10 | 96061619 | DICE | T CD4 TH1 | PLCE1 | A | 1.38155069706E-05 | 0.838348523246 | 0.049 |
| rs12217792 | 10 | 96062386 | DICE | T CD4 TH1 | PLCE1 | C | 1.38155069706E-05 | 0.838348523246 | 0.049 |
| rs11187847 | 10 | 96063440 | DICE | T CD4 TH1 | PLCE1 | G | 1.38155069706E-05 | 0.838348523246 | 0.049 |
| rs75409190 | 10 | 96072425 | DICE | T CD4 TH1 | PLCE1 | T | 1.38155069706E-05 | 0.838348523246 | 0.049 |
| rs771106747 | 10 | 96072873 | DICE | T CD4 TH1 | PLCE1 | C | 1.38155069706E-05 | 0.838348523246 | 0.049 |
| rs749295068 | 10 | 96073508 | DICE | T CD4 TH1 | PLCE1 | A | 1.38155069706E-05 | 0.838348523246 | 0.049 |
| rs11187856 | 10 | 96076869 | DICE | T CD4 TH1 | PLCE1 | A | 1.38155069706E-05 | 0.838348523246 | 0.049 |
| rs11187837 | 10 | 96035980 | eQTLGen | eQTLGen cis eQTLs | NOC3L | C | 1.4029E-05 | 5,6111E+05 | 0.0391420118343195 |
| rs11187838 | 10 | 96038686 | GTEx/v8 | Thyroid | TBC1D12 | A | 1.41454E-05 | -0.12905 | 1.4449E-10 |
| rs10786156 | 10 | 96014622 | GTEx/v8 | Brain Hippocampus | NOC3L | G | 1.45003E-05 | 0.237209 | 0.000170124 |
| rs3891783 | 10 | 96015793 | GTEx/v8 | Brain Hippocampus | NOC3L | G | 1.45003E-05 | 0.237209 | 0.000170124 |
| rs57866767 | 10 | 96023077 | GTEx/v8 | Brain Hippocampus | NOC3L | C | 1.45003E-05 | 0.237209 | 0.000170124 |
| rs34309017 | 10 | 96061619 | GTEx/v8 | Whole Blood | TBC1D12 | A | 1.51594E-05 | 0.2357 | 0.00686763 |
| rs12217792 | 10 | 96062386 | GTEx/v8 | Whole Blood | TBC1D12 | C | 1.51594E-05 | 0.2357 | 0.00686763 |
| rs11187838 | 10 | 96038686 | GTEx/v8 | Heart Atrial Appendage | NOC3L | A | 1.52536E-05 | 0.162842 | 2.17353E-13 |
| rs75409190 | 10 | 96072425 | GTEx/v8 | Whole Blood | TBC1D12 | T | 1.57792E-05 | 0.240261 | 0.00686763 |
| rs771106747 | 10 | 96072873 | GTEx/v8 | Whole Blood | TBC1D12 | C | 1.57792E-05 | 0.240261 | 0.00686763 |
| rs11187856 | 10 | 96076869 | GTEx/v8 | Whole Blood | TBC1D12 | A | 1.57792E-05 | 0.240261 | 0.00686763 |
| rs2274224 | 10 | 96039597 | GTEx/v8 | Skin Sun Exposed Lower leg | RP11-76P2.4 | C | 1.60059E-05 | 0.205582 | 1.123E-06 |
| rs10786156 | 10 | 96014622 | GTEx/v8 | Artery Tibial | PLCE1-AS1 | G | 1.67528E-05 | 0.132008 | 2.47435E-06 |
| rs10786156 | 10 | 96014622 | GTEx/v8 | Heart Atrial Appendage | NOC3L | G | 1.7044E-05 | 0.160855 | 2.17353E-13 |
| rs3891783 | 10 | 96015793 | GTEx/v8 | Heart Atrial Appendage | NOC3L | G | 1.7044E-05 | 0.160855 | 2.17353E-13 |

| | | | | | | | | | |
|---|---|---|---|---|---|---|---|---|---|
| rs57866767 | 10 | 96023077 | GTEx/v8 | Heart Atrial Appendage | NOC3L | C | 1.7044E-05 | 0.160855 | 2.17353E-13 |
| rs778275808 | 10 | 96114212 | DICE | T CD4 TH1 17 | PLCE1 | T | 1.74378734524E-05 | 0.871571813104 | 0.049 |
| rs10509672 | 10 | 96115957 | DICE | T CD4 TH1 17 | PLCE1 | C | 1.74378734524E-05 | 0.871571813104 | 0.049 |
| rs74825688 | 10 | 96119687 | DICE | T CD4 TH1 17 | PLCE1 | A | 1.74378734524E-05 | 0.871571813104 | 0.049 |
| rs79193572 | 10 | 96119713 | DICE | T CD4 TH1 17 | PLCE1 | G | 1.74378734524E-05 | 0.871571813104 | 0.049 |
| rs11187906 | 10 | 96124838 | DICE | T CD4 TH1 17 | PLCE1 | A | 1.74378734524E-05 | 0.871571813104 | 0.049 |
| rs11187907 | 10 | 96125361 | DICE | T CD4 TH1 17 | PLCE1 | T | 1.74378734524E-05 | 0.871571813104 | 0.049 |
| rs113544984 | 10 | 96127448 | DICE | T CD4 TH1 17 | PLCE1 | A | 1.74378734524E-05 | 0.871571813104 | 0.049 |
| rs11187838 | 10 | 96038686 | GTEx/v8 | Adipose Subcutaneo us | NOC3L | A | 1.74632E-05 | 0.109597 | 3.06299E-11 |
| rs11187877 | 10 | 96092121 | GTEx/v8 | Thyroid | NOC3L | A | 1.74843E-05 | -0.162551 | 0.00220997 |
| rs7080472 | 10 | 96012950 | GTEx/v8 | Spleen | NOC3L | T | 1.75545E-05 | 0.224477 | 1.49159E-15 |
| rs11187882 | 10 | 96095861 | GTEx/v8 | Whole Blood | TBC1D12 | C | 1.77453E-05 | 0.239411 | 0.00686763 |
| rs11187844 | 10 | 96056629 | eQTLGen | eQTLGen cis eQTLs | PDLIM1 | A | 1.7948E-05 | -4.2889 | 0.0491932634554234 |
| rs7100626 | 10 | 96069674 | GTEx/v8 | Thyroid | NOC3L | A | 1.81872E-05 | -0.162883 | 0.00220997 |
| rs11187852 | 10 | 96070132 | GTEx/v8 | Thyroid | NOC3L | A | 1.81872E-05 | -0.162883 | 0.00220997 |
| rs753724 | 10 | 96051417 | GTEx/v8 | Muscle Skeletal | NOC3L | T | 1.82552E-05 | 0.20526 | 7.35621E-14 |
| rs11187842 | 10 | 96052511 | GTEx/v8 | Muscle Skeletal | NOC3L | T | 1.82552E-05 | 0.20526 | 7.35621E-14 |
| rs3781266 | 10 | 96052747 | GTEx/v8 | Muscle Skeletal | NOC3L | G | 1.82552E-05 | 0.20526 | 7.35621E-14 |
| rs3740365 | 10 | 96053239 | GTEx/v8 | Muscle Skeletal | NOC3L | A | 1.82552E-05 | 0.20526 | 7.35621E-14 |
| rs12220091 | 10 | 96053689 | GTEx/v8 | Muscle Skeletal | NOC3L | T | 1.82552E-05 | 0.20526 | 7.35621E-14 |
| rs3891783 | 10 | 96015793 | GTEx/v8 | Skin Sun Exposed Lower leg | RP11-76P2.4 | G | 1.96428E-05 | 0.204197 | 1.123E-06 |
| rs57866767 | 10 | 96023077 | GTEx/v8 | Skin Sun Exposed Lower leg | RP11-76P2.4 | C | 1.96428E-05 | 0.204197 | 1.123E-06 |
| rs7080472 | 10 | 96012950 | GTEx/v8 | Prostate | NOC3L | T | 1.9685E-05 | 0.188107 | 7.10224E-08 |
| rs11187838 | 10 | 96038686 | GTEx/v8 | Artery Tibial | PLCE1 | A | 2.06124E-05 | 0.106027 | 1.21915E-07 |
| rs2274224 | 10 | 96039597 | GTEx/v8 | Skin Not Sun Exposed Suprapubic | RP11-76P2.4 | C | 2.0648E-05 | 0.214 | 5.98468E-07 |
| rs11187845 | 10 | 96060198 | GTEx/v8 | Whole Blood | TBC1D12 | A | 2.10554E-05 | 0.225987 | 0.00686763 |
| rs11187847 | 10 | 96063440 | GTEx/v8 | Whole Blood | TBC1D12 | G | 2.10554E-05 | 0.225987 | 0.00686763 |
| rs10786156 | 10 | 96014622 | GTEx/v8 | Adipose Subcutaneo us | NOC3L | G | 2.14308E-05 | 0.107454 | 3.06299E-11 |
| rs11187895 | 10 | 96106240 | GTEx/v8 | Thyroid | NOC3L | C | 2.14366E-05 | -0.162118 | 0.00220997 |
| rs11187897 | 10 | 96106603 | GTEx/v8 | Thyroid | NOC3L | G | 2.14366E-05 | -0.162118 | 0.00220997 |
| rs3891783 | 10 | 96015793 | GTEx/v8 | Adipose Subcutaneo us | NOC3L | G | 2.14918E-05 | 0.107701 | 3.06299E-11 |
| rs57866767 | 10 | 96023077 | GTEx/v8 | Adipose Subcutaneo us | NOC3L | C | 2.14918E-05 | 0.107701 | 3.06299E-11 |
| rs10786156 | 10 | 96014622 | GTEx/v8 | Skin Sun Exposed Lower leg | RP11-76P2.4 | G | 2.15253E-05 | 0.203092 | 1.123E-06 |
| rs2274224 | 10 | 96039597 | GTEx/v8 | Artery Tibial | PLCE1-AS1 | C | 2.18917E-05 | 0.13008 | 2.47435E-06 |
| rs7096883 | 10 | 96069149 | GTEx/v8 | Thyroid | NOC3L | A | 2.23774E-05 | -0.157473 | 0.00220997 |
| rs11187840 | 10 | 96050351 | GTEx/v8 | Muscle Skeletal | NOC3L | G | 2.264E-05 | 0.204891 | 7.35621E-14 |
| rs75017201 | 10 | 96055152 | GTEx/v8 | Muscle Skeletal | NOC3L | T | 2.264E-05 | 0.204891 | 7.35621E-14 |
| rs7080472 | 10 | 96012950 | GTEx/v8 | Nerve Tibial | PLCE1 | T | 2.35053E-05 | 0.140728 | 1.91043E-25 |
| rs7080472 | 10 | 96012950 | GTEx/v8 | Colon Transverse | RP11-76P2.4 | T | 2.41297E-05 | 0.26705 | 2.2326E-06 |
| rs7080472 | 10 | 96012950 | GTEx/v8 | Skin Not Sun Exposed Suprapubic | RP11-76P2.4 | T | 2.42629E-05 | 0.214514 | 5.98468E-07 |

| | | | | | | | | | |
|---|---|---|---|---|---|---|---|---|---|
| rs11187877 | 10 | 96092121 | GTEx/v8 | Whole Blood | TBC1D12 | A | 2.46351E-05 | 0.232016 | 0.00686763 |
| rs11187838 | 10 | 96038686 | GTEx/v8 | Muscle Skeletal | NOC3L | A | 2.53713E-05 | 0.114667 | 7.35621E-14 |
| rs11187907 | 10 | 96125361 | BIOSQTL | BIOS eQTL geneLevel | HELLS | T | 2.59241E-05 | 4.2066091 | 0.0160794899241402 |
| rs11187907 | 10 | 96125361 | BIOSQTL | BIOS eQTL geneLevel | RP11-119K6.6 | T | 2.59241E-05 | 4.2066091 | 0.0160794899241402 |
| rs113544984 | 10 | 96127448 | BIOSQTL | BIOS eQTL geneLevel | HELLS | A | 2.67387E-05 | 4.1997961 | 0.0165349572331722 |
| rs113544984 | 10 | 96127448 | BIOSQTL | BIOS eQTL geneLevel | RP11-119K6.6 | A | 2.67387E-05 | 4.1997961 | 0.0165349572331722 |
| rs10786156 | 10 | 96014622 | GTEx/v8 | Brain Cerebellar Hemisphere | PLCE1 | G | 2.68601E-05 | 0.243306 | 0.0391634 |
| rs3891783 | 10 | 96015793 | GTEx/v8 | Brain Cerebellar Hemisphere | PLCE1 | G | 2.68601E-05 | 0.243306 | 0.0391634 |
| rs57866767 | 10 | 96023077 | GTEx/v8 | Brain Cerebellar Hemisphere | PLCE1 | C | 2.68601E-05 | 0.243306 | 0.0391634 |
| rs11187838 | 10 | 96038686 | GTEx/v8 | Skin Not Sun Exposed Suprapubic | PLCE1 | A | 2.70779E-05 | 0.131307 | 9.09957E-06 |
| rs10786156 | 10 | 96014622 | GTEx/v8 | Skin Not Sun Exposed Suprapubic | PLCE1 | G | 2.85818E-05 | 0.129644 | 9.09957E-06 |
| rs3891783 | 10 | 96015793 | GTEx/v8 | Skin Not Sun Exposed Suprapubic | PLCE1 | G | 2.85818E-05 | 0.129644 | 9.09957E-06 |
| rs57866767 | 10 | 96023077 | GTEx/v8 | Skin Not Sun Exposed Suprapubic | PLCE1 | C | 2.85818E-05 | 0.129644 | 9.09957E-06 |
| rs7080472 | 10 | 96012950 | GTEx/v8 | Muscle Skeletal | NOC3L | T | 2.88251E-05 | 0.114279 | 7.35621E-14 |
| rs11187864 | 10 | 96082506 | GTEx/v8 | Whole Blood | TBC1D12 | T | 2.9558E-05 | 0.230494 | 0.00686763 |
| rs145707916 | 10 | 96092992 | GTEx/v8 | Whole Blood | TBC1D12 | GA | 2.9558E-05 | 0.230494 | 0.00686763 |
| rs750148755 | 10 | 96095271 | GTEx/v8 | Whole Blood | TBC1D12 | C | 2.9558E-05 | 0.230494 | 0.00686763 |
| rs11187838 | 10 | 96038686 | GTEx/v8 | Brain Cerebellar Hemisphere | PLCE1 | A | 3.05367E-05 | 0.241508 | 0.0391634 |
| rs2274224 | 10 | 96039597 | GTEx/v8 | Brain Cerebellar Hemisphere | PLCE1 | C | 3.05367E-05 | 0.241508 | 0.0391634 |
| rs2274224 | 10 | 96039597 | GTEx/v8 | Heart Atrial Appendage | NOC3L | C | 3.21677E-05 | 0.156215 | 2.17353E-13 |
| rs2274224 | 10 | 96039597 | GTEx/v8 | Muscle Skeletal | NOC3L | C | 3.2592E-05 | 0.112813 | 7.35621E-14 |
| rs11187870 | 10 | 96087866 | GTEx/v8 | Thyroid | NOC3L | C | 3.43653E-05 | -0.158631 | 0.00220997 |
| rs11187864 | 10 | 96082506 | GTEx/v8 | Thyroid | NOC3L | T | 3.55691E-05 | -0.157449 | 0.00220997 |
| rs145707916 | 10 | 96092992 | GTEx/v8 | Thyroid | NOC3L | GA | 3.55691E-05 | -0.157449 | 0.00220997 |
| rs750148755 | 10 | 96095271 | GTEx/v8 | Thyroid | NOC3L | C | 3.55691E-05 | -0.157449 | 0.00220997 |
| rs11187882 | 10 | 96095861 | GTEx/v8 | Thyroid | NOC3L | C | 3.55691E-05 | -0.157449 | 0.00220997 |
| rs2077218 | 10 | 96071561 | BIOSQTL | BIOS eQTL geneLevel | NOC3L | G | 3.60874E-05 | -4.13111 | 0.0217668071075289 |
| rs3891783 | 10 | 96015793 | GTEx/v8 | Muscle Skeletal | NOC3L | G | 3.63252E-05 | 0.112317 | 7.35621E-14 |
| rs57866767 | 10 | 96023077 | GTEx/v8 | Muscle Skeletal | NOC3L | C | 3.63252E-05 | 0.112317 | 7.35621E-14 |
| rs75409190 | 10 | 96072425 | GTEx/v8 | Thyroid | NOC3L | T | 3.71677E-05 | -0.157685 | 0.00220997 |
| rs771106747 | 10 | 96072873 | GTEx/v8 | Thyroid | NOC3L | C | 3.71677E-05 | -0.157685 | 0.00220997 |
| rs749295068 | 10 | 96073508 | GTEx/v8 | Thyroid | NOC3L | A | 3.71677E-05 | -0.157685 | 0.00220997 |
| rs11187856 | 10 | 96076869 | GTEx/v8 | Thyroid | NOC3L | A | 3.71677E-05 | -0.157685 | 0.00220997 |
| rs3740359 | 10 | 96100119 | GTEx/v8 | Whole Blood | TBC1D12 | T | 3.72385E-05 | 0.207278 | 0.00686763 |
| rs10786156 | 10 | 96014622 | GTEx/v8 | Muscle Skeletal | NOC3L | G | 3.72999E-05 | 0.112118 | 7.35621E-14 |
| rs7080472 | 10 | 96012950 | GTEx/v8 | Brain Hippocampus | NOC3L | T | 3.88052E-05 | 0.225972 | 0.000170124 |
| 10:96112261_AAAAAAAC_A | 10 | 96112261 | DICE | T CD4 TH1 17 | PLCE1 | A | 4.05134739452E-05 | 0.825353456446 | 0.049 |
| rs10786156 | 10 | 96014622 | GTEx/v8 | Nerve Tibial | PLCE1 | G | 4.08489E-05 | 0.136054 | 1.91043E-25 |
| rs769015184 | 10 | 96049708 | DICE | T CD4 TH1 | PLCE1 | A | 4.16589029341E-05 | 0.760076218492 | 0.049 |

| | | | | | | | | | |
|---|---|---|---|---|---|---|---|---|---|
| rs11187840 | 10 | 96050351 | DICE | T CD4 TH1 | PLCE1 | G | 4.16589029341E-05 | 0.760076218492 | 0.049 |
| rs753724 | 10 | 96051417 | DICE | T CD4 TH1 | PLCE1 | T | 4.16589029341E-05 | 0.760076218492 | 0.049 |
| rs11187842 | 10 | 96052511 | DICE | T CD4 TH1 | PLCE1 | T | 4.16589029341E-05 | 0.760076218492 | 0.049 |
| rs3781266 | 10 | 96052747 | DICE | T CD4 TH1 | PLCE1 | G | 4.16589029341E-05 | 0.760076218492 | 0.049 |
| rs3740365 | 10 | 96053239 | DICE | T CD4 TH1 | PLCE1 | A | 4.16589029341E-05 | 0.760076218492 | 0.049 |
| rs12220091 | 10 | 96053689 | DICE | T CD4 TH1 | PLCE1 | T | 4.16589029341E-05 | 0.760076218492 | 0.049 |
| rs2274224 | 10 | 96039597 | GTEx/v8 | Skin Not Sun Exposed Suprapubic | PLCE1 | C | 4.22949E-05 | 0.1268 | 9.09957E-06 |
| rs7080472 | 10 | 96012950 | GTEx/v8 | Spleen | RP11-76P2.4 | T | 4.24579E-05 | 0.346219 | 9.37816E-08 |
| rs7080472 | 10 | 96012950 | GTEx/v8 | Brain Cerebellar Hemisphere | PLCE1 | T | 4.31362E-05 | 0.242741 | 0.0391634 |
| rs7078844 | 10 | 96081140 | GTEx/v8 | Brain Cerebellum | HELLS | C | 4.34404E-05 | -0.28832 | 1.07902E-06 |
| rs11187907 | 10 | 96125361 | BIOSQTL | BIOS eQTL geneLevel | NOC3L | T | 4.38874E-05 | 4.0861408 | 0.0259650235592606 |
| rs113544984 | 10 | 96127448 | BIOSQTL | BIOS eQTL geneLevel | NOC3L | A | 4.38874E-05 | 4.0861408 | 0.0259650235592606 |
| rs3740360 | 10 | 96025491 | DICE | T CD4 TH1 | PLCE1 | C | 4.42723585059E-05 | 0.755815010248 | 0.049 |
| rs17109875 | 10 | 96026575 | DICE | T CD4 TH1 | PLCE1 | C | 4.42723585059E-05 | 0.755815010248 | 0.049 |
| rs11187837 | 10 | 96035980 | DICE | T CD4 TH1 | PLCE1 | C | 4.42723585059E-05 | 0.755815010248 | 0.049 |
| rs34309017 | 10 | 96061619 | GTEx/v8 | Thyroid | NOC3L | A | 4.48906E-05 | -0.152426 | 0.00220997 |
| rs12217792 | 10 | 96062386 | GTEx/v8 | Thyroid | NOC3L | C | 4.48906E-05 | -0.152426 | 0.00220997 |
| rs11187826 | 10 | 95988042 | PsychENCODE | PsychENCODE eQTLs | NOC3L | G | 4.65866E-05 | 0.0668799 | 0.0043080936142294 |
| rs75017201 | 10 | 96055152 | DICE | T CD4 TH1 | PLCE1 | T | 4.7192585625E-05 | 0.749469540464 | 0.049 |
| rs11187840 | 10 | 96050351 | GTEx/v8 | Whole Blood | TBC1D12 | G | 4.78334E-05 | 0.218164 | 0.00686763 |
| rs75017201 | 10 | 96055152 | GTEx/v8 | Whole Blood | TBC1D12 | T | 4.78334E-05 | 0.218164 | 0.00686763 |
| rs11187863 | 10 | 96081457 | DICE | T CD4 TH1 17 | PLCE1 | G | 4.90263652512E-05 | 0.764200029398 | 0.049 |
| rs11187866 | 10 | 96085991 | DICE | T CD4 TH1 17 | PLCE1 | G | 4.90263652512E-05 | 0.764200029398 | 0.049 |
| rs11187877 | 10 | 96092121 | DICE | T CD4 TH1 17 | PLCE1 | A | 4.90263652512E-05 | 0.764200029398 | 0.049 |
| rs11187883 | 10 | 96096866 | DICE | T CD4 TH1 17 | PLCE1 | C | 4.90263652512E-05 | 0.764200029398 | 0.049 |
| rs11187890 | 10 | 96102758 | DICE | T CD4 TH1 17 | PLCE1 | C | 4.90263652512E-05 | 0.764200029398 | 0.049 |
| rs3891783 | 10 | 96015793 | GTEx/v8 | Nerve Tibial | PLCE1 | G | 5.14442E-05 | 0.134396 | 1.91043E-25 |
| rs57866767 | 10 | 96023077 | GTEx/v8 | Nerve Tibial | PLCE1 | C | 5.14442E-05 | 0.134396 | 1.91043E-25 |
| rs778275808 | 10 | 96114212 | DICE | T CD4 naive TREG | PLCE1 | T | 5.26981619237E-05 | 0.803940206822 | 0.049 |
| rs10509672 | 10 | 96115957 | DICE | T CD4 naive TREG | PLCE1 | C | 5.26981619237E-05 | 0.803940206822 | 0.049 |
| rs74825688 | 10 | 96119687 | DICE | T CD4 naive TREG | PLCE1 | A | 5.26981619237E-05 | 0.803940206822 | 0.049 |
| rs79193572 | 10 | 96119713 | DICE | T CD4 naive TREG | PLCE1 | G | 5.26981619237E-05 | 0.803940206822 | 0.049 |
| rs11187906 | 10 | 96124838 | DICE | T CD4 naive TREG | PLCE1 | A | 5.26981619237E-05 | 0.803940206822 | 0.049 |
| rs11187907 | 10 | 96125361 | DICE | T CD4 naive TREG | PLCE1 | T | 5.26981619237E-05 | 0.803940206822 | 0.049 |
| rs113544984 | 10 | 96127448 | DICE | T CD4 naive TREG | PLCE1 | A | 5.26981619237E-05 | 0.803940206822 | 0.049 |
| rs3891783 | 10 | 96015793 | GTEx/v8 | Nerve Tibial | RP11-76P2.4 | G | 5.28035E-05 | 0.188273 | 5.10248E-05 |
| rs57866767 | 10 | 96023077 | GTEx/v8 | Nerve Tibial | RP11-76P2.4 | C | 5.28035E-05 | 0.188273 | 5.10248E-05 |
| rs7101207 | 10 | 96122857 | BIOSQTL | BIOS eQTL geneLevel | NOC3L | A | 5.52101E-05 | 4.0325847 | 0.0318882681564246 |
| rs778275808 | 10 | 96114212 | GTEx/v8 | Colon Transverse | CYP2C19 | T | 5.57377E-05 | 0.212138 | 0.023149 |
| rs10509672 | 10 | 96115957 | GTEx/v8 | Colon Transverse | CYP2C19 | C | 5.57377E-05 | 0.212138 | 0.023149 |
| rs74825688 | 10 | 96119687 | GTEx/v8 | Colon Transverse | CYP2C19 | A | 5.57377E-05 | 0.212138 | 0.023149 |
| rs79193572 | 10 | 96119713 | GTEx/v8 | Colon Transverse | CYP2C19 | G | 5.57377E-05 | 0.212138 | 0.023149 |

| | | | | | | | | | |
|---|---|---|---|---|---|---|---|---|---|
| rs113544984 | 10 | 96127448 | GTEx/v8 | Colon Transverse | CYP2C19 | A | 5.57377E-05 | 0.212138 | 0.023149 |
| rs79746649 | 10 | 96120218 | GTEx/v8 | Colon Transverse | CYP2C19 | T | 5.59505E-05 | 0.209954 | 0.023149 |
| rs7101207 | 10 | 96122857 | GTEx/v8 | Colon Transverse | CYP2C19 | A | 5.59505E-05 | 0.209954 | 0.023149 |
| rs112252876 | 10 | 96123355 | GTEx/v8 | Colon Transverse | CYP2C19 | T | 5.59505E-05 | 0.209954 | 0.023149 |
| rs3740359 | 10 | 96100119 | GTEx/v8 | Thyroid | NOC3L | T | 5.65973E-05 | -0.141322 | 0.00220997 |
| rs11187845 | 10 | 96060198 | DICE | T CD4 TH1 17 | PLCE1 | A | 5.72412574339E-05 | 0.773056370538 | 0.049 |
| rs34309017 | 10 | 96061619 | DICE | T CD4 TH1 17 | PLCE1 | A | 5.72412574339E-05 | 0.773056370538 | 0.049 |
| rs12217792 | 10 | 96062386 | DICE | T CD4 TH1 17 | PLCE1 | C | 5.72412574339E-05 | 0.773056370538 | 0.049 |
| rs11187847 | 10 | 96063440 | DICE | T CD4 TH1 17 | PLCE1 | G | 5.72412574339E-05 | 0.773056370538 | 0.049 |
| rs75409190 | 10 | 96072425 | DICE | T CD4 TH1 17 | PLCE1 | T | 5.72412574339E-05 | 0.773056370538 | 0.049 |
| rs771106747 | 10 | 96072873 | DICE | T CD4 TH1 17 | PLCE1 | C | 5.72412574339E-05 | 0.773056370538 | 0.049 |
| rs749295068 | 10 | 96073508 | DICE | T CD4 TH1 17 | PLCE1 | A | 5.72412574339E-05 | 0.773056370538 | 0.049 |
| rs11187856 | 10 | 96076869 | DICE | T CD4 TH1 17 | PLCE1 | A | 5.72412574339E-05 | 0.773056370538 | 0.049 |
| rs7078844 | 10 | 96081140 | GTEx/v8 | Brain Cerebellar Hemisphere | HELLS | C | 5.74834E-05 | -0.278659 | 8.15147E-08 |
| rs112252876 | 10 | 96123355 | BIOSQTL | BIOS eQTL geneLevel | NOC3L | T | 5.86444E-05 | 4.0183398 | 0.0336986057285402 |
| rs3740360 | 10 | 96025491 | GTEx/v8 | Thyroid | NOC3L | C | 5.9122E-05 | -0.136651 | 0.00220997 |
| rs17109875 | 10 | 96026575 | GTEx/v8 | Thyroid | NOC3L | C | 5.9122E-05 | -0.136651 | 0.00220997 |
| rs7101207 | 10 | 96122857 | GTEx/v8 | Adipose Visceral Omentum | NOC3L | A | 5.96319E-05 | 0.160985 | 0.000238073 |
| rs112252876 | 10 | 96123355 | GTEx/v8 | Adipose Visceral Omentum | NOC3L | T | 5.96319E-05 | 0.160985 | 0.000238073 |
| rs753724 | 10 | 96051417 | GTEx/v8 | Whole Blood | TBC1D12 | T | 6.22965E-05 | 0.209754 | 0.00686763 |
| rs11187842 | 10 | 96052511 | GTEx/v8 | Whole Blood | TBC1D12 | T | 6.22965E-05 | 0.209754 | 0.00686763 |
| rs3781266 | 10 | 96052747 | GTEx/v8 | Whole Blood | TBC1D12 | G | 6.22965E-05 | 0.209754 | 0.00686763 |
| rs3740365 | 10 | 96053239 | GTEx/v8 | Whole Blood | TBC1D12 | A | 6.22965E-05 | 0.209754 | 0.00686763 |
| rs12220091 | 10 | 96053689 | GTEx/v8 | Whole Blood | TBC1D12 | T | 6.22965E-05 | 0.209754 | 0.00686763 |
| rs11187845 | 10 | 96060198 | GTEx/v8 | Thyroid | NOC3L | A | 6.31024E-05 | -0.147662 | 0.00220997 |
| rs11187847 | 10 | 96063440 | GTEx/v8 | Thyroid | NOC3L | G | 6.31024E-05 | -0.147662 | 0.00220997 |
| rs11187838 | 10 | 96038686 | GTEx/v8 | Spleen | RP11-76P2.4 | A | 6.34948E-05 | 0.344413 | 9.37816E-08 |
| rs7080472 | 10 | 96012950 | GTEx/v8 | Artery Tibial | NOC3L | T | 6.38049E-05 | 0.099248 | 1.31505E-07 |
| rs11187890 | 10 | 96102758 | GTEx/v8 | Whole Blood | TBC1D12 | C | 6.51491E-05 | 0.222292 | 0.00686763 |
| rs79746649 | 10 | 96120218 | DICE | T CD4 TH1 17 | PLCE1 | T | 6.7940652152E-05 | 0.800331492037 | 0.049 |
| rs7101207 | 10 | 96122857 | DICE | T CD4 TH1 17 | PLCE1 | A | 6.7940652152E-05 | 0.800331492037 | 0.049 |
| rs112252876 | 10 | 96123355 | DICE | T CD4 TH1 17 | PLCE1 | T | 6.7940652152E-05 | 0.800331492037 | 0.049 |
| rs2274224 | 10 | 96039597 | GTEx/v8 | Artery Tibial | RP11-76P2.4 | C | 6.81555E-05 | 0.188221 | 5.41051E-05 |
| rs113544984 | 10 | 96127448 | GTEx/v8 | Adipose Visceral Omentum | NOC3L | A | 6.84112E-05 | 0.161311 | 0.000238073 |
| rs11187870 | 10 | 96087866 | GTEx/v8 | Whole Blood | TBC1D12 | C | 6.8925E-05 | 0.222498 | 0.00686763 |
| rs11187907 | 10 | 96125361 | GTEx/v8 | Colon Transverse | CYP2C19 | T | 7.07444E-05 | 0.203081 | 0.023149 |
| rs79746649 | 10 | 96120218 | GTEx/v8 | Colon Transverse | CYP2C9 | T | 7.09602E-05 | 0.229437 | 0.0239192 |
| rs7101207 | 10 | 96122857 | GTEx/v8 | Colon Transverse | CYP2C9 | A | 7.09602E-05 | 0.229437 | 0.0239192 |
| rs112252876 | 10 | 96123355 | GTEx/v8 | Colon Transverse | CYP2C9 | T | 7.09602E-05 | 0.229437 | 0.0239192 |
| rs11187870 | 10 | 96087866 | GTEx/v8 | Colon Transverse | CYP2C19 | C | 7.10595E-05 | 0.207839 | 0.023149 |
| rs2274224 | 10 | 96039597 | GTEx/v8 | Colon Transverse | RP11-76P2.4 | C | 7.1415E-05 | 0.251217 | 2.2326E-06 |
| rs2077218 | 10 | 96071561 | GTEx/v8 | Thyroid | PLCE1 | A | 7.35307E-05 | -0.140544 | 1.91428E-05 |
| rs10786156 | 10 | 96014622 | GTEx/v8 | Colon Transverse | RP11-76P2.4 | G | 7.37059E-05 | 0.251868 | 2.2326E-06 |
| rs3891783 | 10 | 96015793 | GTEx/v8 | Colon Transverse | RP11-76P2.4 | G | 7.37059E-05 | 0.251868 | 2.2326E-06 |

| | | | | | | | | | |
|---|---|---|---|---|---|---|---|---|---|
| rs57866767 | 10 | 96023077 | GTEx/v8 | Colon Transverse | RP11-76P2.4 | C | 7.37059E-05 | 0.251868 | 2.2326E-06 |
| rs11187838 | 10 | 96038686 | GTEx/v8 | Nerve Tibial | PLCE1 | A | 7.57587E-05 | 0.131345 | 1.91043E-25 |
| rs11187838 | 10 | 96038686 | GTEx/v8 | Skin Sun Exposed Lower leg | RP11-76P2.4 | A | 7.65039E-05 | 0.189406 | 1.123E-06 |
| rs11187895 | 10 | 96106240 | GTEx/v8 | Whole Blood | TBC1D12 | C | 7.77315E-05 | 0.220691 | 0.00686763 |
| rs11187897 | 10 | 96106603 | GTEx/v8 | Whole Blood | TBC1D12 | G | 7.77315E-05 | 0.220691 | 0.00686763 |
| rs3891783 | 10 | 96015793 | GTEx/v8 | Artery Tibial | RP11-76P2.4 | G | 7.82492E-05 | 0.18693 | 5.41051E-05 |
| rs57866767 | 10 | 96023077 | GTEx/v8 | Artery Tibial | RP11-76P2.4 | C | 7.82492E-05 | 0.18693 | 5.41051E-05 |
| rs2274224 | 10 | 96039597 | GTEx/v8 | Nerve Tibial | PLCE1 | C | 7.91228E-05 | 0.130521 | 1.91043E-25 |
| rs11187838 | 10 | 96038686 | GTEx/v8 | Whole Blood | NOC3L | A | 7.92195E-05 | 0.0850059 | 0.000592991 |
| rs79746649 | 10 | 96120218 | GTEx/v8 | Whole Blood | TBC1D12 | T | 8.02709E-05 | 0.220546 | 0.00686763 |
| rs11187838 | 10 | 96038686 | GTEx/v8 | Colon Transverse | RP11-76P2.4 | A | 8.22798E-05 | 0.247262 | 2.2326E-06 |
| rs7080472 | 10 | 96012950 | GTEx/v8 | Nerve Tibial | RP11-76P2.4 | T | 8.43931E-05 | 0.183804 | 5.10248E-05 |
| rs10786156 | 10 | 96014622 | GTEx/v8 | Artery Tibial | RP11-76P2.4 | G | 8.49316E-05 | 0.18596 | 5.41051E-05 |
| rs11187895 | 10 | 96106240 | BIOSQTL | BIOS eQTL geneLevel | NOC3L | C | 8.53708E-05 | 3.9289554 | 0.0465583340594232 |
| rs11187897 | 10 | 96106603 | BIOSQTL | BIOS eQTL geneLevel | NOC3L | G | 8.53708E-05 | 3.9289554 | 0.0465583340594232 |
| rs10509672 | 10 | 96115957 | BIOSQTL | BIOS eQTL geneLevel | NOC3L | C | 8.53708E-05 | 3.9289785 | 0.0465583340594232 |
| rs74825688 | 10 | 96119687 | BIOSQTL | BIOS eQTL geneLevel | NOC3L | A | 8.53708E-05 | 3.9289785 | 0.0465583340594232 |
| rs79193572 | 10 | 96119713 | BIOSQTL | BIOS eQTL geneLevel | NOC3L | G | 8.53708E-05 | 3.9289785 | 0.0465583340594232 |
| rs79746649 | 10 | 96120218 | BIOSQTL | BIOS eQTL geneLevel | NOC3L | T | 8.53708E-05 | 3.9289785 | 0.0465583340594232 |
| rs7078844 | 10 | 96081140 | GTEx/v8 | Cells Cultured fibroblasts | PLCE1-AS1 | C | 8.71297E-05 | -0.147185 | 3.86001E-30 |
| rs2274224 | 10 | 96039597 | GTEx/v8 | Whole Blood | NOC3L | C | 8.78308E-05 | 0.084549 | 0.000592991 |
| rs11187890 | 10 | 96102758 | BIOSQTL | BIOS eQTL geneLevel | NOC3L | C | 9.03293E-05 | 3.9152898 | 0.0491618170763485 |
| rs2077218 | 10 | 96071561 | GTEx/v8 | Thyroid | PLCE1-AS1 | A | 9.06102E-05 | -0.143826 | 0.00721889 |
| rs10786156 | 10 | 96014622 | GTEx/v8 | Whole Blood | NOC3L | G | 9.22693E-05 | 0.0845518 | 0.000592991 |
| rs778275808 | 10 | 96114212 | GTEx/v8 | Whole Blood | TBC1D12 | T | 9.58649E-05 | 0.218869 | 0.00686763 |
| rs10509672 | 10 | 96115957 | GTEx/v8 | Whole Blood | TBC1D12 | C | 9.58649E-05 | 0.218869 | 0.00686763 |
| rs11187895 | 10 | 96106240 | GTEx/v8 | Colon Transverse | CYP2C19 | C | 9.87863E-05 | 0.202956 | 0.023149 |
| rs11187897 | 10 | 96106603 | GTEx/v8 | Colon Transverse | CYP2C19 | G | 9.87863E-05 | 0.202956 | 0.023149 |
| rs11187890 | 10 | 96102758 | GTEx/v8 | Colon Transverse | CYP2C19 | C | 9.91947E-05 | 0.200806 | 0.023149 |
| rs11187840 | 10 | 96050351 | GTEx/v8 | Thyroid | NOC3L | G | 9.98719E-05 | -0.143413 | 0.00220997 |
| rs75017201 | 10 | 96055152 | GTEx/v8 | Thyroid | NOC3L | T | 9.98719E-05 | -0.143413 | 0.00220997 |
| rs74825688 | 10 | 96119687 | GTEx/v8 | Whole Blood | TBC1D12 | A | 0.000101837 | 0.217022 | 0.00686763 |
| rs10786156 | 10 | 96014622 | GTEx/v8 | Nerve Tibial | RP11-76P2.4 | G | 0.00010197 | 0.180963 | 5.10248E-05 |
| rs3740360 | 10 | 96025491 | GTEx/v8 | Whole Blood | TBC1D12 | C | 0.000102034 | 0.192773 | 0.00686763 |
| rs17109875 | 10 | 96026575 | GTEx/v8 | Whole Blood | TBC1D12 | C | 0.000102034 | 0.192773 | 0.00686763 |
| rs11187837 | 10 | 96035980 | GTEx/v8 | Whole Blood | TBC1D12 | C | 0.000102034 | 0.192773 | 0.00686763 |
| rs7080472 | 10 | 96012950 | GTEx/v8 | Artery Tibial | RP11-76P2.4 | T | 0.000105243 | 0.184919 | 5.41051E-05 |
| rs79193572 | 10 | 96119713 | GTEx/v8 | Whole Blood | TBC1D12 | G | 0.000110383 | 0.218162 | 0.00686763 |
| rs11187907 | 10 | 96125361 | GTEx/v8 | Colon Transverse | CYP2C9 | T | 0.000117747 | 0.218194 | 0.0239192 |
| rs11187837 | 10 | 96035980 | GTEx/v8 | Thyroid | NOC3L | C | 0.000120608 | -0.13038 | 0.00220997 |
| rs11187838 | 10 | 96038686 | GTEx/v8 | Artery Tibial | RP11-76P2.4 | A | 0.000121227 | 0.181833 | 5.41051E-05 |
| rs3891783 | 10 | 96015793 | GTEx/v8 | Whole Blood | NOC3L | G | 0.000124756 | 0.0828281 | 0.000592991 |
| rs57866767 | 10 | 96023077 | GTEx/v8 | Whole Blood | NOC3L | C | 0.000124756 | 0.0828281 | 0.000592991 |
| rs778275808 | 10 | 96114212 | GTEx/v8 | Colon Transverse | CYP2C9 | T | 0.000130747 | 0.223355 | 0.0239192 |
| rs10509672 | 10 | 96115957 | GTEx/v8 | Colon Transverse | CYP2C9 | C | 0.000130747 | 0.223355 | 0.0239192 |
| rs74825688 | 10 | 96119687 | GTEx/v8 | Colon Transverse | CYP2C9 | A | 0.000130747 | 0.223355 | 0.0239192 |

| | | | | | | | | | |
|---|---|---|---|---|---|---|---|---|---|
| rs79193572 | 10 | 96119713 | GTEx/v8 | Colon Transverse | CYP2C9 | G | 0.000130747 | 0.223355 | 0.0239192 |
| rs113544984 | 10 | 96127448 | GTEx/v8 | Colon Transverse | CYP2C9 | A | 0.000130747 | 0.223355 | 0.0239192 |
| rs11187838 | 10 | 96038686 | GTEx/v8 | Colon Transverse | CYP2C9 | A | 0.000131607 | 0.121358 | 0.0239192 |
| rs7080472 | 10 | 96012950 | GTEx/v8 | Whole Blood | NOC3L | T | 0.000132449 | 0.0827353 | 0.000592991 |
| rs11187864 | 10 | 96082506 | GTEx/v8 | Colon Transverse | CYP2C19 | T | 0.000136789 | 0.197872 | 0.023149 |
| rs145707916 | 10 | 96092992 | GTEx/v8 | Colon Transverse | CYP2C19 | GA | 0.000136789 | 0.197872 | 0.023149 |
| rs750148755 | 10 | 96095271 | GTEx/v8 | Colon Transverse | CYP2C19 | C | 0.000136789 | 0.197872 | 0.023149 |
| rs11187882 | 10 | 96095861 | GTEx/v8 | Colon Transverse | CYP2C19 | C | 0.000136789 | 0.197872 | 0.023149 |
| rs11187877 | 10 | 96092121 | GTEx/v8 | Colon Transverse | CYP2C19 | A | 0.00013711 | 0.195787 | 0.023149 |
| rs7078844 | 10 | 96081140 | GTEx/v8 | Skin Not Sun Exposed Suprapubic | NOC3L | C | 0.000137394 | 0.132453 | 2.01564E-23 |
| rs753724 | 10 | 96051417 | GTEx/v8 | Thyroid | NOC3L | T | 0.000138111 | -0.138724 | 0.00220997 |
| rs11187842 | 10 | 96052511 | GTEx/v8 | Thyroid | NOC3L | T | 0.000138111 | -0.138724 | 0.00220997 |
| rs3781266 | 10 | 96052747 | GTEx/v8 | Thyroid | NOC3L | G | 0.000138111 | -0.138724 | 0.00220997 |
| rs3740365 | 10 | 96053239 | GTEx/v8 | Thyroid | NOC3L | A | 0.000138111 | -0.138724 | 0.00220997 |
| rs12220091 | 10 | 96053689 | GTEx/v8 | Thyroid | NOC3L | T | 0.000138111 | -0.138724 | 0.00220997 |
| rs11187838 | 10 | 96038686 | GTEx/v8 | Colon Transverse | MTND4P20 | A | 0.000139292 | 0.116351 | 0.0452717 |
| rs11187863 | 10 | 96081457 | GTEx/v8 | Colon Transverse | CYP2C9 | G | 0.000141772 | 0.203738 | 0.0239192 |
| rs11187866 | 10 | 96085991 | GTEx/v8 | Colon Transverse | CYP2C9 | G | 0.000141772 | 0.203738 | 0.0239192 |
| rs11187883 | 10 | 96096866 | GTEx/v8 | Colon Transverse | CYP2C9 | C | 0.000141772 | 0.203738 | 0.0239192 |
| rs11187845 | 10 | 96060198 | GTEx/v8 | Thyroid | PLCE1 | A | 0.000142332 | -0.183373 | 1.91428E-05 |
| rs11187847 | 10 | 96063440 | GTEx/v8 | Thyroid | PLCE1 | G | 0.000142332 | -0.183373 | 1.91428E-05 |
| rs2274224 | 10 | 96039597 | GTEx/v8 | Nerve Tibial | RP11-76P2.4 | C | 0.000156152 | 0.175538 | 5.10248E-05 |
| rs2274224 | 10 | 96039597 | GTEx/v8 | Colon Transverse | CYP2C9 | C | 0.000157209 | 0.1209 | 0.0239192 |
| rs752140 | 10 | 96071396 | GTEx/v8 | Colon Transverse | CYP2C9 | T | 0.00016468 | 0.202017 | 0.0239192 |
| rs11187838 | 10 | 96038686 | GTEx/v8 | Nerve Tibial | RP11-76P2.4 | A | 0.000182236 | 0.174418 | 5.10248E-05 |
| rs2274224 | 10 | 96039597 | GTEx/v8 | Muscle Skeletal | CYP2C8 | C | 0.00020384 | 0.15337 | 5.87392E-21 |
| rs753724 | 10 | 96051417 | GTEx/v8 | Thyroid | PLCE1 | T | 0.000252113 | -0.173953 | 1.91428E-05 |
| rs11187842 | 10 | 96052511 | GTEx/v8 | Thyroid | PLCE1 | T | 0.000252113 | -0.173953 | 1.91428E-05 |
| rs3781266 | 10 | 96052747 | GTEx/v8 | Thyroid | PLCE1 | G | 0.000252113 | -0.173953 | 1.91428E-05 |
| rs3740365 | 10 | 96053239 | GTEx/v8 | Thyroid | PLCE1 | A | 0.000252113 | -0.173953 | 1.91428E-05 |
| rs12220091 | 10 | 96053689 | GTEx/v8 | Thyroid | PLCE1 | T | 0.000252113 | -0.173953 | 1.91428E-05 |
| rs79746649 | 10 | 96120218 | GTEx/v8 | Thyroid | NOC3L | T | 0.000260305 | -0.141234 | 0.00220997 |
| rs10786156 | 10 | 96014622 | GTEx/v8 | Muscle Skeletal | CYP2C8 | G | 0.000285424 | 0.150035 | 5.87392E-21 |
| rs3891783 | 10 | 96015793 | GTEx/v8 | Muscle Skeletal | CYP2C8 | G | 0.000318235 | 0.148928 | 5.87392E-21 |
| rs57866767 | 10 | 96023077 | GTEx/v8 | Muscle Skeletal | CYP2C8 | C | 0.000318235 | 0.148928 | 5.87392E-21 |
| rs11187826 | 10 | 95988042 | CMC | CMC SVA cis | NOC3L | G | | 1.0 | 0.049 |
| rs7080472 | 10 | 96012950 | CMC | CMC SVA cis | NOC3L | T | | 1.0 | 0.009 |
| rs10786156 | 10 | 96014622 | CMC | CMC SVA cis | NOC3L | G | | 1.0 | 0.009 |
| rs3891783 | 10 | 96015793 | CMC | CMC SVA cis | NOC3L | G | | 1.0 | 0.009 |
| rs57866767 | 10 | 96023077 | CMC | CMC SVA cis | NOC3L | C | | 1.0 | 0.009 |
| rs3740360 | 10 | 96025491 | CMC | CMC SVA cis | NOC3L | C | | 1.0 | 0.049 |
| rs17109875 | 10 | 96026575 | CMC | CMC SVA cis | NOC3L | C | | 1.0 | 0.049 |
| rs11187837 | 10 | 96035980 | CMC | CMC SVA cis | NOC3L | C | | 1.0 | 0.049 |
| rs11187838 | 10 | 96038686 | CMC | CMC SVA cis | NOC3L | A | | 1.0 | 0.009 |
| rs2274224 | 10 | 96039597 | CMC | CMC SVA cis | NOC3L | C | | 1.0 | 0.009 |

| | | | | | | | | | |
|---|---|---|---|---|---|---|---|---|---|
| rs769015184 | 10 | 96049708 | CMC | CMC SVA cis | NOC3L | A | | 1.0 | 0.009 |
| rs11187840 | 10 | 96050351 | CMC | CMC SVA cis | NOC3L | G | | 1.0 | 0.009 |
| rs753724 | 10 | 96051417 | CMC | CMC SVA cis | NOC3L | T | | 1.0 | 0.009 |
| rs11187842 | 10 | 96052511 | CMC | CMC SVA cis | NOC3L | T | | 1.0 | 0.009 |
| rs3781266 | 10 | 96052747 | CMC | CMC SVA cis | NOC3L | G | | 1.0 | 0.009 |
| rs3740365 | 10 | 96053239 | CMC | CMC SVA cis | NOC3L | A | | 1.0 | 0.009 |
| rs12220091 | 10 | 96053689 | CMC | CMC SVA cis | NOC3L | T | | 1.0 | 0.009 |
| rs75017201 | 10 | 96055152 | CMC | CMC SVA cis | NOC3L | T | | 1.0 | 0.009 |
| rs200197176 | 10 | 96055963 | CMC | CMC SVA cis | NOC3L | CA | | 1.0 | 0.009 |
| rs11187844 | 10 | 96056629 | CMC | CMC SVA cis | NOC3L | A | | 1.0 | 0.009 |
| rs11187845 | 10 | 96060198 | CMC | CMC SVA cis | NOC3L | A | | 1.0 | 0.009 |
| rs34309017 | 10 | 96061619 | CMC | CMC SVA cis | NOC3L | A | | 1.0 | 0.009 |
| rs12217792 | 10 | 96062386 | CMC | CMC SVA cis | NOC3L | C | | 1.0 | 0.009 |
| rs11187847 | 10 | 96063440 | CMC | CMC SVA cis | NOC3L | G | | 1.0 | 0.009 |
| rs7096883 | 10 | 96069149 | CMC | CMC SVA cis | NOC3L | A | | 1.0 | 0.009 |
| rs7100626 | 10 | 96069674 | CMC | CMC SVA cis | NOC3L | A | | 1.0 | 0.009 |
| rs11187852 | 10 | 96070132 | CMC | CMC SVA cis | NOC3L | A | | 1.0 | 0.009 |
| rs752140 | 10 | 96071396 | CMC | CMC SVA cis | NOC3L | T | | 1.0 | 0.009 |
| rs75409190 | 10 | 96072425 | CMC | CMC SVA cis | NOC3L | T | | 1.0 | 0.009 |
| rs771106747 | 10 | 96072873 | CMC | CMC SVA cis | NOC3L | C | | 1.0 | 0.009 |
| rs749295068 | 10 | 96073508 | CMC | CMC SVA cis | NOC3L | A | | 1.0 | 0.009 |
| rs11187856 | 10 | 96076869 | CMC | CMC SVA cis | NOC3L | A | | 1.0 | 0.009 |
| rs7078844 | 10 | 96081140 | CMC | CMC SVA cis | HELLS | T | | 1.0 | 0.049 |
| rs11187863 | 10 | 96081457 | CMC | CMC SVA cis | NOC3L | G | | 1.0 | 0.009 |
| rs11187864 | 10 | 96082506 | CMC | CMC SVA cis | NOC3L | T | | 1.0 | 0.009 |
| rs11187866 | 10 | 96085991 | CMC | CMC SVA cis | NOC3L | G | | 1.0 | 0.009 |
| rs11187870 | 10 | 96087866 | CMC | CMC SVA cis | NOC3L | C | | 1.0 | 0.009 |
| rs11187877 | 10 | 96092121 | CMC | CMC SVA cis | NOC3L | A | | 1.0 | 0.009 |
| rs145707916 | 10 | 96092992 | CMC | CMC SVA cis | NOC3L | GA | | 1.0 | 0.009 |
| rs750148755 | 10 | 96095271 | CMC | CMC SVA cis | NOC3L | C | | 1.0 | 0.009 |
| rs11187882 | 10 | 96095861 | CMC | CMC SVA cis | NOC3L | C | | 1.0 | 0.009 |
| rs11187883 | 10 | 96096866 | CMC | CMC SVA cis | NOC3L | C | | 1.0 | 0.009 |
| rs3740359 | 10 | 96100119 | CMC | CMC SVA cis | NOC3L | T | | 1.0 | 0.009 |
| rs11187890 | 10 | 96102758 | CMC | CMC SVA cis | NOC3L | C | | 1.0 | 0.009 |
| rs11187895 | 10 | 96106240 | CMC | CMC SVA cis | NOC3L | C | | 1.0 | 0.009 |
| rs11187897 | 10 | 96106603 | CMC | CMC SVA cis | NOC3L | G | | 1.0 | 0.009 |
| 10:96112261_AAAAAAAC_A | 10 | 96112261 | CMC | CMC SVA cis | NOC3L | A | | 1.0 | 0.009 |
| rs778275808 | 10 | 96114212 | CMC | CMC SVA cis | NOC3L | T | | 1.0 | 0.009 |
| rs10509672 | 10 | 96115957 | CMC | CMC SVA cis | NOC3L | C | | 1.0 | 0.009 |
| rs74825688 | 10 | 96119687 | CMC | CMC SVA cis | NOC3L | A | | 1.0 | 0.009 |
| rs79193572 | 10 | 96119713 | CMC | CMC SVA cis | NOC3L | G | | 1.0 | 0.009 |
| rs79746649 | 10 | 96120218 | CMC | CMC SVA cis | NOC3L | T | | 1.0 | 0.009 |
| rs7101207 | 10 | 96122857 | CMC | CMC SVA cis | NOC3L | A | | 1.0 | 0.009 |

| | | | | | | | | | |
|---|---|---|---|---|---|---|---|---|---|
| rs112252876 | 10 | 96123355 | CMC | CMC SVA cis | NOC3L | T | | 1.0 | 0.009 |
| rs11187906 | 10 | 96124838 | CMC | CMC SVA cis | NOC3L | A | | 1.0 | 0.009 |
| rs11187907 | 10 | 96125361 | CMC | CMC SVA cis | NOC3L | T | | 1.0 | 0.009 |
| rs113544984 | 10 | 96127448 | CMC | CMC SVA cis | NOC3L | A | | 1.0 | 0.009 |
| rs7080472 | 10 | 96012950 | CMC | CMC NoSVA cis | PLCE1 | T | | 1.0 | 0.049 |
| rs7080472 | 10 | 96012950 | CMC | CMC NoSVA cis | NOC3L | T | | 1.0 | 0.049 |
| rs10786156 | 10 | 96014622 | CMC | CMC NoSVA cis | PLCE1 | G | | 1.0 | 0.049 |
| rs10786156 | 10 | 96014622 | CMC | CMC NoSVA cis | NOC3L | G | | 1.0 | 0.009 |
| rs3891783 | 10 | 96015793 | CMC | CMC NoSVA cis | PLCE1 | G | | 1.0 | 0.049 |
| rs3891783 | 10 | 96015793 | CMC | CMC NoSVA cis | NOC3L | G | | 1.0 | 0.009 |
| rs57866767 | 10 | 96023077 | CMC | CMC NoSVA cis | PLCE1 | C | | 1.0 | 0.049 |
| rs57866767 | 10 | 96023077 | CMC | CMC NoSVA cis | NOC3L | C | | 1.0 | 0.009 |
| rs11187838 | 10 | 96038686 | CMC | CMC NoSVA cis | PLCE1 | A | | 1.0 | 0.049 |
| rs11187838 | 10 | 96038686 | CMC | CMC NoSVA cis | NOC3L | A | | 1.0 | 0.009 |
| rs2274224 | 10 | 96039597 | CMC | CMC NoSVA cis | PLCE1 | C | | 1.0 | 0.049 |
| rs2274224 | 10 | 96039597 | CMC | CMC NoSVA cis | NOC3L | C | | 1.0 | 0.009 |
| rs769015184 | 10 | 96049708 | CMC | CMC NoSVA cis | NOC3L | A | | 1.0 | 0.049 |
| rs11187840 | 10 | 96050351 | CMC | CMC NoSVA cis | NOC3L | G | | 1.0 | 0.049 |
| rs753724 | 10 | 96051417 | CMC | CMC NoSVA cis | NOC3L | T | | 1.0 | 0.049 |
| rs11187842 | 10 | 96052511 | CMC | CMC NoSVA cis | NOC3L | T | | 1.0 | 0.049 |
| rs3781266 | 10 | 96052747 | CMC | CMC NoSVA cis | NOC3L | G | | 1.0 | 0.049 |
| rs3740365 | 10 | 96053239 | CMC | CMC NoSVA cis | NOC3L | A | | 1.0 | 0.049 |
| rs12220091 | 10 | 96053689 | CMC | CMC NoSVA cis | NOC3L | T | | 1.0 | 0.049 |
| rs75017201 | 10 | 96055152 | CMC | CMC NoSVA cis | NOC3L | T | | 1.0 | 0.049 |
| rs200197176 | 10 | 96055963 | CMC | CMC NoSVA cis | NOC3L | CA | | 1.0 | 0.009 |
| rs11187845 | 10 | 96060198 | CMC | CMC NoSVA cis | NOC3L | A | | 1.0 | 0.049 |
| rs34309017 | 10 | 96061619 | CMC | CMC NoSVA cis | NOC3L | A | | 1.0 | 0.049 |
| rs12217792 | 10 | 96062386 | CMC | CMC NoSVA cis | NOC3L | C | | 1.0 | 0.049 |
| rs11187847 | 10 | 96063440 | CMC | CMC NoSVA cis | NOC3L | G | | 1.0 | 0.049 |
| rs7096883 | 10 | 96069149 | CMC | CMC NoSVA cis | NOC3L | A | | 1.0 | 0.049 |
| rs7100626 | 10 | 96069674 | CMC | CMC NoSVA cis | NOC3L | A | | 1.0 | 0.049 |
| rs11187852 | 10 | 96070132 | CMC | CMC NoSVA cis | NOC3L | A | | 1.0 | 0.049 |
| rs752140 | 10 | 96071396 | CMC | CMC NoSVA cis | NOC3L | T | | 1.0 | 0.049 |
| rs75409190 | 10 | 96072425 | CMC | CMC NoSVA cis | NOC3L | T | | 1.0 | 0.049 |
| rs771106747 | 10 | 96072873 | CMC | CMC NoSVA cis | NOC3L | C | | 1.0 | 0.049 |
| rs749295068 | 10 | 96073508 | CMC | CMC NoSVA cis | NOC3L | A | | 1.0 | 0.049 |
| rs11187856 | 10 | 96076869 | CMC | CMC NoSVA cis | NOC3L | A | | 1.0 | 0.049 |
| rs11187863 | 10 | 96081457 | CMC | CMC NoSVA cis | NOC3L | G | | 1.0 | 0.049 |
| rs11187864 | 10 | 96082506 | CMC | CMC NoSVA cis | NOC3L | T | | 1.0 | 0.049 |
| rs11187866 | 10 | 96085991 | CMC | CMC NoSVA cis | NOC3L | G | | 1.0 | 0.049 |
| rs11187870 | 10 | 96087866 | CMC | CMC NoSVA cis | NOC3L | C | | 1.0 | 0.049 |
| rs11187877 | 10 | 96092121 | CMC | CMC NoSVA cis | NOC3L | A | | 1.0 | 0.049 |
| rs145707916 | 10 | 96092992 | CMC | CMC NoSVA cis | NOC3L | GA | | 1.0 | 0.049 |

| | | | | | | | | | |
|---|---|---|---|---|---|---|---|---|---|
| rs750148755 | 10 | 96095271 | CMC | CMC NoSVA cis | NOC3L | C | | 1.0 | 0.049 |
| rs11187882 | 10 | 96095861 | CMC | CMC NoSVA cis | NOC3L | C | | 1.0 | 0.049 |
| rs11187883 | 10 | 96096866 | CMC | CMC NoSVA cis | NOC3L | C | | 1.0 | 0.049 |
| rs3740359 | 10 | 96100119 | CMC | CMC NoSVA cis | NOC3L | T | | 1.0 | 0.049 |
| rs11187890 | 10 | 96102758 | CMC | CMC NoSVA cis | NOC3L | C | | 1.0 | 0.049 |
| rs11187895 | 10 | 96106240 | CMC | CMC NoSVA cis | NOC3L | C | | 1.0 | 0.049 |
| rs11187897 | 10 | 96106603 | CMC | CMC NoSVA cis | NOC3L | G | | 1.0 | 0.049 |
| 10:96112261_AAAAAAAC_A | 10 | 96112261 | CMC | CMC NoSVA cis | NOC3L | A | | 1.0 | 0.049 |
| rs778275808 | 10 | 96114212 | CMC | CMC NoSVA cis | NOC3L | T | | 1.0 | 0.049 |
| rs10509672 | 10 | 96115957 | CMC | CMC NoSVA cis | NOC3L | C | | 1.0 | 0.049 |
| rs74825688 | 10 | 96119687 | CMC | CMC NoSVA cis | NOC3L | A | | 1.0 | 0.049 |
| rs79193572 | 10 | 96119713 | CMC | CMC NoSVA cis | NOC3L | G | | 1.0 | 0.049 |
| rs79746649 | 10 | 96120218 | CMC | CMC NoSVA cis | NOC3L | T | | 1.0 | 0.049 |
| rs7101207 | 10 | 96122857 | CMC | CMC NoSVA cis | NOC3L | A | | 1.0 | 0.049 |
| rs112252876 | 10 | 96123355 | CMC | CMC NoSVA cis | NOC3L | T | | 1.0 | 0.049 |
| rs11187906 | 10 | 96124838 | CMC | CMC NoSVA cis | NOC3L | A | | 1.0 | 0.049 |
| rs11187907 | 10 | 96125361 | CMC | CMC NoSVA cis | NOC3L | T | | 1.0 | 0.049 |
| rs113544984 | 10 | 96127448 | CMC | CMC NoSVA cis | NOC3L | A | | 1.0 | 0.049 |
| rs16968876 | 15 | 39636877 | eQTLGen | eQTLGen cis eQTLs | THBS1 | C | 3.7769E-210 | -30.9371 | 0.0 |
| rs16968890 | 15 | 39638991 | eQTLGen | eQTLGen cis eQTLs | THBS1 | G | 1.0259E-208 | -30.8304 | 0.0 |
| rs3910645 | 15 | 39648251 | eQTLGen | eQTLGen cis eQTLs | THBS1 | G | 2.2440999999999998E-203 | -30.4294 | 0.0 |
| rs3910644 | 15 | 39648234 | eQTLGen | eQTLGen cis eQTLs | THBS1 | C | 1.0002E-196 | -29.9225 | 0.0 |
| rs11638679 | 15 | 39640148 | eQTLGen | eQTLGen cis eQTLs | THBS1 | T | 1.1882E-85 | -19.6133 | 0.0 |
| rs34680120 | 15 | 39664000 | eQTLGen | eQTLGen cis eQTLs | THBS1 | A | 3.1042E-38 | -12.9284 | 0.0 |
| rs4566132 | 15 | 39661138 | eQTLGen | eQTLGen cis eQTLs | THBS1 | G | 6.7617E-38 | -12.8684 | 0.0 |
| rs1837636 | 15 | 39661779 | eQTLGen | eQTLGen cis eQTLs | THBS1 | C | 3.4541999999999998E-37 | -12.7419 | 0.0 |
| rs72722976 | 15 | 39624864 | eQTLGen | eQTLGen cis eQTLs | THBS1 | G | 3.3560000000000004E-36 | 1,3649E+06 | 0.0 |
| rs12592015 | 15 | 39628131 | eQTLGen | eQTLGen cis eQTLs | THBS1 | A | 9.8428E-36 | 1,0526E+06 | 0.0 |
| rs3910644 | 15 | 39648234 | BIOSQTL | BIOS eQTL geneLevel | THBS1 | C | 1.35871E-34 | -12.2671 | 0.0 |
| rs3910644 | 15 | 39648234 | BIOSQTL | BIOS eQTL geneLevel | CTD-2033D15.1 | C | 1.35871E-34 | -12.2671 | 0.0 |
| rs3910645 | 15 | 39648251 | BIOSQTL | BIOS eQTL geneLevel | THBS1 | G | 1.7692400000000001E-34 | -12.2457 | 0.0 |
| rs3910645 | 15 | 39648251 | BIOSQTL | BIOS eQTL geneLevel | CTD-2033D15.1 | G | 1.7692400000000001E-34 | -12.2457 | 0.0 |
| rs1440802 | 15 | 39635124 | eQTLGen | eQTLGen cis eQTLs | THBS1 | C | 2.7421E-34 | -12.2101 | 0.0 |
| rs28520337 | 15 | 39647894 | eQTLGen | eQTLGen cis eQTLs | THBS1 | C | 6.0685000000000004E-34 | -12.1453 | 0.0 |
| rs71471500 | 15 | 39632013 | eQTLGen | eQTLGen cis eQTLs | THBS1 | G | 2.7098000000000002E-33 | -12.0223 | 0.0 |
| rs11639391 | 15 | 39631771 | eQTLGen | eQTLGen cis eQTLs | THBS1 | T | 7.3753E-33 | -11.9392 | 0.0 |
| rs7182018 | 15 | 39632269 | eQTLGen | eQTLGen cis eQTLs | THBS1 | G | 1.5586E-32 | -11.8769 | 0.0 |
| rs17694988 | 15 | 39636227 | eQTLGen | eQTLGen cis eQTLs | THBS1 | C | 1.9747E-32 | -11.857 | 0.0 |
| rs4924346 | 15 | 39639992 | eQTLGen | eQTLGen cis eQTLs | THBS1 | A | 4.3746E-32 | -11.7902 | 0.0 |
| rs4924345 | 15 | 39639898 | eQTLGen | eQTLGen cis eQTLs | THBS1 | C | 7.2122E-31 | -11.5519 | 0.0 |
| rs1080066 | 15 | 39634222 | eQTLGen | eQTLGen cis eQTLs | THBS1 | G | 7.6265000000000005E-31 | -11.547 | 0.0 |
| rs2033939 | 15 | 39633904 | eQTLGen | eQTLGen cis eQTLs | THBS1 | A | 9.0583E-31 | -11.5322 | 0.0 |

| | | | | | | | | | |
|---|---|---|---|---|---|---|---|---|---|
| rs4923822 | 15 | 39662277 | eQTLGen | eQTLGen cis eQTLs | THBS1 | A | 1.8796000000000002E-30 | -11.4694 | 0.0 |
| rs978740 | 15 | 39655920 | eQTLGen | eQTLGen cis eQTLs | THBS1 | C | 6.1505E-28 | -10.9568 | 0.0 |
| rs16968890 | 15 | 39638991 | BIOSQTL | BIOS eQTL geneLevel | THBS1 | G | 9.96844E-28 | -10.913 | 0.0 |
| rs16968890 | 15 | 39638991 | BIOSQTL | BIOS eQTL geneLevel | CTD-2033D15.1 | G | 9.96844E-28 | -10.913 | 0.0 |
| rs16968876 | 15 | 39636877 | BIOSQTL | BIOS eQTL geneLevel | THBS1 | C | 1.23112E-27 | -10.8939 | 0.0 |
| rs16968876 | 15 | 39636877 | BIOSQTL | BIOS eQTL geneLevel | CTD-2033D15.1 | C | 1.23112E-27 | -10.8939 | 0.0 |
| rs3910644 | 15 | 39648234 | eQTLcatalogue | Fairfax 2014 naive | THBS1 | C | 3.20905E-14 | 0.848997 | 1.60452E-10 |
| rs3910645 | 15 | 39648251 | eQTLcatalogue | Fairfax 2014 naive | THBS1 | G | 2.42235E-13 | 0.823476 | 1.21117E-09 |
| rs16968890 | 15 | 39638991 | eQTLcatalogue | Fairfax 2014 naive | THBS1 | G | 4.96976E-13 | 0.793931 | 2.48488E-09 |
| rs16968876 | 15 | 39636877 | eQTLcatalogue | Fairfax 2014 naive | THBS1 | C | 5.2124E-13 | 0.796074 | 2.6062E-09 |
| rs773225188 | 15 | 39631994 | eQTLcatalogue | Fairfax 2014 naive | THBS1 | AAGAG | 7.77307E-13 | 0.78881 | 3.88653E-09 |
| rs56003663 | 15 | 39617295 | eQTLGen | eQTLGen cis eQTLs | THBS1 | C | 9.5054E-12 | -6.8136 | 0.0 |
| rs4923822 | 15 | 39662277 | BIOSQTL | BIOS eQTL geneLevel | CTD-2033D15.1 | A | 1.42704E-09 | -6.05233 | 6.12782645995465E-06 |
| rs4923822 | 15 | 39662277 | BIOSQTL | BIOS eQTL geneLevel | THBS1 | A | 1.98781E-09 | -5.99861 | 6.08309507877608E-06 |
| rs4566132 | 15 | 39661138 | BIOSQTL | BIOS eQTL geneLevel | THBS1 | G | 2.82253E-09 | -5.94143 | 6.03937673632081E-06 |
| rs4566132 | 15 | 39661138 | BIOSQTL | BIOS eQTL geneLevel | CTD-2033D15.1 | G | 2.82253E-09 | -5.94143 | 6.03937673632081E-06 |
| rs1837636 | 15 | 39661779 | BIOSQTL | BIOS eQTL geneLevel | THBS1 | C | 2.82253E-09 | -5.94143 | 6.03937673632081E-06 |
| rs1837636 | 15 | 39661779 | BIOSQTL | BIOS eQTL geneLevel | CTD-2033D15.1 | C | 2.82253E-09 | -5.94143 | 6.03937673632081E-06 |
| rs34680120 | 15 | 39664000 | BIOSQTL | BIOS eQTL geneLevel | THBS1 | A | 2.82253E-09 | -5.94143 | 6.03937673632081E-06 |
| rs34680120 | 15 | 39664000 | BIOSQTL | BIOS eQTL geneLevel | CTD-2033D15.1 | A | 2.82253E-09 | -5.94143 | 6.03937673632081E-06 |
| rs11638679 | 15 | 39640148 | eQTLGen | eQTLGen cis eQTLs | C15orf54 | T | 8.0695E-09 | -5.7669 | 5.79822187862389E-05 |
| rs11638016 | 15 | 39621773 | eQTLGen | eQTLGen cis eQTLs | THBS1 | C | 1.8572E-08 | -5.6248 | 9.58466453674121E-05 |
| rs56113415 | 15 | 39621418 | eQTLGen | eQTLGen cis eQTLs | THBS1 | A | 3.7254E-08 | -5.5033 | 0.0001523035918263 |
| rs11638679 | 15 | 39640148 | BIOSQTL | BIOS eQTL geneLevel | THBS1 | T | 6.41607E-08 | -5.40673 | 5.63443768311922E-05 |
| rs11638679 | 15 | 39640148 | BIOSQTL | BIOS eQTL geneLevel | CTD-2033D15.1 | T | 6.41607E-08 | -5.40673 | 5.63443768311922E-05 |
| rs3910645 | 15 | 39648251 | eQTLcatalogue | BLUEPRINT ge monocyte | THBS1 | G | 7.11506E-08 | 0.845224 | 0.000355753 |
| rs3910644 | 15 | 39648234 | eQTLcatalogue | BLUEPRINT ge monocyte | THBS1 | C | 7.90024E-08 | 0.84269 | 0.000395012 |
| rs2033939 | 15 | 39633904 | BIOSQTL | BIOS eQTL geneLevel | THBS1 | A | 1.39884E-07 | -5.26522 | 8.847600088476E-05 |
| rs8032326 | 15 | 39619456 | eQTLGen | eQTLGen cis eQTLs | THBS1 | A | 1.8283E-07 | -5.2159 | 0.0006118881118881 |
| rs8033007 | 15 | 39619661 | eQTLGen | eQTLGen cis eQTLs | THBS1 | C | 3.8506E-07 | -5.0762 | 0.0012003464917708 |
| rs1080066 | 15 | 39634222 | BIOSQTL | BIOS eQTL geneLevel | THBS1 | G | 5.84104E-07 | -4.99636 | 0.0005052116570942 |
| rs28520337 | 15 | 39647894 | BIOSQTL | BIOS eQTL geneLevel | THBS1 | C | 6.5653E-07 | -4.97369 | 0.0005301664722722 |
| rs28520337 | 15 | 39647894 | BIOSQTL | BIOS eQTL geneLevel | CTD-2033D15.1 | C | 6.5653E-07 | -4.97369 | 0.0005301664722722 |
| rs16968890 | 15 | 39638991 | eQTLcatalogue | BLUEPRINT ge monocyte | THBS1 | G | 7.00785E-07 | 0.796662 | 0.00350393 |
| rs12591944 | 15 | 39627982 | eQTLGen | eQTLGen cis eQTLs | THBS1 | C | 1.2149E-06 | -4.853 | 0.0036831175177131 |
| rs1440802 | 15 | 39635124 | BIOSQTL | BIOS eQTL geneLevel | THBS1 | C | 1.53212E-06 | -4.80687 | 0.0012847743874009 |
| rs72722978 | 15 | 39628675 | eQTLGen | eQTLGen cis eQTLs | THBS1 | T | 1.7935E-06 | -4.7753 | 0.0053233558435237 |
| rs8025239 | 15 | 39623588 | eQTLGen | eQTLGen cis eQTLs | THBS1 | A | 2.0892E-06 | -4.7445 | 0.0060943006440636 |

| | | | | | | | | | |
|---|---|---|---|---|---|---|---|---|---|
| rs16968890 | 15 | 39638991 | eQTLcatalogue | Fairfax 2014 LPS24 | THBS1 | G | 2.28413E-06 | 0.227449 | 0.0114206 |
| rs4924345 | 15 | 39639898 | BIOSQTL | BIOS eQTL geneLevel | THBS1 | C | 2.33078E-06 | -4.72221 | 0.0017824310520939 |
| rs4924345 | 15 | 39639898 | BIOSQTL | BIOS eQTL geneLevel | CTD-2033D15.1 | C | 2.33078E-06 | -4.72221 | 0.0017824310520939 |
| rs16968876 | 15 | 39636877 | eQTLcatalogue | Fairfax 2014 LPS24 | THBS1 | C | 2.85595E-06 | 0.225868 | 0.0142798 |
| rs16968876 | 15 | 39636877 | eQTLcatalogue | BLUEPRINT ge monocyte | THBS1 | C | 2.8667E-06 | 0.752794 | 0.0143335 |
| rs773225188 | 15 | 39631994 | eQTLcatalogue | Fairfax 2014 LPS24 | THBS1 | AAGAG | 3.7226E-06 | 0.222135 | 0.018613 |
| rs3910644 | 15 | 39648234 | GTEx/v8 | Whole Blood | THBS1 | T | 4.45129E-06 | 0.127755 | 3.8153199999999997E-20 |
| rs1837636 | 15 | 39661779 | PsychENCODE | PsychENCODE eQTLs | RP11-37C7.1 | T | 5.27257E-06 | -0.222586 | 0.0006116329885143 |
| rs3910644 | 15 | 39648234 | eQTLcatalogue | Fairfax 2014 IFN24 | THBS1 | C | 5.33505E-06 | 0.461593 | 0.0266752 |
| rs10520128 | 15 | 39627146 | eQTLGen | eQTLGen cis eQTLs | THBS1 | C | 7.0029E-06 | -4.4934 | 0.0199808623886131 |
| rs11629938 | 15 | 39626935 | eQTLGen | eQTLGen cis eQTLs | THBS1 | A | 8.6916E-06 | -4.4473 | 0.0246107963018193 |
| rs34680120 | 15 | 39664000 | PsychENCODE | PsychENCODE eQTLs | RP11-37C7.1 | A | 9.31429E-06 | 0.216218 | 0.0010235897899314 |
| rs16968890 | 15 | 39638991 | eQTLcatalogue | Fairfax 2014 IFN24 | THBS1 | G | 9.51089E-06 | 0.431862 | 0.0475544 |
| rs3910644 | 15 | 39648234 | eQTLcatalogue | BLUEPRINT ge monocyte | RP11-624L4.1 | C | 9.5575E-06 | 0.327274 | 0.0477875 |
| rs4924346 | 15 | 39639992 | BIOSQTL | BIOS eQTL geneLevel | THBS1 | A | 2.08303E-05 | -4.25569 | 0.0133721749753324 |
| rs4924346 | 15 | 39639992 | BIOSQTL | BIOS eQTL geneLevel | CTD-2033D15.1 | A | 2.08303E-05 | -4.25569 | 0.0133721749753324 |
| rs16968876 | 15 | 39636877 | GTEx/v8 | Whole Blood | THBS1 | T | 2.227E-05 | 0.119692 | 3.8153199999999997E-20 |
| rs17694988 | 15 | 39636227 | BIOSQTL | BIOS eQTL geneLevel | THBS1 | C | 2.25922E-05 | -4.23748 | 0.014281031980147 |
| rs17694988 | 15 | 39636227 | BIOSQTL | BIOS eQTL geneLevel | CTD-2033D15.1 | C | 2.25922E-05 | -4.23748 | 0.014281031980147 |
| rs12592015 | 15 | 39628131 | DICE | T CD4 naive | PAK6 | A | 2.38152860034E-05 | 1.15860040249 | 0.049 |
| rs773225188 | 15 | 39631994 | GTEx/v8 | Whole Blood | THBS1 | A | 2.69055E-05 | 0.11834 | 3.8153199999999997E-20 |
| rs16968890 | 15 | 39638991 | GTEx/v8 | Whole Blood | THBS1 | C | 3.04173E-05 | 0.118313 | 3.8153199999999997E-20 |
| rs3910645 | 15 | 39648251 | GTEx/v8 | Whole Blood | THBS1 | C | 3.82701E-05 | 0.115051 | 3.8153199999999997E-20 |
| rs4923822 | 15 | 39662277 | PsychENCODE | PsychENCODE eQTLs | RP11-37C7.1 | G | 4.82356E-05 | -0.204656 | 0.0044418712164 |
| rs8032326 | 15 | 39619456 | PsychENCODE | PsychENCODE eQTLs | RP11-37C7.1 | A | 0.000314673 | 0.162609 | 0.0224872458661352 |
| rs2033939 | 15 | 39633904 | PsychENCODE | PsychENCODE eQTLs | RP11-37C7.1 | A | 0.000399035 | 0.157412 | 0.0274319167847911 |

| Center of mass of the language processing regions of interests retained in both left and right hemispheres. Each ROIs is characterized by their abbreviated anatomical label and their center of mass stereotactic coordinates (x, y, z, in mm) | | | | |
|---|---|---|---|---|
| Hemisphere | ROIs | x | y | z |
| Left hemisphere | Prec | -46.763359 | 2.010178 | 25.857506 |
| | RolS | -45.287982 | -9.405896 | 44.176871 |
| | F3orb | -35.821290 | 35.303807 | -10.772339 |
| | T3p | -45.903226 | -56.483871 | -7.359447 |
| | F3td | -42.150235 | 25.107981 | 14.544601 |
| | T2p | -39.306773 | -65.254980 | 2.051793 |
| | SMG | -40.684932 | -50.252055 | 40.865753 |
| | T1a | -54.076294 | -11.858311 | -5.858311 |
| | F3tv | -45.202073 | 14.242228 | 4.231865 |
| | STSp | -47.122995 | -56.926916 | 24.071301 |
| | PrF3op | -39.616883 | 2.909091 | 39.344156 |
| | Fusa | -35.273273 | -37.173173 | -12.890891 |
| | T1 | -50.465171 | -38.564345 | 14.323495 |
| | T2ml | -54.990991 | -39.625225 | -2.846847 |
| | F2p | -34.726790 | 10.944297 | 48.801061 |
| | F3opd | -43.908772 | 21.976608 | 25.702924 |
| | PT | -59.899083 | -28.119266 | 8.678899 |
| | Pole | -45.000000 | 5.196759 | -26.796296 |
| | AG | -43.936872 | -70.852224 | 22.203730 |
| Right hemisphere | T1R | 51.842553 | -38.519149 | 3.404255 |
| | F3opdR | 49.679389 | 14.931298 | 27.480916 |
| | F2antR | 46.641026 | 39.807692 | 5.064103 |
| | T2pR | 54.768421 | -42.115789 | -7.736842 |
| | T1a/HeschlR | 56.540260 | -13.387013 | 0.815584 |
| | PrecR | 52.686275 | -1.490196 | 24.156863 |
