## Supplemental Figures for "The genetic architecture of language functional connectivity"

Supplementary information

List of sections:

1. List of Supplementary Tables (separate Excel file)
2. Supplementary Figures

### 1. Supplementary tables (separate Excel file)

**Supplementary Table S1.** Overview of the regions obtained from the meta-analysis. Each ROIs is characterised by their abbreviated anatomical label defined by (Vigneau et al., 2006, 2011) and is labelled according to the language component they belong to : phonology, semantic, and syntax.

**Supplementary Table S2.** Heritabilities of the 300 brain functional connectivity, estimated using the genotyped SNPs information using genome-based restricted maximum likelihood (GREML) (Yang et al., 2010) as implemented in GCTA (Yang et al., 2011) software (version 1.93.2beta). A 0.05 threshold on False Discovery Rate (FDR) adjusted p-values was applied to account for multiple testing.

**Supplementary Table S3.** SNPs associated with the 142 heritable functional connectivity measures using MOSTest (van der Meer et al., 2020) at the genome-wide significance threshold $p = 5\text{e}{-8}$.

**Supplementary Table S4.** Replication of the 20 lead SNP association using an independent non-British replication dataset (N=4,754) using MOSTest (van der Meer et al., 2020). We considered the nominal significance threshold $pvalue < 0.05$.

**Supplementary Table S5.** For each of the 20 lead SNPs identified in the multivariate GWAS, the corresponding univariate summary statistics for FCs identified as central FC (threshold on the genome-wide significance threshold $p = 5\text{e}{-8}$).

**Supplementary Table S6.** The SNP-based genetic correlation analysis was estimated (using GCTA (Lee et al., 2012) software, version 1.93.2beta) for each pair of central FCs associated to 15q14 or 3p11.1 genetic loci.

**Supplementary Table S7.** Univariate associations of 2 lead SNPs (rs1440802 on 15q14, rs35124509 on 3p11.1) using PLINK 1.9 (Purcell et al., 2007) with diffusion MRI indices on the following 7 white matter tracts: the corpus callosum, the left frontal aslant tract, the left arcuate anterior/long/posterior segment, the left inferior fronto-occipital fasciculus, the left uncinate tract. Significant results were considered at the Bonferroni-corrected threshold

$p = 6.94e{-3} (0.05/(3 * 9 + 5 * 9))$.

**Supplementary Table S8.** eQTLs association, performed by FUMA, between the SNPs in the three replicated genomic risk loci and all mapped genes in the following databases : GTEx/v8 (Consortium et al., 2017), PsychENCODE (Wang et al., 2018), eQTLGen (Võsa et al., 2018), eQTLcatalogue, DICE (Schmiedel et al., 2018), BIOSQTL (Zhernakova et al., 2017). A 0.05 threshold on False Discovery Rate (FDR) adjusted p-values was applied to account for multiple testing.

**Supplementary Table S9.** Center of mass of the language processing regions of interests retained in both left and right hemispheres. Each ROIs is characterised by their abbreviated anatomical label defined by (Vigneau et al., 2006, 2011) and their center of mass MNI stereotactic coordinates (x, y, z, in mm).

### 2. Supplementary figures

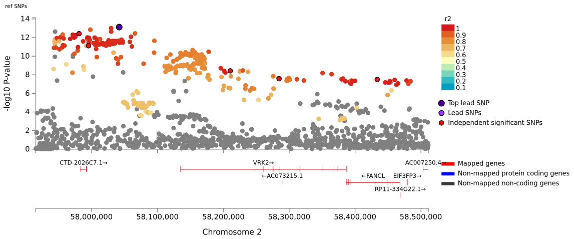


lead SNP: rs2717046

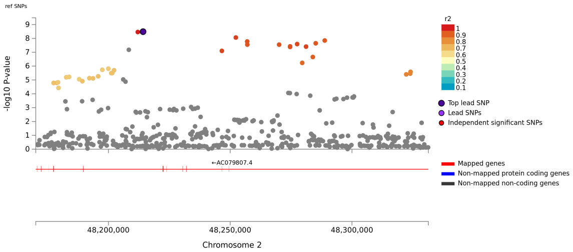


lead SNP: rs62141276

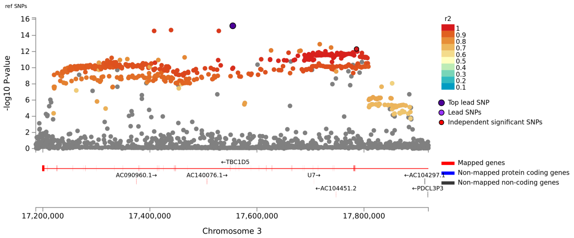


lead SNP: rs67851870

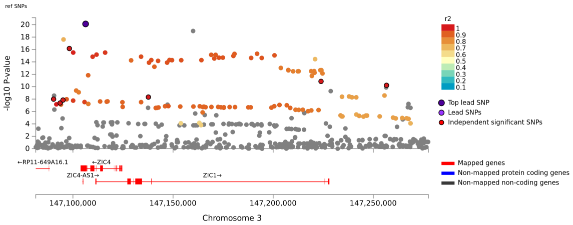


lead SNP: rs2279829

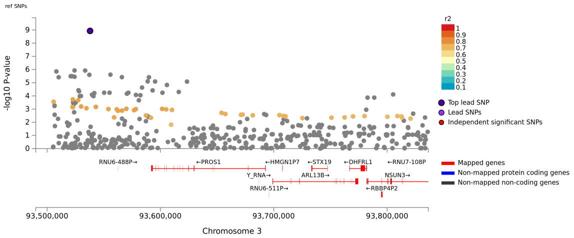


lead SNP: rs62266110

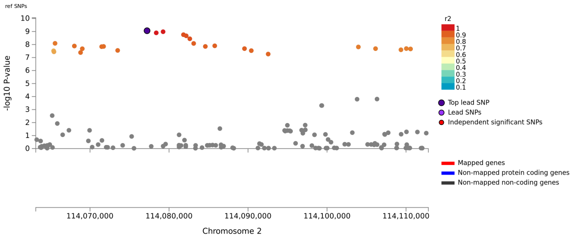


lead SNP: rs62158166

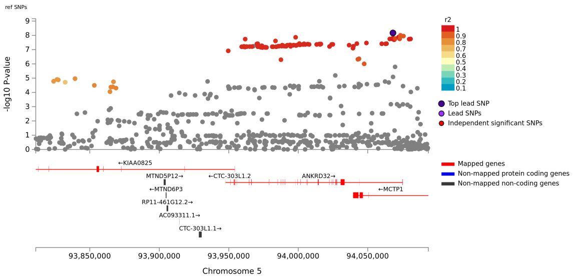


lead SNP: 5:94068140_AC_A

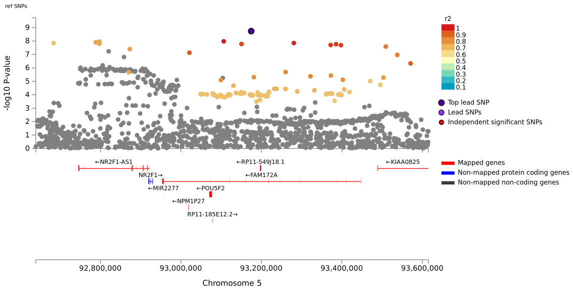


lead SNP: rs145120402

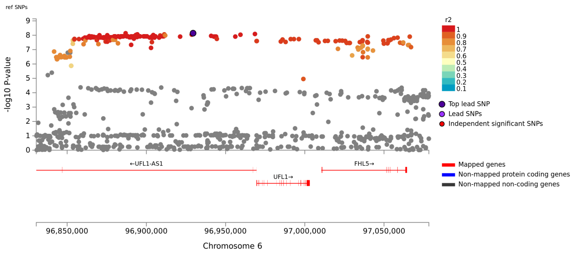


lead SNP: rs4262195

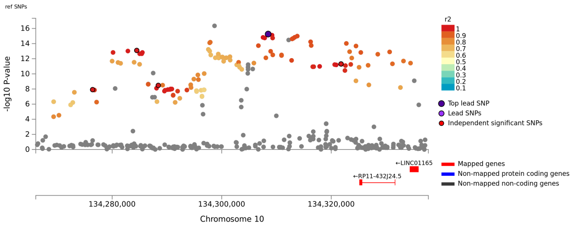


lead SNP: rs11146399

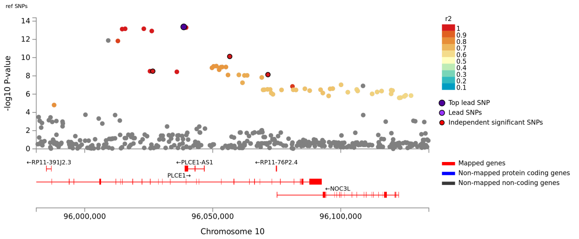


lead SNP: rs11187838

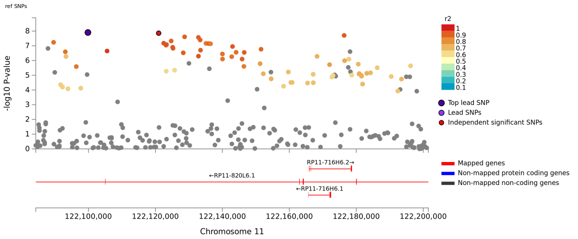


lead SNP: rs11218557

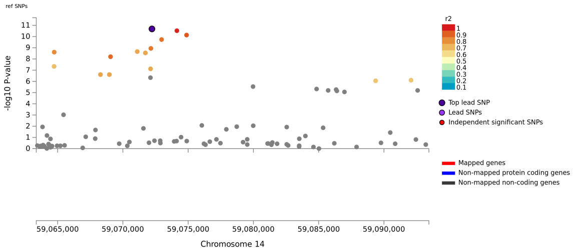


lead SNP: rs186347

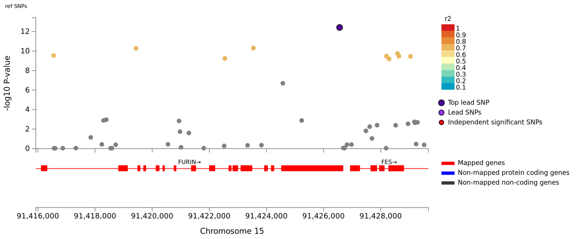


lead SNP: rs4702

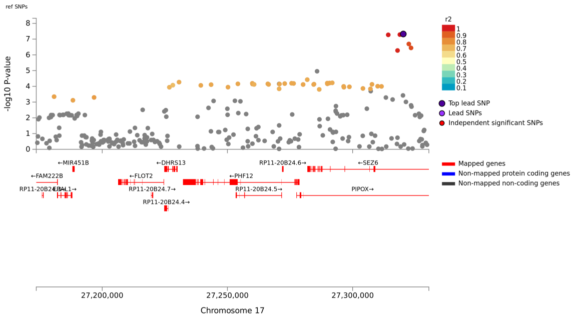


lead SNP: rs34039488

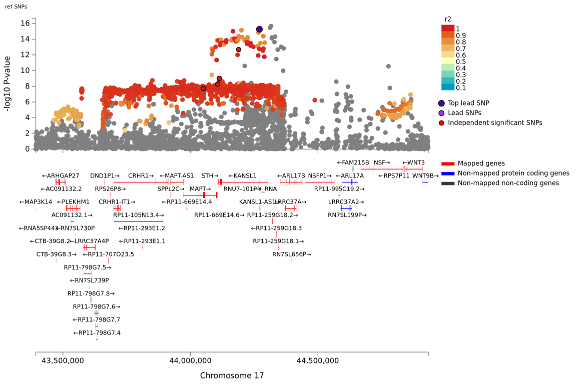


lead SNP: 17:44270659_G_A

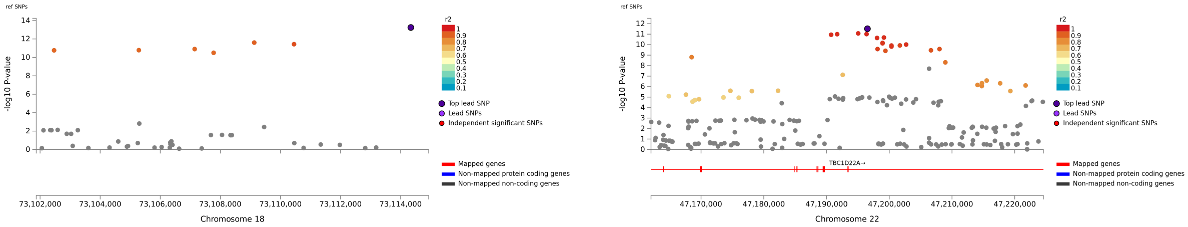


lead SNP: rs7234875 lead SNP: rs2542028

Figure SI1: Locus Zoom of the significant loci identified by the multiariate GWAS for functional connectivity.

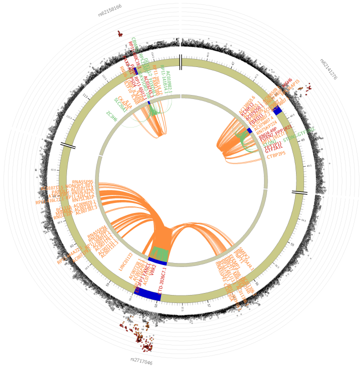

chromosome 2

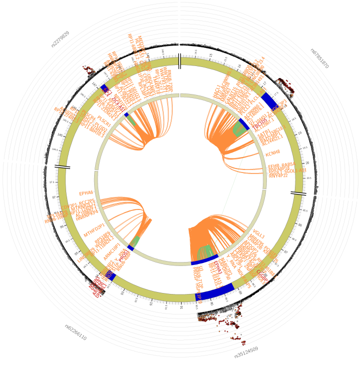

chromosome 3

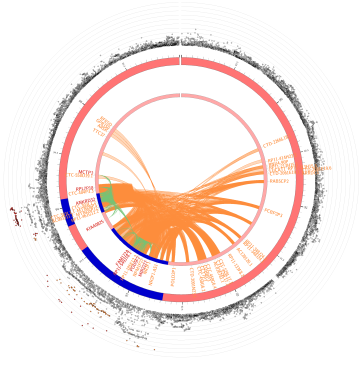

chromosome 5

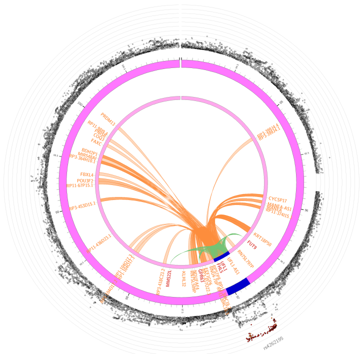

chromosome 6

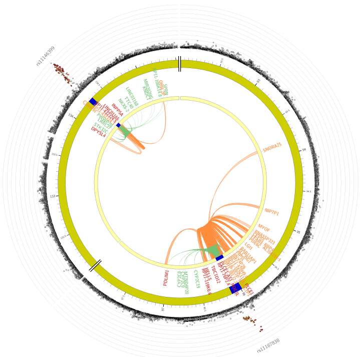

chromosome 10

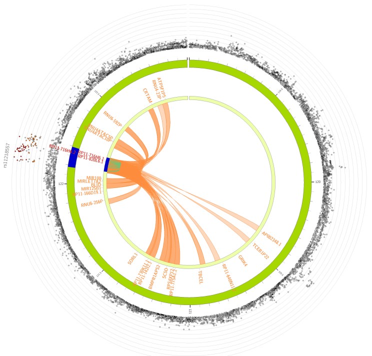

chromosome 11

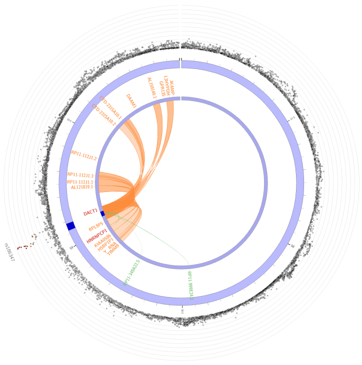

chromosome 14

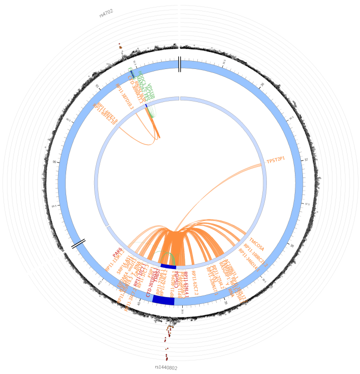

chromosome 15

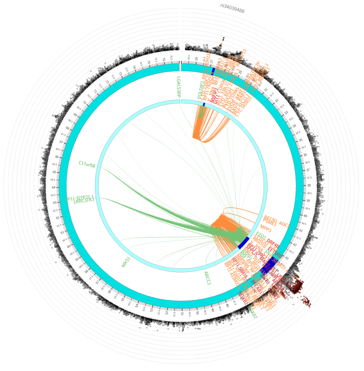

chromosome 17

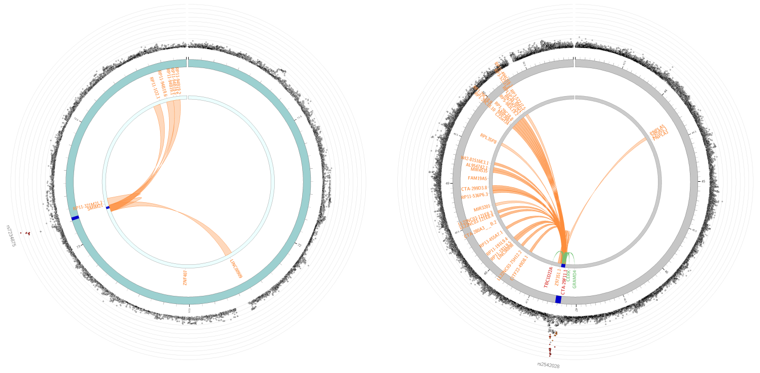

chromosome 18 chromosome 22

Figure SI2: **Genomic loci, eQTL associations and chromatin interactions identified via multivariate GWAS for functional connectivity.** Circos plot representing the genomic risk loci, and the genes associated with the loci by chromatin interactions and eQTLs. From outer layer to inner layer: Manhattan plot. Genomic risk loci are in blue. Genes mapped by chromatin interaction are in orange. Genes mapped by eQTL are in green. Genes mapped by both are in red. Chromatin interaction and eQTLs links follows the same color coding presented above.

rs4262195 (chr. 6)

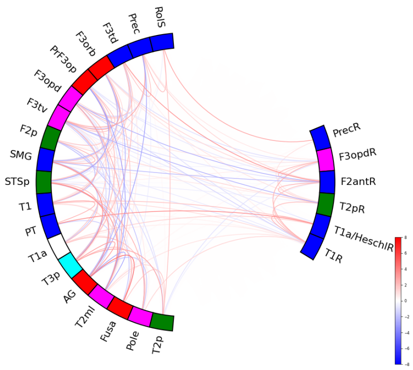


rs7234875 (chr. 18)

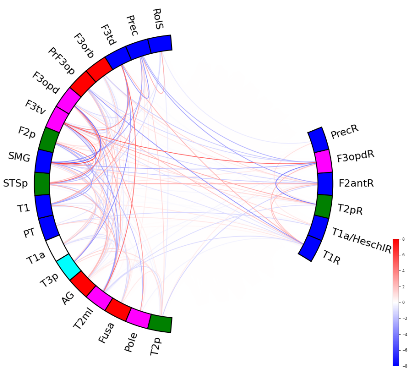


rs11146399 (chr. 10)

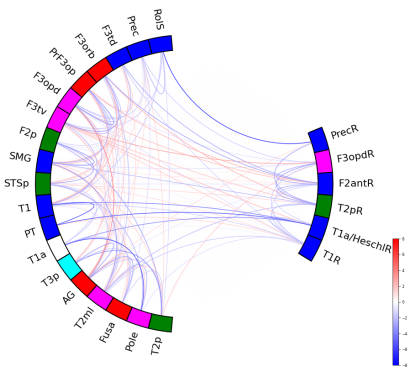


rs62141276 (chr. 2)

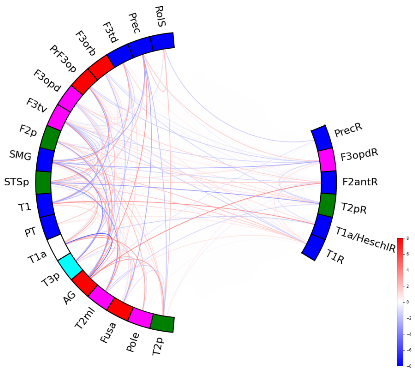


rs11218557 (chr. 11)

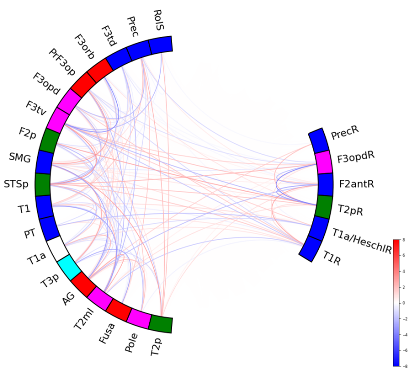


rs34039488 (chr. 17)

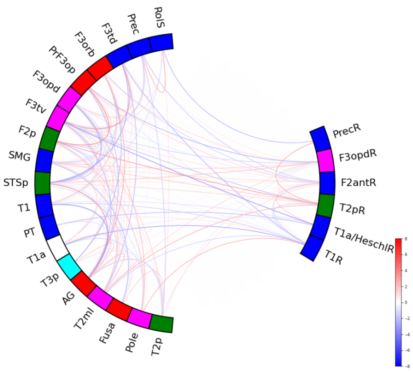


17:44270659_G_A (chr. 17)

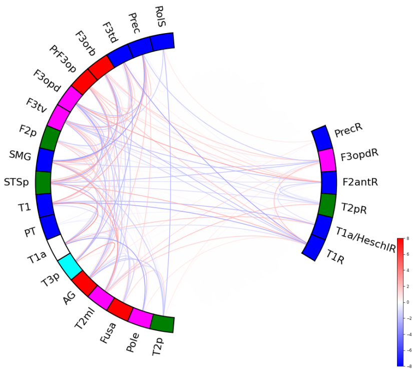


rs62158166 (chr. 2)

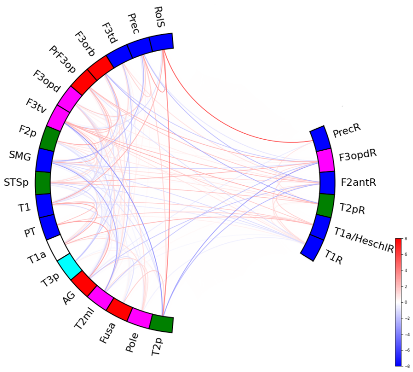


rs62266110 (chr. 3)

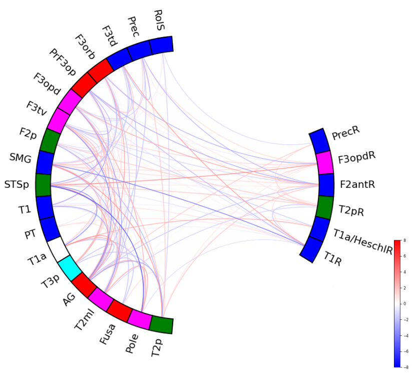

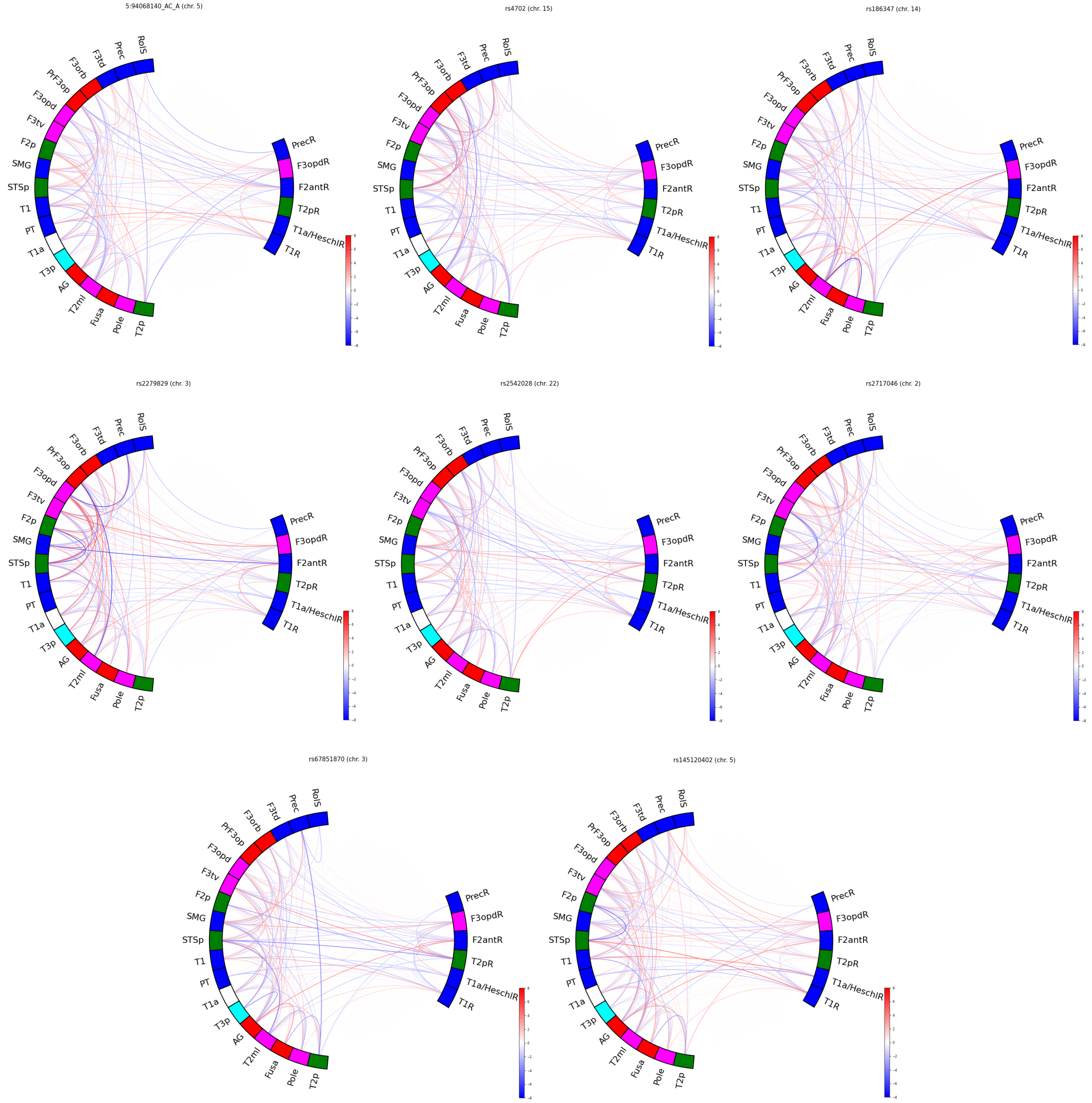


Figure SI3: **Regional effects.** Circle plot illustrating the lead SNPs identified from the multivariate GWAS for functional connectivity. Z-values from the univariate GWAS for each FCs are mapped. The absolute Z-values scaling is clipped at 8 ($p = 1.2\text{e}{-15}$). Positif effects of carrying the minor allele are shown in red, and negative in blue.

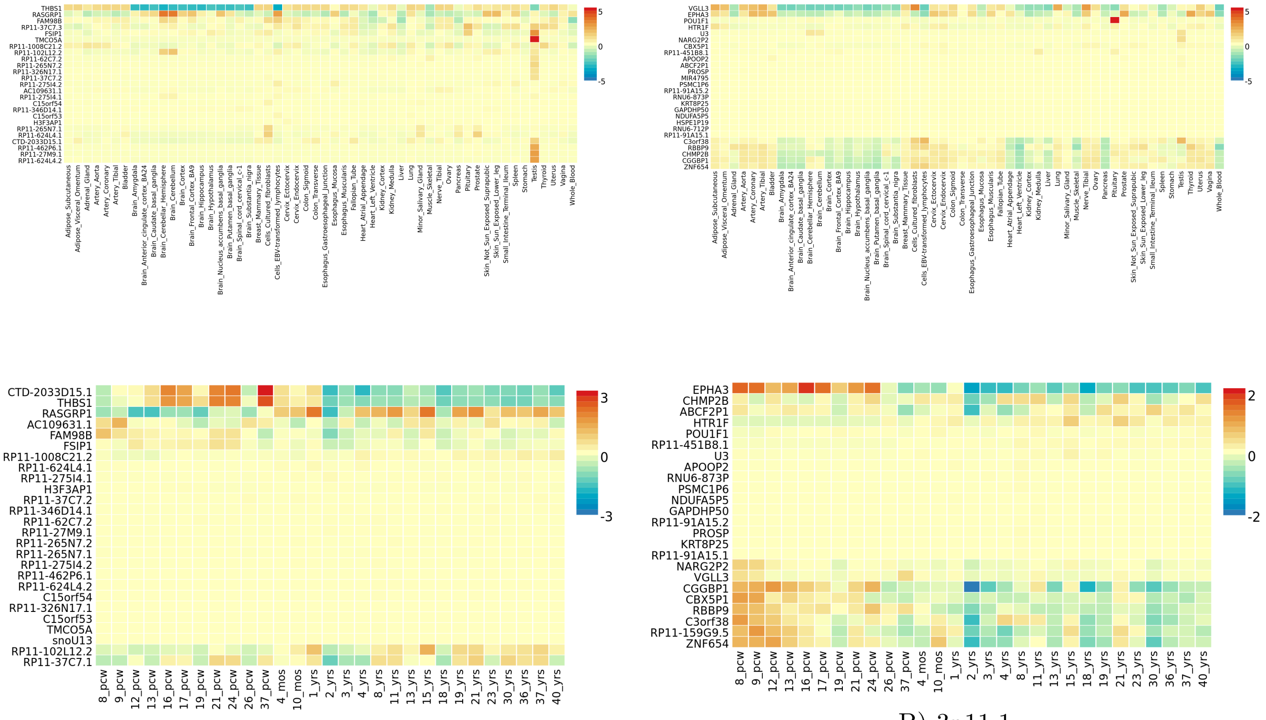


A) 15q14

B) 3p11.1

Figure SI4: **Functional annotation of both genomic risk loci 15q14 and 3p11.1.** A) Gene expression heatmap constructed with GTEx/v8 (54 tissue types) and B) BrainSpan 29 different ages of brain samples. (Average of normalized expression per label).
